# E cadherin appears to be an essential on/off switch for initiating bile canaliculi formation

**DOI:** 10.1101/2024.06.04.597350

**Authors:** Mireille Chevallet, Thierry Rabilloud, Hélène Diemer, Fabrice Bertile, Alexandra Fuchs, Aurélien Deniaud

## Abstract

The mechanisms underlying cell polarization are fundamental in biology, yet they are not fully understood. This is particularly true for hepatocytes, which exhibit a complex polarization, enabling the formation of the bile canaliculi (BCs) network that is essential for liver excretion functions. To identify key proteins involved in hepatocyte polarization and BC formation, we performed a proteomic approach to compare the human hepatocyte cell line HepG2 to its sub clone HepG2/C3A which shows much greater efficiency in forming mature BCs. We localized LimA1 and Espin to the BC for the first time, suggesting their important role there, and confirmed the presence of NHE-RF1. Using a protein repression strategy directed against selected proteins, we highlighted the essential role of E cadherin in the initiation of BC formation. Our data showed, for the first time, that in absence of E cadherin, hepatocytes lose their ability to form BCs.

## Introduction

Cellular polarity is fundamental for the formation and maintenance of appropriate structure and function of almost all tissues. Epithelial cells polarize along an apical-basal axis to form selectively permeable barriers, preventing the free flow of molecules and establishing distinct molecular and functional domains in their apical, basal, and lateral membranes (Riga et al., 2020). Separation of the internal from the external environment is mediated by the organization of uniformly polarized cells into sheets, with specialized cell-cell junctions holding the cells together, in particular adherens and tight junctions. In higher vertebrates, there are over one hundred and fifty different types of epithelia that constitute the essential functional elements of most organs in the body.

The liver is organized in functional units or lobules, 80% of which consist of specific epithelial cells, the hepatocytes. In each lobule, hepatocytes are arranged in hepatic cords separated by adjacent capillaries or sinusoids. Local membrane invaginations between adjacent hepatocytes form a network of microchannels called bile canaliculi (BC) that are contained within each cord. BCs collect bile and are connected to a larger interlobular bile duct hierarchical tree formed by cholangiocytes. Even though hepatocytes and cholangiocytes derive from the common progenitor hepatoblasts, cholangiocytes present a typical columnar epithelial polarization (as described below), whereas hepatocytes display a unique polarity phenotype. A single hepatocyte can form BC lumina with up to three neighbours and can have two basal domains that face the adjacent sinusoids (Gissen and Arias, 2015).The canalicular surface forms microvilli, which considerably increase the apical surface area of the plasma membrane available for bile secretion.

Actin microfilaments, normally found around BCs (pericanalicular web), play an active role in bile secretion. Many experimental studies have indeed shown their importance for the dilatation and contraction of bile canaliculi, suggesting their potential role in control of bile flow (Karpen and Crawford, 1999). The canalicular surface is segregated from the rest of the intercellular surface by junctional complexes: desmosomes, adherens junctions, tight junctions and gap junctions. Among these complexes, tight junctions hold particular significance as they prevent the leakage of plasma into the bile as well as the reflux of bile from BCs into the blood.

During mouse embryogenesis, the hepatic bud emerges around day 11, made-up of non-polarized hepatoblasts, with BCs only becoming apparent in late gestation, around day 17. These early BC domains, formed by multiple hepatocytes, appear dilated; they are often referred to as acini. The emergence of true elongated BC profiles becomes evident by embryonic day 21 and undergoes further development for approximately two weeks after birth (Müsch, 2018). One of the unique characteristics of the liver is its remarkable ability to regenerate, even after losing up to 70% of its mass (Treyer and Müsch, 2013). Significant parallels can be drawn between the morphogenetic events leading to the organization of hepatocytes into cords during embryonic development and liver regeneration. This suggests that the regenerating liver follows a differentiation program similar to that of embryonic development (ref hepatocyte polarity). Hepatocyte polarization and the formation of the BC network require the synchronized expression of numerous key elements of the extracellular matrix (ECM), cell-cell junctions, intracellular protein trafficking machinery (including recycling endosomes), the cytoskeleton, and energy production (Gissen and Arias, 2015). The signalling pathways leading to hepatocyte polarization are partially known, and the key proteins involved in BC formation have yet to be fully identified. A better understanding of the key players in the establishment of hepatocyte polarity and BC formation could be a first step for improving knowledge on liver embryogenesis, regeneration, pathologies and toxicity. Indeed, establishment and maintenance of hepatocyte polarity is essential for many liver functions, and the polarized state of hepatic cells is frequently compromised in liver disease. For example, cholestasis, which can be caused by either a toxin or a genetic mutation, prevents hepatocytes from transporting and secreting the various components of bile into BC.

Using a combined siRNA and proteomic approach in human hepatocyte cell lines, we have identified key protein players in hepatocyte polarization and highlighted certain proteins that are particularly important for BC formation, structure and/or function. We focused on adherens junction proteins and confirmed the pivotal role of E cadherin in initiating the formation as well as the growth of BCs. We also highlighted the hitherto unknown role of three different proteins in BC function, namely the bundling protein Espin and the actin binding proteins LimA1 and Myo1D.

## Results and discussion

### Differential proteomics provides insight into the increased BC formation potential of HepG2/C3A cells compared to HepG2 cells

The human hepatocyte cell line HepG2/C3A is a clonal derivative of HepG2 that has been selected for its enhanced phenotypic characteristics, including strong contact inhibition of growth, high albumin and alpha-fetoprotein production, and the ability to grow in glucose-deprived medium. HepG2/C3A cell line is a valuable tool in early drug discovery and pharmaceutical development (Coltman et al., 2021, Štampar et al., 2021). In a previous study, we described that HepG2/C3A is much more efficient than HepG2 in forming BCs. This can be confirmed by immunofluorescence (IF) labelling of a canalicular transporter (MRP2) and the major cytoskeletal protein (F-actin), as well as staining of nuclei in standard HepG2 and HepG2/C3A 2D cultures (Fig. S1). Indeed, BCs (characterized by colocalization of actin and MRP2) are rarely seen in HepG2 cells, while they are abundant in HepG2/C3A cells (Fig. S1a-b). Images of HepG2/C3A at higher magnification showed BCs at various stages of maturity, with either a patch of actin (immature) or surrounded by a ring of actin (mature) (Fig. S1c-d).

To strengthen the analysis, we compared the proteome of HepG2 cells to that of its subclone HepG2/C3A, both grown in 2D. A total of 3,465 proteins were identified (Table S1), from various cellular compartments, and representing multiple biological processes. Of the 1150 differentially expressed proteins (p-value < 0.05), pathway analysis revealed four main classes of proteins implicated in metabolism, cytoskeleton, trafficking, and in cell-cell and cell-matrix adhesion (Fig. 1).

**Figure 1.**
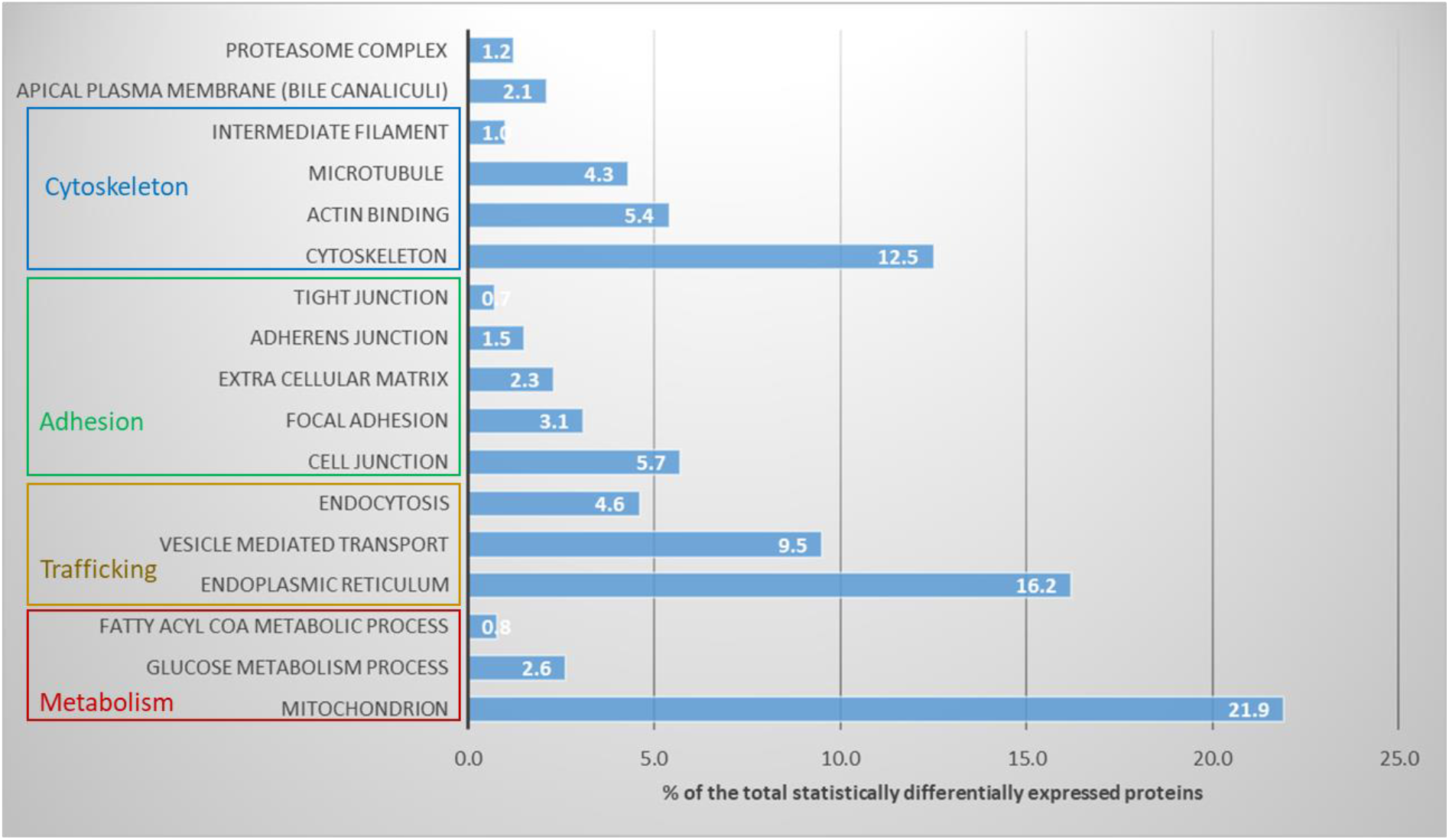
Representation of the different classes of proteins differentially expressed between HepG2 and HepG2/C3A cell lines. Each class is expressed as a percentage of the total proteins, statistically differentially expressed (1150).

These results are consistent with the literature as described in (Gissen and Arias, 2015) and show that HepG2/C3A cells have a more liver-specific phenotype than HepG2 cells as reflected by the expression of hepatic metabolism proteins or morphological components. We notably found that a high percentage (22.3%) of the differentially expressed proteins are involved in mitochondrial functions. Hepatocyte polarization is energy-dependent but the mechanism is unclear (Gissen and Arias, 2015).The fact that the HepG2/C3A cells were selected on the basis of their improved metabolic capacity could also explain this result.

Within the cytoskeleton class (12.5%), we found proteins implicated or associated with the actin network (5.4%), microtubule network (4.3%) and to a lesser extent with intermediate filaments (1%). Cytoskeletal actin can play a structural role by forming a ring around BCs, as seen in HepG2/C3A cells (see Fig. S1c-d) and is also important for trafficking. The role of the microtubule network is essentially in link with the intracellular trafficking, as these dynamic structures mediate trafficking of secreted and canalicular proteins (Gissen and Arias, 2015). The role of intermediate filament is not described in this context, they are considered as mechanical components of the cell that provide resistance to deformation stress. In the case of the adhesion class, proteins involved in adherens junctions (1.5%), tight junctions (0.7%), focal adhesion (3.1%) and ECM (2.3%) were highlighted, reflecting possible differences in the establishment of cell polarity between HepG2 and HepG2/C3A cells.

In the case of intracellular trafficking, our proteomics data also revealed great differences between the two cell lines with regard to endoplasmic reticulum (16.2%), vesicle-mediated transport (9.5%) and endocytosis (4.6%). We also identified proteins specific to the apical pole such as some BC transporters (2.4%).

We next examined in more details the differences in abundance between the two cell lines of selected differentially expressed proteins known to be involved in hepatocyte polarization and BC formation and function (Table 1). We also selected proteins not previously described within this context, such as Lima1 and Myo1D, as their role in other cell types or organs suggest that they could be involved in BCs (Table 1). It should be noted that our selection is by no means exhaustive, and numerous other proteins could potentially be of interest (Table S1).

**Table 1.**
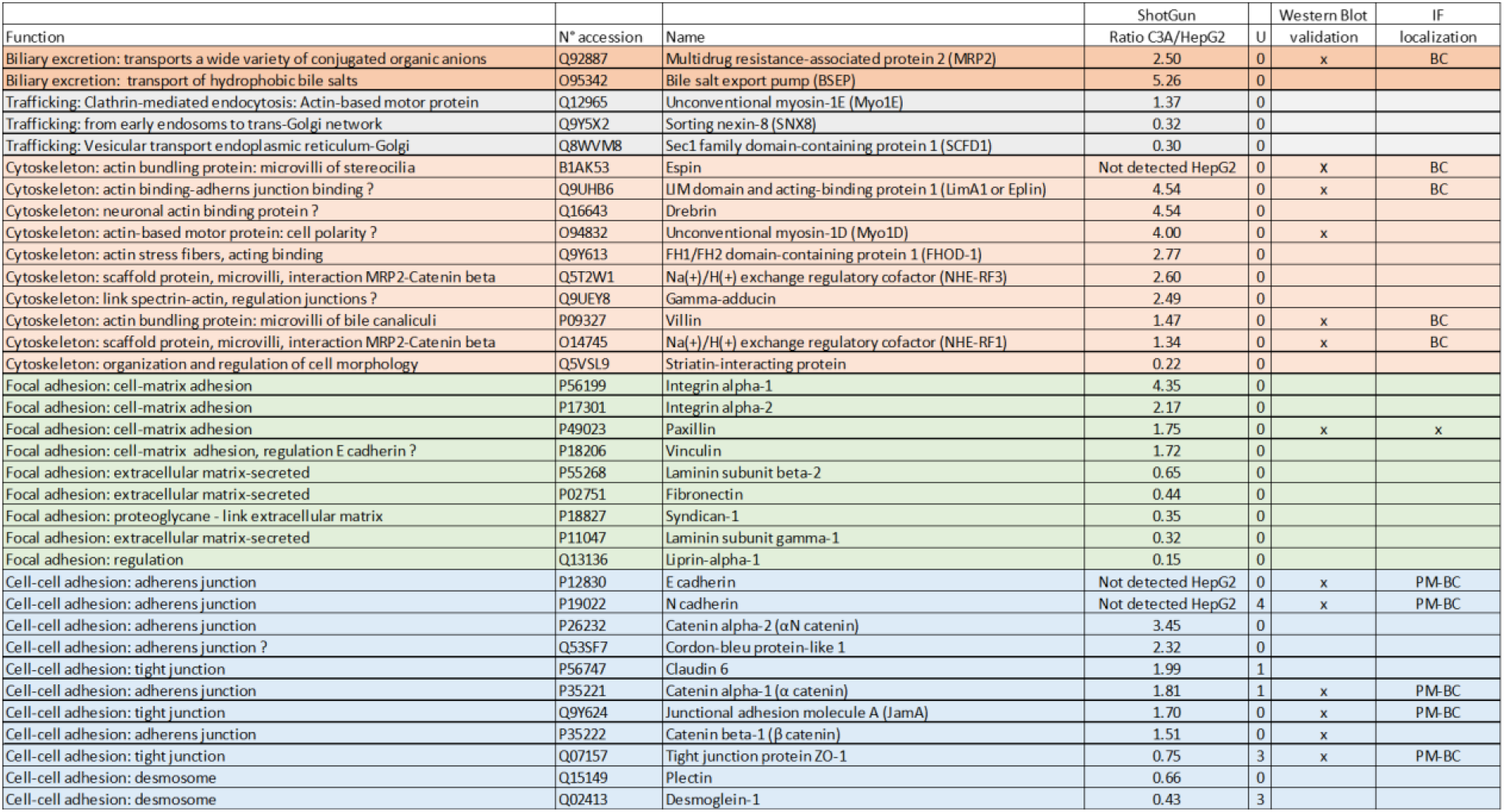
List of differentially expressed proteins between HepG2 and HepG2/C3A cell lines selected and classified by their functions. Summary of shotgun and Western Blot results. PM: plasma membrane, BC: bile canaliculi.

We identified cytoskeleton proteins, particularly actin-binding proteins, which are predominantly overexpressed in HepG2/C3A compared to HepG2 cells, with only the Striatin-interacting protein showing a higher level of expression in HepG2 cells. Very interestingly, the actin cytoskeleton proteins induced in HepG2/C3A cells are often annotated in databases with cellular movement, protrusions, conversion into mechanical force; functions we expect to deform the membrane and create a channel.

We also identified proteins involved in cell-cell adhesion and focal adhesion. Interestingly, within the cell-cell adhesion category, proteins involved in adherens junctions were mainly upregulated in HepG2/C3A cells, while proteins of desmosomes were mostly upregulated in HepG2 cells. Two transmembrane proteins of tight junctions, JamA and Claudin 6, were more abundant in the HepG2/C3A cell line, while the scaffold protein ZO-1, which connects them to actin filaments, was slightly decreased in this cell line. Among the focal adhesion proteins, those implicated in cell-matrix adhesion were upregulated in HepG2/C3A cells, while excreted proteins of the ECM were upregulated in HepG2 cells. Proteins involved in intracellular trafficking were predominantly overexpressed in the HepG2 cell line and, not surprisingly, BC-specific transporters were on the contrary downregulated in HepG2 cells. These results are strongly supportive of the more hepatocyte-like phenotype of HepG2/C3A cells and their higher potential for BC formation and polarization establishment. Therefore, we decided to more specifically study some of these proteins, among which were intriguing specific candidates, to further understand their role in BC formation and structure.

### The higher potential of HepG2/C3A cells for BC formation involves biliary secretion, intracellular trafficking and focal adhesion proteins

**Amongst biliary excretion proteins**, two BC specific transporters implicated in biliary excretion, BSEP and MRP2, were less abundant in the HepG2 cell line compared to the HepG2/C3A clone (Table 1). We confirmed this result obtained via our proteomics approach using Western Blotting (WB) and IF for MRP2 (Fig. S1b-d and S2a). This result is not surprising, as these proteins are typically targeted to BCs, which are virtually absent in HepG2 cells (Fig. S1a). Interestingly, MRP2 was present very early during BC formation (Fig. S1c), and could already be observed at the actin patch stage at 5 hours, and colocalized with actin during BC formation (Fig. S1c). This very early targeting of a transporter to the BC supports a potential role in inducing their formation, probably related to intracellular trafficking.

**The intracellular trafficking machinery** of a hepatocyte is particularly complex and highly developed compared to a typical epithelial cell. To maintain stability of blood chemistry, hepatocytes simultaneously secrete and internalize a wide variety of proteins. Secretion of proteins into the bloodstream takes place at the basal pole, while detoxification and secretion of various compounds into the bile takes place at the apical pole (Schulze et al., 2019). Apical proteins are either directly targeted from the trans-Golgi to the BCs or they transit through the basolateral membrane. Precise regulation of addressing and intracellular trafficking is necessary to maintain hepatocyte polarity. In light of this, it is once again not surprising to observe different expression levels between the two cell lines for proteins involved in intracellular trafficking. More specifically, we found Myo1E overexpressed in HepG2/C3A cells (Table 1). This protein has been previously identified to play various roles, such as in clathrin-mediated endocytosis, glomerular filtration in the kidney, and cell-cell adhesion (Ouderkirk and Krendel, 2014). It is therefore possible that Myo1E also acts in hepatocytes, in particular HepG2/C3A cells, perhaps in connection with biliary secretion.

Both SNX8 and SCFD1 are overexpressed in HepG2 cells. SNX8 is believed to be involved in retrograde transport from endosomes to the trans-Golgi network compartments (van Weering et al., 2012). SCFD1 is involved in transport from the endoplasmic reticulum (ER) to the Golgi apparatus (Dascher and Balch, 1996) and loss of SCFD1 has been found to hinder ER to Golgi transport of extracellular matrix (ECM) proteins during chondrogenesis in Drosophila. Conversely, one could consider that the overexpression of this protein enables the secretion of ECM proteins that are also overexpressed in the HepG2 cell line.

In our study, we highlighted numerous ER proteins. This could be explained by their role in intracellular trafficking, as well as their involvement in several metabolic processes such as lipid synthesis and calcium storage, which are linked to bile excretion and adherens (Gissen and Arias, 2015). Recently, (Chung et al., 2022) uncovered an unexpected role of the cortical ER network and ER-plasma membrane contacts in the regulation of hepatocyte polarity via phospholipid regulation. This finding could also explain the importance of ER proteins in our results.

### Several cytoskeletal proteins appear to be implicated in BC formation and/or function, including Espin, Villin, LimA1, Myo1D, NHE-RF1 and NHE-RF3

Within the **cytoskeleton** group, two proteins known to be involved in microvilli structure, the actin-bundling proteins **Espin** and **Villin**, were highlighted as highly abundant in HepG2/C3A cells in proteomics (Table 1). Western Blot analysis confirmed that Espin was expressed in HepG2/C3A but remained undetectable in HepG2 cells, as well as the lower expression of Villin in the latter (Fig. S2b-c). We carried out IF assays and confirmed that both Villin and Espin were localized in the BC microvilli of HepG2/C3A cells (Fig. 2a-b and Fig. S3a-b). They appeared to colocalize with actin at microvillus.

**Figure 2.**
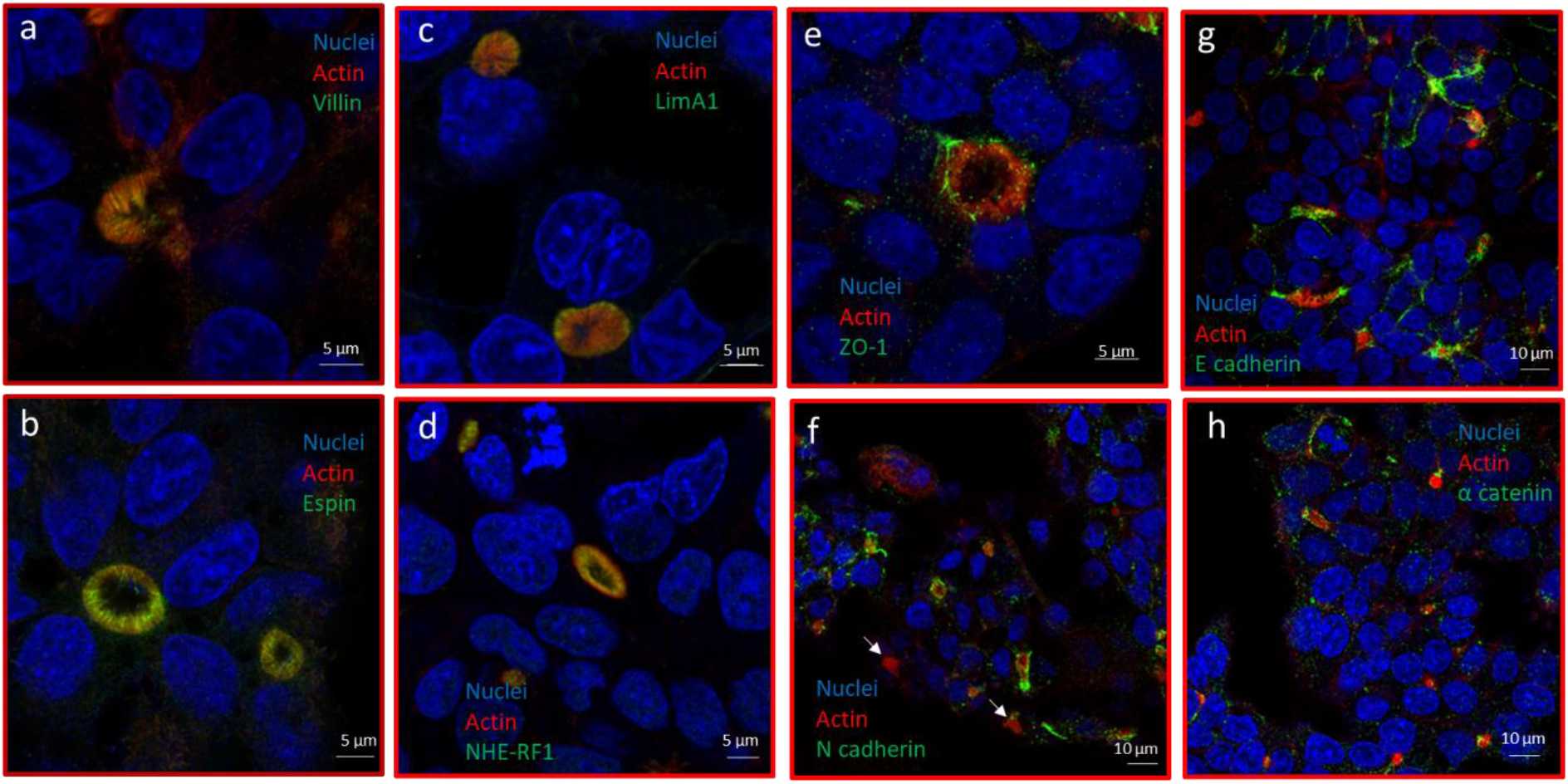
Proteins localized at the BC in HepG2/C3A cells. Fluorescence confocal microscope images acquired at 63x magnification of HepG2/C3A cells immunolabelled with antibody directed against the indicated proteins (a-h), white arrows: BC not surrounded by N cadherin.

Espin, Villin and Fimbrin are described as actin microfilament cross-linkers, necessary for the formation of microvilli (Friederich and Louvard, 2006). These bundling proteins bind to and cross-link actin filaments into parallel actin bundles. Microvilli are crucial in increasing lipid membrane surface area for exchange with BCs and play a crucial role in bile excretion. In various types of cholestasis, abnormal microvilli are commonly observed. Specifically, a decrease in the quantity of canalicular microvilli, as well as thickening of the pericanalicular actin filament zone, are typical of the ultrastructural pathology observed in cholestasis (Phillips et al., 2003). It was previously known that Villin is present in the microvilli of BCs in the liver (Tsukada et al., 1995). (Phillips et al., 2003) already showed an abnormal gene expression of Villin in some patients with progressive cholestasis and hepatic failure. The presence of Espin in BCs, however, had not yet been described. Espins, a family of actin-bundling proteins, have been characterized by Bartles’ group (Bartles et al., 1996). Espins are present in the microvilli of the intestinal and renal brush borders and at the level of Sertoli cell-spermatid junctions. More recently, they demonstrated the crucial role of Espin in the stereocilia of hair cells in the cochlea and vestibular system, by showing that a recessive mutation in its gene causes hair cell degeneration, deafness and vestibular dysfunction in the Jerker Mouse (Zheng et al., 2000). Thus, Espin may play an important role in the BC function. Therefore, we decided to further investigate the role of Espin in this particular context. In order to decipher its role in the liver, we followed BC formation in HepG2/C3A cells, where Espin gene expression was specifically silenced. Western blot analysis confirmed the efficiency of the siRNA approach, with a strongly diminished expression of Espin 48 hours after transfection, which was maintained up to 96 hours (Fig. 3a). Conversely, Espin expression levels after transfection with CTL siRNA remained similar to that of the control without transfection.

**Figure 3.**
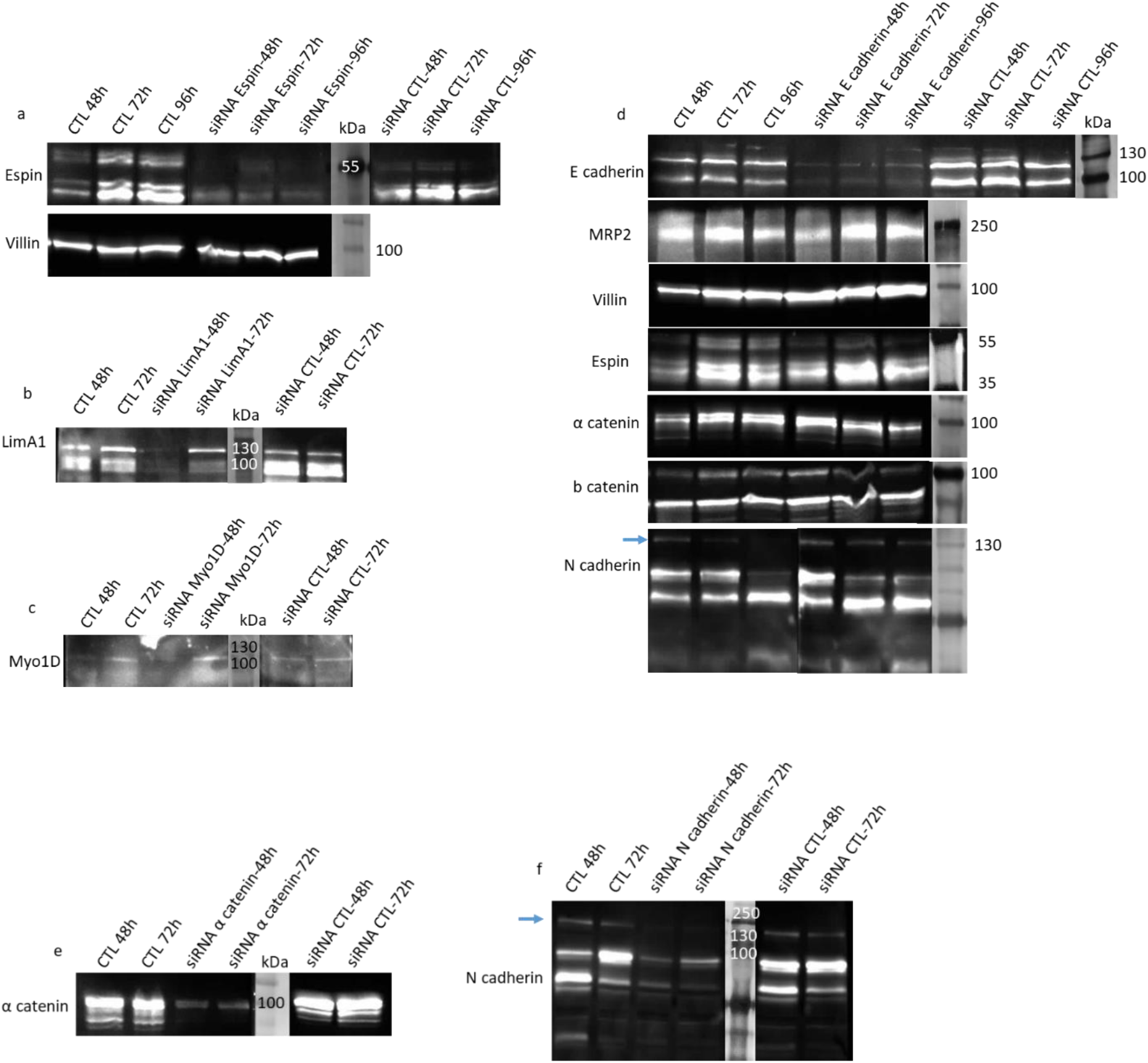
Validation of the efficiency of siRNA in knocking-down protein expression. Representative immunoblots of protein extracts of HepG2/C3A after treatment with the different siRNA at various time of incubation versus siRNA CTL and cells without treatment (CTL), antibody against the protein which is invalidated, a: Espin, b: LimA1, c: Myo1D, d: N cadherin (mature form indicated with the arrow), e: α catenin, f: N cadherin (the blue arrow corresponds to the expected size for N cadherin), for a and d: other proteins of interest were detected, as indicated on the left of the blot.

Using an IF approach with a labelling of actin and MRP2 as markers of BCs, we observed no impact of the absence of Espin on BC formation in terms of the number and presence of microvilli (Fig. S4a). This would suggest that Espin does not play a key role in BC formation.

Using Western blot analysis, we next assessed the expression level of Villin in the absence of Espin (Fig. 3a). Compared with the control, the amount of Villin remained constant over time after transfection.

Proteomics revealed that the expression of another actin-binding protein, **LimA1** (or EPLIN) is five times lower in HepG2 compared to HepG2/C3A cells (Table 1), and this was confirmed by Western-blot (Fig. S2d). IF enabled us to localize LimA1 which is particularly abundant at the periphery of BCs, around the actin ring (Fig. 2c and Fig. S3c), highlighting a potential role of this protein in BC formation or structure. LimA1 has been extensively studied in recent years and was first shown to enhance bundling and stabilization of actin filaments by inhibiting actin filament depolymerisation (Maul et al., 2003). Zang et *al*.’s research, conducted on both mice and humans, successfully identified LimA1 as a crucial protein involved in the regulation of intestinal cholesterol uptake (Zhang et al., 2018). Therefore, we conducted experiments to suppress its expression by transfecting HepG2/C3A cells with a specific siRNA targeting LimA1. This siRNA strongly diminished the expression of its target protein 48 hours after transfection, but its expression was restored 72h post-transfection (Fig. 3b). We did not observe any impact on the formation of BCs, even after 48 hours of LimA1 repression, thus indicating no direct influence of this protein on the establishment of BCs (Fig. S4b).

LimA1 was recently described as mediating linkage of the Cadherin-Catenin complex junction to F-actin, which results in the stabilization of the circumferential Actin belt (Abe and Takeichi, 2008), functioning as a mechanosensor for junction remodelling in epithelial cells and facilitating the interaction between α catenin and vinculin (Taguchi et al., 2011). In the case of the vascular endothelium, LimA1 reinforces the cohesion of cell-cell junctions, by connecting the VE cadherin-catenin complex to the actin cortical ring and promoting vinculin junctional recruitment (Chervin-Pétinot et al., 2012). As we localized LimA1 at the periphery of BCs, we can hypothesize that it could also be a linker between the adherens junction and the actin ring in hepatocytes. However, IF labelling showed a more punctiform distribution for α catenin around BCs than for limA1 (Fig. 2h), also suggesting another role for this protein. The role of LimA1 might affect BC function, but this remains to be elucidated.

We also investigated the role of the protein **Myo1D**, which was scarcely present in the HepG2 cell line and four-fold more expressed in the proteome of the HepG2/C3A cell line (Table 1), which was confirmed by WB (Fig. S2e). The antibody did not yield good results in IF, so we could not localize this protein. Myosins are a superfamily of molecular motors that bind to actin filaments to generate force and movement. In recent years, it was shown that Myo1D, an unconventional myosin, is a chiral determinant, breaking symmetry at all biological scales from molecular to organism level. Its role was first described in drosophila (Hozumi et al., 2006, Lebreton et al., 2018), later in vertebrates such as zebrafish (Juan et al., 2018) and xenopus (Schulze et al., 2019), but its implication in a chiral process in mammals has not yet been described. (Petzoldt et al., 2012) showed that DE cadherin, a core component of adherens junctions, co-immunoprecipitates with Myo1D and is required for Myo1D left-right asymmetric activity in drosophila. Recent data also revealed that a growing number of unconventional myosins play important roles in cell junctions (Liu and Cheney, 2012). For example, drosophila Myo31DF, a member of the Myo1D family interacts and co-localizes with β Catenin, suggesting that these situs inversus genes can direct left-right development through the adherens junction (Spéder et al., 2006). Its potential role in the specific hepatocyte polarization seemed interesting to explore. Therefore, we investigated the impact of its suppression using siRNA. Western blot indicated that the suppression was effective at 48 hours post-transfection, but the protein was re-expressed 24 hours later (Fig. 3c). We did not observe any effect on the number or shape of formed BCs up to 48 hours post-transfection. Altogether, these data did not allow us to identify differences in BC structure (Fig. S4c), and further investigations will be necessary to understand the role of Myo1D in BC formation and/or function. Conversely, one could consider that the overexpression of this protein enables the secretion of ECM proteins that are also overexpressed in the HepG2 cell line.

From our proteomics data, we also observed lower expression in HepG2 cells for two scaffold proteins of the NHERF family, **NHE-RF3** (or PDZK1) and to a lower extent **NHE-RF1** (Table 1). We confirmed this result by WB for NHE-RF1 (Fig. S2f) and localized this protein to BCs by IF, with a labelling on the inner part of the microvilli (Fig. 2d and Fig. S3d), and global co-localization with actin, in particular in the microvilli (Fig. 2d). NHERF scaffold proteins contain several specific protein-binding domains (PDZ modules) that serve to assemble proteins together into functional complexes. They were initially viewed as ‘‘passive linkers’’ between transmembrane proteins and the cortical cytoskeleton underlying the plasma membrane. Very interestingly, in a review from 2011, NHE-RF1 was described beneath the apical pole of both hepatocytes and it was reported to potentially regulate cell surface expression and functional activity of transporters such as MRP2 in hepatocytes and CFTR (cystic fibrosis transmembrane conductance regulator) in cholangiocytes (Clapéron et al., 2011). The interaction of NHE-RF1 with MRP2 in both primary hepatocytes and the WIF-B cell line has been shown to involve its PDZ1 domain and, furthermore, NHE-RF1 was reported to be essential for maintaining the localization and function of MRP2 (Karvar et al., 2014). NHE-RF3 is strongly detected in liver with a subcellular distribution similar to that of NHE-RF1 (Karvar et al., 2014) and using the yeast two-hybrid system, a group has also determined that NHE-RF3 interacts with the carboxy-terminal extremity of MRP2 (Kocher et al., 1999). NHE-RF1 is also described to link radixin (an EMR protein) in hepatocytes microvilli and to interact with β catenin through its PDZ 2 domain (Clapéron et al., 2011). To our knowledge, there is no additional recent literature on this protein family and its role in the liver. Here we confirmed the co-localisation of NHE-RF1 and MRP2 in BC microvilli. This protein could therefore have an impact on both BC structure and the biliary excretion function, underlying the interest of examining in more details its mechanism of action.

**FHOD-1**, another actin binding protein, is also more abundant in HepG2/C3A than in HepG2 cells as revealed using proteomics (Table 1). It belongs to the formin family, which forms a group of eukaryotic proteins known for their role as organizers of the cytoskeleton, regulating processes such as morphogenesis, cell polarity, and cytokinesis. However, since we were unable to validate the proteomic result through WB or localize the protein via IF, we did not pursue further our investigation.

### Focal adhesion and ECM

These proteins were also well represented in our differential proteomics dataset, which is in accordance with the literature on their implication in polarisation. In HepG2 cells, various excreted proteins of the ECM, such as Fibronectin and different Laminins, were more abundant than in HepG2/C3A cells (Table 1). Mature hepatocytes are surrounded by an ECM devoid of Laminin, which contrasts with its presence during hepatocyte differentiation. The high abundance of Laminins in HepG2 cells is in accordance with their lower level of polarisation. Conversely, HepG2/C3A cells exhibited a higher level of integrin transmembrane receptors and of the adaptor proteins Paxillin and Vinculin, present in focal adhesion plaques. Vinculin acts as a linker between Integrins and the actin cytoskeleton. Paxillin can bind directly Integrins but not actin filaments and is one of the binding partner of Vinculin (Turner et al., 1990).Vinculin is also known to be involved in cell-cell adhesion and to bind α and β catenin (Bays and DeMali, 2017). It was also shown that Vinculin cooperates with LimA1 to maintain adherens junction (Taguchi et al., 2011). We confirmed by western-blot the lower level of expression of Paxillin in HepG2 cells (Fig. S2g).

The ECM serves as a mechanical framework for cell adhesion and movement, while also functioning as a signalling platform by sequestering or releasing cytokines. Direct interaction between ECM molecules and cell surface receptors, particularly Integrins, triggers activation, initiating intracellular signalling pathways. Intracellular signalling events can influence the interaction of Integrins with the ECM, resulting in an intricate feedback of signalling cascades. A main process that can be controlled by interactions between the ECM and cell surface receptors is epithelial cell polarization (Treyer and Müsch, 2013). In classical columnar polarization, the presence of a basal lamina causes cells to have their apical poles in the opposite direction. However, for hepatocytes, the presence of low density ECM (without a basal lamina) on both sides of the cells, within the Disse spaces, prompts cells to establish apical poles by interacting with neighbouring cells. In *in vitro* settings, primary hepatocytes polarize effectively when cultured in a collagen sandwich, replicating the *in vivo* situation (Dunn et al., 1991). In our study, the difference in ECM composition between the two cell lines and the higher expression of focal adhesion proteins in HepG2/C3A cells appear consistent.

### Cell-cell adhesion: E cadherin, an adherens junction protein essential for inducing BC formation

The fact that proteins of cell-cell adhesion, including adherens junction, were well represented in the differential proteomics dataset comparing HepG2 to HepG2/C3A cell lines, with the notable absence of both E and N cadherins in HepG2 (Table 1). The role of cell junctions in maintaining epithelial cell polarization is known. However, much remains to be elucidated about the establishment of BCs by hepatocytes, in particular the implication of tight versus adherens junctions (Treyer and Müsch, 2013). For tight junctions, proteomics and WB analyses revealed a similar expression level between the two cell lines of the key tight junction protein ZO-1, which cross-links and anchors tight junction to the actin cytoskeleton (Fig. S2h). Through IF, we localized ZO-1 in dense spots around the actin ring of BCs formed in the HepG2/C3A cell line (Fig. 2e, S5b). In the case of the HepG2 cell line, in absence of BCs, the protein ZO-1 formed lines between two cells in certain areas of the plasma membrane (Fig. S5a). Among the different types of integral membrane proteins in tight junctions, we confirmed the lower expression of JamA in HepG2 cells (Fig. S2i) with a localization similar to that of ZO-1 in both cell lines (Fig. S5c-d). The proteomics approach highlighted Claudin 1 and Claudin 6, two proteins of the Claudin family (small proteins around 20 kDa), to be expressed at similar levels in the two cell lines and two times more in the HepG2/C3A cell line, respectively. Using WB, Occludin was also shown significantly less expressed in HepG2 cells (Fig. S2j). In conclusion, tight junctions showed significant differences between the two cell lines, with similar expression levels but different localization of the intracellular ZO-1, and lower expression of the integral membrane protein Claudin 6.

For adherens junctions, we confirmed by WB the near absence of **E** and **N cadherin** in the HepG2 cell line (Fig. S2k-l), as well as the lower expression of **α catenin** and to a lesser extent of **β catenin**, (Fig. S2m-n). In the case of N cadherin, WB showed a faint band at the expected size of 130 kDa and two very intense bands at lower molecular weights. It is challenging to determine whether this is due to an artefact of protein extraction or the presence of truncated forms. Via IF analysis, E and N cadherin were localized at the membrane of HepG2/C3A cells, between some adjacent cells, and N cadherin could be found surrounding some BCs but not others (white arrows on Fig. 2f show some BCs which are not surrounded by N cadherin, whereas other BCs show a crown of N cadherin in the same field of view). E cadherin however was always present on both sides of BCs (Fig. 2g). α catenin showed a distribution pattern closer to that of E cadherin but with a less intense labelling (Fig. 2h).

The absence of cadherins in HepG2 cells prompted us to further investigate their role in BC formation, focusing first on E cadherin. E cadherin’s role in epithelial cell junctions is much more extensively documented than that of N cadherin, which is primarily studied in neuronal cells. We tracked BC formation in the HepG2/C3A cell line subsequently to protein repression via specific siRNA targeting E cadherin. WB analysis showed that E cadherin was almost absent 48 hours after siRNA transfection and remained so until 96 hours (Fig. 3d), while the level of expression of E cadherin after transfection with the CTL siRNA was similar to that of the control without transfection. In absence of E cadherin, no significant changes were observed in the expression level of partner proteins such as N cadherin, α and β catenin and some of the proteins previously identified as differentially expressed between HepG2/C3A and HepG2 cells such as Espin, Villin and MRP2 (Fig. 3d). Western blotting of N cadherin appeared to show a slight induction of its 130 kDa form while E cadherin was silenced.

Using our IF approach, we observed a strong impact of the absence of E cadherin on BC formation. Indeed, 48 fours after transfection with the siRNA, the HepG2/C3A cells almost lost their capacity to form BCs (fig. 4). IF of MRP2, and actin-labelled cells closely resembled those obtained with the HepG2 cell line (Fig. S1a).

**Figure 4.**
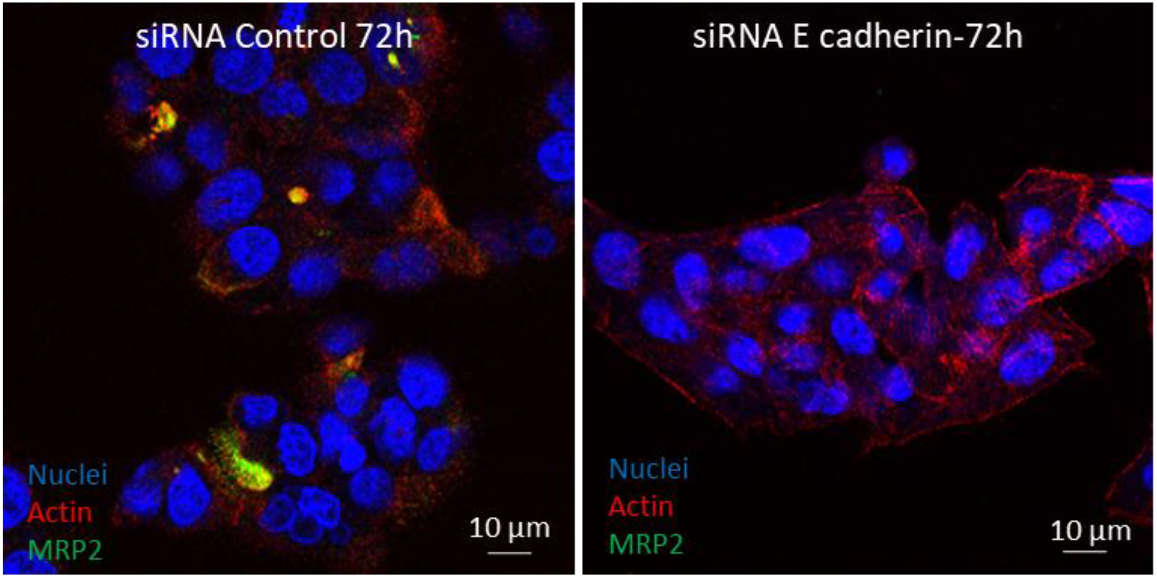
BC disruption in absence of E cadherin. Immunofluorescence labelling of BC (MRP2, actin) in HepG2/C3A cells upon treatment with control versus E cadherin-targeting siRNA. Confocal fluorescence images acquired at 63x magnification.

To ascertain whether adherens junctions trigger BC formation or only E cadherin by itself, we conducted similar experiments by knocking-down α catenin expression. Despite a strong repression of its expression (Fig. 3e), we did not observe any impact on the formation of BCs (Fig. S4d). This result is surprising given the key role of α catenin in connecting the E cadherin–β catenin complex to the actin cytoskeleton. We thus might have expected differences in the shape of the pericanalicular actin ring. However, it is conceivable that tight junctions, which are also linked to the actin cytoskeleton, allow the maintenance of the actin ring within the timeframe of our observation. Moreover, α catenin is known to have not only a structural role but is also important in coordinating actin dynamics (Kobielak and Fuchs, 2004). However, this new role of α catenin does not appear to be involved in the formation of BCs.

This confirms the particular role of E cadherin in the establishment of BCs. To delve deeper, we also investigated the impact of suppressing N cadherin expression. Western blot analysis reveals a significant reduction in protein quantity at 48 and 72 hours post-transfection, particularly for the active form (Fig. 3f indicated by an arrow). However, no effect on BC formation was observed (Fig. S4e).

At the organism scale, tissues can be subdivided into five groups according to the expression profile of E and N cadherin (Tsuchiya et al., 2006): (1) tissues that express both E and N cadherin at the same subcellular location (liver and thymus), (2) tissues that express both E and N cadherin but with different subcellular location (the pituitary, kidney, pancreas, stomach, lung, testis, prostate, mammary gland, salivary gland and endometrium), (3) tissues that express N cadherin but not E cadherin (the cerebrum, cerebellum, peripheral nerve, heart, ovary and adrenal), (4) tissues that express E cadherin but not N cadherin (the epithelium of the thyroid, tonsil, esophagus, duodenum, small intestine, colon, gall bladder, urinary bladder, epidermis and epidermal appendages), and (5) tissues that express neither E nor N cadherin (skeletal muscle, smooth muscle, fat and fibroblasts). As the liver expresses both E and N cadherin at the same subcellular location, this raises questions about the respective roles of E and N cadherins in hepatocytes. Our study demonstrates that they are not exchangeable. In the literature, the role of N cadherin has been much less studied in a healthy context and rather in the context of epithelial-mesenchymal transition (EMT). Importantly, E and N cadherin both play a role in EMT as well as in mesenchymal-epithelial transition which are important processes during embryogenesis or tissue regeneration but also in pathological processes, notably tissue fibrosis, tumor invasiveness and metastasis (Choi and Diehl, 2009).

Interestingly, we previously demonstrated that the localization of E and N cadherin is not strictly similar, with E cadherin alone being consistently present around bile canaliculi. In accordance with this observation, we also demonstrated that their roles are not interchangeable in hepatic tissue, and that only E cadherin seems to play a key role in the initiation of BC formation.

To further demonstrate significant effects of the different siRNAs, we acquired images of various randomly selected fields at lower magnification (20x) on a confocal microscope. We then semi-automatically counted the number of BCs (labelled with MRP2) and the number of nuclei (Hoescht labelling). The ratio of the number of BCs over the number of cells was drastically low only when E cadherin was silenced (Fig. 5), confirming its key importance for BC formation.

**Figure 5.**
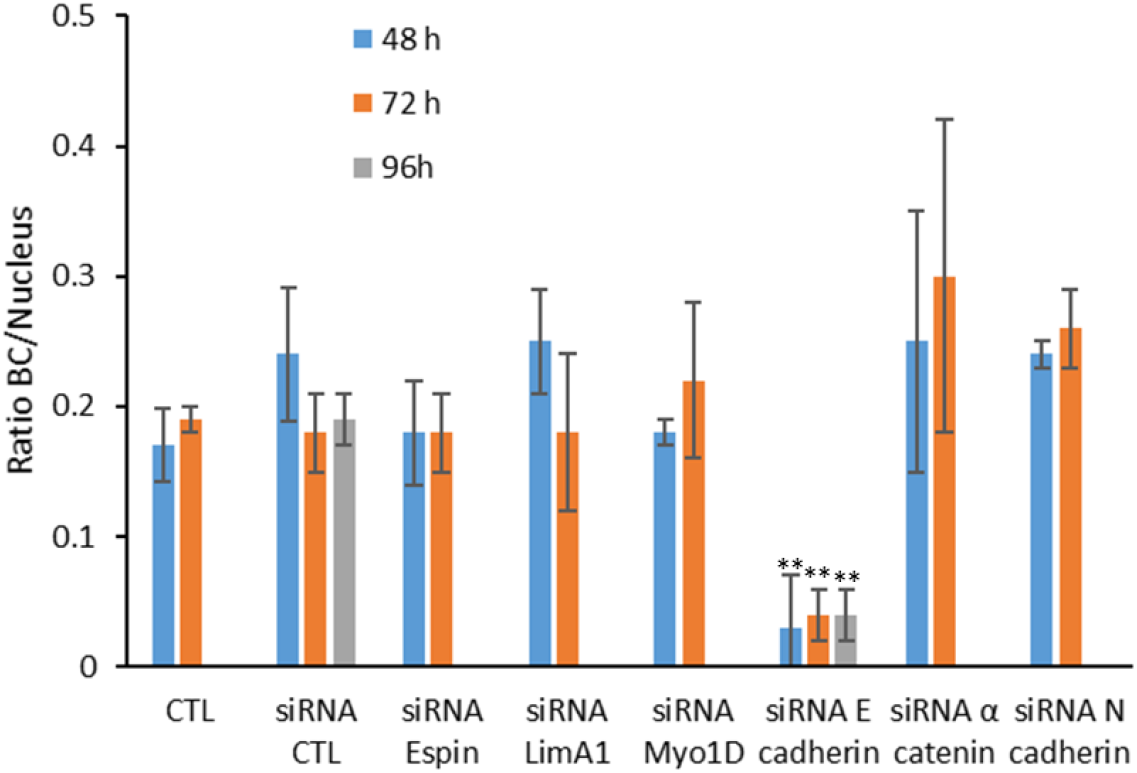
Effect of selected protein repression on BC formation. The expression of selected proteins was repressed in HepG2/C3A cells using specific siRNA. Nuclei and BC were stained and confocal fluorescence images acquired at 20x. Nuclei and BC were segmented from confocal fluorescence images in FiJi (ImageJ)(Schindelin et al., 2012). Results are expressed as the ratio of the number of BC to the number of nuclei. Each value represents the mean of relative expression ± SD from three independent experiments. **: p<0.01 vs control.

**To follow the early stages of BC formation**, we studied the effect of siRNA E cadherin, in 2D culture, **during the first 48 hours**. We showed earlier that repression was nearly total at 48 hours. However, western blot analysis (Fig. 6a) revealed overexpression of E cadherin precursor (130 kDa) 5h after transfection with both control and E cadherin-targeting siRNAs. Cell detachment and resuspension before transfection disrupt extracellular proteins of the junctions and likely trigger the neo-synthesis of proteins necessary for the formation of new junctions upon replating. This could explain our result. Twenty-four hours after transfection, siRNA was effective and E cadherin expression levels decreased drastically as expected given its half-life time of around 5 hours (Shore and Nelson, 1991). In control cells, the band corresponding to the mature form of E cadherin increased and the one corresponding to the precursor decreased, as seen previously, starting from 48 hours. After labelling cells for actin and MRP2, we checked the presence of BCs 5 hours after transfection and replating. At this early stage, cell density was low but we found a few labelled structures, which looked like immature BCs with actin organized as patches in comparison with a mature BC presenting an organized actin ring (Fig. 6b). We were unable to consistently quantify the number of BCs due to low cell density during short time intervals and variability in kinetics up to 24 hours but these structures were visible in both control and siRNA treated samples. This finding is in accordance with the presence of E cadherin at this early stage. We can hypothesize that the few BCs observed after 48 hours of transfection were established very early, when E cadherin was still present to initiate their formation.

**Figure 6.**
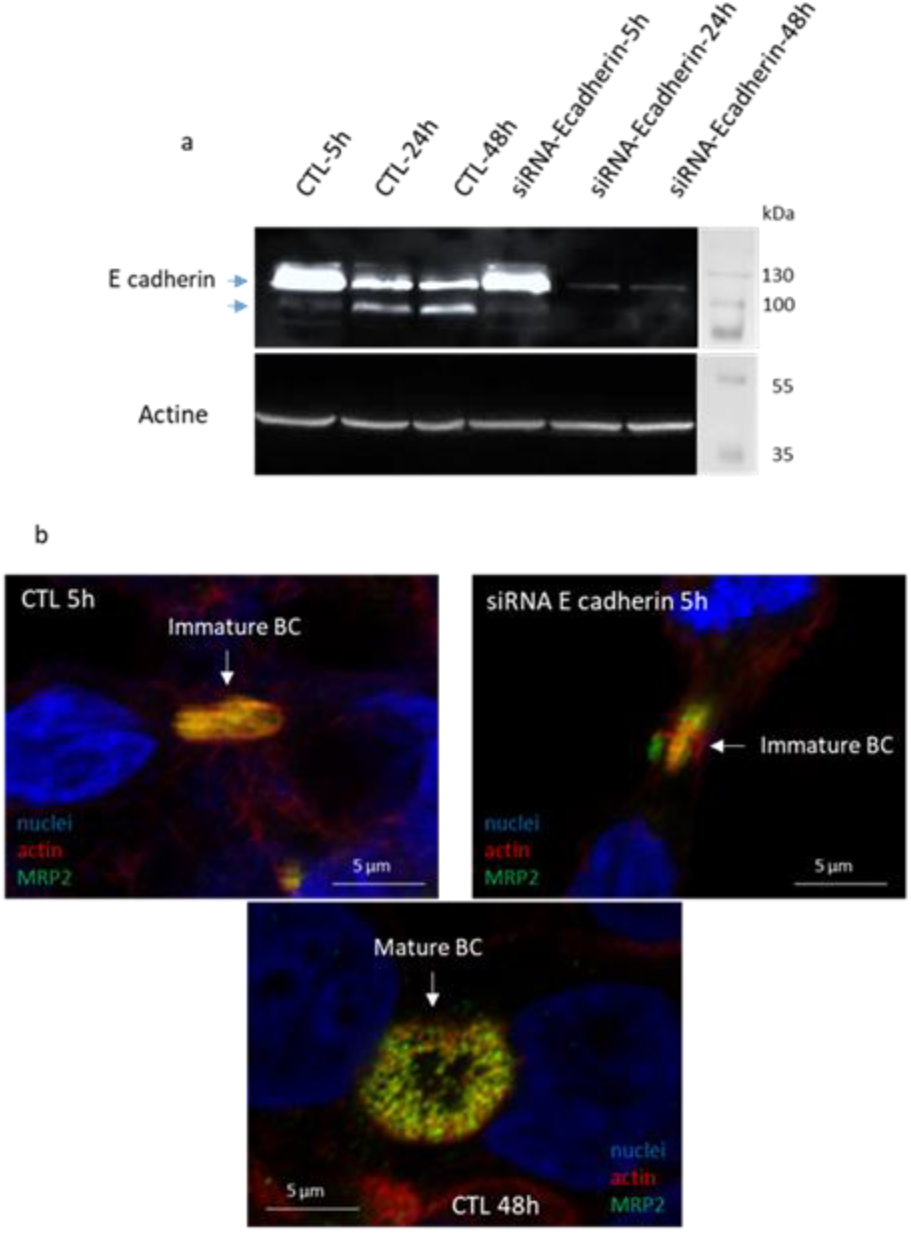
Impact of E cadherin repression on early stages BC formation. (a) Representative immunoblots of protein extracts of HepG2/C3A after treatment with the E cadherin siRNA versus CTL at various time of incubation (b) Immunofluorescence labelling of BC (MRP2, actin) in HepG2/C3A cells upon treatment with control (CTL) versus E cadherin-targeting siRNA. Confocal fluorescence images acquired at 63x magnification in airyscan mode.

In light of these interesting results, we **analysed the effects of E cadherin repression in a 3D spheroid model**. We have previously shown that when grown in 3D as matrix-free hepatic spheroids, HepG2/C3A cells develop more mature and functional BCs than in 2D cell culture (Sharma et al., 2020). We have also developed a highly parallelized on-chip method to fabricate hundreds of hepatocyte spheroids regularly arrayed on a glass coverslip in order to easily study BC formation by immunofluorescence imaging (Fig. 7a). We carried out E cadherin down-regulation experiments on this spheroid model and confirmed that this protein is also essential for BC formation in a 3D model (Fig. 7b-c). Grown in spheroids, HepG2/C3A cells can be maintained in culture longer and recovery after removing siRNA can therefore be monitored. We showed that after 3 days of recovery the restoration of E cadherin expression triggered BC formation in the spheroids (Fig. 7b-c). This result confirmed that the expression of E cadherin is essential, and could be one of the triggers for BC formation.

**Figure 7.**
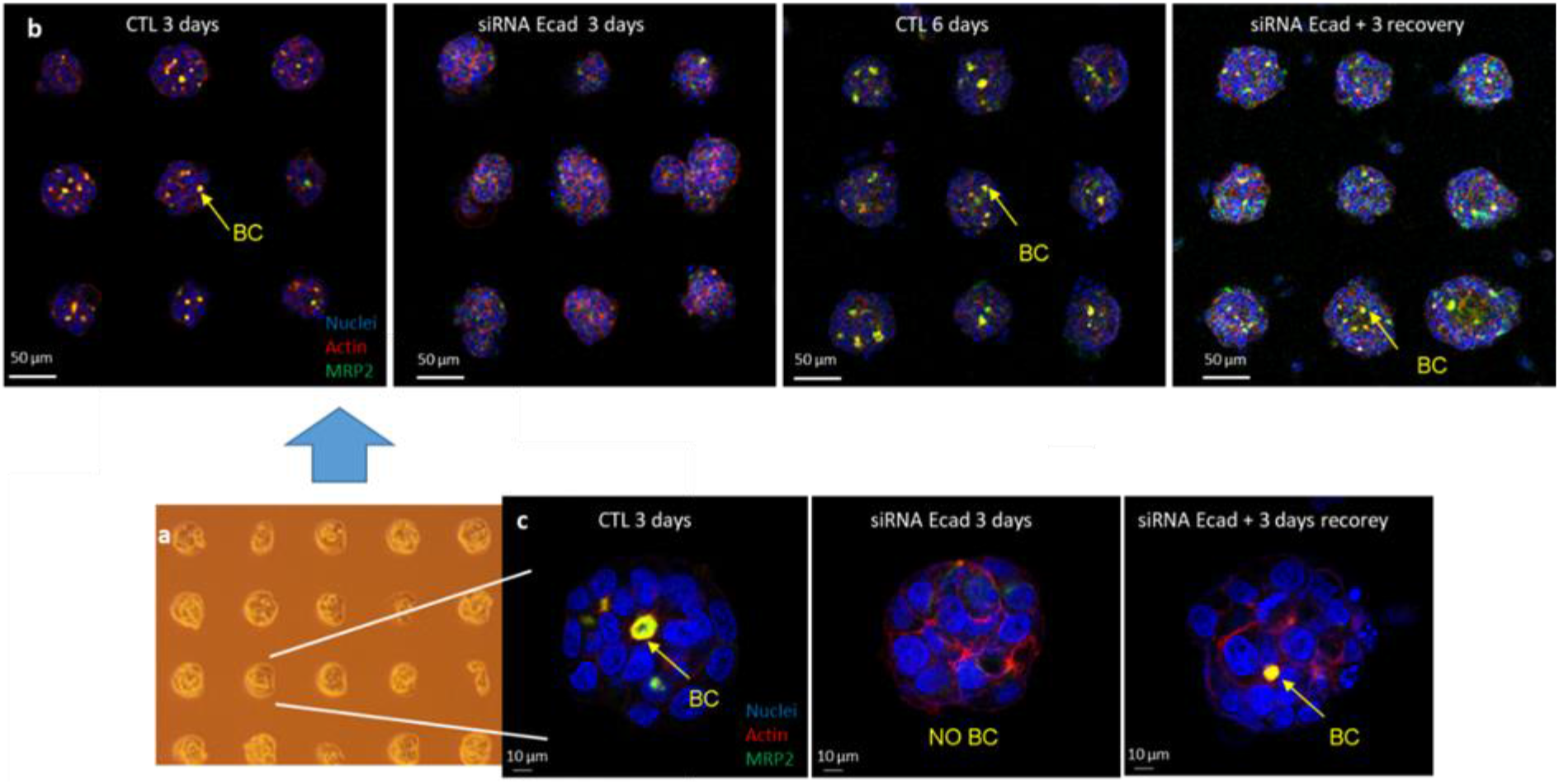
Validation of the role of E cadherin in hepatocyte spheroids. (a) On-chip HepG2/C3A spheroids. Immunofluorescence labelling of BC (MRP2, actin) in HepG2/C3A spheroids after 3 days of treatment with control (CTL) versus E cadherin-targeting siRNA and after 6 days treatment with CTL siRNA or 3 days with E cadherin-targeting siRNA + 3days recovery. Confocal images were acquired at 20x magnification (b) and at 63x (c).

The important role of adherens junctions and especially of E cadherin in BC formation demonstrated here is also supported by the fact that several of the proteins we have studied (Vinculin, NHE-RF1), particularly those with poorly understood roles in BCs (LimA1, Myo1D), are known to be associated with adherens junctions, as mentioned previously.

At the boundary of BCs, while adherens junctions mediate mechanical linkage between neighbouring hepatocytes, tight junctions regulate the blood-biliary barrier, which prevents the leakage of molecules between BC lumina and sinusoids. In the literature, both tight and adherens junctions are highlighted for their important role in BC structure and maintenance (Gissen and Arias, 2015, Treyer and Müsch, 2013). Compared to these previous data, our data clearly demonstrate that E cadherin is critically involved in BC formation. To follow BC formation, Virgile Viasnoff’s group (Li et al., 2016) has developed artificial microniches to accommodate two or just a few primary hepatocytes in a highly defined microenvironment (various ECM or E cadherin coating configurations). This work provides strong evidence that environmental scaffolding by ECM adhesion plays a guidance role in the elongation process of BCs. They notably showed that ECM scaffolding influences the directed elongation of intercellular lumens. Accordingly, we identified numerous proteins involved in focal adhesion. In addition, they observed that seeding hepatocytes in wells coated with E cadherin (walls) and fibronectin (bottom) resulted in thin and large tubular BCs. With this microniche model, they also demonstrated no drastic effect of α catenin expression inhibition on BC formation, a result in accordance with our data. However, they did not test E cadherin expression inhibition in this former study, but recently published a potential crucial role of E cadherin in the initiation of BC formation (Zhang et al., 2020). Indeed, they patterned single primary hepatocytes on E cadherin coated small motifs, in order to mimic cell-cell interaction and they demonstrated that this is sufficient to initiate the spontaneous formation of a hemi lumen (hemi BC) between the cell and the coated E cadherin motif. They detailed the different phases of BC formation in this model, starting with the appearance of a diffuse actin patch at 4 hours in the middle of the contact area that finally organizes into an actin ring after 14 hours. At 4 hours, the attachment of the cell’s cadherins to the substrate remains homogenous over the entire contact area, however, from 7 hours, in parallel of hemi-lumen formation, cadherins detach from the underlying substrate beneath the central patch region. However, in this study, they used a pan-cadherin antibody for immunofluorescence, which targets both E and N cadherin, making it impossible to differentiate between the two cadherins. Finally, using this simplified model, E cadherin was found to be the minimal element to trigger apical hemi lumen formation, with the role of N cadherin remaining unclear.

## Conclusions

In this study, we confirmed the implication of various classes of proteins in the BC structure such as proteins of focal adhesions, cell-cell junctions, the cytoskeleton, cell trafficking machinery and metabolism. Based on a non-targeted extensive proteomic approach, we identified proteins such as Espin, LimA1 and potentially Myo1D, which had not been previously described to be localized in BCs and we also shed light on the involvement of NHE-RF1 and NHE-RF3. Our siRNA approach did not allow us to identify their potential roles in BC formation. We can hypothesize that either their roles are not crucial for BC formation or other proteins with similar functions compensate their absence. We raised the question of the roles of all these proteins in BC formation, function and structure.

In classical 2D cultures and in a 3D model, our work demonstrates the specific and crucial role of E cadherin in the initiation of BC formation. Although, only a few patches of actin were observed at five hours post-silencing, which likely corresponds to the first step of BC formation, after the total depletion of E cadherin, hepatocytes were no longer capable of initiating the formation of BCs. As N cadherin and α catenin suppression did not show the same impact on BC formation and because the restoration of E cadherin expression allowed re-initiation of BC formation, we propose that E cadherin can orchestrate BC formation without the other adherens junction proteins.

## Materials and methods

### Cell lines and cell culture

HepG2 and HepG2/C3A cell lines were obtained from the American Type Culture collection (ATCC) (Manassas, Virginia, USA). They were grown in modified Eagle’s medium (MEM) supplemented with 10% v/v FBS, 20 mM L-glutamine, 10 mM sodium pyruvate, 100 µg/mL streptomycin and 100 U/mL penicillin. Cells were cultured at 37°C in a humidified atmosphere with 5% CO2.

### Spheroids on chip culture

Micropatterns were produced on glass slides by deep-UV photolithography that degrades a non-toxic cell repellent PEG-PLL coating locally thanks to a photomask (as described in (Azioune et al., 2010)). Micropatterns were then coated with fibronectin (20 μg/ml in PBS) to form adhesive disks of 50 or 70 μm in diameter, surrounded by a cytophobic surface with a gap between disks of 100 or 150 µm. HepG2/C3A cells were seeded on micropatterned surfaces at a density of 0.5.10^5^ cells per cm^2^ (0.5 million in a well of a six wells plate) and were allowed to attach.

Cells on micropatterns were maintained in culture for up to one week, to allow spheroids to grow.

### siRNA forward transfection

siRNA experiments were performed in 24 well plates with 12 mm glass coverslips for IF labelling, in 6 well plates for micropattern chips (20x20 mm glass coverslips) and in 60 mm dishes for protein extraction according to the following protocol (Table 2).

**Table 2.** Volume of reagents for siRNA forward transfection.

|  | 24 well plate | 6 well plate | 60 mm dish |
| --- | --- | --- | --- |
| Opti-MEM | 100 µl | 500 µl | 1 ml |
| lipofectamine RNAiMax | 1 µl | 5 µl | 10 µl |
| siRNA (10μM) | 1 μl | 5 μl | 10 μl |
| Cells (10<sup>6</sup>) | 0.15-0.2 | 0.5-1 | 1.5-2 |

A same volume of lipofectamine RNAiMax (Invitrogen, thermos Fisher Scientific) and siRNA solution Santa Cruz Biotechnology: siRNA CTL (sc-37007), siRNA E cadherin (sc-35242), siRNA N cadherin (sc-29403), siRNA LimA1 (sc-60593), siRNA α catenin (sc-29190), siRNA Myo1D (sc-44608), siRNA Espin (sc-78697)) at 10 µM were mixed in opti-MEM medium (Gibco, Thermo Fisher Scientific) and incubated for at least 30 min. Then, transfection mix was deposited on the well or the dish and cells were added at the attempted concentration in MEM supplemented with 5% v/v FBS, without antibiotics. The cells were cultured in the transfection medium during the time of the experiment (48, 72 or 96h). In the case of recovery, the medium was removed and replaced with regular medium after washing with PBS.

### Immunofluorescence

All the steps were made at room temperature. The cells grown as 2D or 3D culture on glass coverslips were fixed with a formalin solution (Sigma) for 15-20 min at room temperature and washed three times with phosphate-buffer saline (PBS). Fixed cells were permeabilized with 0.5% Triton X-100 and 0.05% Tween-20 in PBS for 15 min for 2D culture and 60 minutes for 3D culture and then blocked to prevent the non-specific binding of the antibodies, with a solution 3% BSA, 0.1% Triton X-100, 0.05% Tween20 in PBS for 1 hour. The cells were then incubated with primary antibodies (listed in Table 3) diluted in solution 1% BSA, 0.1% Triton X-100, 0.05% Tween20 in PBS for 1 hour at room temperature. After PBS washes, Phalloïdin-TRITC and secondary antibody Alexa Fluor 488 anti-mouse or anti-rabbit (1:1,000; Thermo Fisher Scientific) were incubated for 1 h at room temperature, followed by final PBS washes. Hoechst (1 μg/ml in PBS) was added in the last wash. The coverslips were mounted on glass slides with mounting medium (Sigma) and sealed with varnish. Images were collected at room temperature on a Zeiss LSM880 confocal laser scanning microscope equipped with a Zeiss Plan-APO ×63–numerical aperture 1.46 oil immersion objective and a Plan-Apo x20 0.8NA.

**Table 3.** List and conditions for using antibodies in IF and WB.

| Antibody against | reference | Dilution in IF | Dilution in WB |
| --- | --- | --- | --- |
| MRP2 | Abcam 3373 | 1/50 | 1/50 (Protein extracts not boiled) |
| Paxillin | Santa Cruz 365379 | 1/50 | 1/500 |
| Espin | Gift from Bartles JR | 1/10 | 1/200 |
| Villin | Santa Cruz 58897 | 1/50 | 1/500 |
| LimA1 | Santa Cruz 136399 | 1/50 | 1/500 |
| Myo1D | Santa Cruz 515292 | ∅ | 1/100 |
| NHE-RF1 | Santa Cruz 36271552 | 1/50 | 1/500 |
| E cadherin | Takara M106 | 1/50 | 1/500 |
| N cadherin | Santa Cruz 59987 | 1/50 | 1/500 |
| $\alpha$ catenin | Santa Cruz 9988 | 1/25 | 1/500 |
| ZO-1 | Invitrogen 40-2200 | 1/50 | 1/500 |
| Jam A | Sigma SAB4200468 | 1/50 | 1/500 |
| Occluding | Sigma WH0004950M1 | 1/50 | 1/250 |
| $\beta$ catenin | BD transduction 610154 | 1/25 | 1/1000 |
| $\beta$ actin | A5441 | ∅ | 1/5000 |
| Phalloïdin-TRITC | Sigma P1951 | 1/1000 | ∅ |

### IF images statistical analysis

For each transfection condition, a minimum of three field of view were imaged without bias (based on nuclei) at 20x magnification. Each multi-colour image are converted to jpeg by separating the channels using the Zen Blue software, in order to obtain an image for nucleus, an image for BC and one for actin labelling. The analysis is then conducted using the FiJi (ImajeJ) software(Schindelin et al., 2012). For each field of view, the image corresponding to nuclear staining and the one corresponding to CB staining are transformed into binary images after thresholding. The “particle analysis” module is then used to count objects based on their sizes and circularity indices (nuclei and BC). The mean and the standard deviation of, at least three ratio BC on nuclei, were then calculated using Excel. This was performed for at least three independent experiments. Finally, the mean and standard deviation of these three independent experiments were calculated and statistical student t-tests were run using Excel.

### Western blotting

Cells were rinsed twice with PBS (pellets could be frozen at - 80°C for subsequent extraction) and lysed on ice in at least 10 times the estimated volume of the cell pellet in the extraction buffer RIPA (10 mM HEPES, pH 7.5, 3 mM MgCl2, 40 mM KCl, 2,5% Glycerol, 1% triton X-100 and supplemented (freshly added) with protease inhibitors) for 20-30 minutes. The lysate was then centrifuged at 15,000 rpm for 30 minutes at 4°C. The supernatant was collected and protein concentration was measured using the Bradford assay. The samples were then diluted in Laemmli buffer (to improve storage) and stored at - 20°C. Fifty μg of proteins for each extract was boiled, separated by SDS-PAGE and then electrotransferred (Bio-Rad system) onto nitrocellulose membranes (Bio-Rad). Membranes were stained with indian ink (1/1000) to evaluated the migration and verified the quantity of proteins between samples. They were blocked with milk 3% in PBS - 0.1% Tween for 1 hour or overnight. They were then probed with primary antibodies (see Table 3) at room temperature for at least 1 hour. This was followed by incubation with appropriate horse-radish peroxidase-conjugated secondary antibodies. Bands were detected by chemiluminescence (ECL, Pierce) using a Fusion Fx7 apparatus (Vilbert Lourmat).

### Shotgun proteomics

#### Sample preparation

HepG2 and HepG2/C3A cells were seeded in 100 mm diameter culture plates at a density around 60 000 cells per cm^2^, in complete culture medium. Cells were allowed to grow up to 80% confluency. Briefly, after removing the medium and washing with PBS buffer, cells were collected using 0.25% trypsin, 1 mM EDTA, centrifuged at 1,200 rpm for 5 min. The cell pellets were then rinsed three times with PBS, and finally lysed in 200 µl of lysis buffer (7M urea, 2M thiourea, 4% CHAPS, 15mM spermine base, 15 mM spermine tetrahydrochloride, 10 mM tris(carboxyethyl) phosphine hydrochloride) at room temperature. After extraction for 1 hour, the extracts were clarified by centrifugation (15,000 g 30 minutes, 4°C). The supernatants were collected and their protein concentration measured by a dye-binding assay (Bradford, 1976). The extracts were kept frozen at -20°C until use.

For shotgun proteomic analysis, protein extracts (50 µg) were included in polyacrylamide gel plugs (20 µl) as described in (Cavazza et al., 2022), using photopolymerization (Lyubimova et al., 1993) to effect acrylamide polymerization. After fixing and rinsing the gel plugs as described previously (Cavazza et al., 2022), they were washed three times with 50 µL of 25 mM ammonium hydrogen carbonate (NH4HCO3) and 50 µL of acetonitrile. The cysteine residues were reduced using 50 µL of 10 mM dithiothreitol at 57°C and alkylated using 50 µL of 55 mM iodoacetamide. After two washes with NH4HCO3 and acetonitrile, gel plugs were dehydrated using acetonitrile. Protein digestion was performed in gel, overnight at room temperature, using 15 µL of 7 ng/µL of modified porcine trypsin (Promega, Madison, WI, USA) in 25 mM NH4HCO3. The peptides obtained were extracted with 50 µL of 60% acetonitrile in 0.1% formic acid. Acetonitrile was then evaporated under vacuum and samples were resuspended in a solution of 1% acetonitrile and 0.1% formic acid in order to obtain a final concentration of 150 ng/µL.

### Mass spectrometry analysis

NanoLC-MS/MS analysis was performed using a nanoACQUITY Ultra-Performance-LC (Waters Corporation, Milford, USA) coupled to a Q Exactive HF-X Hybrid Quadrupole-Orbitrap Mass Spectrometer (Thermo Fisher Scientific, Bremen, Germany).

The peptide digest (300 ng) was trapped on a nanoEase^TM^ M/Z Symmetry C18 trap precolumn (100Å, 5 µm, 180 µm x 20 mm, Waters) and the peptides were subsequently separated on a nanoEase^TM^ M/Z Peptide BEH C18 column (130Å, 1.7 µm, 75 µm × 250 mm, Waters). The solvent system consisted of 0.1% formic acid in water (solvent A) and 0.1% formic acid in acetonitrile (solvent B). Peptide trapping was performed during 3 min at a flow rate of 5 µL/min with 99% A and 1% B. Peptide elution was performed at 60°C at a flow rate of 400 nL/min from 1 to 8% of B in 2 min then from 8 to 35% of B in 77 min.

MS and MS/MS acquisitions were carried out in positive mode, with the spray voltage set to 1800 V and the capillary temperature to 250°C. The MS scan was acquired with a resolution of 60,000, an AGC target value of 3×10^6^ and a maximum injection time of 50 ms over m/z [300-1800] range. The MS/MS scans were acquired with a resolution of 15,000, an AGC target value of 1×10^5^ and the maximum injection time was 100 ms with fixed first mass of 100 m/z over m/z [200-2000] range and isolation window of 2 m/z. Lock-mass option was enabled (polysiloxane, 445.12002 m/z). Top 20 HCD was selected with intensity threshold of 2×10^5^ and dynamic exclusion of 60 s. Peptide match selection option was turned on. The normalized collision energy (NCE) was fixed at 27 V. The complete system was fully controlled by Thermo Scientific™ Xcalibur™ software (v4.0). Raw data were converted into mgf files using the MSConvert tool from ProteomeWizard (v3.0.6090; http://proteowizard.sourceforge.net/).

### Protein identification and quantification

For protein identification, MS/MS data were searched using a local Mascot server (v2.6.2; Matrix Science, London, UK) against a database containing the sequences from all *Homo sapiens* entries as found in UniProtKB/SwissProt (version 2019_09, 20,410 sequences) and the corresponding 20,410 reverse sequences. The database was generated using MSDA software (Carapito et al., 2014). Spectra were searched with a mass tolerance of 10 ppm for MS and 0.07 Da for MS/MS data, allowing a maximum of one trypsin missed cleavage. Acetylation of protein N-termini, carbamidomethylation of cysteine residues and oxidation of methionine residues were specified as variable modifications. Identification results were imported into Proline v2.1 (www.profiproteomics.fr/proline) for validation. Peptide Spectrum Matches (PSM) with pretty rank equal to one and with a minimum sequence length of 7 were retained. False Discovery Rate was then optimized to be below 1% at PSM level using Mascot Adjusted E-value and below 1% at the protein level using the Mascot score. Peptide abundances, were extracted using a m/z tolerance of 10 ppm. Alignment of the LC-MS runs was performed using Loess smoothing. Cross Assignment was performed between all runs. Protein abundances were computed as the mean of all peptide abundances (normalized using the median). Peptides sharing peakels were discarded.

The mass spectrometry proteomics data have been deposited to the ProteomeXchange Consortium via the PRIDE (Perez-Riverol et al., 2022) partner repository with the dataset identifier PXD052062 and 10.6019/PXD052062.

### Statistical and pathway analysis

Differentially expressed proteins were identified via a Mann-Whitney U test, with a p value lesser than 0.05 (i.e. U=0 in the Mann-Whitney statistics). Pathway analysis was performed using the Uniprot and QuickGO databases, and we finalized the classification of proteins based on their Gene Ontology (GO) terms by hand to achieve a level of detail unattainable using conventional algorithm.

## Acknowledgments

We thank Bartles JR for kindly providing the antibody against espin. We thank the microscopy facility MuLife of IRIG/DBSCI, funded by CEA Nanobio and GRAL LabEX (ANR-10-LABX-49-01) financed within the University Grenoble Alpes graduate school CBH-EUR-GS (ANR-17-EURE-0003). This project received funding from the LabEx Arcane, a program from the Chemistry Biology Health Graduate School of University Grenoble Alpes (ANR-17-EURE-0003).

## Author contributions

T. Rabilloud performed the sample preparation for shotgun analyses. Hélène Diemer and Fabrice Bertile performed and analysed shotgun experiments. M. Chevallet performed and analysed all the biological experiments. M. Chevallet, A. Deniaud and A. Fuchs conceived and coordinated the project. M. Chevallet wrote the paper, with contributions from all the authors.

## References

Abe, K. & Takeichi, M. 2008. EPLIN mediates linkage of the cadherin–catenin complex to F-actin and stabilizes the circumferential actin belt. Proceedings of the National Academy of Sciences, 105, 13–19.

Azioune, A., Carpi, N., Tseng, Q., Théry, M. & Piel, M. 2010. Chapter 8 - Protein Micropatterns: A Direct Printing Protocol Using Deep UVs. In: Cassimeris, L. & Tran, P. (eds.) Methods in Cell Biology. Academic Press.

Bartles, J., Wierda, A. & Zheng, L. 1996. Identification and characterization of espin, an actin-binding protein localized to the F-actin-rich junctional plaques of Sertoli cell ectoplasmic specializations. Journal of cell science, 109 (Pt 6), 1229–39.

Bays, J. L. & Demali, K. A. 2017. Vinculin in cell–cell and cell–matrix adhesions. Cellular and Molecular Life Sciences, 74, 2999–3009.

Bradford, M. M. 1976. A rapid and sensitive method for the quantitation of microgram quantities of protein utilizing the principle of protein-dye binding. Analytical Biochemistry, 72, 248–254.

Carapito, C., Burel, A., Guterl, P., Walter, A., Varrier, F., Bertile, F. & Van Dorsselaer, A. 2014. MSDA, a proteomics software suite for in-depth Mass Spectrometry Data Analysis using grid computing. PROTEOMICS, 14, 1014–1019.

Cavazza, C., Collin-Faure, V., Pérard, J., Diemer, H., Cianférani, S., Rabilloud, T. & Darrouzet, E. 2022. Proteomic analysis of Rhodospirillum rubrum after carbon monoxide exposure reveals an important effect on metallic cofactor biosynthesis. Journal of Proteomics, 250, 104389.

Chervin-Pétinot, A., Courçon, M., Almagro, S., Nicolas, A., Grichine, A., Grunwald, D., Prandini, M.-H., Huber, P. & Gulino-Debrac, D. 2012. Epithelial Protein Lost In Neoplasm (EPLIN) Interacts with α-Catenin and Actin Filaments in Endothelial Cells and Stabilizes Vascular Capillary Network in Vitro. Journal of Biological Chemistry, 287, 7556–7572.

Choi, S. S. & Diehl, A. M. 2009. Epithelial-to-mesenchymal transitions in the liver. Hepatology, 50, 2007–13.

Chung, G. H. C., Lorvellec, M., Gissen, P., Pichaud, F., Burden, J. J. & Stefan, C. J. 2022. The ultrastructural organization of endoplasmic reticulum-plasma membrane contacts is conserved in epithelial cells. Molecular Biology of the Cell, 33, ar113.

Clapéron, A., Mergey, M. & Fouassier, L. 2011. Roles of the scaffolding proteins NHERF in liver biology. Clinics and Research in Hepatology and Gastroenterology, 35, 176–181.

Coltman, N. J., Coke, B. A., Chatzi, K., Shepherd, E. L., Lalor, P. F., Schulz-Utermoehl, T. & Hodges, N. J. 2021. Application of HepG2/C3A liver spheroids as a model system for genotoxicity studies. Toxicol Lett, 345, 34–45.

Dascher, C. & Balch, W. E. 1996. Mammalian Sly1 Regulates Syntaxin 5 Function in Endoplasmic Reticulum to Golgi Transport*. Journal of Biological Chemistry, 271, 15866–15869.

Dunn, J. C. Y., Tompkins, R. G. & Yarmush, M. L. 1991. Long-Term in Vitro Function of Adult Hepatocytes in a Collagen Sandwich Configuration. Biotechnology Progress, 7, 237–245.

Friederich, E. & Louvard, D. 2006. Microvilli. Encyclopedic Reference of Genomics and Proteomics in Molecular Medicine. Berlin, Heidelberg: Springer Berlin Heidelberg.

Gissen, P. & Arias, I. M. 2015. Structural and functional hepatocyte polarity and liver disease. Journal of Hepatology, 63, 1023–1037.

Hozumi, S., Maeda, R., Taniguchi, K., Kanai, M., Shirakabe, S., Sasamura, T., Spéder, P., Noselli, S., Aigaki, T., Murakami, R. & Matsuno, K. 2006. An unconventional myosin in Drosophila reverses the default handedness in visceral organs. Nature, 440, 798–802.

Juan, T., Géminard, C., Coutelis, J.-B., Cerezo, D., PolÉs, S., Noselli, S. & Fürthauer, M. 2018. Myosin1D is an evolutionarily conserved regulator of animal left–right asymmetry. Nature Communications, 9, 1942.

Karpen, S. J. & Crawford, J. M. 1999. Cellular and molecular biology of the liver. Current Opinion in Gastroenterology, 15.

Karvar, S., Suda, J., Zhu, L. & Rockey, D. C. 2014. Distribution dynamics and functional importance of NHERF1 in regulation of Mrp-2 trafficking in hepatocytes. American Journal of Physiology-Cell Physiology, 307, C727–C737.

Kobielak, A. & Fuchs, E. 2004. α-catenin: at the junction of intercellular adhesion and actin dynamics. Nature Reviews Molecular Cell Biology, 5, 614–625.

Kocher, O., Comella, N., Gilchrist, A., Pal, R., Tognazzi, K., Brown, L. F. & Knoll, J. H. 1999. PDZK1, a novel PDZ domain-containing protein up-regulated in carcinomas and mapped to chromosome 1q21, interacts with cMOAT (MRP2), the multidrug resistance-associated protein. Lab Invest, 79, 1161–70.

Lebreton, G., Géminard, C., Lapraz, F., Pyrpassopoulos, S., Cerezo, D., Spéder, P., Ostap, E. M. & Noselli, S. 2018. Molecular to organismal chirality is induced by the conserved myosin 1D. Science, 362, 949–952.

Li, Q., Zhang, Y., Pluchon, P., Robens, J., Herr, K., Mercade, M., Thiery, J.-P., Yu, H. & Viasnoff, V. 2016. Extracellular matrix scaffolding guides lumen elongation by inducing anisotropic intercellular mechanical tension. Nature Cell Biology, 18, 311–318.

Liu, K. C. & Cheney, R. E. 2012. Myosins in cell junctions. Bioarchitecture, 2, 158–70.

Lyubimova, T., Caglio, S., Gelfi, C., Righetti, P. G. & Rabilloud, T. 1993. Photopolymerization of polyacrylamide gels with methylene blue. ELECTROPHORESIS, 14, 40–50.

Maul, R. S., Song, Y., Amann, K. J., Gerbin, S. C., Pollard, T. D. & Chang, D. D. 2003. EPLIN regulates actin dynamics by cross-linking and stabilizing filaments. J Cell Biol, 160, 399–407.

Müsch, A. 2018. From a common progenitor to distinct liver epithelial phenotypes. Curr Opin Cell Biol, 54, 18–23.

Ouderkirk, J. L. & Krendel, M. 2014. Myosin 1e is a component of the invadosome core that contributes to regulation of invadosome dynamics. Experimental Cell Research, 322, 265–276.

Perez-Riverol, Y., Bai, J., Bandla, C., García-Seisdedos, D., Hewapathirana, S., Kamatchinathan, S., Kundu Deepti J., Prakash, A., Frericks-Zipper, A., Eisenacher, M., Walzer, M., Wang, S., Brazma, A. & Vizcaíno Juan A. 2022. The PRIDE database resources in 2022: a hub for mass spectrometry-based proteomics evidences. Nucleic Acids Research, 50, D543–D552.

Petzoldt, A. G., Coutelis, J.-B., Géminard, C., Spéder, P., Suzanne, M., Cerezo, D. & Noselli, S. 2012. DE-Cadherin regulates unconventional Myosin ID and Myosin IC in Drosophila left-right asymmetry establishment. Development, 139, 1874–1884.

Phillips, M. J., Azuma, T., Meredith, S.-L. M., Squire, J. A., Ackerley, C. A., Pluthero, F. G., Roberts, E. A., Superina, R. A., Levy, G. A. & Marsden, P. A. 2003. Abnormalities in villin gene expression and canalicular microvillus structure in progressive cholestatic liver disease of childhood. The Lancet, 362, 1112–1119.

Riga, A., Castiglioni, V. G. & Boxem, M. 2020. New insights into apical-basal polarization in epithelia. Current Opinion in Cell Biology, 62, 1–8.

Schindelin, J., Arganda-Carreras, I., Frise, E., Kaynig, V., Longair, M., Pietzsch, T., Preibisch, S., Rueden, C., Saalfeld, S., Schmid, B., Tinevez, J.-Y., White, D. J., Hartenstein, V., Eliceiri, K., Tomancak, P. & Cardona, A. 2012. Fiji: an open-source platform for biological-image analysis. Nature Methods, 9, 676–682.

Schulze, R. J., Schott, M. B., Casey, C. A., Tuma, P. L. & Mcniven, M. A. 2019. The cell biology of the hepatocyte: A membrane trafficking machine. Journal of Cell Biology, 218, 2096–2112.

Sharma, V. R., Shrivastava, A., Gallet, B., Karepina, E., Charbonnier, P., Chevallet, M., Jouneau, P.-H. & Deniaud, A. 2020. Canalicular domain structure and function in matrix-free hepatic spheroids. Biomaterials Science, 8, 485–496.

Shore, E. & Nelson, W. 1991. Biosynthesis of the cell adhesion molecule uvomorulin (E-cadherin) in Madin-Darby canine kidney epithelial cells. The Journal of biological chemistry, 266, 19672–80.

Spéder, P., ádám, G. & Noselli, S. 2006. Type ID unconventional myosin controls left–right asymmetry in Drosophila. Nature, 440, 803–807.

Štampar, M., Sedighi Frandsen, H., Rogowska-Wrzesinska, A., Wrzesinski, K., Filipič, M. & Žegura, B. 2021. Hepatocellular carcinoma (HepG2/C3A) cell-based 3D model for genotoxicity testing of chemicals. Sci Total Environ, 755, 143255.

Taguchi, K., Ishiuchi, T. & Takeichi, M. 2011. Mechanosensitive EPLIN-dependent remodeling of adherens junctions regulates epithelial reshaping. Journal of Cell Biology, 194, 643–656.

Treyer, A. & Müsch, A. 2013. Hepatocyte Polarity. Comprehensive Physiology.

Tsuchiya, B., Sato, Y., Kameya, T., Okayasu, I. & Mukai, K. 2006. Differential expression of N-cadherin and E-cadherin in normal human tissues. Archives of histology and cytology, 69, 135–145.

Tsukada, N., Ackerley, C. A. & Phillips, M. J. 1995. The structure and organization of the bile canalicular cytoskeleton with special reference to actin and actin-binding proteins. Hepatology, 21, 1106–1113.

Turner, C. E., Glenney, J. R., JR & Burridge, K. 1990. Paxillin: a new vinculin-binding protein present in focal adhesions. Journal of Cell Biology, 111, 1059–1068.

Van Weering, J. R. T., Verkade, P. & Cullen, P. J. 2012. SNX–BAR-Mediated Endosome Tubulation is Co-ordinated with Endosome Maturation. Traffic, 13, 94–107.

Zhang, Y.-Y., Fu, Z.-Y., Wei, J., Qi, W., Baituola, G., Luo, J., Meng, Y.-J., Guo, S.-Y., Yin, H., Jiang, S.-Y., Li, Y.-F., Miao, H.-H., Liu, Y., Wang, Y., Li, B.-L. MA Y.-T. & Song, B.-L. 2018. A LIMA1 variant promotes low plasma LDL cholesterol and decreases intestinal cholesterol absorption. Science, 360, 1087–1092.

Zhang, Y., De Mets, R., Monzel, C., Acharya, V., Toh, P., Chin, J. F. L., Van Hul, N., Ng, I. C., Yu, H., Ng, S. S., Tamir Rashid, S. & Viasnoff, V. 2020. Biomimetic niches reveal the minimal cues to trigger apical lumen formation in single hepatocytes. Nature Materials, 19, 1026–1035.

Zheng, L., Sekerková, G., Vranich, K., Tilney, L. G., Mugnaini, E. & Bartles, J. R. 2000. The Deaf Jerker Mouse Has a Mutation in the Gene Encoding the Espin Actin-Bundling Proteins of Hair Cell Stereocilia and Lacks Espins. Cell, 102, 377–385.

