## Supplementary material for "E cadherin appears to be an essential on/off switch for initiating bile canaliculi formation": Table S1

| accession | N° accession | gene_name | description | HepG2/A | HepG2/B | HepG2/C | HepG2/D | C3A/1 | C3A/1 | C3A/3 | C3A/4 | C3A/hepG2 | U |
|---|---|---|---|---|---|---|---|---|---|---|---|---|---|
| sp\|B1AK5 | B1AK53 | ESPN | Espin OS=Homo sapiens (Human) GN=ESPN PE=1 SV=1 | 0 | 0 | 0 | 0 | 1908465 | 2764452.25 | 1987987.62 | 1496092.12 | #DIV/0! | 0 |
| sp\|O4377 | O43772 | SLC25A20 | Mitochondrial carnitine/acylcarnitine carrier protein OS= | 0 | 0 | 0 | 0 | 811015.62 | 0 | 0 | 779974 | #DIV/0! | 4 |
| sp\|O9547 | O95479 | H6PD | GDH/6PGL endoplasmic bifunctional protein OS=Homo s | 0 | 0 | 0 | 0 | 2958574 | 2355119.5 | 2368595.25 | 2476773.5 | #DIV/0! | 0 |
| sp\|P04003 | P04003 | C4BPA | C4b-binding protein alpha chain OS=Homo sapiens (Hum | 0 | 0 | 0 | 0 | 696544.62 | 673287.81 | 1043348.44 | 802446.75 | #DIV/0! | 0 |
| sp\|P12830 | P12830 | CDH1 | Cadherin-1 OS=Homo sapiens (Human) GN=CDH1 PE=1 S | 0 | 0 | 0 | 0 | 1154234.12 | 1294069.25 | 0 | 1189502.38 | #DIV/0! | 2 |
| sp\|P19022 | P19022 | CDH2 | Cadherin-2 OS=Homo sapiens (Human) GN=CDH2 PE=1 S | 0 | 0 | 0 | 0 | 732604.75 | 0 | 0 | 455703.41 | #DIV/0! | 4 |
| sp\|P26447 | P26447 | S100A4 | Protein S100-A4 OS=Homo sapiens (Human) GN=S100A4 | 0 | 0 | 0 | 0 | 28140340 | 24040756 | 27760456 | 20764890 | #DIV/0! | 0 |
| sp\|P28300 | P28300 | LOX | Protein-lysine 6-oxidase OS=Homo sapiens (Human) GN= | 0 | 0 | 0 | 0 | 2011000.75 | 4246583.5 | 2071924.88 | 1610688.5 | #DIV/0! | 0 |
| sp\|P45877 | P45877 | PPIC | Peptidyl-prolyl cis-trans isomerase C OS=Homo sapiens ( | 0 | 0 | 0 | 0 | 0 | 2260498 | 0 | 0 | #DIV/0! | 6 |
| sp\|P54619 | P54619 | PRKAG1 | 5'-AMP-activated protein kinase subunit gamma-1 OS=H | 0 | 0 | 0 | 0 | 0 | 450412.06 | 0 | 643203.88 | #DIV/0! | 4 |
| sp\|Q0141 | Q01415 | GALK2 | N-acetylgalactosamine kinase OS=Homo sapiens (Humar | 0 | 0 | 0 | 0 | 1033687.19 | 986572.62 | 1355547.12 | 1021825.5 | #DIV/0! | 0 |
| sp\|Q0233 | Q02338 | BDH1 | D-beta-hydroxybutyrate dehydrogenase, mitochondrial | 0 | 0 | 0 | 0 | 887347.75 | 0 | 0 | 961209.94 | #DIV/0! | 4 |
| sp\|Q0494 | Q04941 | PLP2 | Proteolipid protein 2 OS=Homo sapiens (Human) GN=PLI | 0 | 0 | 0 | 0 | 646311.94 | 830417.94 | 0 | 924163.56 | #DIV/0! | 2 |
| sp\|Q1349 | Q13496 | MTM1 | Myotubularin OS=Homo sapiens (Human) GN=MTM1 PE | 0 | 0 | 0 | 0 | 193897.95 | 237671.36 | 0 | 0 | #DIV/0! | 4 |
| sp\|Q1679 | Q16790 | CA9 | Carbonic anhydrase 9 OS=Homo sapiens (Human) GN=C/ | 0 | 0 | 0 | 0 | 0 | 440721.34 | 0 | 0 | #DIV/0! | 6 |
| sp\|Q68CQ | Q68CQ4 | DIEXF | Digestive organ expansion factor homolog OS=Homo sap | 0 | 0 | 0 | 0 | 0 | 0 | 0 | 538072.62 | #DIV/0! | 6 |
| sp\|Q6ZWJ | Q6ZWJ1 | STXBP4 | Syntaxin-binding protein 4 OS=Homo sapiens (Human) G | 0 | 0 | 0 | 0 | 808332.38 | 758525.31 | 1417983.38 | 1494972 | #DIV/0! | 0 |
| sp\|Q8N8A | Q8N8A6 | DDX51 | ATP-dependent RNA helicase DDX51 OS=Homo sapiens ( | 0 | 0 | 0 | 0 | 0 | 0 | 0 | 393188.25 | #DIV/0! | 6 |
| sp\|Q8WV7 | Q8WV74 | NUDT8 | Nucleoside diphosphate-linked moiety X motif 8 OS=Hor | 0 | 0 | 0 | 0 | 0 | 1215944.5 | 0 | 959746.19 | #DIV/0! | 4 |
| sp\|Q8WXE | Q8WXE9 | STON2 | Stonin-2 OS=Homo sapiens (Human) GN=STON2 PE=1 SV | 0 | 0 | 0 | 0 | 1443645 | 1950862.75 | 1331275.75 | 1289709.25 | #DIV/0! | 0 |
| sp\|Q96MI | Q96MM6 | HSPA12B | Heat shock 70 kDa protein 12B OS=Homo sapiens (Huma | 0 | 0 | 0 | 0 | 1725221.75 | 1426756 | 1533494.5 | 1111711.5 | #DIV/0! | 0 |
| sp\|Q9BRP | Q9BRP8 | PYM1 | Partner of Y14 and mago OS=Homo sapiens (Human) GN | 0 | 0 | 0 | 0 | 2692647 | 0 | 1602866.25 | 793578.81 | #DIV/0! | 2 |
| sp\|Q9H0T | Q9H0T7 | RAB17 | Ras-related protein Rab-17 OS=Homo sapiens (Human) ( | 0 | 0 | 0 | 0 | 361042.28 | 621488.38 | 0 | 428622.44 | #DIV/0! | 2 |
| sp\|Q9H2H | Q9H2H8 | PPIL3 | Peptidyl-prolyl cis-trans isomerase-like 3 OS=Homo sapie | 0 | 0 | 0 | 0 | 3604556.25 | 3456922.75 | 3084631.5 | 0 | #DIV/0! | 2 |
| sp\|Q9H9H | Q9H9H4 | VPS37B | Vacuolar protein sorting-associated protein 37B OS=Hon | 0 | 0 | 0 | 0 | 110334.77 | 0 | 220197.45 | 169826.81 | #DIV/0! | 2 |
| sp\|Q9NPE | Q9NPE3 | NOP10 | H/ACA ribonucleoprotein complex subunit 3 OS=Homo s | 0 | 0 | 0 | 0 | 0 | 560557.75 | 856217.5 | 901651.12 | #DIV/0! | 2 |
| sp\|Q9UBV | Q9UBV7 | B4GALT7 | Beta-1,4-galactosyltransferase 7 OS=Homo sapiens (Hun | 0 | 0 | 0 | 0 | 558074.44 | 488286 | 0 | 603932.88 | #DIV/0! | 2 |
| sp\|Q9UHK | Q9UHK6 | AMACR | Alpha-methylacyl-CoA racemase OS=Homo sapiens (Hun | 0 | 0 | 0 | 0 | 468573.5 | 0 | 0 | 467077.19 | #DIV/0! | 4 |
| sp\|Q9UN7 | Q9UN76 | SLC6A14 | Sodium- and chloride-dependent neutral and basic amin | 0 | 0 | 0 | 0 | 3123292.5 | 2696305.5 | 2309328 | 2608354 | #DIV/0! | 0 |
| sp\|Q9Y2Z | Q9Y2Z9 | COQ6 | Ubiquinone biosynthesis monooxygenase COQ6, mitoch | 0 | 0 | 0 | 0 | 796207.88 | 742697.5 | 808382.38 | 770921.38 | #DIV/0! | 0 |
| sp\|Q9308 | Q93088 | BHMT | Betaine--homocysteine S-methyltransferase 1 OS=Homo | 0 | 0 | 0 | 115166.28 | 124380.49 | 612988 | 665873.94 | 520268.25 | 16.70203014 | 0 |
| sp\|Q9HAN | Q9HAN9 | NMNAT1 | Nicotinamide/nicotinic acid mononucleotide adenylyltra | 144437.73 | 121285.45 | 78449.05 | 79684.85 | 1646645.88 | 1218367.75 | 1509816 | 1383157.88 | 13.5847383 | 0 |
| sp\|Q6UX5 | Q6UX53 | METTL7B | Methyltransferase-like protein 7B OS=Homo sapiens (Hu | 0 | 739170.88 | 0 | 0 | 2682103.5 | 2790991 | 2011023.25 | 1974576.5 | 12.79635671 | 0 |
| sp\|Q9BVN | Q9BVM2 | DPCD | Protein DPCD OS=Homo sapiens (Human) GN=DPCD PE= | 191620.45 | 0 | 208364.58 | 0 | 1271966 | 1429098.5 | 1119015.62 | 339489.81 | 10.39931402 | 0 |
| sp\|P16401 | P16401 | HIST1H1B | Histone H1.5 OS=Homo sapiens (Human) GN=HIST1H1B | 3792671.5 | 4297052 | 2838022.5 | 4971669.5 | 39437316 | 47144536 | 37063704 | 34647848 | 9.95592599 | 0 |
| sp\|Q5HYK | Q5HYK3 | COQ5 | 2-methoxy-6-polyprenyl-1,4-benzoquinol methylase, mit | 0 | 0 | 0 | 345793.38 | 983145.69 | 1202599.25 | 750651.69 | 245874.34 | 9.202810563 | 1 |
| sp\|Q9P1F | Q9P1F3 | ABRACL | Costars family protein ABRACL OS=Homo sapiens (Huma | 0 | 0 | 229621.61 | 0 | 469402.97 | 637709.19 | 435356.84 | 543230.44 | 9.083201882 | 0 |
| sp\|Q1529 | Q15293 | RCN1 | Reticulocalbin-1 OS=Homo sapiens (Human) GN=RCN1 P | 527072.75 | 1969118.25 | 430072.94 | 1719015.5 | 9563640 | 8707646 | 11999824 | 9723310 | 8.609690874 | 0 |
| sp\|P16455 | P16455 | MGMT | Methylated-DNA--protein-cysteine methyltransferase O: | 445377.59 | 0 | 0 | 0 | 670625 | 0 | 889830.06 | 1914744.38 | 7.802816123 | 2.5 |
| sp\|O9552 | O95528 | SLC2A10 | Solute carrier family 2, facilitated glucose transporter me | 0 | 0 | 208099.2 | 0 | 327121.06 | 446482.41 | 353138.09 | 332418.91 | 7.011850454 | 0 |
| sp\|Q8N1S | Q8N1S5 | SLC39A11 | Zinc transporter ZIP11 OS=Homo sapiens (Human) GN=S | 0 | 0 | 0 | 570417.12 | 933130 | 1046985.69 | 1072582.38 | 881119.12 | 6.896386963 | 0 |
| sp\|Q9H9P | Q9H9P8 | L2HGDH | L-2-hydroxyglutarate dehydrogenase, mitochondrial OS= | 0 | 333506.38 | 0 | 0 | 403771.75 | 645875.62 | 555215.81 | 641242.94 | 6.734822044 | 0 |
| sp\|Q1552 | Q15526 | SURF1 | Surfeit locus protein 1 OS=Homo sapiens (Human) GN=S | 329488.38 | 0 | 0 | 0 | 456440.5 | 652092.31 | 441309.09 | 664802.44 | 6.721464168 | 0 |
| sp\|Q86VU | Q86VU5 | COMTD1 | Catechol O-methyltransferase domain-containing proteir | 0 | 144233.17 | 0 | 214601.89 | 678793.12 | 443652.31 | 111064.4 | 1140447.38 | 6.61573373 | 2 |

| | | | | | | | | | | | | | |
|---|---|---|---|---|---|---|---|---|---|---|---|---|---|
| sp\|Q9H47 | Q9H477 | RBKS | Ribokinase OS=Homo sapiens (Human) GN=RBKS PE=1 S' | 0 | 0 | 0 | 469100.66 | 624982.25 | 710477.81 | 826371.5 | 846948.56 | **6.413932822** | 0 |
| sp\|A8MXV | A8MXV4 | NUDT19 | Nucleoside diphosphate-linked moiety X motif 19 OS=Hc | 283603.84 | 301434.03 | 357048.34 | 485903.25 | 2197929 | 2036101.5 | 2451645 | 2160388.25 | **6.194768237** | 0 |
| sp\|L0R6Q1 | L0R6Q1 | SLC35A4 | SLC35A4 upstream open reading frame protein OS=Hom | 642917.12 | 0 | 0 | 0 | 1257118.75 | 1175074.75 | 347347.84 | 1194141.62 | **6.180707958** | 1 |
| sp\|Q9GZP | Q9GZP4 | PITHD1 | PITH domain-containing protein 1 OS=Homo sapiens (Hu | 0 | 312092.53 | 292481.66 | 454250.28 | 925701.06 | 1889953 | 605344.44 | 3029564.5 | **6.092192977** | 0 |
| sp\|Q0199 | Q01995 | TAGLN | Transgelin OS=Homo sapiens (Human) GN=TAGLN PE=1 | 4714150.5 | 4196114 | 5637111 | 4653550 | 28969708 | 27786190 | 31434258 | 28263676 | **6.065011397** | 0 |
| sp\|Q9H3K | Q9H3K2 | GHITM | Growth hormone-inducible transmembrane protein OS= | 0 | 0 | 1062266.12 | 0 | 2312774.5 | 1913306.12 | 0 | 2159903.75 | **6.011661532** | 2.5 |
| sp\|O43301 | O43301 | HSPA12A | Heat shock 70 kDa protein 12A OS=Homo sapiens (Huma | 2526798.25 | 1248239.12 | 2571760.5 | 2442376.75 | 14121052 | 13437114 | 11871804 | 9352308 | **5.550268382** | 0 |
| sp\|Q66K1 | Q66K14 | TBC1D9B | TBC1 domain family member 9B OS=Homo sapiens (Hun | 0 | 0 | 351722.66 | 0 | 421456.81 | 541658 | 432013.91 | 465166.25 | **5.289096159** | 0 |
| sp\|O9598 | O95989 | NUDT3 | Diphosphoinositol polyphosphate phosphohydrolase 1 C | 596409.25 | 0 | 0 | 0 | 931816.62 | 586955.75 | 705561.62 | 926390.94 | **5.282823715** | 1 |
| sp\|Q1406 | Q14061 | COX17 | Cytochrome c oxidase copper chaperone OS=Homo sapi | 0 | 0 | 0 | 470840.91 | 948937.81 | 0 | 783240.75 | 728782.19 | **5.226735183** | 2.5 |
| sp\|O9534 | O95342 | ABCB11 | Bile salt export pump OS=Homo sapiens (Human) GN=AE | 372662.44 | 0 | 0 | 0 | 464797.75 | 485576.44 | 416180.12 | 550620.31 | **5.144534072** | 0 |
| sp\|Q96DX | Q96DX5 | ASB9 | Ankyrin repeat and SOCS box protein 9 OS=Homo sapien | 0 | 290057.94 | 264187.97 | 0 | 563853.62 | 719230.5 | 776719.06 | 744026.94 | **5.058819685** | 0 |
| sp\|Q1401 | Q14019 | COTL1 | Coactosin-like protein OS=Homo sapiens (Human) GN=C | 134584.03 | 0 | 88718.66 | 0 | 308770.25 | 243030.83 | 335808.88 | 235446.28 | **5.029300095** | 0 |
| sp\|Q0225 | Q02252 | ALDH6A1 | Methylmalonate-semialdehyde dehydrogenase [acylatin | 3127067.5 | 2925258.25 | 3073416.5 | 3458688.5 | 14790995 | 15730565 | 16164997 | 15578190 | **4.947760311** | 0 |
| sp\|Q9C07 | Q9C075 | KRT23 | Keratin, type I cytoskeletal 23 OS=Homo sapiens (Humar | 2043801.62 | 5589553.5 | 3274368.5 | 2625910.75 | 15681121 | 17663118 | 17083504 | 15109452 | **4.842542159** | 0 |
| sp\|Q96FX | Q96FX7 | TRMT61A | tRNA (adenine(58)-N(1))-methyltransferase catalytic subu | 0 | 577585.94 | 0 | 0 | 548123.81 | 1308266.75 | 517972.75 | 377146.22 | **4.763809746** | 3 |
| sp\|Q8WZA | Q8WZA0 | LZIC | Protein LZIC OS=Homo sapiens (Human) GN=LZIC PE=1 S | 0 | 0 | 120084.95 | 394811.34 | 390785.34 | 716138.62 | 700732.06 | 643718.94 | **4.760910124** | 1 |
| sp\|Q9BW | Q9BWM7 | SFXN3 | Sideroflexin-3 OS=Homo sapiens (Human) GN=SFXN3 PE | 2369179.5 | 2530861.5 | 2650147 | 3524567 | 13488156 | 11792444 | 12832942 | 13019106 | **4.617045524** | 0 |
| sp\|O0023 | O00233 | PSMD9 | 26S proteasome non-ATPase regulatory subunit 9 OS=Hc | 1267615.25 | 820815.62 | 871465.88 | 0 | 3113662.5 | 3121672.5 | 4006249.5 | 3366934 | **4.59763284** | 0 |
| sp\|E9PQ5 | E9PQ53 | NDUFC2-K | NADH dehydrogenase [ubiquinone] 1 subunit C2, isoforn | 0 | 0 | 0 | 1770736 | 1641023.12 | 2195519 | 2320512.75 | 1901151 | **4.550766388** | 1 |
| sp\|Q1664 | Q16643 | DBN1 | Drebrin OS=Homo sapiens (Human) GN=DBN1 PE=1 SV= | 2688307.75 | 2875008.25 | 2873674.25 | 3440561 | 14426514 | 14729209 | 13108507 | 11373550 | **4.51589548** | 0 |
| sp\|Q8IV08 | Q8IV08 | PLD3 | Phospholipase D3 OS=Homo sapiens (Human) GN=PLD3 | 542847.38 | 0 | 173965.55 | 641843.06 | 1467975.5 | 1346115.5 | 1827292.88 | 1450401.75 | **4.483685109** | 0 |
| sp\|Q9UHB | Q9UHB6 | LIMA1 | LIM domain and actin-binding protein 1 OS=Homo sapie | 1602720.5 | 568082 | 527400.56 | 1901141 | 4143505.5 | 4545209 | 5880014.5 | 6012259.5 | **4.474766017** | 0 |
| sp\|Q9267 | Q92673 | SORL1 | Sortilin-related receptor OS=Homo sapiens (Human) GN | 724646.88 | 647402.06 | 637206.5 | 570311.62 | 3233266 | 2865809 | 2020272.75 | 3210330 | **4.392084984** | 0 |
| sp\|P56199 | P56199 | ITGA1 | Integrin alpha-1 OS=Homo sapiens (Human) GN=ITGA1 F | 976470 | 1555815.62 | 0 | 1204263.38 | 4202960 | 4021012.5 | 3727756 | 4095155 | **4.294573281** | 0 |
| sp\|Q8IV48 | Q8IV48 | ERI1 | 3'-5' exoribonuclease 1 OS=Homo sapiens (Human) GN= | 0 | 0 | 526040.31 | 0 | 0 | 621962.12 | 581622.44 | 1036857.88 | **4.25906988** | 2.5 |
| sp\|Q6P4A | Q6P4A7 | SFXN4 | Sideroflexin-4 OS=Homo sapiens (Human) GN=SFXN4 PE | 0 | 0 | 65329.05 | 0 | 41718.48 | 0 | 84387.32 | 147965.16 | **4.195238718** | 3.5 |
| sp\|Q9BV7 | Q9BV79 | MECR | Enoyl-[acyl-carrier-protein] reductase, mitochondrial OS | 0 | 120364.16 | 179601.3 | 162390.14 | 279461.91 | 491940 | 563326 | 600654.81 | **4.185918198** | 0 |
| sp\|Q6JQN | Q6JQN1 | ACAD10 | Acyl-CoA dehydrogenase family member 10 OS=Homo s | 331219.31 | 253089.09 | 472605.31 | 414293.72 | 1479769.5 | 1605174.5 | 1592915 | 1458509.88 | **4.17097464** | 0 |
| sp\|Q96EY | Q96EY8 | MMAB | Corrinoid adenosyltransferase OS=Homo sapiens (Huma | 1238137.38 | 1135830.12 | 1005778.88 | 1135537.12 | 4759008.5 | 3824713 | 5169311 | 5000664 | **4.153381842** | 0 |
| sp\|Q53EL | Q53EL6 | PDCD4 | Programmed cell death protein 4 OS=Homo sapiens (Hu | 2080747 | 2593173.75 | 2090019.25 | 1963733 | 8711892 | 7945302.5 | 9972239 | 9475916 | **4.136881561** | 0 |
| sp\|Q9Y6X | Q9Y6X4 | FAM169A | Soluble lamin-associated protein of 75 kDa OS=Homo sa | 2743394.5 | 2632627 | 1725017.25 | 2555307.25 | 10514769 | 8100400 | 10131081 | 11063804 | **4.122683052** | 0 |
| sp\|P36405 | P36405 | ARL3 | ADP-ribosylation factor-like protein 3 OS=Homo sapiens | 3708625.75 | 2217666.5 | 1926569.75 | 3847348.75 | 13848389 | 12496136 | 11498094 | 10316835 | **4.116118507** | 0 |
| sp\|P21589 | P21589 | NT5E | 5'-nucleotidase OS=Homo sapiens (Human) GN=NT5E PE | 2261539.5 | 2559232.75 | 1040403.25 | 2711339.25 | 8906690 | 8144391 | 8844703 | 9037868 | **4.075076336** | 0 |
| sp\|Q9NQ | Q9NQH7 | XPNPEP3 | Xaa-Pro aminopeptidase 3 OS=Homo sapiens (Human) G | 355868.62 | 374062.88 | 867430.38 | 735552 | 2038303 | 2178758.25 | 2955021.5 | 2300167.5 | **4.060265718** | 0 |
| sp\|Q8N5I4 | Q8N5I4 | DHRSX | Dehydrogenase/reductase SDR family member on chrom | 0 | 64599.52 | 0 | 74022.55 | 114369.81 | 135515.39 | 134466.41 | 177257.44 | **4.051368227** | 0 |
| sp\|O9483 | O94832 | MYO1D | Unconventional myosin-Id OS=Homo sapiens (Human) G | 1147121 | 554012.25 | 861375.06 | 1558830.62 | 4112810 | 3074553.75 | 3842515.5 | 5650455 | **4.047309511** | 0 |
| sp\|O7576 | O75764 | TCEA3 | Transcription elongation factor A protein 3 OS=Homo sa | 3546846.5 | 1086491.25 | 1976512.88 | 1839161.25 | 9875713 | 8292721 | 8349576 | 7292088 | **4.001662973** | 0 |
| sp\|O9537 | O95372 | LYPLA2 | Acyl-protein thioesterase 2 OS=Homo sapiens (Human) ( | 4435650.5 | 3509321 | 2352547.75 | 2761186.75 | 15026369 | 13481737 | 13647236 | 9963379 | **3.99110915** | 0 |
| sp\|Q8N4F | Q8N4F0 | BPIFB2 | BPI fold-containing family B member 2 OS=Homo sapien | 118239.92 | 0 | 79408.53 | 0 | 170233.55 | 208073.08 | 182465.95 | 221835.58 | **3.959596749** | 0 |
| sp\|Q9282 | Q92823 | NRCAM | Neuronal cell adhesion molecule OS=Homo sapiens (Hur | 541285.88 | 0 | 0 | 339507.66 | 1289873.88 | 706879.56 | 700529.44 | 738389.75 | **3.900656027** | 0 |
| sp\|Q1437 | Q14376 | GALE | UDP-glucose 4-epimerase OS=Homo sapiens (Human) GI | 5612641 | 6637734 | 4331794.5 | 6042219 | 22809012 | 22328400 | 20789608 | 20945248 | **3.839762034** | 0 |
| sp\|O7517 | O75175 | CNOT3 | CCR4-NOT transcription complex subunit 3 OS=Homo sa | 0 | 0 | 625281.56 | 0 | 791190.69 | 0 | 869019.69 | 588459.75 | **3.596252111** | 3.5 |
| sp\|Q9H8H | Q9H8H3 | METTL7A | Methyltransferase-like protein 7A OS=Homo sapiens (Hu | 8256284.5 | 7700495 | 8140532 | 6921724.5 | 27852964 | 24452920 | 29158120 | 29390568 | **3.573759417** | 0 |
| sp\|P04066 | P04066 | FUCA1 | Tissue alpha-L-fucosidase OS=Homo sapiens (Human) GI | 0 | 0 | 0 | 155660.39 | 123496.62 | 158815.34 | 105436.33 | 155448.22 | **3.489625781** | 3 |

| | | | | | | | | | | | | | |
|---|---|---|---|---|---|---|---|---|---|---|---|---|---|
| sp\|Q9BX6 | Q9BX68 | HINT2 | Histidine triad nucleotide-binding protein 2, mitochondri | 0 | 1719383.88 | 1684246 | 0 | 3130698.5 | 2538411.25 | 3374048.75 | 2765617.25 | **3.469465296** | **0** |
| sp\|P26232 | P26232 | CTNNA2 | Catenin alpha-2 OS=Homo sapiens (Human) GN=CTNNA | 844492.19 | 509548.44 | 2530000.5 | 691574 | 4323055.5 | 2247579.5 | 4798847.5 | 4439583 | **3.455068893** | **1** |
| sp\|P05166 | P05166 | PCCB | Propionyl-CoA carboxylase beta chain, mitochondrial OS | 3648119.75 | 3662768.5 | 1982919.5 | 1715652 | 7943956 | 6711949 | 11202236 | 11791723 | **3.419773981** | **0** |
| sp\|O0027 | O00273 | DFFA | DNA fragmentation factor subunit alpha OS=Homo sapie | 1050523.75 | 891167 | 997120.62 | 2554311 | 5435456.5 | 3724118 | 4916605 | 4505210.5 | **3.382664494** | **0** |
| sp\|P52926 | P52926 | HMGA2 | High mobility group protein HMGI-C OS=Homo sapiens ( | 6272304.5 | 5305896 | 2953235.75 | 4927511.5 | 11442451 | 17968972 | 20893432 | 14552256 | **3.333022517** | **0** |
| sp\|Q1424 | Q14241 | ELOA | Elongin-A OS=Homo sapiens (Human) GN=ELOA PE=1 SV | 247088.84 | 263281.03 | 325744.16 | 316234.41 | 795517.88 | 1073884.25 | 1037241.5 | 913977.12 | **3.315508242** | **0** |
| sp\|Q8IYU | Q8IYU8 | MICU2 | Calcium uptake protein 2, mitochondrial OS=Homo sapie | 528643.38 | 0 | 0 | 0 | 585987 | 660964.62 | 503617.62 | 0 | **3.311436984** | **3.5** |
| sp\|P35914 | P35914 | HMGCL | Hydroxymethylglutaryl-CoA lyase, mitochondrial OS=Ho | 2165033 | 2784923.75 | 5590937 | 4650248 | 13257032 | 9189960 | 14523186 | 12371952 | **3.248085681** | **0** |
| sp\|Q9NQT | Q9NQT8 | KIF13B | Kinesin-like protein KIF13B OS=Homo sapiens (Human) G | 0 | 107328.1 | 298947.78 | 0 | 327418.41 | 310939.19 | 319133.25 | 347716.56 | **3.212613582** | **0** |
| sp\|Q9NVZ | Q9NVZ3 | NECAP2 | Adaptin ear-binding coat-associated protein 2 OS=Homo | 1567181.25 | 682139.5 | 1650218.88 | 1750496.12 | 4152752.75 | 4175162.5 | 4463163 | 5308033 | **3.203362253** | **0** |
| sp\|P30038 | P30038 | ALDH4A1 | Delta-1-pyrroline-5-carboxylate dehydrogenase, mitochc | 42554156 | 37448900 | 36165828 | 38472080 | 136424432 | 124146752 | 118922528 | 109743232 | **3.163695643** | **0** |
| sp\|P17174 | P17174 | GOT1 | Aspartate aminotransferase, cytoplasmic OS=Homo sapi | 57406048 | 56415140 | 50395928 | 57221572 | 176066208 | 172954672 | 187604688 | 159597264 | **3.144088498** | **0** |
| sp\|Q8IZV5 | Q8IZV5 | RDH10 | Retinol dehydrogenase 10 OS=Homo sapiens (Human) G | 390454.06 | 0 | 371911.75 | 326470.5 | 708663.38 | 896861.31 | 740539.31 | 1036345.88 | **3.106444788** | **0** |
| sp\|Q9261 | Q92615 | LARP4B | La-related protein 4B OS=Homo sapiens (Human) GN=LA | 1219397.75 | 0 | 758759.94 | 849551.94 | 2913744.5 | 2153756.25 | 1918664.5 | 1783650.75 | **3.101384918** | **0** |
| sp\|P23786 | P23786 | CPT2 | Carnitine O-palmitoyltransferase 2, mitochondrial OS=H | 1260726.88 | 1137535.5 | 1382466.75 | 1349556.12 | 2561967 | 3109424 | 5066390 | 5063093 | **3.079921141** | **0** |
| sp\|Q7Z73 | Q7Z739 | YTHDF3 | YTH domain-containing family protein 3 OS=Homo sapie | 0 | 0 | 0 | 601421.94 | 1018073.88 | 0 | 0 | 829598.94 | **3.072173955** | **5** |
| sp\|O6021 | O60218 | AKR1B10 | Aldo-keto reductase family 1 member B10 OS=Homo sap | 28307538 | 29797264 | 29962506 | 30846442 | 95682960 | 83952440 | 99298952 | 85395040 | **3.06381215** | **0** |
| sp\|Q96AQ | Q96AQ6 | PBXIP1 | Pre-B-cell leukemia transcription factor-interacting prote | 1125008.38 | 1473037.88 | 1112393.12 | 756331.88 | 3735029.25 | 3677161.5 | 3182046.25 | 2991696.75 | **3.041555737** | **0** |
| sp\|O4314 | O43148 | RNMT | mRNA cap guanine-N7 methyltransferase OS=Homo sap | 806177.5 | 0 | 0 | 895390.19 | 1364936.12 | 1487228.25 | 469818 | 1801502.5 | **3.011037939** | **2** |
| sp\|P16083 | P16083 | NQO2 | Ribosyldihydronicotinamide dehydrogenase [quinone] C | 0 | 1207282.5 | 1226028.12 | 448014.12 | 2019272.88 | 2263434 | 2044592.62 | 2340735.5 | **3.008350596** | **0** |
| sp\|Q9Y4K | Q9Y4K1 | CRYBG1 | Beta/gamma crystallin domain-containing protein 1 OS= | 1222940 | 965831.38 | 1369441.62 | 1176348.75 | 3310157.25 | 3705553 | 3938035.5 | 3288687.75 | **3.008184126** | **0** |
| sp\|Q1476 | Q14764 | MVP | Major vault protein OS=Homo sapiens (Human) GN=MVI | 2381824.75 | 1920563.12 | 1092737 | 1891471 | 5699163.5 | 6242869 | 4973753 | 4886010.5 | **2.99204133** | **0** |
| sp\|P22612 | P22612 | PRKACG | cAMP-dependent protein kinase catalytic subunit gamma | 404349.84 | 0 | 0 | 1288892.25 | 1268735.75 | 2590389 | 0 | 1206840.25 | **2.991872828** | **5** |
| sp\|P00966 | P00966 | ASS1 | Argininosuccinate synthase OS=Homo sapiens (Human) | 6784523 | 11669274 | 11007104 | 9289934 | 29425788 | 22811244 | 29618568 | 33338302 | **2.972681802** | **0** |
| sp\|Q86WA | Q86WA6 | BPHL | Valacyclovir hydrolase OS=Homo sapiens (Human) GN=B | 388953.56 | 380621.22 | 300393.28 | 455926.41 | 628121.62 | 774253.38 | 2464115 | 653218.19 | **2.962005747** | **0** |
| sp\|Q96AB | Q96AB3 | ISOC2 | Isochorismatase domain-containing protein 2 OS=Homo | 597826.38 | 2389363 | 779286.5 | 1203238.12 | 4139824.5 | 2685389 | 4716559 | 3122061 | **2.950639312** | **0** |
| sp\|P07225 | P07225 | PROS1 | Vitamin K-dependent protein S OS=Homo sapiens (Huma | 784772.25 | 0 | 0 | 724879.88 | 1256063.25 | 1330890.88 | 939995.88 | 876756.31 | **2.91703382** | **0** |
| sp\|P34913 | P34913 | EPHX2 | Bifunctional epoxide hydrolase 2 OS=Homo sapiens (Hur | 770270.88 | 818596.56 | 790380.12 | 741731.81 | 2131867 | 2120632 | 2241639.25 | 2572983 | **2.905216656** | **0** |
| sp\|P51690 | P51690 | ARSE | Arylsulfatase E OS=Homo sapiens (Human) GN=ARSE PE= | 705568.88 | 1465778.75 | 475930.84 | 1497869 | 3037785 | 2780248 | 2777154 | 3350456 | **2.881837881** | **0** |
| sp\|P42892 | P42892 | ECE1 | Endothelin-converting enzyme 1 OS=Homo sapiens (Hun | 385798.44 | 1027340.62 | 723287.62 | 795827.69 | 2674448.75 | 2077030.5 | 1310059.62 | 2344665.25 | **2.866805897** | **0** |
| sp\|Q9NW | Q9NWS0 | PIH1D1 | PIH1 domain-containing protein 1 OS=Homo sapiens (Hu | 726332.19 | 718958.94 | 660626 | 562749.25 | 2381741.5 | 1206711.5 | 2110428.5 | 1941882.75 | **2.863139547** | **0** |
| sp\|Q9UL4 | Q9UL40 | ZNF346 | Zinc finger protein 346 OS=Homo sapiens (Human) GN=Z | 0 | 1671459.75 | 0 | 0 | 2055642 | 0 | 2704321 | 0 | **2.847787989** | **5** |
| sp\|Q8IUR | Q8IUR7 | ARMC8 | Armadillo repeat-containing protein 8 OS=Homo sapiens | 0 | 0 | 522688.81 | 0 | 481297.06 | 0 | 429151.66 | 569081.69 | **2.830614281** | **4.5** |
| sp\|Q9BSV | Q9BSV6 | TSEN34 | tRNA-splicing endonuclease subunit Sen34 OS=Homo sa | 0 | 0 | 0 | 218799.75 | 365025.16 | 253767.58 | 0 | 0 | **2.828123615** | **5** |
| sp\|P06213 | P06213 | INSR | Insulin receptor OS=Homo sapiens (Human) GN=INSR PE | 106612.85 | 172985.98 | 0 | 214562.08 | 298078.06 | 131139.34 | 425630.03 | 537012.75 | **2.816613277** | **2** |
| sp\|P05121 | P05121 | SERPINE1 | Plasminogen activator inhibitor 1 OS=Homo sapiens (Hu | 877078.38 | 633437.25 | 475196.62 | 720418 | 2164460 | 3825302 | 773230.56 | 843249.12 | **2.810744856** | **2** |
| sp\|Q9Y61 | Q9Y613 | FHOD1 | FH1/FH2 domain-containing protein 1 OS=Homo sapiens | 138391.81 | 141475.02 | 144459.62 | 0 | 343347.09 | 311960.72 | 272861 | 260985.88 | **2.802452428** | **0** |
| sp\|Q9HD4 | Q9HD42 | CHMP1A | Charged multivesicular body protein 1a OS=Homo sapie | 1027203.56 | 0 | 1624597.62 | 1541496.5 | 3043774.5 | 2056443.75 | 3894245 | 2599789.75 | **2.764948707** | **0** |
| sp\|P07311 | P07311 | ACYP1 | Acylphosphatase-1 OS=Homo sapiens (Human) GN=ACYI | 505818.03 | 0 | 457712.09 | 528169.44 | 1063926.38 | 914550.56 | 1184556.25 | 943847.19 | **2.753155186** | **0** |
| sp\|P04792 | P04792 | HSPB1 | Heat shock protein beta-1 OS=Homo sapiens (Human) G | 31893816 | 31452740 | 27545484 | 28092532 | 96173192 | 77045456 | 82697416 | 69611944 | **2.735884178** | **0** |
| sp\|Q9Y2Q | Q9Y2Q3 | GSTK1 | Glutathione S-transferase kappa 1 OS=Homo sapiens (Hu | 1234328.25 | 857584 | 2515727.25 | 2745574.5 | 4443058 | 4004972 | 6075618 | 5515377 | **2.725206284** | **0** |
| sp\|P52758 | P52758 | RIDA | 2-iminobutanoate/2-iminopropanoate deaminase OS=H | 253061.53 | 464150.56 | 819537.69 | 350928.28 | 1355497.88 | 1243712.75 | 1416569.5 | 1111194.75 | **2.716021862** | **0** |
| sp\|Q1342 | Q13423 | NNT | NAD(P) transhydrogenase, mitochondrial OS=Homo sapi | 11402562 | 10994519 | 8947141 | 11862530 | 31667080 | 27546158 | 24387716 | 33394128 | **2.707796272** | **0** |
| sp\|O9516 | O95169 | NDUFB8 | NADH dehydrogenase [ubiquinone] 1 beta subcomplex s | 1397725.12 | 0 | 1384498.88 | 0 | 1947861.25 | 1867023.75 | 1574543.25 | 2127256.25 | **2.701682** | **0** |
| sp\|P49773 | P49773 | HINT1 | Histidine triad nucleotide-binding protein 1 OS=Homo sa | 12597216 | 10146362 | 10108583 | 8642506 | 25548900 | 33883416 | 28476638 | 21774422 | **2.643312597** | **0** |

| | | | | | | | | | | | | | |
|---|---|---|---|---|---|---|---|---|---|---|---|---|---|
| sp\|O1497 | O14975 | SLC27A2 | Very long-chain acyl-CoA synthetase OS=Homo sapiens ( | 1310466.75 | 896959.12 | 710983.5 | 1287659.75 | 2699496 | 2444614 | 2992484.5 | 2931531.75 | **2.631465612** | 0 |
| sp\|Q96TA | Q96TA2 | YME1L1 | ATP-dependent zinc metalloprotease YME1L1 OS=Homo | 0 | 582456.75 | 0 | 0 | 662893.38 | 869683.44 | 0 | 0 | **2.631228533** | 5 |
| sp\|Q9ULZ | Q9ULZ3 | PYCARD | Apoptosis-associated speck-like protein containing a CAR | 489800.28 | 751869.88 | 421372.81 | 425849.59 | 1268334 | 1446903.75 | 1460176.25 | 1299278.88 | **2.620859007** | 0 |
| sp\|Q5T2W | Q5T2W1 | PDZK1 | Na(+)/H(+) exchange regulatory cofactor NHE-RF3 OS=H | 0 | 622531.06 | 762640.31 | 894130.19 | 1088748.62 | 1253281.12 | 1226144.88 | 2349720.75 | **2.596363497** | 0 |
| sp\|Q9NRY | Q9NRY5 | FAM114A2 | Protein FAM114A2 OS=Homo sapiens (Human) GN=FAM | 0 | 381739.38 | 229003.67 | 603521.88 | 752126.62 | 803491.75 | 739038 | 846309.5 | **2.586722051** | 0 |
| sp\|P04233 | P04233 | CD74 | HLA class II histocompatibility antigen gamma chain OS= | 2654465.5 | 1712148.38 | 1882297.88 | 3237352.5 | 6457551.5 | 4448807 | 7054829 | 6535310 | **2.582312365** | 0 |
| sp\|O4382 | O43824 | GTPBP6 | Putative GTP-binding protein 6 OS=Homo sapiens (Huma | 0 | 0 | 0 | 885294.31 | 1113923 | 0 | 1161089.12 | 0 | **2.569780574** | 5 |
| sp\|Q9H29 | Q9H299 | SH3BGRL3 | SH3 domain-binding glutamic acid-rich-like protein 3 OS | 1022063.25 | 1427213 | 2022849.38 | 1758592.75 | 3515237.5 | 3930771.25 | 4349451 | 4207067 | **2.568327723** | 0 |
| sp\|P55157 | P55157 | MTTP | Microsomal triglyceride transfer protein large subunit O | 35355724 | 38526944 | 34633100 | 32639672 | 88494632 | 85237648 | 94440000 | 94075488 | **2.566303984** | 0 |
| sp\|Q6IN84 | Q6IN84 | MRM1 | rRNA methyltransferase 1, mitochondrial OS=Homo sapi | 1252772 | 1327911.75 | 1273699.88 | 702132.25 | 3259361.25 | 3115951.75 | 2679497.75 | 2621952.5 | **2.562651718** | 0 |
| sp\|Q9BSE | Q9BSE5 | AGMAT | Agmatinase, mitochondrial OS=Homo sapiens (Human) ( | 41165700 | 42034460 | 39046840 | 35609724 | 115655616 | 105301720 | 74782144 | 108167176 | **2.558691488** | 0 |
| sp\|P52209 | P52209 | PGD | 6-phosphogluconate dehydrogenase, decarboxylating O | 96118104 | 99048808 | 110003328 | 108864368 | 262629568 | 254916192 | 290986432 | 248844160 | **2.553835674** | 0 |
| sp\|Q7L3T | Q7L3T8 | PARS2 | Probable proline--tRNA ligase, mitochondrial OS=Homo | 0 | 0 | 325266.47 | 361410.25 | 478451.25 | 363548.03 | 494836 | 394538.06 | **2.52138057** | 0 |
| sp\|Q5VW | Q5VWZ2 | LYPLAL1 | Lysophospholipase-like protein 1 OS=Homo sapiens (Hu | 481996.78 | 0 | 363556.84 | 0 | 361588.62 | 713810.88 | 600867.19 | 455576.25 | **2.521239209** | 3 |
| sp\|P50225 | P50225 | SULT1A1 | Sulfotransferase 1A1 OS=Homo sapiens (Human) GN=SU | 47459464 | 50503716 | 51072836 | 46988840 | 138737216 | 119953552 | 126671008 | 108857360 | **2.521206474** | 0 |
| sp\|Q9288 | Q92887 | ABCC2 | Canalicular multispecific organic anion transporter 1 OS= | 2899740.75 | 4049746.25 | 4816422 | 2236307.5 | 9249059 | 8516045 | 7988039.5 | 9501023 | **2.517756135** | 0 |
| sp\|O0013 | O00139 | KIF2A | Kinesin-like protein KIF2A OS=Homo sapiens (Human) G | 0 | 0 | 0 | 633527.62 | 437940.84 | 498884.5 | 0 | 650600.94 | **2.50569388** | 4.5 |
| sp\|P13804 | P13804 | ETFA | Electron transfer flavoprotein subunit alpha, mitochondr | 44517736 | 46374120 | 36962144 | 43196320 | 105877048 | 101959464 | 116782128 | 103830776 | **2.504815051** | 0 |
| sp\|P16949 | P16949 | STMN1 | Stathmin OS=Homo sapiens (Human) GN=STMN1 PE=1 S | 1647199.75 | 1369843.5 | 1570072 | 1592098.88 | 3942722.5 | 3805711.25 | 3819088 | 3903853.5 | **2.503777167** | 0 |
| sp\|O4348 | O43488 | AKR7A2 | Aflatoxin B1 aldehyde reductase member 2 OS=Homo sa | 11748556 | 11511897 | 11487182 | 9857156 | 28732592 | 28022344 | 28603674 | 25896234 | **2.494235294** | 0 |
| sp\|P21912 | P21912 | SDHB | Succinate dehydrogenase [ubiquinone] iron-sulfur subun | 5710995 | 7080867.5 | 3113497.5 | 7396638 | 14536979 | 13236604 | 15355888 | 14772253 | **2.484839455** | 0 |
| sp\|Q8NHP | Q8NHP8 | PLBD2 | Putative phospholipase B-like 2 OS=Homo sapiens (Hum | 708406.44 | 541368.5 | 486351.47 | 534572.5 | 1584230.75 | 1575295.5 | 854023.19 | 1625068.12 | **2.483207939** | 0 |
| sp\|Q6KB6 | Q6KB66 | KRT80 | Keratin, type II cytoskeletal 80 OS=Homo sapiens (Huma | 1468213.12 | 1607699.12 | 0 | 0 | 2119209.75 | 1867971.5 | 2055017.62 | 1564948 | **2.473135212** | 1 |
| sp\|Q8N0X | Q8N0X4 | CLYBL | Citramalyl-CoA lyase, mitochondrial OS=Homo sapiens (I | 0 | 0 | 0 | 218942.25 | 235889.16 | 304379.84 | 0 | 0 | **2.467632446** | 5 |
| sp\|Q9UEY | Q9UEY8 | ADD3 | Gamma-adducin OS=Homo sapiens (Human) GN=ADD3 I | 459717.22 | 1998014.75 | 1698414.5 | 1972735.5 | 4120721 | 3745302 | 3980122.75 | 3219108 | **2.458075359** | 0 |
| sp\|Q5K4L | Q5K4L6 | SLC27A3 | Long-chain fatty acid transport protein 3 OS=Homo sapie | 137243.83 | 65562.52 | 0 | 81650.89 | 146775.69 | 207694.38 | 192210.59 | 150819.34 | **2.452038134** | 0 |
| sp\|Q96IU4 | Q96IU4 | ABHD14B | Protein ABHD14B OS=Homo sapiens (Human) GN=ABHD | 7070621.5 | 7127798 | 7244941.5 | 7749254 | 17226536 | 27145558 | 13429303 | 13753919 | **2.451144442** | 0 |
| sp\|O75381 | O75381 | PEX14 | Peroxisomal membrane protein PEX14 OS=Homo sapien | 1186778 | 905055.81 | 975086.19 | 990442.94 | 2368813.25 | 2232787.25 | 2284686.25 | 3025840.5 | **2.442997434** | 0 |
| sp\|Q9H4M | Q9H4M3 | FBXO44 | F-box only protein 44 OS=Homo sapiens (Human) GN=FE | 127263.05 | 119725.61 | 0 | 146484.61 | 247824.33 | 300851.59 | 249947.06 | 159334.59 | **2.43461918** | 0 |
| sp\|Q9BRX | Q9BRX8 | PRXL2A | Peroxiredoxin-like 2A OS=Homo sapiens (Human) GN=PI | 9145632 | 7836277.5 | 9418179 | 10474832 | 21421588 | 22108694 | 23438138 | 21665676 | **2.403641684** | 0 |
| sp\|Q7Z4S | Q7Z4S6 | KIF21A | Kinesin-like protein KIF21A OS=Homo sapiens (Human) ( | 90640.52 | 100560.96 | 0 | 423667.38 | 81041.16 | 597200.31 | 695026 | 103454.93 | **2.40168676** | 4 |
| sp\|Q9NZJ | Q9NZJ7 | MTCH1 | Mitochondrial carrier homolog 1 OS=Homo sapiens (Hur | 0 | 0 | 0 | 1128530.75 | 1374715.5 | 1330012.5 | 0 | 0 | **2.396680817** | 5 |
| sp\|Q9BY4 | Q9BY49 | PECR | Peroxisomal trans-2-enoyl-CoA reductase OS=Homo sap | 5786097 | 5786219 | 4123698.5 | 6608819 | 13542629 | 12586054 | 13848450 | 13418670 | **2.39391175** | 0 |
| sp\|Q7LBC | Q7LBC6 | KDM3B | Lysine-specific demethylase 3B OS=Homo sapiens (Huma | 1864654.5 | 930106.81 | 1646611 | 1094853.75 | 1999351.75 | 2971479.25 | 3906462.5 | 4302192.5 | **2.38059029** | 0 |
| sp\|Q8NI6 | Q8NI60 | COQ8A | Atypical kinase COQ8A, mitochondrial OS=Homo sapiens | 1180317 | 1555865.88 | 803267.38 | 921058.69 | 3435892 | 1376467.12 | 4266887 | 1499390.25 | **2.371620927** | 2 |
| sp\|Q6KC7 | Q6KC79 | NIPBL | Nipped-B-like protein OS=Homo sapiens (Human) GN=N | 152609.62 | 611604.06 | 155407.98 | 586565 | 813826.88 | 823227.69 | 927375 | 1002351.38 | **2.368086934** | 0 |
| sp\|Q9BY3 | Q9BY32 | ITPA | Inosine triphosphate pyrophosphatase OS=Homo sapien | 9433744 | 8882926 | 6561798 | 8403102 | 18144470 | 19925250 | 20050824 | 20511002 | **2.362615285** | 0 |
| sp\|Q9H33 | Q9H330 | TMEM245 | Transmembrane protein 245 OS=Homo sapiens (Human) | 1109262.25 | 613930.56 | 603206.25 | 469059.03 | 1484010.25 | 1380068.75 | 1850835.88 | 1884345.12 | **2.360707901** | 0 |
| sp\|P38117 | P38117 | ETFB | Electron transfer flavoprotein subunit beta OS=Homo sa | 69914688 | 68991712 | 75938976 | 74488200 | 170688608 | 159047216 | 181039456 | 172028944 | **2.359920454** | 0 |
| sp\|Q0281 | Q02818 | NUCB1 | Nucleobindin-1 OS=Homo sapiens (Human) GN=NUCB1 I | 2161966.5 | 2899795.5 | 1293133.25 | 1586204.25 | 4980148 | 3848731 | 5208517 | 4650030 | **2.353254231** | 0 |
| sp\|Q5SSJ5 | Q5SSJ5 | HP1BP3 | Heterochromatin protein 1-binding protein 3 OS=Homo | 19446698 | 14438088 | 13766743 | 16835306 | 37705964 | 35456988 | 41446384 | 36667564 | **2.345857104** | 0 |
| sp\|Q96HR | Q96HR9 | REEP6 | Receptor expression-enhancing protein 6 OS=Homo sapi | 11567235 | 9760877 | 10012294 | 16845402 | 31647286 | 27763962 | 34187864 | 19417702 | **2.345437769** | 0 |
| sp\|P15954 | P15954 | COX7C | Cytochrome c oxidase subunit 7C, mitochondrial OS=Ho | 3894424 | 3472328.25 | 2387742.5 | 183869.06 | 4944957 | 5024684 | 6086099.5 | 7203362 | **2.340335184** | 0 |
| sp\|Q9Y69 | Q9Y696 | CLIC4 | Chloride intracellular channel protein 4 OS=Homo sapier | 17891608 | 23262018 | 23848112 | 21778716 | 54226372 | 41256516 | 57239300 | 49978752 | **2.335790269** | 0 |
| sp\|Q53SF7 | Q53SF7 | COBLL1 | Cordon-bleu protein-like 1 OS=Homo sapiens (Human) G | 1151674.38 | 2359829.5 | 2331081 | 892821.38 | 4192412 | 2592267 | 4613077 | 4277886 | **2.327349145** | 0 |

| | | | | | | | | | | | | |
|---|---|---|---|---|---|---|---|---|---|---|---|---|
| sp\|P00439 | P00439 | PAH | Phenylalanine-4-hydroxylase OS=Homo sapiens (Human | 16701266 | 15391943 | 16292966 | 15997817 | 42937080 | 38826104 | 36173232 | 31379586 | **2.319147934** | 0 |
| sp\|O4359 | O43598 | DNPH1 | 2'-deoxynucleoside 5'-phosphate N-hydrolase 1 OS=Hon | 755326.88 | 686605.75 | 990763.44 | 2678275.5 | 5420720.5 | 4230446 | 1073740.62 | 1032092.25 | **2.300345288** | 2 |
| sp\|P48200 | P48200 | IREB2 | Iron-responsive element-binding protein 2 OS=Homo sa | 1437140.88 | 1383564 | 1152939 | 1003832.81 | 3039717 | 3499422 | 2867932 | 2014122.25 | **2.294574935** | 0 |
| sp\|P30047 | P30047 | GCHFR | GTP cyclohydrolase 1 feedback regulatory protein OS=H | 1194735.5 | 1194297.38 | 1467268.5 | 1282899.5 | 2815294 | 3271051 | 2816667 | 2847151 | **2.286379395** | 0 |
| sp\|Q6NXE | Q6NXE6 | ARMC6 | Armadillo repeat-containing protein 6 OS=Homo sapiens | 409485.34 | 347551.75 | 689304.06 | 0 | 1050587.75 | 549789.44 | 1167870.25 | 534813.88 | **2.283735978** | 2 |
| sp\|P16219 | P16219 | ACADS | Short-chain specific acyl-CoA dehydrogenase, mitochond | 2156898.5 | 3774117.25 | 3993055.5 | 4897284 | 7785983 | 9544925 | 7881223 | 8605276 | **2.281667663** | 0 |
| sp\|Q9HCE | Q9HCE6 | ARHGEF10 | Rho guanine nucleotide exchange factor 10-like protein | 4829365 | 3843013.25 | 5091319 | 4891450 | 10248046 | 9904857 | 11801190 | 10427190 | **2.271827846** | 0 |
| sp\|P16930 | P16930 | FAH | Fumarylacetoacetase OS=Homo sapiens (Human) GN=F | 22074278 | 21238416 | 22731658 | 20693678 | 51009072 | 50443992 | 50446932 | 45074068 | **2.270907744** | 0 |
| sp\|Q9H0W | Q9H0W9 | C11orf54 | Ester hydrolase C11orf54 OS=Homo sapiens (Human) GN | 3114213 | 2951037.5 | 2694861.5 | 3118630.5 | 6538714 | 6199572.5 | 7653116.5 | 6446681 | **2.259337131** | 0 |
| sp\|Q9NQ9 | Q9NQ94 | A1CF | APOBEC1 complementation factor OS=Homo sapiens (H | 4494075.5 | 2805968.5 | 5360244 | 3698850.25 | 8831234 | 9310072 | 10254569 | 8475708 | **2.253882964** | 0 |
| sp\|Q1676 | Q16762 | TST | Thiosulfate sulfurtransferase OS=Homo sapiens (Human | 1175881.5 | 1657336.75 | 879257.69 | 533332.81 | 1672375.62 | 2088696.75 | 2559676 | 3229728.25 | **2.249389264** | 0 |
| sp\|Q969H | Q969H8 | MYDGF | Myeloid-derived growth factor OS=Homo sapiens (Hum | 9189610 | 9571614 | 4170254.5 | 10981892 | 17716512 | 20961686 | 18448696 | 19080252 | **2.247112124** | 0 |
| sp\|Q0565 | Q05655 | PRKCD | Protein kinase C delta type OS=Homo sapiens (Human) | 0 | 0 | 1568089.62 | 0 | 1375648.62 | 0 | 1239618.25 | 904809.44 | **2.244818322** | 5.5 |
| sp\|Q9H83 | Q9H832 | UBE2Z | Ubiquitin-conjugating enzyme E2 Z OS=Homo sapiens (H | 1778435.25 | 0 | 3491031 | 0 | 2694355.25 | 2633484 | 3744283.75 | 2746995.75 | **2.242944198** | 3 |
| sp\|Q96CM | Q96CM8 | ACSF2 | Acyl-CoA synthetase family member 2, mitochondrial O | 10549171 | 11470152 | 11463405 | 9086690 | 22071132 | 22321478 | 25290058 | 24833998 | **2.220295001** | 0 |
| sp\|P55061 | P55061 | TMBIM6 | Bax inhibitor 1 OS=Homo sapiens (Human) GN=TMBIM6 | 498690.59 | 0 | 648925.44 | 702437.62 | 1073035.62 | 1142326.62 | 996527.94 | 888377.06 | **2.216296398** | 0 |
| sp\|P0DMN | P0DMM9 | SULT1A3 | Sulfotransferase 1A3 OS=Homo sapiens (Human) GN=SU | 71524320 | 70314472 | 70247392 | 69055064 | 170050592 | 151598080 | 160010400 | 141230816 | **2.215576307** | 0 |
| sp\|P30084 | P30084 | ECHS1 | Enoyl-CoA hydratase, mitochondrial OS=Homo sapiens ( | 36987312 | 39020540 | 41428836 | 45602424 | 96909672 | 77580304 | 99087640 | 86651904 | **2.209466892** | 0 |
| sp\|P20339 | P20339 | RAB5A | Ras-related protein Rab-5A OS=Homo sapiens (Human) | 5553175 | 6398609 | 6303481 | 6951441 | 5840679 | 4688450.5 | 38498348 | 6624522 | **2.207825152** | 8 |
| sp\|P20962 | P20962 | PTMS | Parathymosin OS=Homo sapiens (Human) GN=PTMS PE= | 5651358 | 6813549 | 10151145 | 6683497 | 17278818 | 14942946 | 18600858 | 13826084 | **2.206474441** | 0 |
| sp\|P08670 | P08670 | VIM | Vimentin OS=Homo sapiens (Human) GN=VIM PE=1 SV= | 257347424 | 372447584 | 367803744 | 258032032 | 784428352 | 539655936 | 891311808 | 536842880 | **2.19191741** | 0 |
| sp\|Q96AZ | Q96AZ6 | ISG20 | Interferon-stimulated gene 20 kDa protein OS=Homo sa | 0 | 327608.62 | 0 | 297363.66 | 368916.16 | 0 | 464636.19 | 535795.5 | **2.191053738** | 3 |
| sp\|P19623 | P19623 | SRM | Spermidine synthase OS=Homo sapiens (Human) GN=SR | 14360671 | 16217913 | 14794184 | 17077218 | 37260112 | 32562656 | 34401240 | 32329248 | **2.186601867** | 0 |
| sp\|P11310 | P11310 | ACADM | Medium-chain specific acyl-CoA dehydrogenase, mitoch | 15098281 | 15177802 | 13479075 | 16226994 | 31642206 | 30368758 | 38054600 | 31061444 | **2.186100425** | 0 |
| sp\|O1526 | O15269 | SPTLC1 | Serine palmitoyltransferase 1 OS=Homo sapiens (Huma | 1197054.38 | 0 | 1385728.38 | 0 | 1717980.12 | 1222546.12 | 1333380.25 | 1339644.25 | **2.1734506** | 3 |
| sp\|Q9H6R | Q9H6R3 | ACSS3 | Acyl-CoA synthetase short-chain family member 3, mitoc | 27040812 | 32467758 | 24911962 | 28311954 | 51828432 | 59014584 | 68151328 | 65906976 | **2.172411243** | 0 |
| sp\|Q0315 | Q03154 | ACY1 | Aminoacylase-1 OS=Homo sapiens (Human) GN=ACY1 P | 19143308 | 18146180 | 14964736 | 21079198 | 37303128 | 42866552 | 37648724 | 41216496 | **2.168655105** | 0 |
| sp\|Q8WW | Q8WW59 | SPRYD4 | SPRY domain-containing protein 4 OS=Homo sapiens (H | 0 | 0 | 233642.62 | 413529.66 | 311512.69 | 0 | 500490.38 | 588236.19 | **2.163626755** | 4 |
| sp\|P17301 | P17301 | ITGA2 | Integrin alpha-2 OS=Homo sapiens (Human) GN=ITGA2 | 18335614 | 18473962 | 14040973 | 15870274 | 34586548 | 41565108 | 29199730 | 38894748 | **2.161935772** | 0 |
| sp\|Q9H3G | Q9H3G5 | CPVL | Probable serine carboxypeptidase CPVL OS=Homo sapie | 4141188 | 5125689.5 | 3899086.75 | 4931358 | 8872972 | 8617568 | 9741247 | 11802504 | **2.156909761** | 0 |
| sp\|O7591 | O75911 | DHRS3 | Short-chain dehydrogenase/reductase 3 OS=Homo sapie | 230158.98 | 262901.88 | 305905.94 | 307031.34 | 619342.75 | 582350.5 | 561529 | 621768.62 | **2.156414901** | 0 |
| sp\|P27338 | P27338 | MAOB | Amine oxidase [flavin-containing] B OS=Homo sapiens ( | 1987480.12 | 1995057.38 | 1632021 | 4128063.75 | 3137785 | 5633059.5 | 6841828.5 | 5368217.5 | **2.153515754** | 1 |
| sp\|P07602 | P07602 | PSAP | Prosaposin OS=Homo sapiens (Human) GN=PSAP PE=1 S | 3104561.5 | 2421459 | 1576051.5 | 2777760.25 | 5095345 | 4741925 | 6286211.5 | 5027012.5 | **2.140774607** | 0 |
| sp\|P14927 | P14927 | UQCRB | Cytochrome b-c1 complex subunit 7 OS=Homo sapiens ( | 934159.06 | 0 | 0 | 1011743.25 | 1105110 | 934619.06 | 1157057.88 | 965601.25 | **2.139053008** | 2 |
| sp\|Q7L8L6 | Q7L8L6 | FASTKD5 | FAST kinase domain-containing protein 5, mitochondrial | 4534256.5 | 2222120.25 | 3470682 | 4442967 | 10485870 | 5203160 | 2784170.5 | 12869379 | **2.136504737** | 3 |
| sp\|P00367 | P00367 | GLUD1 | Glutamate dehydrogenase 1, mitochondrial OS=Homo s | 457732000 | 522932128 | 525673728 | 491263520 | 1096616320 | 1030165440 | 1075343744 | 1056213760 | **2.131726237** | 0 |
| sp\|Q8N1N | Q8N1N4 | KRT78 | Keratin, type II cytoskeletal 78 OS=Homo sapiens (Huma | 174117568 | 273158624 | 231614864 | 165675168 | 509762016 | 350595296 | 587432128 | 346278848 | **2.124248208** | 0 |
| sp\|Q96C0 | Q96C01 | FAM136A | Protein FAM136A OS=Homo sapiens (Human) GN=FAM1 | 1913274.62 | 3943344.5 | 1429318.38 | 1473068.38 | 1663913.12 | 4901191 | 5854597.5 | 6174466 | **2.122862785** | 2 |
| sp\|P54802 | P54802 | NAGLU | Alpha-N-acetylglucosaminidase OS=Homo sapiens (Hum | 210844.42 | 248242 | 609806.75 | 248844.66 | 617028.94 | 769491.56 | 705462.06 | 699168.56 | **2.118138416** | 0 |
| sp\|Q9H1E | Q9H1E5 | TMX4 | Thioredoxin-related transmembrane protein 4 OS=Hom | 1312513.88 | 1089523.25 | 1510237.38 | 1175397 | 2571008.75 | 3294257.5 | 2477011.5 | 2408290.75 | **2.113062622** | 0 |
| sp\|P62316 | P62316 | SNRPD2 | Small nuclear ribonucleoprotein Sm D2 OS=Homo sapier | 15485808 | 16441843 | 16013880 | 16214629 | 35105784 | 30471676 | 33028104 | 36641864 | **2.108097305** | 0 |
| sp\|P50995 | P50995 | ANXA11 | Annexin A11 OS=Homo sapiens (Human) GN=ANXA11 PE | 14222517 | 15237560 | 15193527 | 14431725 | 31794620 | 28716704 | 32178662 | 31419688 | **2.100515917** | 0 |
| sp\|Q9Y2U | Q9Y2U8 | LEMD3 | Inner nuclear membrane protein Man1 OS=Homo sapier | 417382.22 | 0 | 494538.31 | 616229.81 | 811325.12 | 859252.94 | 951379.75 | 587242.38 | **2.10005528** | 1 |
| sp\|P30043 | P30043 | BLVRB | Flavin reductase (NADPH) OS=Homo sapiens (Human) G | 489660.25 | 935769.38 | 672481.5 | 610974.12 | 1413177.88 | 1758595.5 | 1562678.5 | 954158.06 | **2.09998188** | 0 |
| sp\|Q8TAQ | Q8TAQ2 | SMARCC2 | SWI/SNF complex subunit SMARCC2 OS=Homo sapiens ( | 404095.59 | 254559.19 | 303279.88 | 3087715.5 | 574765.94 | 593789.62 | 3823678.25 | 3489146.75 | **2.09434895** | 2 |

| | | | | | | | | | | | | | |
|---|---|---|---|---|---|---|---|---|---|---|---|---|---|
| sp\|P06733 | P06733 | ENO1 | Alpha-enolase OS=Homo sapiens (Human) GN=ENO1 PE= | 1319959552 | 1456898304 | 1450442240 | 1324270720 | 2891070720 | 3114878208 | 2830483712 | 2779604480 | **2.092387453** | 0 |
| sp\|P16220 | P16220 | CREB1 | Cyclic AMP-responsive element-binding protein 1 OS=Ho | 1382198.12 | 1438352.75 | 1629370 | 1492715.38 | 2455855.5 | 2030575.75 | 3825588.75 | 4118409.5 | **2.091736559** | 0 |
| sp\|Q5TH6 | Q5TH69 | ARFGEF3 | Brefeldin A-inhibited guanine nucleotide-exchange prote | 99997.88 | 310861.25 | 294930.25 | 97607.74 | 353441.94 | 548528.19 | 425583.28 | 344740.47 | **2.081528348** | 0 |
| sp\|P05546 | P05546 | SERPIND1 | Heparin cofactor 2 OS=Homo sapiens (Human) GN=SERP | 952274.75 | 0 | 0 | 892055.25 | 492996.56 | 840802 | 1375258.5 | 1116693 | **2.074330548** | 4 |
| sp\|P31327 | P31327 | CPS1 | Carbamoyl-phosphate synthase [ammonia], mitochondri | 1473621.75 | 1480849.75 | 1447485.25 | 1353907.25 | 3061472 | 2984194.5 | 2862336.5 | 3029628 | **2.073994625** | 0 |
| sp\|Q0612 | Q06124 | PTPN11 | Tyrosine-protein phosphatase non-receptor type 11 OS= | 2235776 | 2341241.25 | 1557163.5 | 1018357.25 | 5067894 | 3720163.5 | 3864023.75 | 2163671.5 | **2.071397978** | 2 |
| sp\|P63208 | P63208 | SKP1 | S-phase kinase-associated protein 1 OS=Homo sapiens (I | 4418499.5 | 5455304 | 4703743.5 | 4955267.5 | 9691822 | 9761106 | 10624287 | 10008045 | **2.052200926** | 0 |
| sp\|C9JRZ8 | C9JRZ8 | AKR1B15 | Aldo-keto reductase family 1 member B15 OS=Homo sap | 16561145 | 20088884 | 18735230 | 17969706 | 39858868 | 36892944 | 43332756 | 29910600 | **2.044785489** | 0 |
| sp\|P13929 | P13929 | ENO3 | Beta-enolase OS=Homo sapiens (Human) GN=ENO3 PE= | 128271664 | 137820192 | 141670672 | 147141168 | 268387648 | 314738080 | 299121120 | 252329520 | **2.044636531** | 0 |
| sp\|P49748 | P49748 | ACADVL | Very long-chain specific acyl-CoA dehydrogenase, mitoch | 40999840 | 44425224 | 39908012 | 40718496 | 82823152 | 88360128 | 81272856 | 86342088 | **2.040319281** | 0 |
| sp\|Q9NR2 | Q9NR28 | DIABLO | Diablo homolog, mitochondrial OS=Homo sapiens (Huma | 1903048.62 | 1734754.62 | 3035271 | 2135589.25 | 4935873 | 4318933.5 | 3863062.75 | 4772248 | **2.030968406** | 0 |
| sp\|P27487 | P27487 | DPP4 | Dipeptidyl peptidase 4 OS=Homo sapiens (Human) GN=D | 22316900 | 24419116 | 29066900 | 30451030 | 49999240 | 52588196 | 50182736 | 62734944 | **2.028208119** | 0 |
| sp\|P05161 | P05161 | ISG15 | Ubiquitin-like protein ISG15 OS=Homo sapiens (Human) | 1562527.62 | 2008124.75 | 2293802.5 | 3402822 | 4108515 | 4503951.5 | 5190786 | 4920124 | **2.020375215** | 0 |
| sp\|Q9Y5N | Q9Y5N5 | N6AMT1 | Methyltransferase N6AMT1 OS=Homo sapiens (Human) | 0 | 101710.66 | 174113.7 | 0 | 182435.91 | 165704.83 | 0 | 208999.42 | **2.019909191** | 4 |
| sp\|Q5T4S | Q5T4S7 | UBR4 | E3 ubiquitin-protein ligase UBR4 OS=Homo sapiens (Hun | 16208767 | 16038217 | 17346606 | 17468792 | 30981068 | 30647146 | 32030384 | 41043104 | **2.008603005** | 0 |
| sp\|O4367 | O43678 | NDUFA2 | NADH dehydrogenase [ubiquinone] 1 alpha subcomplex | 2565062 | 1963579.38 | 1140126 | 2346545 | 4538523 | 2879718.5 | 4355929.5 | 4287763 | **2.003906178** | 0 |
| sp\|P09104 | P09104 | ENO2 | Gamma-enolase OS=Homo sapiens (Human) GN=ENO2 F | 140290656 | 148399760 | 152728384 | 156062384 | 282874400 | 331783808 | 317274848 | 265207392 | **2.003645437** | 0 |
| sp\|Q8IYB8 | Q8IYB8 | SUPV3L1 | ATP-dependent RNA helicase SUPV3L1, mitochondrial O | 723545.81 | 740180.75 | 1437383.62 | 1331886 | 1612892.75 | 2030313 | 2451141.25 | 2374488.75 | **2.00067172** | 0 |
| sp\|O9547 | O95470 | SGPL1 | Sphingosine-1-phosphate lyase 1 OS=Homo sapiens (Hu | 4834897 | 8304021 | 6142134 | 7264701.5 | 12530894 | 14059462 | 10890197 | 15491035 | **1.995482554** | 0 |
| sp\|P56747 | P56747 | CLDN6 | Claudin-6 OS=Homo sapiens (Human) GN=CLDN6 PE=1 S | 828396.38 | 0 | 994069.44 | 996468.12 | 1476582.5 | 1170040.5 | 995123.62 | 1968824.38 | **1.990316595** | 1 |
| sp\|P50135 | P50135 | HNMT | Histamine N-methyltransferase OS=Homo sapiens (Hum | 19738102 | 18774802 | 20736540 | 19257078 | 42540548 | 28937488 | 44834256 | 39828872 | **1.988894171** | 0 |
| sp\|P04181 | P04181 | OAT | Ornithine aminotransferase, mitochondrial OS=Homo sa | 21857506 | 18698484 | 18827248 | 22236470 | 40276288 | 47345252 | 36327644 | 38242912 | **1.987168295** | 0 |
| sp\|Q9Y2I8 | Q9Y2I8 | WDR37 | WD repeat-containing protein 37 OS=Homo sapiens (Hu | 0 | 267643.22 | 202234.02 | 217529.86 | 316566 | 288493.09 | 280203.38 | 476900.38 | **1.981595549** | 0 |
| sp\|Q9Y6N | Q9Y6N5 | SQOR | Sulfide:quinone oxidoreductase, mitochondrial OS=Hom | 4023577 | 7414994 | 5346402 | 5371414 | 10364300 | 10486837 | 11276453 | 11734916 | **1.979677734** | 0 |
| sp\|Q5TDH | Q5TDH0 | DDI2 | Protein DDI1 homolog 2 OS=Homo sapiens (Human) GN= | 12079679 | 11668754 | 9160231 | 8873610 | 25861960 | 18248840 | 20760744 | 17723752 | **1.976802316** | 0 |
| sp\|Q9Y47 | Q9Y478 | PRKAB1 | 5'-AMP-activated protein kinase subunit beta-1 OS=Hom | 371264.06 | 234671.44 | 238754.97 | 315846.12 | 510715.75 | 690315.38 | 567341 | 524231.97 | **1.975469037** | 0 |
| sp\|Q9NW | Q9NWS8 | RMND1 | Required for meiotic nuclear division protein 1 homolog | 773488.19 | 1193990.75 | 560617.44 | 998366.5 | 2018024.5 | 2193770 | 828457.06 | 1917235.38 | **1.972936389** | 2 |
| sp\|O0074 | O00748 | CES2 | Cocaine esterase OS=Homo sapiens (Human) GN=CES2 P | 10775631 | 11365808 | 10180519 | 9142353 | 19099776 | 19594638 | 22963996 | 20141864 | **1.972787489** | 0 |
| sp\|P36957 | P36957 | DLST | Dihydrolipoyllysine-residue succinyltransferase compone | 21023272 | 25373354 | 25342652 | 25404846 | 44550048 | 44134432 | 52022352 | 50842184 | **1.971802391** | 0 |
| sp\|O9567 | O95678 | KRT75 | Keratin, type II cytoskeletal 75 OS=Homo sapiens (Huma | 401233984 | 562608640 | 515207168 | 386621600 | 980107264 | 815276544 | 1129785088 | 750099904 | **1.969944341** | 0 |
| sp\|P55263 | P55263 | ADK | Adenosine kinase OS=Homo sapiens (Human) GN=ADK P | 10998537 | 11188166 | 12849111 | 15203887 | 24539000 | 21702592 | 27504888 | 25170960 | **1.968909807** | 0 |
| sp\|Q9HAB | Q9HAB8 | PPCS | Phosphopantothenate--cysteine ligase OS=Homo sapien | 1823577.38 | 1576521.75 | 3033340.75 | 3281044.5 | 3470652.25 | 4953021 | 3799411.5 | 6862144.5 | **1.964615774** | 0 |
| sp\|P07741 | P07741 | APRT | Adenine phosphoribosyltransferase OS=Homo sapiens (I | 42493916 | 42348744 | 48383888 | 47936404 | 89665520 | 89137424 | 87692304 | 89354712 | **1.964253486** | 0 |
| sp\|Q9UBI1 | Q9UBI1 | COMMD3 | COMM domain-containing protein 3 OS=Homo sapiens ( | 277148.5 | 0 | 345532.91 | 343669.38 | 467761.88 | 470298.59 | 492785.59 | 464694.09 | **1.961544575** | 0 |
| sp\|O95721 | O95721 | SNAP29 | Synaptosomal-associated protein 29 OS=Homo sapiens ( | 371099.28 | 0 | 0 | 324878.97 | 0 | 454021.38 | 497343.12 | 409487.59 | **1.955308359** | 3 |
| sp\|Q9NP9 | Q9NP97 | DYNLRB1 | Dynein light chain roadblock-type 1 OS=Homo sapiens (H | 1564739.25 | 1161627.75 | 0 | 1011975.5 | 1415848.25 | 2176682.5 | 1997539.88 | 1714345.88 | **1.953918484** | 1 |
| sp\|Q1687 | Q16877 | PFKFB4 | 6-phosphofructo-2-kinase/fructose-2,6-bisphosphatase 4 | 261438.83 | 414516.88 | 276452.41 | 316650.84 | 371725.94 | 1358112 | 305212.66 | 441791.47 | **1.951715522** | 3 |
| sp\|Q9UIJ7 | Q9UIJ7 | AK3 | GTP:AMP phosphotransferase AK3, mitochondrial OS=H | 15824702 | 14598196 | 10860296 | 14500737 | 28724520 | 28117668 | 27743778 | 24211880 | **1.950343836** | 0 |
| sp\|P08397 | P08397 | HMBS | Porphobilinogen deaminase OS=Homo sapiens (Human) | 2249904.5 | 2264183 | 1917839.5 | 4037988.5 | 3416007.5 | 9408686 | 3722364 | 3857150.5 | **1.948841707** | 3 |
| sp\|P05787 | P05787 | KRT8 | Keratin, type II cytoskeletal 8 OS=Homo sapiens (Human | 1965442944 | 2156036352 | 2522395392 | 2006861056 | 4713939968 | 3733259776 | 4362250240 | 4008085248 | **1.94405837** | 0 |
| sp\|Q8TDY | Q8TDY4 | ASAP3 | Arf-GAP with SH3 domain, ANK repeat and PH domain-cc | 0 | 188811.34 | 0 | 0 | 0 | 366629.31 | 0 | 0 | **1.941775902** | 7.5 |
| sp\|Q96EF | Q96EF6 | FBXO17 | F-box only protein 17 OS=Homo sapiens (Human) GN=FE | 166319.5 | 118887.66 | 118099.23 | 127215.56 | 238430.2 | 374001.53 | 227312.56 | 189765.97 | **1.940561102** | 0 |
| sp\|Q9Y3C | Q9Y3C6 | PPIL1 | Peptidyl-prolyl cis-trans isomerase-like 1 OS=Homo sapie | 5740214 | 3758047 | 5274761 | 4127691.5 | 5315796.5 | 4465911 | 22162652 | 4718870.5 | **1.93978021** | 5 |
| sp\|Q1496 | Q14966 | ZNF638 | Zinc finger protein 638 OS=Homo sapiens (Human) GN=Z | 0 | 0 | 868791.62 | 650736 | 1002376.38 | 943101.75 | 999703.19 | 0 | **1.938221643** | 3 |
| sp\|Q6P1X | Q6P1X6 | C8orf82 | UPF0598 protein C8orf82 OS=Homo sapiens (Human) GN | 739222.38 | 498215.12 | 653263.62 | 413515.94 | 1061981.25 | 1123368.25 | 1149232 | 1124776.88 | **1.935303083** | 0 |

| | | | | | | | | | | | | |
|---|---|---|---|---|---|---|---|---|---|---|---|---|
| sp\|Q1457 | Q14573 | ITPR3 | Inositol 1,4,5-trisphosphate receptor type 3 OS=Homo s | 1133483.62 | 1010027.88 | 1431275.75 | 2694849.25 | 2245859.5 | 2874937 | 3389380.5 | 3564370.25 | **1.925876763** | 1 |
| sp\|Q9252 | Q92520 | FAM3C | Protein FAM3C OS=Homo sapiens (Human) GN=FAM3C | 0 | 81924.06 | 0 | 121609.02 | 68684.67 | 91562.51 | 108477.11 | 122694.9 | **1.923123209** | 4 |
| sp\|P08236 | P08236 | GUSB | Beta-glucuronidase OS=Homo sapiens (Human) GN=GUS | 749046.25 | 745827.88 | 0 | 0 | 898440.12 | 941199.38 | 1034064.06 | 0 | **1.922371591** | 3 |
| sp\|P35754 | P35754 | GLRX | Glutaredoxin-1 OS=Homo sapiens (Human) GN=GLRX PE | 5269204 | 4016947 | 3169070.5 | 4376946.5 | 8516103 | 7019781 | 9188454 | 7546943 | **1.917238528** | 0 |
| sp\|Q9Y5J9 | Q9Y5J9 | TIMM8B | Mitochondrial import inner membrane translocase subu | 1738451.38 | 1675322.38 | 1280504.75 | 0 | 7179944 | 0 | 0 | 1810164.25 | **1.915120339** | 7 |
| sp\|P34897 | P34897 | SHMT2 | Serine hydroxymethyltransferase, mitochondrial OS=Ho | 132280024 | 144773408 | 129549696 | 144910240 | 279450560 | 263129920 | 256024976 | 256186288 | **1.912540666** | 0 |
| sp\|Q8N98 | Q8N983 | MRPL43 | 39S ribosomal protein L43, mitochondrial OS=Homo sap | 2674224 | 2693912 | 1530986.75 | 4420412 | 6528447 | 4188234 | 5870225 | 5042594 | **1.91081175** | 1 |
| sp\|Q9942 | Q99426 | TBCB | Tubulin-folding cofactor B OS=Homo sapiens (Human) G | 3069728.5 | 2844749.75 | 2558584 | 3239598.75 | 10339548 | 4992657.5 | 3636502.75 | 3362328.25 | **1.906572426** | 0 |
| sp\|Q1420 | Q14203 | DCTN1 | Dynactin subunit 1 OS=Homo sapiens (Human) GN=DCT | 5952056 | 4077813 | 5112711 | 3561889.5 | 11206641 | 9690899 | 5592091.5 | 9161487 | **1.906021366** | 1 |
| sp\|P52630 | P52630 | STAT2 | Signal transducer and activator of transcription 2 OS=Ho | 197167.31 | 270265.59 | 208851 | 240767.11 | 350099.91 | 519404.44 | 363320.28 | 512776.53 | **1.903494071** | 0 |
| sp\|P21796 | P21796 | VDAC1 | Voltage-dependent anion-selective channel protein 1 O | 121914704 | 211426960 | 200210448 | 206670048 | 339048960 | 332437792 | 330776480 | 405216896 | **1.901429333** | 0 |
| sp\|Q9NPH | Q9NPH2 | ISYNA1 | Inositol-3-phosphate synthase 1 OS=Homo sapiens (Hum | 6764158.5 | 5645315 | 5816541 | 7627429.5 | 12247442 | 12090554 | 11625659 | 13007077 | **1.894166673** | 0 |
| sp\|Q6IAN | Q6IAN0 | DHRS7B | Dehydrogenase/reductase SDR family member 7B OS=H | 945740.94 | 1463469 | 970069.38 | 1665120.75 | 1900150.5 | 2678770.75 | 2345620.5 | 2624948 | **1.893087308** | 0 |
| sp\|Q9NPJ | Q9NPJ3 | ACOT13 | Acyl-coenzyme A thioesterase 13 OS=Homo sapiens (Hu | 4731357 | 4849142 | 4775622 | 5335972 | 9345457 | 8980032 | 10503334 | 8399105 | **1.89050133** | 0 |
| sp\|O7596 | O75962 | TRIO | Triple functional domain protein OS=Homo sapiens (Hu | 0 | 133958.47 | 104227.06 | 204285.73 | 113400.22 | 190734.44 | 287912.88 | 243996.17 | **1.889487037** | 3 |
| sp\|Q9NYL | Q9NYL4 | FKBP11 | Peptidyl-prolyl cis-trans isomerase FKBP11 OS=Homo sa | 2366547.25 | 1857228.62 | 1903507 | 2049282 | 3572910 | 3794372.75 | 4665638 | 3307764.5 | **1.876177282** | 0 |
| sp\|O9533 | O95336 | PGLS | 6-phosphogluconolactonase OS=Homo sapiens (Human) | 8561142 | 7855025 | 7230503 | 8125465 | 16108057 | 14392549 | 15496368 | 13518992 | **1.873212675** | 0 |
| sp\|P0DN7 | P0DN79 | CBSL | Cystathionine beta-synthase-like protein OS=Homo sapi | 2501367 | 4813899 | 2775838 | 3956934 | 6378578.5 | 5559543 | 7640070 | 6736286 | **1.873178126** | 0 |
| sp\|O14981 | O14981 | BTAF1 | TATA-binding protein-associated factor 172 OS=Homo s | 2312179.5 | 1267487 | 1496247.25 | 1445758 | 2667132.5 | 3545794.5 | 2928125 | 3057728.75 | **1.87049904** | 0 |
| sp\|P05783 | P05783 | KRT18 | Keratin, type I cytoskeletal 18 OS=Homo sapiens (Huma | 1144753280 | 1168168576 | 1176185344 | 1231371904 | 2077995520 | 2138548480 | 2331776512 | 2262234624 | **1.866453583** | 0 |
| sp\|P50748 | P50748 | KNTC1 | Kinetochore-associated protein 1 OS=Homo sapiens (Hu | 0 | 574102.25 | 0 | 462981.09 | 515599 | 415921.03 | 479521.81 | 522808.91 | **1.864701394** | 5 |
| sp\|O95081 | O95081 | AGFG2 | Arf-GAP domain and FG repeat-containing protein 2 OS= | 4802582 | 1979773.62 | 1275319.5 | 1718986.62 | 4194184 | 6157724 | 3072480 | 4783784.5 | **1.862412036** | 3 |
| sp\|P30048 | P30048 | PRDX3 | Thioredoxin-dependent peroxide reductase, mitochondr | 110717592 | 108458560 | 95977648 | 104800752 | 196504976 | 201065872 | 196657344 | 187365024 | **1.861137621** | 0 |
| sp\|P61457 | P61457 | PCBD1 | Pterin-4-alpha-carbinolamine dehydratase OS=Homo sa | 246710.41 | 330291.28 | 272493.47 | 524553.25 | 532858.12 | 653075.81 | 617949.69 | 751893.94 | **1.860034582** | 0 |
| sp\|Q9P0M | Q9P0M6 | H2AFY2 | Core histone macro-H2A.2 OS=Homo sapiens (Human) G | 22081578 | 23736684 | 22977780 | 21127386 | 47099428 | 35728004 | 40386040 | 43969440 | **1.859169693** | 0 |
| sp\|Q9NQV | Q9NQW7 | XPNPEP1 | Xaa-Pro aminopeptidase 1 OS=Homo sapiens (Human) G | 4208530.5 | 4067291.5 | 4049327 | 4650569.5 | 7976890 | 8009937 | 8106198 | 7405109 | **1.85548164** | 0 |
| sp\|P02538 | P02538 | KRT6A | Keratin, type II cytoskeletal 6A OS=Homo sapiens (Huma | 310047872 | 543825472 | 393934560 | 307728384 | 767274880 | 653333888 | 874072064 | 588354240 | **1.853402646** | 0 |
| sp\|A0MZ6 | A0MZ66 | SHTN1 | Shootin-1 OS=Homo sapiens (Human) GN=SHTN1 PE=1 S | 6687259.5 | 6964947 | 9497178 | 5179845 | 12141081 | 11425069 | 15268030 | 13616556 | **1.851470616** | 0 |
| sp\|P36955 | P36955 | SERPINF1 | Pigment epithelium-derived factor OS=Homo sapiens (H | 1405641.25 | 200877.59 | 795269.88 | 886999.88 | 1116804.25 | 727044.69 | 2064296.5 | 2177842.5 | **1.850525735** | 4 |
| sp\|P62253 | P62253 | UBE2G1 | Ubiquitin-conjugating enzyme E2 G1 OS=Homo sapiens ( | 0 | 623612.75 | 1173072 | 976163.12 | 1453274.12 | 1245603.75 | 1342783.38 | 1085591.88 | **1.849092835** | 1 |
| sp\|Q8N33 | Q8N335 | GPD1L | Glycerol-3-phosphate dehydrogenase 1-like protein OS= | 853236.44 | 794051.5 | 811163.19 | 910413.88 | 1271616 | 1333372.38 | 1328074 | 2286800.75 | **1.846278527** | 0 |
| sp\|P10606 | P10606 | COX5B | Cytochrome c oxidase subunit 5B, mitochondrial OS=Ho | 4461644 | 3870066.75 | 4269113 | 4967355.5 | 8526809 | 7011999.5 | 8507369 | 8349807 | **1.844014911** | 0 |
| sp\|O7556 | O75569 | PRKRA | Interferon-inducible double-stranded RNA-dependent pr | 3975953.5 | 4178149.75 | 4189408.5 | 3553628.5 | 7677243.5 | 7075761 | 7688838.5 | 6807962 | **1.839941306** | 0 |
| sp\|Q4G17 | Q4G176 | ACSF3 | Acyl-CoA synthetase family member 3, mitochondrial O | 1810468.62 | 1851931 | 2055345 | 2007246.62 | 2969355 | 3487130 | 3746191.75 | 4008808.25 | **1.83967652** | 0 |
| sp\|Q9P2J5 | Q9P2J5 | LARS | Leucine--tRNA ligase, cytoplasmic OS=Homo sapiens (Hu | 22948930 | 28939322 | 24003650 | 25844032 | 44176972 | 42469648 | 48934980 | 51247736 | **1.836414418** | 0 |
| sp\|O6076 | O60762 | DPM1 | Dolichol-phosphate mannosyltransferase subunit 1 OS= | 2851096 | 2675597.5 | 2157317.75 | 3406289.75 | 4296659.5 | 5401125 | 4509102.5 | 6110019 | **1.831952622** | 0 |
| sp\|P13647 | P13647 | KRT5 | Keratin, type II cytoskeletal 5 OS=Homo sapiens (Human | 320724896 | 549904448 | 406996544 | 315682752 | 776515200 | 657231616 | 885407936 | 596577792 | **1.829986025** | 0 |
| sp\|Q8WW | Q8WW12 | PCNP | PEST proteolytic signal-containing nuclear protein OS=H | 5431484 | 4737924 | 4099612 | 7830189.5 | 12939603 | 6444011 | 11864797 | 9176296 | **1.829237693** | 1 |
| sp\|P46952 | P46952 | HAAO | 3-hydroxyanthranilate 3,4-dioxygenase OS=Homo sapie | 1652205.75 | 1989631.75 | 1913740.12 | 916710.62 | 2882462.25 | 3506020.25 | 2587572 | 2852591 | **1.827583238** | 0 |
| sp\|Q9Y5X | Q9Y5X3 | SNX5 | Sorting nexin-5 OS=Homo sapiens (Human) GN=SNX5 PE | 16008206 | 12386973 | 15557932 | 14438269 | 27501430 | 27189078 | 25609894 | 26258958 | **1.824915938** | 0 |
| sp\|Q0057 | Q00577 | PURA | Transcriptional activator protein Pur-alpha OS=Homo sa | 1507744.62 | 2597381 | 2223290.5 | 2340002.25 | 3739681 | 4154869 | 3939205.5 | 3984590.75 | **1.824824965** | 0 |
| sp\|P21964 | P21964 | COMT | Catechol O-methyltransferase OS=Homo sapiens (Huma | 34406760 | 34437764 | 111543896 | 30488304 | 46823008 | 47226876 | 139166016 | 151042032 | **1.822192249** | 2 |
| sp\|Q8WV | Q8WVC6 | DCAKD | Dephospho-CoA kinase domain-containing protein OS=H | 138944.78 | 910815.31 | 1030656.5 | 850488.25 | 1627525.5 | 1309377.75 | 1190068.25 | 1211300.12 | **1.821373232** | 0 |
| sp\|Q9BZF1 | Q9BZF1 | OSBPL8 | Oxysterol-binding protein-related protein 8 OS=Homo s | 2438558.5 | 2441677.75 | 3159568.5 | 2654803.75 | 4743726.5 | 4995942.5 | 4745146 | 4981873.5 | **1.820233859** | 0 |
| sp\|Q9UKD | Q9UKD2 | MRTO4 | mRNA turnover protein 4 homolog OS=Homo sapiens (H | 11584360 | 11667854 | 13150028 | 11318282 | 21443436 | 19934962 | 21773108 | 23672504 | **1.819427004** | 0 |

| | | | | | | | | | | | | | |
|---|---|---|---|---|---|---|---|---|---|---|---|---|---|
| sp\|P35527 | P35527 | KRT9 | Keratin, type I cytoskeletal 9 OS=Homo sapiens (Human) | 0 | 278066.5 | 177466.61 | 462295.66 | 0 | 1669884.88 | 0 | 0 | **1.819386071** | **5.5** |
| sp\|P30086 | P30086 | PEBP1 | Phosphatidylethanolamine-binding protein 1 OS=Homo | 196655328 | 147403936 | 154920096 | 203851040 | 384565184 | 273974976 | 321043296 | 298949984 | **1.819120858** | **0** |
| sp\|Q8WX | Q8WXX5 | DNAJC9 | DnaJ homolog subfamily C member 9 OS=Homo sapiens | 4308424.5 | 3301101.5 | 1444879.62 | 4132696 | 6055692 | 6256943 | 6417473.5 | 5228826 | **1.816846127** | **0** |
| sp\|Q9NVS | Q9NVS9 | PNPO | Pyridoxine-5'-phosphate oxidase OS=Homo sapiens (Hu | 3593831.5 | 2561620.75 | 4081558.25 | 3573203.5 | 7864950 | 6394787.5 | 5715280 | 5096996 | **1.815468862** | **0** |
| sp\|P29966 | P29966 | MARCKS | Myristoylated alanine-rich C-kinase substrate OS=Homo | 11877940 | 13527608 | 13648606 | 13495084 | 22774324 | 16616292 | 30421476 | 25494560 | **1.813663825** | **0** |
| sp\|O1465 | O14653 | GOSR2 | Golgi SNAP receptor complex member 2 OS=Homo sapie | 262799.28 | 0 | 211667.72 | 341857.91 | 452215.84 | 597271.75 | 0 | 428432.06 | **1.810455165** | **3.5** |
| sp\|P35237 | P35237 | SERPINB6 | Serpin B6 OS=Homo sapiens (Human) GN=SERPINB6 PE= | 17884118 | 20470934 | 18962340 | 18780908 | 33256628 | 35059916 | 33942628 | 35420844 | **1.80923905** | **0** |
| sp\|P49006 | P49006 | MARCKSL1 | MARCKS-related protein OS=Homo sapiens (Human) GN | 8683297 | 10731774 | 7579448.5 | 7514060.5 | 16071700 | 13141201 | 19923000 | 13205470 | **1.80654698** | **0** |
| sp\|P35221 | P35221 | CTNNA1 | Catenin alpha-1 OS=Homo sapiens (Human) GN=CTNNA | 6746225.5 | 5841217 | 12317818 | 4919285 | 17264402 | 10404407 | 12841788 | 13354010 | **1.806049551** | **1** |
| sp\|Q96DA | Q96DA6 | DNAJC19 | Mitochondrial import inner membrane translocase subu | 0 | 0 | 0 | 1654582.5 | 1131712 | 788405.19 | 1061693.25 | 0 | **1.802152773** | **5.5** |
| sp\|Q96EL | Q96EL3 | MRPL53 | 39S ribosomal protein L53, mitochondrial OS=Homo sapi | 165306.55 | 122667.77 | 84018.74 | 0 | 158467.2 | 171810.2 | 198111.56 | 140497.81 | **1.798116261** | **2** |
| sp\|Q9H6V | Q9H6V9 | LDAH | Lipid droplet-associated hydrolase OS=Homo sapiens (H | 1591351.5 | 1590478 | 1561461.75 | 1486518.62 | 2812159.75 | 2632380 | 3007514.25 | 2749671.75 | **1.79808469** | **0** |
| sp\|Q9260 | Q92604 | LPGAT1 | Acyl-CoA:lysophosphatidylglycerol acyltransferase 1 OS= | 956536.06 | 1139244.75 | 1086228.38 | 1033439.56 | 1857260.25 | 1742378.88 | 1935649 | 2039595.88 | **1.796934196** | **0** |
| sp\|P49441 | P49441 | INPP1 | Inositol polyphosphate 1-phosphatase OS=Homo sapien | 928398.62 | 704962.12 | 926718.25 | 977976.19 | 1651895.75 | 1452683.5 | 1875739 | 1376411 | **1.796673293** | **0** |
| sp\|Q8IY17 | Q8IY17 | PNPLA6 | Neuropathy target esterase OS=Homo sapiens (Human) | 256851.59 | 247794.75 | 240589.95 | 225668.28 | 408952.56 | 414256.62 | 557480.12 | 363228.72 | **1.796178609** | **0** |
| sp\|Q8IXM | Q8IXM3 | MRPL41 | 39S ribosomal protein L41, mitochondrial OS=Homo sapi | 3396933 | 2002328.38 | 1664491.62 | 3067920.75 | 4576913 | 4581993 | 4608785 | 4423237 | **1.795451418** | **0** |
| sp\|Q8IVH | Q8IVH2 | FOXP4 | Forkhead box protein P4 OS=Homo sapiens (Human) GN | 0 | 185932.52 | 0 | 220479.17 | 0 | 250135.97 | 251410.53 | 228024.41 | **1.795152374** | **3** |
| sp\|Q9Y3D | Q9Y3D6 | FIS1 | Mitochondrial fission 1 protein OS=Homo sapiens (Huma | 1948310.62 | 1432830 | 1881905.38 | 1716571 | 3424247.25 | 3584571 | 3989224.5 | 1518636.25 | **1.793318888** | **3** |
| sp\|Q9P0S | Q9P0S9 | TMEM14C | Transmembrane protein 14C OS=Homo sapiens (Human | 2069727.62 | 2157500.75 | 958341.38 | 2408396.75 | 3695176 | 3507950 | 2111755.75 | 4297233 | **1.792490756** | **2** |
| sp\|Q8NCN | Q8NCN5 | PDPR | Pyruvate dehydrogenase phosphatase regulatory subuni | 881034.94 | 842677 | 800808.88 | 860211.44 | 1346052.5 | 1285876.25 | 1668592.75 | 1760218.12 | **1.790611237** | **0** |
| sp\|O9568 | O95684 | FGFR1OP | FGFR1 oncogene partner OS=Homo sapiens (Human) GN | 115427.7 | 99106.94 | 69502.12 | 0 | 101535.77 | 215341.23 | 93798.3 | 97035.96 | **1.787484338** | **5** |
| sp\|Q96BS | Q96BS2 | TESC | Calcineurin B homologous protein 3 OS=Homo sapiens (I | 2836323 | 1418424.12 | 2474466 | 1474262 | 4225913.5 | 3932276 | 3404245.5 | 3085697.75 | **1.785600923** | **0** |
| sp\|P82979 | P82979 | SARNP | SAP domain-containing ribonucleoprotein OS=Homo sap | 15361050 | 8363806 | 10473867 | 13096378 | 22989302 | 19743410 | 21953774 | 19657552 | **1.783356758** | **0** |
| sp\|P33240 | P33240 | CSTF2 | Cleavage stimulation factor subunit 2 OS=Homo sapiens | 2701944.5 | 2640027.75 | 2407188.75 | 4471182 | 6065237.5 | 4781268.5 | 5903400.5 | 4977750 | **1.777990724** | **0** |
| sp\|P04259 | P04259 | KRT6B | Keratin, type II cytoskeletal 6B OS=Homo sapiens (Huma | 310047872 | 610677184 | 393934560 | 307728384 | 767274880 | 653333888 | 875080960 | 588354240 | **1.777653661** | **1** |
| sp\|Q1654 | Q16540 | MRPL23 | 39S ribosomal protein L23, mitochondrial OS=Homo sapi | 4266730 | 2921176 | 1468919.75 | 3213562.5 | 5144151.5 | 5501426.5 | 5916273.5 | 4535558 | **1.777314192** | **0** |
| sp\|Q1674 | Q16740 | CLPP | ATP-dependent Clp protease proteolytic subunit, mitoch | 2110754.5 | 2229533.5 | 1109951.38 | 2048182.25 | 3295613.5 | 3280629 | 3307895.25 | 3433045.5 | **1.775998191** | **0** |
| sp\|P45973 | P45973 | CBX5 | Chromobox protein homolog 5 OS=Homo sapiens (Huma | 4340697 | 4035619 | 3543227 | 3174597.25 | 6988659 | 5629865 | 7673500 | 6476283 | **1.773423763** | **0** |
| sp\|O6072 | O60725 | ICMT | Protein-S-isoprenylcysteine O-methyltransferase OS=Ho | 1284260.88 | 1560862 | 1426352 | 1547473.62 | 2149300.5 | 2467096.75 | 2244633.75 | 3456770.5 | **1.77313848** | **0** |
| sp\|Q8IW | Q8IWW6 | ARHGAP1 | Rho GTPase-activating protein 12 OS=Homo sapiens (Hu | 818425.25 | 1518755.25 | 1884985.25 | 557643.75 | 2645271.5 | 2081620.75 | 2880189.25 | 860891.31 | **1.771613034** | **2** |
| sp\|P55957 | P55957 | BID | BH3-interacting domain death agonist OS=Homo sapien | 521813.31 | 1546723.25 | 1371525.62 | 1811889.25 | 2471913.75 | 2031957.38 | 2920869.25 | 1876425.38 | **1.770992341** | **0** |
| sp\|Q1671 | Q16719 | KYNU | Kynureninase OS=Homo sapiens (Human) GN=KYNU PE= | 6859843 | 7056854.5 | 6258786.5 | 6902092.5 | 11559312 | 12882631 | 11936173 | 11559726 | **1.770388942** | **0** |
| sp\|Q86TX | Q86TX2 | ACOT1 | Acyl-coenzyme A thioesterase 1 OS=Homo sapiens (Hum | 26217924 | 26985056 | 24100632 | 26641968 | 44215016 | 43492976 | 48807128 | 47293012 | **1.768311187** | **0** |
| sp\|Q9UJA | Q9UJA5 | TRMT6 | tRNA (adenine(58)-N(1))-methyltransferase non-catalytic | 1002648.31 | 1202612.62 | 1323001.75 | 2433823.5 | 2023570.88 | 2730002 | 3087824 | 2691662.5 | **1.766673453** | **1** |
| sp\|Q96JJ3 | Q96JJ3 | ELMO2 | Engulfment and cell motility protein 2 OS=Homo sapiens | 967311 | 1184132.25 | 1060030.62 | 1108413.88 | 1652809 | 1368429.75 | 1655556 | 2940032 | **1.763200155** | **0** |
| sp\|Q9ULR | Q9ULR3 | PPM1H | Protein phosphatase 1H OS=Homo sapiens (Human) GN= | 603079.12 | 593865.19 | 674259.31 | 673511 | 1425946.25 | 883222.56 | 1262371.38 | 905585.94 | **1.759382406** | **0** |
| sp\|Q8TAV | Q8TAV3 | CYP2W1 | Cytochrome P450 2W1 OS=Homo sapiens (Human) GN= | 2660057.5 | 3281283.5 | 3272355 | 2741300.5 | 5324131.5 | 4905832 | 4745605 | 6038554.5 | **1.757769063** | **0** |
| sp\|Q1382 | Q13825 | AUH | Methylglutaconyl-CoA hydratase, mitochondrial OS=Hor | 781905.12 | 1133840.38 | 1057747.5 | 903028.94 | 1763358.38 | 1239479 | 2016906.5 | 1786534.62 | **1.755769374** | **0** |
| sp\|P08183 | P08183 | ABCB1 | Multidrug resistance protein 1 OS=Homo sapiens (Huma | 5787934 | 7295928 | 6874614 | 6284704 | 11064586 | 11954652 | 10802022 | 12250030 | **1.755552871** | **0** |
| sp\|P78509 | P78509 | RELN | Reelin OS=Homo sapiens (Human) GN=RELN PE=1 SV=3 | 977804.25 | 1898997.5 | 1918430.12 | 1632944.5 | 3201271.25 | 1887209.5 | 3205251 | 2962169.5 | **1.751025579** | **2** |
| sp\|Q9UK2 | Q9UK22 | FBXO2 | F-box only protein 2 OS=Homo sapiens (Human) GN=FB | 27584578 | 26365142 | 22610570 | 22721974 | 39544072 | 44383740 | 44502024 | 45153788 | **1.748385029** | **0** |
| sp\|P53634 | P53634 | CTSC | Dipeptidyl peptidase 1 OS=Homo sapiens (Human) GN= | 9919352 | 13855439 | 11897696 | 13974263 | 23076632 | 20297028 | 22341436 | 21061340 | **1.747877474** | **0** |
| sp\|P49023 | P49023 | PXN | Paxillin OS=Homo sapiens (Human) GN=PXN PE=1 SV=3 | 706022.94 | 721160.81 | 685628.06 | 422474.72 | 1091261.62 | 1023751.56 | 1493523.25 | 821170.75 | **1.747221518** | **0** |
| sp\|Q9H0R | Q9H0R4 | HDHD2 | Haloacid dehalogenase-like hydrolase domain-containin | 774767.75 | 1463972.25 | 851611.56 | 1498456.38 | 2216699.5 | 1921308 | 2121167 | 1756113.75 | **1.746703796** | **0** |
| sp\|P86790 | P86790 | CCZ1B | Vacuolar fusion protein CCZ1 homolog B OS=Homo sapie | 0 | 686723.38 | 947865.56 | 0 | 877257.69 | 0 | 893018 | 1082929.25 | **1.74551832** | **5** |

| | | | | | | | | | | | | | |
|---|---|---|---|---|---|---|---|---|---|---|---|---|---|
| sp\|P50416 | P50416 | CPT1A | Carnitine O-palmitoyltransferase 1, liver isoform OS=Hor | 1911040.75 | 2353922.75 | 2931138.75 | 2395337.5 | 3533353.5 | 3928038.75 | 5199258 | 4028802.75 | **1.740036265** | 0 |
| sp\|O9580 | O95807 | TMEM50A | Transmembrane protein 50A OS=Homo sapiens (Human | 277508.59 | 383809.31 | 0 | 387694.62 | 463272.25 | 493886.03 | 404189.75 | 460784.59 | **1.736997972** | 0 |
| sp\|Q9Y5M | Q9Y5M8 | SRPRB | Signal recognition particle receptor subunit beta OS=Hor | 13864291 | 13466397 | 17818360 | 15381939 | 24463824 | 23276164 | 25954234 | 31321094 | **1.734901762** | 0 |
| sp\|P15289 | P15289 | ARSA | Arylsulfatase A OS=Homo sapiens (Human) GN=ARSA PE | 339492.06 | 359245.94 | 385284.5 | 0 | 439750.78 | 464774.69 | 536389.5 | 439605.59 | **1.734761557** | 0 |
| sp\|P18206 | P18206 | VCL | Vinculin OS=Homo sapiens (Human) GN=VCL PE=1 SV=4 | 30523922 | 35363624 | 45339848 | 34233152 | 68641888 | 55652508 | 66017592 | 61988964 | **1.73449749** | 0 |
| sp\|Q9BTU | Q9BTU6 | PI4K2A | Phosphatidylinositol 4-kinase type 2-alpha OS=Homo sa | 333745.09 | 424875.44 | 502026.69 | 494940.62 | 483538.5 | 881628.88 | 404308.44 | 1273828.25 | **1.733495756** | 5 |
| sp\|P14174 | P14174 | MIF | Macrophage migration inhibitory factor OS=Homo sapie | 60274624 | 39741148 | 48515936 | 59933288 | 93931152 | 80886224 | 114382752 | 71692528 | **1.73119067** | 0 |
| sp\|O7536 | O75367 | H2AFY | Core histone macro-H2A.1 OS=Homo sapiens (Human) G | 77820152 | 77975592 | 74183016 | 76955904 | 144247280 | 125541864 | 134284464 | 126593760 | **1.728926153** | 0 |
| sp\|Q6UW | Q6UWM9 | UGT2A3 | UDP-glucuronosyltransferase 2A3 OS=Homo sapiens (Hu | 647524.44 | 559698.19 | 652463.12 | 450485.84 | 857096.62 | 1005623.06 | 978302.06 | 1146916.88 | **1.726252127** | 0 |
| sp\|Q9Y6K | Q9Y6K1 | DNMT3A | DNA (cytosine-5)-methyltransferase 3A OS=Homo sapier | 252445.33 | 576958.88 | 300106.84 | 393049.97 | 445241.75 | 780146.75 | 667337.12 | 734128.81 | **1.7252868** | 1 |
| sp\|P30040 | P30040 | ERP29 | Endoplasmic reticulum resident protein 29 OS=Homo sa | 59738840 | 50260872 | 51980432 | 56893844 | 107653240 | 82848480 | 100091560 | 86955768 | **1.724960793** | 0 |
| sp\|P03905 | P03905 | MT-ND4 | NADH-ubiquinone oxidoreductase chain 4 OS=Homo sap | 0 | 0 | 0 | 829434.81 | 441542.34 | 540519.19 | 0 | 448455.09 | **1.72468843** | 5.5 |
| sp\|Q9972 | Q99729 | HNRNPAB | Heterogeneous nuclear ribonucleoprotein A/B OS=Hom | 27637286 | 25931918 | 31434522 | 31574602 | 52374648 | 45815300 | 55491952 | 47211412 | **1.723247498** | 0 |
| sp\|Q5T9A | Q5T9A4 | ATAD3B | ATPase family AAA domain-containing protein 3B OS=Hc | 26429650 | 30903484 | 31249908 | 25990240 | 53086512 | 47248112 | 43966992 | 53004036 | **1.722091299** | 0 |
| sp\|Q96TC | Q96TC7 | RMDN3 | Regulator of microtubule dynamics protein 3 OS=Homo | 1257037 | 1054608.38 | 0 | 1255174.25 | 1342790.88 | 1362457.75 | 1691441.12 | 1738643.12 | **1.720113016** | 0 |
| sp\|Q6PIU | Q6PIU2 | NCEH1 | Neutral cholesterol ester hydrolase 1 OS=Homo sapiens | 439913.12 | 1468103.25 | 326246.59 | 587775.25 | 525662.75 | 623362.69 | 1715407.88 | 1986056.88 | **1.718789697** | 3 |
| sp\|Q9962 | Q99624 | SLC38A3 | Sodium-coupled neutral amino acid transporter 3 OS=Hc | 660731 | 726504.44 | 775837.31 | 1073697.25 | 1588894.88 | 1223959.88 | 1334056.5 | 1410593 | **1.716990784** | 0 |
| sp\|P78417 | P78417 | GSTO1 | Glutathione S-transferase omega-1 OS=Homo sapiens (H | 75997208 | 73309088 | 79026912 | 72167776 | 135159152 | 125379440 | 129129744 | 124808272 | **1.712062973** | 0 |
| sp\|Q1390 | Q13907 | IDI1 | Isopentenyl-diphosphate Delta-isomerase 1 OS=Homo s | 4005087.25 | 4240713.5 | 4873835 | 5092314 | 7168557 | 6813817.5 | 7567044 | 9578378 | **1.709196265** | 0 |
| sp\|Q7Z2E | Q7Z2E3 | APTX | Aprataxin OS=Homo sapiens (Human) GN=APTX PE=1 SV | 0 | 667659.38 | 562163.88 | 533221.06 | 594717.5 | 597923 | 714481.75 | 1106201 | **1.709159104** | 2 |
| sp\|Q9NYB | Q9NYB0 | TERF2IP | Telomeric repeat-binding factor 2-interacting protein 1 ( | 1953489.75 | 914481.56 | 888277.25 | 1214876 | 1934808 | 2550946 | 2472443.25 | 1534883.75 | **1.708482839** | 2 |
| sp\|Q0342 | Q03426 | MVK | Mevalonate kinase OS=Homo sapiens (Human) GN=MVK | 4068913.5 | 4445928 | 4184284 | 3904850.25 | 7828145.5 | 6536595.5 | 6147926 | 7845880 | **1.707937149** | 0 |
| sp\|Q3SXM | Q3SXM5 | HSDL1 | Inactive hydroxysteroid dehydrogenase-like protein 1 OS | 993573.88 | 1157099.38 | 1082698.5 | 1137013.62 | 1661585.75 | 1752962.5 | 1734384.12 | 2310698.5 | **1.706858829** | 0 |
| sp\|O0023 | O00231 | PSMD11 | 26S proteasome non-ATPase regulatory subunit 11 OS=H | 54866208 | 83838632 | 59951200 | 57679164 | 53035988 | 119569328 | 128318392 | 135093296 | **1.700964195** | 4 |
| sp\|P35244 | P35244 | RPA3 | Replication protein A 14 kDa subunit OS=Homo sapiens ( | 1778636.38 | 0 | 466631.56 | 1240967.12 | 2051024.12 | 1896680.12 | 762332.5 | 1208636.75 | **1.697726455** | 4 |
| sp\|Q96FQ | Q96FQ6 | S100A16 | Protein S100-A16 OS=Homo sapiens (Human) GN=S100A | 364184.97 | 530406.38 | 501673.28 | 371882.69 | 518550.06 | 681427.12 | 794249.69 | 1005754.88 | **1.69668088** | 1 |
| sp\|Q9UM | Q9UM22 | EPDR1 | Mammalian ependymin-related protein 1 OS=Homo sap | 2825617.5 | 3634027.25 | 1543805.62 | 3222500.5 | 3943185.5 | 6455307 | 4010719.25 | 4634691 | **1.696417789** | 0 |
| sp\|Q8TEM | Q8TEM1 | NUP210 | Nuclear pore membrane glycoprotein 210 OS=Homo sap | 31282554 | 10098393 | 7311363 | 6546221.5 | 35581396 | 33855332 | 10942045 | 13248648 | **1.694965787** | 2 |
| sp\|Q1527 | Q15274 | QPRT | Nicotinate-nucleotide pyrophosphorylase [carboxylating | 32184728 | 32229596 | 31548478 | 30708030 | 55638888 | 53966276 | 56717956 | 48217704 | **1.693687652** | 0 |
| sp\|Q0026 | Q00266 | MAT1A | S-adenosylmethionine synthase isoform type-1 OS=Hom | 15108800 | 14168745 | 16313719 | 13488586 | 31400990 | 24141742 | 24538000 | 19953630 | **1.693206093** | 0 |
| sp\|Q1510 | Q15102 | PAFAH1B3 | Platelet-activating factor acetylhydrolase IB subunit gam | 6918878 | 6176580.5 | 5318157 | 6405984 | 10763500 | 9657418 | 12255690 | 9338415 | **1.692816316** | 0 |
| sp\|Q9Y62 | Q9Y624 | F11R | Junctional adhesion molecule A OS=Homo sapiens (Hum | 3963759.5 | 3849363 | 1766471.62 | 2321054.5 | 5248359 | 5319485 | 4799359.5 | 4770285 | **1.692133693** | 0 |
| sp\|P30711 | P30711 | GSTT1 | Glutathione S-transferase theta-1 OS=Homo sapiens (Hu | 7411906.5 | 6280197 | 5358989.5 | 6426939 | 12186323 | 9932486 | 11138572 | 9843576 | **1.691690983** | 0 |
| sp\|P30405 | P30405 | PPIF | Peptidyl-prolyl cis-trans isomerase F, mitochondrial OS= | 26502532 | 24392990 | 20393940 | 23242156 | 42338084 | 38325924 | 42610964 | 36602872 | **1.691263171** | 0 |
| sp\|P18669 | P18669 | PGAM1 | Phosphoglycerate mutase 1 OS=Homo sapiens (Human) | 146777408 | 121511352 | 118552584 | 125721552 | 232474288 | 254102528 | 196511088 | 183476400 | **1.690649695** | 0 |
| sp\|P45983 | P45983 | MAPK8 | Mitogen-activated protein kinase 8 OS=Homo sapiens (H | 237278.47 | 225935.48 | 259625.02 | 228850.89 | 390461.34 | 342256.16 | 407049.06 | 468944.44 | **1.690373164** | 0 |
| sp\|Q0212 | Q02127 | DHODH | Dihydroorotate dehydrogenase (quinone), mitochondrial | 798384.38 | 743459.5 | 372900.84 | 1142038.12 | 1118498.25 | 1386566.62 | 1333611.5 | 1327121 | **1.689945816** | 1 |
| sp\|P08727 | P08727 | KRT19 | Keratin, type I cytoskeletal 19 OS=Homo sapiens (Humar | 325097280 | 401262048 | 203990064 | 354812448 | 362527296 | 578171520 | 645219392 | 579694080 | **1.685089162** | 1 |
| sp\|P61088 | P61088 | UBE2N | Ubiquitin-conjugating enzyme E2 N OS=Homo sapiens (H | 39956836 | 35216924 | 35667860 | 42902796 | 64660104 | 61916768 | 69748208 | 62731484 | **1.684981938** | 0 |
| sp\|Q9ULV | Q9ULV4 | CORO1C | Coronin-1C OS=Homo sapiens (Human) GN=CORO1C PE= | 40214100 | 38475912 | 32418056 | 38885088 | 67696824 | 71181360 | 53506764 | 60173152 | **1.683797493** | 0 |
| sp\|Q8NF3 | Q8NF37 | LPCAT1 | Lysophosphatidylcholine acyltransferase 1 OS=Homo sa | 3401674.25 | 1032221.56 | 2975921.25 | 2764920 | 5243104 | 3800751.25 | 3679485.75 | 4397262 | **1.682658028** | 0 |
| sp\|O4323 | O43237 | DYNC1LI2 | Cytoplasmic dynein 1 light intermediate chain 2 OS=Hom | 4972367 | 4947325.5 | 4328949.5 | 5601475.5 | 9936160 | 5555552 | 12023354 | 5850262 | **1.680862997** | 1 |
| sp\|Q9HBH | Q9HBH1 | PDF | Peptide deformylase, mitochondrial OS=Homo sapiens ( | 3340099 | 2207824.5 | 3053410.75 | 2436951.75 | 5105287.5 | 3509539.75 | 5124164.5 | 4806484 | **1.68010466** | 0 |
| sp\|Q9H4A | Q9H4A6 | GOLPH3 | Golgi phosphoprotein 3 OS=Homo sapiens (Human) GN= | 3678122.75 | 4678043 | 2652072.75 | 4683014 | 6239396.5 | 6756198 | 6852486.5 | 6498023 | **1.679031295** | 0 |
| sp\|P20073 | P20073 | ANXA7 | Annexin A7 OS=Homo sapiens (Human) GN=ANXA7 PE=1 | 18839544 | 21251588 | 26098512 | 19765370 | 40698228 | 32992024 | 32370662 | 38213808 | **1.678491053** | 0 |

| | | | | | | | | | | | | | |
|---|---|---|---|---|---|---|---|---|---|---|---|---|---|
| sp\|Q9949 | Q99497 | PARK7 | Protein/nucleic acid deglycase DJ-1 OS=Homo sapiens (H | 143817584 | 160176912 | 144196400 | 116008768 | 264948832 | 204860032 | 241698160 | 235132608 | **1.677845083** | **0** |
| sp\|P61081 | P61081 | UBE2M | NEDD8-conjugating enzyme Ubc12 OS=Homo sapiens (H | 5031151 | 5031667 | 7791269 | 5288553 | 10453177 | 7952323 | 10937402 | 9463092 | **1.676817943** | **0** |
| sp\|P49903 | P49903 | SEPHS1 | Selenide, water dikinase 1 OS=Homo sapiens (Human) G | 4057925.25 | 3194693.5 | 2878398 | 4770755 | 7311419 | 5975195.5 | 6089833 | 5581403 | **1.674824371** | **0** |
| sp\|P19013 | P19013 | KRT4 | Keratin, type II cytoskeletal 4 OS=Homo sapiens (Human | 106729408 | 164128288 | 137151808 | 124620464 | 210334448 | 258707072 | 216082128 | 206476480 | **1.673957872** | **0** |
| sp\|P49590 | P49590 | HARS2 | Probable histidine--tRNA ligase, mitochondrial OS=Homo | 9957348 | 11504029 | 11613240 | 10907678 | 19257858 | 16506219 | 16687480 | 21135510 | **1.673106576** | **0** |
| sp\|P15586 | P15586 | GNS | N-acetylglucosamine-6-sulfatase OS=Homo sapiens (Hur | 6580480 | 5559704 | 8618186 | 7114623 | 14593988 | 12033189 | 10357518 | 9642333 | **1.672838938** | **0** |
| sp\|Q6RW1 | Q6RW13 | AGTRAP | Type-1 angiotensin II receptor-associated protein OS=Hc | 641843.88 | 886242.38 | 551113.5 | 0 | 873050.56 | 1809711 | 0 | 792115.94 | **1.671257167** | **5.5** |
| sp\|Q96C2 | Q96C23 | GALM | Aldose 1-epimerase OS=Homo sapiens (Human) GN=GAI | 34267328 | 36195280 | 35194732 | 33785256 | 54244960 | 54602464 | 61209440 | 62926464 | **1.670818923** | **0** |
| sp\|Q0005 | Q00059 | TFAM | Transcription factor A, mitochondrial OS=Homo sapiens | 5902634 | 10951894 | 11972084 | 11116262 | 18759270 | 13225503 | 20340958 | 14402897 | **1.67060157** | **0** |
| sp\|Q5VTR | Q5VTR2 | RNF20 | E3 ubiquitin-protein ligase BRE1A OS=Homo sapiens (Hu | 0 | 629561.62 | 782160.25 | 540885.44 | 860981.94 | 900753.38 | 754948.38 | 741680.81 | **1.668724937** | **2** |
| sp\|Q9NSK | Q9NSK0 | KLC4 | Kinesin light chain 4 OS=Homo sapiens (Human) GN=KLC | 4787033.5 | 3816935.5 | 2525568 | 3121638.75 | 5721138 | 5937816.5 | 5893581 | 6213996 | **1.667689173** | **0** |
| sp\|Q1575 | Q15750 | TAB1 | TGF-beta-activated kinase 1 and MAP3K7-binding proteir | 0 | 0 | 847187.56 | 909184.81 | 959085.12 | 1038782.62 | 931075.38 | 0 | **1.667609426** | **3** |
| sp\|P07858 | P07858 | CTSB | Cathepsin B OS=Homo sapiens (Human) GN=CTSB PE=1 S | 24436296 | 25327124 | 19249412 | 21764758 | 39872796 | 35848512 | 41258016 | 34142388 | **1.664746905** | **0** |
| sp\|Q96JB3 | Q96JB3 | HIC2 | Hypermethylated in cancer 2 protein OS=Homo sapiens | 8778828 | 6803626 | 6869933 | 7421694 | 14410863 | 10703138 | 12353768 | 12237296 | **1.663819048** | **0** |
| sp\|Q5JS37 | Q5JS37 | NHLRC3 | NHL repeat-containing protein 3 OS=Homo sapiens (Hun | 173134.66 | 240304.97 | 0 | 236160.69 | 315892.59 | 299730.84 | 0 | 463230 | **1.660795718** | **3.5** |
| sp\|Q9Y6Y | Q9Y6Y8 | SEC23IP | SEC23-interacting protein OS=Homo sapiens (Human) GI | 5128213 | 6901129.5 | 6031502 | 6870317.5 | 10216382 | 9921223 | 10734068 | 10522082 | **1.66032193** | **0** |
| sp\|Q9NZL | Q9NZL9 | MAT2B | Methionine adenosyltransferase 2 subunit beta OS=Hom | 8640330 | 7973654 | 8220404 | 8884421 | 14093954 | 13832276 | 14706102 | 13299748 | **1.658779822** | **0** |
| sp\|Q0842 | Q08426 | EHHADH | Peroxisomal bifunctional enzyme OS=Homo sapiens (Hu | 5865616.5 | 1988950.25 | 4417266 | 3350432 | 8262877.5 | 5696177.5 | 6415003.5 | 5514103.5 | **1.657132459** | **2** |
| sp\|Q9HB0 | Q9HB07 | C12orf10 | UPF0160 protein MYG1, mitochondrial OS=Homo sapien | 6431161.5 | 5937900 | 5388218 | 4696761.5 | 9797134 | 8551146 | 10319578 | 8540424 | **1.657086223** | **0** |
| sp\|P02649 | P02649 | APOE | Apolipoprotein E OS=Homo sapiens (Human) GN=APOE I | 7061567 | 9644160 | 11496413 | 10795677 | 17814012 | 11832212 | 18445450 | 16505023 | **1.656418281** | **0** |
| sp\|P06396 | P06396 | GSN | Gelsolin OS=Homo sapiens (Human) GN=GSN PE=1 SV=1 | 2967772.5 | 2665819 | 1125135 | 3693900.25 | 4853488 | 2568141 | 5294849 | 4587298.5 | **1.655447661** | **3** |
| sp\|O15084 | O15084 | ANKRD28 | Serine/threonine-protein phosphatase 6 regulatory anky | 142852.2 | 723753 | 206020.8 | 565492.81 | 924499 | 784753.25 | 257793.91 | 744178.12 | **1.655084029** | **2** |
| sp\|Q9UNZ | Q9UNZ2 | NSFL1C | NSFL1 cofactor p47 OS=Homo sapiens (Human) GN=NSFI | 21738018 | 21216868 | 22424308 | 15340044 | 35495064 | 32682636 | 35000484 | 30352228 | **1.654257589** | **0** |
| sp\|O6042 | O60427 | FADS1 | Fatty acid desaturase 1 OS=Homo sapiens (Human) GN=I | 125099.83 | 0 | 142424.84 | 170892.5 | 122152.76 | 177799.55 | 191295.55 | 232758.27 | **1.651409159** | **3** |
| sp\|P20810 | P20810 | CAST | Calpastatin OS=Homo sapiens (Human) GN=CAST PE=1 S | 1013498.62 | 2045772.88 | 1059686.38 | 1304470.75 | 3146175 | 848865.69 | 3169911.5 | 1786785.5 | **1.650567989** | **5** |
| sp\|P22307 | P22307 | SCP2 | Non-specific lipid-transfer protein OS=Homo sapiens (Hu | 6968169 | 6194798 | 4750550.5 | 7227321.5 | 10804343 | 10074517 | 9418685 | 11168980 | **1.64936918** | **0** |
| sp\|Q8IZ21 | Q8IZ21 | PHACTR4 | Phosphatase and actin regulator 4 OS=Homo sapiens (Hu | 0 | 338584.62 | 0 | 0 | 0 | 0 | 558074.31 | 0 | **1.648256527** | **7.5** |
| sp\|Q1507 | Q15075 | EEA1 | Early endosome antigen 1 OS=Homo sapiens (Human) G | 22900842 | 25504836 | 24977184 | 20405844 | 38440708 | 35715592 | 39983016 | 40391292 | **1.647646232** | **0** |
| sp\|P05165 | P05165 | PCCA | Propionyl-CoA carboxylase alpha chain, mitochondrial O | 1834383.75 | 2549793.25 | 2150287 | 3370297 | 4257608.5 | 3175961 | 3767277.25 | 5106839 | **1.646449192** | **1** |
| sp\|P61163 | P61163 | ACTR1A | Alpha-centractin OS=Homo sapiens (Human) GN=ACTR1 | 14516177 | 14088640 | 17168510 | 20745660 | 26692876 | 28928016 | 32937240 | 20892304 | **1.645401425** | **0** |
| sp\|P48735 | P48735 | IDH2 | Isocitrate dehydrogenase [NADP], mitochondrial OS=Ho | 26063570 | 31412820 | 30867844 | 31283812 | 46655684 | 47052000 | 52140860 | 50791868 | **1.643765142** | **0** |
| sp\|Q9Y69 | Q9Y697 | NFS1 | Cysteine desulfurase, mitochondrial OS=Homo sapiens ( | 4916234.5 | 2483526 | 2629241 | 4684270.5 | 6405550.5 | 6549089.5 | 5757757.5 | 5443587 | **1.641781957** | **0** |
| sp\|Q1357 | Q13576 | IQGAP2 | Ras GTPase-activating-like protein IQGAP2 OS=Homo sa | 42881188 | 47734680 | 44721804 | 41465332 | 74943208 | 69611624 | 66218308 | 79424592 | **1.641361999** | **0** |
| sp\|Q9BRT | Q9BRT6 | LLPH | Protein LLP homolog OS=Homo sapiens (Human) GN=LLI | 0 | 1068531.62 | 1213807.75 | 1172742 | 1710847.5 | 1383986.12 | 1208987.38 | 1364547.88 | **1.640589113** | **1** |
| sp\|O9588 | O95881 | TXNDC12 | Thioredoxin domain-containing protein 12 OS=Homo sa | 8206959.5 | 5485684.5 | 3331778.75 | 5823360.5 | 9737434 | 9805112 | 9830288 | 8069890 | **1.638790231** | **1** |
| sp\|Q96HQ | Q96HQ2 | CDKN2AIPI | CDKN2AIP N-terminal-like protein OS=Homo sapiens (Hu | 637005.25 | 561940 | 433031.56 | 561283.5 | 814150.69 | 865924.62 | 1049649 | 860782.44 | **1.63706366** | **0** |
| sp\|P41214 | P41214 | EIF2D | Eukaryotic translation initiation factor 2D OS=Homo sapi | 443577.91 | 986353.12 | 364495.25 | 296908.41 | 820237.81 | 843647.31 | 941705.56 | 818012.25 | **1.637042099** | **4** |
| sp\|O9561 | O95619 | YEATS4 | YEATS domain-containing protein 4 OS=Homo sapiens (H | 617105.94 | 592823.75 | 0 | 636714.5 | 869277.5 | 876961.75 | 664326.56 | 612355.69 | **1.636981026** | **2** |
| sp\|Q8NBU | Q8NBU5 | ATAD1 | ATPase family AAA domain-containing protein 1 OS=Hor | 0 | 431945.34 | 389815.38 | 230848.22 | 736023.62 | 545760 | 0 | 437134.19 | **1.633007041** | **3.5** |
| sp\|P27824 | P27824 | CANX | Calnexin OS=Homo sapiens (Human) GN=CANX PE=1 SV= | 169838864 | 179671904 | 176386464 | 173911904 | 261516960 | 285880512 | 313246208 | 281814752 | **1.632528604** | **0** |
| sp\|Q9Y3B | Q9Y3B4 | SF3B6 | Splicing factor 3B subunit 6 OS=Homo sapiens (Human) ( | 2697444.5 | 1753365.25 | 1386084.12 | 4036560 | 3864578 | 1906957.38 | 4288966 | 6013646 | **1.628016659** | **3** |
| sp\|Q0795 | Q07954 | LRP1 | Prolow-density lipoprotein receptor-related protein 1 O! | 616474.94 | 935111.88 | 858213.12 | 915357.62 | 1080098.38 | 1768211.12 | 1484896.25 | 1077767.12 | **1.62728315** | **0** |
| sp\|A0A0B | A0A0B4J2D5 | GATD3B | Glutamine amidotransferase-like class 1 domain-contain | 4628316 | 3623053 | 3615543.25 | 4278854.5 | 6324891 | 6561556 | 7423692.5 | 5948716 | **1.626361628** | **0** |
| sp\|Q96GK | Q96GK7 | FAHD2A | Fumarylacetoacetate hydrolase domain-containing prote | 1143400.38 | 1154926.38 | 1271944.88 | 1201156.75 | 1815728.38 | 1826828 | 2191610.5 | 1924661.25 | **1.626101766** | **0** |
| sp\|Q9BPW | Q9BPW8 | NIPSNAP1 | Protein NipSnap homolog 1 OS=Homo sapiens (Human) | 10123038 | 9422284 | 9590264 | 10551414 | 18369530 | 13602377 | 18201610 | 14354236 | **1.625916623** | **0** |

| | | | | | | | | | | | | | |
|---|---|---|---|---|---|---|---|---|---|---|---|---|---|
| sp\|Q969S3 | Q969S3 | ZNF622 | Zinc finger protein 622 OS=Homo sapiens (Human) GN=Z | 581727.75 | 704961.62 | 516165.69 | 1217102.5 | 1289504 | 1177158.75 | 1312563.75 | 1127375.12 | 1.624725355 | 2 |
| sp\|Q93034 | Q93034 | CUL5 | Cullin-5 OS=Homo sapiens (Human) GN=CUL5 PE=1 SV=4 | 1191113.38 | 587196.5 | 1345350.25 | 1238443 | 1925277.5 | 1586875.75 | 1931536.25 | 1642764.75 | 1.624549911 | 0 |
| sp\|Q9C0D9 | Q9C0D9 | SELENOI | Ethanolaminephosphotransferase 1 OS=Homo sapiens (I | 2286066.25 | 1757000.25 | 1233735 | 2317191.75 | 3239029.75 | 3976583.75 | 2036977.38 | 3080669 | 1.624080964 | 2 |
| sp\|P51687 | P51687 | SUOX | Sulfite oxidase, mitochondrial OS=Homo sapiens (Human | 0 | 281093.62 | 0 | 0 | 219066.8 | 237072.94 | 0 | 0 | 1.622732455 | 7 |
| sp\|Q99735 | Q99735 | MGST2 | Microsomal glutathione S-transferase 2 OS=Homo sapie | 14282628 | 15022179 | 15255148 | 13522133 | 23610904 | 25769322 | 22923082 | 21910388 | 1.622078325 | 0 |
| sp\|P52815 | P52815 | MRPL12 | 39S ribosomal protein L12, mitochondrial OS=Homo sapi | 1680552.75 | 633894.12 | 1037654.12 | 1952582 | 2222116 | 2423301 | 2878927 | 1074080.25 | 1.620911988 | 2 |
| sp\|Q92845 | Q92845 | KIFAP3 | Kinesin-associated protein 3 OS=Homo sapiens (Human) | 220695.67 | 293499.91 | 252351.22 | 92414.14 | 268796.06 | 355903.16 | 377034.75 | 389516.03 | 1.619689482 | 1 |
| sp\|Q6P5W | Q6P5W5 | SLC39A4 | Zinc transporter ZIP4 OS=Homo sapiens (Human) GN=SL | 0 | 275415.12 | 418380.88 | 352972.97 | 462952.72 | 484908.25 | 359010.41 | 387788.12 | 1.618943194 | 2 |
| sp\|Q8WY2 | Q8WY22 | BRI3BP | BRI3-binding protein OS=Homo sapiens (Human) GN=BR | 1312605.5 | 1879022.5 | 1251385.5 | 2252638 | 3003311.75 | 1702509 | 2959916 | 3167807.25 | 1.617996994 | 2 |
| sp\|Q4KMC | Q4KMQ2 | ANO6 | Anoctamin-6 OS=Homo sapiens (Human) GN=ANO6 PE=1 | 1646739.5 | 2186242.75 | 1385866.62 | 1890209.25 | 3055475.25 | 3101689.5 | 2107265.75 | 3222686.25 | 1.61584229 | 1 |
| sp\|Q92522 | Q92522 | H1FX | Histone H1x OS=Homo sapiens (Human) GN=H1FX PE=1 | 13046840 | 12446961 | 9115971 | 11562919 | 20975890 | 19816238 | 18334446 | 15453042 | 1.615232173 | 0 |
| sp\|Q9BXS5 | Q9BXS5 | AP1M1 | AP-1 complex subunit mu-1 OS=Homo sapiens (Human) | 1315654.38 | 290143.06 | 342831.28 | 1600737.25 | 1596678.38 | 1761067.75 | 357433.03 | 2011632 | 1.613474409 | 3 |
| sp\|P42025 | P42025 | ACTR1B | Beta-centractin OS=Homo sapiens (Human) GN=ACTR1B | 10860785 | 10915331 | 14035560 | 17407766 | 21356880 | 22577112 | 27044568 | 14836065 | 1.612467583 | 1 |
| sp\|Q5BJH7 | Q5BJH7 | YIF1B | Protein YIF1B OS=Homo sapiens (Human) GN=YIF1B PE= | 1319873.38 | 0 | 1270224.88 | 1712905 | 1824480.5 | 2090517.88 | 1410580.25 | 1607293.12 | 1.611170462 | 2 |
| sp\|Q16204 | Q16204 | CCDC6 | Coiled-coil domain-containing protein 6 OS=Homo sapie | 574999.75 | 1021091.94 | 932562.75 | 1070057.25 | 1621519.25 | 1387862.88 | 1523216 | 1258786.12 | 1.609293755 | 0 |
| sp\|Q08495 | Q08495 | DMTN | Dematin OS=Homo sapiens (Human) GN=DMTN PE=1 SV | 732031.88 | 753501.31 | 883668.62 | 751500.81 | 979647.25 | 1196858.25 | 1599054.12 | 1242982 | 1.608144777 | 0 |
| sp\|Q9UKK9 | Q9UKK9 | NUDT5 | ADP-sugar pyrophosphatase OS=Homo sapiens (Human) | 21243570 | 27783858 | 22704744 | 26774562 | 34585168 | 43187172 | 33873376 | 46741176 | 1.607878828 | 0 |
| sp\|Q86SX6 | Q86SX6 | GLRX5 | Glutaredoxin-related protein 5, mitochondrial OS=Homo | 2378155 | 2318365 | 1502352.5 | 2484122 | 3460289.75 | 3285843.5 | 3706160 | 3507656.25 | 1.607734463 | 0 |
| sp\|Q9BW92 | Q9BW92 | TARS2 | Threonine--tRNA ligase, mitochondrial OS=Homo sapien | 1646935.62 | 3432595.25 | 2259550 | 2336702.25 | 4078082.75 | 3955673.5 | 4146373 | 3369488.5 | 1.607065553 | 1 |
| sp\|P15559 | P15559 | NQO1 | NAD(P)H dehydrogenase [quinone] 1 OS=Homo sapiens | 156674160 | 150138944 | 146258768 | 151859200 | 260369152 | 236355392 | 224061728 | 251038752 | 1.606505384 | 0 |
| sp\|Q13952 | Q13952 | NFYC | Nuclear transcription factor Y subunit gamma OS=Homo | 0 | 619499.94 | 0 | 528275.62 | 457523.56 | 0 | 683345.81 | 701579.88 | 1.605234781 | 5 |
| sp\|A6NCS6 | A6NCS6 | C2orf72 | Uncharacterized protein C2orf72 OS=Homo sapiens (Hur | 5074218 | 8174268.5 | 2628588.5 | 5931638 | 9915942 | 6672361 | 10401784 | 8015108.5 | 1.605101388 | 2 |
| sp\|Q9H6T3 | Q9H6T3 | RPAP3 | RNA polymerase II-associated protein 3 OS=Homo sapie | 1180236.5 | 454035.31 | 568999.62 | 974870.62 | 1368865.5 | 1368834.25 | 1152663.75 | 1210569.88 | 1.605004842 | 1 |
| sp\|Q9NVI7 | Q9NVI7 | ATAD3A | ATPase family AAA domain-containing protein 3A OS=Hc | 33717440 | 37293104 | 38898076 | 34317588 | 62536932 | 55725120 | 51096776 | 62117052 | 1.60495019 | 0 |
| sp\|P55145 | P55145 | MANF | Mesencephalic astrocyte-derived neurotrophic factor OS | 2638156 | 3405828 | 1678008.62 | 4606368 | 4538201 | 4947000 | 5656521 | 4618085.5 | 1.602792789 | 1 |
| sp\|P20823 | P20823 | HNF1A | Hepatocyte nuclear factor 1-alpha OS=Homo sapiens (Hu | 518862.28 | 1341256.75 | 404310.66 | 1419370.25 | 1674447 | 611761.12 | 2005401.25 | 1611988.25 | 1.602583668 | 2 |
| sp\|Q15369 | Q15369 | ELOC | Elongin-C OS=Homo sapiens (Human) GN=ELOC PE=1 SV | 3787568.5 | 2641646 | 1363164 | 3527002.5 | 5116882.5 | 3933704.5 | 6115754.5 | 2952395.25 | 1.600682648 | 2 |
| sp\|P12081 | P12081 | HARS | Histidine--tRNA ligase, cytoplasmic OS=Homo sapiens (H | 28860816 | 32208168 | 30097718 | 29043844 | 47857052 | 46415608 | 46899132 | 51217372 | 1.60043499 | 0 |
| sp\|Q9NVH | Q9NVH1 | DNAJC11 | DnaJ homolog subfamily C member 11 OS=Homo sapien | 4121764.5 | 3662612.5 | 3413533.5 | 3631303.5 | 5508738.5 | 6246061 | 6162159 | 5815523.5 | 1.600387047 | 0 |
| sp\|O95571 | O95571 | ETHE1 | Persulfide dioxygenase ETHE1, mitochondrial OS=Homo | 3413064.25 | 2655308 | 2630449.5 | 2136990.5 | 4496884.5 | 2862084.75 | 4789577 | 5190908 | 1.600198845 | 1 |
| sp\|Q13561 | Q13561 | DCTN2 | Dynactin subunit 2 OS=Homo sapiens (Human) GN=DCTI | 12703440 | 11910134 | 12151341 | 11516202 | 20632040 | 19181316 | 19879852 | 17548964 | 1.599842274 | 0 |
| sp\|P39210 | P39210 | MPV17 | Protein Mpv17 OS=Homo sapiens (Human) GN=MPV17 F | 426413.12 | 456602.94 | 269351.19 | 521253.81 | 715007.69 | 780160 | 552876.25 | 626560.94 | 1.598094661 | 0 |
| sp\|Q9Y2B0 | Q9Y2B0 | CNPY2 | Protein canopy homolog 2 OS=Homo sapiens (Human) G | 13555077 | 12868014 | 13975245 | 13340001 | 20843822 | 19126284 | 24248716 | 21646288 | 1.597837127 | 0 |
| sp\|Q14677 | Q14677 | CLINT1 | Clathrin interactor 1 OS=Homo sapiens (Human) GN=CLI | 5902165.5 | 7393568.5 | 6053579 | 8046435 | 11138897 | 9516429 | 12102753 | 11005327 | 1.597452495 | 0 |
| sp\|P26885 | P26885 | FKBP2 | Peptidyl-prolyl cis-trans isomerase FKBP2 OS=Homo sapi | 2313862 | 2186570 | 1822979.38 | 2786531 | 3670823 | 3898419.5 | 4058755.5 | 2910851.25 | 1.595932075 | 0 |
| sp\|Q9UBV | Q9UBV8 | PEF1 | Peflin OS=Homo sapiens (Human) GN=PEF1 PE=1 SV=1 | 1038453.75 | 381965.53 | 1414743.38 | 1364568.25 | 1638608 | 1750339.25 | 1425474.75 | 1882149 | 1.594523826 | 0 |
| sp\|Q8N4Z0 | Q8N4Z0 | RAB42 | Ras-related protein Rab-42 OS=Homo sapiens (Human) ( | 535748.38 | 469732.47 | 518992.84 | 585633.81 | 1513755.12 | 1045000.88 | 294284.47 | 509688.31 | 1.59362913 | 7 |
| sp\|Q9Y4C8 | Q9Y4C8 | RBM19 | Probable RNA-binding protein 19 OS=Homo sapiens (Hu | 1188446.25 | 1969286.25 | 1104735.5 | 990794.75 | 2024143.25 | 2134410 | 2180766.25 | 2031070.5 | 1.593369759 | 0 |
| sp\|O43592 | O43592 | XPOT | Exportin-T OS=Homo sapiens (Human) GN=XPOT PE=1 S\ | 7686131 | 8681679 | 7523872.5 | 7791775 | 10825398 | 12600198 | 12617261 | 14410604 | 1.592422828 | 0 |
| sp\|Q96KP1 | Q96KP1 | EXOC2 | Exocyst complex component 2 OS=Homo sapiens (Huma | 966061.12 | 702285.5 | 1221854.25 | 1211571.25 | 1115152.25 | 1679792.5 | 1891397.25 | 1844213.62 | 1.592130286 | 2 |
| sp\|Q1KMD | Q1KMD3 | HNRNPUL2 | Heterogeneous nuclear ribonucleoprotein U-like protein | 12381615 | 13458897 | 13968953 | 10215518 | 18786276 | 20678334 | 17844248 | 22304576 | 1.591473484 | 0 |
| sp\|Q8IUH3 | Q8IUH3 | RBM45 | RNA-binding protein 45 OS=Homo sapiens (Human) GN= | 0 | 159564.36 | 679489.12 | 584748.75 | 682454.62 | 427819.69 | 412245.47 | 742954.44 | 1.591143891 | 4 |
| sp\|P42858 | P42858 | HTT | Huntingtin OS=Homo sapiens (Human) GN=HTT PE=1 SV | 0 | 177712.03 | 248459.34 | 220164.36 | 202457.48 | 314112.69 | 220777.44 | 289982.66 | 1.589468479 | 3 |
| sp\|P49756 | P49756 | RBM25 | RNA-binding protein 25 OS=Homo sapiens (Human) GN= | 3386349.75 | 3327775.5 | 611495 | 5227098 | 7512995 | 6159435 | 3184563.5 | 3082279.25 | 1.588442627 | 6 |
| sp\|P62841 | P62841 | RPS15 | 40S ribosomal protein S15 OS=Homo sapiens (Human) G | 1930261 | 320978.84 | 1045190.88 | 3154019.5 | 3718455.5 | 2092636.38 | 2528907.5 | 1898149.25 | 1.587199076 | 4 |

| | | | | | | | | | | | | | |
|---|---|---|---|---|---|---|---|---|---|---|---|---|---|
| sp\|Q9259 | Q92597 | NDRG1 | Protein NDRG1 OS=Homo sapiens (Human) GN=NDRG1 | 7955292 | 6938828 | 6433460 | 7479714 | 12330016 | 18335540 | 6863965 | 8187468 | **1.586993523** | **3** |
| sp\|O1532 | O15321 | TM9SF1 | Transmembrane 9 superfamily member 1 OS=Homo sap | 2550272.25 | 3457081 | 3015714 | 2717787.75 | 1013902.94 | 1198239.5 | 15095260 | 1304852.12 | **1.585255466** | **4** |
| sp\|Q7KZN | Q7KZN9 | COX15 | Cytochrome c oxidase assembly protein COX15 homolog | 148576.72 | 187487.09 | 90804.41 | 236895.08 | 216147.84 | 360202.5 | 212947.7 | 262758 | **1.584986757** | **2** |
| sp\|P63218 | P63218 | GNG5 | Guanine nucleotide-binding protein G(I)/G(S)/G(O) subur | 0 | 1407466.5 | 0 | 1088299.38 | 1068436 | 834914.06 | 841551.69 | 1204337.5 | **1.582375687** | **7** |
| sp\|P53992 | P53992 | SEC24C | Protein transport protein Sec24C OS=Homo sapiens (Hur | 24670118 | 26797740 | 24535984 | 23289446 | 39229368 | 35854876 | 42193476 | 39807500 | **1.582032614** | **0** |
| sp\|Q6GM | Q6GMV3 | PTRHD1 | Putative peptidyl-tRNA hydrolase PTRHD1 OS=Homo sap | 0 | 654240 | 0 | 698607.19 | 0 | 0 | 1178603.12 | 959728.81 | **1.580616012** | **6** |
| sp\|Q9H9E | Q9H9E3 | COG4 | Conserved oligomeric Golgi complex subunit 4 OS=Hom | 0 | 449656.72 | 681445 | 467063.47 | 676347.56 | 521510.41 | 701754.62 | 626315.69 | **1.580517643** | **3** |
| sp\|Q9BT1 | Q9BT17 | MTG1 | Mitochondrial ribosome-associated GTPase 1 OS=Homo | 596825.56 | 589650.5 | 499128.66 | 860405 | 979375.88 | 1395103.25 | 1271987.62 | 374975.09 | **1.579507654** | **4** |
| sp\|O0026 | O00264 | PGRMC1 | Membrane-associated progesterone receptor componen | 7304515.5 | 7741584 | 7475446 | 9513032 | 16525238 | 11573870 | 11245302 | 11214166 | **1.578250127** | **0** |
| sp\|Q9Y5K | Q9Y5K8 | ATP6V1D | V-type proton ATPase subunit D OS=Homo sapiens (Hum | 601411.75 | 0 | 699435.19 | 609355.38 | 682371.06 | 745784.19 | 811486.62 | 775045.44 | **1.578203146** | **1** |
| sp\|Q1526 | Q15262 | PTPRK | Receptor-type tyrosine-protein phosphatase kappa OS= | 668772.25 | 1399044.5 | 0 | 972284.75 | 1839256.75 | 765545.5 | 1543194.88 | 647424.19 | **1.577388558** | **5** |
| sp\|Q1290 | Q12907 | LMAN2 | Vesicular integral-membrane protein VIP36 OS=Homo sa | 13704397 | 12905517 | 11625585 | 12983322 | 22873800 | 21451374 | 18928960 | 17510742 | **1.576859335** | **0** |
| sp\|Q9NR1 | Q9NR12 | PDLIM7 | PDZ and LIM domain protein 7 OS=Homo sapiens (Huma | 1059197.75 | 835049.94 | 1009408.56 | 832637.62 | 1749003.62 | 1423575.5 | 1403179.75 | 1314635.75 | **1.57653408** | **0** |
| sp\|P49419 | P49419 | ALDH7A1 | Alpha-aminoadipic semialdehyde dehydrogenase OS=H | 38576420 | 46119968 | 45785100 | 39920124 | 72054304 | 66666288 | 66179460 | 63724044 | **1.576417575** | **0** |
| sp\|P49750 | P49750 | YLPM1 | YLP motif-containing protein 1 OS=Homo sapiens (Huma | 3499020.5 | 3149030.5 | 2415091.5 | 201782.78 | 3663000.75 | 3071855 | 4401891 | 3464945.25 | **1.576018323** | **3** |
| sp\|P05543 | P05543 | SERPINA7 | Thyroxine-binding globulin OS=Homo sapiens (Human) | 0 | 463902.84 | 692890.31 | 498835.12 | 662532.88 | 572064.5 | 766391.25 | 603540.94 | **1.573136686** | **3** |
| sp\|P31040 | P31040 | SDHA | Succinate dehydrogenase [ubiquinone] flavoprotein sub | 28400708 | 27767132 | 30713842 | 26379972 | 42187300 | 49411132 | 44242260 | 41854624 | **1.568892116** | **0** |
| sp\|P46013 | P46013 | MKI67 | Proliferation marker protein Ki-67 OS=Homo sapiens (Hu | 3960125.5 | 2453253 | 4579886.5 | 2518809 | 3864975.25 | 5540613.5 | 5496300.5 | 6295885.5 | **1.568802447** | **2** |
| sp\|Q9Y2Q | Q9Y2Q5 | LAMTOR2 | Ragulator complex protein LAMTOR2 OS=Homo sapiens | 663711.38 | 773711.19 | 1140411.88 | 836808.62 | 1782859.5 | 958405.62 | 1166112.62 | 1438480.25 | **1.565568606** | **1** |
| sp\|Q9UGK | Q9UGK3 | STAP2 | Signal-transducing adaptor protein 2 OS=Homo sapiens | 0 | 809285.5 | 1158725.88 | 636850 | 1200048.88 | 662992 | 1389600.5 | 820625.88 | **1.563717475** | **3** |
| sp\|Q1340 | Q13405 | MRPL49 | 39S ribosomal protein L49, mitochondrial OS=Homo sapi | 3061628.5 | 2822970 | 2658850 | 3503834.5 | 5314532 | 3722279.5 | 5415968 | 4367747 | **1.562221664** | **0** |
| sp\|Q8NE6 | Q8NE62 | CHDH | Choline dehydrogenase, mitochondrial OS=Homo sapier | 1368228.12 | 3724633 | 4769243 | 4502348 | 5023105.5 | 5605818 | 5585671 | 6197070.5 | **1.560217181** | **0** |
| sp\|Q1342 | Q13428 | TCOF1 | Treacle protein OS=Homo sapiens (Human) GN=TCOF1 P | 45812288 | 40445352 | 49692392 | 37416288 | 71223816 | 58194924 | 80011832 | 61042460 | **1.560124435** | **0** |
| sp\|Q6PGP | Q6PGP7 | TTC37 | Tetratricopeptide repeat protein 37 OS=Homo sapiens ( | 1134772.75 | 653635.19 | 990291.38 | 1277188.5 | 1446068.75 | 2089905.5 | 1412247.5 | 1377832.62 | **1.559721238** | **0** |
| sp\|P43897 | P43897 | TSFM | Elongation factor Ts, mitochondrial OS=Homo sapiens (H | 8672376 | 9150249 | 8189322.5 | 8022305.5 | 12902287 | 12713357 | 14371662 | 13085187 | **1.559384688** | **0** |
| sp\|P07148 | P07148 | FABP1 | Fatty acid-binding protein, liver OS=Homo sapiens (Hum | 139976512 | 136660896 | 140989376 | 157831504 | 218338608 | 191512928 | 246105936 | 241383168 | **1.559349581** | **0** |
| sp\|P61970 | P61970 | NUTF2 | Nuclear transport factor 2 OS=Homo sapiens (Human) G | 18787226 | 19016212 | 16395529 | 16449586 | 26901178 | 29781620 | 26832806 | 26649426 | **1.559338802** | **0** |
| sp\|P45954 | P45954 | ACADSB | Short/branched chain specific acyl-CoA dehydrogenase, | 48363668 | 48880576 | 47851236 | 47754628 | 76950688 | 66280568 | 73365768 | 83122792 | **1.554159441** | **0** |
| sp\|Q96HE | Q96HE7 | ERO1A | ERO1-like protein alpha OS=Homo sapiens (Human) GN= | 27752200 | 29225362 | 29386148 | 28074608 | 42875876 | 52787572 | 43062160 | 39048880 | **1.553452472** | **0** |
| sp\|Q86YH | Q86YH6 | PDSS2 | Decaprenyl-diphosphate synthase subunit 2 OS=Homo s | 888511.5 | 832858.75 | 688163.19 | 350043.69 | 909340 | 887408.75 | 1077326.88 | 1412492.62 | **1.553342432** | **1** |
| sp\|Q70J99 | Q70J99 | UNC13D | Protein unc-13 homolog D OS=Homo sapiens (Human) G | 255671.84 | 250495.22 | 233634.45 | 247203.98 | 389836.5 | 356624.12 | 442457.06 | 340979.31 | **1.550038987** | **0** |
| sp\|P23588 | P23588 | EIF4B | Eukaryotic translation initiation factor 4B OS=Homo sapi | 12241245 | 10103193 | 15395134 | 14200620 | 22155944 | 20443582 | 21827504 | 16029841 | **1.549029141** | **0** |
| sp\|Q9NX6 | Q9NX63 | CHCHD3 | MICOS complex subunit MIC19 OS=Homo sapiens (Huma | 5254918.5 | 8374708 | 9485823 | 8770640 | 12304661 | 10542854 | 13542460 | 12982585 | **1.54840436** | **0** |
| sp\|P62942 | P62942 | FKBP1A | Peptidyl-prolyl cis-trans isomerase FKBP1A OS=Homo sa | 12846996 | 11435190 | 10735686 | 10186901 | 16807836 | 18957636 | 20972852 | 13191586 | **1.546958548** | **0** |
| sp\|P20674 | P20674 | COX5A | Cytochrome c oxidase subunit 5A, mitochondrial OS=Ho | 10199939 | 8383573 | 8177079 | 9703730 | 11523098 | 12626499 | 15276068 | 16903760 | **1.544781953** | **0** |
| sp\|P54886 | P54886 | ALDH18A1 | Delta-1-pyrroline-5-carboxylate synthase OS=Homo sapi | 124598072 | 139185792 | 132702688 | 133550584 | 200820208 | 181621904 | 230242064 | 205385840 | **1.543420188** | **0** |
| sp\|Q1677 | Q16775 | HAGH | Hydroxyacylglutathione hydrolase, mitochondrial OS=H | 2279760.25 | 2333924.75 | 2557962.75 | 2813987 | 3482159.75 | 3064916.25 | 3774661 | 5087402.5 | **1.543130696** | **0** |
| sp\|Q8IYS2 | Q8IYS2 | KIAA2013 | Uncharacterized protein KIAA2013 OS=Homo sapiens (H | 2443064.5 | 1941935.75 | 1688493.62 | 2150712.5 | 2952551 | 3656096.25 | 3174174.75 | 2902728.5 | **1.542465003** | **0** |
| sp\|Q1416 | Q14165 | MLEC | Malectin OS=Homo sapiens (Human) GN=MLEC PE=1 SV | 9387322 | 10481678 | 6859543 | 7084012 | 10829242 | 14949457 | 10859117 | 15470414 | **1.541091172** | **0** |
| sp\|Q9P25 | Q9P258 | RCC2 | Protein RCC2 OS=Homo sapiens (Human) GN=RCC2 PE=1 | 17042268 | 21326172 | 17278116 | 19044980 | 27451552 | 27998052 | 27514486 | 32115736 | **1.540734495** | **0** |
| sp\|O7543 | O75431 | MTX2 | Metaxin-2 OS=Homo sapiens (Human) GN=MTX2 PE=1 S | 1701445.5 | 2176568.75 | 1752994.88 | 2149476.25 | 2527544.5 | 2667438.75 | 2662903.5 | 4127984.75 | **1.540504341** | **0** |
| sp\|Q5T44 | Q5T440 | IBA57 | Putative transferase CAF17, mitochondrial OS=Homo sa | 3716420 | 5594183 | 5352353.5 | 3751815.25 | 6603880 | 7046651 | 6369344.5 | 8329815 | **1.539508113** | **0** |
| sp\|O7594 | O75947 | ATP5PD | ATP synthase subunit d, mitochondrial OS=Homo sapien | 13083924 | 9921452 | 8683382 | 10706406 | 17970364 | 13663197 | 19589788 | 14022996 | **1.539004425** | **0** |
| sp\|Q1583 | Q15836 | VAMP3 | Vesicle-associated membrane protein 3 OS=Homo sapie | 8184892.5 | 10299139 | 11280118 | 9684828 | 17489996 | 13680073 | 15708976 | 13786254 | **1.537816766** | **0** |
| sp\|O4392 | O43920 | NDUFS5 | NADH dehydrogenase [ubiquinone] iron-sulfur protein 5 | 2720048.75 | 2729766 | 2197893 | 2930275.75 | 4394834.5 | 3533089.25 | 4248820.5 | 4084477 | **1.537270431** | **0** |

| | | | | | | | | | | | | | |
|---|---|---|---|---|---|---|---|---|---|---|---|---|---|
| sp\|P35908 | P35908 | KRT2 | Keratin, type II cytoskeletal 2 epidermal OS=Homo sapie | 141009392 | 266653136 | 166482656 | 145958080 | 259684592 | 310776576 | 289500128 | 247024656 | **1.537260012** | 2 |
| sp\|Q9UGI | Q9UGI8 | TES | Testin OS=Homo sapiens (Human) GN=TES PE=1 SV=1 | 1375971.38 | 2574480 | 3769949.5 | 1810523.88 | 4273382 | 3286700.25 | 3681283.5 | 3406961.5 | **1.536926124** | 3 |
| sp\|P23297 | P23297 | S100A1 | Protein S100-A1 OS=Homo sapiens (Human) GN=S100A1 | 15995628 | 8065817.5 | 5926969.5 | 6034289.5 | 8576419 | 22073280 | 12028258 | 12652009 | **1.535974791** | 3 |
| sp\|Q9UBL | Q9UBL3 | ASH2L | Set1/Ash2 histone methyltransferase complex subunit AS | 464607.34 | 1131428.75 | 407594.31 | 538566.94 | 1407254.88 | 1530434.5 | 497912.94 | 466917.12 | **1.535096972** | 4 |
| sp\|P47985 | P47985 | UQCRFS1 | Cytochrome b-c1 complex subunit Rieske, mitochondrial | 20774232 | 19186314 | 76151240 | 20288578 | 88187640 | 71585664 | 26251096 | 23190326 | **1.533828209** | 3 |
| sp\|Q0825 | Q08257 | CRYZ | Quinone oxidoreductase OS=Homo sapiens (Human) GN | 52004896 | 47382992 | 50699616 | 52501020 | 81073920 | 77772344 | 81413440 | 70425952 | **1.53357974** | 0 |
| sp\|O9556 | O95563 | MPC2 | Mitochondrial pyruvate carrier 2 OS=Homo sapiens (Hur | 2626601 | 2549434 | 2259364.5 | 3204301.5 | 3722847.25 | 3642508 | 3254587.25 | 5693187 | **1.533231949** | 0 |
| sp\|Q96CN | Q96CN7 | ISOC1 | Isochorismatase domain-containing protein 1 OS=Homo | 17760816 | 23286162 | 23241770 | 23529784 | 34918344 | 31510922 | 33567268 | 34539600 | **1.531978854** | 0 |
| sp\|O7531 | O75319 | DUSP11 | RNA/RNP complex-1-interacting phosphatase OS=Homo | 0 | 0 | 394677.62 | 358958.19 | 391805.88 | 369576.28 | 391071.25 | 0 | **1.529191414** | 6 |
| sp\|Q9UNV | Q9UNW1 | MINPP1 | Multiple inositol polyphosphate phosphatase 1 OS=Hom | 6058548 | 6089594.5 | 6418663 | 6355055 | 10420537 | 7438001.5 | 9899292 | 10322426 | **1.527986103** | 0 |
| sp\|O7514 | O75146 | HIP1R | Huntingtin-interacting protein 1-related protein OS=Hon | 1675315.12 | 2243555.5 | 2025306.75 | 1346137 | 2544326 | 2637322.5 | 3263081.5 | 2692736.75 | **1.527707337** | 0 |
| sp\|P14735 | P14735 | IDE | Insulin-degrading enzyme OS=Homo sapiens (Human) GI | 13641655 | 10828638 | 13065968 | 11636019 | 20386144 | 19133806 | 17519808 | 18068940 | **1.527460146** | 0 |
| sp\|O4339 | O43399 | TPD52L2 | Tumor protein D54 OS=Homo sapiens (Human) GN=TPD! | 31214272 | 32476916 | 31904006 | 33533738 | 46575064 | 45681544 | 55646312 | 49235632 | **1.526679954** | 0 |
| sp\|Q9Y6B | Q9Y6B6 | SAR1B | GTP-binding protein SAR1b OS=Homo sapiens (Human) ( | 22553078 | 23023270 | 27532492 | 25234160 | 40279672 | 37398608 | 34245772 | 38197616 | **1.526510967** | 0 |
| sp\|Q96I36 | Q96I36 | COX14 | Cytochrome c oxidase assembly protein COX14 OS=Hom | 1362855.88 | 1266253.88 | 1641228.12 | 1089464 | 2053762.25 | 1893578.38 | 1854176.25 | 2359588.5 | **1.522650569** | 0 |
| sp\|Q9UL4 | Q9UL45 | BLOC1S6 | Biogenesis of lysosome-related organelles complex 1 sub | 105988.3 | 102613.98 | 67099.26 | 111316.58 | 89773.31 | 176206.69 | 135455.72 | 187434.64 | **1.521557595** | 3 |
| sp\|P32189 | P32189 | GK | Glycerol kinase OS=Homo sapiens (Human) GN=GK PE=1 | 10674218 | 16156540 | 8875786 | 12598972 | 18602780 | 16462062 | 17778260 | 20652000 | **1.521463967** | 0 |
| sp\|Q9BRA | Q9BRA2 | TXNDC17 | Thioredoxin domain-containing protein 17 OS=Homo sa | 18018868 | 17085212 | 16844584 | 17495454 | 30572228 | 27528826 | 23393236 | 24109846 | **1.520706707** | 0 |
| sp\|O6074 | O60749 | SNX2 | Sorting nexin-2 OS=Homo sapiens (Human) GN=SNX2 PE | 14807946 | 14182534 | 13270351 | 13557469 | 23876276 | 19870814 | 21098752 | 19966826 | **1.519441975** | 0 |
| sp\|Q9ULC | Q9ULC4 | MCTS1 | Malignant T-cell-amplified sequence 1 OS=Homo sapien | 811302.12 | 345493.19 | 308073.25 | 491985.91 | 681212.19 | 1344455 | 511547.44 | 428863.19 | **1.51573756** | 4 |
| sp\|Q9278 | Q92783 | STAM | Signal transducing adapter molecule 1 OS=Homo sapien | 180059.48 | 1838589.62 | 1558244.25 | 1607687.25 | 1985909.5 | 2091471.12 | 2038194.25 | 1742027.62 | **1.515571479** | 1 |
| sp\|P17096 | P17096 | HMGA1 | High mobility group protein HMG-I/HMG-Y OS=Homo sa | 830507.38 | 1046853.5 | 946392.94 | 1059041.62 | 1201383.38 | 1514496.88 | 1673161.38 | 1494355.88 | **1.51524787** | 0 |
| sp\|P35222 | P35222 | CTNNB1 | Catenin beta-1 OS=Homo sapiens (Human) GN=CTNNB1 | 86504184 | 96442088 | 85621144 | 74805896 | 136720352 | 119971656 | 127820176 | 135482256 | **1.514370575** | 0 |
| sp\|Q9BQ3 | Q9BQ39 | DDX50 | ATP-dependent RNA helicase DDX50 OS=Homo sapiens ( | 3523591.5 | 3434960.5 | 2760715.25 | 2745198.75 | 2765331.25 | 4817992 | 4770472.5 | 6506669.5 | **1.51313865** | 2 |
| sp\|Q96PU | Q96PU5 | NEDD4L | E3 ubiquitin-protein ligase NEDD4-like OS=Homo sapien | 773254.06 | 769150.88 | 799966.69 | 723533.5 | 1106327.62 | 1036750.31 | 1352935.25 | 1140308.12 | **1.512219427** | 0 |
| sp\|P56385 | P56385 | ATP5ME | ATP synthase subunit e, mitochondrial OS=Homo sapien | 7387187 | 7377838 | 6563812 | 7452568 | 11538828 | 10572104 | 11368814 | 10012689 | **1.511129669** | 0 |
| sp\|Q9BTW | Q9BTW9 | TBCD | Tubulin-specific chaperone D OS=Homo sapiens (Human | 5803117 | 6569033 | 5709169 | 6596800.5 | 9474571 | 9870886 | 8696186 | 9236195 | **1.510562342** | 0 |
| sp\|O1473 | O14737 | PDCD5 | Programmed cell death protein 5 OS=Homo sapiens (Hu | 1126273.88 | 1020612.19 | 1204974.62 | 1892367.88 | 2014349.25 | 1737048 | 2383101.5 | 1786719 | **1.510463864** | 2 |
| sp\|Q96CT | Q96CT7 | CCDC124 | Coiled-coil domain-containing protein 124 OS=Homo sa | 0 | 534092.06 | 611558.06 | 769290.19 | 1083691 | 715646.31 | 1084531.62 | 0 | **1.505983719** | 4.5 |
| sp\|O7552 | O75521 | ECI2 | Enoyl-CoA delta isomerase 2, mitochondrial OS=Homo s | 20250094 | 20464628 | 18681764 | 19222424 | 32524302 | 27510736 | 29744230 | 28421198 | **1.503461012** | 0 |
| sp\|Q5MIZ | Q5MIZ7 | PPP4R3B | Serine/threonine-protein phosphatase 4 regulatory subu | 0 | 217028.08 | 226592.48 | 241249.31 | 160221.22 | 319296.25 | 253127.77 | 296505.16 | **1.502694811** | 3 |
| sp\|Q9253 | Q92530 | PSMF1 | Proteasome inhibitor PI31 subunit OS=Homo sapiens (Hu | 9451732 | 8202988.5 | 7655829 | 7884053 | 14101979 | 12143823 | 11686923 | 11945421 | **1.502598081** | 0 |
| sp\|Q6P3W | Q6P3W7 | SCYL2 | SCY1-like protein 2 OS=Homo sapiens (Human) GN=SCYL | 618117.25 | 786023.25 | 899724.12 | 844180.5 | 1066687.75 | 1383127.88 | 1066292.5 | 1213294.75 | **1.502330081** | 0 |
| sp\|Q8N4T | Q8N4T8 | CBR4 | Carbonyl reductase family member 4 OS=Homo sapiens | 825511.38 | 1169100.25 | 1108604.5 | 888895.19 | 1090416.38 | 1136265.88 | 1505263.25 | 2259950.5 | **1.5009341** | 3 |
| sp\|Q1345 | Q13451 | FKBP5 | Peptidyl-prolyl cis-trans isomerase FKBP5 OS=Homo sapi | 8606622 | 9141305 | 12297442 | 11433560 | 15180057 | 15250788 | 16859854 | 14930611 | **1.500070313** | 0 |
| sp\|Q9BX6 | Q9BX66 | SORBS1 | Sorbin and SH3 domain-containing protein 1 OS=Homo s | 1389263.62 | 1191046.12 | 949175.81 | 979119.81 | 1650985.12 | 1560351.75 | 1711759.75 | 1839161.62 | **1.499855876** | 0 |
| sp\|Q1547 | Q15477 | SKIV2L | Helicase SKI2W OS=Homo sapiens (Human) GN=SKIV2L F | 992360.38 | 488680.16 | 1417918 | 883776.06 | 1178135.75 | 1118311 | 1719487.5 | 1654486.25 | **1.499026789** | 2 |
| sp\|Q96C1 | Q96C19 | EFHD2 | EF-hand domain-containing protein D2 OS=Homo sapier | 7932592 | 7377961.5 | 7822271 | 8014277 | 12885041 | 15246242 | 9432467 | 9113863 | **1.498618194** | 0 |
| sp\|Q9Y58 | Q9Y580 | RBM7 | RNA-binding protein 7 OS=Homo sapiens (Human) GN=R | 482915.34 | 369666.25 | 0 | 573197.19 | 545949.44 | 610421.38 | 550808.19 | 427674.69 | **1.497324641** | 4 |
| sp\|P23434 | P23434 | GCSH | Glycine cleavage system H protein, mitochondrial OS=Hc | 925448.06 | 610743.25 | 1129434.75 | 698983.56 | 1049326 | 1023220.31 | 1548059.5 | 1417177.25 | **1.497286054** | 2 |
| sp\|O0015 | O00154 | ACOT7 | Cytosolic acyl coenzyme A thioester hydrolase OS=Homc | 10561850 | 11069782 | 10129520 | 9601188 | 16000478 | 15845090 | 14893987 | 15174944 | **1.496880955** | 0 |
| sp\|Q9Y2R | Q9Y2R0 | COA3 | Cytochrome c oxidase assembly factor 3 homolog, mitocl | 3036699.5 | 4264843 | 3704513.75 | 3663454.5 | 5265289 | 4959445 | 5275830.5 | 6452715.5 | **1.496524347** | 0 |
| sp\|Q96CV | Q96CV9 | OPTN | Optineurin OS=Homo sapiens (Human) GN=OPTN PE=1 S | 0 | 207676.56 | 226079.86 | 239550.67 | 0 | 324247.34 | 324714.81 | 356403.12 | **1.493174934** | 3.5 |
| sp\|Q96ER | Q96ER9 | CCDC51 | Coiled-coil domain-containing protein 51 OS=Homo sapi | 692098.19 | 617759.44 | 777836.25 | 638107.06 | 1060263.25 | 958118.94 | 860349.12 | 1188607 | **1.492162634** | 0 |
| sp\|P55084 | P55084 | HADHB | Trifunctional enzyme subunit beta, mitochondrial OS=Hc | 130416816 | 140500720 | 145523232 | 147975072 | 215428576 | 187855088 | 223417952 | 215193280 | **1.491621667** | 0 |

| | | | | | | | | | | | | |
|---|---|---|---|---|---|---|---|---|---|---|---|---|
| sp\|P40123 | P40123 | CAP2 | Adenylyl cyclase-associated protein 2 OS=Homo sapiens | 16545921 | 15929111 | 21476492 | 18017914 | 28053212 | 23465604 | 26951416 | 28840004 | **1.491052855** | 0 |
| sp\|P21127 | P21127 | CDK11B | Cyclin-dependent kinase 11B OS=Homo sapiens (Human | 1130026 | 1589136.25 | 461890.22 | 3287234.5 | 3652633 | 2871259.75 | 1716111 | 1396285.75 | **1.489774579** | 4 |
| sp\|P36871 | P36871 | PGM1 | Phosphoglucomutase-1 OS=Homo sapiens (Human) GN= | 27715008 | 30964380 | 32893930 | 31738108 | 63547904 | 41851756 | 41339832 | 36914548 | **1.48935136** | 0 |
| sp\|Q1500 | Q15003 | NCAPH | Condensin complex subunit 2 OS=Homo sapiens (Humar | 3074493.5 | 2855845 | 2987578.5 | 2742668.5 | 4051830.75 | 4412127 | 3606355 | 5292570.5 | **1.489023278** | 0 |
| sp\|Q9BQ6 | Q9BQ61 | TRIR | Telomerase RNA component interacting RNase OS=Hom | 1442729.62 | 1507891.75 | 0 | 1315143.38 | 1855008.5 | 1279147.38 | 1717229.62 | 1496513 | **1.488103276** | 4 |
| sp\|Q9Y2V | Q9Y2V2 | CARHSP1 | Calcium-regulated heat-stable protein 1 OS=Homo sapie | 3264360 | 2199535.75 | 1428681.38 | 2101334.5 | 3524402.5 | 4182060 | 3511819.75 | 2146352.25 | **1.485964622** | 2 |
| sp\|P55769 | P55769 | SNU13 | NHP2-like protein 1 OS=Homo sapiens (Human) GN=SNU | 11884596 | 10652520 | 14118688 | 15981468 | 20567184 | 19518326 | 20406082 | 17551828 | **1.482664603** | 0 |
| sp\|P30740 | P30740 | SERPINB1 | Leukocyte elastase inhibitor OS=Homo sapiens (Human) | 13077282 | 16271590 | 16895176 | 14264476 | 21910780 | 23846024 | 21618130 | 22288662 | **1.48183413** | 0 |
| sp\|Q1423 | Q14232 | EIF2B1 | Translation initiation factor eIF-2B subunit alpha OS=Hor | 5059701.5 | 4585791.5 | 5455318 | 4382265 | 7319555 | 6704947 | 7046510.5 | 7794686.5 | **1.481578114** | 0 |
| sp\|Q0224 | Q02241 | KIF23 | Kinesin-like protein KIF23 OS=Homo sapiens (Human) GN | 0 | 428166.25 | 686090.38 | 622528.75 | 538460.44 | 658862.31 | 699147 | 673428.12 | **1.479686494** | 4 |
| sp\|P49902 | P49902 | NT5C2 | Cytosolic purine 5'-nucleotidase OS=Homo sapiens (Hum | 2079594 | 2584372 | 2322956 | 2610914.75 | 2941104 | 3519451 | 3549862 | 4189465.25 | **1.479487787** | 0 |
| sp\|P09110 | P09110 | ACAA1 | 3-ketoacyl-CoA thiolase, peroxisomal OS=Homo sapiens | 22523406 | 18495500 | 29153744 | 28589994 | 41666584 | 37716028 | 40206924 | 26502428 | **1.479222893** | 2 |
| sp\|Q96DI7 | Q96DI7 | SNRNP40 | U5 small nuclear ribonucleoprotein 40 kDa protein OS=H | 6781745 | 3255052 | 3034948 | 3552549 | 7119898 | 6618280 | 4293933 | 6508547 | **1.476192493** | 3 |
| sp\|P53602 | P53602 | MVD | Diphosphomevalonate decarboxylase OS=Homo sapiens | 2941599.5 | 2689445.75 | 5190198.5 | 3528773.5 | 6258413.5 | 5411587 | 5123114.5 | 4365266 | **1.474449865** | 2 |
| sp\|Q8N5N | Q8N5N7 | MRPL50 | 39S ribosomal protein L50, mitochondrial OS=Homo sapi | 2670394.75 | 727217.56 | 2164805.5 | 2226767.5 | 3110215.5 | 2590615 | 3293624 | 2480184.25 | **1.473150053** | 2 |
| sp\|Q9Y5Z | Q9Y5Z9 | UBIAD1 | UbiA prenyltransferase domain-containing protein 1 OS= | 1333689.88 | 633952.62 | 0 | 1063433.88 | 0 | 1436287.12 | 1585436.12 | 1442143.88 | **1.472700309** | 3.5 |
| sp\|P34932 | P34932 | HSPA4 | Heat shock 70 kDa protein 4 OS=Homo sapiens (Human) | 96419976 | 105014360 | 120876896 | 96305512 | 163845728 | 152712272 | 151498848 | 148282016 | **1.472322531** | 0 |
| sp\|P51649 | P51649 | ALDH5A1 | Succinate-semialdehyde dehydrogenase, mitochondrial | 17902532 | 16993294 | 20074464 | 18470522 | 23557920 | 24325878 | 32506526 | 27692970 | **1.471706141** | 0 |
| sp\|P61020 | P61020 | RAB5B | Ras-related protein Rab-5B OS=Homo sapiens (Human) ( | 4104050.25 | 5894090 | 6119253 | 6353972 | 8814321 | 7408433 | 7937405 | 8897190 | **1.471087699** | 0 |
| sp\|Q1308 | Q13084 | MRPL28 | 39S ribosomal protein L28, mitochondrial OS=Homo sapi | 5402894 | 3842377 | 3643003.75 | 4583206 | 5832402 | 6255675.5 | 7046540 | 6561491 | **1.47074589** | 0 |
| sp\|P51858 | P51858 | HDGF | Hepatoma-derived growth factor OS=Homo sapiens (Hu | 14829880 | 14249402 | 14825792 | 15511528 | 21299878 | 20168770 | 25516738 | 20383578 | **1.470446997** | 0 |
| sp\|Q8TC1 | Q8TC12 | RDH11 | Retinol dehydrogenase 11 OS=Homo sapiens (Human) G | 2651456.5 | 5174450.5 | 5248453.5 | 4615343 | 5707802 | 6425358.5 | 6934223 | 6942844 | **1.470359721** | 0 |
| sp\|Q9BTT | Q9BTT6 | LRRC1 | Leucine-rich repeat-containing protein 1 OS=Homo sapie | 5609629 | 5018909 | 4746079.5 | 3552221.5 | 6895091 | 7665666.5 | 6177139 | 7074464 | **1.469466745** | 0 |
| sp\|P45880 | P45880 | VDAC2 | Voltage-dependent anion-selective channel protein 2 OS | 89705784 | 90443888 | 103915712 | 95009000 | 143418704 | 101531576 | 146584752 | 164785728 | **1.46757677** | 1 |
| sp\|O1461 | O14618 | CCS | Copper chaperone for superoxide dismutase OS=Homo s | 0 | 0 | 0 | 561315.06 | 467541.81 | 0 | 0 | 355851.91 | **1.466901173** | 7 |
| sp\|P46939 | P46939 | UTRN | Utrophin OS=Homo sapiens (Human) GN=UTRN PE=1 SV | 31531376 | 32453916 | 23714074 | 24932326 | 40120276 | 33938112 | 46068616 | 45067576 | **1.466679378** | 0 |
| sp\|P21695 | P21695 | GPD1 | Glycerol-3-phosphate dehydrogenase [NAD(+)], cytoplasi | 9326788 | 12434853 | 11906752 | 11280542 | 16196466 | 15520420 | 17761422 | 16369111 | **1.464938357** | 0 |
| sp\|P53999 | P53999 | SUB1 | Activated RNA polymerase II transcriptional coactivator p | 52809728 | 49197352 | 40789568 | 45514736 | 66408844 | 61141836 | 78627680 | 69531688 | **1.464117793** | 0 |
| sp\|P09327 | P09327 | VIL1 | Villin-1 OS=Homo sapiens (Human) GN=VIL1 PE=1 SV=4 | 224926016 | 221301328 | 230800672 | 213014544 | 335151296 | 306720576 | 336120256 | 322814944 | **1.46151109** | 0 |
| sp\|Q9H49 | Q9H490 | PIGU | Phosphatidylinositol glycan anchor biosynthesis class U p | 0 | 1708527.5 | 1396165.88 | 1870968.75 | 1755874.88 | 2049950.75 | 1472048.25 | 1992260.38 | **1.461139054** | 3 |
| sp\|O7602 | O76024 | WFS1 | Wolframin OS=Homo sapiens (Human) GN=WFS1 PE=1 S | 8740817 | 11153552 | 7028800.5 | 8227409.5 | 13686200 | 12150114 | 12903260 | 12560345 | **1.459433115** | 0 |
| sp\|O7539 | O75390 | CS | Citrate synthase, mitochondrial OS=Homo sapiens (Hum | 36181396 | 37029304 | 36992000 | 46403304 | 54724816 | 54303492 | 54511584 | 64991036 | **1.459273094** | 0 |
| sp\|Q8N6C | Q8N6C5 | IGSF1 | Immunoglobulin superfamily member 1 OS=Homo sapie | 4359325.5 | 4565247.5 | 4579919 | 3802539.5 | 6469970 | 5309055.5 | 6716677 | 6758823.5 | **1.459206104** | 0 |
| sp\|P22694 | P22694 | PRKACB | cAMP-dependent protein kinase catalytic subunit beta C | 3293215 | 3248980 | 4229201 | 5520708 | 5285669 | 6707252 | 4927724.5 | 6830828 | **1.457851822** | 2 |
| sp\|Q9NYH | Q9NYH9 | UTP6 | U3 small nucleolar RNA-associated protein 6 homolog O | 5740384 | 5548819 | 4723621 | 6230545 | 7913809.5 | 6105924 | 8105819 | 10293742 | **1.457481306** | 1 |
| sp\|P38646 | P38646 | HSPA9 | Stress-70 protein, mitochondrial OS=Homo sapiens (Hun | 497881088 | 516803552 | 493853408 | 519688800 | 802866688 | 678280128 | 806202624 | 667256448 | **1.456743308** | 0 |
| sp\|P20338 | P20338 | RAB4A | Ras-related protein Rab-4A OS=Homo sapiens (Human) ( | 2779279 | 1177898 | 851460.88 | 2782756.5 | 2624956.25 | 3369471.75 | 2099556.5 | 2961588.5 | **1.456329687** | 4 |
| sp\|O7595 | O75955 | FLOT1 | Flotillin-1 OS=Homo sapiens (Human) GN=FLOT1 PE=1 S\ | 6306224.5 | 7439181 | 9130707 | 8137910.5 | 10978217 | 11582928 | 10052545 | 12530513 | **1.455606162** | 0 |
| sp\|Q8N5M | Q8N5M9 | JAGN1 | Protein jagunal homolog 1 OS=Homo sapiens (Human) G | 3666928.5 | 4129899 | 3509180.25 | 3901855.25 | 4708496 | 5109160 | 6325789 | 5972804 | **1.454264087** | 0 |
| sp\|Q9NQP | Q9NQP4 | PFDN4 | Prefoldin subunit 4 OS=Homo sapiens (Human) GN=PFDI | 639470.19 | 1308086.5 | 446042.75 | 1517316.25 | 1753628.25 | 714416.69 | 1662606.75 | 1556587.38 | **1.454196286** | 2 |
| sp\|O6044 | O60443 | GSDME | Gasdermin-E OS=Homo sapiens (Human) GN=GSDME PE | 2281358 | 1568119.75 | 1994758.75 | 2326393 | 2630044 | 3356734.25 | 2378414 | 3507484.25 | **1.453092017** | 0 |
| sp\|P04004 | P04004 | VTN | Vitronectin OS=Homo sapiens (Human) GN=VTN PE=1 S\ | 552227.31 | 482372 | 654072.19 | 552478.5 | 666809.19 | 1069895.88 | 841061.12 | 678039.12 | **1.452738688** | 0 |
| sp\|Q9NX2 | Q9NX24 | NHP2 | H/ACA ribonucleoprotein complex subunit 2 OS=Homo s | 1804336.5 | 2011480 | 1781391.25 | 1496206.62 | 2403750.5 | 2156890.75 | 2955649 | 2787375.75 | **1.452567898** | 0 |
| sp\|P19404 | P19404 | NDUFV2 | NADH dehydrogenase [ubiquinone] flavoprotein 2, mitoc | 3594912 | 3404907.75 | 2641446 | 2842822 | 4390174.5 | 4078345.25 | 5324893 | 4340551.5 | **1.452566228** | 0 |
| sp\|Q8N0V | Q8N0V3 | RBFA | Putative ribosome-binding factor A, mitochondrial OS=H | 0 | 96223.24 | 78428.88 | 208677.06 | 120306.87 | 142226.77 | 142149.08 | 151838.25 | **1.451809565** | 4 |

| | | | | | | | | | | | | | |
|---|---|---|---|---|---|---|---|---|---|---|---|---|---|
| sp\|Q9294 | Q92947 | GCDH | Glutaryl-CoA dehydrogenase, mitochondrial OS=Homo s | 757472.62 | 1088641.12 | 1891576.25 | 1274144.5 | 1834355.5 | 1219548.12 | 2140053.25 | 2080961.25 | 1.451547958 | 3 |
| sp\|O6081 | O60814 | HIST1H2BK | Histone H2B type 1-K OS=Homo sapiens (Human) GN=HI | 346377952 | 316260864 | 342220512 | 429763200 | 611639296 | 591522560 | 470839360 | 407594752 | 1.450971198 | 1 |
| sp\|Q1283 | Q12830 | BPTF | Nucleosome-remodeling factor subunit BPTF OS=Homo | 238176.84 | 0 | 202327.34 | 0 | 0 | 304360.78 | 0 | 334743.38 | 1.450846982 | 6 |
| sp\|P26639 | P26639 | TARS | Threonine--tRNA ligase, cytoplasmic OS=Homo sapiens ( | 46560676 | 48777292 | 44515092 | 47644924 | 70843888 | 65904368 | 68072696 | 67197552 | 1.450780953 | 0 |
| sp\|P17612 | P17612 | PRKACA | cAMP-dependent protein kinase catalytic subunit alpha | 3293215 | 3248980 | 4229201 | 5243497 | 5285669 | 6707252 | 4927724.5 | 6310936 | 1.450623585 | 1 |
| sp\|Q8TAE | Q8TAE8 | GADD45GI | Growth arrest and DNA damage-inducible proteins-intera | 936828.38 | 935834.19 | 966238.19 | 1329853 | 1381896.75 | 1300645.88 | 1475651.88 | 1886236.88 | 1.449937257 | 1 |
| sp\|O9516 | O95168 | NDUFB4 | NADH dehydrogenase [ubiquinone] 1 beta subcomplex s | 2657011.5 | 2111848.25 | 2088842.75 | 2613016.75 | 3545930.25 | 2766149.5 | 3701231.25 | 3717278.5 | 1.449793742 | 0 |
| sp\|P10644 | P10644 | PRKAR1A | cAMP-dependent protein kinase type I-alpha regulatory | 5201789.5 | 7577555 | 5841445.5 | 4090629 | 8639562 | 8293399.5 | 8046538 | 7935171 | 1.449256451 | 0 |
| sp\|Q9BRQ | Q9BRQ8 | AIFM2 | Apoptosis-inducing factor 2 OS=Homo sapiens (Human) | 4144235 | 4847012 | 5809132 | 5099785 | 6326386 | 6605147.5 | 6601366 | 9268102 | 1.44727458 | 0 |
| sp\|Q9961 | Q99611 | SEPHS2 | Selenide, water dikinase 2 OS=Homo sapiens (Human) G | 2687318 | 1544191.5 | 1341382.88 | 1282582 | 2324257.5 | 2043603.5 | 3086124 | 2463199.5 | 1.446608061 | 3 |
| sp\|Q9Y2Q | Q9Y2Q9 | MRPS28 | 28S ribosomal protein S28, mitochondrial OS=Homo sap | 6982305 | 6640347 | 2811713.5 | 2728385.25 | 6971938 | 6303665 | 5948065 | 8493531 | 1.446410245 | 5 |
| sp\|O6061 | O60610 | DIAPH1 | Protein diaphanous homolog 1 OS=Homo sapiens (Huma | 7365785.5 | 8302765 | 8592054 | 6251292.5 | 10967411 | 11228409 | 11082101 | 10820082 | 1.445272413 | 0 |
| sp\|Q9UMI | Q9UMR2 | DDX19B | ATP-dependent RNA helicase DDX19B OS=Homo sapiens | 11433342 | 14174732 | 14735682 | 13739839 | 18339388 | 19658382 | 18204152 | 21899166 | 1.444080927 | 0 |
| sp\|Q9UL1 | Q9UL12 | SARDH | Sarcosine dehydrogenase, mitochondrial OS=Homo sapi | 3745094.5 | 2774762.5 | 1529361.5 | 4071231 | 3887421 | 4875052 | 4391657 | 4348409 | 1.444050322 | 1 |
| sp\|O7534 | O75340 | PDCD6 | Programmed cell death protein 6 OS=Homo sapiens (Hu | 9140045 | 9470796 | 8173219 | 9962290 | 14601542 | 13188536 | 14359828 | 10872337 | 1.442925433 | 0 |
| sp\|Q9970 | Q99700 | ATXN2 | Ataxin-2 OS=Homo sapiens (Human) GN=ATXN2 PE=1 SV | 537311.12 | 906966.12 | 991981.75 | 683226.44 | 1646589.12 | 0 | 1397729.25 | 1456715.75 | 1.442877109 | 4 |
| sp\|Q86VP | Q86VP6 | CAND1 | Cullin-associated NEDD8-dissociated protein 1 OS=Homo | 64106680 | 58885172 | 63405880 | 64561756 | 87105552 | 89242960 | 89042544 | 96384728 | 1.441570458 | 0 |
| sp\|Q1502 | Q15020 | SART3 | Squamous cell carcinoma antigen recognized by T-cells 3 | 16461000 | 18800778 | 21157642 | 15330319 | 23173322 | 23135226 | 28070378 | 29018472 | 1.44108396 | 0 |
| sp\|P30041 | P30041 | PRDX6 | Peroxiredoxin-6 OS=Homo sapiens (Human) GN=PRDX6 | 116316880 | 112600608 | 121663296 | 120288584 | 173459984 | 162175840 | 174891024 | 167832000 | 1.440651896 | 0 |
| sp\|Q1324 | Q13242 | SRSF9 | Serine/arginine-rich splicing factor 9 OS=Homo sapiens ( | 16945550 | 14345246 | 16035829 | 17179184 | 23085842 | 21330466 | 25663386 | 22780800 | 1.439567931 | 0 |
| sp\|Q9Y4W | Q9Y4W6 | AFG3L2 | AFG3-like protein 2 OS=Homo sapiens (Human) GN=AFG | 4488226 | 4149089 | 3167597 | 8415682 | 8497574 | 8167773 | 2763895.5 | 9679392 | 1.439553878 | 5 |
| sp\|Q9NVH | Q9NVH0 | EXD2 | Exonuclease 3'-5' domain-containing protein 2 OS=Homo | 246868.27 | 214609.98 | 287101.94 | 244824.14 | 311497.03 | 410537.56 | 400784.28 | 305923.09 | 1.438228038 | 0 |
| sp\|P49588 | P49588 | AARS | Alanine--tRNA ligase, cytoplasmic OS=Homo sapiens (Hu | 68046472 | 63515400 | 63160736 | 66009760 | 97112616 | 92022248 | 92547568 | 92818272 | 1.436341437 | 0 |
| sp\|Q8NE8 | Q8NE86 | MCU | Calcium uniporter protein, mitochondrial OS=Homo sapi | 3052047.75 | 3692346.5 | 2686316.75 | 3071650.5 | 4959392 | 3421863.25 | 4159603.25 | 5408122 | 1.435647218 | 1 |
| sp\|O0062 | O00625 | PIR | Pirin OS=Homo sapiens (Human) GN=PIR PE=1 SV=1 | 1725611.88 | 1976122.62 | 2142562 | 1613762.62 | 2752814.5 | 3067992.5 | 1868114 | 3017393.5 | 1.435536287 | 2 |
| sp\|O9487 | O94874 | UFL1 | E3 UFM1-protein ligase 1 OS=Homo sapiens (Human) GN | 10188502 | 8666387 | 10678649 | 10455634 | 13485161 | 14197250 | 14182639 | 15537222 | 1.435445375 | 0 |
| sp\|P35611 | P35611 | ADD1 | Alpha-adducin OS=Homo sapiens (Human) GN=ADD1 PE | 6282637.5 | 5151073.5 | 5823097.5 | 6640476.5 | 8226528 | 8393905 | 9297337 | 8374070 | 1.434968031 | 0 |
| sp\|P01024 | P01024 | C3 | Complement C3 OS=Homo sapiens (Human) GN=C3 PE=1 | 56477588 | 63556620 | 62673068 | 63516984 | 101728480 | 71901832 | 92349896 | 86863856 | 1.433019086 | 0 |
| sp\|P14625 | P14625 | HSP90B1 | Endoplasmin OS=Homo sapiens (Human) GN=HSP90B1 F | 837642432 | 900220992 | 971772224 | 849272960 | 1213478016 | 1276050816 | 1264521216 | 1344465664 | 1.43260653 | 0 |
| sp\|Q9H9B | Q9H9B4 | SFXN1 | Sideroflexin-1 OS=Homo sapiens (Human) GN=SFXN1 PE | 27069458 | 24674204 | 28995604 | 27600832 | 42747980 | 36003860 | 36967964 | 39377520 | 1.431578214 | 0 |
| sp\|Q1671 | Q16718 | NDUFA5 | NADH dehydrogenase [ubiquinone] 1 alpha subcomplex | 5830132 | 4704206 | 3381919.75 | 4747163 | 7040454.5 | 6512916 | 6487699.5 | 6643186 | 1.429762333 | 0 |
| sp\|Q9BTY | Q9BTY2 | FUCA2 | Plasma alpha-L-fucosidase OS=Homo sapiens (Human) G | 1163539.25 | 1724305.25 | 1446544.25 | 1682204 | 2023082.38 | 1401131 | 2208868.5 | 2961240 | 1.428436698 | 3 |
| sp\|Q8N5V | Q8N5V2 | NGEF | Ephexin-1 OS=Homo sapiens (Human) GN=NGEF PE=1 S\ | 240825.3 | 289327.44 | 234855.25 | 278804.06 | 294094.28 | 354350.72 | 410072.34 | 430876.91 | 1.426879724 | 0 |
| sp\|O7600 | O76003 | GLRX3 | Glutaredoxin-3 OS=Homo sapiens (Human) GN=GLRX3 P | 34662272 | 33915072 | 39814408 | 33422024 | 50077412 | 53241272 | 50623868 | 48366956 | 1.426585722 | 0 |
| sp\|P61916 | P61916 | NPC2 | NPC intracellular cholesterol transporter 2 OS=Homo sap | 1822354.5 | 1699701.5 | 1561659.5 | 1758191.25 | 2068279 | 2014864.75 | 2885833.5 | 2781347.75 | 1.425088847 | 0 |
| sp\|P55209 | P55209 | NAP1L1 | Nucleosome assembly protein 1-like 1 OS=Homo sapiens | 32638300 | 29511572 | 28951860 | 28173920 | 42193952 | 43876936 | 41644136 | 42180920 | 1.424397529 | 0 |
| sp\|Q2T9J0 | Q2T9J0 | TYSND1 | Peroxisomal leader peptide-processing protease OS=Ho | 241243.17 | 0 | 213063.81 | 221005.08 | 236396 | 281989.91 | 228857.28 | 214607.3 | 1.424305217 | 4 |
| sp\|Q0032 | Q00325 | SLC25A3 | Phosphate carrier protein, mitochondrial OS=Homo sapi | 89694576 | 97030368 | 85015760 | 109714832 | 155416544 | 149334480 | 109227664 | 128938784 | 1.423278523 | 1 |
| sp\|Q8NBF | Q8NBF2 | NHLRC2 | NHL repeat-containing protein 2 OS=Homo sapiens (Hun | 1036272 | 1746184.5 | 1341144 | 1518561.75 | 1439229.62 | 1546407.62 | 2500279 | 2538732 | 1.422264707 | 3 |
| sp\|O6022 | O60220 | TIMM8A | Mitochondrial import inner membrane translocase subu | 1859552 | 0 | 1796718.75 | 1751690.88 | 1638912.62 | 1821430.25 | 2017974.62 | 2211519.25 | 1.421947356 | 4 |
| sp\|P45984 | P45984 | MAPK9 | Mitogen-activated protein kinase 9 OS=Homo sapiens (H | 315893.56 | 328180.62 | 357086.47 | 313996.28 | 429565.69 | 502262.22 | 397339.88 | 538427.19 | 1.420054852 | 0 |
| sp\|P05556 | P05556 | ITGB1 | Integrin beta-1 OS=Homo sapiens (Human) GN=ITGB1 PE | 34327768 | 33576208 | 34503428 | 32725180 | 53228464 | 52175652 | 43959452 | 42514952 | 1.419927854 | 0 |
| sp\|Q9BRG | Q9BRG1 | VPS25 | Vacuolar protein-sorting-associated protein 25 OS=Homo | 363272.34 | 311423.97 | 420499.62 | 265733.5 | 429221.88 | 534843.81 | 494526.62 | 472124.66 | 1.418675302 | 0 |
| sp\|Q9250 | Q92506 | HSD17B8 | Estradiol 17-beta-dehydrogenase 8 OS=Homo sapiens (H | 1216790.75 | 1992404.38 | 1298262.75 | 1451678 | 1973945.25 | 2357816.5 | 1965063.38 | 2151458 | 1.417702717 | 2 |
| sp\|P19440 | P19440 | GGT1 | Glutathione hydrolase 1 proenzyme OS=Homo sapiens (I | 2052467 | 2068868.62 | 2332722.5 | 2121152.25 | 3864750.5 | 2718959.75 | 2439430.25 | 3130999.25 | 1.417357619 | 0 |

| | | | | | | | | | | | | | |
|---|---|---|---|---|---|---|---|---|---|---|---|---|---|
| sp\|P05114 | P05114 | HMGN1 | Non-histone chromosomal protein HMG-14 OS=Homo sa | 825965.69 | 676218 | 1265247.62 | 640921.19 | 1312305.12 | 1031093.75 | 1360499.5 | 1125664.12 | **1.416978581** | 2 |
| sp\|P49189 | P49189 | ALDH9A1 | 4-trimethylaminobutyraldehyde dehydrogenase OS=Hor | 4948760 | 5615596 | 4806463 | 4842051 | 8846529 | 6776328.5 | 6997234.5 | 5985873.5 | **1.415235219** | 0 |
| sp\|O1497/ | O14974 | PPP1R12A | Protein phosphatase 1 regulatory subunit 12A OS=Homo | 2828592 | 2563821 | 1208376.5 | 2114433.5 | 3044509 | 2302613.25 | 3853045.75 | 3132187.5 | **1.415036138** | 2 |
| sp\|P09622 | P09622 | DLD | Dihydrolipoyl dehydrogenase, mitochondrial OS=Homo s | 25013176 | 29443014 | 23392004 | 24308738 | 37715660 | 34097796 | 39620444 | 33028342 | **1.414120796** | 0 |
| sp\|P13674 | P13674 | P4HA1 | Prolyl 4-hydroxylase subunit alpha-1 OS=Homo sapiens ( | 30961788 | 28750930 | 29241026 | 27375160 | 42008256 | 51486064 | 37742696 | 33120548 | **1.412869531** | 0 |
| sp\|O4316! | O43169 | CYB5B | Cytochrome b5 type B OS=Homo sapiens (Human) GN=C | 22971336 | 20355388 | 19195190 | 18623424 | 29706546 | 34189080 | 25885416 | 24831490 | **1.412435204** | 0 |
| sp\|Q96H2( | Q96H20 | SNF8 | Vacuolar-sorting protein SNF8 OS=Homo sapiens (Huma | 726599.06 | 542159.44 | 1072580.62 | 666910.81 | 927385.69 | 1314800.88 | 1000963.81 | 1004726.12 | **1.412075658** | 3 |
| sp\|Q9306; | Q93062 | RBPMS | RNA-binding protein with multiple splicing OS=Homo sa | 1323349.75 | 1228498.88 | 1150154.12 | 1314918.62 | 1594967.5 | 1797710.5 | 1889867.5 | 1799447.75 | **1.411621337** | 0 |
| sp\|Q9H00 | Q9H008 | LHPP | Phospholysine phosphohistidine inorganic pyrophosphat | 355029.53 | 279626.59 | 138833.03 | 241663.83 | 334419.78 | 469331.19 | 385433.19 | 243647.14 | **1.411443722** | 3 |
| sp\|P57678 | P57678 | GEMIN4 | Gem-associated protein 4 OS=Homo sapiens (Human) GI | 1502015.75 | 29303890 | 1627590.12 | 30415434 | 1440115.12 | 30248178 | 24553758 | 32460090 | **1.411354836** | 7 |
| sp\|Q969U | Q969U7 | PSMG2 | Proteasome assembly chaperone 2 OS=Homo sapiens (H | 0 | 949323.31 | 1085495.38 | 990170.69 | 1022368.38 | 1340525.5 | 999988.88 | 903015.44 | **1.410219232** | 5 |
| sp\|O60341 | O60341 | KDM1A | Lysine-specific histone demethylase 1A OS=Homo sapier | 5549020.5 | 3662302.75 | 4347061.5 | 4074230.25 | 7244189.5 | 6093398 | 4996155 | 6529875 | **1.410092462** | 1 |
| sp\|Q9Y31! | Q9Y315 | DERA | Deoxyribose-phosphate aldolase OS=Homo sapiens (Hur | 3630312.75 | 5472202 | 4175102 | 6311843 | 6954318.5 | 7478827.5 | 6404366 | 6784953 | **1.410067728** | 0 |
| sp\|P35573 | P35573 | AGL | Glycogen debranching enzyme OS=Homo sapiens (Huma | 1273081.62 | 1378786.62 | 1034287.94 | 1037955.94 | 1367767 | 2545069.25 | 1174379 | 1571029.12 | **1.409417093** | 3 |
| sp\|Q8WVJ | Q8WVJ2 | NUDCD2 | NudC domain-containing protein 2 OS=Homo sapiens (H | 2413564.25 | 2877353.75 | 3067575 | 2371905.5 | 3784911.25 | 3695613.5 | 3912155.5 | 3730730 | **1.409398752** | 0 |
| sp\|O7535( | O75356 | ENTPD5 | Ectonucleoside triphosphate diphosphohydrolase 5 OS= | 15728617 | 14144105 | 17947692 | 2418050.25 | 17013026 | 18821616 | 17329452 | 17634086 | **1.409242521** | 3 |
| sp\|Q9UHY | Q9UHY7 | ENOPH1 | Enolase-phosphatase E1 OS=Homo sapiens (Human) GN | 1041884.38 | 1005275.62 | 1073248.12 | 1978713.25 | 1878769.38 | 2268312.25 | 1799879.12 | 1237557 | **1.408971709** | 3 |
| sp\|Q9NR3 | Q9NR31 | SAR1A | GTP-binding protein SAR1a OS=Homo sapiens (Human) ( | 20506562 | 19205904 | 24133276 | 23068144 | 30497996 | 30069662 | 29109752 | 32639712 | **1.407336936** | 0 |
| sp\|P14406 | P14406 | COX7A2 | Cytochrome c oxidase subunit 7A2, mitochondrial OS=H | 15193570 | 16887292 | 14291464 | 18404962 | 23693154 | 21807584 | 22445666 | 23211886 | **1.407256969** | 0 |
| sp\|P61106 | P61106 | RAB14 | Ras-related protein Rab-14 OS=Homo sapiens (Human) ( | 17384596 | 18432840 | 17759084 | 23942416 | 19284690 | 27969460 | 48004396 | 13828690 | **1.407233402** | 5 |
| sp\|Q9964: | Q99643 | SDHC | Succinate dehydrogenase cytochrome b560 subunit, mit | 2156808.75 | 2190513.75 | 1820100.62 | 1742646.62 | 2328032 | 2994214.25 | 2358794.25 | 3443232 | **1.406343163** | 0 |
| sp\|O9542! | O95425 | SVIL | Supervillin OS=Homo sapiens (Human) GN=SVIL PE=1 SV | 0 | 363253 | 480337.19 | 438929.41 | 416405.72 | 383455.09 | 404806.09 | 598393.25 | **1.405873368** | 6 |
| sp\|P84090 | P84090 | ERH | Enhancer of rudimentary homolog OS=Homo sapiens (H | 14476173 | 12805902 | 9966582 | 11789148 | 19867840 | 15238180 | 19703188 | 14094688 | **1.405117868** | 1 |
| sp\|Q9UKG | Q9UKG1 | APPL1 | DCC-interacting protein 13-alpha OS=Homo sapiens (Hu | 605122.44 | 0 | 434060.16 | 454444.94 | 382299.28 | 547046.44 | 660277.94 | 508855.75 | **1.404954953** | 5 |
| sp\|P34896 | P34896 | SHMT1 | Serine hydroxymethyltransferase, cytosolic OS=Homo sa | 12368908 | 12367680 | 12050208 | 13566738 | 17665740 | 16761294 | 16839216 | 19411882 | **1.403637965** | 0 |
| sp\|P61604 | P61604 | HSPE1 | 10 kDa heat shock protein, mitochondrial OS=Homo sapi | 205377088 | 153968976 | 134753328 | 172979136 | 262548864 | 218821728 | 257490464 | 197123712 | **1.403110322** | 1 |
| sp\|Q9NP7 | Q9NP72 | RAB18 | Ras-related protein Rab-18 OS=Homo sapiens (Human) ( | 3770338 | 4840457 | 4758869.5 | 4416003 | 6632317 | 6363105 | 5815050 | 6137065.5 | **1.402676481** | 0 |
| sp\|P23919 | P23919 | DTYMK | Thymidylate kinase OS=Homo sapiens (Human) GN=DTY | 17119102 | 15189018 | 16128825 | 17099200 | 23726936 | 22589344 | 23538718 | 21943554 | **1.400731642** | 0 |
| sp\|P29317 | P29317 | EPHA2 | Ephrin type-A receptor 2 OS=Homo sapiens (Human) GN | 893772.75 | 755862.94 | 532771.62 | 553222.62 | 964085.31 | 901274.12 | 1030710.38 | 935589.25 | **1.400649634** | 0 |
| sp\|Q7Z43 | Q7Z434 | MAVS | Mitochondrial antiviral-signaling protein OS=Homo sapie | 12258900 | 12649264 | 13038196 | 13032673 | 18421886 | 20191390 | 15316321 | 17454702 | **1.400267812** | 0 |
| sp\|Q9Y4A! | Q9Y4A5 | TRRAP | Transformation/transcription domain-associated protein | 151251.17 | 572142.62 | 165957.42 | 449409.03 | 445282.38 | 359105.31 | 319658.97 | 749771.5 | **1.399666724** | 6 |
| sp\|Q0820! | Q08209 | PPP3CA | Serine/threonine-protein phosphatase 2B catalytic subur | 3264706.25 | 3935196 | 4597611.5 | 3988527.25 | 5638806 | 6720809.5 | 3964677.75 | 5769084.5 | **1.399551525** | 2 |
| sp\|P56556 | P56556 | NDUFA6 | NADH dehydrogenase [ubiquinone] 1 alpha subcomplex | 8916004 | 8075644.5 | 8229498.5 | 4429264.5 | 10993374 | 10101452 | 10158462 | 10182553 | **1.397479458** | 0 |
| sp\|Q9UNF | Q9UNF1 | MAGED2 | Melanoma-associated antigen D2 OS=Homo sapiens (Hu | 1697305.62 | 3589694.75 | 4880192 | 3159477.5 | 4764187 | 3406915.5 | 5124621 | 5308562.5 | **1.396019124** | 3 |
| sp\|Q9P0J7 | Q9P0J7 | KCMF1 | E3 ubiquitin-protein ligase KCMF1 OS=Homo sapiens (Hu | 441505.34 | 588145.5 | 505180.38 | 707907.62 | 838983.44 | 792151.06 | 679872.38 | 819696.88 | **1.39592881** | 1 |
| sp\|Q0VDF! | Q0VDF9 | HSPA14 | Heat shock 70 kDa protein 14 OS=Homo sapiens (Human | 1757100.25 | 4708763 | 3120459 | 3112643 | 4227695.5 | 3948977.5 | 4207582 | 5340052 | **1.395728443** | 3 |
| sp\|Q8TBX | Q8TBX8 | PIP4K2C | Phosphatidylinositol 5-phosphate 4-kinase type-2 gamm | 830292.06 | 731587.38 | 858312.06 | 852623.5 | 1207006.25 | 1173168.12 | 1022417.62 | 1159419.5 | **1.393910591** | 0 |
| sp\|Q8NCV | Q8NCW5 | NAXE | NAD(P)H-hydrate epimerase OS=Homo sapiens (Human) | 6720527 | 5709269 | 4802570 | 5723125 | 7602471 | 8188325 | 8811688 | 7379899 | **1.393234542** | 0 |
| sp\|P40939 | P40939 | HADHA | Trifunctional enzyme subunit alpha, mitochondrial OS=H | 133401472 | 128612008 | 139392832 | 144635056 | 181019936 | 175777168 | 205211488 | 198493216 | **1.392754931** | 0 |
| sp\|Q96M2 | Q96M27 | PRRC1 | Protein PRRC1 OS=Homo sapiens (Human) GN=PRRC1 PE | 8737761 | 9004615 | 5840046 | 8949710 | 11935116 | 7078329.5 | 11838228 | 14448063 | **1.392461352** | 3 |
| sp\|Q9268! | Q92685 | ALG3 | Dol-P-Man:Man(5)GlcNAc(2)-PP-Dol alpha-1,3-mannosyl | 0 | 570273.88 | 0 | 613702.56 | 521857.94 | 463319.75 | 0 | 660043.94 | **1.389572946** | 7 |
| sp\|O9586! | O95865 | DDAH2 | N(G),N(G)-dimethylarginine dimethylaminohydrolase 2 ( | 3604541.5 | 3323191.25 | 3236832.25 | 3455985.5 | 4590677 | 4308676.5 | 5133317 | 4891689 | **1.38939755** | 0 |
| sp\|P50452 | P50452 | SERPINB8 | Serpin B8 OS=Homo sapiens (Human) GN=SERPINB8 PE= | 8621255 | 10470842 | 8919385 | 9187368 | 10434650 | 14016200 | 12927328 | 14291900 | **1.389023532** | 1 |
| sp\|P36776 | P36776 | LONP1 | Lon protease homolog, mitochondrial OS=Homo sapiens | 39479280 | 39863968 | 39218476 | 39718528 | 50549824 | 56144460 | 54916444 | 58087740 | **1.388034611** | 0 |
| sp\|Q96DG | Q96DG6 | CMBL | Carboxymethylenebutenolidase homolog OS=Homo sap | 1899916 | 1611332.62 | 1733008.5 | 2030827 | 2651651.75 | 2885351.5 | 2291973 | 2262713.75 | **1.387157844** | 0 |

| | | | | | | | | | | | | | |
|---|---|---|---|---|---|---|---|---|---|---|---|---|---|
| sp\|P24928 | P24928 | POLR2A | DNA-directed RNA polymerase II subunit RPB1 OS=Homo | 2900529.5 | 1607080.75 | 2756932.5 | 2762899 | 2914771 | 3281079.75 | 3625669.75 | 4073020 | **1.38565158** | 0 |
| sp\|O4317 | O43175 | PHGDH | D-3-phosphoglycerate dehydrogenase OS=Homo sapien | 198960576 | 202764464 | 192986000 | 199772944 | 275807296 | 231960928 | 307121440 | 284870880 | **1.384245077** | 0 |
| sp\|P5774 | P57740 | NUP107 | Nuclear pore complex protein Nup107 OS=Homo sapien | 3543838 | 3552456.75 | 3779453 | 3410993.5 | 3403484.5 | 6706555 | 3277505 | 6377843 | **1.38347767** | 8 |
| sp\|Q9NRX | Q9NRX2 | MRPL17 | 39S ribosomal protein L17, mitochondrial OS=Homo sapi | 343497.88 | 990982 | 710041.81 | 411704.78 | 1075897.12 | 1044936.12 | 940093.62 | 335850.94 | **1.382925329** | 5 |
| sp\|Q9UQ8 | Q9UQ80 | PA2G4 | Proliferation-associated protein 2G4 OS=Homo sapiens ( | 79756992 | 86408152 | 100767232 | 84974864 | 120659104 | 122918304 | 121108024 | 121899416 | **1.382707693** | 0 |
| sp\|Q9NZN | Q9NZN8 | CNOT2 | CCR4-NOT transcription complex subunit 2 OS=Homo sa | 1096276.38 | 1044215.31 | 1214240.62 | 1013031.44 | 1395088.25 | 1712378 | 1577178.38 | 1351902.88 | **1.382068229** | 0 |
| sp\|Q9UI12 | Q9UI12 | ATP6V1H | V-type proton ATPase subunit H OS=Homo sapiens (Hum | 2843540.75 | 4331682.5 | 5418382.5 | 3672064.5 | 4222923.5 | 9656309 | 4333302 | 4266700.5 | **1.382004839** | 5 |
| sp\|Q1578 | Q15785 | TOMM34 | Mitochondrial import receptor subunit TOM34 OS=Hom | 7966060.5 | 8724782 | 8962588 | 8968626 | 11706906 | 13483886 | 13261305 | 9386197 | **1.381728841** | 0 |
| sp\|P15144 | P15144 | ANPEP | Aminopeptidase N OS=Homo sapiens (Human) GN=ANPI | 19059198 | 22329312 | 21983272 | 21648958 | 29938260 | 29151412 | 26173570 | 32176410 | **1.381305926** | 0 |
| sp\|Q8IYK4 | Q8IYK4 | COLGALT2 | Procollagen galactosyltransferase 2 OS=Homo sapiens (H | 561892.25 | 675148.31 | 477165.84 | 776721.31 | 730626.56 | 771238.44 | 980653.88 | 955099.06 | **1.380055281** | 2 |
| sp\|P09668 | P09668 | CTSH | Pro-cathepsin H OS=Homo sapiens (Human) GN=CTSH PI | 1433583 | 2467356.25 | 2716610.25 | 2685261 | 3451595.5 | 3404690.25 | 3179366.5 | 2795015.25 | **1.379224859** | 0 |
| sp\|P51654 | P51654 | GPC3 | Glypican-3 OS=Homo sapiens (Human) GN=GPC3 PE=1 S | 6722262 | 5891288 | 4580050 | 5783815.5 | 7917857.5 | 7497066 | 9524306 | 6741699.5 | **1.378785573** | 0 |
| sp\|P60520 | P60520 | GABARAPL | Gamma-aminobutyric acid receptor-associated protein-li | 658049.06 | 830941.25 | 813908.25 | 548705.31 | 1270009.62 | 1112017.5 | 837143.88 | 709123.19 | **1.377573593** | 2 |
| sp\|P42167 | P42167 | TMPO | Lamina-associated polypeptide 2, isoforms beta/gamma | 34142388 | 33062912 | 42072044 | 36841456 | 61069960 | 44207128 | 52046532 | 43899980 | **1.377123272** | 0 |
| sp\|P08133 | P08133 | ANXA6 | Annexin A6 OS=Homo sapiens (Human) GN=ANXA6 PE=1 | 86008304 | 92863456 | 84795432 | 90317192 | 124395208 | 124236216 | 110795816 | 127820936 | **1.376467997** | 0 |
| sp\|O9600 | O96000 | NDUFB10 | NADH dehydrogenase [ubiquinone] 1 beta subcomplex s | 6202604.5 | 5981266 | 6516674.5 | 7401767 | 8815490 | 8138219 | 10369040 | 8596374 | **1.376089712** | 0 |
| sp\|Q5XKP | Q5XKP0 | MIC13 | MICOS complex subunit MIC13 OS=Homo sapiens (Huma | 2238181.25 | 1913298.25 | 1932578.5 | 2290613 | 3176223.5 | 3139143.25 | 2721429.75 | 2486161.5 | **1.375929634** | 0 |
| sp\|P06748 | P06748 | NPM1 | Nucleophosmin OS=Homo sapiens (Human) GN=NPM1 F | 415553440 | 422217728 | 461098848 | 428747840 | 702365248 | 405378368 | 705981504 | 562906624 | **1.375669819** | 4 |
| sp\|Q9Y3C | Q9Y3C8 | UFC1 | Ubiquitin-fold modifier-conjugating enzyme 1 OS=Homo | 1996500.25 | 1310620.75 | 1602874.25 | 1468482.25 | 2332407.25 | 1954996.62 | 2476876.25 | 2010188.62 | **1.375636857** | 1 |
| sp\|P51688 | P51688 | SGSH | N-sulphoglucosamine sulphohydrolase OS=Homo sapien | 2867403.5 | 1427672.12 | 2798356 | 2418490.25 | 3046979.25 | 2513917.25 | 3445560.75 | 4077847 | **1.375568936** | 2 |
| sp\|Q9GZN | Q9GZM5 | YIPF3 | Protein YIPF3 OS=Homo sapiens (Human) GN=YIPF3 PE=: | 0 | 863785.5 | 0 | 841781.88 | 983160.06 | 803773.44 | 558918.25 | 0 | **1.375408429** | 7 |
| sp\|Q96HS | Q96HS1 | PGAM5 | Serine/threonine-protein phosphatase PGAM5, mitochor | 9892920 | 11287799 | 11027226 | 10108692 | 14059310 | 14470726 | 13605751 | 16037722 | **1.37471957** | 0 |
| sp\|Q96N6 | Q96N66 | MBOAT7 | Lysophospholipid acyltransferase 7 OS=Homo sapiens (H | 603376.88 | 483882.75 | 361957 | 561970.19 | 651273.19 | 814752.06 | 567121.44 | 727253.06 | **1.372522792** | 1 |
| sp\|Q1315 | Q13151 | HNRNPA0 | Heterogeneous nuclear ribonucleoprotein A0 OS=Homo | 42165424 | 39266072 | 44684912 | 44822208 | 60451608 | 54568620 | 63604172 | 55785656 | **1.371311302** | 0 |
| sp\|Q8IYB3 | Q8IYB3 | SRRM1 | Serine/arginine repetitive matrix protein 1 OS=Homo sap | 9693045 | 9064496 | 10616414 | 10703708 | 11391717 | 16232267 | 12971274 | 14358660 | **1.37118569** | 0 |
| sp\|Q96AT | Q96AT9 | RPE | Ribulose-phosphate 3-epimerase OS=Homo sapiens (Hu | 2359761.5 | 2236515.75 | 2943862.5 | 2759160 | 4112090.25 | 3297878.75 | 3661260 | 3035472 | **1.369675739** | 0 |
| sp\|P06132 | P06132 | UROD | Uroporphyrinogen decarboxylase OS=Homo sapiens (Hu | 5082925 | 4755820.5 | 5835302.5 | 5681442 | 8064632.5 | 8297822.5 | 5748521 | 7135229.5 | **1.369493535** | 1 |
| sp\|Q1350 | Q13501 | SQSTM1 | Sequestosome-1 OS=Homo sapiens (Human) GN=SQSTN | 3948403.5 | 1733711.75 | 3646567 | 1906511.5 | 3546480 | 5774792 | 3100062 | 2956534.5 | **1.368723036** | 6 |
| sp\|Q9Y60 | Q9Y606 | PUS1 | tRNA pseudouridine synthase A OS=Homo sapiens (Hum | 9707079 | 9329448 | 10009027 | 10272841 | 13397215 | 11717777 | 14654933 | 14000709 | **1.367569404** | 0 |
| sp\|O7553 | O75531 | BANF1 | Barrier-to-autointegration factor OS=Homo sapiens (Hur | 6829642 | 5690062 | 6224676 | 7216478.5 | 7995988.5 | 8331205 | 10107798 | 9007504 | **1.365228176** | 0 |
| sp\|P00450 | P00450 | CP | Ceruloplasmin OS=Homo sapiens (Human) GN=CP PE=1 | 831311.88 | 828580.06 | 650372.12 | 917666.31 | 914118.25 | 1591316.75 | 1096356.5 | 803169.75 | **1.364639489** | 4 |
| sp\|Q7Z2W | Q7Z2W9 | MRPL21 | 39S ribosomal protein L21, mitochondrial OS=Homo sapi | 2735948.25 | 2247600.5 | 2818501 | 2775438 | 3311041 | 3864689.5 | 3168407.5 | 4084356.5 | **1.364075747** | 0 |
| sp\|Q9UL2 | Q9UL25 | RAB21 | Ras-related protein Rab-21 OS=Homo sapiens (Human) ( | 9314678 | 8561700 | 10439938 | 10083515 | 13665850 | 12847552 | 11819294 | 14041627 | **1.363920664** | 0 |
| sp\|Q96EY1 | Q96EY1 | DNAJA3 | DnaJ homolog subfamily A member 3, mitochondrial OS | 3610034.5 | 2894777.25 | 3148798.25 | 2925716.25 | 4045928 | 4274512 | 4805444 | 4029543.25 | **1.363779499** | 0 |
| sp\|Q1296 | Q12965 | MYO1E | Unconventional myosin-Ie OS=Homo sapiens (Human) G | 2493367.25 | 2577306.5 | 2647669 | 1992156.75 | 3173568 | 3420800.5 | 3104782.25 | 3538823.75 | **1.363264011** | 0 |
| sp\|P50213 | P50213 | IDH3A | Isocitrate dehydrogenase [NAD] subunit alpha, mitochon | 11881054 | 13415401 | 14671418 | 11138573 | 18496016 | 15396768 | 16947140 | 18738980 | **1.361450648** | 0 |
| sp\|Q9994 | Q99943 | AGPAT1 | 1-acyl-sn-glycerol-3-phosphate acyltransferase alpha OS | 217340.8 | 209289.17 | 125487.64 | 210388.84 | 223054.86 | 281148.59 | 299589.91 | 234269.48 | **1.361382373** | 0 |
| sp\|Q8WU | Q8WUH6 | TMEM263 | Transmembrane protein 263 OS=Homo sapiens (Human) | 7417250 | 7096146 | 5774555.5 | 6300065.5 | 9473649 | 10288042 | 8257680 | 8171166 | **1.361159691** | 0 |
| sp\|Q1543 | Q15437 | SEC23B | Protein transport protein Sec23B OS=Homo sapiens (Hur | 13559665 | 14702277 | 13864416 | 13080466 | 16471295 | 17742018 | 19108120 | 21793594 | **1.360611272** | 0 |
| sp\|Q9Y2V | Q9Y2V7 | COG6 | Conserved oligomeric Golgi complex subunit 6 OS=Homo | 0 | 836956.38 | 1271358.88 | 0 | 0 | 0 | 1726601.88 | 1140377.12 | **1.359843594** | 7 |
| sp\|Q1290 | Q12904 | AIMP1 | Aminoacyl tRNA synthase complex-interacting multifunct | 12631445 | 12161428 | 14500692 | 14551496 | 18817400 | 15108638 | 20020402 | 19274392 | **1.35984305** | 0 |
| sp\|O1494 | O14949 | UQCRQ | Cytochrome b-c1 complex subunit 8 OS=Homo sapiens ( | 6349047 | 6956426 | 5836190.5 | 7222283 | 9578714 | 10162831 | 8542851 | 7547357 | **1.35911947** | 0 |
| sp\|P50395 | P50395 | GDI2 | Rab GDP dissociation inhibitor beta OS=Homo sapiens (H | 216459232 | 219885520 | 210203680 | 210633328 | 300015744 | 285467520 | 284870528 | 292699744 | **1.35683421** | 0 |
| sp\|P38571 | P38571 | LIPA | Lysosomal acid lipase/cholesteryl ester hydrolase OS=Hc | 821537.25 | 908021.38 | 945938.38 | 762020.25 | 1046441.25 | 1133022 | 1075467 | 1406419.88 | **1.35602232** | 0 |
| sp\|Q9UHV | Q9UHV9 | PFDN2 | Prefoldin subunit 2 OS=Homo sapiens (Human) GN=PFDI | 3388614 | 2605881.75 | 2703818.5 | 3279702.5 | 3947444.5 | 3723901 | 4632049.5 | 3936841.5 | **1.35583685** | 0 |

| | | | | | | | | | | | | | |
|---|---|---|---|---|---|---|---|---|---|---|---|---|---|
| sp\|P09960 | P09960 | LTA4H | Leukotriene A-4 hydrolase OS=Homo sapiens (Human) G | 21346784 | 24131276 | 24002864 | 22455864 | 34056604 | 33288840 | 29423828 | 27800088 | **1.354945748** | **0** |
| sp\|P40616 | P40616 | ARL1 | ADP-ribosylation factor-like protein 1 OS=Homo sapiens | 6112812 | 6895872 | 6244819 | 5978808.5 | 6793057 | 7969712 | 6576944.5 | 12812498 | **1.353511013** | **2** |
| sp\|Q9Y343 | Q9Y343 | SNX24 | Sorting nexin-24 OS=Homo sapiens (Human) GN=SNX24 | 0 | 0 | 457525.22 | 320984 | 376428.84 | 0 | 344039.62 | 333090.66 | **1.35330333** | **6** |
| sp\|P49840 | P49840 | GSK3A | Glycogen synthase kinase-3 alpha OS=Homo sapiens (Hu | 0 | 433550.59 | 839762.44 | 638962.25 | 732862.12 | 623691.25 | 670131.69 | 559504.69 | **1.352414988** | **6** |
| sp\|P62877 | P62877 | RBX1 | E3 ubiquitin-protein ligase RBX1 OS=Homo sapiens (Hum | 685473.81 | 980554.38 | 1104655.12 | 887898.69 | 1089928.75 | 947244.44 | 1277175.12 | 1627652.88 | **1.350796891** | **3** |
| sp\|Q9NVM | Q9NVM6 | DNAJC17 | DnaJ homolog subfamily C member 17 OS=Homo sapien | 276939.44 | 190859.44 | 125258.02 | 217099.73 | 293617.69 | 304077.97 | 283387 | 213128.14 | **1.350616362** | **2** |
| sp\|O43747 | O43747 | AP1G1 | AP-1 complex subunit gamma-1 OS=Homo sapiens (Hum | 7096557 | 6807302 | 5929597.5 | 6682516 | 6874221 | 9564929 | 8729581 | 10641532 | **1.350516674** | **1** |
| sp\|P61626 | P61626 | LYZ | Lysozyme C OS=Homo sapiens (Human) GN=LYZ PE=1 SV | 7651882 | 7052933.5 | 4426967 | 5068914.5 | 9522233 | 5271073.5 | 9761756 | 8104763.5 | **1.349540718** | **2** |
| sp\|Q9Y6K0 | Q9Y6K0 | CEPT1 | Choline/ethanolaminephosphotransferase 1 OS=Homo s | 2325181.75 | 126645.62 | 1975075.25 | 1931093.25 | 2065160.5 | 2235572 | 2428884.25 | 1848851.75 | **1.349240968** | **5** |
| sp\|Q8WUI | Q8WUD6 | CHPT1 | Cholinephosphotransferase 1 OS=Homo sapiens (Humar | 1042710 | 843819.81 | 1324554 | 1234711.5 | 1616274.62 | 1927941.88 | 1128982 | 1321910.38 | **1.348489632** | **3** |
| sp\|Q04446 | Q04446 | GBE1 | 1,4-alpha-glucan-branching enzyme OS=Homo sapiens (I | 7496458 | 6270968 | 5738433 | 6227648.5 | 9365803 | 10389026 | 7283700.5 | 7643861 | **1.347752167** | **1** |
| sp\|O75436 | O75436 | VPS26A | Vacuolar protein sorting-associated protein 26A OS=Hon | 11188403 | 9455300 | 6929857 | 10622895 | 12629244 | 15382062 | 9081100 | 14376838 | **1.347487457** | **3** |
| sp\|P16615 | P16615 | ATP2A2 | Sarcoplasmic/endoplasmic reticulum calcium ATPase 2 ( | 32457618 | 36322012 | 36996576 | 34091860 | 45824768 | 49476160 | 43097604 | 49984096 | **1.346859461** | **0** |
| sp\|Q13616 | Q13616 | CUL1 | Cullin-1 OS=Homo sapiens (Human) GN=CUL1 PE=1 SV=2 | 4323851.5 | 5800312 | 5896435 | 4166337 | 5327826 | 7053102 | 6830974.5 | 7969651 | **1.346492314** | **2** |
| sp\|O43464 | O43464 | HTRA2 | Serine protease HTRA2, mitochondrial OS=Homo sapien: | 4435929 | 4582425 | 5612426.5 | 4844095.5 | 6979369 | 5342196.5 | 6639651.5 | 7260133.5 | **1.346419382** | **1** |
| sp\|Q9UI09 | Q9UI09 | NDUFA12 | NADH dehydrogenase [ubiquinone] 1 alpha subcomplex | 2572973.25 | 2596984.75 | 2282497.5 | 2549420 | 3713710.5 | 3784874 | 3054938 | 2884575 | **1.343557766** | **0** |
| sp\|Q9BYC9 | Q9BYC9 | MRPL20 | 39S ribosomal protein L20, mitochondrial OS=Homo sapi | 2091984.12 | 2096567.25 | 2332316 | 2440057 | 2689212 | 2625434.25 | 2861407.25 | 3859581.25 | **1.343124242** | **0** |
| sp\|P53367 | P53367 | ARFIP1 | Arfaptin-1 OS=Homo sapiens (Human) GN=ARFIP1 PE=1 | 989645.19 | 889131.06 | 965712.56 | 1194815.88 | 1400655.88 | 1435234.25 | 1201382.12 | 1387718.5 | **1.34305064** | **0** |
| sp\|Q969X5 | Q969X5 | ERGIC1 | Endoplasmic reticulum-Golgi intermediate compartment | 6654550.5 | 6187795 | 5718768 | 6174081 | 9295401 | 8537838 | 8638871 | 6737609.5 | **1.34261** | **0** |
| sp\|Q6UX04 | Q6UX04 | CWC27 | Spliceosome-associated protein CWC27 homolog OS=Ho | 0 | 351179.72 | 345595.88 | 412589.97 | 368904.12 | 381081.66 | 341361.22 | 396870.34 | **1.34150309** | **6** |
| sp\|O75170 | O75170 | PPP6R2 | Serine/threonine-protein phosphatase 6 regulatory subu | 645276.75 | 699593.5 | 746771.88 | 677733.75 | 1085372.88 | 1135805.75 | 727263.38 | 764903.81 | **1.340860172** | **1** |
| sp\|Q9NRW | Q9NRW7 | VPS45 | Vacuolar protein sorting-associated protein 45 OS=Hom | 1292807.62 | 1269357.25 | 1249085.5 | 1592512.25 | 1788694.25 | 1667716.25 | 2127609 | 1660730.5 | **1.340686205** | **0** |
| sp\|O60888 | O60888 | CUTA | Protein CutA OS=Homo sapiens (Human) GN=CUTA PE=1 | 4017073 | 3223584.25 | 1384820.38 | 3453664.5 | 3360099.75 | 5010716 | 4140503.75 | 3659399 | **1.338730708** | **3** |
| sp\|O00767 | O00767 | SCD | Acyl-CoA desaturase OS=Homo sapiens (Human) GN=SCI | 5258854 | 4629091 | 6080896 | 4903906.5 | 7920727 | 6094361.5 | 7347658.5 | 6568060.5 | **1.338147146** | **0** |
| sp\|P11168 | P11168 | SLC2A2 | Solute carrier family 2, facilitated glucose transporter me | 273617.22 | 264269.22 | 371550.75 | 0 | 412362.16 | 455181.78 | 349213.69 | 0 | **1.337923766** | **4.5** |
| sp\|O14745 | O14745 | SLC9A3R1 | Na(+)/H(+) exchange regulatory cofactor NHE-RF1 OS=H | 44513528 | 31195696 | 41304728 | 37730308 | 55604864 | 48809260 | 57401336 | 45153776 | **1.337492169** | **0** |
| sp\|Q13425 | Q13425 | SNTB2 | Beta-2-syntrophin OS=Homo sapiens (Human) GN=SNTB | 608537.81 | 692239.06 | 719933 | 891174.12 | 874519.56 | 858261.69 | 1033048.75 | 1125575.38 | **1.336387505** | **2** |
| sp\|P31937 | P31937 | HIBADH | 3-hydroxyisobutyrate dehydrogenase, mitochondrial OS | 9801256 | 9578114 | 8594577 | 10795650 | 13035985 | 11101535 | 14750571 | 12886798 | **1.335450792** | **0** |
| sp\|Q86VR | Q86VR2 | RETREG3 | Reticulophagy regulator 3 OS=Homo sapiens (Human) GI | 1599361.5 | 1589425.25 | 1504677.38 | 2017023.25 | 2284993.25 | 2144469.5 | 2314506.25 | 2207787.75 | **1.33399502** | **0** |
| sp\|Q6P1A | Q6P1A2 | LPCAT3 | Lysophospholipid acyltransferase 5 OS=Homo sapiens (H | 1619436.5 | 1979788.38 | 1287395 | 1784661.62 | 2182495.25 | 2272145.75 | 1928027.75 | 2513261.25 | **1.333466441** | **1** |
| sp\|Q96CS3 | Q96CS3 | FAF2 | FAS-associated factor 2 OS=Homo sapiens (Human) GN= | 2614550 | 2422134.5 | 1829264.5 | 2372067.5 | 2842741 | 2948613 | 3130494.5 | 3361562.5 | **1.329658915** | **0** |
| sp\|P62826 | P62826 | RAN | GTP-binding nuclear protein Ran OS=Homo sapiens (Hur | 230448032 | 204550720 | 223576576 | 216060960 | 290129184 | 298384480 | 281046400 | 291618240 | **1.327612769** | **0** |
| sp\|Q96CW | Q96CW1 | AP2M1 | AP-2 complex subunit mu OS=Homo sapiens (Human) GI | 9864331 | 9877016 | 8921486 | 8550833 | 13274366 | 12486012 | 11506146 | 12126088 | **1.327270793** | **0** |
| sp\|Q9UHD | Q9UHD2 | TBK1 | Serine/threonine-protein kinase TBK1 OS=Homo sapiens | 354570 | 306131.91 | 302627.44 | 362108.47 | 398418.56 | 503274.09 | 456406.44 | 400968.22 | **1.327159436** | **0** |
| sp\|Q99614 | Q99614 | TTC1 | Tetratricopeptide repeat protein 1 OS=Homo sapiens (H | 720615.5 | 939930.62 | 615296.94 | 562209.69 | 971758.94 | 721473.19 | 939424.88 | 1133685.25 | **1.327086771** | **2** |
| sp\|Q8WW | Q8WWY3 | PRPF31 | U4/U6 small nuclear ribonucleoprotein Prp31 OS=Homo | 5285200 | 6628606.5 | 6168831 | 5740559 | 6736961.5 | 5675220 | 10602858 | 8587864 | **1.326560166** | **3** |
| sp\|O00483 | O00483 | NDUFA4 | Cytochrome c oxidase subunit NDUFA4 OS=Homo sapier | 8091506 | 10322780 | 7920328.5 | 8230276 | 9994936 | 9713537 | 12348257 | 13774988 | **1.325961614** | **2** |
| sp\|O60502 | O60502 | OGA | Protein O-GlcNAcase OS=Homo sapiens (Human) GN=OC | 447982.03 | 426887.69 | 346207.81 | 574536.5 | 652015 | 830956.75 | 430011.56 | 464035.44 | **1.323791589** | **3** |
| sp\|Q9BYD | Q9BYD2 | MRPL9 | 39S ribosomal protein L9, mitochondrial OS=Homo sapie | 1679800.75 | 1716490.38 | 1338328.75 | 2045807 | 2501154.5 | 2011591.25 | 2282746.5 | 2172082.75 | **1.322567909** | **1** |
| sp\|P54819 | P54819 | AK2 | Adenylate kinase 2, mitochondrial OS=Homo sapiens (Hu | 36430404 | 32843784 | 25536530 | 32560082 | 45009852 | 45346060 | 42291472 | 35550940 | **1.320540689** | **1** |
| sp\|P22033 | P22033 | MUT | Methylmalonyl-CoA mutase, mitochondrial OS=Homo sa | 7496874 | 5373805 | 5285133 | 6255826 | 8738047 | 6932192 | 8314650.5 | 8228268.5 | **1.319581996** | **1** |
| sp\|Q9NUU | Q9NUU7 | DDX19A | ATP-dependent RNA helicase DDX19A OS=Homo sapiens | 14733240 | 17252664 | 16876188 | 14976790 | 20285458 | 20215130 | 20214160 | 23513564 | **1.319388895** | **0** |
| sp\|Q96T76 | Q96T76 | MMS19 | MMS19 nucleotide excision repair protein homolog OS= | 8870047 | 9960656 | 9679920 | 9372018 | 11877437 | 12286408 | 12281421 | 13526791 | **1.31912812** | **0** |
| sp\|Q96MU | Q96MU7 | YTHDC1 | YTH domain-containing protein 1 OS=Homo sapiens (Hu | 0 | 0 | 625953.19 | 792465.75 | 373878.44 | 0 | 848598.25 | 648525.88 | **1.319076133** | **6** |
| sp\|P53004 | P53004 | BLVRA | Biliverdin reductase A OS=Homo sapiens (Human) GN=B | 22897946 | 24199648 | 26908252 | 24554896 | 33339356 | 31080572 | 32707812 | 32797074 | **1.318220737** | **0** |

| | | | | | | | | | | | | | |
|---|---|---|---|---|---|---|---|---|---|---|---|---|---|
| sp\|P42126 | P42126 | ECI1 | Enoyl-CoA delta isomerase 1, mitochondrial OS=Homo sa | 15690940 | 14275527 | 15699417 | 16643588 | 21220686 | 17607554 | 23391812 | 19894644 | **1.317852541** | 0 |
| sp\|P10412 | P10412 | HIST1H1E | Histone H1.4 OS=Homo sapiens (Human) GN=HIST1H1E | 569511424 | 568051520 | 520678976 | 601355328 | 775264960 | 752958656 | 722804096 | 725734592 | **1.317386232** | 0 |
| sp\|Q6YN1 | Q6YN16 | HSDL2 | Hydroxysteroid dehydrogenase-like protein 2 OS=Homo | 12779684 | 12381840 | 11206071 | 13464867 | 17149640 | 16001609 | 14618918 | 17865352 | **1.317123746** | 0 |
| sp\|O6094 | O60942 | RNGTT | mRNA-capping enzyme OS=Homo sapiens (Human) GN= | 0 | 397288.16 | 545967.56 | 550738.31 | 496035.31 | 473820.44 | 484081.84 | 511344.47 | **1.315455096** | 8 |
| sp\|Q9NPH | Q9NPH0 | ACP6 | Lysophosphatidic acid phosphatase type 6 OS=Homo sap | 136290.22 | 146758.27 | 79880.77 | 147967.14 | 112208.27 | 163712.31 | 215105.86 | 181020.47 | **1.315426983** | 3 |
| sp\|O1490 | O14907 | TAX1BP3 | Tax1-binding protein 3 OS=Homo sapiens (Human) GN=T | 1066874.25 | 1082373.62 | 1554506.5 | 1097530.75 | 1751273 | 1588132 | 1526789 | 1449045.38 | **1.315322715** | 2 |
| sp\|P58546 | P58546 | MTPN | Myotrophin OS=Homo sapiens (Human) GN=MTPN PE=1 | 6265316 | 5124500.5 | 5714359 | 5509402 | 8481394 | 6490522 | 8606362 | 6156579.5 | **1.314911694** | 1 |
| sp\|Q96EK | Q96EK5 | KIF1BP | KIF1-binding protein OS=Homo sapiens (Human) GN=KIF | 398358.31 | 0 | 133844.38 | 0 | 107311.12 | 122220.05 | 168442.31 | 301800.66 | **1.314863967** | 6 |
| sp\|Q9H85 | Q9H857 | NT5DC2 | 5'-nucleotidase domain-containing protein 2 OS=Homo s | 933081.62 | 595999.56 | 1342197.38 | 645897.5 | 1217879.25 | 1188524.62 | 1023490.12 | 1194407.75 | **1.314776872** | 4 |
| sp\|Q9UJY | Q9UJY5 | GGA1 | ADP-ribosylation factor-binding protein GGA1 OS=Homo | 122584.67 | 128218.05 | 73896.99 | 121958.98 | 124351.05 | 230646.38 | 116996.13 | 114255.1 | **1.312520439** | 7 |
| sp\|P36542 | P36542 | ATP5F1C | ATP synthase subunit gamma, mitochondrial OS=Homo s | 22309998 | 24922148 | 23767830 | 28342072 | 32819516 | 26474574 | 34925756 | 36160524 | **1.312438918** | 1 |
| sp\|P24941 | P24941 | CDK2 | Cyclin-dependent kinase 2 OS=Homo sapiens (Human) G | 5651302 | 4189590 | 4325925 | 5290206.5 | 6825259 | 6554134 | 5815171 | 6337786 | **1.312243365** | 0 |
| sp\|P31942 | P31942 | HNRNPH3 | Heterogeneous nuclear ribonucleoprotein H3 OS=Homo | 24644136 | 25924652 | 18868046 | 21423024 | 29449306 | 31035906 | 29877708 | 28838566 | **1.311926836** | 0 |
| sp\|P50750 | P50750 | CDK9 | Cyclin-dependent kinase 9 OS=Homo sapiens (Human) G | 577363.44 | 459509.47 | 616496.12 | 550891.88 | 605412.56 | 817338 | 716943.06 | 751368.06 | **1.31157871** | 1 |
| sp\|Q9Y3D | Q9Y3D0 | CIAO2B | Cytosolic iron-sulfur assembly component 2B OS=Homo | 4163097.25 | 4451932.5 | 1044197.12 | 3617063 | 5066865.5 | 4824302 | 4768226 | 2748245.5 | **1.311182504** | 3 |
| sp\|Q9H2F | Q9H2F3 | HSD3B7 | 3 beta-hydroxysteroid dehydrogenase type 7 OS=Homo | 1288000.62 | 363215.38 | 1270401 | 421369.97 | 303091.44 | 1340483.25 | 1635456.12 | 1101082.5 | **1.31023942** | 6 |
| sp\|P20700 | P20700 | LMNB1 | Lamin-B1 OS=Homo sapiens (Human) GN=LMNB1 PE=1 S | 148906272 | 151125888 | 182481248 | 146897632 | 215411840 | 191211888 | 206704656 | 210949632 | **1.309602094** | 0 |
| sp\|Q1361 | Q13618 | CUL3 | Cullin-3 OS=Homo sapiens (Human) GN=CUL3 PE=1 SV=2 | 4904431.5 | 4899566.5 | 4593638.5 | 4522530.5 | 4116906 | 7216071.5 | 6980794 | 6460147.5 | **1.309392195** | 4 |
| sp\|O7596 | O75964 | ATP5MG | ATP synthase subunit g, mitochondrial OS=Homo sapien | 14749620 | 13764063 | 22686900 | 15516664 | 22193316 | 17445084 | 27373652 | 20342078 | **1.309318563** | 3 |
| sp\|A1L0T0 | A1L0T0 | ILVBL | Acetolactate synthase-like protein OS=Homo sapiens (Hu | 14373565 | 14516934 | 10761800 | 15656532 | 18647332 | 17066120 | 18394806 | 18259266 | **1.308426208** | 0 |
| sp\|Q9981 | Q99816 | TSG101 | Tumor susceptibility gene 101 protein OS=Homo sapiens | 444065.12 | 460330.94 | 323633.59 | 469251.41 | 671538.56 | 598991.31 | 565014.81 | 384730.31 | **1.308136314** | 3 |
| sp\|Q32MZ | Q32MZ4 | LRRFIP1 | Leucine-rich repeat flightless-interacting protein 1 OS=H | 1601481.75 | 1320479.38 | 5307276 | 4147446.5 | 4033555.25 | 5576504 | 2735862.75 | 3844298.5 | **1.308122675** | 6 |
| sp\|Q9NX4 | Q9NX46 | ADPRHL2 | Poly(ADP-ribose) glycohydrolase ARH3 OS=Homo sapien | 536284 | 646286.69 | 1313758.38 | 536508.12 | 531663.94 | 1363328.12 | 703566.81 | 1368621.12 | **1.308075489** | 5 |
| sp\|Q0108 | Q01085 | TIAL1 | Nucleolysin TIAR OS=Homo sapiens (Human) GN=TIAL1 P | 13865965 | 15463923 | 15371048 | 16307637 | 20549376 | 16089553 | 22182596 | 20918860 | **1.307035734** | 1 |
| sp\|Q1340 | Q13404 | UBE2V1 | Ubiquitin-conjugating enzyme E2 variant 1 OS=Homo sap | 24019448 | 21642532 | 19142194 | 23766660 | 31319624 | 27968422 | 29616640 | 26841116 | **1.306816214** | 0 |
| sp\|P30613 | P30613 | PKLR | Pyruvate kinase PKLR OS=Homo sapiens (Human) GN=PK | 89313904 | 85101656 | 114177080 | 85141376 | 132616064 | 118195224 | 117640464 | 119924216 | **1.306747438** | 0 |
| sp\|P0C0S5 | P0C0S5 | H2AFZ | Histone H2A.Z OS=Homo sapiens (Human) GN=H2AFZ PE | 842149888 | 714997632 | 736696128 | 759962496 | 1119213952 | 908576704 | 1041382400 | 921236224 | **1.306700259** | 0 |
| sp\|Q08J23 | Q08J23 | NSUN2 | tRNA (cytosine(34)-C(5))-methyltransferase OS=Homo sa | 11105986 | 13658306 | 9258696 | 10572998 | 13535207 | 15034493 | 15156170 | 14544986 | **1.306639032** | 1 |
| sp\|Q4G14 | Q4G148 | GXYLT1 | Glucoside xylosyltransferase 1 OS=Homo sapiens (Huma | 574568.62 | 1167817.25 | 1278529.38 | 1179084.75 | 1211532.5 | 1305138.5 | 1333465.5 | 1635486 | **1.306100595** | 1 |
| sp\|Q641Q | Q641Q2 | WASHC2A | WASH complex subunit 2A OS=Homo sapiens (Human) G | 422381.25 | 363227.53 | 100975.17 | 147285.28 | 262470 | 446225.59 | 342867.16 | 298330.72 | **1.305671386** | 6 |
| sp\|Q6P2E | Q6P2E9 | EDC4 | Enhancer of mRNA-decapping protein 4 OS=Homo sapie | 3717769.5 | 3894645.5 | 3378134.25 | 2919476.25 | 4440826 | 4016545.25 | 4987735.5 | 4716768 | **1.305667969** | 0 |
| sp\|Q1518 | Q15185 | PTGES3 | Prostaglandin E synthase 3 OS=Homo sapiens (Human) G | 41286636 | 36122328 | 30162144 | 34215044 | 57879736 | 43316712 | 48281168 | 35572712 | **1.305136823** | 2 |
| sp\|Q96I59 | Q96I59 | NARS2 | Probable asparagine--tRNA ligase, mitochondrial OS=Ho | 818098.69 | 754848.31 | 740734.12 | 807949.44 | 823325.75 | 823717.94 | 872974.38 | 1553766.75 | **1.30501824** | 0 |
| sp\|Q8IXK0 | Q8IXK0 | PHC2 | Polyhomeotic-like protein 2 OS=Homo sapiens (Human) | 458310.72 | 435683.62 | 275031.22 | 438709.72 | 493034.78 | 431611.62 | 528979.88 | 644204.06 | **1.304835669** | 3 |
| sp\|O4382 | O43822 | CFAP410 | Cilia- and flagella-associated protein 410 OS=Homo sapi | 212514.03 | 236111.28 | 251194.7 | 0 | 198840.09 | 224912.2 | 233368.75 | 255884.59 | **1.304629215** | 7 |
| sp\|Q709C | Q709C8 | VPS13C | Vacuolar protein sorting-associated protein 13C OS=Hon | 662106.12 | 681036.81 | 362144.91 | 382634 | 726570.31 | 849905.5 | 703749.62 | 443579.56 | **1.304553139** | 2 |
| sp\|P16104 | P16104 | H2AFX | Histone H2AX OS=Homo sapiens (Human) GN=H2AFX PE | 848261504 | 720217344 | 739304640 | 763764864 | 1121646592 | 914438208 | 1044883392 | 923707328 | **1.303796998** | 0 |
| sp\|Q9HD2 | Q9HD20 | ATP13A1 | Manganese-transporting ATPase 13A1 OS=Homo sapien | 5681609.5 | 7728424 | 5736769.5 | 5752276 | 8851092 | 9228287 | 7252178 | 7131274 | **1.303776377** | 2 |
| sp\|P51572 | P51572 | BCAP31 | B-cell receptor-associated protein 31 OS=Homo sapiens | 31956712 | 36518936 | 36427704 | 29848416 | 57149216 | 36921812 | 44768956 | 36834404 | **1.303688928** | 0 |
| sp\|Q9GZU | Q9GZU7 | CTDSP1 | Carboxy-terminal domain RNA polymerase II polypeptide | 2157386.5 | 0 | 0 | 0 | 0 | 2810577.5 | 0 | 0 | **1.302769578** | 7.5 |
| sp\|Q8NBL | Q8NBL1 | POGLUT1 | Protein O-glucosyltransferase 1 OS=Homo sapiens (Hum | 927894.25 | 0 | 795687.31 | 1047216.19 | 798314.19 | 811820.31 | 1012386.19 | 986814.25 | **1.30263385** | 6 |
| sp\|Q1469 | Q14697 | GANAB | Neutral alpha-glucosidase AB OS=Homo sapiens (Human | 143505552 | 133004520 | 128150232 | 136424256 | 175616864 | 174407392 | 177802544 | 176969424 | **1.302562069** | 0 |
| sp\|Q9UEW | Q9UEW8 | STK39 | STE20/SPS1-related proline-alanine-rich protein kinase C | 1621950.38 | 1337801.88 | 1821317.88 | 1834374.75 | 2051202.38 | 2184761 | 1931806.25 | 2447069.5 | **1.302231259** | 0 |
| sp\|P53990 | P53990 | IST1 | IST1 homolog OS=Homo sapiens (Human) GN=IST1 PE=1 | 28407802 | 23011004 | 25660994 | 25922540 | 39677392 | 30674376 | 32592248 | 31172472 | **1.302072244** | 0 |
| sp\|Q9255 | Q92556 | ELMO1 | Engulfment and cell motility protein 1 OS=Homo sapiens | 465232.91 | 577857.81 | 559467.44 | 456638.91 | 700223.88 | 678535.44 | 772788.5 | 529675.38 | **1.302072171** | 2 |

| | | | | | | | | | | | | | |
|---|---|---|---|---|---|---|---|---|---|---|---|---|---|
| sp\|P48506 | P48506 | GCLC | Glutamate--cysteine ligase catalytic subunit OS=Homo sa | 7333800.5 | 8839859 | 8544564 | 8247635 | 11446434 | 9709019 | 10599565 | 11152449 | **1.301572868** | 0 |
| sp\|P43307 | P43307 | SSR1 | Translocon-associated protein subunit alpha OS=Homo s | 18333720 | 21200536 | 20534094 | 21060654 | 28222134 | 25184340 | 28562158 | 23612910 | **1.301403158** | 0 |
| sp\|Q9H0E9 | Q9H0E9 | BRD8 | Bromodomain-containing protein 8 OS=Homo sapiens (H | 296433.94 | 279035.44 | 0 | 0 | 0 | 0 | 480407.59 | 268347.03 | **1.301119827** | 8 |
| sp\|O7543! | O75439 | PMPCB | Mitochondrial-processing peptidase subunit beta OS=Hc | 9778188 | 16462182 | 22406326 | 18763324 | 22447378 | 19429378 | 24411202 | 21416434 | **1.301058685** | 2 |
| sp\|O00217 | O00217 | NDUFS8 | NADH dehydrogenase [ubiquinone] iron-sulfur protein 8, | 4200533.5 | 3705335 | 3672158.25 | 4506819.5 | 4941272.5 | 5387648 | 5186812 | 5399470 | **1.300304782** | 0 |
| sp\|P41252 | P41252 | IARS | Isoleucine--tRNA ligase, cytoplasmic OS=Homo sapiens ( | 38679096 | 43695528 | 39192004 | 37875672 | 49708632 | 47431480 | 52021704 | 58112540 | **1.299996024** | 0 |
| sp\|Q8IXI2\| | Q8IXI2 | RHOT1 | Mitochondrial Rho GTPase 1 OS=Homo sapiens (Human) | 2074759.75 | 1883468.12 | 1976067.12 | 1796995.5 | 2478008.25 | 2613581.25 | 2339084.25 | 2615793.25 | **1.299455377** | 0 |
| sp\|Q9UDV | Q9UDW1 | UQCR10 | Cytochrome b-c1 complex subunit 9 OS=Homo sapiens ( | 1607792.62 | 1802429.75 | 1860083 | 1781332.88 | 1786560.62 | 1967323.88 | 2067414.88 | 3338477.75 | **1.298957321** | 2 |
| sp\|P28331 | P28331 | NDUFS1 | NADH-ubiquinone oxidoreductase 75 kDa subunit, mitoc | 40733720 | 41429180 | 37570796 | 43849112 | 53291216 | 50200156 | 55955044 | 52995912 | **1.29868371** | 0 |
| sp\|P31150 | P31150 | GDI1 | Rab GDP dissociation inhibitor alpha OS=Homo sapiens ( | 62670500 | 57810996 | 51608520 | 59836856 | 75709656 | 76318208 | 72710648 | 76314936 | **1.298053328** | 0 |
| sp\|P56192 | P56192 | MARS | Methionine--tRNA ligase, cytoplasmic OS=Homo sapiens | 59678720 | 70626408 | 43758408 | 58332528 | 75994776 | 73446104 | 78289120 | 73908896 | **1.297951828** | 0 |
| sp\|P25815 | P25815 | S100P | Protein S100-P OS=Homo sapiens (Human) GN=S100P PE | 34782816 | 44806404 | 47876704 | 39572152 | 67368456 | 43409800 | 50615976 | 55405820 | **1.297907981** | 2 |
| sp\|Q1280( | Q12800 | TFCP2 | Alpha-globin transcription factor CP2 OS=Homo sapiens | 731487.31 | 918629.25 | 711896.75 | 621663.81 | 1092125 | 843493.06 | 909698.94 | 1025684.38 | **1.297392856** | 2 |
| sp\|O15347 | O15347 | HMGB3 | High mobility group protein B3 OS=Homo sapiens (Huma | 9933856 | 11773659 | 13093772 | 18274676 | 19872128 | 16173190 | 15850338 | 16954084 | **1.297192479** | 3 |
| sp\|Q9BW9 | Q9BW91 | NUDT9 | ADP-ribose pyrophosphatase, mitochondrial OS=Homo s | 0 | 188232.39 | 0 | 214445.23 | 183368.89 | 0 | 194385.03 | 144494.48 | **1.296939224** | 8 |
| sp\|O95816 | O95816 | BAG2 | BAG family molecular chaperone regulator 2 OS=Homo s | 12920111 | 17312298 | 13275890 | 12498902 | 17870804 | 17085350 | 18117320 | 19516742 | **1.296087194** | 1 |
| sp\|Q5T1J5 | Q5T1J5 | CHCHD2P9 | Putative coiled-coil-helix-coiled-coil-helix domain-contai | 1289571.62 | 1118430.75 | 1606251.12 | 1562247.75 | 1525880.38 | 2461015.25 | 1585319.62 | 1653860.75 | **1.295808194** | 3 |
| sp\|P30044 | P30044 | PRDX5 | Peroxiredoxin-5, mitochondrial OS=Homo sapiens (Hum | 45520564 | 47042628 | 38367032 | 48437172 | 59500572 | 51287428 | 63639932 | 57945520 | **1.295516672** | 0 |
| sp\|O43252 | O43252 | PAPSS1 | Bifunctional 3'-phosphoadenosine 5'-phosphosulfate syn | 12916535 | 15930569 | 13038682 | 14919048 | 19478318 | 18079264 | 17867476 | 18103606 | **1.294408571** | 0 |
| sp\|P56962 | P56962 | STX17 | Syntaxin-17 OS=Homo sapiens (Human) GN=STX17 PE=1 | 94536.37 | 81545.79 | 86813.64 | 138300.55 | 92659.54 | 121647.42 | 136595.28 | 168370.2 | **1.294309981** | 4 |
| sp\|Q7L014 | Q7L014 | DDX46 | Probable ATP-dependent RNA helicase DDX46 OS=Homo | 24877144 | 26479030 | 32606164 | 26167158 | 39543364 | 32645240 | 35565688 | 34722592 | **1.293721384** | 0 |
| sp\|O7574( | O75746 | SLC25A12 | Calcium-binding mitochondrial carrier protein Aralar1 O | 17176160 | 18838884 | 19840604 | 18048188 | 23613772 | 23012948 | 24100220 | 24804080 | **1.292639532** | 0 |
| sp\|Q6ZXV! | Q6ZXV5 | TMTC3 | Transmembrane and TPR repeat-containing protein 3 O | 6448025.5 | 4789261 | 3838831.25 | 5073435.5 | 6858413 | 6412723 | 7516092.5 | 5258016 | **1.292596624** | 2 |
| sp\|P56381 | P56381 | ATP5F1E | ATP synthase subunit epsilon, mitochondrial OS=Homo s | 12325034 | 7966265.5 | 17263166 | 12669174 | 16283686 | 13555884 | 17271958 | 17801712 | **1.292483791** | 2 |
| sp\|Q1679! | Q16795 | NDUFA9 | NADH dehydrogenase [ubiquinone] 1 alpha subcomplex | 16674725 | 15877859 | 15212265 | 17259446 | 19963030 | 22409640 | 18968164 | 22675122 | **1.292070233** | 0 |
| sp\|P06899 | P06899 | HIST1H2BJ | Histone H2B type 1-J OS=Homo sapiens (Human) GN=HIS | 254359968 | 198947504 | 224668192 | 322517568 | 419981312 | 425521984 | 240772576 | 206426416 | **1.292065** | 5 |
| sp\|P01011 | P01011 | SERPINA3 | Alpha-1-antichymotrypsin OS=Homo sapiens (Human) G | 8112178.5 | 11603072 | 7853590 | 9745818 | 9608026 | 10998485 | 14869723 | 12736581 | **1.292061001** | 3 |
| sp\|P22059 | P22059 | OSBP | Oxysterol-binding protein 1 OS=Homo sapiens (Human) | 4428136.5 | 4149540.5 | 4699438.5 | 4676399 | 6230797 | 5476641 | 6593493.5 | 4891114.5 | **1.291783066** | 0 |
| sp\|P62495 | P62495 | ETF1 | Eukaryotic peptide chain release factor subunit 1 OS=Ho | 21748030 | 24148614 | 22600344 | 24683106 | 29114160 | 28658476 | 27647946 | 34938520 | **1.291682556** | 0 |
| sp\|Q9BUT | Q9BUT1 | BDH2 | 3-hydroxybutyrate dehydrogenase type 2 OS=Homo sap | 298941.22 | 254413.39 | 226261.7 | 326427.62 | 367226.75 | 332242.25 | 361409.62 | 367642.53 | **1.291559143** | 0 |
| sp\|P28838 | P28838 | LAP3 | Cytosol aminopeptidase OS=Homo sapiens (Human) GN: | 64145280 | 67801856 | 67454032 | 64970452 | 91025296 | 79224960 | 82408528 | 88741464 | **1.291364966** | 0 |
| sp\|Q86W4 | Q86W42 | THOC6 | THO complex subunit 6 homolog OS=Homo sapiens (Hur | 4637620 | 4281391 | 3321667 | 4955382 | 3983493.25 | 5600170 | 6064445.5 | 6539273 | **1.290259615** | 3 |
| sp\|O1477( | O14776 | TCERG1 | Transcription elongation regulator 1 OS=Homo sapiens ( | 8145928.5 | 8794561 | 9059403 | 8369316.5 | 10193344 | 9910944 | 12333183 | 11881872 | **1.289507216** | 0 |
| sp\|O6092! | O60925 | PFDN1 | Prefoldin subunit 1 OS=Homo sapiens (Human) GN=PFDI | 8580058 | 6190438.5 | 0 | 0 | 6987005.5 | 7258212 | 0 | 4786521 | **1.288496869** | 7 |
| sp\|O95831 | O95831 | AIFM1 | Apoptosis-inducing factor 1, mitochondrial OS=Homo sa | 35942488 | 37525964 | 38230532 | 28592668 | 45104144 | 42762732 | 45818904 | 47031848 | **1.288156675** | 0 |
| sp\|Q9C00! | Q9C005 | DPY30 | Protein dpy-30 homolog OS=Homo sapiens (Human) GN: | 572940 | 416174.94 | 360514.19 | 512483.47 | 602827.5 | 611082.06 | 647600.88 | 535761.19 | **1.287393485** | 1 |
| sp\|Q9HAT | Q9HAT2 | SIAE | Sialate O-acetylesterase OS=Homo sapiens (Human) GN: | 152545.47 | 155139.27 | 55701.79 | 169682.27 | 0 | 258051.88 | 200531.27 | 227550.55 | **1.287139108** | 4 |
| sp\|Q9UM\ | Q9UMY4 | SNX12 | Sorting nexin-12 OS=Homo sapiens (Human) GN=SNX12 | 2805950.75 | 2209487.75 | 2026090.75 | 1820104.25 | 3159983 | 2335491.5 | 3222216.5 | 2684480.25 | **1.286689553** | 2 |
| sp\|Q96I99 | Q96I99 | SUCLG2 | Succinate--CoA ligase [GDP-forming] subunit beta, mitocl | 25124834 | 19562888 | 20468380 | 18163888 | 26197968 | 26991906 | 24008976 | 29848380 | **1.284772478** | 1 |
| sp\|Q0706! | Q07065 | CKAP4 | Cytoskeleton-associated protein 4 OS=Homo sapiens (Hu | 148461776 | 154504128 | 168087696 | 160808368 | 210368992 | 188375360 | 212265920 | 200233488 | **1.283893953** | 0 |
| sp\|P53007 | P53007 | SLC25A1 | Tricarboxylate transport protein, mitochondrial OS=Hom | 26334480 | 22195102 | 26868518 | 28454152 | 37706676 | 34234656 | 29763556 | 31622458 | **1.283817572** | 0 |
| sp\|O1488( | O14880 | MGST3 | Microsomal glutathione S-transferase 3 OS=Homo sapie | 3152441 | 2770847 | 2055294.12 | 2205308 | 2534092.25 | 3511903.75 | 3268845.5 | 3755057.75 | **1.283389657** | 2 |
| sp\|Q9Y4P1 | Q9Y4P1 | ATG4B | Cysteine protease ATG4B OS=Homo sapiens (Human) GN | 1297163.62 | 1268291.38 | 1857852.75 | 1826086.38 | 1986430.75 | 1671333 | 1734943.5 | 2624284.5 | **1.282843038** | 4 |
| sp\|Q8IX01 | Q8IX01 | SUGP2 | SURP and G-patch domain-containing protein 2 OS=Hom | 4882129 | 5144986 | 6084655.5 | 17685692 | 6126114 | 5008381 | 5773792 | 26428600 | **1.282252684** | 6 |
| sp\|Q1542! | Q15428 | SF3A2 | Splicing factor 3A subunit 2 OS=Homo sapiens (Human) ( | 2257163 | 1776010.25 | 1615933.62 | 1876118.75 | 2557949.75 | 2149725.25 | 2689692.5 | 2251141.75 | **1.282155478** | 2 |

| | | | | | | | | | | | | | |
|---|---|---|---|---|---|---|---|---|---|---|---|---|---|
| sp\|P41223 | P41223 | BUD31 | Protein BUD31 homolog OS=Homo sapiens (Human) GN | 4251648 | 3675163.5 | 2065226.5 | 3009283.25 | 3598780 | 4241539 | 4812002 | 4014217.5 | **1.281911137** | 4 |
| sp\|Q0280 | Q02809 | PLOD1 | Procollagen-lysine,2-oxoglutarate 5-dioxygenase 1 OS=H | 24876236 | 20116788 | 20644718 | 24064850 | 27848100 | 35154828 | 28125122 | 23839362 | **1.281650947** | 2 |
| sp\|Q1305 | Q13057 | COASY | Bifunctional coenzyme A synthase OS=Homo sapiens (Hu | 13194396 | 13194487 | 12855211 | 14453722 | 17230538 | 17552296 | 18028916 | 15964437 | **1.280800452** | 0 |
| sp\|Q9ULW | Q9ULW0 | TPX2 | Targeting protein for Xklp2 OS=Homo sapiens (Human) ( | 625323.56 | 0 | 2856462.5 | 1457825.25 | 2019548.5 | 2320946 | 777617.38 | 1205022 | **1.280087335** | 6 |
| sp\|Q1293 | Q12931 | TRAP1 | Heat shock protein 75 kDa, mitochondrial OS=Homo sap | 53268208 | 52337484 | 53497440 | 52688888 | 61649764 | 70591616 | 62680412 | 75992040 | **1.279150329** | 0 |
| sp\|Q1583 | Q15833 | STXBP2 | Syntaxin-binding protein 2 OS=Homo sapiens (Human) G | 2696549 | 3601589.5 | 3757025.25 | 3832206 | 4797041.5 | 4976864 | 3345105.75 | 4636413 | **1.278530389** | 3 |
| sp\|P13646 | P13646 | KRT13 | Keratin, type I cytoskeletal 13 OS=Homo sapiens (Humar | 53446536 | 130927840 | 77422584 | 80455472 | 101926232 | 106630680 | 118891912 | 110105224 | **1.278454167** | 4 |
| sp\|O4381 | O43819 | SCO2 | Protein SCO2 homolog, mitochondrial OS=Homo sapiens | 439764.91 | 398008.69 | 578938.12 | 406407.56 | 633019.75 | 636480.75 | 505039.41 | 556175.44 | **1.278421755** | 2 |
| sp\|Q1504 | Q15046 | KARS | Lysine--tRNA ligase OS=Homo sapiens (Human) GN=KAR | 41277540 | 41748816 | 42443636 | 38665196 | 55852904 | 55255680 | 48852060 | 49702924 | **1.277383421** | 0 |
| sp\|P68402 | P68402 | PAFAH1B2 | Platelet-activating factor acetylhydrolase IB subunit beta | 11619648 | 10041531 | 9060387 | 11798618 | 13828052 | 13473965 | 14408582 | 12564607 | **1.276457458** | 0 |
| sp\|P36915 | P36915 | GNL1 | Guanine nucleotide-binding protein-like 1 OS=Homo sap | 4148715 | 4484045 | 4984134.5 | 4822090 | 6380587 | 5743925.5 | 5931466.5 | 5479091 | **1.276375605** | 0 |
| sp\|P09234 | P09234 | SNRPC | U1 small nuclear ribonucleoprotein C OS=Homo sapiens | 1140536.25 | 1209242.62 | 403800.28 | 855823.94 | 1242873.88 | 1099508.75 | 1302698.38 | 960729.69 | **1.276058834** | 4 |
| sp\|Q1581 | Q15819 | UBE2V2 | Ubiquitin-conjugating enzyme E2 variant 2 OS=Homo sa | 11078792 | 10361060 | 9897506 | 11515078 | 12254796 | 13732838 | 14782181 | 13908023 | **1.275956354** | 0 |
| sp\|Q96P1 | Q96P16 | RPRD1A | Regulation of nuclear pre-mRNA domain-containing prot | 3288460.75 | 2596212 | 1435141 | 3625209.75 | 3136673 | 3064908.25 | 3555959.75 | 4204526 | **1.275654365** | 5 |
| sp\|P48047 | P48047 | ATP5PO | ATP synthase subunit O, mitochondrial OS=Homo sapien | 66914960 | 73306960 | 75159064 | 65780644 | 97256336 | 81875984 | 94055528 | 85469696 | **1.27562764** | 0 |
| sp\|P36543 | P36543 | ATP6V1E1 | V-type proton ATPase subunit E 1 OS=Homo sapiens (Hu | 1437019.5 | 1518212.88 | 1313194.62 | 1439488.62 | 1721203.25 | 1896175.88 | 1838896.88 | 1824314.75 | **1.275525296** | 0 |
| sp\|Q6UW | Q6UW02 | CYP20A1 | Cytochrome P450 20A1 OS=Homo sapiens (Human) GN= | 2235313.25 | 2251518.5 | 4140500.5 | 2616659 | 3352746.5 | 3396645 | 3072734 | 4517194.5 | **1.275287367** | 3 |
| sp\|O7560 | O75608 | LYPLA1 | Acyl-protein thioesterase 1 OS=Homo sapiens (Human) ( | 5467973 | 7432145.5 | 4897119 | 5257599 | 7132890 | 6899543 | 7158070 | 8205262 | **1.275036802** | 3 |
| sp\|Q9UQE | Q9UQE7 | SMC3 | Structural maintenance of chromosomes protein 3 OS=H | 8241093.5 | 10568805 | 9946182 | 6511237 | 11454215 | 10400358 | 12007665 | 11103219 | **1.274989429** | 1 |
| sp\|P09429 | P09429 | HMGB1 | High mobility group protein B1 OS=Homo sapiens (Huma | 68083952 | 52455716 | 64548016 | 76043304 | 95113024 | 72368976 | 63534448 | 101809304 | **1.274554792** | 4 |
| sp\|P49821 | P49821 | NDUFV1 | NADH dehydrogenase [ubiquinone] flavoprotein 1, mitoc | 14784548 | 13192344 | 12749977 | 14753078 | 16834936 | 15302730 | 17789952 | 20744318 | **1.27382847** | 0 |
| sp\|Q969M | Q969M3 | YIPF5 | Protein YIPF5 OS=Homo sapiens (Human) GN=YIPF5 PE= | 2798366.75 | 3136271 | 3071605.5 | 3376921.5 | 4253095 | 4662116 | 4163956.5 | 2691775.75 | **1.273579377** | 4 |
| sp\|P40937 | P40937 | RFC5 | Replication factor C subunit 5 OS=Homo sapiens (Humar | 1797699.75 | 1411393.12 | 2035365.75 | 1955355.75 | 2260608.75 | 2467614.25 | 1766684.75 | 2671212.5 | **1.27310508** | 3 |
| sp\|Q96F8 | Q96F86 | EDC3 | Enhancer of mRNA-decapping protein 3 OS=Homo sapie | 693906.06 | 658163.06 | 465176.53 | 0 | 694239.62 | 476028.31 | 706178.5 | 436591.44 | **1.272826197** | 5 |
| sp\|P61077 | P61077 | UBE2D3 | Ubiquitin-conjugating enzyme E2 D3 OS=Homo sapiens ( | 12837838 | 11219744 | 8977066 | 13655275 | 14789621 | 17213938 | 14190943 | 13179634 | **1.271669178** | 1 |
| sp\|Q9NXA | Q9NXA8 | SIRT5 | NAD-dependent protein deacylase sirtuin-5, mitochondri | 731822.88 | 777417.81 | 887937.88 | 604315.38 | 896554.75 | 1056603.5 | 1086840 | 774762 | **1.270953836** | 2 |
| sp\|Q8WX | Q8WXA9 | SREK1 | Splicing regulatory glutamine/lysine-rich protein 1 OS=H | 997668.25 | 759142 | 585613.31 | 1007797.12 | 1236264.75 | 1323322.62 | 796460.5 | 901160.81 | **1.270724853** | 4 |
| sp\|P09661 | P09661 | SNRPA1 | U2 small nuclear ribonucleoprotein A' OS=Homo sapiens | 27312154 | 22973254 | 20890048 | 25434548 | 31402876 | 29514154 | 32041400 | 29741530 | **1.270054393** | 0 |
| sp\|Q9965 | Q99653 | CHP1 | Calcineurin B homologous protein 1 OS=Homo sapiens (I | 1703399 | 2060660.25 | 2477264.25 | 1851928.38 | 2058742.25 | 3951471.5 | 2259816.25 | 2004377.25 | **1.269502964** | 5 |
| sp\|P16403 | P16403 | HIST1H1C | Histone H1.2 OS=Homo sapiens (Human) GN=HIST1H1C | 620685952 | 612317760 | 518462720 | 650053056 | 768693376 | 832754240 | 780333120 | 666430080 | **1.269284231** | 0 |
| sp\|Q9C0E | Q9C0E8 | LNPK | Endoplasmic reticulum junction formation protein lunap | 2458851 | 2136107 | 1144234 | 935274.06 | 2347481.75 | 2332399.5 | 1972556.75 | 1815562.38 | **1.268715775** | 6 |
| sp\|Q9282 | Q92820 | GGH | Gamma-glutamyl hydrolase OS=Homo sapiens (Human) | 22862852 | 23088208 | 17769094 | 29311104 | 21635704 | 32979710 | 31520696 | 31840468 | **1.268139124** | 3 |
| sp\|P04908 | P04908 | HIST1H2AB | Histone H2A type 1-B/E OS=Homo sapiens (Human) GN= | 1073596800 | 898954112 | 880794048 | 1006591232 | 1340897024 | 1119082496 | 1286850432 | 1147200000 | **1.267904366** | 0 |
| sp\|P61421 | P61421 | ATP6V0D1 | V-type proton ATPase subunit d 1 OS=Homo sapiens (Hu | 3925229.5 | 3855199.5 | 4451782.5 | 1922040.25 | 5532857 | 4993632.5 | 3309648.75 | 4094971.5 | **1.266835582** | 4 |
| sp\|Q8N39 | Q8N392 | ARHGAP18 | Rho GTPase-activating protein 18 OS=Homo sapiens (Hu | 17250428 | 12851552 | 19115786 | 15364913 | 21647528 | 19214842 | 19795952 | 21124314 | **1.266324613** | 0 |
| sp\|Q9UNX | Q9UNX4 | WDR3 | WD repeat-containing protein 3 OS=Homo sapiens (Hum | 2083573.62 | 3162951.75 | 2631204.5 | 2693181.25 | 2238578.5 | 3674404 | 3056697.5 | 4414260 | **1.266110352** | 4 |
| sp\|P06865 | P06865 | HEXA | Beta-hexosaminidase subunit alpha OS=Homo sapiens (H | 1007938.5 | 1229050.38 | 1127148.88 | 1396583.25 | 1087879.25 | 1389855 | 1506801.38 | 2041138.75 | **1.265706259** | 4 |
| sp\|P62310 | P62310 | LSM3 | U6 snRNA-associated Sm-like protein LSm3 OS=Homo sa | 8378324.5 | 8772155 | 9456188 | 8125636 | 9583788 | 9734077 | 10219012 | 14389092 | **1.264700713** | 0 |
| sp\|Q1425 | Q14257 | RCN2 | Reticulocalbin-2 OS=Homo sapiens (Human) GN=RCN2 P | 6905855.5 | 5926020 | 6370341.5 | 6988528 | 7763866.5 | 8557701 | 7903845.5 | 8885071 | **1.264205505** | 0 |
| sp\|Q9NW | Q9NWU5 | MRPL22 | 39S ribosomal protein L22, mitochondrial OS=Homo sapi | 3715823.5 | 4223615 | 2801339.75 | 3531740.25 | 4662381 | 5035386 | 4466211 | 3868727.75 | **1.263456464** | 1 |
| sp\|Q1479 | Q14790 | CASP8 | Caspase-8 OS=Homo sapiens (Human) GN=CASP8 PE=1 S | 3212086.25 | 691133.06 | 3017810.5 | 1067682 | 2864215.5 | 2342560.5 | 2543117.5 | 2340959.5 | **1.263138944** | 8 |
| sp\|P0C0S8 | P0C0S8 | HIST1H2AG | Histone H2A type 1 OS=Homo sapiens (Human) GN=HIST | 1099294208 | 916623552 | 891958848 | 1030256768 | 1371336960 | 1133165824 | 1308496896 | 1158096128 | **1.262297473** | 0 |
| sp\|Q1677 | Q16777 | HIST2H2AC | Histone H2A type 2-C OS=Homo sapiens (Human) GN=HI | 1098257152 | 915158656 | 891369088 | 1029092480 | 1370009088 | 1130922240 | 1307078400 | 1157160576 | **1.262156857** | 0 |
| sp\|Q9Y58 | Q9Y584 | TIMM22 | Mitochondrial import inner membrane translocase subu | 1053401.62 | 1091181.38 | 1256617.38 | 1074700 | 1315394.25 | 1381191.38 | 1457552.88 | 1493763.25 | **1.261847066** | 0 |
| sp\|P49959 | P49959 | MRE11 | Double-strand break repair protein MRE11 OS=Homo sa | 5352527 | 4964854.5 | 6919632.5 | 6350713 | 6000121.5 | 8102221 | 7407374 | 8253485.5 | **1.261808821** | 2 |

| | | | | | | | | | | | | |
|---|---|---|---|---|---|---|---|---|---|---|---|---|
| sp\|P26572 | P26572 | MGAT1 | Alpha-1,3-mannosyl-glycoprotein 2-beta-N-acetylglucosa | 533862.75 | 484649.75 | 423432.91 | 716826.81 | 664671.25 | 728923.75 | 586107.81 | 740596.56 | **1.260114126** | 2 |
| sp\|Q1425 | Q14254 | FLOT2 | Flotillin-2 OS=Homo sapiens (Human) GN=FLOT2 PE=1 SV | 7588033 | 7427005 | 6205415 | 7077487.5 | 10611193 | 10016182 | 6806884 | 8219282.5 | **1.259934146** | 3 |
| sp\|P63010 | P63010 | AP2B1 | AP-2 complex subunit beta OS=Homo sapiens (Human) ( | 39703472 | 39754292 | 44184032 | 40814384 | 49340164 | 52621704 | 50375864 | 54773712 | **1.259371609** | 0 |
| sp\|Q3B7T | Q3B7T1 | EDRF1 | Erythroid differentiation-related factor 1 OS=Homo sapi | 115478.97 | 133061.23 | 156667.44 | 152163.91 | 147377.59 | 147191.11 | 197186.48 | 209954.56 | **1.258962249** | 4 |
| sp\|Q9954 | Q99543 | DNAJC2 | DnaJ homolog subfamily C member 2 OS=Homo sapiens | 1278399.25 | 1205292.5 | 543254.31 | 1306580.38 | 1205954.5 | 1541947.75 | 1432661.25 | 1272928 | **1.258441958** | 4 |
| sp\|Q4VC3 | Q4VC31 | CCDC58 | Coiled-coil domain-containing protein 58 OS=Homo sapi | 2711831.5 | 2481887.5 | 2271855.25 | 2912039 | 2886060.75 | 3477363.75 | 3152465 | 3541420.75 | **1.258219008** | 1 |
| sp\|P06454 | P06454 | PTMA | Prothymosin alpha OS=Homo sapiens (Human) GN=PTM | 43485336 | 41946568 | 63402456 | 39266112 | 65583568 | 48735332 | 70589992 | 51352044 | **1.256035849** | 2 |
| sp\|Q96FC | Q96FC7 | PHYHIPL | Phytanoyl-CoA hydroxylase-interacting protein-like OS=I | 6897138 | 6380043 | 6447899 | 4604411.5 | 8682030 | 7614487 | 7243013 | 7016392 | **1.255921111** | 0 |
| sp\|A4D1E | A4D1E9 | GTPBP10 | GTP-binding protein 10 OS=Homo sapiens (Human) GN= | 1243119.5 | 1195320.5 | 1293183.88 | 1929171 | 1572829.62 | 1680528.12 | 1864327.25 | 1983963 | **1.254531941** | 3 |
| sp\|Q9UI3 | Q9UI30 | TRMT112 | Multifunctional methyltransferase subunit TRM112-like | 2761478.5 | 2322332.5 | 1316741.12 | 2681462.5 | 2743044.25 | 2997355 | 2884596.5 | 2763977.25 | **1.254013947** | 1 |
| sp\|P39019 | P39019 | RPS19 | 40S ribosomal protein S19 OS=Homo sapiens (Human) G | 100271632 | 91555712 | 98183104 | 111865280 | 137917360 | 115307744 | 139123088 | 111470952 | **1.253669005** | 1 |
| sp\|P01023 | P01023 | A2M | Alpha-2-macroglobulin OS=Homo sapiens (Human) GN=/ | 29018382 | 34377644 | 31588436 | 33191472 | 37681256 | 25543466 | 50347552 | 46980536 | **1.252597153** | 4 |
| sp\|Q9Y23 | Q9Y230 | RUVBL2 | RuvB-like 2 OS=Homo sapiens (Human) GN=RUVBL2 PE= | 18566058 | 19193740 | 23497208 | 20443986 | 28954862 | 25817950 | 22558136 | 24930910 | **1.251659931** | 1 |
| sp\|O4383 | O43837 | IDH3B | Isocitrate dehydrogenase [NAD] subunit beta, mitochonc | 4727963.5 | 6388480 | 7599564.5 | 7643804 | 7558053 | 7262095 | 8525044 | 9633887 | **1.251112072** | 4 |
| sp\|Q9BQ9 | Q9BQ95 | ECSIT | Evolutionarily conserved signaling intermediate in Toll pa | 569444.88 | 520901.75 | 663118.69 | 393255.69 | 629200.5 | 518980.84 | 727031.69 | 808802.25 | **1.250286026** | 4 |
| sp\|Q9UKN | Q9UKM9 | RALY | RNA-binding protein Raly OS=Homo sapiens (Human) GN | 114252208 | 144095632 | 155853296 | 152572784 | 175291632 | 155562160 | 190519984 | 187037184 | **1.249900419** | 1 |
| sp\|P50336 | P50336 | PPOX | Protoporphyrinogen oxidase OS=Homo sapiens (Human) | 1968070 | 1511268 | 706069.06 | 1604338.25 | 1579963.62 | 1831501.38 | 1836455.5 | 1984529.25 | **1.249182712** | 4 |
| sp\|Q9BZE | Q9BZE1 | MRPL37 | 39S ribosomal protein L37, mitochondrial OS=Homo sapi | 8306116.5 | 8082551 | 5728294 | 8994122 | 10591577 | 9896240 | 8444526 | 9930912 | **1.249177162** | 1 |
| sp\|Q9NT6 | Q9NT62 | ATG3 | Ubiquitin-like-conjugating enzyme ATG3 OS=Homo sapie | 2159882.25 | 949924.81 | 2262330 | 847554.88 | 1852755 | 2196068 | 2114505.75 | 1606013.88 | **1.249152322** | 7 |
| sp\|Q9NQ5 | Q9NQ50 | MRPL40 | 39S ribosomal protein L40, mitochondrial OS=Homo sapi | 1484566.88 | 1290940.62 | 1401158.25 | 2016476.62 | 2016201.88 | 1861405.5 | 2096651 | 1761777.5 | **1.249129346** | 3 |
| sp\|A3KMH | A3KMH1 | VWA8 | von Willebrand factor A domain-containing protein 8 OS | 0 | 845739.75 | 286212.22 | 1014591.62 | 1246930.75 | 0 | 317202.62 | 1115210.5 | **1.248213119** | 5.5 |
| sp\|P15121 | P15121 | AKR1B1 | Aldose reductase OS=Homo sapiens (Human) GN=AKR1E | 29018068 | 29161050 | 29052340 | 28974772 | 38724392 | 36883432 | 36298096 | 33050038 | **1.247402639** | 0 |
| sp\|Q9944 | Q99442 | SEC62 | Translocation protein SEC62 OS=Homo sapiens (Human) | 696781.5 | 1234717.38 | 1233039.12 | 0 | 951181.06 | 964443.56 | 706519.44 | 1324537.62 | **1.247158884** | 6 |
| sp\|O0058 | O00584 | RNASET2 | Ribonuclease T2 OS=Homo sapiens (Human) GN=RNASE | 3309258 | 2786625.25 | 3039529 | 3546003.25 | 4242382 | 4187757.75 | 4006028.5 | 3375337 | **1.24682495** | 1 |
| sp\|P49354 | P49354 | FNTA | Protein farnesyltransferase/geranylgeranyltransferase ty | 1154687.62 | 1582711.5 | 129874.13 | 1440580.75 | 1176185.62 | 1002038.38 | 1409434.75 | 1782616.38 | **1.246624219** | 7 |
| sp\|Q0068 | Q00688 | FKBP3 | Peptidyl-prolyl cis-trans isomerase FKBP3 OS=Homo sapi | 9595859 | 10088330 | 10579618 | 9949330 | 15090143 | 10865652 | 13237977 | 10935754 | **1.246595758** | 0 |
| sp\|Q9NQ5 | Q9NQ55 | PPAN | Suppressor of SWI4 1 homolog OS=Homo sapiens (Huma | 659670.19 | 2844225.75 | 901390.31 | 3475501 | 3805740.25 | 657770.5 | 863856.94 | 4496080.5 | **1.246505949** | 7 |
| sp\|P38159 | P38159 | RBMX | RNA-binding motif protein, X chromosome OS=Homo sa | 78340520 | 66165860 | 84998144 | 71597424 | 103322184 | 86494368 | 91827464 | 93674328 | **1.246482617** | 0 |
| sp\|Q9Y36 | Q9Y365 | STARD10 | START domain-containing protein 10 OS=Homo sapiens | 7638079.5 | 7859672 | 8384323 | 5446476 | 8550744 | 8181096 | 9914312 | 9890268 | **1.245762896** | 1 |
| sp\|Q86VN | Q86VN1 | VPS36 | Vacuolar protein-sorting-associated protein 36 OS=Hom | 0 | 898428.19 | 2104910 | 2165660 | 1800435.62 | 2048917.75 | 2589779.25 | 0 | **1.245721585** | 7.5 |
| sp\|P50552 | P50552 | VASP | Vasodilator-stimulated phosphoprotein OS=Homo sapie | 7535689 | 6076921.5 | 6668934 | 7161046 | 8278023.5 | 7441503.5 | 9783935 | 8672732 | **1.245370549** | 1 |
| sp\|O6071 | O60716 | CTNND1 | Catenin delta-1 OS=Homo sapiens (Human) GN=CTNND1 | 24585444 | 20988828 | 24775374 | 20237312 | 29714450 | 29142694 | 28028692 | 25921238 | **1.245290453** | 0 |
| sp\|Q0469 | Q04695 | KRT17 | Keratin, type I cytoskeletal 17 OS=Homo sapiens (Humar | 65716180 | 175691008 | 94878112 | 89984360 | 126427232 | 132894736 | 141119008 | 130368136 | **1.245242535** | 4 |
| sp\|Q9NPF | Q9NPF5 | DMAP1 | DNA methyltransferase 1-associated protein 1 OS=Homo | 0 | 838324.12 | 949034.69 | 1272303 | 1018855.94 | 763063.94 | 1000165.62 | 1020546.25 | **1.242827471** | 6 |
| sp\|P18031 | P18031 | PTPN1 | Tyrosine-protein phosphatase non-receptor type 1 OS=H | 0 | 3277030.5 | 3500242.75 | 1950364.12 | 3912751.75 | 1990482 | 1009136.5 | 3933579 | **1.242713095** | 5 |
| sp\|Q9GZZ | Q9GZZ1 | NAA50 | N-alpha-acetyltransferase 50 OS=Homo sapiens (Human | 1757310 | 0 | 2217674 | 2837350.5 | 2240248.5 | 1981011.25 | 1620596.62 | 2621957.25 | **1.242424843** | 7 |
| sp\|Q9H55 | Q9H553 | ALG2 | Alpha-1,3/1,6-mannosyltransferase ALG2 OS=Homo sapi | 2613158.75 | 3606867.5 | 1886393.38 | 3029923.5 | 3327112.75 | 2970168.75 | 4178756.5 | 3354666.5 | **1.241943099** | 4 |
| sp\|O7539 | O75396 | SEC22B | Vesicle-trafficking protein SEC22b OS=Homo sapiens (Hu | 26120004 | 29914786 | 25475656 | 26408458 | 36262024 | 31980612 | 35532428 | 30234392 | **1.241760721** | 0 |
| sp\|P53680 | P53680 | AP2S1 | AP-2 complex subunit sigma OS=Homo sapiens (Human) | 3829769 | 3836693.25 | 5026841 | 4178215.25 | 5683021.5 | 5617940.5 | 4535712.5 | 5090968 | **1.240412503** | 1 |
| sp\|Q0837 | Q08379 | GOLGA2 | Golgin subfamily A member 2 OS=Homo sapiens (Humar | 2939358.5 | 2603728.5 | 2256917 | 2471070.25 | 2142234 | 2659379 | 4458161.5 | 3476489 | **1.240012796** | 5 |
| sp\|Q9NY1 | Q9NY12 | GAR1 | H/ACA ribonucleoprotein complex subunit 1 OS=Homo s | 6098393 | 6710463 | 5589159 | 3747003.75 | 8083808 | 3735348.5 | 8493142 | 7141168.5 | **1.239712972** | 4 |
| sp\|Q9NP6 | Q9NP66 | HMG20A | High mobility group protein 20A OS=Homo sapiens (Hum | 310480 | 284947.41 | 159773.91 | 215143.61 | 218469.86 | 320655.19 | 345712.28 | 316417.78 | **1.237967111** | 2 |
| sp\|P52597 | P52597 | HNRNPF | Heterogeneous nuclear ribonucleoprotein F OS=Homo s | 85222176 | 82730752 | 77271632 | 77139896 | 99690600 | 102643952 | 101477312 | 95245040 | **1.23790603** | 0 |
| sp\|Q1571 | Q15717 | ELAVL1 | ELAV-like protein 1 OS=Homo sapiens (Human) GN=ELA\ | 50256352 | 46309232 | 54494872 | 49662916 | 59909428 | 63390868 | 63058880 | 62020844 | **1.237424509** | 0 |
| sp\|Q8N5K | Q8N5K1 | CISD2 | CDGSH iron-sulfur domain-containing protein 2 OS=Hom | 4771524.5 | 4877041 | 4605525 | 4290889 | 5390454.5 | 5426310.5 | 5898420.5 | 6223325 | **1.236912152** | 0 |

| | | | | | | | | | | | | |
|---|---|---|---|---|---|---|---|---|---|---|---|---|
| sp\|P60059 | P60059 | SEC61G | Protein transport protein Sec61 subunit gamma OS=Hom | 3521462.5 | 3780608 | 3643213 | 3346648 | 4268456 | 3905286.5 | 4310650 | 5191711 | **1.236788988** | 0 |
| sp\|Q86V2 | Q86V21 | AACS | Acetoacetyl-CoA synthetase OS=Homo sapiens (Human) | 4517569 | 3448938.75 | 4549763 | 3710275 | 6271970 | 4487759.5 | 4552365 | 4750989 | **1.236435888** | 2 |
| sp\|P61966 | P61966 | AP1S1 | AP-1 complex subunit sigma-1A OS=Homo sapiens (Hum | 1035570 | 1250120.38 | 1079639.38 | 998475.06 | 1232449.38 | 1564533.38 | 1491224 | 1105875.25 | **1.236096075** | 2 |
| sp\|P18887 | P18887 | XRCC1 | DNA repair protein XRCC1 OS=Homo sapiens (Human) G | 3397495.5 | 2281743.75 | 4538224 | 3540674 | 4681568 | 3535363.5 | 4675167 | 4112605.5 | **1.235974296** | 3 |
| sp\|Q8NBQ | Q8NBQ5 | HSD17B11 | Estradiol 17-beta-dehydrogenase 11 OS=Homo sapiens ( | 10525288 | 12144356 | 6580651.5 | 12008176 | 14581430 | 13171624 | 10860479 | 12379875 | **1.235950004** | 2 |
| sp\|P09651 | P09651 | HNRNPA1 | Heterogeneous nuclear ribonucleoprotein A1 OS=Homo | 310079520 | 256831808 | 287394368 | 260699792 | 361713600 | 324067680 | 369205920 | 322353536 | **1.235277091** | 0 |
| sp\|O7597 | O75976 | CPD | Carboxypeptidase D OS=Homo sapiens (Human) GN=CPI | 9654066 | 8872526 | 8332867 | 7986245.5 | 11457278 | 11661798 | 9623846 | 10289445 | **1.234940364** | 1 |
| sp\|P62861 | P62861 | FAU | 40S ribosomal protein S30 OS=Homo sapiens (Human) G | 18033528 | 17016716 | 16022908 | 20987504 | 20456726 | 19875714 | 30339226 | 18313838 | **1.234869469** | 3 |
| sp\|P05026 | P05026 | ATP1B1 | Sodium/potassium-transporting ATPase subunit beta-1 ( | 24937490 | 34264956 | 26902496 | 23223992 | 36840760 | 26321404 | 36756620 | 35029064 | **1.234328764** | 2 |
| sp\|Q7RTV | Q7RTV0 | PHF5A | PHD finger-like domain-containing protein 5A OS=Homo | 2923039.5 | 2736591.5 | 0 | 2928706 | 0 | 2577330 | 4894251.5 | 3127464 | **1.23412082** | 6.5 |
| sp\|P00387 | P00387 | CYB5R3 | NADH-cytochrome b5 reductase 3 OS=Homo sapiens (Hu | 63717224 | 67814480 | 62236928 | 66719936 | 93181280 | 90094672 | 66641424 | 71403952 | **1.23353332** | 2 |
| sp\|Q8WT\ | Q8WTV0 | SCARB1 | Scavenger receptor class B member 1 OS=Homo sapiens | 1406482.38 | 1364792.38 | 753906.56 | 1472818 | 1426258 | 1870746.12 | 1380667.12 | 1487045 | **1.23343679** | 3 |
| sp\|Q6IBS0 | Q6IBS0 | TWF2 | Twinfilin-2 OS=Homo sapiens (Human) GN=TWF2 PE=1 S | 8178838.5 | 6011951.5 | 5334484.5 | 7312351 | 7877588 | 8316537 | 7812171 | 9073826 | **1.232602415** | 2 |
| sp\|O9563 | O95630 | STAMBP | STAM-binding protein OS=Homo sapiens (Human) GN=S | 1661475.38 | 945103.69 | 1229475 | 1441962.25 | 1930859.5 | 1367699.5 | 1342700.75 | 1862363.38 | **1.232209742** | 4 |
| sp\|P0C0L4 | P0C0L4 | C4A | Complement C4-A OS=Homo sapiens (Human) GN=C4A I | 24804986 | 26499096 | 22593244 | 25405168 | 27473528 | 25837224 | 33953400 | 35063456 | **1.231868436** | 1 |
| sp\|P10398 | P10398 | ARAF | Serine/threonine-protein kinase A-Raf OS=Homo sapien | 203352.3 | 230932.38 | 124749.38 | 294962.5 | 242504.64 | 331000.5 | 297600.94 | 180727.14 | **1.231659786** | 4 |
| sp\|Q9309 | Q93099 | HGD | Homogentisate 1,2-dioxygenase OS=Homo sapiens (Hun | 14614950 | 14139896 | 12128852 | 13509525 | 16007992 | 17259530 | 18105990 | 15620242 | **1.231656267** | 0 |
| sp\|P62304 | P62304 | SNRPE | Small nuclear ribonucleoprotein E OS=Homo sapiens (Hu | 16996962 | 19139078 | 27236268 | 23735432 | 24642684 | 20813814 | 25038860 | 36721832 | **1.230857212** | 4 |
| sp\|P54920 | P54920 | NAPA | Alpha-soluble NSF attachment protein OS=Homo sapien | 7767942.5 | 6643443.5 | 7242486 | 7280115.5 | 8714016 | 9262076 | 8507889 | 9126850 | **1.230761263** | 0 |
| sp\|Q9NP8 | Q9NP81 | SARS2 | Serine--tRNA ligase, mitochondrial OS=Homo sapiens (H | 4929500 | 4724313.5 | 5347022 | 5419353 | 6642009 | 6792570 | 6791670 | 4896206 | **1.230275372** | 3 |
| sp\|E9PRG | E9PRG8 | C11orf98 | Uncharacterized protein C11orf98 OS=Homo sapiens (Hu | 3806751.75 | 2688427.25 | 3254278.25 | 3020968 | 4118443 | 3965758 | 3904589.5 | 3716972.75 | **1.229854366** | 1 |
| sp\|Q9HAV | Q9HAV7 | GRPEL1 | GrpE protein homolog 1, mitochondrial OS=Homo sapier | 15864581 | 13026793 | 11472202 | 12700626 | 15133323 | 16169278 | 18268984 | 15669624 | **1.22947687** | 2 |
| sp\|Q9BPZ | Q9BPZ7 | MAPKAP1 | Target of rapamycin complex 2 subunit MAPKAP1 OS=Hc | 214168.75 | 181324.06 | 192268.14 | 213855.3 | 267043.91 | 219771.73 | 272371.69 | 226284.12 | **1.22935563** | 0 |
| sp\|Q9H2W | Q9H2W6 | MRPL46 | 39S ribosomal protein L46, mitochondrial OS=Homo sapi | 4033211.5 | 2657272 | 1759681.12 | 4158840.5 | 4068721.75 | 4113602.5 | 3253606.5 | 4060957 | **1.229033346** | 5 |
| sp\|P25685 | P25685 | DNAJB1 | DnaJ homolog subfamily B member 1 OS=Homo sapiens | 28939188 | 28114138 | 31053490 | 30331302 | 39800052 | 32163018 | 38986620 | 34545040 | **1.228445136** | 0 |
| sp\|Q9GZS | Q9GZS3 | WDR61 | WD repeat-containing protein 61 OS=Homo sapiens (Hu | 3895674 | 3743615 | 3994736 | 4206677.5 | 4824991.5 | 5324112 | 4044287.5 | 5257332 | **1.227895228** | 1 |
| sp\|P61978 | P61978 | HNRNPK | Heterogeneous nuclear ribonucleoprotein K OS=Homo s | 343633184 | 337612032 | 345969664 | 258571008 | 398574240 | 394307008 | 387321376 | 398225024 | **1.227597583** | 0 |
| sp\|Q9Y26 | Q9Y265 | RUVBL1 | RuvB-like 1 OS=Homo sapiens (Human) GN=RUVBL1 PE= | 40560608 | 36314900 | 41423776 | 31765436 | 47972064 | 47497464 | 44334424 | 44324864 | **1.226996032** | 0 |
| sp\|Q9NYL | Q9NYL9 | TMOD3 | Tropomodulin-3 OS=Homo sapiens (Human) GN=TMOD3 | 4178931.5 | 3440087 | 3203652.25 | 3567482.5 | 4737061 | 4334569.5 | 4955754.5 | 3619029.25 | **1.22628397** | 1 |
| sp\|P10155 | P10155 | TROVE2 | 60 kDa SS-A/Ro ribonucleoprotein OS=Homo sapiens (Hu | 6917678.5 | 6274744.5 | 5412994.5 | 6556381 | 7301401.5 | 7491478.5 | 8427766 | 7622995 | **1.225812257** | 0 |
| sp\|Q9H3P | Q9H3P2 | NELFA | Negative elongation factor A OS=Homo sapiens (Human) | 7761083 | 6815916 | 8697410 | 10836520 | 10860694 | 9321525 | 8672495 | 12928318 | **1.224916272** | 3 |
| sp\|Q96IX5 | Q96IX5 | ATP5MD | Up-regulated during skeletal muscle growth protein 5 O | 8450874 | 8379883 | 10332626 | 9859108 | 10121948 | 12051806 | 10819983 | 12350916 | **1.224786657** | 1 |
| sp\|P06756 | P06756 | ITGAV | Integrin alpha-V OS=Homo sapiens (Human) GN=ITGAV I | 3517022.5 | 6908184 | 6648605 | 5826382 | 7989316 | 7794949 | 4904791 | 7346960.5 | **1.224269852** | 3 |
| sp\|Q53GQ | Q53GQ0 | HSD17B12 | Very-long-chain 3-oxoacyl-CoA reductase OS=Homo sapi | 16235198 | 16331252 | 13333082 | 16920522 | 15889828 | 20560166 | 16684909 | 23764248 | **1.224117875** | 4 |
| sp\|Q9P2R | Q9P2R3 | ANKFY1 | Rabankyrin-5 OS=Homo sapiens (Human) GN=ANKFY1 PI | 1915405.62 | 2241203.75 | 3278526.75 | 1956259 | 3044593 | 2973461.25 | 3490631.5 | 1984994 | **1.223852218** | 4 |
| sp\|Q9H7Z | Q9H7Z7 | PTGES2 | Prostaglandin E synthase 2 OS=Homo sapiens (Human) ( | 4843313.5 | 6619399.5 | 3468988.75 | 6548981 | 6892183.5 | 7983008 | 4432170 | 6977585.5 | **1.2236551** | 3 |
| sp\|P82932 | P82932 | MRPS6 | 28S ribosomal protein S6, mitochondrial OS=Homo sapie | 603778 | 724575.56 | 451631.53 | 799201.25 | 665278.12 | 757046.06 | 821182.56 | 911933.62 | **1.223424733** | 3 |
| sp\|P13073 | P13073 | COX4I1 | Cytochrome c oxidase subunit 4 isoform 1, mitochondria | 16377473 | 21789188 | 17423676 | 21614632 | 28597568 | 29670034 | 15650523 | 20529102 | **1.223330936** | 6 |
| sp\|P83916 | P83916 | CBX1 | Chromobox protein homolog 1 OS=Homo sapiens (Huma | 8322008 | 8005551 | 8349696.5 | 7970249 | 10158140 | 9844839 | 10847810 | 9082557 | **1.223166873** | 0 |
| sp\|Q9BUJ | Q9BUJ2 | HNRNPUL1 | Heterogeneous nuclear ribonucleoprotein U-like protein | 14959048 | 13116852 | 14223022 | 13265576 | 15101248 | 15753547 | 19600980 | 17506624 | **1.223126303** | 0 |
| sp\|O6076 | O60763 | USO1 | General vesicular transport factor p115 OS=Homo sapier | 25357640 | 28309052 | 27231682 | 28151538 | 33236516 | 31947970 | 35139988 | 33041778 | **1.222983582** | 0 |
| sp\|P00441 | P00441 | SOD1 | Superoxide dismutase [Cu-Zn] OS=Homo sapiens (Huma | 23116112 | 21502894 | 21109308 | 20164928 | 26802536 | 23486802 | 25664016 | 29068720 | **1.222704739** | 0 |
| sp\|P49755 | P49755 | TMED10 | Transmembrane emp24 domain-containing protein 10 C | 30714454 | 32651738 | 32066236 | 29597084 | 39381612 | 36260392 | 39517296 | 37409744 | **1.220264252** | 0 |
| sp\|O9567 | O95674 | CDS2 | Phosphatidate cytidylyltransferase 2 OS=Homo sapiens ( | 2141890.75 | 1875081 | 2199625.25 | 2090995 | 2415201.5 | 2577039.75 | 2603588.5 | 2540502.25 | **1.220128769** | 0 |
| sp\|Q9NYU | Q9NYU2 | UGGT1 | UDP-glucose:glycoprotein glucosyltransferase 1 OS=Hon | 66956536 | 69657504 | 67501856 | 78309376 | 86344552 | 85120864 | 86923824 | 86154888 | **1.219947937** | 0 |

| | | | | | | | | | | | | | |
|---|---|---|---|---|---|---|---|---|---|---|---|---|---|
| sp\|Q9NR4 | Q9NR45 | NANS | Sialic acid synthase OS=Homo sapiens (Human) GN=NAN | 29853242 | 28612364 | 27150944 | 26778684 | 34622556 | 30215546 | 38480156 | 33673120 | **1.218836183** | 0 |
| sp\|P61619 | P61619 | SEC61A1 | Protein transport protein Sec61 subunit alpha isoform 1 | 16136570 | 20837748 | 20996722 | 19488050 | 21503814 | 19744470 | 26310942 | 26826724 | **1.218526451** | 2 |
| sp\|Q7Z7H | Q7Z7H8 | MRPL10 | 39S ribosomal protein L10, mitochondrial OS=Homo sapi | 682785.19 | 623378.06 | 487943.62 | 722386.5 | 949984.81 | 670627 | 749084.19 | 696680.25 | **1.218511555** | 3 |
| sp\|Q8IWZ | Q8IWZ3 | ANKHD1 | Ankyrin repeat and KH domain-containing protein 1 OS= | 1793718.5 | 1722626 | 974263.75 | 1970469.5 | 2300180 | 1583519 | 2047147.62 | 1941557.5 | **1.218435132** | 4 |
| sp\|Q5ST3C | Q5ST30 | VARS2 | Valine--tRNA ligase, mitochondrial OS=Homo sapiens (H | 677939.94 | 1136745.88 | 1181878.38 | 867367.94 | 1098388.88 | 1427798.12 | 899855.12 | 1281545.38 | **1.218341143** | 4 |
| sp\|P30837 | P30837 | ALDH1B1 | Aldehyde dehydrogenase X, mitochondrial OS=Homo sa | 186833552 | 191095392 | 215486656 | 197560480 | 250136704 | 227747200 | 252500384 | 233214416 | **1.218240005** | 0 |
| sp\|P24666 | P24666 | ACP1 | Low molecular weight phosphotyrosine protein phospha | 30391242 | 30052254 | 29179936 | 30086496 | 40154588 | 33715264 | 35007636 | 36903416 | **1.217784577** | 0 |
| sp\|Q9UJSC | Q9UJS0 | SLC25A13 | Calcium-binding mitochondrial carrier protein Aralar2 O | 53346720 | 60091488 | 55689364 | 57832632 | 71419312 | 63203296 | 70035600 | 71725368 | **1.217762282** | 0 |
| sp\|Q86X2 | Q86X29 | LSR | Lipolysis-stimulated lipoprotein receptor OS=Homo sapi | 1382254.62 | 3059942 | 3152986.75 | 3105619.25 | 4452670 | 2178687.5 | 2914627 | 3482038.75 | **1.217480942** | 6 |
| sp\|Q1301 | Q13011 | ECH1 | Delta(3,5)-Delta(2,4)-dienoyl-CoA isomerase, mitochondi | 54738112 | 55505996 | 65702492 | 48499284 | 60541256 | 63777296 | 63997464 | 84907160 | **1.217323174** | 3 |
| sp\|O1496 | O14967 | CLGN | Calmegin OS=Homo sapiens (Human) GN=CLGN PE=1 SV | 3499155 | 2307692 | 2403162.25 | 1469906.88 | 3333221 | 3205089.5 | 3174011.25 | 2070278.75 | **1.217221342** | 6 |
| sp\|P62993 | P62993 | GRB2 | Growth factor receptor-bound protein 2 OS=Homo sapie | 1049975.75 | 1259556.88 | 1392988.5 | 1154565.25 | 1358472.5 | 1219615.5 | 1555943.12 | 1775221.75 | **1.216625031** | 3 |
| sp\|Q9HCC | Q9HCC0 | MCCC2 | Methylcrotonoyl-CoA carboxylase beta chain, mitochond | 19641182 | 19564826 | 19269336 | 19335626 | 25089462 | 22419352 | 22022212 | 25134974 | **1.216615087** | 0 |
| sp\|O9492 | O94925 | GLS | Glutaminase kidney isoform, mitochondrial OS=Homo sa | 45930116 | 43177196 | 37457648 | 37938660 | 52837716 | 42243712 | 52704244 | 52247448 | **1.21598005** | 2 |
| sp\|Q9H47 | Q9H479 | FN3K | Fructosamine-3-kinase OS=Homo sapiens (Human) GN=I | 6185625.5 | 8484100 | 9654020 | 11149366 | 15226361 | 12507179 | 10251226 | 5144176 | **1.215820665** | 5 |
| sp\|O9515 | O95155 | UBE4B | Ubiquitin conjugation factor E4 B OS=Homo sapiens (Hu | 281648.28 | 337624.62 | 348064.34 | 325578.19 | 333273.38 | 377388.78 | 406494.22 | 453592.53 | **1.214889136** | 2 |
| sp\|P38435 | P38435 | GGCX | Vitamin K-dependent gamma-carboxylase OS=Homo sap | 7788539.5 | 8527612 | 7659031 | 7860495 | 9367577 | 9384789 | 10306076 | 9605713 | **1.214491352** | 0 |
| sp\|P38432 | P38432 | COIL | Coilin OS=Homo sapiens (Human) GN=COIL PE=1 SV=1 | 4386838 | 4602096 | 4778064.5 | 3988905.5 | 4852087.5 | 5619461.5 | 5780576.5 | 5282318 | **1.212804682** | 0 |
| sp\|Q70UQ | Q70UQ0 | IKBIP | Inhibitor of nuclear factor kappa-B kinase-interacting pro | 13641058 | 14336153 | 21183546 | 18955094 | 17270046 | 20612426 | 19060436 | 25644862 | **1.212460371** | 4 |
| sp\|Q9NTJ | Q9NTJ3 | SMC4 | Structural maintenance of chromosomes protein 4 OS=H | 5163718 | 4148700 | 5659720 | 22483428 | 2081344.5 | 5319136.5 | 20344174 | 17667990 | **1.212440495** | 8 |
| sp\|P21980 | P21980 | TGM2 | Protein-glutamine gamma-glutamyltransferase 2 OS=Ho | 1677362.75 | 1970871.12 | 2257413.75 | 2301318.5 | 1755546.12 | 3186540.75 | 1939835 | 3066808 | **1.212229918** | 6 |
| sp\|P25325 | P25325 | MPST | 3-mercaptopyruvate sulfurtransferase OS=Homo sapien | 17246840 | 14437696 | 12777294 | 13723563 | 15136274 | 18578588 | 20383154 | 16420779 | **1.211967323** | 2 |
| sp\|Q13442 | Q13442 | PDAP1 | 28 kDa heat- and acid-stable phosphoprotein OS=Homo | 4050936.5 | 3302240.5 | 3565655 | 5031796 | 5585094 | 5065922.5 | 4644489.5 | 4033368.5 | **1.211793949** | 3 |
| sp\|Q9BRT | Q9BRT9 | GINS4 | DNA replication complex GINS protein SLD5 OS=Homo sa | 730523.69 | 0 | 0 | 0 | 0 | 885082.56 | 0 | 0 | **1.211572701** | 7.5 |
| sp\|Q6P1L | Q6P1L8 | MRPL14 | 39S ribosomal protein L14, mitochondrial OS=Homo sapi | 3331149 | 4204534.5 | 2418415.75 | 3869328.5 | 4024757.5 | 3674804.5 | 4591193 | 4440315.5 | **1.210341661** | 3 |
| sp\|O4380 | O43809 | NUDT21 | Cleavage and polyadenylation specificity factor subunit 5 | 31230182 | 20934756 | 29402770 | 37365340 | 38241968 | 32798266 | 36951356 | 35943292 | **1.210217718** | 3 |
| sp\|Q8TBC | Q8TBC4 | UBA3 | NEDD8-activating enzyme E1 catalytic subunit OS=Homo | 4532368 | 5334631 | 1787022.62 | 4954557.5 | 3672733 | 5301147 | 5379099 | 5742230 | **1.20992945** | 4 |
| sp\|Q96AE | Q96AE4 | FUBP1 | Far upstream element-binding protein 1 OS=Homo sapie | 57319068 | 61011216 | 62869396 | 57932176 | 65362016 | 68736536 | 82086096 | 73131488 | **1.209860287** | 0 |
| sp\|P49914 | P49914 | MTHFS | 5-formyltetrahydrofolate cyclo-ligase OS=Homo sapiens | 1929887.25 | 2202519 | 2193079 | 2071843.25 | 2286289.75 | 2060640.88 | 2646444.75 | 3165942.75 | **1.209827403** | 3 |
| sp\|O7587 | O75874 | IDH1 | Isocitrate dehydrogenase [NADP] cytoplasmic OS=Homo | 203402432 | 248333440 | 234603920 | 229877008 | 298626720 | 247903536 | 298400992 | 263520336 | **1.209813642** | 1 |
| sp\|P54136 | P54136 | RARS | Arginine--tRNA ligase, cytoplasmic OS=Homo sapiens (H | 51106676 | 51795252 | 51143032 | 49925148 | 59777836 | 62248564 | 61002292 | 63603232 | **1.209157197** | 0 |
| sp\|O9529 | O95299 | NDUFA10 | NADH dehydrogenase [ubiquinone] 1 alpha subcomplex | 9943337 | 10725814 | 10676710 | 10096653 | 12789907 | 11097387 | 11858008 | 14353182 | **1.208866914** | 0 |
| sp\|P32119 | P32119 | PRDX2 | Peroxiredoxin-2 OS=Homo sapiens (Human) GN=PRDX2 | 313767520 | 368546848 | 361555840 | 316696800 | 482281792 | 344889696 | 433171808 | 383889600 | **1.208490935** | 2 |
| sp\|P56134 | P56134 | ATP5MF | ATP synthase subunit f, mitochondrial OS=Homo sapiens | 17205246 | 16613313 | 15980524 | 18048804 | 20032094 | 19934544 | 20847296 | 21153784 | **1.208110107** | 0 |
| sp\|P06493 | P06493 | CDK1 | Cyclin-dependent kinase 1 OS=Homo sapiens (Human) G | 17764298 | 15947647 | 20257688 | 17809028 | 24067954 | 22514728 | 19400194 | 20732328 | **1.208091692** | 1 |
| sp\|Q9BTT | Q9BTT0 | ANP32E | Acidic leucine-rich nuclear phosphoprotein 32 family me | 6888627 | 11152671 | 11787114 | 10361689 | 12582388 | 12876813 | 12302357 | 10781009 | **1.207823961** | 2 |
| sp\|Q96DZ | Q96DZ1 | ERLEC1 | Endoplasmic reticulum lectin 1 OS=Homo sapiens (Huma | 2156152.25 | 971111.31 | 2061902 | 1835107.62 | 2019073.88 | 2123496.25 | 1916380.38 | 2422796.5 | **1.207491052** | 5 |
| sp\|P51970 | P51970 | NDUFA8 | NADH dehydrogenase [ubiquinone] 1 alpha subcomplex | 4549600 | 3740402 | 3264483.5 | 4903884.5 | 4709918.5 | 5072056.5 | 5280123.5 | 4809461.5 | **1.207383234** | 2 |
| sp\|Q9BYC | Q9BYC8 | MRPL32 | 39S ribosomal protein L32, mitochondrial OS=Homo sapi | 1256376.62 | 1048798.25 | 1126784.62 | 1227229.75 | 1493816.25 | 1461908.75 | 1158812.38 | 1510656.75 | **1.207333259** | 2 |
| sp\|P13995 | P13995 | MTHFD2 | Bifunctional methylenetetrahydrofolate dehydrogenase/ | 13650265 | 12717278 | 17545644 | 16425919 | 23333964 | 16450080 | 24385576 | 8662948 | **1.207054145** | 5 |
| sp\|Q9UDY | Q9UDY8 | MALT1 | Mucosa-associated lymphoid tissue lymphoma transloca | 1475182.12 | 1458418.88 | 2430513.75 | 1345145.5 | 1421286.12 | 3061625.25 | 1944321.12 | 1670618 | **1.206966221** | 5 |
| sp\|Q96E3 | Q96E39 | RBMXL1 | RNA binding motif protein, X-linked-like-1 OS=Homo sap | 63926692 | 56965240 | 71083936 | 60005928 | 81542840 | 69839120 | 72860496 | 79863456 | **1.206856673** | 1 |
| sp\|P52272 | P52272 | HNRNPM | Heterogeneous nuclear ribonucleoprotein M OS=Homo | 334227904 | 344883328 | 411105728 | 349973280 | 443549088 | 382254144 | 482581536 | 429700224 | **1.206844029** | 1 |
| sp\|P32321 | P32321 | DCTD | Deoxycytidylate deaminase OS=Homo sapiens (Human) | 1415330.12 | 1128017 | 450213.84 | 1218186.75 | 1468058.75 | 1349616.12 | 1174615.25 | 1086499.25 | **1.205862677** | 6 |
| sp\|P15374 | P15374 | UCHL3 | Ubiquitin carboxyl-terminal hydrolase isozyme L3 OS=Ho | 6518359.5 | 6513635 | 5499163.5 | 6157063.5 | 7458995 | 7348620 | 7622554.5 | 7334850.5 | **1.205636461** | 0 |

| | | | | | | | | | | | | | |
|---|---|---|---|---|---|---|---|---|---|---|---|---|---|
| sp\|Q1528( | Q15286 | RAB35 | Ras-related protein Rab-35 OS=Homo sapiens (Human) ( | 6799280 | 5325051 | 1508931.38 | 1993652.62 | 2394615.75 | 7253223 | 1537812.88 | 7642819 | **1.204874451** | 5 |
| sp\|Q9UBU | Q9UBU6 | FAM8A1 | Protein FAM8A1 OS=Homo sapiens (Human) GN=FAM8A | 1118381 | 1225347.75 | 1310729.25 | 985993.25 | 1591062.25 | 1464593.38 | 1267503.62 | 1263021.12 | **1.203801111** | 2 |
| sp\|Q9294! | Q92945 | KHSRP | Far upstream element-binding protein 2 OS=Homo sapie | 58503540 | 57670384 | 60197812 | 55641360 | 69770568 | 67073040 | 76970896 | 65090988 | **1.202110988** | 0 |
| sp\|Q5T2E( | Q5T2E6 | ARMH3 | Armadillo-like helical domain-containing protein 3 OS=H | 389157.72 | 385966.19 | 492697.28 | 422404.59 | 397887.81 | 516683.84 | 510639.84 | 606125.38 | **1.20181392** | 2 |
| sp\|Q96JB5 | Q96JB5 | CDK5RAP3 | CDK5 regulatory subunit-associated protein 3 OS=Homo | 2773555.25 | 2894194 | 3315719.5 | 3424807 | 3543326.5 | 2946411.5 | 3649606.25 | 4772195 | **1.201741447** | 2 |
| sp\|P14618 | P14618 | PKM | Pyruvate kinase PKM OS=Homo sapiens (Human) GN=PK | 735042240 | 790612608 | 819702272 | 722524224 | 952441984 | 966928832 | 884387904 | 882084288 | **1.201429454** | 0 |
| sp\|O7534; | O75347 | TBCA | Tubulin-specific chaperone A OS=Homo sapiens (Human | 7883298 | 6898560.5 | 5943549 | 8200339.5 | 8183080.5 | 8443654 | 9693896 | 8421782 | **1.201089552** | 1 |
| sp\|Q0795! | Q07955 | SRSF1 | Serine/arginine-rich splicing factor 1 OS=Homo sapiens ( | 62322672 | 52982928 | 64765624 | 57561312 | 77711208 | 59329860 | 81704704 | 66668508 | **1.201074082** | 2 |
| sp\|Q9NS8( | Q9NS86 | LANCL2 | LanC-like protein 2 OS=Homo sapiens (Human) GN=LAN( | 436721.03 | 0 | 900934.69 | 684521.56 | 515471.84 | 631694.31 | 636461.44 | 644732.75 | **1.200864219** | 8 |
| sp\|Q9P01! | Q9P015 | MRPL15 | 39S ribosomal protein L15, mitochondrial OS=Homo sapi | 10592194 | 9947398 | 8805358 | 9593274 | 11635267 | 11791125 | 13358968 | 9971867 | **1.200805332** | 1 |
| sp\|Q9HD4 | Q9HD45 | TM9SF3 | Transmembrane 9 superfamily member 3 OS=Homo sap | 3970059 | 5173888 | 4436191.5 | 4200275.5 | 5904967 | 5698745 | 4244078.5 | 5502499 | **1.200775724** | 2 |
| sp\|O9578; | O95782 | AP2A1 | AP-2 complex subunit alpha-1 OS=Homo sapiens (Huma | 18140292 | 17019330 | 19782892 | 16826448 | 20262522 | 21176024 | 20805160 | 23918804 | **1.200553939** | 0 |
| sp\|Q0178( | Q01780 | EXOSC10 | Exosome component 10 OS=Homo sapiens (Human) GN: | 6807539 | 6775361 | 7242541 | 7065398 | 7071900 | 6424482 | 12328333 | 7657256 | **1.200464819** | 5 |
| sp\|Q1341( | Q13416 | ORC2 | Origin recognition complex subunit 2 OS=Homo sapiens | 194808.77 | 0 | 248802.84 | 0 | 0 | 0 | 208659.8 | 323736.12 | **1.200139735** | 7 |
| sp\|Q0781; | Q07812 | BAX | Apoptosis regulator BAX OS=Homo sapiens (Human) GN: | 20300580 | 22035306 | 22301012 | 11154831 | 22008358 | 21309896 | 24994772 | 22633182 | **1.199948981** | 4 |
| sp\|P42166 | P42166 | TMPO | Lamina-associated polypeptide 2, isoform alpha OS=Hor | 41788960 | 38032332 | 49917736 | 39495744 | 58139132 | 45790768 | 53492820 | 45644868 | **1.199916457** | 2 |
| sp\|Q8WV\ | Q8WVV9 | HNRNPLL | Heterogeneous nuclear ribonucleoprotein L-like OS=Hor | 16413792 | 16967676 | 15766957 | 15639058 | 22482400 | 16831074 | 20891062 | 17478404 | **1.199042414** | 1 |
| sp\|P23528 | P23528 | CFL1 | Cofilin-1 OS=Homo sapiens (Human) GN=CFL1 PE=1 SV=: | 340113120 | 316187136 | 292754208 | 336875232 | 398667392 | 387821120 | 391690336 | 363184608 | **1.198637422** | 0 |
| sp\|Q1350! | Q13505 | MTX1 | Metaxin-1 OS=Homo sapiens (Human) GN=MTX1 PE=1 S | 2993428.25 | 3325892.5 | 2922373.25 | 3262137.25 | 4248379.5 | 4167807.5 | 3530050.5 | 3037959 | **1.19836842** | 2 |
| sp\|P40926 | P40926 | MDH2 | Malate dehydrogenase, mitochondrial OS=Homo sapien | 179720448 | 185715456 | 182152048 | 185451728 | 218932544 | 204622016 | 226686864 | 227915568 | **1.197966517** | 0 |
| sp\|Q0297! | Q02978 | SLC25A11 | Mitochondrial 2-oxoglutarate/malate carrier protein OS: | 13975389 | 15056768 | 15793102 | 14640332 | 16811036 | 17955898 | 15371168 | 21090958 | **1.197819761** | 1 |
| sp\|O9548( | O95486 | SEC24A | Protein transport protein Sec24A OS=Homo sapiens (Hu | 14152814 | 11280277 | 14990210 | 13666684 | 15173505 | 14437520 | 19051708 | 16122821 | **1.197736587** | 1 |
| sp\|Q1056; | Q10567 | AP1B1 | AP-1 complex subunit beta-1 OS=Homo sapiens (Human | 32202320 | 34006992 | 39763184 | 34463516 | 39096048 | 43186348 | 40622620 | 45240712 | **1.19731204** | 1 |
| sp\|O4361! | O43615 | TIMM44 | Mitochondrial import inner membrane translocase subu | 9189959 | 12497008 | 10481070 | 11427619 | 11385362 | 13166231 | 14655090 | 12981106 | **1.197086907** | 2 |
| sp\|P11498 | P11498 | PC | Pyruvate carboxylase, mitochondrial OS=Homo sapiens ( | 40395408 | 38448900 | 33846628 | 34805600 | 41915736 | 43685768 | 47932580 | 42969392 | **1.196661839** | 0 |
| sp\|Q9C0I1 | Q9C0I1 | MTMR12 | Myotubularin-related protein 12 OS=Homo sapiens (Hur | 465499.03 | 426865.31 | 424846.47 | 540929.56 | 570297.38 | 480830.81 | 697941.94 | 474318.28 | **1.196566441** | 2 |
| sp\|O9542; | O95427 | PIGN | GPI ethanolamine phosphate transferase 1 OS=Homo sa | 117383.78 | 93136.95 | 123713.4 | 163393.08 | 142214.89 | 151449.23 | 153000.98 | 148604.48 | **1.196215898** | 4 |
| sp\|Q49AR: | Q49AR2 | C5orf22 | UPF0489 protein C5orf22 OS=Homo sapiens (Human) GN | 1600093.5 | 1376527.25 | 822816.25 | 1328975.38 | 1612321.25 | 1812448.25 | 1183395.12 | 1525257.38 | **1.195968956** | 4 |
| sp\|O9574; | O95747 | OXSR1 | Serine/threonine-protein kinase OSR1 OS=Homo sapiens | 4822196.5 | 4794636 | 5343224 | 6370521.5 | 5889252.5 | 6225881.5 | 6759574.5 | 6620731 | **1.195253101** | 2 |
| sp\|P00403 | P00403 | MT-CO2 | Cytochrome c oxidase subunit 2 OS=Homo sapiens (Hum | 39062880 | 48808480 | 36054016 | 65392260 | 75956664 | 67339680 | 32307172 | 50521532 | **1.194421464** | 5 |
| sp\|P12429 | P12429 | ANXA3 | Annexin A3 OS=Homo sapiens (Human) GN=ANXA3 PE=1 | 9983936 | 11485311 | 10838097 | 12754444 | 11102681 | 14157704 | 14590326 | 13949479 | **1.193920445** | 2 |
| sp\|Q1555! | Q15555 | MAPRE2 | Microtubule-associated protein RP/EB family member 2 | 10054717 | 9674569 | 8106506.5 | 9086092 | 10747499 | 9914293 | 11887529 | 11520865 | **1.193606085** | 1 |
| sp\|O1496( | O14964 | HGS | Hepatocyte growth factor-regulated tyrosine kinase subs | 5966748.5 | 5754836 | 8156780 | 4994700 | 7142281.5 | 5597592.5 | 8496930 | 8410551 | **1.191946212** | 4 |
| sp\|P19012 | P19012 | KRT15 | Keratin, type I cytoskeletal 15 OS=Homo sapiens (Humar | 61646000 | 186321664 | 87935328 | 86795512 | 118587504 | 125495264 | 135031712 | 124067232 | **1.190403342** | 4 |
| sp\|O4339( | O43390 | HNRNPR | Heterogeneous nuclear ribonucleoprotein R OS=Homo s | 78649072 | 89294032 | 86482536 | 76220424 | 96283144 | 99006424 | 100893056 | 97069224 | **1.189343805** | 0 |
| sp\|Q9NXR | Q9NXR7 | BABAM2 | BRISC and BRCA1-A complex member 2 OS=Homo sapie | 1098475.38 | 1221507.12 | 1185135.88 | 1453814.12 | 1348422.12 | 1620273.38 | 1472174.38 | 1455034.25 | **1.188946236** | 1 |
| sp\|Q9BXS! | Q9BXS9 | SLC26A6 | Solute carrier family 26 member 6 OS=Homo sapiens (Hu | 100258.95 | 198901.02 | 123729.38 | 189012.11 | 182007.7 | 184931.2 | 151083.55 | 208867.33 | **1.187919669** | 6 |
| sp\|A5YKK( | A5YKK6 | CNOT1 | CCR4-NOT transcription complex subunit 1 OS=Homo sa | 7161994 | 9069741 | 11200296 | 8367517 | 8881465 | 9530940 | 11692726 | 12396292 | **1.187205576** | 3 |
| sp\|Q9NW( | Q9NW64 | RBM22 | Pre-mRNA-splicing factor RBM22 OS=Homo sapiens (Hu | 1815376.38 | 2063864 | 1883892.62 | 2005343.25 | 2319722.5 | 1777518.38 | 2407708.75 | 2715566.5 | **1.186914375** | 4 |
| sp\|Q8TD4 | Q8TD47 | RPS4Y2 | 40S ribosomal protein S4, Y isoform 2 OS=Homo sapiens | 115869952 | 114293216 | 118825312 | 104107856 | 141277136 | 146155696 | 119947720 | 130082896 | **1.186201267** | 0 |
| sp\|Q9BVK | Q9BVK6 | TMED9 | Transmembrane emp24 domain-containing protein 9 OS | 10555952 | 9943623 | 9977886 | 11122567 | 13275473 | 11510507 | 11699311 | 12846742 | **1.18586538** | 0 |
| sp\|Q9BV8( | Q9BV86 | NTMT1 | N-terminal Xaa-Pro-Lys N-methyltransferase 1 OS=Homo | 6853197.5 | 6675112 | 6624190 | 6688727.5 | 9796391 | 7451268 | 7461911.5 | 7116912 | **1.185731282** | 0 |
| sp\|Q9NUB | Q9NUB1 | ACSS1 | Acetyl-coenzyme A synthetase 2-like, mitochondrial OS= | 6179077.5 | 6642380.5 | 6172607 | 6188214.5 | 7237490 | 7063510 | 7655689.5 | 7901036.5 | **1.185664149** | 0 |
| sp\|Q6NVY | Q6NVY1 | HIBCH | 3-hydroxyisobutyryl-CoA hydrolase, mitochondrial OS=H | 13685492 | 10029416 | 12458650 | 13520871 | 15594508 | 12175734 | 16457546 | 14658062 | **1.184958781** | 3 |
| sp\|Q9BST! | Q9BST9 | RTKN | Rhotekin OS=Homo sapiens (Human) GN=RTKN PE=1 SV= | 1440176.25 | 1286581.75 | 1154442.5 | 1219075.75 | 1578551.75 | 1539876.38 | 1348361.38 | 1576609.12 | **1.184915941** | 1 |

| | | | | | | | | | | | | | |
|---|---|---|---|---|---|---|---|---|---|---|---|---|---|
| sp\|Q8TDD | Q8TDD1 | DDX54 | ATP-dependent RNA helicase DDX54 OS=Homo sapiens ( | 5557886 | 5684894 | 8850416 | 6208318 | 7588981 | 6670319.5 | 9009779 | 7882869.5 | **1.18441657** | 3 |
| sp\|P02765 | P02765 | AHSG | Alpha-2-HS-glycoprotein OS=Homo sapiens (Human) GN | 1358858.88 | 1195394.12 | 1044169.38 | 1468533 | 1833414.5 | 1946516.5 | 1221727.38 | 995393.69 | **1.183561255** | 6 |
| sp\|P13688 | P13688 | CEACAM1 | Carcinoembryonic antigen-related cell adhesion molecul | 8539084 | 10252300 | 9177266 | 8505510 | 11500050 | 9719825 | 9748514 | 12200708 | **1.183552877** | 2 |
| sp\|Q0113( | Q01130 | SRSF2 | Serine/arginine-rich splicing factor 2 OS=Homo sapiens ( | 50380552 | 49046068 | 47735352 | 49457056 | 62910764 | 53778912 | 61812496 | 54171416 | **1.18337269** | 0 |
| sp\|P46977 | P46977 | STT3A | Dolichyl-diphosphooligosaccharide--protein glycosyltrans | 25074574 | 32210144 | 30420196 | 31981356 | 34235068 | 35391312 | 31194124 | 40796096 | **1.183231794** | 2 |
| sp\|P84103 | P84103 | SRSF3 | Serine/arginine-rich splicing factor 3 OS=Homo sapiens ( | 73941416 | 66702096 | 75938360 | 73404112 | 92721416 | 76984976 | 96237872 | 76910528 | **1.182315046** | 0 |
| sp\|Q96HY( | Q96HY6 | DDRGK1 | DDRGK domain-containing protein 1 OS=Homo sapiens ( | 5214294.5 | 4988403.5 | 6251478.5 | 6306605 | 7196245.5 | 6927935 | 6240287.5 | 6527057.5 | **1.181485157** | 2 |
| sp\|Q9287{ | Q92878 | RAD50 | DNA repair protein RAD50 OS=Homo sapiens (Human) G | 2551849.25 | 6139843 | 3271597.5 | 3229290 | 3528033.25 | 4175074 | 3625410 | 6618553.5 | **1.181305022** | 3 |
| sp\|O4349: | O43493 | TGOLN2 | Trans-Golgi network integral membrane protein 2 OS=H | 10125552 | 11940012 | 14953559 | 11381530 | 15326700 | 12223756 | 15334575 | 14290248 | **1.18129148** | 2 |
| sp\|Q9P28: | Q9P287 | BCCIP | BRCA2 and CDKN1A-interacting protein OS=Homo sapie | 15919481 | 16375306 | 17901696 | 17160268 | 20491374 | 22420094 | 15788732 | 20867346 | **1.181285392** | 4 |
| sp\|Q7Z7H! | Q7Z7H5 | TMED4 | Transmembrane emp24 domain-containing protein 4 OS | 3831533 | 5552228 | 5245603 | 5388956 | 6793154 | 5752806 | 5383151 | 5705916.5 | **1.180669881** | 2 |
| sp\|Q9UJW | Q9UJW0 | DCTN4 | Dynactin subunit 4 OS=Homo sapiens (Human) GN=DCTI | 609852.62 | 609588.19 | 3830096 | 669248.75 | 744213.5 | 994243.25 | 4341720 | 671335.31 | **1.180584932** | 3 |
| sp\|Q9Y3C: | Q9Y3C1 | NOP16 | Nucleolar protein 16 OS=Homo sapiens (Human) GN=NC | 583560.25 | 467496.75 | 365004.53 | 635548.31 | 527471 | 614581.56 | 662320.94 | 617516.62 | **1.180482796** | 4 |
| sp\|P02647 | P02647 | APOA1 | Apolipoprotein A-I OS=Homo sapiens (Human) GN=APO/ | 4638932.5 | 4897589.5 | 7672019.5 | 4449986.5 | 8212490 | 4344621 | 6505262 | 6498456.5 | **1.180173902** | 6 |
| sp\|O6066< | O60664 | PLIN3 | Perilipin-3 OS=Homo sapiens (Human) GN=PLIN3 PE=1 S | 39685736 | 35434436 | 28438752 | 37268884 | 42518472 | 38758356 | 46283592 | 38626292 | **1.180070288** | 2 |
| sp\|Q1361: | Q13617 | CUL2 | Cullin-2 OS=Homo sapiens (Human) GN=CUL2 PE=1 SV=2 | 440709.38 | 341335.72 | 798030.38 | 0 | 0 | 404216.81 | 592495.5 | 867483.38 | **1.179814328** | 6.5 |
| sp\|Q8N1G | Q8N1G4 | LRRC47 | Leucine-rich repeat-containing protein 47 OS=Homo sap | 30328118 | 25866860 | 27446932 | 21045176 | 36859264 | 28955888 | 29986132 | 27592536 | **1.178691897** | 3 |
| sp\|O4368: | O43681 | ASNA1 | ATPase ASNA1 OS=Homo sapiens (Human) GN=ASNA1 P | 3638609 | 3900591.75 | 3561417 | 4029759 | 5093134 | 5372278.5 | 4137997.75 | 3229521.5 | **1.178617826** | 4 |
| sp\|Q9UBP | Q9UBP6 | METTL1 | tRNA (guanine-N(7)-)-methyltransferase OS=Homo sapie | 1578738.75 | 1615572 | 1639665.25 | 1754348.25 | 1678267.75 | 1885580.25 | 2162565.5 | 2034245.38 | **1.177941247** | 1 |
| sp\|O7541( | O75410 | TACC1 | Transforming acidic coiled-coil-containing protein 1 OS= | 229127.89 | 0 | 0 | 264550.88 | 0 | 0 | 581515.88 | 0 | **1.177923612** | 7 |
| sp\|P15924 | P15924 | DSP | Desmoplakin OS=Homo sapiens (Human) GN=DSP PE=1 S | 190393408 | 238639136 | 195770304 | 181146336 | 240325696 | 259438864 | 235048816 | 214382992 | **1.177737241** | 2 |
| sp\|O7525: | O75251 | NDUFS7 | NADH dehydrogenase [ubiquinone] iron-sulfur protein 7, | 707409.19 | 983965.75 | 1424329.88 | 847533.94 | 840989.25 | 997990.94 | 1337336.88 | 1488944.88 | **1.177133711** | 5 |
| sp\|Q9300{ | Q93008 | USP9X | Probable ubiquitin carboxyl-terminal hydrolase FAF-X O! | 4694591 | 4343529 | 2586625 | 4048854 | 4232716 | 5053167 | 5090065 | 4068479.75 | **1.176783185** | 4 |
| sp\|Q9UH9 | Q9UH99 | SUN2 | SUN domain-containing protein 2 OS=Homo sapiens (Hu | 3837335.5 | 1501249 | 2026705.25 | 2690701.75 | 2664259 | 4866791 | 1337740 | 2963363 | **1.176627188** | 7 |
| sp\|P07686 | P07686 | HEXB | Beta-hexosaminidase subunit beta OS=Homo sapiens (H | 7357706 | 5406937.5 | 8170815 | 7673855 | 8435016 | 6333258 | 10293199 | 8592738 | **1.176337594** | 3 |
| sp\|O0040: | O00401 | WASL | Neural Wiskott-Aldrich syndrome protein OS=Homo sapi | 1237296.25 | 1969446.5 | 1875287.25 | 1686475.25 | 1989051.5 | 1741573.75 | 1931076 | 2300223.25 | **1.176319469** | 3 |
| sp\|Q9Y39< | Q9Y394 | DHRS7 | Dehydrogenase/reductase SDR family member 7 OS=Ho | 615092.12 | 402049.75 | 408842.5 | 509375.59 | 567981.56 | 653204.81 | 533380 | 520999.41 | **1.175784261** | 3 |
| sp\|O9521{ | O95218 | ZRANB2 | Zinc finger Ran-binding domain-containing protein 2 OS= | 2161406.5 | 1917751.5 | 1923649 | 777481.12 | 1893746.5 | 1836606.25 | 2227419.5 | 2011076.75 | **1.175296515** | 7 |
| sp\|Q9Y3B: | Q9Y3B3 | TMED7 | Transmembrane emp24 domain-containing protein 7 OS | 5053771 | 5480049 | 5189798 | 5695705.5 | 6765688.5 | 6712542 | 5715504 | 5972336 | **1.174923685** | 0 |
| sp\|O6021( | O60216 | RAD21 | Double-strand-break repair protein rad21 homolog OS=I | 1664450 | 3048145 | 3421301 | 3252848.75 | 3288964 | 3270446.5 | 2990847.5 | 3823727.5 | **1.174522288** | 5 |
| sp\|Q9NQT | Q9NQT5 | EXOSC3 | Exosome complex component RRP40 OS=Homo sapiens | 5558442 | 5602384.5 | 5575092 | 6337948 | 7101806 | 6278324 | 7280215.5 | 6433113 | **1.174205394** | 1 |
| sp\|P82675 | P82675 | MRPS5 | 28S ribosomal protein S5, mitochondrial OS=Homo sapie | 2013594.38 | 2632837.25 | 2765850 | 2585376 | 3271788.75 | 2151997 | 3490837 | 2822130.25 | **1.173950283** | 3 |
| sp\|P2002( | P20020 | ATP2B1 | Plasma membrane calcium-transporting ATPase 1 OS=H | 8380817 | 9310960 | 7589552.5 | 9870477 | 10461559 | 11768555 | 8619174 | 10377179 | **1.172812185** | 2 |
| sp\|P1455( | P14550 | AKR1A1 | Alcohol dehydrogenase [NADP(+)] OS=Homo sapiens (Hu | 55835628 | 53626604 | 52177184 | 50352620 | 59321968 | 56219920 | 66333728 | 66597404 | **1.172086578** | 0 |
| sp\|Q9UKY | Q9UKY7 | CDV3 | Protein CDV3 homolog OS=Homo sapiens (Human) GN=( | 3332542.5 | 2499397 | 2353687.25 | 3325052 | 3885633 | 3176949.5 | 3827242 | 2600865.5 | **1.17201516** | 4 |
| sp\|P63092 | P63092 | GNAS | Guanine nucleotide-binding protein G(s) subunit alpha is | 5310941 | 5708188 | 4705620 | 5114551.5 | 5865677 | 6646234 | 4724645.5 | 7187007 | **1.17199536** | 3 |
| sp\|P30085 | P30085 | CMPK1 | UMP-CMP kinase OS=Homo sapiens (Human) GN=CMPK | 18645894 | 12422710 | 15495858 | 19200872 | 19797120 | 18791896 | 20546020 | 17928300 | **1.171792665** | 3 |
| sp\|P0DP2: | P0DP23 | CALM1 | Calmodulin-1 OS=Homo sapiens (Human) GN=CALM1 PE | 19087934 | 21492500 | 24295556 | 28790060 | 26184716 | 19770536 | 34388352 | 29404346 | **1.171694013** | 4 |
| sp\|O0017{ | O00178 | GTPBP1 | GTP-binding protein 1 OS=Homo sapiens (Human) GN=G | 1621154.38 | 1729789.5 | 1404950.25 | 1791202.5 | 1854374.75 | 2046487.75 | 2053399.62 | 1716275.25 | **1.171593731** | 2 |
| sp\|O7547: | O75477 | ERLIN1 | Erlin-1 OS=Homo sapiens (Human) GN=ERLIN1 PE=1 SV= | 28920620 | 26638688 | 31722512 | 25876502 | 36097128 | 36330132 | 28396938 | 31747970 | **1.171563573** | 2 |
| sp\|P2644( | P26440 | IVD | Isovaleryl-CoA dehydrogenase, mitochondrial OS=Homo | 9720307 | 8548890 | 6662472 | 7645472 | 11039970 | 9679945 | 9878598 | 7557523 | **1.17125183** | 4 |
| sp\|P42226 | P42226 | STAT6 | Signal transducer and activator of transcription 6 OS=Ho | 359290.78 | 221010.02 | 213996.45 | 244533.17 | 244470.39 | 350585.97 | 295737.31 | 325690.69 | **1.171013417** | 5 |
| sp\|Q96DV | Q96DV4 | MRPL38 | 39S ribosomal protein L38, mitochondrial OS=Homo sapi | 8125067 | 8990635 | 8112477 | 8235210 | 10632530 | 8139143 | 11139808 | 9272222 | **1.170942459** | 2 |
| sp\|Q1287< | Q12874 | SF3A3 | Splicing factor 3A subunit 3 OS=Homo sapiens (Human) ( | 24177362 | 23166430 | 25713742 | 26245656 | 31095666 | 24964066 | 28957852 | 31234708 | **1.170680338** | 2 |
| sp\|Q9Y45( | Q9Y450 | HBS1L | HBS1-like protein OS=Homo sapiens (Human) GN=HBS1L | 1539437.62 | 1603256.5 | 2252571 | 3430259 | 2645889.5 | 3491072.5 | 2330796.5 | 1857104 | **1.169886611** | 4 |

| | | | | | | | | | | | | | |
|---|---|---|---|---|---|---|---|---|---|---|---|---|---|
| sp\|Q6NUK | Q6NUK1 | SLC25A24 | Calcium-binding mitochondrial carrier protein SCaMC-1 | 1946386.88 | 1791684 | 1633180 | 1832113.5 | 2235102.5 | 2345453.75 | 1851920.12 | 1993100.12 | **1.169672398** | 1 |
| sp\|Q0165 | Q01658 | DR1 | Protein Dr1 OS=Homo sapiens (Human) GN=DR1 PE=1 SV | 1153870.75 | 1259612 | 1216459.75 | 1202894.38 | 1432698.75 | 1438743 | 1646479.62 | 1133158 | **1.16930894** | 4 |
| sp\|Q9NX2 | Q9NX20 | MRPL16 | 39S ribosomal protein L16, mitochondrial OS=Homo sapi | 6019518 | 5977784 | 5487078 | 6154799.5 | 7324178.5 | 6883092.5 | 7659484.5 | 5773797.5 | **1.169268713** | 3 |
| sp\|P40925 | P40925 | MDH1 | Malate dehydrogenase, cytoplasmic OS=Homo sapiens ( | 119680296 | 120944064 | 112612680 | 115754904 | 146140880 | 137182592 | 133597784 | 131299256 | **1.16893375** | 0 |
| sp\|Q96H7 | Q96H79 | ZC3HAV1L | Zinc finger CCCH-type antiviral protein 1-like OS=Homo s | 1310495.25 | 0 | 0 | 696642.88 | 788700.38 | 1036615.25 | 520765.38 | 0 | **1.168868737** | 7 |
| sp\|P26583 | P26583 | HMGB2 | High mobility group protein B2 OS=Homo sapiens (Huma | 31601986 | 31267066 | 33607132 | 46311256 | 46643468 | 40551196 | 42339988 | 37338120 | **1.168679626** | 3 |
| sp\|Q1518 | Q15181 | PPA1 | Inorganic pyrophosphatase OS=Homo sapiens (Human) | 40262312 | 44614328 | 45426520 | 46705328 | 49638920 | 48241252 | 54287348 | 54595356 | **1.168095826** | 0 |
| sp\|Q9288 | Q92882 | OSTF1 | Osteoclast-stimulating factor 1 OS=Homo sapiens (Huma | 762303.94 | 724738.69 | 888700.69 | 857597.06 | 858635.12 | 947881.5 | 978646.75 | 988121.56 | **1.166992796** | 1 |
| sp\|P09012 | P09012 | SNRPA | U1 small nuclear ribonucleoprotein A OS=Homo sapiens | 23187080 | 20186450 | 20028458 | 20313292 | 26953008 | 22080282 | 24702560 | 23959034 | **1.166989873** | 1 |
| sp\|Q1386 | Q13868 | EXOSC2 | Exosome complex component RRP4 OS=Homo sapiens (I | 2460675.75 | 2728158 | 1937369.75 | 2682846.25 | 2941889.75 | 2888689 | 2648883 | 2966999.25 | **1.166928631** | 2 |
| sp\|Q9UJC | Q9UJC5 | SH3BGRL2 | SH3 domain-binding glutamic acid-rich-like protein 2 OS | 2891905 | 2921375.5 | 3015723.75 | 3068159 | 3042868.75 | 3597671.75 | 4002944.25 | 3238346 | **1.166818548** | 1 |
| sp\|O9476 | O94760 | DDAH1 | N(G),N(G)-dimethylarginine dimethylaminohydrolase 1 ( | 6264270 | 6997275.5 | 5482725 | 7077444.5 | 7943873.5 | 6688519 | 7909474.5 | 7573239.5 | **1.166270579** | 2 |
| sp\|P16422 | P16422 | EPCAM | Epithelial cell adhesion molecule OS=Homo sapiens (Hur | 4267123 | 5116635.5 | 4512786.5 | 5030629 | 5158667 | 5224817.5 | 5263685.5 | 6414206.5 | **1.165592735** | 0 |
| sp\|P03915 | P03915 | MT-ND5 | NADH-ubiquinone oxidoreductase chain 5 OS=Homo sap | 898710.56 | 0 | 449251.19 | 1367986.88 | 1291516.5 | 1580526.12 | 293287 | 0 | **1.165460048** | 7.5 |
| sp\|Q9NYV | Q9NYV4 | CDK12 | Cyclin-dependent kinase 12 OS=Homo sapiens (Human) | 950150.06 | 1047530.38 | 0 | 883414 | 874849.56 | 731561.62 | 995541.06 | 754045.19 | **1.164834232** | 6 |
| sp\|O1493 | O14936 | CASK | Peripheral plasma membrane protein CASK OS=Homo sa | 1100667.25 | 1459725 | 897132.19 | 1207474.38 | 1520088.88 | 1367586.88 | 550411.69 | 1991319.75 | **1.163860359** | 5 |
| sp\|Q9Y3D | Q9Y3D7 | PAM16 | Mitochondrial import inner membrane translocase subu | 1503922.75 | 1300611.25 | 1165328.75 | 924811.56 | 1449139.25 | 1472004 | 1454679.62 | 1320041.5 | **1.163686082** | 4 |
| sp\|Q9H84 | Q9H845 | ACAD9 | Acyl-CoA dehydrogenase family member 9, mitochondria | 5627906 | 6030904.5 | 3249817 | 4851354.5 | 5543671 | 5823854 | 5115434.5 | 6510725.5 | **1.163649086** | 5 |
| sp\|P06753 | P06753 | TPM3 | Tropomyosin alpha-3 chain OS=Homo sapiens (Human) | 105606240 | 101796840 | 96929320 | 106801856 | 135817888 | 90940472 | 144842960 | 106796416 | **1.16360466** | 5 |
| sp\|P08621 | P08621 | SNRNP70 | U1 small nuclear ribonucleoprotein 70 kDa OS=Homo sa | 22174346 | 20996636 | 19873056 | 22135222 | 24718592 | 19544738 | 32544224 | 22293418 | **1.163440161** | 4 |
| sp\|P07951 | P07951 | TPM2 | Tropomyosin beta chain OS=Homo sapiens (Human) GN= | 97805872 | 95158848 | 90941192 | 99702872 | 125262576 | 81857368 | 138360832 | 100796224 | **1.163364914** | 4 |
| sp\|Q9BVL | Q9BVL4 | SELENOO | Selenoprotein O OS=Homo sapiens (Human) GN=SELENC | 273309.16 | 192654.09 | 224376.48 | 215630.48 | 329804.69 | 226548.66 | 254959.2 | 242473.27 | **1.163157252** | 3 |
| sp\|P49458 | P49458 | SRP9 | Signal recognition particle 9 kDa protein OS=Homo sapie | 8161429.5 | 9318331 | 8733590 | 9839448 | 11096410 | 10246378 | 10417793 | 10161616 | **1.162800081** | 0 |
| sp\|P17252 | P17252 | PRKCA | Protein kinase C alpha type OS=Homo sapiens (Human) | 3285735.25 | 2851430.25 | 2989745.5 | 3307330.5 | 3483071.5 | 3544975 | 3405393.25 | 4024857.5 | **1.162780798** | 0 |
| sp\|O7560 | O75607 | NPM3 | Nucleoplasmin-3 OS=Homo sapiens (Human) GN=NPM3 | 3232515.5 | 8970663 | 8248825 | 7051072 | 14353993 | 3846909.25 | 3270297.5 | 10459637 | **1.160991495** | 6 |
| sp\|Q1662 | Q16629 | SRSF7 | Serine/arginine-rich splicing factor 7 OS=Homo sapiens ( | 39098260 | 33832588 | 36473768 | 33388604 | 43466144 | 38013780 | 47990096 | 36310060 | **1.160980052** | 3 |
| sp\|P32754 | P32754 | HPD | 4-hydroxyphenylpyruvate dioxygenase OS=Homo sapien | 33044706 | 32559798 | 40613156 | 33498130 | 41920020 | 44850104 | 37278128 | 38153268 | **1.160939075** | 2 |
| sp\|Q4G0J | Q4G0J3 | LARP7 | La-related protein 7 OS=Homo sapiens (Human) GN=LAR | 2871302.25 | 1570383.5 | 3096398.5 | 1752502.75 | 2635876 | 2836394.75 | 2678442 | 2633184.5 | **1.160733681** | 8 |
| sp\|Q96QF | Q96QF0 | RAB3IP | Rab-3A-interacting protein OS=Homo sapiens (Human) G | 97301.34 | 62644.36 | 0 | 86525.12 | 0 | 108441.66 | 73899.53 | 103629.3 | **1.16026104** | 5.5 |
| sp\|Q96QK | Q96QK1 | VPS35 | Vacuolar protein sorting-associated protein 35 OS=Hom | 50526532 | 48364116 | 50932656 | 50150340 | 55252892 | 57461904 | 53457416 | 65745920 | **1.159743491** | 0 |
| sp\|Q9NSE | Q9NSE4 | IARS2 | Isoleucine--tRNA ligase, mitochondrial OS=Homo sapien | 32643434 | 31862934 | 30437354 | 26830200 | 41341232 | 37792416 | 31007580 | 31071002 | **1.159626196** | 4 |
| sp\|P36873 | P36873 | PPP1CC | Serine/threonine-protein phosphatase PP1-gamma catal | 35884392 | 37962460 | 38097984 | 37264672 | 40437204 | 45243840 | 41331796 | 46000192 | **1.159530879** | 0 |
| sp\|Q96FW | Q96FW1 | OTUB1 | Ubiquitin thioesterase OTUB1 OS=Homo sapiens (Huma | 16742373 | 17500392 | 19027668 | 17166868 | 18989798 | 18390096 | 19601224 | 24682898 | **1.159385934** | 2 |
| sp\|Q9UHD | Q9UHD9 | UBQLN2 | Ubiquilin-2 OS=Homo sapiens (Human) GN=UBQLN2 PE= | 912137.81 | 861978.12 | 944640.38 | 880849 | 733863.88 | 1294064.62 | 1083353.25 | 1060233.5 | **1.158881291** | 4 |
| sp\|Q8NC5 | Q8NC56 | LEMD2 | LEM domain-containing protein 2 OS=Homo sapiens (Hu | 4555267.5 | 3178750 | 5566909 | 4964075 | 6034449 | 3968933 | 5924105 | 5237346.5 | **1.1587644** | 4 |
| sp\|P25705 | P25705 | ATP5F1A | ATP synthase subunit alpha, mitochondrial OS=Homo sa | 543393728 | 536609728 | 547233792 | 553953600 | 641347200 | 607008896 | 636973184 | 641468864 | **1.158448902** | 0 |
| sp\|Q86UP | Q86UP2 | KTN1 | Kinectin OS=Homo sapiens (Human) GN=KTN1 PE=1 SV= | 26940250 | 25562422 | 27363494 | 22920396 | 28312992 | 27282382 | 31527442 | 31945134 | **1.158399967** | 1 |
| sp\|Q9H44 | Q9H444 | CHMP4B | Charged multivesicular body protein 4b OS=Homo sapie | 853126.19 | 731727.75 | 667982 | 726525.69 | 1030294.88 | 742422.5 | 999540 | 678348.56 | **1.158169557** | 4 |
| sp\|Q9262 | Q92620 | DHX38 | Pre-mRNA-splicing factor ATP-dependent RNA helicase P | 3673971 | 4163365 | 3050019 | 2638772.75 | 3710089.5 | 3372691.25 | 4618901.5 | 3953630.5 | **1.157412753** | 4 |
| sp\|Q5JTH | Q5JTH9 | RRP12 | RRP12-like protein OS=Homo sapiens (Human) GN=RRP1 | 30388636 | 31576220 | 35507092 | 30478404 | 36884124 | 33591508 | 40693908 | 36800672 | **1.156465845** | 1 |
| sp\|O9507 | O95070 | YIF1A | Protein YIF1A OS=Homo sapiens (Human) GN=YIF1A PE= | 2444110.5 | 2562784.75 | 1839073.25 | 2439381.5 | 2343645 | 2420526 | 3422511.5 | 2549266.5 | **1.156224483** | 7 |
| sp\|O0056 | O00567 | NOP56 | Nucleolar protein 56 OS=Homo sapiens (Human) GN=NC | 60721144 | 61331276 | 66283460 | 66753120 | 70144360 | 64996460 | 77659176 | 82053160 | **1.15588346** | 2 |
| sp\|P23246 | P23246 | SFPQ | Splicing factor, proline- and glutamine-rich OS=Homo sa | 174643488 | 174259008 | 189422432 | 175902080 | 215908352 | 188089888 | 216274928 | 205249520 | **1.155826759** | 1 |
| sp\|Q9BZH | Q9BZH6 | WDR11 | WD repeat-containing protein 11 OS=Homo sapiens (Hu | 132809.7 | 175248.7 | 142267.33 | 205950.52 | 142963.11 | 231352.03 | 184463.03 | 199593.67 | **1.15556801** | 4 |
| sp\|P27144 | P27144 | AK4 | Adenylate kinase 4, mitochondrial OS=Homo sapiens (Hu | 27280844 | 24118800 | 19566464 | 21506286 | 28511634 | 27205320 | 27010232 | 24045256 | **1.154641265** | 4 |

| | | | | | | | | | | | | | |
|---|---|---|---|---|---|---|---|---|---|---|---|---|---|
| sp\|O9601: | O96013 | PAK4 | Serine/threonine-protein kinase PAK 4 OS=Homo sapien | 0 | 191648.55 | 161670.12 | 277126.97 | 176227.22 | 149151.09 | 193903.03 | 208532.75 | **1.154443847** | 7 |
| sp\|P06576 | P06576 | ATP5F1B | ATP synthase subunit beta, mitochondrial OS=Homo sap | 387047872 | 368861664 | 353341952 | 400405952 | 449740320 | 430105216 | 435845376 | 426733408 | **1.15418523** | 0 |
| sp\|O1523! | O15235 | MRPS12 | 28S ribosomal protein S12, mitochondrial OS=Homo sap | 656063.06 | 1031017.44 | 561322.69 | 959630.88 | 1199060.62 | 672052.19 | 745753.12 | 1085297.12 | **1.154028595** | 4 |
| sp\|A6NHQ | A6NHQ2 | FBLL1 | rRNA/tRNA 2'-O-methyltransferase fibrillarin-like protein | 13328226 | 13316993 | 11530986 | 13398741 | 13955043 | 13228800 | 14545001 | 17783760 | **1.153905309** | 3 |
| sp\|Q96GD | Q96GD0 | PDXP | Pyridoxal phosphate phosphatase OS=Homo sapiens (Hu | 3451226.75 | 4431795 | 3564661.25 | 4256819.5 | 4858994 | 4188649.25 | 4433564 | 4621544 | **1.152710902** | 2 |
| sp\|A8MW | A8MWD9 | SNRPGP15 | Putative small nuclear ribonucleoprotein G-like protein 1 | 24784674 | 23745586 | 27767014 | 21656346 | 34734664 | 24565320 | 25148646 | 28390540 | **1.151965287** | 3 |
| sp\|P51610 | P51610 | HCFC1 | Host cell factor 1 OS=Homo sapiens (Human) GN=HCFC1 | 10472494 | 7448033 | 7975809.5 | 7289632.5 | 9272088 | 9629683 | 10801588 | 8525298 | **1.151952411** | 3 |
| sp\|Q8WW | Q8WWQ0 | PHIP | PH-interacting protein OS=Homo sapiens (Human) GN=P | 665447.94 | 940799.88 | 410762.12 | 655635.38 | 817696.25 | 730671.75 | 794804.44 | 734686.38 | **1.151615142** | 4 |
| sp\|Q9979! | Q99798 | ACO2 | Aconitate hydratase, mitochondrial OS=Homo sapiens (H | 23890604 | 24690094 | 22268976 | 23466268 | 28332388 | 28410036 | 28186952 | 23684280 | **1.151593821** | 2 |
| sp\|Q0221! | Q02218 | OGDH | 2-oxoglutarate dehydrogenase, mitochondrial OS=Homo | 6420742 | 7279612 | 5947732 | 7573480.5 | 7323256.5 | 8137745 | 6706709 | 9165380 | **1.151039214** | 3 |
| sp\|P08579 | P08579 | SNRPB2 | U2 small nuclear ribonucleoprotein B'' OS=Homo sapien | 27396488 | 25973802 | 26269972 | 25749802 | 32771056 | 26704612 | 33551712 | 28270858 | **1.150945672** | 1 |
| sp\|Q9UM! | Q9UMS4 | PRPF19 | Pre-mRNA-processing factor 19 OS=Homo sapiens (Hum | 46967320 | 42907632 | 46452760 | 41073612 | 51436080 | 47978248 | 52407456 | 52310676 | **1.150681716** | 0 |
| sp\|Q9UM\ | Q9UMY1 | NOL7 | Nucleolar protein 7 OS=Homo sapiens (Human) GN=NOL | 8692170 | 6150398 | 5624798.5 | 5964428 | 7892173 | 7779206.5 | 7512349 | 7223480 | **1.150402728** | 4 |
| sp\|O1495: | O14957 | UQCR11 | Cytochrome b-c1 complex subunit 10 OS=Homo sapiens | 1921127.88 | 2213195 | 2056685.88 | 2074213.88 | 2222362.75 | 2421496.25 | 2612256.25 | 2252197.5 | **1.150400076** | 0 |
| sp\|A0FGR! | A0FGR8 | ESYT2 | Extended synaptotagmin-2 OS=Homo sapiens (Human) ( | 3765034.5 | 3820730.5 | 3973924.25 | 3646544.75 | 4203287 | 4317026.5 | 4507236.5 | 4456611 | **1.14980218** | 0 |
| sp\|P22090 | P22090 | RPS4Y1 | 40S ribosomal protein S4, Y isoform 1 OS=Homo sapiens | 140074240 | 132831376 | 135102144 | 123462872 | 161455552 | 168968768 | 137432112 | 142670016 | **1.148749171** | 1 |
| sp\|Q9BQ7 | Q9BQ75 | CMSS1 | Protein CMSS1 OS=Homo sapiens (Human) GN=CMSS1 P | 377327.44 | 1059864.5 | 1840400.25 | 1855153.75 | 1466245.88 | 1182067.12 | 1856876.88 | 1390480.5 | **1.148638652** | 6 |
| sp\|P24539 | P24539 | ATP5PB | ATP synthase F(0) complex subunit B1, mitochondrial OS | 31759540 | 32604006 | 37439812 | 31979638 | 41138072 | 35334036 | 40464200 | 36608292 | **1.147713869** | 2 |
| sp\|Q70CQ | Q70CQ2 | USP34 | Ubiquitin carboxyl-terminal hydrolase 34 OS=Homo sapi | 1669209.12 | 1857533.88 | 2285221.5 | 1907895.75 | 1827538.25 | 2263732.5 | 2035213.25 | 2730197 | **1.14725924** | 5 |
| sp\|P22087 | P22087 | FBL | rRNA 2'-O-methyltransferase fibrillarin OS=Homo sapien | 45878788 | 45432968 | 41612192 | 45723788 | 53409500 | 51743896 | 48064260 | 51674272 | **1.1469047** | 0 |
| sp\|Q0796( | Q07960 | ARHGAP1 | Rho GTPase-activating protein 1 OS=Homo sapiens (Hum | 8224288 | 10406407 | 10184920 | 10711429 | 11494495 | 11334386 | 11043822 | 11446007 | **1.146524137** | 0 |
| sp\|P49406 | P49406 | MRPL19 | 39S ribosomal protein L19, mitochondrial OS=Homo sapi | 8222615.5 | 8565097 | 8028754.5 | 8751306 | 9052016 | 8329409.5 | 9890545 | 11203507 | **1.146202862** | 2 |
| sp\|Q86V4! | Q86V48 | LUZP1 | Leucine zipper protein 1 OS=Homo sapiens (Human) GN= | 604410.06 | 663309.88 | 595993.56 | 481174.62 | 0 | 1326870.62 | 1360293.25 | 0 | **1.145966772** | 8 |
| sp\|P61224 | P61224 | RAP1B | Ras-related protein Rap-1b OS=Homo sapiens (Human) ( | 23506680 | 18696452 | 15881560 | 23215956 | 29848148 | 23265940 | 19795748 | 20247320 | **1.145835344** | 5 |
| sp\|P61006 | P61006 | RAB8A | Ras-related protein Rab-8A OS=Homo sapiens (Human) ( | 3717520 | 3896545 | 4535310.5 | 3507275 | 4340974.5 | 5245682.5 | 4021421.5 | 4331548.5 | **1.145815128** | 3 |
| sp\|Q1290! | Q12905 | ILF2 | Interleukin enhancer-binding factor 2 OS=Homo sapiens | 94652928 | 89303344 | 96888296 | 95055592 | 98664592 | 107758768 | 104678296 | 119485568 | **1.14548295** | 0 |
| sp\|P60981 | P60981 | DSTN | Destrin OS=Homo sapiens (Human) GN=DSTN PE=1 SV=3 | 29817102 | 30890636 | 30557604 | 34508968 | 37940692 | 35933588 | 37097912 | 33070166 | **1.145244669** | 1 |
| sp\|P48729 | P48729 | CSNK1A1 | Casein kinase I isoform alpha OS=Homo sapiens (Human | 2079730 | 2008017.75 | 2044914.25 | 2458633.75 | 2359195 | 2620808.25 | 2521414.25 | 2332194.25 | **1.144601703** | 2 |
| sp\|P29218 | P29218 | IMPA1 | Inositol monophosphatase 1 OS=Homo sapiens (Human) | 6007123.5 | 6430855 | 4368999 | 6366162 | 6126011.5 | 7330368 | 6655521 | 6405496 | **1.144316095** | 3 |
| sp\|Q96EY4 | Q96EY4 | TMA16 | Translation machinery-associated protein 16 OS=Homo s | 2919637.25 | 1515196.62 | 3797915.5 | 1695546.38 | 3327593 | 1424377.75 | 3497836 | 3110278 | **1.144212968** | 7 |
| sp\|Q9H6Z4 | Q9H6Z4 | RANBP3 | Ran-binding protein 3 OS=Homo sapiens (Human) GN=R. | 9107394 | 8797985 | 8266306.5 | 9706008 | 10092127 | 9775894 | 11284001 | 9883907 | **1.143772773** | 0 |
| sp\|Q9UM) | Q9UMX0 | UBQLN1 | Ubiquilin-1 OS=Homo sapiens (Human) GN=UBQLN1 PE= | 3689485.5 | 3079167 | 3851517.5 | 3266443.25 | 3399562 | 4645401 | 3944552 | 3892841 | **1.143717025** | 2 |
| sp\|Q9NZI8 | Q9NZI8 | IGF2BP1 | Insulin-like growth factor 2 mRNA-binding protein 1 OS= | 216915744 | 212589936 | 227417344 | 200642976 | 265211488 | 232329968 | 252713264 | 230527088 | **1.143680846** | 0 |
| sp\|O1521: | O15213 | WDR46 | WD repeat-containing protein 46 OS=Homo sapiens (Hu | 3726708 | 3911480.5 | 3704160.5 | 2318668.25 | 4228777.5 | 4126952.75 | 3008290.5 | 4251624 | **1.14308067** | 3 |
| sp\|Q8WU\ | Q8WUY1 | THEM6 | Protein THEM6 OS=Homo sapiens (Human) GN=THEM6 I | 4161015 | 4211966.5 | 3916810 | 5616521 | 4722058 | 4231981 | 4664430.5 | 6846243 | **1.142876988** | 3 |
| sp\|Q6PJT7 | Q6PJT7 | ZC3H14 | Zinc finger CCCH domain-containing protein 14 OS=Hom | 955610.19 | 1054496.25 | 1514908.12 | 971519.5 | 1223897.62 | 1052898.5 | 1414093.5 | 1446979.88 | **1.142628841** | 5 |
| sp\|P37108 | P37108 | SRP14 | Signal recognition particle 14 kDa protein OS=Homo sapi | 24110144 | 18672142 | 17251138 | 27881684 | 23910316 | 23960240 | 27708034 | 24870168 | **1.142565371** | 6 |
| sp\|P21283 | P21283 | ATP6V1C1 | V-type proton ATPase subunit C 1 OS=Homo sapiens (Hu | 1517518.12 | 2372934 | 2358353.5 | 2332796.5 | 2005442.38 | 2824627.75 | 1834693.12 | 3139988.25 | **1.142531588** | 6 |
| sp\|Q1564: | Q15642 | TRIP10 | Cdc42-interacting protein 4 OS=Homo sapiens (Human) ( | 356628.09 | 1952284.25 | 357709.97 | 427394.94 | 1590219.25 | 401863.94 | 355078.25 | 1185542.62 | **1.141785509** | 8 |
| sp\|Q1318! | Q13185 | CBX3 | Chromobox protein homolog 3 OS=Homo sapiens (Huma | 29439026 | 24586346 | 28261884 | 24187328 | 31332168 | 27741338 | 30097440 | 32287090 | **1.140723273** | 2 |
| sp\|P78329 | P78329 | CYP4F2 | Phylloquinone omega-hydroxylase CYP4F2 OS=Homo sa | 171823.64 | 216191.27 | 218708.08 | 243873.41 | 224445.59 | 235202.92 | 250408.45 | 259903.25 | **1.14032955** | 2 |
| sp\|Q9H1Y( | Q9H1Y0 | ATG5 | Autophagy protein 5 OS=Homo sapiens (Human) GN=AT | 619518.69 | 436147.94 | 565142.62 | 529277.38 | 438499.66 | 773437.25 | 795435.25 | 444082.44 | **1.140165501** | 6 |
| sp\|Q9UDR | Q9UDR5 | AASS | Alpha-aminoadipic semialdehyde synthase, mitochondri | 1674716.25 | 1477005.12 | 1276996.25 | 1764413 | 1837025.75 | 1173635 | 1626210.62 | 2414607 | **1.138596746** | 6 |
| sp\|P39656 | P39656 | DDOST | Dolichyl-diphosphooligosaccharide--protein glycosyltran | 74025944 | 84632112 | 85611472 | 77130128 | 84361928 | 88611208 | 93614560 | 99316968 | **1.138472482** | 2 |
| sp\|Q96C3( | Q96C36 | PYCR2 | Pyrroline-5-carboxylate reductase 2 OS=Homo sapiens (I | 16926528 | 15210733 | 15272858 | 19502150 | 20029732 | 18988938 | 18090790 | 19059344 | **1.138338382** | 3 |

| | | | | | | | | | | | | | |
|---|---|---|---|---|---|---|---|---|---|---|---|---|---|
| sp\|P51571 | P51571 | SSR4 | Translocon-associated protein subunit delta OS=Homo s | 33413764 | 31416872 | 30723740 | 33748316 | 39560240 | 38480376 | 36959372 | 32181710 | **1.138272496** | 2 |
| sp\|P67936 | P67936 | TPM4 | Tropomyosin alpha-4 chain OS=Homo sapiens (Human) ( | 123265352 | 118007056 | 113539944 | 117657256 | 147750176 | 104196864 | 168511744 | 117295440 | **1.138177387** | 7 |
| sp\|Q9956! | Q99569 | PKP4 | Plakophilin-4 OS=Homo sapiens (Human) GN=PKP4 PE=1 | 507269.38 | 656630 | 494952.44 | 542677.12 | 571390.25 | 633116.38 | 695671.62 | 605315.88 | **1.138070041** | 3 |
| sp\|Q0083! | Q00839 | HNRNPU | Heterogeneous nuclear ribonucleoprotein U OS=Homo s | 426907200 | 427381536 | 446566592 | 370838400 | 466654304 | 453707872 | 516159264 | 465457760 | **1.137755779** | 0 |
| sp\|Q9UIA! | Q9UIA9 | XPO7 | Exportin-7 OS=Homo sapiens (Human) GN=XPO7 PE=1 S\ | 2428598.5 | 4305107 | 4723589.5 | 4249997 | 5030215 | 4268341 | 4531872 | 4037081.75 | **1.13752961** | 6 |
| sp\|Q1315! | Q13155 | AIMP2 | Aminoacyl tRNA synthase complex-interacting multifunct | 7469643 | 7570777 | 7313053 | 8294334.5 | 8701558 | 9411208 | 9103499 | 7618522 | **1.136615955** | 1 |
| sp\|Q9ULP! | Q9ULP9 | TBC1D24 | TBC1 domain family member 24 OS=Homo sapiens (Hum | 0 | 1094231.75 | 0 | 783347.19 | 1205819 | 928025.44 | 0 | 0 | **1.136487204** | 7 |
| sp\|P14314 | P14314 | PRKCSH | Glucosidase 2 subunit beta OS=Homo sapiens (Human) ( | 54691744 | 46829560 | 45597220 | 35938980 | 53083308 | 51365704 | 49860124 | 53688864 | **1.136244051** | 4 |
| sp\|P09001 | P09001 | MRPL3 | 39S ribosomal protein L3, mitochondrial OS=Homo sapie | 6596537 | 6511447 | 6595913 | 7002186.5 | 8281163.5 | 7355141 | 6774392 | 7932783 | **1.136201027** | 1 |
| sp\|Q9Y5S! | Q9Y5S9 | RBM8A | RNA-binding protein 8A OS=Homo sapiens (Human) GN= | 5353471 | 7391386 | 7817819.5 | 9530864 | 9609342 | 7754230 | 8519038 | 8304064 | **1.13601369** | 4 |
| sp\|P15311 | P15311 | EZR | Ezrin OS=Homo sapiens (Human) GN=EZR PE=1 SV=4 | 93964400 | 94852520 | 113682176 | 90188024 | 111546088 | 96256904 | 126260904 | 111807064 | **1.135435662** | 3 |
| sp\|Q9BV1( | Q9BV10 | ALG12 | Dol-P-Man:Man(7)GlcNAc(2)-PP-Dol alpha-1,6-mannosyl | 239539.05 | 270551.16 | 289023.22 | 248248.59 | 227399.61 | 332898.41 | 275107.94 | 353774.25 | **1.13540513** | 5 |
| sp\|P20645 | P20645 | M6PR | Cation-dependent mannose-6-phosphate receptor OS=H | 2308657.75 | 2081138.12 | 1930398.88 | 2102397.75 | 2247038 | 2648497.5 | 2687322.75 | 1979105.12 | **1.135275554** | 4 |
| sp\|Q9253! | Q92538 | GBF1 | Golgi-specific brefeldin A-resistance guanine nucleotide ( | 6099817.5 | 4522561.5 | 3529058.75 | 4291078 | 5811507 | 5266458.5 | 4516690 | 5336902 | **1.134962159** | 5 |
| sp\|Q9H9Q | Q9H9Q2 | COPS7B | COP9 signalosome complex subunit 7b OS=Homo sapien | 8643873 | 9569753 | 8059530.5 | 9347960 | 11804052 | 10880414 | 10850717 | 6892809.5 | **1.134944563** | 4 |
| sp\|P08240 | P08240 | SRPRA | Signal recognition particle receptor subunit alpha OS=Hc | 11695213 | 11668023 | 12226756 | 11124722 | 13399627 | 9353418 | 14102842 | 16152374 | **1.134723013** | 4 |
| sp\|P42356 | P42356 | PI4KA | Phosphatidylinositol 4-kinase alpha OS=Homo sapiens (H | 202273.25 | 206686.62 | 187752.33 | 196249.83 | 196990.75 | 226293.31 | 263104.28 | 213402.84 | **1.134721646** | 2 |
| sp\|P25398 | P25398 | RPS12 | 40S ribosomal protein S12 OS=Homo sapiens (Human) G | 38336020 | 35079228 | 33881748 | 33778560 | 42875172 | 35055924 | 40983932 | 41070088 | **1.134038529** | 2 |
| sp\|P50895 | P50895 | BCAM | Basal cell adhesion molecule OS=Homo sapiens (Human) | 32608376 | 36037304 | 30382716 | 36252824 | 43461520 | 39119448 | 34302020 | 36485948 | **1.13370456** | 2 |
| sp\|P48426 | P48426 | PIP4K2A | Phosphatidylinositol 5-phosphate 4-kinase type-2 alpha | 1190197.88 | 841743.38 | 1289200.88 | 870868.69 | 1185610 | 1177768.88 | 935223.31 | 1452492 | **1.133368778** | 6 |
| sp\|P11182 | P11182 | DBT | Lipoamide acyltransferase component of branched-chair | 984696.88 | 733763.69 | 601874.94 | 762790.69 | 1001517.62 | 861992.38 | 994758.94 | 633900.25 | **1.132671504** | 4 |
| sp\|Q1465: | Q14653 | IRF3 | Interferon regulatory factor 3 OS=Homo sapiens (Humar | 1430900.25 | 684355.5 | 700758.56 | 1446425.88 | 519214.62 | 1424014.25 | 1332351 | 1549086.5 | **1.131902421** | 8 |
| sp\|Q8NC6( | Q8NC60 | NOA1 | Nitric oxide-associated protein 1 OS=Homo sapiens (Hun | 449631.75 | 510467.91 | 1029951.25 | 1073352.25 | 485962.94 | 574980.75 | 1191216.25 | 1215227.62 | **1.131874382** | 5 |
| sp\|P60866 | P60866 | RPS20 | 40S ribosomal protein S20 OS=Homo sapiens (Human) G | 19904522 | 13516074 | 7915224 | 13397265 | 17783728 | 17054184 | 13169468 | 13939080 | **1.131791859** | 6 |
| sp\|Q9Y2R! | Q9Y2R5 | MRPS17 | 28S ribosomal protein S17, mitochondrial OS=Homo sap | 5221848 | 6286407 | 5269890.5 | 5570575.5 | 5970150.5 | 5610809 | 6694001 | 7013029.5 | **1.131518443** | 2 |
| sp\|P14678 | P14678 | SNRPB | Small nuclear ribonucleoprotein-associated proteins B ar | 44882036 | 44427672 | 47438924 | 50288696 | 53570736 | 53433412 | 54571412 | 50016644 | **1.131283291** | 1 |
| sp\|Q8N2U | Q8N2U0 | TMEM256 | Transmembrane protein 256 OS=Homo sapiens (Human) | 4760254 | 5026691.5 | 3512315.5 | 4140876.5 | 5374573 | 5470642.5 | 4410552 | 4473335 | **1.131246958** | 4 |
| sp\|Q1536: | Q15363 | TMED2 | Transmembrane emp24 domain-containing protein 2 OS | 1855511 | 2266598.75 | 1985151.75 | 2777741 | 4035893.5 | 2630648.75 | 1797560.25 | 1586243.38 | **1.131158475** | 7 |
| sp\|Q9UIV1 | Q9UIV1 | CNOT7 | CCR4-NOT transcription complex subunit 7 OS=Homo sa | 794277.81 | 787047.94 | 1013091.5 | 891877.06 | 895237.69 | 892445.88 | 1049996.75 | 1103925.38 | **1.130600388** | 2 |
| sp\|P07737 | P07737 | PFN1 | Profilin-1 OS=Homo sapiens (Human) GN=PFN1 PE=1 SV: | 174011584 | 164681248 | 171886240 | 182915104 | 207393824 | 196789040 | 190670880 | 189145968 | **1.130506555** | 0 |
| sp\|P10253 | P10253 | GAA | Lysosomal alpha-glucosidase OS=Homo sapiens (Human | 18883776 | 18035978 | 12592033 | 17637074 | 17826416 | 17509784 | 19319408 | 21248090 | **1.130379531** | 5 |
| sp\|O1495( | O14950 | MYL12B | Myosin regulatory light chain 12B OS=Homo sapiens (Hu | 14134570 | 13438556 | 12526872 | 16509047 | 15916950 | 13723221 | 17369592 | 16925300 | **1.129414266** | 3 |
| sp\|Q8N44 | Q8N442 | GUF1 | Translation factor GUF1, mitochondrial OS=Homo sapier | 586224.31 | 662636.31 | 539723.38 | 0 | 410798.03 | 594292 | 520111.97 | 494526.75 | **1.129233377** | 6 |
| sp\|Q96S5! | Q96S55 | WRNIP1 | ATPase WRNIP1 OS=Homo sapiens (Human) GN=WRNIP | 3155573.5 | 2981268.5 | 4151158.5 | 3359446.5 | 2576145.75 | 4751575 | 4001277.25 | 4078894.25 | **1.128994474** | 6 |
| sp\|Q99471 | Q99471 | PFDN5 | Prefoldin subunit 5 OS=Homo sapiens (Human) GN=PFDI | 4509252.5 | 3655001.5 | 3767458 | 3914469.5 | 4532534.5 | 4332226.5 | 5132813 | 3890966 | **1.128886477** | 3 |
| sp\|Q5T65: | Q5T653 | MRPL2 | 39S ribosomal protein L2, mitochondrial OS=Homo sapie | 4244069 | 3645891.75 | 2528704.75 | 3789929.75 | 5173564 | 4480197 | 4878663.5 | 1505933.88 | **1.128778609** | 4 |
| sp\|Q1312: | Q13123 | IK | Protein Red OS=Homo sapiens (Human) GN=IK PE=1 SV= | 2668516.25 | 2192610.5 | 3136678.75 | 2317785.75 | 3595935.5 | 2575468 | 2965457.75 | 2505013 | **1.128570721** | 5 |
| sp\|Q9H9J2 | Q9H9J2 | MRPL44 | 39S ribosomal protein L44, mitochondrial OS=Homo sapi | 4570620 | 4385932 | 3995439 | 4303967 | 5006687.5 | 4594365.5 | 5231085.5 | 4642418.5 | **1.128570028** | 0 |
| sp\|Q9NTJ! | Q9NTJ5 | SACM1L | Phosphatidylinositide phosphatase SAC1 OS=Homo sapi | 4443838 | 4636351.5 | 4251143.5 | 4671156 | 4673267 | 4787681 | 5008359.5 | 5838534.5 | **1.128057459** | 0 |
| sp\|Q1542 | Q15427 | SF3B4 | Splicing factor 3B subunit 4 OS=Homo sapiens (Human) ( | 3007169.25 | 3534437.75 | 2625085.75 | 3945813 | 3982380.5 | 3401594.5 | 3939727.25 | 3463239 | **1.127697599** | 5 |
| sp\|P05091 | P05091 | ALDH2 | Aldehyde dehydrogenase, mitochondrial OS=Homo sapi | 253280512 | 247661712 | 266616400 | 270560960 | 307468160 | 292550912 | 281917920 | 288423040 | **1.1273846** | 0 |
| sp\|O1454 | O14548 | COX7A2L | Cytochrome c oxidase subunit 7A-related protein, mitoch | 1278600.25 | 1374854.62 | 1624064 | 1220583.75 | 1635163.75 | 1300124.25 | 1452833.62 | 1809655 | **1.127257356** | 3 |
| sp\|Q86YN | Q86YN1 | DOLPP1 | Dolichyldiphosphatase 1 OS=Homo sapiens (Human) GN | 732912.62 | 794546.81 | 678499.94 | 1053575.25 | 804993.62 | 1041566.5 | 846585.88 | 981144.75 | **1.127243971** | 4 |
| sp\|Q9UIQ( | Q9UIQ6 | LNPEP | Leucyl-cystinyl aminopeptidase OS=Homo sapiens (Hum | 1764051 | 2115151.5 | 1640158.25 | 1894832.25 | 1920507.38 | 1957603.38 | 1982834.5 | 2492787.5 | **1.126721783** | 3 |
| sp\|P35080 | P35080 | PFN2 | Profilin-2 OS=Homo sapiens (Human) GN=PFN2 PE=1 SV: | 9623156 | 8079357 | 9153658 | 8347037 | 10101872 | 9505868 | 10305174 | 9748051 | **1.126629283** | 1 |

| | | | | | | | | | | | | |
|---|---|---|---|---|---|---|---|---|---|---|---|---|
| sp\|O60547 | O60547 | GMDS | GDP-mannose 4,6 dehydratase OS=Homo sapiens (Huma | 34523272 | 34287640 | 33149610 | 31092844 | 41306152 | 38795188 | 33402316 | 36353296 | 1.126292077 | 2 |
| sp\|Q13409 | Q13409 | DYNC1I2 | Cytoplasmic dynein 1 intermediate chain 2 OS=Homo sa | 8341453.5 | 8606058 | 11473680 | 7646527 | 10485354 | 9692176 | 10515995 | 9907053 | 1.125676358 | 4 |
| sp\|P62487 | P62487 | POLR2G | DNA-directed RNA polymerase II subunit RPB7 OS=Homo | 1296768.25 | 1765023.25 | 1886454.5 | 1935312 | 2215185.5 | 1927613.75 | 1463538.62 | 2139361.25 | 1.125246438 | 4 |
| sp\|P07954 | P07954 | FH | Fumarate hydratase, mitochondrial OS=Homo sapiens (H | 24973322 | 24645942 | 27694444 | 25565546 | 29109950 | 26989172 | 31965192 | 27687672 | 1.125124663 | 2 |
| sp\|Q14318 | Q14318 | FKBP8 | Peptidyl-prolyl cis-trans isomerase FKBP8 OS=Homo sapi | 3142205 | 2159428.5 | 3175817 | 2709641.5 | 3361920 | 4054631 | 2968968.75 | 2198842 | 1.124900175 | 5 |
| sp\|O60493 | O60493 | SNX3 | Sorting nexin-3 OS=Homo sapiens (Human) GN=SNX3 PE | 4950122.5 | 4117392 | 4078172.5 | 3475153 | 6425280 | 5017782.5 | 3989704.5 | 3263995.5 | 1.124898772 | 7 |
| sp\|Q12959 | Q12959 | DLG1 | Disks large homolog 1 OS=Homo sapiens (Human) GN=D | 4552109 | 3975867.75 | 4800013 | 4002355.75 | 5043459 | 4618978.5 | 4801219 | 5019170 | 1.124203006 | 1 |
| sp\|Q9Y5Q | Q9Y5Q9 | GTF3C3 | General transcription factor 3C polypeptide 3 OS=Homo | 1479921 | 2388083.75 | 2561376.5 | 2153332.25 | 1818750.5 | 2393315.5 | 2097027.75 | 3335428.75 | 1.123714837 | 7 |
| sp\|Q9GZT8 | Q9GZT8 | NIF3L1 | NIF3-like protein 1 OS=Homo sapiens (Human) GN=NIF3I | 1306440.75 | 1884275.12 | 2123312.75 | 1693732.5 | 2184616 | 1730073.38 | 2399403.5 | 1559011 | 1.123483484 | 5 |
| sp\|Q9BZZ5 | Q9BZZ5 | API5 | Apoptosis inhibitor 5 OS=Homo sapiens (Human) GN=AP | 21551746 | 21029778 | 18880810 | 20341556 | 24923270 | 22687680 | 23866038 | 20401166 | 1.123151405 | 2 |
| sp\|Q9Y3D9 | Q9Y3D9 | MRPS23 | 28S ribosomal protein S23, mitochondrial OS=Homo sap | 9308150 | 8589591 | 8378445 | 9607850 | 9519543 | 8637914 | 10331948 | 11806712 | 1.122953867 | 3 |
| sp\|P14923 | P14923 | JUP | Junction plakoglobin OS=Homo sapiens (Human) GN=JUI | 32265508 | 66029676 | 33643576 | 30637228 | 44842544 | 42542440 | 45899336 | 49205192 | 1.122487486 | 4 |
| sp\|P00734 | P00734 | F2 | Prothrombin OS=Homo sapiens (Human) GN=F2 PE=1 SV | 7524674.5 | 6760475.5 | 5447952 | 6863685.5 | 7579544 | 7744850.5 | 7325587 | 7188012 | 1.121864567 | 2 |
| sp\|Q9NVA | Q9NVA1 | UQCC1 | Ubiquinol-cytochrome-c reductase complex assembly fac | 647186.19 | 841463.25 | 877635.25 | 726300.06 | 912429.69 | 881110.31 | 704452.88 | 969869.31 | 1.121347504 | 3 |
| sp\|P63220 | P63220 | RPS21 | 40S ribosomal protein S21 OS=Homo sapiens (Human) G | 14606713 | 12664887 | 13026111 | 14492366 | 17348210 | 14801392 | 17835942 | 11440544 | 1.121117023 | 4 |
| sp\|O00161 | O00161 | SNAP23 | Synaptosomal-associated protein 23 OS=Homo sapiens ( | 2707659 | 2647143.25 | 2283385.75 | 2392740.5 | 3741533.5 | 2755548.5 | 1853974.75 | 2892536.5 | 1.120892572 | 4 |
| sp\|Q96KA5 | Q96KA5 | CLPTM1L | Cleft lip and palate transmembrane protein 1-like proteir | 741499.31 | 850075.56 | 726372.69 | 802306.75 | 802298.25 | 1141795.62 | 854555.75 | 697527.75 | 1.12047834 | 6 |
| sp\|Q5JVF3 | Q5JVF3 | PCID2 | PCI domain-containing protein 2 OS=Homo sapiens (Hun | 2917796.25 | 2963681 | 2508563 | 3211428.5 | 3030325 | 3507270.5 | 2961634.5 | 3499314 | 1.120422274 | 3 |
| sp\|O00159 | O00159 | MYO1C | Unconventional myosin-Ic OS=Homo sapiens (Human) G | 19468538 | 20739054 | 17337200 | 20419232 | 20669668 | 22061574 | 21104908 | 23507260 | 1.120304026 | 1 |
| sp\|P58511 | P58511 | SMIM11A | Small integral membrane protein 11A OS=Homo sapiens | 115825.16 | 149423.66 | 169730 | 137403.38 | 129777.71 | 157991.45 | 169283.11 | 184135.89 | 1.120209818 | 5 |
| sp\|O43670 | O43670 | ZNF207 | BUB3-interacting and GLEBS motif-containing protein ZN | 6396050 | 7734184 | 8572738 | 5542067.5 | 9249372 | 7457602 | 9305620 | 5604956 | 1.119401869 | 5 |
| sp\|P02545 | P02545 | LMNA | Prelamin-A/C OS=Homo sapiens (Human) GN=LMNA PE= | 220600752 | 253748704 | 270129824 | 227307888 | 278248512 | 248488672 | 279648256 | 281061568 | 1.119017665 | 2 |
| sp\|P11217 | P11217 | PYGM | Glycogen phosphorylase, muscle form OS=Homo sapiens | 17012318 | 18715312 | 19711374 | 19365840 | 20112212 | 19043150 | 19570432 | 24945526 | 1.1185281 | 3 |
| sp\|Q04837 | Q04837 | SSBP1 | Single-stranded DNA-binding protein, mitochondrial OS= | 21194388 | 22975394 | 22008572 | 24526808 | 23205256 | 23489448 | 26969324 | 27764208 | 1.118219005 | 2 |
| sp\|Q9BW7 | Q9BW72 | HIGD2A | HIG1 domain family member 2A, mitochondrial OS=Hom | 5326261 | 4559365 | 4377641 | 6928977.5 | 5194978 | 7101756.5 | 4886150 | 6511989.5 | 1.118091762 | 5 |
| sp\|Q9BYD | Q9BYD1 | MRPL13 | 39S ribosomal protein L13, mitochondrial OS=Homo sapi | 1638797.88 | 1461607.75 | 1518086.62 | 1151739.12 | 1686821.25 | 1642948.38 | 1380694.12 | 1737656.38 | 1.117480343 | 3 |
| sp\|Q9Y3A6 | Q9Y3A6 | TMED5 | Transmembrane emp24 domain-containing protein 5 OS | 640384.44 | 762858.38 | 85682.02 | 749455.25 | 479449.59 | 1018451.38 | 424269.12 | 579159.62 | 1.117473177 | 7 |
| sp\|P09493 | P09493 | TPM1 | Tropomyosin alpha-1 chain OS=Homo sapiens (Human) ( | 93660840 | 89624216 | 86685392 | 93874088 | 117142920 | 73209064 | 126179160 | 89897208 | 1.117038493 | 6 |
| sp\|Q9H9Y2 | Q9H9Y2 | RPF1 | Ribosome production factor 1 OS=Homo sapiens (Humai | 0 | 338694.41 | 596049.25 | 487548.62 | 442576.09 | 332326.59 | 391192.5 | 422337.66 | 1.116811827 | 7 |
| sp\|Q13595 | Q13595 | TRA2A | Transformer-2 protein homolog alpha OS=Homo sapiens | 4441971 | 1856562 | 4733142 | 2510229.25 | 2465507.75 | 1986542 | 5198239 | 5472093 | 1.116710137 | 6 |
| sp\|Q99959 | Q99959 | PKP2 | Plakophilin-2 OS=Homo sapiens (Human) GN=PKP2 PE=1 | 7277969 | 8000693 | 7448114.5 | 8007576.5 | 9587457 | 10190645 | 7963263.5 | 6579554 | 1.116695689 | 6 |
| sp\|Q68CQ | Q68CQ7 | GLT8D1 | Glycosyltransferase 8 domain-containing protein 1 OS=H | 973043.94 | 929050.88 | 917586.25 | 906030 | 1055211.38 | 703097.75 | 1333073.5 | 1068958.62 | 1.116656974 | 4 |
| sp\|P45974 | P45974 | USP5 | Ubiquitin carboxyl-terminal hydrolase 5 OS=Homo sapie | 29222014 | 29807514 | 21959132 | 29522758 | 26557778 | 33507566 | 29189322 | 34115904 | 1.116360393 | 6 |
| sp\|P02771 | P02771 | AFP | Alpha-fetoprotein OS=Homo sapiens (Human) GN=AFP P | 30484792 | 36087932 | 35431652 | 34473636 | 33327432 | 29024618 | 45156508 | 44827328 | 1.116193618 | 7 |
| sp\|Q9Y224 | Q9Y224 | RTRAF | RNA transcription, translation and transport factor prote | 13452236 | 12463001 | 10592556 | 13460154 | 14969130 | 14815869 | 14302990 | 11671192 | 1.115898978 | 3 |
| sp\|O95793 | O95793 | STAU1 | Double-stranded RNA-binding protein Staufen homolog 1 | 14032913 | 13188960 | 20182134 | 14831113 | 19431320 | 18215478 | 15744379 | 16048853 | 1.115769199 | 4 |
| sp\|P14866 | P14866 | HNRNPL | Heterogeneous nuclear ribonucleoprotein L OS=Homo sa | 157379392 | 162343520 | 150937344 | 151553552 | 177143248 | 167884976 | 182323440 | 166700144 | 1.11545549 | 0 |
| sp\|Q8NBN | Q8NBN7 | RDH13 | Retinol dehydrogenase 13 OS=Homo sapiens (Human) G | 1987706.25 | 1582086.12 | 1048608.25 | 2355294.75 | 1811678.12 | 1872768.25 | 2008311.12 | 2082159 | 1.114891901 | 6 |
| sp\|P60174 | P60174 | TPI1 | Triosephosphate isomerase OS=Homo sapiens (Human) | 491603552 | 449441728 | 413048992 | 469407616 | 529247872 | 558114496 | 500525344 | 444981280 | 1.114815951 | 3 |
| sp\|O94826 | O94826 | TOMM70 | Mitochondrial import receptor subunit TOM70 OS=Hom | 20003102 | 19437158 | 19876098 | 19503248 | 21380558 | 20708714 | 22979598 | 22778842 | 1.114541374 | 0 |
| sp\|Q9Y6M | Q9Y6M1 | IGF2BP2 | Insulin-like growth factor 2 mRNA-binding protein 2 OS= | 57641076 | 55638052 | 50865576 | 49946652 | 65732228 | 57070032 | 61122244 | 54497848 | 1.113647727 | 3 |
| sp\|Q14191 | Q14191 | WRN | Werner syndrome ATP-dependent helicase OS=Homo sa | 1001461.69 | 1401775.25 | 1997846.38 | 1811914.5 | 1319447.88 | 2220001 | 1543658.25 | 1835944.12 | 1.113641345 | 6 |
| sp\|Q49A26 | Q49A26 | GLYR1 | Putative oxidoreductase GLYR1 OS=Homo sapiens (Huma | 2122973 | 2202830.5 | 1967351.5 | 1808349.12 | 2301685.5 | 2007532 | 2594178 | 2114878.75 | 1.113160484 | 4 |
| sp\|P54577 | P54577 | YARS | Tyrosine--tRNA ligase, cytoplasmic OS=Homo sapiens (H | 42629012 | 50535408 | 40406608 | 50399880 | 51978608 | 49057948 | 52032176 | 51670276 | 1.112887957 | 2 |
| sp\|P61964 | P61964 | WDR5 | WD repeat-containing protein 5 OS=Homo sapiens (Hum | 4269088 | 4398313 | 2575268.25 | 4208281 | 4272716 | 4231368 | 4151412 | 4538801.5 | 1.11283107 | 6 |

| | | | | | | | | | | | | |
|---|---|---|---|---|---|---|---|---|---|---|---|---|
| sp\|O1497 | O14976 | GAK | Cyclin-G-associated kinase OS=Homo sapiens (Human) G | 414911.78 | 346887.5 | 330419.97 | 374118 | 351237.34 | 439165.78 | 417165.19 | 424026.25 | **1.112700751** | 2 |
| sp\|O0029 | O00299 | CLIC1 | Chloride intracellular channel protein 1 OS=Homo sapier | 208282560 | 211039984 | 195802016 | 205496800 | 230798416 | 204127376 | 231787696 | 246371168 | **1.112674737** | 3 |
| sp\|P51003 | P51003 | PAPOLA | Poly(A) polymerase alpha OS=Homo sapiens (Human) GI | 6047595 | 4441290.5 | 5035025.5 | 4376603 | 5049596 | 6483520.5 | 5777593.5 | 4830415 | **1.112590609** | 4 |
| sp\|Q9BT7 | Q9BT73 | PSMG3 | Proteasome assembly chaperone 3 OS=Homo sapiens (H | 1420228.25 | 1628161.5 | 1580375.38 | 1408423.88 | 1662998.5 | 1854660.25 | 1706198.5 | 1493046.75 | **1.112587992** | 2 |
| sp\|Q1284 | Q12846 | STX4 | Syntaxin-4 OS=Homo sapiens (Human) GN=STX4 PE=1 SV | 796694.12 | 539403.56 | 692337.5 | 427543.56 | 928372.81 | 819813.06 | 510206.66 | 473606.91 | **1.112387251** | 6 |
| sp\|Q9Y3B | Q9Y3B7 | MRPL11 | 39S ribosomal protein L11, mitochondrial OS=Homo sapi | 11700464 | 10852857 | 9581991 | 11591937 | 12758644 | 9019601 | 14766686 | 12081578 | **1.11204135** | 4 |
| sp\|Q0621 | Q06210 | GFPT1 | Glutamine--fructose-6-phosphate aminotransferase [isor | 67608600 | 69053152 | 61799436 | 61731876 | 70254176 | 65452392 | 76056400 | 77533968 | **1.111854911** | 2 |
| sp\|P82930 | P82930 | MRPS34 | 28S ribosomal protein S34, mitochondrial OS=Homo sap | 7203775 | 8083830 | 8387507 | 9042908 | 8980096 | 7877831 | 9458967 | 10059860 | **1.111826266** | 4 |
| sp\|Q6IN85 | Q6IN85 | PPP4R3A | Serine/threonine-protein phosphatase 4 regulatory subu | 185051.62 | 657749.75 | 720764.25 | 420654.59 | 533154.75 | 633891.12 | 462211.47 | 576503.56 | **1.111651262** | 8 |
| sp\|P60468 | P60468 | SEC61B | Protein transport protein Sec61 subunit beta OS=Homo | 26092634 | 28661974 | 32941656 | 27925936 | 41726664 | 29548484 | 27620004 | 29612356 | **1.1114432** | 5 |
| sp\|P31946 | P31946 | YWHAB | 14-3-3 protein beta/alpha OS=Homo sapiens (Human) G | 192204752 | 211683072 | 269467232 | 223123008 | 256192288 | 231090544 | 249187536 | 259491424 | **1.110971737** | 4 |
| sp\|Q1290 | Q12906 | ILF3 | Interleukin enhancer-binding factor 3 OS=Homo sapiens | 116219512 | 127989584 | 111245256 | 114935968 | 135501536 | 122585872 | 128608216 | 135391216 | **1.109901326** | 1 |
| sp\|Q1541 | Q15418 | RPS6KA1 | Ribosomal protein S6 kinase alpha-1 OS=Homo sapiens ( | 35366984 | 38442548 | 38097568 | 38934220 | 41414080 | 39409740 | 43246008 | 43335964 | **1.109813889** | 0 |
| sp\|Q9H91 | Q9H910 | JPT2 | Jupiter microtubule associated homolog 2 OS=Homo sap | 2554040.25 | 1890749.88 | 2484579.5 | 2268263.5 | 2722206 | 2540757 | 2697467.5 | 2245700.25 | **1.109647515** | 4 |
| sp\|Q8TBF | Q8TBF2 | PRXL2B | Prostamide/prostaglandin F synthase OS=Homo sapiens | 322162.41 | 366876.66 | 424248.53 | 344509.81 | 301769.59 | 479944.22 | 401889.53 | 433625.09 | **1.109364318** | 5 |
| sp\|Q9Y5K | Q9Y5K6 | CD2AP | CD2-associated protein OS=Homo sapiens (Human) GN= | 3485563.25 | 3533647 | 2605864.75 | 3521246.5 | 3547581.5 | 4374994 | 4098110.5 | 2553855.75 | **1.108640295** | 4 |
| sp\|Q9276 | Q92769 | HDAC2 | Histone deacetylase 2 OS=Homo sapiens (Human) GN=H | 7225878 | 9898503 | 10976926 | 7646443 | 10849845 | 9003571 | 10905941 | 8871112 | **1.10861436** | 6 |
| sp\|P27694 | P27694 | RPA1 | Replication protein A 70 kDa DNA-binding subunit OS=H | 15218895 | 16719001 | 16157239 | 16351850 | 18560922 | 14837433 | 18270720 | 19729580 | **1.107866489** | 4 |
| sp\|Q9Y23 | Q9Y237 | PIN4 | Peptidyl-prolyl cis-trans isomerase NIMA-interacting 4 O | 735780.19 | 743695.88 | 725909 | 756345.81 | 695204.81 | 718044.44 | 1046249.69 | 820956.56 | **1.107614308** | 8 |
| sp\|P21549 | P21549 | AGXT | Serine--pyruvate aminotransferase OS=Homo sapiens (H | 6653041 | 6473344.5 | 6063704 | 6305617 | 6074689 | 6664272.5 | 6632543 | 8852255 | **1.107000487** | 4 |
| sp\|Q9HD3 | Q9HD33 | MRPL47 | 39S ribosomal protein L47, mitochondrial OS=Homo sapi | 3328650.5 | 2728100.5 | 2566788.5 | 3571731.5 | 3552545 | 3511933.25 | 3298741 | 3135676 | **1.106895882** | 6 |
| sp\|P13716 | P13716 | ALAD | Delta-aminolevulinic acid dehydratase OS=Homo sapien | 1700369 | 1536644.38 | 1513529.25 | 1397301.5 | 1753862.5 | 1569118.5 | 1792202.62 | 1689255.75 | **1.106800892** | 2 |
| sp\|Q9UID | Q9UID3 | VPS51 | Vacuolar protein sorting-associated protein 51 homolog | 256393.78 | 357006.88 | 426204.47 | 380825.69 | 282222.97 | 439197.03 | 424955.22 | 425710.5 | **1.106766833** | 5 |
| sp\|Q9962 | Q99623 | PHB2 | Prohibitin-2 OS=Homo sapiens (Human) GN=PHB2 PE=1 | 131158632 | 146063344 | 147554048 | 142014752 | 166417328 | 146664272 | 157007120 | 157056448 | **1.106484429** | 1 |
| sp\|P62750 | P62750 | RPL23A | 60S ribosomal protein L23a OS=Homo sapiens (Human) | 91360136 | 77444008 | 71518856 | 81778936 | 99530624 | 94068104 | 87768600 | 74937152 | **1.106185465** | 4 |
| sp\|Q9Y2X | Q9Y2X3 | NOP58 | Nucleolar protein 58 OS=Homo sapiens (Human) GN=NC | 26130610 | 30526828 | 33936504 | 26430154 | 32272496 | 29796800 | 32309012 | 35029900 | **1.105825317** | 4 |
| sp\|P48634 | P48634 | PRRC2A | Protein PRRC2A OS=Homo sapiens (Human) GN=PRRC2A | 2216121.25 | 3076884.25 | 1587439.88 | 2752326.5 | 1898953.5 | 2452835.75 | 3552700.25 | 2740719.25 | **1.105103379** | 7 |
| sp\|Q9P1Y | Q9P1Y5 | CAMSAP3 | Calmodulin-regulated spectrin-associated protein 3 OS= | 479406.38 | 487568.62 | 454022.75 | 449519.31 | 458320.16 | 493886.66 | 575869.38 | 538823.44 | **1.104988393** | 2 |
| sp\|O7583 | O75832 | PSMD10 | 26S proteasome non-ATPase regulatory subunit 10 OS=H | 4362413 | 3112775 | 3619499.5 | 3532572.5 | 4315201.5 | 4075506 | 3656719 | 4114132.5 | **1.104893124** | 4 |
| sp\|Q1314 | Q13144 | EIF2B5 | Translation initiation factor eIF-2B subunit epsilon OS=H | 2558378.5 | 3224229.5 | 3480026.75 | 1241652 | 2895401 | 2597731.5 | 3362124.25 | 2749143.5 | **1.104729957** | 7 |
| sp\|Q9ULX | Q9ULX6 | AKAP8L | A-kinase anchor protein 8-like OS=Homo sapiens (Huma | 2353271.5 | 2563199.25 | 1803113 | 1712630.25 | 1843053 | 2067168.38 | 2602408 | 2798137.5 | **1.10419006** | 4 |
| sp\|Q8WW | Q8WWC4 | MAIP1 | m-AAA protease-interacting protein 1, mitochondrial OS | 4986870 | 7989054 | 6504157.5 | 5727160.5 | 6863069 | 6614617.5 | 7011918.5 | 7337806 | **1.103945088** | 4 |
| sp\|O7548 | O75489 | NDUFS3 | NADH dehydrogenase [ubiquinone] iron-sulfur protein 3, | 15965083 | 17016134 | 17354158 | 18908214 | 18275112 | 19262024 | 18906068 | 19982636 | **1.103724418** | 2 |
| sp\|Q1314 | Q13148 | TARDBP | TAR DNA-binding protein 43 OS=Homo sapiens (Human) | 21425292 | 21347912 | 18481136 | 22696936 | 22302384 | 23748236 | 21136538 | 25455746 | **1.103531815** | 4 |
| sp\|P35659 | P35659 | DEK | Protein DEK OS=Homo sapiens (Human) GN=DEK PE=1 S | 22815670 | 19814278 | 20694498 | 22092956 | 27763360 | 23465094 | 23478506 | 19481444 | **1.102684017** | 4 |
| sp\|Q1328 | Q13283 | G3BP1 | Ras GTPase-activating protein-binding protein 1 OS=Hon | 34609248 | 31168946 | 22457956 | 32718296 | 41659248 | 31775020 | 36809248 | 23111264 | **1.102520696** | 5 |
| sp\|Q4G0N | Q4G0N4 | NADK2 | NAD kinase 2, mitochondrial OS=Homo sapiens (Human) | 1100986.12 | 1256549.12 | 1148508.38 | 1276840.12 | 1226948.5 | 1332733.12 | 1282863 | 1430525 | **1.102487517** | 2 |
| sp\|Q96HC | Q96HC4 | PDLIM5 | PDZ and LIM domain protein 5 OS=Homo sapiens (Huma | 18686750 | 15934407 | 23256444 | 16242400 | 20483242 | 17402996 | 24560642 | 19260866 | **1.102371086** | 4 |
| sp\|Q1276 | Q12768 | WASHC5 | WASH complex subunit 5 OS=Homo sapiens (Human) GN | 576559.31 | 609829.88 | 636334.94 | 586368.56 | 706122.12 | 679053 | 597008.25 | 673496.94 | **1.10235705** | 2 |
| sp\|P60953 | P60953 | CDC42 | Cell division control protein 42 homolog OS=Homo sapie | 36113200 | 33424420 | 35285304 | 36761600 | 43649592 | 43436500 | 35975000 | 32953580 | **1.101918964** | 6 |
| sp\|Q2TB9 | Q2TB90 | HKDC1 | Putative hexokinase HKDC1 OS=Homo sapiens (Human) | 3727539.75 | 3406747.25 | 2532626.75 | 3481076.5 | 5775086.5 | 5199692 | 1713773.5 | 1794645.25 | **1.101552174** | 8 |
| sp\|Q1474 | Q14746 | COG2 | Conserved oligomeric Golgi complex subunit 2 OS=Homo | 0 | 0 | 1235919.5 | 2619430 | 1392324.75 | 1667585.25 | 1186516.25 | 0 | **1.101437431** | 7 |
| sp\|Q9Y2L1 | Q9Y2L1 | DIS3 | Exosome complex exonuclease RRP44 OS=Homo sapiens | 6906104 | 7368443 | 3609347 | 6903867.5 | 6253857.5 | 7576935 | 6140216 | 7329967 | **1.101389309** | 7 |
| sp\|P61981 | P61981 | YWHAG | 14-3-3 protein gamma OS=Homo sapiens (Human) GN=Y | 180074464 | 196476576 | 237327488 | 218295840 | 233652208 | 214072048 | 230314640 | 238471536 | **1.101344222** | 4 |
| sp\|P11387 | P11387 | TOP1 | DNA topoisomerase 1 OS=Homo sapiens (Human) GN=T | 9576764 | 9635049 | 7649083 | 18374610 | 16417622 | 14402016 | 9349350 | 9644506 | **1.101203422** | 6 |

| | | | | | | | | | | | | | |
|---|---|---|---|---|---|---|---|---|---|---|---|---|---|
| sp\|Q1685( | Q16850 | CYP51A1 | Lanosterol 14-alpha demethylase OS=Homo sapiens (Hu | 5532354.5 | 5389225 | 5197530 | 5762826.5 | 5904134.5 | 3868348.5 | 6995529 | 7323596.5 | **1.100981581** | 4 |
| sp\|O95861 | O95861 | BPNT1 | 3'(2'),5'-bisphosphate nucleotidase 1 OS=Homo sapiens | 7289383 | 10601065 | 10250036 | 10095772 | 10716122 | 9446955 | 10680887 | 11252694 | **1.10096182** | 3 |
| sp\|Q96T6( | Q96T60 | PNKP | Bifunctional polynucleotide phosphatase/kinase OS=Hor | 944044.31 | 990056.06 | 1566470.38 | 1088966.25 | 918256.06 | 1194806.75 | 1338626.25 | 1599506.12 | **1.100589271** | 6 |
| sp\|Q9290( | Q92900 | UPF1 | Regulator of nonsense transcripts 1 OS=Homo sapiens (H | 40851120 | 38791468 | 40579224 | 35350584 | 45717744 | 40335696 | 44600084 | 40557380 | **1.100522383** | 4 |
| sp\|P05387 | P05387 | RPLP2 | 60S acidic ribosomal protein P2 OS=Homo sapiens (Hum | 29489160 | 23734898 | 26751596 | 24787062 | 30913470 | 25496460 | 30496414 | 28379310 | **1.100445449** | 3 |
| sp\|O4379! | O43795 | MYO1B | Unconventional myosin-Ib OS=Homo sapiens (Human) G | 21509968 | 23449288 | 24531100 | 19090576 | 24892592 | 23347004 | 24635564 | 24594870 | **1.100350017** | 2 |
| sp\|O1514! | O15145 | ARPC3 | Actin-related protein 2/3 complex subunit 3 OS=Homo s | 16468960 | 15396539 | 15684089 | 15397554 | 18154720 | 17131944 | 18243902 | 15729510 | **1.100289446** | 1 |
| sp\|Q9UBB | Q9UBB4 | ATXN10 | Ataxin-10 OS=Homo sapiens (Human) GN=ATXN10 PE=1 | 10053383 | 10987469 | 12400092 | 10302254 | 13764544 | 11568060 | 10422339 | 12361487 | **1.099975132** | 4 |
| sp\|Q86Y3! | Q86Y39 | NDUFA11 | NADH dehydrogenase [ubiquinone] 1 alpha subcomplex | 3774848.5 | 4282988 | 3762393.5 | 4747143 | 4259999 | 4666114 | 3845762.5 | 5442339.5 | **1.099402724** | 5 |
| sp\|P13051 | P13051 | UNG | Uracil-DNA glycosylase OS=Homo sapiens (Human) GN=U | 775790.12 | 959422.38 | 1276904.12 | 1073093.25 | 1110211.25 | 1063400.75 | 1136371.12 | 1178877.62 | **1.098807866** | 5 |
| sp\|Q0184 | Q01844 | EWSR1 | RNA-binding protein EWS OS=Homo sapiens (Human) GI | 15498016 | 15718067 | 13143528 | 13516760 | 17339752 | 12328310 | 20161392 | 13747184 | **1.098490401** | 6 |
| sp\|Q1516! | Q15165 | PON2 | Serum paraoxonase/arylesterase 2 OS=Homo sapiens (H | 9396988 | 9117173 | 9714565 | 10449120 | 11086570 | 11719822 | 10044498 | 9623867 | **1.098167592** | 3 |
| sp\|P31947 | P31947 | SFN | 14-3-3 protein sigma OS=Homo sapiens (Human) GN=SFI | 133311488 | 140399760 | 175412816 | 147322448 | 170593840 | 150237360 | 164115760 | 170018272 | **1.098112268** | 4 |
| sp\|Q96F0 | Q96F07 | CYFIP2 | Cytoplasmic FMR1-interacting protein 2 OS=Homo sapie | 8425630 | 7981738 | 8975279 | 7899520 | 8391725 | 9100618 | 8876414 | 10174785 | **1.097991666** | 3 |
| sp\|Q1279 | Q12797 | ASPH | Aspartyl/asparaginyl beta-hydroxylase OS=Homo sapien | 6971687 | 6771885 | 5238380 | 5439599.5 | 6558154 | 6893356 | 6468279 | 6893529.5 | **1.097936734** | 6 |
| sp\|Q8IZ83 | Q8IZ83 | ALDH16A1 | Aldehyde dehydrogenase family 16 member A1 OS=Hom | 8238285.5 | 7104189 | 7978770 | 7319378 | 8471557 | 9002098 | 8379325.5 | 7767986.5 | **1.097267753** | 2 |
| sp\|Q0652( | Q06520 | SULT2A1 | Bile salt sulfotransferase OS=Homo sapiens (Human) GN | 19075094 | 17941912 | 20557060 | 20975266 | 21491732 | 17533602 | 24277632 | 22886340 | **1.097263386** | 4 |
| sp\|P50502 | P50502 | ST13 | Hsc70-interacting protein OS=Homo sapiens (Human) GI | 34080968 | 31286324 | 36302064 | 37229028 | 41572640 | 34265032 | 42615744 | 33930896 | **1.097092044** | 5 |
| sp\|Q9H99 | Q9H993 | ARMT1 | Protein-glutamate O-methyltransferase OS=Homo sapie | 13744688 | 10686756 | 11560619 | 12693470 | 13347280 | 13642406 | 12974365 | 13440871 | **1.096936168** | 4 |
| sp\|P80303 | P80303 | NUCB2 | Nucleobindin-2 OS=Homo sapiens (Human) GN=NUCB2 I | 2414318.75 | 2985053.5 | 2252871 | 2592834.75 | 2447169.25 | 2210009.75 | 3457917.75 | 3122157 | **1.096844138** | 6 |
| sp\|O9482 | O94822 | LTN1 | E3 ubiquitin-protein ligase listerin OS=Homo sapiens (Hu | 923288.88 | 1027562.62 | 907960.19 | 844090.69 | 903985.62 | 1181766.25 | 977812.75 | 997017.75 | **1.096594496** | 5 |
| sp\|Q1655! | Q16555 | DPYSL2 | Dihydropyrimidinase-related protein 2 OS=Homo sapien | 78409912 | 76390664 | 83366408 | 80041952 | 89501480 | 84493216 | 91333232 | 83593584 | **1.096517013** | 0 |
| sp\|Q9P0Z! | Q9P0Z9 | PIPOX | Peroxisomal sarcosine oxidase OS=Homo sapiens (Huma | 9819764 | 9300358 | 9350582 | 10814229 | 11170016 | 11113452 | 10225519 | 10558297 | **1.09627994** | 2 |
| sp\|Q9NTX! | Q9NTX5 | ECHDC1 | Ethylmalonyl-CoA decarboxylase OS=Homo sapiens (Hur | 2153726.25 | 2196371 | 3743095.5 | 2435853.75 | 2942714.5 | 3713906.25 | 2862871 | 2019296.62 | **1.095900599** | 7 |
| sp\|Q9BWI | Q9BWD1 | ACAT2 | Acetyl-CoA acetyltransferase, cytosolic OS=Homo sapien | 83974208 | 75785416 | 53802304 | 79776240 | 89650216 | 77649456 | 82316072 | 71741120 | **1.095516708** | 6 |
| sp\|O7534 | O75348 | ATP6V1G1 | V-type proton ATPase subunit G 1 OS=Homo sapiens (Hu | 1953119.75 | 2953234 | 3540190.5 | 3374393.75 | 2792195.5 | 2852031.25 | 3594451 | 3709546.25 | **1.095363498** | 6 |
| sp\|Q9BVI4 | Q9BVI4 | NOC4L | Nucleolar complex protein 4 homolog OS=Homo sapiens | 7388034 | 4851251 | 5170681 | 8226790 | 7251745 | 7460233.5 | 5919083.5 | 7441461.5 | **1.095010753** | 6 |
| sp\|O0044 | O00442 | RTCA | RNA 3'-terminal phosphate cyclase OS=Homo sapiens (H | 2953020 | 3045324.5 | 3901795.5 | 3123747 | 3528040.25 | 3281550.25 | 3465611.5 | 3985386 | **1.094956367** | 3 |
| sp\|P21291 | P21291 | CSRP1 | Cysteine and glycine-rich protein 1 OS=Homo sapiens (H | 11769608 | 7291916.5 | 9992320 | 10801475 | 10829064 | 10383098 | 12500960 | 9925409 | **1.094923627** | 6 |
| sp\|Q71RC | Q71RC2 | LARP4 | La-related protein 4 OS=Homo sapiens (Human) GN=LAR | 1655541.5 | 1929257 | 1827539.25 | 1627432.38 | 1532235.5 | 2144492.5 | 1843426.88 | 2185336.75 | **1.094565801** | 5 |
| sp\|Q9P26! | Q9P265 | DIP2B | Disco-interacting protein 2 homolog B OS=Homo sapiens | 5874464.5 | 5949376.5 | 6654148 | 5706082.5 | 7387715 | 7494121.5 | 5316158.5 | 6272903 | **1.094559202** | 5 |
| sp\|P78330 | P78330 | PSPH | Phosphoserine phosphatase OS=Homo sapiens (Human) | 23581918 | 28160168 | 23009972 | 26691070 | 24664866 | 26983508 | 25110936 | 34267844 | **1.094476838** | 5 |
| sp\|P35232 | P35232 | PHB | Prohibitin OS=Homo sapiens (Human) GN=PHB PE=1 SV= | 160947136 | 157774720 | 210160544 | 170614352 | 208989648 | 178296464 | 184359216 | 193632944 | **1.094041209** | 4 |
| sp\|Q9959 | Q99598 | TSNAX | Translin-associated protein X OS=Homo sapiens (Human | 2222085.5 | 2425878.75 | 2289214.75 | 2652740.5 | 2519638.75 | 2950058 | 2432723.5 | 2583776.75 | **1.093460378** | 3 |
| sp\|Q1381 | Q13813 | SPTAN1 | Spectrin alpha chain, non-erythrocytic 1 OS=Homo sapie | 310138528 | 352091104 | 319719040 | 289072768 | 321995744 | 327114176 | 378140288 | 362383200 | **1.093320195** | 2 |
| sp\|Q9BTD | Q9BTD8 | RBM42 | RNA-binding protein 42 OS=Homo sapiens (Human) GN= | 1976054.5 | 2199251 | 3211012.5 | 2469589.5 | 2481841 | 2943465.5 | 2536023.75 | 2810920 | **1.09297396** | 4 |
| sp\|Q9P0V! | Q9P0V9 | SEPT10 | Septin-10 OS=Homo sapiens (Human) GN=SEPT10 PE=1 S | 1765592.38 | 1540441.5 | 2007751.5 | 2101224.25 | 1650029.62 | 2062651.12 | 1830479.5 | 2558318 | **1.092578249** | 6 |
| sp\|O7593 | O75934 | BCAS2 | Pre-mRNA-splicing factor SPF27 OS=Homo sapiens (Hum | 2309329.75 | 1737774.5 | 1977264.38 | 2287517.5 | 2192726.25 | 2327947.5 | 2064057.25 | 2496606 | **1.092572355** | 4 |
| sp\|O75691 | O75691 | UTP20 | Small subunit processome component 20 homolog OS=H | 15156433 | 14341766 | 37491300 | 28679410 | 25166874 | 26731666 | 25248102 | 27375430 | **1.092539604** | 8 |
| sp\|P21281 | P21281 | ATP6V1B2 | V-type proton ATPase subunit B, brain isoform OS=Homo | 21590060 | 20580808 | 24250906 | 19864160 | 24352292 | 23433860 | 23721136 | 22742772 | **1.092299238** | 3 |
| sp\|P82673 | P82673 | MRPS35 | 28S ribosomal protein S35, mitochondrial OS=Homo sap | 3265326 | 2707979 | 1281131.25 | 2901414.25 | 3620729.5 | 2649677.5 | 2616850.5 | 2203306.75 | **1.092036974** | 7 |
| sp\|Q8NBX | Q8NBX0 | SCCPDH | Saccharopine dehydrogenase-like oxidoreductase OS=H | 10594237 | 8544296 | 8325427.5 | 8003269 | 9989195 | 9357089 | 10360784 | 9011863 | **1.091681858** | 4 |
| sp\|Q86WA | Q86WA8 | LONP2 | Lon protease homolog 2, peroxisomal OS=Homo sapiens | 849630.88 | 805861.81 | 1293462.5 | 1049111.25 | 1249421.5 | 1150097 | 1049210.75 | 915847.62 | **1.091671921** | 5 |
| sp\|Q1349 | Q13492 | PICALM | Phosphatidylinositol-binding clathrin assembly protein C | 1072087.12 | 2227941.75 | 1532852.25 | 1438417 | 1936129.88 | 1896108 | 2019017.25 | 993636.69 | **1.091463313** | 7 |
| sp\|P43378 | P43378 | PTPN9 | Tyrosine-protein phosphatase non-receptor type 9 OS=H | 0 | 211070.89 | 202989.67 | 226178.27 | 188183.81 | 0 | 171413.67 | 339168.19 | **1.091414074** | 6.5 |

| | | | | | | | | | | | | | |
|---|---|---|---|---|---|---|---|---|---|---|---|---|---|
| sp\|Q1351( | Q13510 | ASAH1 | Acid ceramidase OS=Homo sapiens (Human) GN=ASAH1 | 8435487 | 11095922 | 9932944 | 11182155 | 12516221 | 12292565 | 9793364 | 9754432 | **1.091276574** | 6 |
| sp\|Q9BRK! | Q9BRK5 | SDF4 | 45 kDa calcium-binding protein OS=Homo sapiens (Hum | 2314467 | 1539591 | 129100.1 | 1141585.25 | 1029539.56 | 1284043.38 | 2235848.5 | 1040117.38 | **1.090698292** | 7 |
| sp\|P67809 | P67809 | YBX1 | Nuclease-sensitive element-binding protein 1 OS=Homo | 97887432 | 43405936 | 64301400 | 65664336 | 100399472 | 62070112 | 74464648 | 58756940 | **1.090069117** | 7 |
| sp\|Q96EK4 | Q96EK4 | THAP11 | THAP domain-containing protein 11 OS=Homo sapiens (I | 457046.66 | 677127.75 | 567769.31 | 575526.75 | 605979.12 | 648726.88 | 591873 | 634956.06 | **1.089601421** | 4 |
| sp\|Q9NP7 | Q9NP71 | MLXIPL | Carbohydrate-responsive element-binding protein OS=H | 745098.62 | 935951.81 | 654963.06 | 715966.5 | 985204.69 | 666797.56 | 845692.69 | 826147.44 | **1.089077383** | 5 |
| sp\|P49411 | P49411 | TUFM | Elongation factor Tu, mitochondrial OS=Homo sapiens (H | 228987984 | 232493424 | 223197824 | 225967392 | 245938432 | 241556432 | 261665920 | 242366496 | **1.088816731** | 0 |
| sp\|Q86UE4 | Q86UE4 | MTDH | Protein LYRIC OS=Homo sapiens (Human) GN=MTDH PE= | 8023879 | 7202018.5 | 7615947.5 | 6020053 | 9285325 | 6820865 | 7805537 | 7502629 | **1.088436942** | 6 |
| sp\|Q9P2I0 | Q9P2I0 | CPSF2 | Cleavage and polyadenylation specificity factor subunit 2 | 2070149.88 | 1892587.25 | 1836710.75 | 1185141.5 | 1603014.75 | 1922063 | 1857088.75 | 2218852.25 | **1.088255635** | 6 |
| sp\|P62318 | P62318 | SNRPD3 | Small nuclear ribonucleoprotein Sm D3 OS=Homo sapier | 19625518 | 21438112 | 21351620 | 26044416 | 25851996 | 21262824 | 24094300 | 25054692 | **1.088222648** | 6 |
| sp\|P62312 | P62312 | LSM6 | U6 snRNA-associated Sm-like protein LSm6 OS=Homo sa | 7467756 | 11943207 | 7905475 | 6853009.5 | 11707250 | 6974111 | 10892306 | 7610082.5 | **1.088216293** | 7 |
| sp\|Q92542 | Q92542 | NCSTN | Nicastrin OS=Homo sapiens (Human) GN=NCSTN PE=1 S\ | 3744524 | 3717749.5 | 4287405.5 | 3645483.5 | 4285167 | 4137443.5 | 3966694.25 | 4358602 | **1.087868137** | 3 |
| sp\|P62851 | P62851 | RPS25 | 40S ribosomal protein S25 OS=Homo sapiens (Human) G | 100703944 | 97361688 | 101255328 | 111881944 | 115668760 | 117776192 | 109994376 | 103637584 | **1.087241621** | 2 |
| sp\|O75351 | O75351 | VPS4B | Vacuolar protein sorting-associated protein 4B OS=Hom | 9512031 | 10200804 | 6713029 | 7507726 | 8495287 | 8938035 | 10140658 | 9318426 | **1.087194311** | 7 |
| sp\|P04844 | P04844 | RPN2 | Dolichyl-diphosphooligosaccharide--protein glycosyltran | 63900920 | 57694348 | 52932848 | 67895792 | 60436604 | 72054280 | 60971888 | 69932520 | **1.086507078** | 4 |
| sp\|P35249 | P35249 | RFC4 | Replication factor C subunit 4 OS=Homo sapiens (Humar | 3528742 | 3008796.5 | 4634040 | 4616268.5 | 3961146.5 | 3664649.5 | 4356306 | 5168493.5 | **1.086316298** | 6 |
| sp\|P50454 | P50454 | SERPINH1 | Serpin H1 OS=Homo sapiens (Human) GN=SERPINH1 PE= | 76854464 | 70893880 | 64694368 | 74312264 | 74961552 | 83964552 | 78734608 | 73832112 | **1.086268243** | 3 |
| sp\|P05141 | P05141 | SLC25A5 | ADP/ATP translocase 2 OS=Homo sapiens (Human) GN=S | 252815600 | 254911040 | 281509120 | 215132880 | 270271584 | 263919840 | 244233472 | 311917760 | **1.08560006** | 5 |
| sp\|P28072 | P28072 | PSMB6 | Proteasome subunit beta type-6 OS=Homo sapiens (Hun | 46831812 | 48050564 | 45693148 | 33357920 | 48762336 | 43970160 | 52052680 | 43947504 | **1.085085626** | 6 |
| sp\|P48637 | P48637 | GSS | Glutathione synthetase OS=Homo sapiens (Human) GN= | 33510462 | 50120316 | 37464596 | 30090634 | 38545552 | 37448956 | 40312840 | 47677776 | **1.084658072** | 5 |
| sp\|Q9HC3 | Q9HC35 | EML4 | Echinoderm microtubule-associated protein-like 4 OS=H | 125373136 | 128770584 | 110300856 | 107332736 | 130042080 | 130922408 | 136352048 | 114323840 | **1.084495509** | 2 |
| sp\|P61009 | P61009 | SPCS3 | Signal peptidase complex subunit 3 OS=Homo sapiens (H | 4151603 | 3580203.5 | 4135175.75 | 4286986.5 | 3784316.25 | 4336104 | 4185034 | 5211037.5 | **1.084346022** | 4 |
| sp\|P62330 | P62330 | ARF6 | ADP-ribosylation factor 6 OS=Homo sapiens (Human) GN | 2147383.5 | 1836385 | 1709833.12 | 2284088.25 | 1884278.62 | 2124530.5 | 1929640 | 2710132.75 | **1.084096024** | 6 |
| sp\|Q96JB2 | Q96JB2 | COG3 | Conserved oligomeric Golgi complex subunit 3 OS=Homo | 1950466.5 | 2051608.5 | 1505461 | 1854975.12 | 1638824.25 | 1974490.5 | 2194249 | 2173540 | **1.084019246** | 4 |
| sp\|Q1656 | Q16563 | SYPL1 | Synaptophysin-like protein 1 OS=Homo sapiens (Human) | 4142706.5 | 5407420 | 4936534.5 | 4917978.5 | 4588777.5 | 6388400.5 | 3823815.5 | 6228656 | **1.083743375** | 7 |
| sp\|P39687 | P39687 | ANP32A | Acidic leucine-rich nuclear phosphoprotein 32 family me | 52665724 | 52208148 | 57220216 | 48144908 | 51477336 | 53831424 | 66724516 | 55807088 | **1.083720757** | 5 |
| sp\|Q9UBX | Q9UBX3 | SLC25A10 | Mitochondrial dicarboxylate carrier OS=Homo sapiens (H | 21358100 | 18572342 | 22346932 | 22049316 | 22943386 | 22660748 | 23262992 | 22519078 | **1.083716247** | 0 |
| sp\|P62140 | P62140 | PPP1CB | Serine/threonine-protein phosphatase PP1-beta catalytic | 33963796 | 36618028 | 35969704 | 35631416 | 35617500 | 41111468 | 36420380 | 40903332 | **1.083482137** | 4 |
| sp\|Q9NUJ | Q9NUJ1 | ABHD10 | Mycophenolic acid acyl-glucuronide esterase, mitochond | 4692527.5 | 4603610.5 | 5166116.5 | 5785453 | 5491732 | 4597313 | 5629088 | 6215644 | **1.083272119** | 6 |
| sp\|P26368 | P26368 | U2AF2 | Splicing factor U2AF 65 kDa subunit OS=Homo sapiens (H | 25591302 | 26556124 | 25522672 | 28632768 | 26708720 | 31083822 | 28038438 | 29298244 | **1.083030292** | 2 |
| sp\|Q9NRX | Q9NRX4 | PHPT1 | 14 kDa phosphohistidine phosphatase OS=Homo sapiens | 970511.56 | 689641.19 | 604282.38 | 1047857.44 | 866481.94 | 904163.81 | 964223.38 | 850913.38 | **1.082568171** | 8 |
| sp\|Q9UPT | Q9UPT8 | ZC3H4 | Zinc finger CCCH domain-containing protein 4 OS=Homo | 545637.25 | 389031.34 | 321403.5 | 352979.56 | 364324.5 | 444772.59 | 487786.78 | 443750.09 | **1.081776312** | 5 |
| sp\|Q9943( | Q99436 | PSMB7 | Proteasome subunit beta type-7 OS=Homo sapiens (Hun | 9828998 | 9588828 | 10215151 | 11164604 | 10433494 | 12146554 | 11675733 | 9852394 | **1.081146821** | 3 |
| sp\|P78527 | P78527 | PRKDC | DNA-dependent protein kinase catalytic subunit OS=Hor | 330577408 | 355244032 | 373611712 | 328495776 | 341820800 | 362234624 | 354897120 | 441456480 | **1.081041683** | 5 |
| sp\|O6033 | O60333 | KIF1B | Kinesin-like protein KIF1B OS=Homo sapiens (Human) GI | 209930.7 | 266173.12 | 313096.28 | 270735.81 | 319800.5 | 284748.81 | 281733.62 | 259470.38 | **1.080964707** | 5 |
| sp\|P35637 | P35637 | FUS | RNA-binding protein FUS OS=Homo sapiens (Human) GN | 23794262 | 28013576 | 30975432 | 28864130 | 31067022 | 28260392 | 33684440 | 27674780 | **1.080962333** | 5 |
| sp\|Q9255 | Q92552 | MRPS27 | 28S ribosomal protein S27, mitochondrial OS=Homo sap | 13141333 | 15186512 | 12254109 | 12805521 | 14533438 | 13313299 | 14704132 | 15154364 | **1.080875861** | 4 |
| sp\|P08574 | P08574 | CYC1 | Cytochrome c1, heme protein, mitochondrial OS=Homo | 11470115 | 11106663 | 9732005 | 13047961 | 15297416 | 14529970 | 8868021 | 10324271 | **1.08075831** | 7 |
| sp\|Q9NYF | Q9NYF8 | BCLAF1 | Bcl-2-associated transcription factor 1 OS=Homo sapiens | 19064272 | 17503830 | 20348208 | 18449470 | 17614072 | 19767398 | 22335900 | 21696072 | **1.080244137** | 4 |
| sp\|Q9HBK | Q9HBK9 | AS3MT | Arsenite methyltransferase OS=Homo sapiens (Human) ( | 2252680 | 2003224 | 2817757.75 | 2565253.75 | 2535159 | 2937281.75 | 2334593 | 2604504.75 | **1.080156632** | 5 |
| sp\|Q9Y2P | Q9Y2P8 | RCL1 | RNA 3'-terminal phosphate cyclase-like protein OS=Hom | 5164265 | 6436733.5 | 6686325 | 6842062.5 | 6870576 | 6249636 | 7186897 | 6833703 | **1.080042783** | 4 |
| sp\|Q8IWE | Q8IWE2 | FAM114A1 | Protein NOXP20 OS=Homo sapiens (Human) GN=FAM11 | 2175636.25 | 1612038 | 1751521.25 | 1900974.25 | 1911758.38 | 1023997 | 2599492.25 | 2497900.75 | **1.07969961** | 5 |
| sp\|Q9NXS | Q9NXS2 | QPCTL | Glutaminyl-peptide cyclotransferase-like protein OS=Ho | 2200178.75 | 2078182.12 | 1957189.75 | 2178864 | 1880683.25 | 2170711.5 | 2469212.5 | 2561482.5 | **1.07934897** | 6 |
| sp\|Q9280 | Q92804 | TAF15 | TATA-binding protein-associated factor 2N OS=Homo sa | 12387254 | 13973486 | 14624374 | 13115795 | 14046074 | 12740672 | 16659459 | 14942730 | **1.079259777** | 4 |
| sp\|P62258 | P62258 | YWHAE | 14-3-3 protein epsilon OS=Homo sapiens (Human) GN=Y | 222510432 | 231126880 | 275375456 | 243584080 | 275888864 | 243557840 | 259554816 | 270638528 | **1.079213911** | 4 |
| sp\|Q8N12 | Q8N122 | RPTOR | Regulatory-associated protein of mTOR OS=Homo sapier | 1055006.25 | 1320156.75 | 1155322.75 | 1138896.12 | 1216708.88 | 1272331 | 1203192.12 | 1346435.25 | **1.079086567** | 3 |

| | | | | | | | | | | | | | |
|---|---|---|---|---|---|---|---|---|---|---|---|---|---|
| sp\|O4385 | O43852 | CALU | Calumenin OS=Homo sapiens (Human) GN=CALU PE=1 S | 30984250 | 23583866 | 21399518 | 24146732 | 28253876 | 25741360 | 27820184 | 26183252 | **1.078752993** | 4 |
| sp\|Q9NX1 | Q9NX14 | NDUFB11 | NADH dehydrogenase [ubiquinone] 1 beta subcomplex s | 1575163.88 | 2270777.75 | 1801926.5 | 1600982.25 | 2171498.25 | 2347367.75 | 1361556.88 | 1938922.12 | **1.078701393** | 6 |
| sp\|Q0110 | Q01105 | SET | Protein SET OS=Homo sapiens (Human) GN=SET PE=1 SV | 52552940 | 59402620 | 58764588 | 57287464 | 64370560 | 62074264 | 61261576 | 58138844 | **1.078232616** | 2 |
| sp\|P48449 | P48449 | LSS | Lanosterol synthase OS=Homo sapiens (Human) GN=LSS | 2943283.5 | 3793599 | 6759477 | 3709282 | 4426065 | 2219366.75 | 5355640.5 | 6548048 | **1.078083619** | 7 |
| sp\|Q96A3 | Q96A33 | CCDC47 | Coiled-coil domain-containing protein 47 OS=Homo sapi | 10658013 | 10345166 | 10396930 | 10900090 | 11172222 | 11209799 | 11309605 | 11905976 | **1.077952423** | 0 |
| sp\|O1504 | O15042 | U2SURP | U2 snRNP-associated SURP motif-containing protein OS= | 6783819.5 | 8008727.5 | 7009198 | 7599036 | 7569091.5 | 7809264.5 | 7913978 | 8382750.5 | **1.077355207** | 4 |
| sp\|Q53S5 | Q53S58 | TMEM177 | Transmembrane protein 177 OS=Homo sapiens (Human | 695308.25 | 1609269.62 | 859137.19 | 2329558 | 941948.62 | 1629855.88 | 1675726.5 | 1670069.25 | **1.077244875** | 5 |
| sp\|O9490 | O94905 | ERLIN2 | Erlin-2 OS=Homo sapiens (Human) GN=ERLIN2 PE=1 SV= | 30160432 | 27835398 | 33114428 | 28470156 | 35759340 | 33411564 | 29059284 | 30586476 | **1.077238819** | 3 |
| sp\|Q1662 | Q16625 | OCLN | Occludin OS=Homo sapiens (Human) GN=OCLN PE=1 SV= | 0 | 0 | 767984 | 0 | 0 | 0 | 0 | 827112.19 | **1.076991435** | 7.5 |
| sp\|P51659 | P51659 | HSD17B4 | Peroxisomal multifunctional enzyme type 2 OS=Homo sa | 92126800 | 88234368 | 88982968 | 98374152 | 93901680 | 103604424 | 97028000 | 101473832 | **1.076932937** | 2 |
| sp\|Q9H9G | Q9H9G7 | AGO3 | Protein argonaute-3 OS=Homo sapiens (Human) GN=AG | 708890.56 | 755473.75 | 629876.25 | 697147.25 | 713692.06 | 639387.31 | 817659.31 | 835310.62 | **1.07690135** | 4 |
| sp\|P82912 | P82912 | MRPS11 | 28S ribosomal protein S11, mitochondrial OS=Homo sap | 1103134.88 | 1208560.12 | 963140.38 | 951530.81 | 990001.19 | 1322200.5 | 933302.31 | 1304532 | **1.076583475** | 6 |
| sp\|Q1415 | Q14151 | SAFB2 | Scaffold attachment factor B2 OS=Homo sapiens (Huma | 8909630 | 5735883.5 | 7768070.5 | 6013976.5 | 6239965 | 8015677.5 | 7792835.5 | 8550688 | **1.07639085** | 5 |
| sp\|O7530 | O75306 | NDUFS2 | NADH dehydrogenase [ubiquinone] iron-sulfur protein 2, | 8876914 | 9163468 | 7275236 | 9402391 | 9901978 | 9117340 | 8994180 | 9351930 | **1.07625492** | 5 |
| sp\|Q9P2N | Q9P2N5 | RBM27 | RNA-binding protein 27 OS=Homo sapiens (Human) GN= | 1148666.5 | 1411294.75 | 1239670.25 | 1255151.12 | 1240524 | 1206594.25 | 1208935.38 | 1783758.38 | **1.076171305** | 8 |
| sp\|P0DN7 | P0DN76 | U2AF1L5 | Splicing factor U2AF 35 kDa subunit-like protein OS=Hon | 21885428 | 23585308 | 23296704 | 25817284 | 26731188 | 24327946 | 28203056 | 22464760 | **1.075511411** | 4 |
| sp\|P62937 | P62937 | PPIA | Peptidyl-prolyl cis-trans isomerase A OS=Homo sapiens ( | 404505728 | 349608384 | 337302592 | 378045632 | 376205504 | 404247488 | 419548160 | 379899424 | **1.075155543** | 4 |
| sp\|Q96I24 | Q96I24 | FUBP3 | Far upstream element-binding protein 3 OS=Homo sapie | 27515736 | 26687516 | 21224552 | 23141966 | 24811634 | 25764600 | 26680828 | 28719272 | **1.075140319** | 6 |
| sp\|Q1683 | Q16836 | HADH | Hydroxyacyl-coenzyme A dehydrogenase, mitochondrial | 23148024 | 22064644 | 21706030 | 23429762 | 25852108 | 23130886 | 23715464 | 24435034 | **1.07509848** | 2 |
| sp\|O9520 | O95202 | LETM1 | Mitochondrial proton/calcium exchanger protein OS=Ho | 14711297 | 15933700 | 14374498 | 16432770 | 17094492 | 15867606 | 15548350 | 17526308 | **1.074602474** | 4 |
| sp\|Q96A2 | Q96A26 | FAM162A | Protein FAM162A OS=Homo sapiens (Human) GN=FAM1 | 6415596 | 5326065 | 3189931 | 6009679.5 | 6897093 | 8434514 | 3204214 | 3948305.5 | **1.073675326** | 6 |
| sp\|Q9UM | Q9UM00 | TMCO1 | Calcium load-activated calcium channel OS=Homo sapie | 17634796 | 18710016 | 17145246 | 18343980 | 19228952 | 19572504 | 18786958 | 19528542 | **1.073543381** | 0 |
| sp\|P24752 | P24752 | ACAT1 | Acetyl-CoA acetyltransferase, mitochondrial OS=Homo s | 72404520 | 70380984 | 69403952 | 65544340 | 75612048 | 66905836 | 78830144 | 76807984 | **1.07353162** | 3 |
| sp\|Q7L59 | Q7L592 | NDUFAF7 | Protein arginine methyltransferase NDUFAF7, mitochond | 2855140.25 | 4240019 | 4422676.5 | 4265947 | 3788343.5 | 3444727.5 | 4656120.5 | 5051100 | **1.073271963** | 6 |
| sp\|Q9NRR | Q9NRR5 | UBQLN4 | Ubiquilin-4 OS=Homo sapiens (Human) GN=UBQLN4 PE= | 4582685 | 2982008.75 | 3732563 | 3753017 | 3133872 | 4702492.5 | 3735676 | 4579970.5 | **1.073203801** | 6 |
| sp\|Q96EK | Q96EK6 | GNPNAT1 | Glucosamine 6-phosphate N-acetyltransferase OS=Hom | 4048432.75 | 5742551.5 | 5527201.5 | 5726125.5 | 5035213.5 | 5516194 | 5967939.5 | 6051463.5 | **1.072537382** | 6 |
| sp\|P23381 | P23381 | WARS | Tryptophan--tRNA ligase, cytoplasmic OS=Homo sapiens | 25354990 | 23294296 | 21044776 | 21870968 | 27050330 | 23104340 | 25353846 | 22694368 | **1.072493331** | 5 |
| sp\|Q9BY6 | Q9BY67 | CADM1 | Cell adhesion molecule 1 OS=Homo sapiens (Human) GN | 444695.84 | 705135.25 | 591358 | 456840.16 | 481908.34 | 850123.69 | 513132.47 | 511999.25 | **1.072398718** | 6 |
| sp\|Q9BSJ2 | Q9BSJ2 | TUBGCP2 | Gamma-tubulin complex component 2 OS=Homo sapien | 830460.56 | 1237029 | 1481630 | 1216152 | 1434843 | 841524.44 | 1385338.5 | 1448247 | **1.072331949** | 6 |
| sp\|Q9Y38 | Q9Y383 | LUC7L2 | Putative RNA-binding protein Luc7-like 2 OS=Homo sapi | 6996381 | 6855637 | 10038692 | 6993017 | 9096716 | 6463440.5 | 9699360 | 7853717.5 | **1.072190348** | 7 |
| sp\|Q1388 | Q13884 | SNTB1 | Beta-1-syntrophin OS=Homo sapiens (Human) GN=SNTB | 27350016 | 27131516 | 22074398 | 25424440 | 28676946 | 25793968 | 27950698 | 26765120 | **1.070664207** | 4 |
| sp\|Q9UNK | Q9UNK0 | STX8 | Syntaxin-8 OS=Homo sapiens (Human) GN=STX8 PE=1 SV | 0 | 692650.06 | 0 | 789384 | 0 | 959375.12 | 0 | 626401.81 | **1.070000328** | 8 |
| sp\|Q1456 | Q14562 | DHX8 | ATP-dependent RNA helicase DHX8 OS=Homo sapiens (H | 383158.88 | 723376.44 | 568149.69 | 0 | 470097.38 | 0 | 668854 | 651554.88 | **1.069160021** | 7.5 |
| sp\|O7532 | O75323 | NIPSNAP2 | Protein NipSnap homolog 2 OS=Homo sapiens (Human) | 2840811.75 | 3784961.5 | 3161342.5 | 3957547 | 3868256 | 3077391.25 | 3695611.5 | 4049758.5 | **1.068852508** | 6 |
| sp\|P13861 | P13861 | PRKAR2A | cAMP-dependent protein kinase type II-alpha regulatory | 19139540 | 15920086 | 16051662 | 16117594 | 16064959 | 17762018 | 17243598 | 20781806 | **1.068772511** | 4 |
| sp\|P82663 | P82663 | MRPS25 | 28S ribosomal protein S25, mitochondrial OS=Homo sap | 2388050.75 | 2251299.75 | 1855133.25 | 2073775 | 2357462.5 | 2131870.5 | 2352615 | 2312492 | **1.068413113** | 5 |
| sp\|Q8WU | Q8WUK0 | PTPMT1 | Phosphatidylglycerophosphatase and protein-tyrosine pl | 658443.38 | 614679.31 | 659602.38 | 580871.81 | 677489.81 | 677319.75 | 703074.81 | 627114.31 | **1.068189852** | 2 |
| sp\|P63241 | P63241 | EIF5A | Eukaryotic translation initiation factor 5A-1 OS=Homo sa | 85634576 | 76011952 | 73668216 | 84157568 | 96528912 | 82548680 | 90741520 | 71320368 | **1.06782174** | 6 |
| sp\|P10809 | P10809 | HSPD1 | 60 kDa heat shock protein, mitochondrial OS=Homo sapi | 2552014592 | 2832610560 | 3420297472 | 2838979072 | 3356428032 | 2947379456 | 3388734208 | 2735888384 | **1.067376761** | 6 |
| sp\|P62857 | P62857 | RPS28 | 40S ribosomal protein S28 OS=Homo sapiens (Human) G | 14197239 | 13823422 | 16162731 | 15807654 | 16532560 | 15083678 | 17160768 | 15250702 | **1.067287742** | 4 |
| sp\|P46782 | P46782 | RPS5 | 40S ribosomal protein S5 OS=Homo sapiens (Human) GN | 102797352 | 104358736 | 105045672 | 102475912 | 128239736 | 104002752 | 110089888 | 100198912 | **1.067169317** | 6 |
| sp\|P10599 | P10599 | TXN | Thioredoxin OS=Homo sapiens (Human) GN=TXN PE=1 S | 52735148 | 58808572 | 64732788 | 53378852 | 78981384 | 44205576 | 66356144 | 55535388 | **1.067157727** | 6 |
| sp\|P43243 | P43243 | MATR3 | Matrin-3 OS=Homo sapiens (Human) GN=MATR3 PE=1 S | 87610520 | 84952032 | 87858048 | 86048568 | 90077368 | 90504896 | 102236792 | 86894592 | **1.067089606** | 2 |
| sp\|P23284 | P23284 | PPIB | Peptidyl-prolyl cis-trans isomerase B OS=Homo sapiens ( | 321946176 | 360138880 | 365621184 | 332413184 | 379885568 | 315277632 | 408423456 | 368807360 | **1.06685986** | 4 |
| sp\|Q9268 | Q92688 | ANP32B | Acidic leucine-rich nuclear phosphoprotein 32 family me | 46022732 | 40765192 | 43784460 | 40917752 | 48373252 | 41964940 | 46565748 | 46016672 | **1.066653839** | 3 |

| | | | | | | | | | | | | |
|---|---|---|---|---|---|---|---|---|---|---|---|---|
| sp\|Q9Y3L5 | Q9Y3L5 | RAP2C | Ras-related protein Rap-2c OS=Homo sapiens (Human) G | 1864571.88 | 1648157.25 | 1951689 | 2344023 | 2196971.5 | 1997030.75 | 1781767.38 | 2352095.25 | **1.0665208** | 5 |
| sp\|Q1515 | Q15154 | PCM1 | Pericentriolar material 1 protein OS=Homo sapiens (Hun | 1776417.75 | 966206.5 | 1889035.38 | 1363724.5 | 1904340 | 1534866 | 1305343.5 | 1649133.5 | **1.066434254** | 7 |
| sp\|P02749 | P02749 | APOH | Beta-2-glycoprotein 1 OS=Homo sapiens (Human) GN=A | 1329851.5 | 1244132.62 | 1768879.75 | 1545516.5 | 1381426 | 1314300.75 | 1806834.62 | 1774421.5 | **1.065994803** | 5 |
| sp\|Q9Y6M | Q9Y6M9 | NDUFB9 | NADH dehydrogenase [ubiquinone] 1 beta subcomplex s | 4832591 | 3292013.5 | 3287753.25 | 4618291.5 | 4642605.5 | 4325954.5 | 4445549 | 3671754 | **1.065824767** | 7 |
| sp\|P16278 | P16278 | GLB1 | Beta-galactosidase OS=Homo sapiens (Human) GN=GLB1 | 13221177 | 14991117 | 12567808 | 14690980 | 13771883 | 13997564 | 14806722 | 16540208 | **1.065715232** | 5 |
| sp\|O1502 | O15027 | SEC16A | Protein transport protein Sec16A OS=Homo sapiens (Hu | 2238905 | 4027224.75 | 1733646 | 3836145.75 | 2870489.5 | 3011365.25 | 3271082.25 | 3457894.25 | **1.065471011** | 8 |
| sp\|Q9NZ0 | Q9NZ01 | TECR | Very-long-chain enoyl-CoA reductase OS=Homo sapiens | 22455940 | 24568104 | 21314216 | 25911608 | 25257142 | 24271640 | 22719744 | 28163696 | **1.065383158** | 5 |
| sp\|P55072 | P55072 | VCP | Transitional endoplasmic reticulum ATPase OS=Homo sa | 342038304 | 373876608 | 394350944 | 356350048 | 394935936 | 379495680 | 400272128 | 387683392 | **1.065300827** | 2 |
| sp\|P62805 | P62805 | HIST1H4A | Histone H4 OS=Homo sapiens (Human) GN=HIST1H4A PE | 2478088960 | 2676660224 | 2852403968 | 2607528192 | 2883038720 | 2633641216 | 2716111872 | 3072765952 | **1.065086873** | 3 |
| sp\|Q9Y63 | Q9Y639 | NPTN | Neuroplastin OS=Homo sapiens (Human) GN=NPTN PE=1 | 1856590 | 2160791.5 | 3137473 | 1761523.25 | 2480362.25 | 2302249.25 | 2234344.75 | 2477934.5 | **1.064882065** | 4 |
| sp\|Q8IWX | Q8IWX8 | CHERP | Calcium homeostasis endoplasmic reticulum protein OS= | 5275089 | 4711683.5 | 5419439.5 | 5396700.5 | 5461648 | 4689840.5 | 6164265 | 5836293.5 | **1.064853155** | 4 |
| sp\|P78316 | P78316 | NOP14 | Nucleolar protein 14 OS=Homo sapiens (Human) GN=NC | 3609326 | 3685476 | 3531166 | 3378853 | 3972553.5 | 3658769.5 | 3227641.5 | 4266298.5 | **1.06479786** | 5 |
| sp\|P04070 | P04070 | PROC | Vitamin K-dependent protein C OS=Homo sapiens (Huma | 767116.62 | 756558.06 | 537348.81 | 618390.81 | 708579.88 | 777037.56 | 695714.31 | 671694 | **1.064794552** | 6 |
| sp\|P62306 | P62306 | SNRPF | Small nuclear ribonucleoprotein F OS=Homo sapiens (Hu | 10505527 | 12356926 | 15039128 | 15695539 | 13627029 | 13711527 | 13626580 | 16103862 | **1.064777324** | 6 |
| sp\|P61586 | P61586 | RHOA | Transforming protein RhoA OS=Homo sapiens (Human) ( | 16115132 | 21785472 | 20332890 | 19050774 | 21861006 | 19859180 | 17862192 | 22695128 | **1.064608725** | 5 |
| sp\|Q9UBQ | Q9UBQ0 | VPS29 | Vacuolar protein sorting-associated protein 29 OS=Hom | 6564622 | 5818244 | 5161278.5 | 6022404 | 5947419.5 | 7009962.5 | 6865232.5 | 5264746 | **1.064532657** | 5 |
| sp\|Q8NB9 | Q8NB90 | SPATA5 | ATPase family protein 2 homolog OS=Homo sapiens (Hu | 101149.79 | 93006.88 | 83879.82 | 121605.88 | 86610.98 | 137222.58 | 95477.88 | 106028.94 | **1.064302516** | 6 |
| sp\|Q9UJZ1 | Q9UJZ1 | STOML2 | Stomatin-like protein 2, mitochondrial OS=Homo sapien | 41547152 | 36539116 | 41454680 | 39945176 | 41634440 | 41704688 | 44130996 | 42224312 | **1.064007525** | 0 |
| sp\|P00846 | P00846 | MT-ATP6 | ATP synthase subunit a OS=Homo sapiens (Human) GN= | 2431591.5 | 3548137.5 | 2155898.75 | 5082312.5 | 4762838.5 | 3840276 | 1790330.12 | 3670419.5 | **1.063998161** | 7 |
| sp\|O1464 | O14647 | CHD2 | Chromodomain-helicase-DNA-binding protein 2 OS=Hon | 701838.69 | 1011426.19 | 524095.66 | 719672.25 | 739280.12 | 724619.94 | 872799.31 | 808746.56 | **1.06371696** | 4 |
| sp\|O7593 | O75937 | DNAJC8 | DnaJ homolog subfamily C member 8 OS=Homo sapiens | 20187930 | 17015214 | 10527604 | 20236572 | 20923302 | 17680002 | 19390104 | 14301373 | **1.063669731** | 7 |
| sp\|Q9Y5L | Q9Y5L0 | TNPO3 | Transportin-3 OS=Homo sapiens (Human) GN=TNPO3 PE | 1947737 | 1853100 | 1598160.5 | 669679.88 | 1553422.62 | 1708709.75 | 1633826 | 1558456.25 | **1.063561995** | 6 |
| sp\|Q1343 | Q13435 | SF3B2 | Splicing factor 3B subunit 2 OS=Homo sapiens (Human) ( | 33702160 | 31185012 | 31207352 | 27223772 | 31963688 | 31727636 | 32941344 | 34512080 | **1.063465457** | 3 |
| sp\|P38606 | P38606 | ATP6V1A | V-type proton ATPase catalytic subunit A OS=Homo sapi | 15088912 | 15253252 | 11833147 | 16653760 | 17637350 | 16077259 | 15390672 | 13448831 | **1.063319732** | 5 |
| sp\|Q9Y5Z | Q9Y5Z4 | HEBP2 | Heme-binding protein 2 OS=Homo sapiens (Human) GN= | 2930843.5 | 1121515.75 | 6421013 | 1347603 | 4934668.5 | 3379959.75 | 2874831.5 | 1375493.88 | **1.06293714** | 6 |
| sp\|P68036 | P68036 | UBE2L3 | Ubiquitin-conjugating enzyme E2 L3 OS=Homo sapiens (l | 16265704 | 17108988 | 16457094 | 18068290 | 19548838 | 16757070 | 19091412 | 16767944 | **1.062815659** | 4 |
| sp\|Q9273 | Q92734 | TFG | Protein TFG OS=Homo sapiens (Human) GN=TFG PE=1 S\ | 10474297 | 12309262 | 13089682 | 13138851 | 11558764 | 11737140 | 13516643 | 15263179 | **1.062507717** | 6 |
| sp\|Q9Y5L | Q9Y5L4 | TIMM13 | Mitochondrial import inner membrane translocase subu | 3836805.75 | 3299960 | 3604213 | 3689945 | 3757335.75 | 3092602.75 | 3803346.5 | 4679624 | **1.062503639** | 6 |
| sp\|P22570 | P22570 | FDXR | NADPH:adrenodoxin oxidoreductase, mitochondrial OS= | 88614600 | 88031016 | 84145800 | 87757872 | 93173488 | 88586616 | 100816992 | 87707256 | **1.062358653** | 4 |
| sp\|Q6P58 | Q6P587 | FAHD1 | Acylpyruvase FAHD1, mitochondrial OS=Homo sapiens (I | 1855187.5 | 2101076.75 | 2539422 | 3508013 | 2853073.5 | 1499938 | 2837259.25 | 3431943 | **1.061828578** | 7 |
| sp\|P06737 | P06737 | PYGL | Glycogen phosphorylase, liver form OS=Homo sapiens (H | 28237468 | 30932622 | 32162250 | 27177614 | 30905324 | 30542982 | 31621118 | 32709170 | **1.061333582** | 5 |
| sp\|Q9Y2W | Q9Y2W2 | WBP11 | WW domain-binding protein 11 OS=Homo sapiens (Hum | 987319.69 | 671197.75 | 660993.81 | 619614.44 | 847578.56 | 721776.75 | 975182.62 | 573788.31 | **1.0609707** | 7 |
| sp\|Q7Z46 | Q7Z460 | CLASP1 | CLIP-associating protein 1 OS=Homo sapiens (Human) GI | 3692461 | 4211132.5 | 3636083.25 | 4085382.5 | 4374433 | 3963389.25 | 4620662 | 3618531.5 | **1.060924985** | 6 |
| sp\|Q9BPX | Q9BPX6 | MICU1 | Calcium uptake protein 1, mitochondrial OS=Homo sapie | 125836.52 | 125145.21 | 53613.06 | 189646.34 | 93678.95 | 206331.88 | 114748.36 | 109515.98 | **1.06076799** | 7 |
| sp\|Q8IY67 | Q8IY67 | RAVER1 | Ribonucleoprotein PTB-binding 1 OS=Homo sapiens (Hu | 8665780 | 8510513 | 9655549 | 8912399 | 10224184 | 7836561.5 | 11498645 | 8348858.5 | **1.060541445** | 8 |
| sp\|Q86V8 | Q86V81 | ALYREF | THO complex subunit 4 OS=Homo sapiens (Human) GN= | 13435906 | 10913644 | 11534442 | 12736980 | 13439852 | 12392946 | 14015418 | 11714944 | **1.060512735** | 4 |
| sp\|Q9945 | Q99459 | CDC5L | Cell division cycle 5-like protein OS=Homo sapiens (Hum | 23461780 | 21574644 | 19790522 | 16986822 | 22885728 | 19956490 | 23685672 | 20234986 | **1.060492361** | 5 |
| sp\|P13987 | P13987 | CD59 | CD59 glycoprotein OS=Homo sapiens (Human) GN=CD59 | 11141277 | 12118960 | 10692248 | 11467534 | 13825609 | 11970346 | 11325688 | 11024980 | **1.060030886** | 6 |
| sp\|P36404 | P36404 | ARL2 | ADP-ribosylation factor-like protein 2 OS=Homo sapiens | 3525599.75 | 1802364.62 | 3502680.5 | 3899869 | 3724796.5 | 3159831.5 | 2409424.25 | 4200435 | **1.060011197** | 7 |
| sp\|Q9P2R | Q9P2R7 | SUCLA2 | Succinate--CoA ligase [ADP-forming] subunit beta, mitocl | 8793949 | 9999794 | 8119503 | 8298332.5 | 8909822 | 8392524 | 10188253 | 9828100 | **1.059841694** | 4 |
| sp\|Q9Y3E | Q9Y3E0 | GOLT1B | Vesicle transport protein GOT1B OS=Homo sapiens (Hun | 5216502.5 | 4570841.5 | 4733871 | 5116027 | 4783161.5 | 5861539.5 | 4755616.5 | 5393127.5 | **1.058878075** | 4 |
| sp\|Q9UHG | Q9UHG3 | PCYOX1 | Prenylcysteine oxidase 1 OS=Homo sapiens (Human) GN | 21370782 | 24977444 | 29699056 | 32258692 | 25080658 | 30136382 | 28173996 | 31276616 | **1.058738016** | 6 |
| sp\|P07099 | P07099 | EPHX1 | Epoxide hydrolase 1 OS=Homo sapiens (Human) GN=EPH | 20086726 | 18947494 | 16522700 | 19286224 | 21090568 | 19291226 | 18124924 | 20721416 | **1.058589067** | 4 |
| sp\|Q1383 | Q13838 | DDX39B | Spliceosome RNA helicase DDX39B OS=Homo sapiens (H | 152470400 | 164965344 | 160389472 | 165582000 | 168762864 | 156262928 | 181482656 | 174091056 | **1.057805208** | 3 |
| sp\|Q9HC3 | Q9HC38 | GLOD4 | Glyoxalase domain-containing protein 4 OS=Homo sapie | 20782708 | 22753966 | 18554232 | 20403522 | 25235100 | 20948320 | 22899974 | 18159584 | **1.05756207** | 5 |

| | | | | | | | | | | | | | |
|---|---|---|---|---|---|---|---|---|---|---|---|---|---|
| sp\|O7549 | O75494 | SRSF10 | Serine/arginine-rich splicing factor 10 OS=Homo sapiens | 15082786 | 12300006 | 11934361 | 14084483 | 15753432 | 11321878 | 18086292 | 11302468 | **1.057347194** | 8 |
| sp\|P11279 | P11279 | LAMP1 | Lysosome-associated membrane glycoprotein 1 OS=Hon | 5803976.5 | 6633128 | 5562598 | 6594939.5 | 5147124 | 5869084 | 5726552 | 9254944 | **1.057047466** | 7 |
| sp\|Q8WU | Q8WU90 | ZC3H15 | Zinc finger CCCH domain-containing protein 15 OS=Hom | 10553617 | 6265345 | 10611132 | 9003368 | 9221738 | 7825463.5 | 10691168 | 10768088 | **1.05689812** | 5 |
| sp\|P30566 | P30566 | ADSL | Adenylosuccinate lyase OS=Homo sapiens (Human) GN= | 17507432 | 18311602 | 15888178 | 14970922 | 18267716 | 20421060 | 17509840 | 14269534 | **1.056840463** | 6 |
| sp\|P09496 | P09496 | CLTA | Clathrin light chain A OS=Homo sapiens (Human) GN=CL | 1890636.25 | 2031553.5 | 2901090.5 | 3032783 | 2779828.5 | 2110948 | 2703026.75 | 2819656.75 | **1.05655369** | 8 |
| sp\|Q9P03 | Q9P035 | HACD3 | Very-long-chain (3R)-3-hydroxyacyl-CoA dehydratase 3 ( | 21884206 | 24036434 | 26123272 | 25650846 | 25251352 | 24138042 | 24122230 | 29702022 | **1.056491137** | 6 |
| sp\|P60983 | P60983 | GMFB | Glia maturation factor beta OS=Homo sapiens (Human) ( | 2220874.75 | 2118058.5 | 1975352.75 | 2770199 | 2237433.5 | 2536759.25 | 2311075.25 | 2510882 | **1.056322951** | 4 |
| sp\|Q9NPD | Q9NPD3 | EXOSC4 | Exosome complex component RRP41 OS=Homo sapiens | 4342000.5 | 3805856 | 4427429 | 4280992.5 | 4995604.5 | 4509296.5 | 4401491 | 3897808.75 | **1.056235591** | 4 |
| sp\|Q9NYK | Q9NYK5 | MRPL39 | 39S ribosomal protein L39, mitochondrial OS=Homo sapi | 4103618 | 4722203 | 5530969 | 5643637.5 | 3818932.5 | 5670765 | 5687165 | 5948074.5 | **1.056224273** | 4 |
| sp\|P63151 | P63151 | PPP2R2A | Serine/threonine-protein phosphatase 2A 55 kDa regulat | 4557900.5 | 4922054.5 | 5130720.5 | 4496193.5 | 5387425 | 4577099 | 4820981 | 5386130 | **1.05572687** | 4 |
| sp\|O0016 | O00165 | HAX1 | HCLS1-associated protein X-1 OS=Homo sapiens (Human | 202316.22 | 212224.08 | 222985.62 | 260492.69 | 258116.19 | 233230.42 | 221284.27 | 235367.42 | **1.055655517** | 5 |
| sp\|P12235 | P12235 | SLC25A4 | ADP/ATP translocase 1 OS=Homo sapiens (Human) GN=S | 193945136 | 190956272 | 200204480 | 146510448 | 189313680 | 191543856 | 167572720 | 223653600 | **1.055312488** | 8 |
| sp\|O6034 | O60343 | TBC1D4 | TBC1 domain family member 4 OS=Homo sapiens (Huma | 6531718 | 7567472.5 | 6822375.5 | 6633152 | 8457455 | 5863964 | 7757199.5 | 6998115 | **1.055236112** | 5 |
| sp\|Q2Q1W | Q2Q1W2 | TRIM71 | E3 ubiquitin-protein ligase TRIM71 OS=Homo sapiens (H | 5091847.5 | 5063720 | 5386246.5 | 5149501.5 | 5396564 | 4930409.5 | 4887876.5 | 6618041.5 | **1.055171746** | 8 |
| sp\|Q9UKK | Q9UKK3 | PARP4 | Poly [ADP-ribose] polymerase 4 OS=Homo sapiens (Hum | 512649.31 | 519800.75 | 492898.66 | 442666.22 | 484217.84 | 477511.22 | 611021.69 | 502785.97 | **1.054634636** | 8 |
| sp\|Q9UM | Q9UMS0 | NFU1 | NFU1 iron-sulfur cluster scaffold homolog, mitochondrial | 1895537.25 | 1738984.88 | 1951481.12 | 1649246.12 | 1993567.25 | 1424605.12 | 2256067.5 | 1955330.25 | **1.054499953** | 4 |
| sp\|O1537 | O15371 | EIF3D | Eukaryotic translation initiation factor 3 subunit D OS=H | 29853416 | 30339336 | 32821560 | 34351796 | 32711806 | 33764556 | 32264772 | 35508944 | **1.05404868** | 5 |
| sp\|O4332 | O43324 | EEF1E1 | Eukaryotic translation elongation factor 1 epsilon-1 OS=I | 18292302 | 16861148 | 12685889 | 15677919 | 16614280 | 16660365 | 18083136 | 15573738 | **1.053753281** | 8 |
| sp\|Q8N16 | Q8N163 | CCAR2 | Cell cycle and apoptosis regulator protein 2 OS=Homo sa | 37525868 | 33476320 | 28019052 | 34663520 | 35104496 | 35094184 | 36601440 | 34059936 | **1.053673253** | 5 |
| sp\|Q9UBU | Q9UBU9 | NXF1 | Nuclear RNA export factor 1 OS=Homo sapiens (Human) | 2166378 | 3399874 | 3611306 | 1698764.12 | 1534478.88 | 3032813.5 | 2895521.75 | 3996892.75 | **1.053638055** | 8 |
| sp\|Q9NZL | Q9NZL4 | HSPBP1 | Hsp70-binding protein 1 OS=Homo sapiens (Human) GN | 2325416 | 2177033.5 | 6438969.5 | 3769781.5 | 3293785.5 | 3565461.25 | 3263987 | 5376972 | **1.053632961** | 7 |
| sp\|P43246 | P43246 | MSH2 | DNA mismatch repair protein Msh2 OS=Homo sapiens (H | 9260083 | 9955047 | 9882052 | 11285123 | 9969741 | 10551565 | 10031776 | 11992300 | **1.053564971** | 3 |
| sp\|Q1357 | Q13573 | SNW1 | SNW domain-containing protein 1 OS=Homo sapiens (Hu | 7614678 | 6109166 | 6332849 | 7923526 | 6760565.5 | 8009650 | 8242506.5 | 6463798.5 | **1.053477119** | 4 |
| sp\|O7504 | O75044 | SRGAP2 | SLIT-ROBO Rho GTPase-activating protein 2 OS=Homo sa | 1589690.62 | 2067628.88 | 338356.12 | 487832.81 | 446078.66 | 1828759.25 | 596571.06 | 1850846.5 | **1.053250048** | 7 |
| sp\|Q1412 | Q14126 | DSG2 | Desmoglein-2 OS=Homo sapiens (Human) GN=DSG2 PE= | 61209516 | 73527656 | 73735840 | 101251496 | 76842216 | 61357100 | 74785408 | 113221024 | **1.053212579** | 5 |
| sp\|Q9299 | Q92995 | USP13 | Ubiquitin carboxyl-terminal hydrolase 13 OS=Homo sapi | 2487220.25 | 2224345.25 | 1943800.62 | 2246484 | 2107173.25 | 2539104 | 2366324 | 2362305.75 | **1.053141411** | 5 |
| sp\|Q1536 | Q15366 | PCBP2 | Poly(rC)-binding protein 2 OS=Homo sapiens (Human) G | 98179920 | 94466600 | 135216048 | 97976464 | 129335640 | 100427552 | 100714904 | 117990232 | **1.053140493** | 4 |
| sp\|Q9974 | Q99747 | NAPG | Gamma-soluble NSF attachment protein OS=Homo sapie | 1770646.75 | 1357245.88 | 1400722.88 | 1408191.12 | 1557484.62 | 1689615.88 | 1470250 | 1533935.12 | **1.052971068** | 4 |
| sp\|Q5RKV | Q5RKV6 | EXOSC6 | Exosome complex component MTR3 OS=Homo sapiens ( | 4931500 | 5231511 | 4671111 | 5678215.5 | 5441969.5 | 5289718.5 | 5409660 | 5452558.5 | **1.05272773** | 4 |
| sp\|Q9P1Z | Q9P1Z9 | CCDC180 | Coiled-coil domain-containing protein 180 OS=Homo sap | 3879709 | 4029877.25 | 4222162.5 | 3685764.75 | 3892762.5 | 4179253.25 | 4382242 | 4192304.25 | **1.052413327** | 4 |
| sp\|P62854 | P62854 | RPS26 | 40S ribosomal protein S26 OS=Homo sapiens (Human) G | 47340964 | 45754196 | 46064380 | 57314288 | 51898136 | 50225468 | 55719804 | 48873140 | **1.052132745** | 4 |
| sp\|Q1360 | Q13601 | KRR1 | KRR1 small subunit processome component homolog O | 2587416 | 2696845 | 2212689 | 2368839.25 | 2345888.5 | 2815937 | 2675288.75 | 2542581.25 | **1.052089725** | 6 |
| sp\|O0014 | O00148 | DDX39A | ATP-dependent RNA helicase DDX39A OS=Homo sapiens | 109488952 | 120362088 | 121919024 | 120970880 | 123732896 | 110240592 | 133510232 | 129736672 | **1.05178195** | 3 |
| sp\|P60228 | P60228 | EIF3E | Eukaryotic translation initiation factor 3 subunit E OS=H | 45635248 | 41122440 | 43873068 | 46817056 | 47635860 | 42555576 | 45025424 | 51418648 | **1.051776891** | 5 |
| sp\|P09132 | P09132 | SRP19 | Signal recognition particle 19 kDa protein OS=Homo sapi | 4025697.25 | 2591819.75 | 1077997.38 | 2291955.5 | 3210628.25 | 993272.12 | 3342575.5 | 2957958.25 | **1.051761282** | 7 |
| sp\|Q9UJX | Q9UJX3 | ANAPC7 | Anaphase-promoting complex subunit 7 OS=Homo sapie | 830969.88 | 829531.06 | 763800 | 1165421.75 | 923980 | 973724.94 | 970808 | 906086.31 | **1.051501627** | 4 |
| sp\|Q96GC | Q96GC5 | MRPL48 | 39S ribosomal protein L48, mitochondrial OS=Homo sapi | 3871519.5 | 3777791.5 | 4553863 | 3967223 | 4695334 | 3997255 | 4267086.5 | 4034793.25 | **1.050961751** | 3 |
| sp\|Q1506 | Q15067 | ACOX1 | Peroxisomal acyl-coenzyme A oxidase 1 OS=Homo sapie | 29785042 | 29231902 | 32683692 | 33630392 | 33356318 | 34461524 | 33745860 | 30147812 | **1.050909069** | 3 |
| sp\|Q1410 | Q14108 | SCARB2 | Lysosome membrane protein 2 OS=Homo sapiens (Huma | 102978.55 | 168397.23 | 71002.08 | 139716.17 | 81615.5 | 153865.14 | 98833.27 | 172127.17 | **1.0505027** | 7 |
| sp\|Q9UJV | Q9UJV9 | DDX41 | Probable ATP-dependent RNA helicase DDX41 OS=Homo | 3798148 | 2288840.5 | 4818811 | 3177797 | 3541408.5 | 3300962 | 4886821 | 3061499 | **1.05020692** | 7 |
| sp\|P47914 | P47914 | RPL29 | 60S ribosomal protein L29 OS=Homo sapiens (Human) G | 52368252 | 39571380 | 64367308 | 53550284 | 60873164 | 57921048 | 49628768 | 51839648 | **1.049583254** | 8 |
| sp\|Q6UXN | Q6UXN9 | WDR82 | WD repeat-containing protein 82 OS=Homo sapiens (Hu | 1124280 | 783728.31 | 0 | 1142149.38 | 1041581.25 | 693107.38 | 437205.19 | 1028806.5 | **1.049355688** | 6 |
| sp\|P18583 | P18583 | SON | Protein SON OS=Homo sapiens (Human) GN=SON PE=1 S | 6352588.5 | 7339705 | 8024420 | 6068258.5 | 7441233.5 | 6345613 | 8097572 | 7264219.5 | **1.049079265** | 6 |
| sp\|Q9UBE | Q9UBE0 | SAE1 | SUMO-activating enzyme subunit 1 OS=Homo sapiens (H | 18921392 | 23378544 | 25132830 | 20792448 | 22602360 | 20391720 | 28521434 | 21024444 | **1.048906019** | 7 |
| sp\|Q1689 | Q16891 | IMMT | MICOS complex subunit MIC60 OS=Homo sapiens (Huma | 35477676 | 42579552 | 37978968 | 39738200 | 40104760 | 41054692 | 42473024 | 39642808 | **1.048152252** | 5 |

| | | | | | | | | | | | | | |
|---|---|---|---|---|---|---|---|---|---|---|---|---|---|
| sp\|P04040 | P04040 | CAT | Catalase OS=Homo sapiens (Human) GN=CAT PE=1 SV=3 | 54256380 | 52222292 | 52434648 | 50885464 | 59410412 | 48287360 | 55459280 | 56734124 | **1.048105102** | 4 |
| sp\|P62136 | P62136 | PPP1CA | Serine/threonine-protein phosphatase PP1-alpha catalyt | 46812576 | 49399536 | 48909868 | 49143648 | 47630676 | 51829560 | 47980000 | 56120824 | **1.047849082** | 6 |
| sp\|Q8NI36 | Q8NI36 | WDR36 | WD repeat-containing protein 36 OS=Homo sapiens (Hu | 11158464 | 11229509 | 13667178 | 13230911 | 13128022 | 13766703 | 10909580 | 13812024 | **1.047280446** | 6 |
| sp\|Q9UBC | Q9UBC2 | EPS15L1 | Epidermal growth factor receptor substrate 15-like 1 OS | 6369805 | 5920646 | 7522997 | 7163503.5 | 6936457 | 7803612.5 | 6503920 | 7002161 | **1.047047532** | 6 |
| sp\|P47224 | P47224 | RABIF | Guanine nucleotide exchange factor MSS4 OS=Homo sap | 1267719.25 | 938870.94 | 1525096.25 | 1123983 | 1461930.25 | 1071238.5 | 1057090.75 | 1492574.38 | **1.046783341** | 8 |
| sp\|Q9Y6V | Q9Y6V7 | DDX49 | Probable ATP-dependent RNA helicase DDX49 OS=Homo | 1682605 | 2069397.75 | 2184411.75 | 1425411 | 1819062.25 | 2050849.75 | 2003058 | 1830744.62 | **1.046440807** | 8 |
| sp\|O6078 | O60784 | TOM1 | Target of Myb protein 1 OS=Homo sapiens (Human) GN= | 4528939 | 3647968.25 | 4636540.5 | 2828076.25 | 3864808.75 | 4395203 | 4473614.5 | 3633924.25 | **1.046416609** | 7 |
| sp\|Q9P0J0 | Q9P0J0 | NDUFA13 | NADH dehydrogenase [ubiquinone] 1 alpha subcomplex | 7856588 | 8741355 | 8418977 | 9198475 | 9448007 | 8476698 | 9134711 | 8725546 | **1.045873122** | 5 |
| sp\|Q9Y49 | Q9Y490 | TLN1 | Talin-1 OS=Homo sapiens (Human) GN=TLN1 PE=1 SV=3 | 116930176 | 116345664 | 109808000 | 103141744 | 115773664 | 108655584 | 125960664 | 116014504 | **1.045221145** | 7 |
| sp\|Q71DI3 | Q71DI3 | HIST2H3A | Histone H3.2 OS=Homo sapiens (Human) GN=HIST2H3A | 1218430848 | 1352533504 | 1586230016 | 1298991104 | 1543352832 | 1386227328 | 1074267776 | 1698675968 | **1.045148471** | 6 |
| sp\|P34949 | P34949 | MPI | Mannose-6-phosphate isomerase OS=Homo sapiens (Hu | 2384593.25 | 2004984.38 | 2112320 | 2381250.5 | 2257453 | 2453375.25 | 2512052.5 | 2060069.25 | **1.045006777** | 5 |
| sp\|P84243 | P84243 | H3F3A | Histone H3.3 OS=Homo sapiens (Human) GN=H3F3A PE= | 1219464704 | 1353125632 | 1586300928 | 1299750528 | 1543947392 | 1386344960 | 1074504576 | 1699111296 | **1.044931769** | 6 |
| sp\|P68431 | P68431 | HIST1H3A | Histone H3.1 OS=Homo sapiens (Human) GN=HIST1H3A | 1220644480 | 1354206464 | 1586332160 | 1301883648 | 1544287616 | 1387810432 | 1075240832 | 1700223360 | **1.044754256** | 6 |
| sp\|Q1523 | Q15233 | NONO | Non-POU domain-containing octamer-binding protein O | 88852384 | 92861024 | 96945712 | 94661760 | 101855720 | 83616152 | 109249264 | 95303792 | **1.044744478** | 5 |
| sp\|Q9NTG | Q9NTG7 | SIRT3 | NAD-dependent protein deacetylase sirtuin-3, mitochon | 617208.5 | 493839.94 | 506977.47 | 832953.38 | 535079.62 | 618715.5 | 692133.06 | 714299.5 | **1.044573363** | 5 |
| sp\|Q969Y | Q969Y2 | GTPBP3 | tRNA modification GTPase GTPBP3, mitochondrial OS=H | 1012884.94 | 448683.19 | 961025.25 | 1162572.38 | 636571.69 | 1072977.38 | 779047.94 | 1255944.25 | **1.044454151** | 7 |
| sp\|Q9BZE | Q9BZE4 | GTPBP4 | Nucleolar GTP-binding protein 1 OS=Homo sapiens (Hum | 14030442 | 13665254 | 15678147 | 15058742 | 15877002 | 13209214 | 15793475 | 16145895 | **1.044375942** | 4 |
| sp\|Q9NX6 | Q9NX61 | TMEM161 | Transmembrane protein 161A OS=Homo sapiens (Huma | 212451.27 | 191318.48 | 245696.8 | 248406.92 | 206405.34 | 267391.81 | 174533.27 | 289054.66 | **1.044005766** | 7 |
| sp\|Q96AA | Q96AA3 | RFT1 | Protein RFT1 homolog OS=Homo sapiens (Human) GN=R | 900682.75 | 821377.62 | 749267.62 | 787447.62 | 729892.62 | 878274.94 | 949395.62 | 844491.06 | **1.043967013** | 6 |
| sp\|Q7Z3B | Q7Z3B4 | NUP54 | Nucleoporin p54 OS=Homo sapiens (Human) GN=NUP54 | 3816369.5 | 3955655 | 3642268.5 | 4529838.5 | 3495874.5 | 4400830 | 4237529.5 | 4494554 | **1.042940972** | 7 |
| sp\|P55039 | P55039 | DRG2 | Developmentally-regulated GTP-binding protein 2 OS=H | 4622733 | 6094492 | 6041170 | 6354505 | 5419500 | 5189475.5 | 6444059 | 7050469 | **1.042859334** | 6 |
| sp\|Q9Y4G | Q9Y4G6 | TLN2 | Talin-2 OS=Homo sapiens (Human) GN=TLN2 PE=1 SV=4 | 12309138 | 12110758 | 14861865 | 12327049 | 11775648 | 12284094 | 15840330 | 13904996 | **1.042555874** | 8 |
| sp\|P20585 | P20585 | MSH3 | DNA mismatch repair protein Msh3 OS=Homo sapiens (H | 289383.75 | 245422.73 | 189429.05 | 259564.5 | 238814.98 | 302203.34 | 250747.98 | 233581.59 | **1.042232017** | 8 |
| sp\|P17480 | P17480 | UBTF | Nucleolar transcription factor 1 OS=Homo sapiens (Hum | 3818517.5 | 3909073 | 4230315 | 3916280.5 | 4589144 | 3830342 | 4205162.5 | 3918848.5 | **1.042163485** | 5 |
| sp\|O6083 | O60832 | DKC1 | H/ACA ribonucleoprotein complex subunit DKC1 OS=Hor | 32277008 | 33980192 | 32969680 | 30094558 | 33179824 | 31653702 | 36916872 | 32940370 | **1.041519257** | 6 |
| sp\|Q9NPF | Q9NPF4 | OSGEP | Probable tRNA N6-adenosine threonylcarbamoyltransfer | 945511.19 | 824957.12 | 760488.94 | 660757.5 | 831720.62 | 723606.88 | 1015414.94 | 752891.56 | **1.041331779** | 7 |
| sp\|P30153 | P30153 | PPP2R1A | Serine/threonine-protein phosphatase 2A 65 kDa regulat | 50054208 | 53493728 | 64652048 | 56003856 | 60771912 | 54866380 | 62715776 | 54977928 | **1.040713647** | 6 |
| sp\|P19823 | P19823 | ITIH2 | Inter-alpha-trypsin inhibitor heavy chain H2 OS=Homo sa | 13286673 | 13468690 | 10252280 | 10248648 | 11202618 | 7388717 | 12688742 | 17881704 | **1.040322462** | 8 |
| sp\|Q14CX | Q14CX7 | NAA25 | N-alpha-acetyltransferase 25, NatB auxiliary subunit OS= | 4045878 | 4158871 | 4764447.5 | 3913591 | 5213899 | 4306009 | 3768488 | 4267134.5 | **1.039847863** | 6 |
| sp\|Q8TDN | Q8TDN6 | BRIX1 | Ribosome biogenesis protein BRX1 homolog OS=Homo s | 11099973 | 12192714 | 14271249 | 13584973 | 10414339 | 14696710 | 11397346 | 16670400 | **1.039685812** | 7 |
| sp\|Q9NR3 | Q9NR30 | DDX21 | Nucleolar RNA helicase 2 OS=Homo sapiens (Human) GN | 115676536 | 104193440 | 126274904 | 114279160 | 125472664 | 123486864 | 122035728 | 107631208 | **1.039534043** | 6 |
| sp\|O4366 | O43660 | PLRG1 | Pleiotropic regulator 1 OS=Homo sapiens (Human) GN=P | 3875201.25 | 2561012.5 | 2249669 | 4062593.75 | 3569144.75 | 3230809.75 | 3390870.25 | 3059322 | **1.039351388** | 8 |
| sp\|Q1279 | Q12792 | TWF1 | Twinfilin-1 OS=Homo sapiens (Human) GN=TWF1 PE=1 S | 4684801.5 | 4192911.5 | 4847842.5 | 3821063 | 4811322.5 | 6034904.5 | 3258219 | 4132442.25 | **1.039339189** | 8 |
| sp\|P30049 | P30049 | ATP5F1D | ATP synthase subunit delta, mitochondrial OS=Homo sap | 15896092 | 16204070 | 23409586 | 21555226 | 20766326 | 17959422 | 20273766 | 21035964 | **1.038545449** | 8 |
| sp\|P16989 | P16989 | YBX3 | Y-box-binding protein 3 OS=Homo sapiens (Human) GN= | 70374800 | 24054542 | 31433992 | 30742316 | 68752128 | 29192046 | 37078348 | 27596952 | **1.038401067** | 8 |
| sp\|Q9257 | Q92575 | UBXN4 | UBX domain-containing protein 4 OS=Homo sapiens (Hu | 8119895 | 7857167 | 8203990.5 | 6857094 | 8280934 | 8552895 | 8130015 | 7256459.5 | **1.038087229** | 4 |
| sp\|O1511 | O15116 | LSM1 | U6 snRNA-associated Sm-like protein LSm1 OS=Homo sa | 838224.31 | 884304.75 | 683582.69 | 719167.88 | 856289 | 879903.56 | 756977.81 | 750788.25 | **1.037973879** | 6 |
| sp\|P55795 | P55795 | HNRNPH2 | Heterogeneous nuclear ribonucleoprotein H2 OS=Homo | 75882496 | 71962424 | 70844288 | 56772640 | 71815920 | 66638244 | 81179024 | 66270892 | **1.037908088** | 8 |
| sp\|P12236 | P12236 | SLC25A6 | ADP/ATP translocase 3 OS=Homo sapiens (Human) GN=S | 232817344 | 234099152 | 257624144 | 197132720 | 236215824 | 236742976 | 212329712 | 271247296 | **1.037825166** | 5 |
| sp\|Q8IZ69 | Q8IZ69 | TRMT2A | tRNA (uracil-5-)-methyltransferase homolog A OS=Homo | 271671.72 | 255594.42 | 266941.84 | 251126.78 | 246088.56 | 366800.12 | 471891.06 | 0 | **1.037734304** | 8 |
| sp\|P51991 | P51991 | HNRNPA3 | Heterogeneous nuclear ribonucleoprotein A3 OS=Homo | 84936296 | 112014600 | 113897864 | 122865160 | 111700528 | 104624000 | 115924472 | 117628312 | **1.037267404** | 8 |
| sp\|P12931 | P12931 | SRC | Proto-oncogene tyrosine-protein kinase Src OS=Homo sa | 15906970 | 18740662 | 16325220 | 15092622 | 18977120 | 15588600 | 14443715 | 19514936 | **1.037219093** | 7 |
| sp\|P28074 | P28074 | PSMB5 | Proteasome subunit beta type-5 OS=Homo sapiens (Hun | 40393944 | 36133668 | 46922332 | 48681632 | 51039420 | 33497346 | 47912696 | 46074888 | **1.03713888** | 7 |
| sp\|P62760 | P62760 | VSNL1 | Visinin-like protein 1 OS=Homo sapiens (Human) GN=VS | 2334299 | 2562085 | 7245007 | 3403000.25 | 3926436 | 4198663 | 4529129 | 3467355 | **1.037131834** | 4 |
| sp\|Q9944 | Q99447 | PCYT2 | Ethanolamine-phosphate cytidylyltransferase OS=Homo | 30025356 | 29580692 | 31204014 | 31705994 | 30506092 | 28933748 | 33402868 | 34214000 | **1.037061689** | 6 |

| | | | | | | | | | | | | |
|---|---|---|---|---|---|---|---|---|---|---|---|---|
| sp\|Q9NS6 | Q9NS69 | TOMM22 | Mitochondrial import receptor subunit TOM22 homolog | 13208082 | 11701509 | 11327667 | 13452056 | 13593780 | 12757206 | 13356592 | 11809823 | **1.036790345** | 5 |
| sp\|Q9954 | Q99541 | PLIN2 | Perilipin-2 OS=Homo sapiens (Human) GN=PLIN2 PE=1 S' | 11277214 | 12721163 | 15138639 | 12945266 | 14137109 | 14111015 | 12792591 | 12950940 | **1.036660702** | 5 |
| sp\|Q1497 | Q14978 | NOLC1 | Nucleolar and coiled-body phosphoprotein 1 OS=Homo | 56868936 | 49633044 | 64384760 | 42594116 | 56633584 | 49824480 | 62376384 | 52423408 | **1.036429496** | 7 |
| sp\|P13473 | P13473 | LAMP2 | Lysosome-associated membrane glycoprotein 2 OS=Hon | 21640624 | 26634284 | 26735718 | 27046488 | 25183076 | 24015706 | 27811444 | 28744364 | **1.036229478** | 6 |
| sp\|Q9Y29 | Q9Y295 | DRG1 | Developmentally-regulated GTP-binding protein 1 OS=H | 19578360 | 20565232 | 22348614 | 21064372 | 21257490 | 21170684 | 22023248 | 22067060 | **1.035447885** | 4 |
| sp\|O0051 | O00515 | LAD1 | Ladinin-1 OS=Homo sapiens (Human) GN=LAD1 PE=1 SV | 14809598 | 14866601 | 16329119 | 13444039 | 15628594 | 14269549 | 16635129 | 15021060 | **1.035407868** | 5 |
| sp\|Q9BXP | Q9BXP5 | SRRT | Serrate RNA effector molecule homolog OS=Homo sapie | 25183580 | 26627294 | 29889308 | 23582730 | 22623688 | 27776350 | 28612152 | 29978278 | **1.035215173** | 6 |
| sp\|Q9254 | Q92544 | TM9SF4 | Transmembrane 9 superfamily member 4 OS=Homo sap | 8189728.5 | 7589879 | 6488388 | 7291199 | 7224322 | 8558364 | 7891539 | 6919928.5 | **1.035013099** | 7 |
| sp\|Q96DH | Q96DH6 | MSI2 | RNA-binding protein Musashi homolog 2 OS=Homo sapi | 3632407.5 | 3855552 | 3624579.25 | 3618042.25 | 1401474.25 | 4412142.5 | 4633566.5 | 4796879 | **1.03485818** | 4 |
| sp\|P35270 | P35270 | SPR | Sepiapterin reductase OS=Homo sapiens (Human) GN=S | 27583012 | 25932318 | 17173628 | 27994084 | 29187858 | 25901684 | 23090100 | 23942136 | **1.034846271** | 7 |
| sp\|Q1537 | Q15370 | ELOB | Elongin-B OS=Homo sapiens (Human) GN=ELOB PE=1 SV | 16154653 | 13328744 | 16699798 | 16960048 | 16108468 | 16251808 | 15821908 | 17161324 | **1.034845613** | 8 |
| sp\|P49366 | P49366 | DHPS | Deoxyhypusine synthase OS=Homo sapiens (Human) GN | 181457.28 | 417587.88 | 324183.38 | 260207.48 | 130237.27 | 493263.94 | 314005 | 286982.53 | **1.034689429** | 8 |
| sp\|Q8WZ8 | Q8WZ82 | OVCA2 | Esterase OVCA2 OS=Homo sapiens (Human) GN=OVCA2 | 500137.53 | 449325.66 | 427581.44 | 411460.84 | 414721.56 | 486207.03 | 465113.72 | 483456.5 | **1.034102965** | 6 |
| sp\|Q6IA86 | Q6IA86 | ELP2 | Elongator complex protein 2 OS=Homo sapiens (Human) | 435656 | 343904.41 | 299486.09 | 459160.09 | 276702.66 | 485512.12 | 326453.94 | 501901.66 | **1.034042105** | 7 |
| sp\|Q0838 | Q08380 | LGALS3BP | Galectin-3-binding protein OS=Homo sapiens (Human) G | 11179840 | 5010905 | 5102955 | 12368630 | 11588570 | 7566680.5 | 6063094.5 | 9588858 | **1.03401051** | 7 |
| sp\|Q9998 | Q99986 | VRK1 | Serine/threonine-protein kinase VRK1 OS=Homo sapiens | 2254075 | 2508219 | 2538596 | 2172543 | 2316457 | 1905889.12 | 2867832 | 2705285.25 | **1.033992996** | 6 |
| sp\|Q9976 | Q99766 | ATP5S | ATP synthase subunit s, mitochondrial OS=Homo sapiens | 694489.62 | 572059.56 | 750155.88 | 963591.44 | 762176.19 | 794951.12 | 807886.62 | 715554.81 | **1.033645055** | 5 |
| sp\|Q9NUC | Q9NUQ8 | ABCF3 | ATP-binding cassette sub-family F member 3 OS=Homo s | 1215015.88 | 1011055.56 | 992567.12 | 1334683 | 1297101.12 | 1122595.12 | 966637.5 | 1319407.88 | **1.033474477** | 8 |
| sp\|Q9ULC | Q9ULC5 | ACSL5 | Long-chain-fatty-acid--CoA ligase 5 OS=Homo sapiens (H | 49428956 | 41716048 | 42418656 | 41444192 | 44258684 | 44299500 | 46033764 | 46219744 | **1.033163312** | 4 |
| sp\|Q9Y3I0 | Q9Y3I0 | RTCB | tRNA-splicing ligase RtcB homolog OS=Homo sapiens (H | 34962528 | 32549576 | 34232444 | 33773960 | 43614500 | 29667092 | 35671768 | 31052460 | **1.033112171** | 8 |
| sp\|Q96EC | Q96EC8 | YIPF6 | Protein YIPF6 OS=Homo sapiens (Human) GN=YIPF6 PE= | 651649.62 | 516390.81 | 0 | 488455.06 | 0 | 923622.81 | 0 | 787653.06 | **1.033070045** | 7 |
| sp\|Q2TAY | Q2TAY7 | SMU1 | WD40 repeat-containing protein SMU1 OS=Homo sapier | 5990865.5 | 5821968.5 | 6446434 | 6242356 | 5809735.5 | 5777483.5 | 6755801.5 | 6955066 | **1.032506519** | 8 |
| sp\|P04843 | P04843 | RPN1 | Dolichyl-diphosphooligosaccharide--protein glycosyltran | 121419272 | 134814544 | 128664560 | 125363176 | 127649760 | 133181408 | 136951424 | 128973776 | **1.032326198** | 4 |
| sp\|Q9BT7 | Q9BT78 | COPS4 | COP9 signalosome complex subunit 4 OS=Homo sapiens | 7015962.5 | 5373439 | 6128807 | 5643454 | 6556877 | 4679122 | 6485775.5 | 7215111 | **1.032084837** | 6 |
| sp\|P05455 | P05455 | SSB | Lupus La protein OS=Homo sapiens (Human) GN=SSB PE | 73103872 | 73455008 | 85653160 | 65104764 | 81380376 | 72744944 | 85979512 | 66697220 | **1.031902832** | 7 |
| sp\|Q8IXB1 | Q8IXB1 | DNAJC10 | DnaJ homolog subfamily C member 10 OS=Homo sapien | 6527827.5 | 7591555.5 | 8241590 | 6675571 | 7843544.5 | 7350359.5 | 7507700.5 | 7247322 | **1.03142187** | 7 |
| sp\|Q9BSJ8 | Q9BSJ8 | ESYT1 | Extended synaptotagmin-1 OS=Homo sapiens (Human) ( | 40830188 | 42095528 | 43140724 | 44082100 | 41490304 | 41139196 | 42708028 | 50057880 | **1.030836985** | 8 |
| sp\|O9583 | O95832 | CLDN1 | Claudin-1 OS=Homo sapiens (Human) GN=CLDN1 PE=1 S | 678297.56 | 764205.38 | 768454.25 | 1087207.62 | 670078.44 | 754046.94 | 666023.75 | 1308435.5 | **1.030447181** | 5 |
| sp\|P63167 | P63167 | DYNLL1 | Dynein light chain 1, cytoplasmic OS=Homo sapiens (Hur | 12204703 | 10701176 | 12091137 | 11183915 | 12353454 | 13442487 | 11515724 | 10273546 | **1.030408222** | 6 |
| sp\|O7609 | O76094 | SRP72 | Signal recognition particle subunit SRP72 OS=Homo sapi | 17710700 | 19194472 | 19982180 | 18409686 | 20140058 | 19148842 | 18994080 | 19249274 | **1.029685311** | 5 |
| sp\|Q1342 | Q13427 | PPIG | Peptidyl-prolyl cis-trans isomerase G OS=Homo sapiens ( | 3720727.5 | 3116519.75 | 3404621 | 3676340.5 | 2034045.88 | 2753921.75 | 5229119 | 4312815 | **1.029579444** | 8 |
| sp\|Q9HBH | Q9HBH5 | RDH14 | Retinol dehydrogenase 14 OS=Homo sapiens (Human) G | 198667.56 | 153844.94 | 126968.7 | 125683.68 | 119555.34 | 158011.12 | 148212.5 | 197181.86 | **1.029406763** | 8 |
| sp\|Q9H6F | Q9H6F5 | CCDC86 | Coiled-coil domain-containing protein 86 OS=Homo sapi | 5596787.5 | 3958998.5 | 4156113.5 | 4913491 | 6048538.5 | 4897412 | 5503051.5 | 2721355.5 | **1.02925936** | 7 |
| sp\|Q1564 | Q15645 | TRIP13 | Pachytene checkpoint protein 2 homolog OS=Homo sapi | 1270934.62 | 1306095.88 | 1786312.38 | 1396940.38 | 1477654.38 | 1787027 | 1202888.12 | 1460484.12 | **1.029125366** | 6 |
| sp\|Q1414 | Q14146 | URB2 | Unhealthy ribosome biogenesis protein 2 homolog OS=H | 1164949.25 | 922010.12 | 1562494 | 1439290 | 1219132 | 1253126.25 | 1294947.5 | 1467344.25 | **1.028652777** | 7 |
| sp\|P33176 | P33176 | KIF5B | Kinesin-1 heavy chain OS=Homo sapiens (Human) GN=K | 60855864 | 59092404 | 63301120 | 47902440 | 64691212 | 51976448 | 61364200 | 59704444 | **1.028485503** | 6 |
| sp\|Q1545 | Q15459 | SF3A1 | Splicing factor 3A subunit 1 OS=Homo sapiens (Human) ( | 41252360 | 45420672 | 59797056 | 51486964 | 48300868 | 50674516 | 48859976 | 55759304 | **1.028478965** | 7 |
| sp\|O4370 | O43707 | ACTN4 | Alpha-actinin-4 OS=Homo sapiens (Human) GN=ACTN4 F | 472101920 | 518618848 | 498844704 | 479709312 | 502972288 | 479129408 | 525374880 | 517261696 | **1.028164423** | 5 |
| sp\|P62995 | P62995 | TRA2B | Transformer-2 protein homolog beta OS=Homo sapiens | 39984916 | 39593480 | 40580692 | 42290076 | 44393300 | 41008936 | 44004528 | 37571028 | **1.027877201** | 5 |
| sp\|P42285 | P42285 | MTREX | Exosome RNA helicase MTR4 OS=Homo sapiens (Human | 15010367 | 18122572 | 18139744 | 18272478 | 17459128 | 18022248 | 17091536 | 18907542 | **1.02782786** | 7 |
| sp\|P62633 | P62633 | CNBP | Cellular nucleic acid-binding protein OS=Homo sapiens ( | 19618232 | 21589320 | 19723596 | 19531164 | 22527408 | 19595248 | 24919040 | 15649388 | **1.027699577** | 7 |
| sp\|Q9NP6 | Q9NP61 | ARFGAP3 | ADP-ribosylation factor GTPase-activating protein 3 OS= | 4012654.5 | 4123350.5 | 3630097.5 | 3008263.25 | 3936584.5 | 3346182.75 | 3910218.75 | 3987410.5 | **1.02748211** | 7 |
| sp\|O4339 | O43395 | PRPF3 | U4/U6 small nuclear ribonucleoprotein Prp3 OS=Homo s | 7120355.5 | 6793054 | 6680377 | 5985040 | 7831470 | 7409145 | 7309112 | 4751175 | **1.027167321** | 4 |
| sp\|Q9NTI5 | Q9NTI5 | PDS5B | Sister chromatid cohesion protein PDS5 homolog B OS=H | 8125679 | 8139178.5 | 7970956 | 7066751 | 7840183 | 7853938.5 | 7900506 | 8550121 | **1.026904633** | 7 |
| sp\|Q9NR0 | Q9NR09 | BIRC6 | Baculoviral IAP repeat-containing protein 6 OS=Homo sa | 1161502.75 | 766350.19 | 1371386.88 | 1189455.88 | 1344714 | 1264398.75 | 823754.25 | 1176383 | **1.026857312** | 7 |

| | | | | | | | | | | | | |
|---|---|---|---|---|---|---|---|---|---|---|---|---|
| sp\|P09874 | P09874 | PARP1 | Poly [ADP-ribose] polymerase 1 OS=Homo sapiens (Hum | 67775664 | 66167728 | 68802120 | 67095180 | 67798824 | 66382304 | 69474592 | 73429328 | **1.026846789** | 4 |
| sp\|Q8TBQ | Q8TBQ9 | TMEM167 | Protein kish-A OS=Homo sapiens (Human) GN=TMEM16 | 3076603.75 | 3161507.5 | 3575127.75 | 3796229.75 | 3331065 | 3372874 | 3961271 | 3308761.75 | **1.026783044** | 6 |
| sp\|Q96S52 | Q96S52 | PIGS | GPI transamidase component PIG-S OS=Homo sapiens (H | 2791897.5 | 2267355.75 | 1259944.38 | 2737394.75 | 2445829 | 2777699 | 1825265.25 | 2248945 | **1.026626557** | 7 |
| sp\|Q1434 | Q14344 | GNA13 | Guanine nucleotide-binding protein subunit alpha-13 OS | 1358695.5 | 3113158 | 2188351.75 | 2092229.88 | 1865050.5 | 1910763.5 | 2809646.5 | 2399082 | **1.026519176** | 8 |
| sp\|Q9Y6C | Q9Y6C9 | MTCH2 | Mitochondrial carrier homolog 2 OS=Homo sapiens (Hur | 29331418 | 27189044 | 27231618 | 30591306 | 28656122 | 30420994 | 26968088 | 31303676 | **1.026284808** | 7 |
| sp\|O4315 | O43156 | TTI1 | TELO2-interacting protein 1 homolog OS=Homo sapiens | 1851966.12 | 1034943.38 | 2229624.5 | 2272996.5 | 1798488.38 | 2299029 | 1782419.88 | 1702808.25 | **1.026147129** | 7 |
| sp\|P47756 | P47756 | CAPZB | F-actin-capping protein subunit beta OS=Homo sapiens ( | 40579528 | 48078336 | 49254240 | 50104260 | 52938104 | 40376844 | 49781464 | 49831036 | **1.026120514** | 6 |
| sp\|Q9GZT | Q9GZT3 | SLIRP | SRA stem-loop-interacting RNA-binding protein, mitocho | 25003574 | 25402502 | 20875846 | 24944138 | 27701488 | 24245612 | 26419744 | 20360102 | **1.025989696** | 7 |
| sp\|Q9H9A | Q9H9A5 | CNOT10 | CCR4-NOT transcription complex subunit 10 OS=Homo s | 739311.44 | 711882.31 | 802880.69 | 780710.69 | 651303.81 | 823952.81 | 756013.81 | 882276.69 | **1.02595307** | 6 |
| sp\|Q9NQG | Q9NQG5 | RPRD1B | Regulation of nuclear pre-mRNA domain-containing prot | 23712788 | 22820174 | 20680016 | 23440362 | 24545824 | 21791882 | 22633938 | 24030200 | **1.025906426** | 6 |
| sp\|P15531 | P15531 | NME1 | Nucleoside diphosphate kinase A OS=Homo sapiens (Hu | 148615552 | 165822032 | 171781312 | 158651392 | 173369312 | 161858784 | 163152864 | 163045712 | **1.025673976** | 6 |
| sp\|Q9BTC | Q9BTC8 | MTA3 | Metastasis-associated protein MTA3 OS=Homo sapiens ( | 3082762.25 | 3366020 | 4089384 | 2826792.5 | 2870709.75 | 2618373.25 | 3570903 | 4645699.5 | **1.025494037** | 8 |
| sp\|Q69YN | Q69YN4 | VIRMA | Protein virilizer homolog OS=Homo sapiens (Human) GN | 655709.69 | 813983.81 | 621185.62 | 617335.38 | 561283 | 608260.25 | 708752.12 | 898855.75 | **1.025454638** | 7 |
| sp\|Q2M38 | Q2M389 | WASHC4 | WASH complex subunit 4 OS=Homo sapiens (Human) GN | 1182608.75 | 1076973.88 | 928303.75 | 1329514.62 | 999295.5 | 1193820.62 | 1108050.12 | 1331134.75 | **1.025434977** | 6 |
| sp\|Q9UNL | Q9UNL2 | SSR3 | Translocon-associated protein subunit gamma OS=Homo | 7035902 | 9810676 | 7262179.5 | 7363840 | 8286935 | 7435685.5 | 8766604 | 7769517.5 | **1.024978698** | 4 |
| sp\|P12956 | P12956 | XRCC6 | X-ray repair cross-complementing protein 6 OS=Homo s | 212695808 | 212534304 | 216766176 | 214594544 | 208450304 | 212124704 | 236856752 | 220426560 | **1.024828059** | 8 |
| sp\|Q5VT6 | Q5VT66 | MARC1 | Mitochondrial amidoxime-reducing component 1 OS=Ho | 923683.88 | 962840.88 | 969373.12 | 1154183.75 | 961375.19 | 1027170.69 | 999145.94 | 1119472.88 | **1.024209749** | 6 |
| sp\|Q1469 | Q14692 | BMS1 | Ribosome biogenesis protein BMS1 homolog OS=Homo | 7899598.5 | 7397853 | 7989922 | 7364437.5 | 7059402 | 7926536 | 8852604 | 7554065.5 | **1.024168115** | 7 |
| sp\|Q9262 | Q92621 | NUP205 | Nuclear pore complex protein Nup205 OS=Homo sapien | 12487018 | 11611033 | 11432155 | 11330350 | 10025378 | 12619876 | 10816025 | 14529913 | **1.024127669** | 8 |
| sp\|O9497 | O94979 | SEC31A | Protein transport protein Sec31A OS=Homo sapiens (Hu | 58564672 | 63511972 | 63637864 | 57346008 | 57640012 | 60177912 | 70218680 | 60863036 | **1.024023334** | 7 |
| sp\|Q9BYD | Q9BYD6 | MRPL1 | 39S ribosomal protein L1, mitochondrial OS=Homo sapie | 2080863.75 | 6931571.5 | 9198919 | 11031226 | 7995856 | 5843728 | 7054754 | 9040829 | **1.023684187** | 7 |
| sp\|P13667 | P13667 | PDIA4 | Protein disulfide-isomerase A4 OS=Homo sapiens (Huma | 406675136 | 424139520 | 466192320 | 415149504 | 472075712 | 416503200 | 459797600 | 404254368 | **1.023639428** | 7 |
| sp\|Q9ULA | Q9ULA0 | DNPEP | Aspartyl aminopeptidase OS=Homo sapiens (Human) GN | 13810154 | 9799291 | 11659306 | 11468128 | 12379054 | 13245177 | 12519991 | 9689522 | **1.02346894** | 7 |
| sp\|P09972 | P09972 | ALDOC | Fructose-bisphosphate aldolase C OS=Homo sapiens (Hu | 253372928 | 251380240 | 258612384 | 239422368 | 239003712 | 315363616 | 230031936 | 241851776 | **1.023397889** | 5 |
| sp\|P62266 | P62266 | RPS23 | 40S ribosomal protein S23 OS=Homo sapiens (Human) G | 34274388 | 71482008 | 73173536 | 66771192 | 70437264 | 67777824 | 58812888 | 54419832 | **1.02338892** | 6 |
| sp\|Q1491 | Q14919 | DRAP1 | Dr1-associated corepressor OS=Homo sapiens (Human) | 1418450 | 1604330.88 | 1914376.5 | 1868341.5 | 1553038 | 1985615 | 1587410.5 | 1837866.75 | **1.023279905** | 8 |
| sp\|Q9NQC | Q9NQC3 | RTN4 | Reticulon-4 OS=Homo sapiens (Human) GN=RTN4 PE=1 S | 31577396 | 31722046 | 29601512 | 33448676 | 33989472 | 35114016 | 29983532 | 30199058 | **1.023240654** | 6 |
| sp\|Q0491 | Q04917 | YWHAH | 14-3-3 protein eta OS=Homo sapiens (Human) GN=YWH | 161938240 | 166173168 | 209797648 | 179302032 | 193232608 | 164863248 | 186027344 | 189655584 | **1.023100167** | 6 |
| sp\|P31483 | P31483 | TIA1 | Nucleolysin TIA-1 isoform p40 OS=Homo sapiens (Huma | 13129203 | 11980872 | 13208142 | 14083647 | 12356797 | 11531709 | 15728639 | 13975598 | **1.02272589** | 8 |
| sp\|P18754 | P18754 | RCC1 | Regulator of chromosome condensation OS=Homo sapie | 21197022 | 20794144 | 21120878 | 21294362 | 20503952 | 19527340 | 29172212 | 17082764 | **1.022271556** | 4 |
| sp\|P61019 | P61019 | RAB2A | Ras-related protein Rab-2A OS=Homo sapiens (Human) | 15155888 | 15676309 | 14675268 | 15688108 | 15989806 | 16539523 | 12739641 | 17285504 | **1.022205871** | 4 |
| sp\|P52701 | P52701 | MSH6 | DNA mismatch repair protein Msh6 OS=Homo sapiens (H | 14452854 | 11453837 | 14740915 | 15016710 | 13147706 | 14533520 | 13450149 | 15749313 | **1.021851917** | 8 |
| sp\|Q1469 | Q14694 | USP10 | Ubiquitin carboxyl-terminal hydrolase 10 OS=Homo sapi | 11276669 | 10135406 | 10983795 | 9551412 | 11174266 | 10541772 | 10284545 | 10860616 | **1.021787276** | 7 |
| sp\|P98179 | P98179 | RBM3 | RNA-binding protein 3 OS=Homo sapiens (Human) GN=R | 5069117 | 4037823.5 | 7352980 | 6999071 | 5706541.5 | 5545231 | 7014966 | 5691325.5 | **1.021274252** | 7 |
| sp\|P35241 | P35241 | RDX | Radixin OS=Homo sapiens (Human) GN=RDX PE=1 SV=1 | 65165912 | 66286740 | 69539736 | 63646892 | 73851520 | 63585596 | 68157088 | 64625412 | **1.021086575** | 8 |
| sp\|Q1539 | Q15393 | SF3B3 | Splicing factor 3B subunit 3 OS=Homo sapiens (Human) | 63087208 | 66328272 | 61335332 | 63349372 | 67589976 | 60920420 | 66644984 | 64282360 | **1.021005715** | 5 |
| sp\|P57721 | P57721 | PCBP3 | Poly(rC)-binding protein 3 OS=Homo sapiens (Human) G | 76034096 | 75045008 | 106116416 | 77301256 | 97744816 | 76662136 | 75101896 | 91781552 | **1.020309983** | 7 |
| sp\|P27348 | P27348 | YWHAQ | 14-3-3 protein theta OS=Homo sapiens (Human) GN=YW | 228479824 | 235109104 | 311293760 | 234035296 | 276270304 | 233353888 | 260004848 | 259694960 | **1.020225644** | 6 |
| sp\|Q9UQ3 | Q9UQ35 | SRRM2 | Serine/arginine repetitive matrix protein 2 OS=Homo sa | 22980648 | 20610342 | 26683122 | 18198294 | 23504894 | 20771428 | 23708976 | 22243044 | **1.019847273** | 6 |
| sp\|Q969Q | Q969Q0 | RPL36AL | 60S ribosomal protein L36a-like OS=Homo sapiens (Hum | 31768868 | 28703812 | 27956206 | 33763064 | 39801088 | 28323542 | 30678372 | 25775244 | **1.019529077** | 7 |
| sp\|Q9Y6D | Q9Y6D5 | ARFGEF2 | Brefeldin A-inhibited guanine nucleotide-exchange prote | 4363789.5 | 4358401 | 3592452.75 | 4075664.75 | 3801836.75 | 4304894.5 | 4310910.5 | 4290370.5 | **1.019383666** | 7 |
| sp\|O0040 | O00400 | SLC33A1 | Acetyl-coenzyme A transporter 1 OS=Homo sapiens (Hu | 701956.31 | 697248.12 | 995122.31 | 779836 | 850163.19 | 0 | 944086.19 | 1440187.38 | **1.018988951** | 6 |
| sp\|Q9Y51 | Q9Y512 | SAMM50 | Sorting and assembly machinery component 50 homolog | 11441647 | 10337309 | 12069701 | 11709599 | 12006326 | 11692015 | 9937249 | 12784444 | **1.018915957** | 7 |
| sp\|O0026 | O00267 | SUPT5H | Transcription elongation factor SPT5 OS=Homo sapiens ( | 3483509.75 | 4048644 | 2669004.5 | 3189975.5 | 1824069.75 | 3441963.75 | 4573367.5 | 3798849 | **1.018453721** | 7 |
| sp\|P61803 | P61803 | DAD1 | Dolichyl-diphosphooligosaccharide--protein glycosyltran | 9109986 | 10722412 | 14148084 | 10907372 | 11188963 | 11359045 | 10739618 | 12426068 | **1.01839785** | 5 |

| | | | | | | | | | | | | |
|---|---|---|---|---|---|---|---|---|---|---|---|---|
| sp\|Q9H85 | Q9H853 | TUBA4B | Putative tubulin-like protein alpha-4B OS=Homo sapiens | 104006696 | 110732808 | 134662240 | 113757760 | 95911608 | 114226128 | 105301960 | 156207968 | **1.018326645** | 8 |
| sp\|Q9Y5Q | Q9Y5Q8 | GTF3C5 | General transcription factor 3C polypeptide 5 OS=Homo | 0 | 397835.06 | 371649.28 | 571806.06 | 0 | 418612.91 | 448911.06 | 498283.62 | **1.018278808** | 6.5 |
| sp\|Q96A7 | Q96A72 | MAGOHB | Protein mago nashi homolog 2 OS=Homo sapiens (Huma | 7991006 | 7461105.5 | 7469127.5 | 8566247 | 7445838 | 8356564.5 | 7729495 | 8524456 | **1.018066463** | 8 |
| sp\|Q9NXH | Q9NXH9 | TRMT1 | tRNA (guanine(26)-N(2))-dimethyltransferase OS=Homo | 1520434.25 | 1714642.75 | 1751764.5 | 1439308.75 | 1506076.62 | 1525503 | 2015508 | 1493912.38 | **1.017872248** | 8 |
| sp\|Q8TCS | Q8TCS8 | PNPT1 | Polyribonucleotide nucleotidyltransferase 1, mitochondr | 9849478 | 11461704 | 12525231 | 11886754 | 11554036 | 10770674 | 12208350 | 12001572 | **1.017747349** | 7 |
| sp\|P84085 | P84085 | ARF5 | ADP-ribosylation factor 5 OS=Homo sapiens (Human) GN | 40200668 | 48872068 | 34614392 | 41903020 | 46446100 | 47025488 | 40067704 | 34934324 | **1.017413282** | 8 |
| sp\|P08779 | P08779 | KRT16 | Keratin, type I cytoskeletal 16 OS=Homo sapiens (Humar | 61646000 | 258350464 | 87935328 | 86795512 | 118587504 | 125495264 | 135156192 | 124067232 | **1.01734064** | 4 |
| sp\|P02533 | P02533 | KRT14 | Keratin, type I cytoskeletal 14 OS=Homo sapiens (Humar | 63655772 | 258837856 | 89493608 | 87439216 | 119452920 | 126476864 | 136014336 | 126134704 | **1.017324617** | 4 |
| sp\|O9524 | O95249 | GOSR1 | Golgi SNAP receptor complex member 1 OS=Homo sapie | 1664496.62 | 1763996.62 | 1961151.12 | 0 | 1441129.12 | 1395571 | 1258317 | 1385911.88 | **1.016937043** | 4 |
| sp\|P46109 | P46109 | CRKL | Crk-like protein OS=Homo sapiens (Human) GN=CRKL PE | 6366649 | 5609391 | 5648810.5 | 6292424.5 | 6501704 | 7019679 | 5554552 | 5244964 | **1.016875836** | 8 |
| sp\|Q8NBZ | Q8NBZ7 | UXS1 | UDP-glucuronic acid decarboxylase 1 OS=Homo sapiens | 1497814.75 | 1409009.38 | 1303247.75 | 1336432.12 | 1256695.12 | 1477186.88 | 1668955.75 | 1234810.5 | **1.016432739** | 7 |
| sp\|Q8TCT | Q8TCT9 | HM13 | Minor histocompatibility antigen H13 OS=Homo sapiens | 26664702 | 30164460 | 29447702 | 32078716 | 32369916 | 29486794 | 30406716 | 28017954 | **1.016271307** | 6 |
| sp\|P61204 | P61204 | ARF3 | ADP-ribosylation factor 3 OS=Homo sapiens (Human) GN | 67369360 | 70763976 | 53065408 | 71825224 | 71420360 | 71568160 | 62802552 | 61428892 | **1.015952904** | 8 |
| sp\|P49591 | P49591 | SARS | Serine--tRNA ligase, cytoplasmic OS=Homo sapiens (Hun | 36916272 | 37377020 | 34260316 | 31958528 | 36361692 | 33981068 | 37797088 | 34594244 | **1.015813268** | 7 |
| sp\|O7540 | O75400 | PRPF40A | Pre-mRNA-processing factor 40 homolog A OS=Homo sa | 13297756 | 12698275 | 15236769 | 12596358 | 11020136 | 14249921 | 14500233 | 14875493 | **1.015170681** | 7 |
| sp\|Q1336 | Q13363 | CTBP1 | C-terminal-binding protein 1 OS=Homo sapiens (Human) | 7809770 | 8407298 | 11067303 | 8057500 | 9897355 | 9264953 | 7535542 | 9175707 | **1.015044082** | 7 |
| sp\|Q9UBD | Q9UBD5 | ORC3 | Origin recognition complex subunit 3 OS=Homo sapiens | 790767 | 770140.88 | 383910.31 | 742131.38 | 585553.38 | 820099.81 | 578875.38 | 740715 | **1.014251849** | 7 |
| sp\|P50897 | P50897 | PPT1 | Palmitoyl-protein thioesterase 1 OS=Homo sapiens (Hun | 8703579 | 9953016 | 10912434 | 9449230 | 10352153 | 8687203 | 10973643 | 9560347 | **1.01422634** | 7 |
| sp\|P08134 | P08134 | RHOC | Rho-related GTP-binding protein RhoC OS=Homo sapien | 16202281 | 21724800 | 21140900 | 18725872 | 20117694 | 18912862 | 17592214 | 22274290 | **1.014181159** | 7 |
| sp\|O6082 | O60826 | CCDC22 | Coiled-coil domain-containing protein 22 OS=Homo sapi | 750058.56 | 830494 | 1179595 | 610558.75 | 431942.56 | 826849.88 | 1049429.25 | 1109846.12 | **1.014050913** | 8 |
| sp\|P42330 | P42330 | AKR1C3 | Aldo-keto reductase family 1 member C3 OS=Homo sapi | 424804800 | 384180992 | 415776992 | 412397408 | 472987296 | 410555808 | 463776832 | 312430112 | **1.013798195** | 7 |
| sp\|P39748 | P39748 | FEN1 | Flap endonuclease 1 OS=Homo sapiens (Human) GN=FEI | 32099092 | 29854008 | 31448816 | 34148384 | 35461360 | 31251832 | 34164564 | 28394652 | **1.013501403** | 7 |
| sp\|P04080 | P04080 | CSTB | Cystatin-B OS=Homo sapiens (Human) GN=CSTB PE=1 SV | 12682190 | 9576454 | 8654646 | 10332839 | 9910568 | 12050636 | 9240723 | 10564336 | **1.012610492** | 7 |
| sp\|Q9NZJ | Q9NZJ9 | NUDT4 | Diphosphoinositol polyphosphate phosphohydrolase 2 ( | 2443941.5 | 2251350.5 | 3282736.5 | 2763518 | 1596724.88 | 3374923.5 | 3928106.5 | 1974392.62 | **1.012344684** | 8 |
| sp\|P00568 | P00568 | AK1 | Adenylate kinase isoenzyme 1 OS=Homo sapiens (Huma | 5250001.5 | 5282354.5 | 4035554.75 | 6111251.5 | 5285485.5 | 5112823.5 | 5706798 | 4825477 | **1.012158217** | 8 |
| sp\|P53597 | P53597 | SUCLG1 | Succinate--CoA ligase [ADP/GDP-forming] subunit alpha, | 21771674 | 29779104 | 28033220 | 24896460 | 26042712 | 20161302 | 35159296 | 24359020 | **1.011886165** | 7 |
| sp\|Q1525 | Q15257 | PTPA | Serine/threonine-protein phosphatase 2A activator OS=I | 3757135.25 | 5828946 | 5099931 | 5505192 | 4946615.5 | 5086202 | 4773373 | 5621373 | **1.01170605** | 6 |
| sp\|Q9BW | Q9BW27 | NUP85 | Nuclear pore complex protein Nup85 OS=Homo sapiens | 2086804.75 | 2224036.75 | 2223054.25 | 2332741.75 | 2079204.75 | 1981598 | 2925233.25 | 1983978.25 | **1.01165907** | 4 |
| sp\|Q8WV | Q8WVY7 | UBLCP1 | Ubiquitin-like domain-containing CTD phosphatase 1 OS | 2759300 | 3156030 | 2877991.25 | 3216423 | 3048119.5 | 2810879 | 3436374.75 | 2853075.5 | **1.01154933** | 8 |
| sp\|Q96BN | Q96BN8 | OTULIN | Ubiquitin thioesterase otulin OS=Homo sapiens (Human | 346548.09 | 349613.91 | 257310.78 | 419924.12 | 297471.56 | 410230.91 | 332139.62 | 349128.41 | **1.011339475** | 7 |
| sp\|P10768 | P10768 | ESD | S-formylglutathione hydrolase OS=Homo sapiens (Huma | 23211252 | 20265052 | 22041594 | 22758300 | 21415442 | 24442888 | 22885038 | 20531368 | **1.01131152** | 7 |
| sp\|Q1663 | Q16630 | CPSF6 | Cleavage and polyadenylation specificity factor subunit 6 | 9572978 | 9194441 | 11569662 | 8835800 | 9428803 | 9417442 | 10319163 | 10444032 | **1.011144419** | 6 |
| sp\|Q9289 | Q92890 | UFD1 | Ubiquitin recognition factor in ER-associated degradatior | 3058971.25 | 3338893.75 | 3285978 | 3002871.75 | 3707181 | 2468533.75 | 3197793.25 | 3450762.75 | **1.010842523** | 6 |
| sp\|O0015 | O00151 | PDLIM1 | PDZ and LIM domain protein 1 OS=Homo sapiens (Huma | 56120660 | 44261308 | 47839224 | 50696920 | 53723684 | 45676124 | 55845760 | 45781864 | **1.010603961** | 8 |
| sp\|Q9BYT | Q9BYT8 | NLN | Neurolysin, mitochondrial OS=Homo sapiens (Human) G | 4085801.5 | 5276688 | 5110132 | 4406242 | 5019219 | 4433881 | 4337460 | 5282576 | **1.010290476** | 7 |
| sp\|Q1473 | Q14739 | LBR | Lamin-B receptor OS=Homo sapiens (Human) GN=LBR PI | 14833813 | 17003238 | 16917294 | 17096742 | 17481248 | 16595260 | 15681066 | 16733990 | **1.009726142** | 7 |
| sp\|P00505 | P00505 | GOT2 | Aspartate aminotransferase, mitochondrial OS=Homo sa | 104889632 | 112904648 | 111444408 | 109908568 | 118198424 | 107727192 | 104593256 | 112845376 | **1.009602683** | 8 |
| sp\|Q9Y6D | Q9Y6D6 | ARFGEF1 | Brefeldin A-inhibited guanine nucleotide-exchange prote | 1512662.62 | 1742404 | 1763894.12 | 1505904.75 | 1609042.25 | 1474490.75 | 1744673.25 | 1759111.5 | **1.009571425** | 8 |
| sp\|P34931 | P34931 | HSPA1L | Heat shock 70 kDa protein 1-like OS=Homo sapiens (Hun | 665399936 | 386147584 | 772374976 | 331835296 | 563393792 | 540678720 | 551947712 | 519730976 | **1.009274422** | 8 |
| sp\|P63104 | P63104 | YWHAZ | 14-3-3 protein zeta/delta OS=Homo sapiens (Human) GN | 259117104 | 280056160 | 364294400 | 290295776 | 314749568 | 265674800 | 308630336 | 315393728 | **1.008950678** | 6 |
| sp\|Q96KG | Q96KG9 | SCYL1 | N-terminal kinase-like protein OS=Homo sapiens (Humai | 8051127 | 7212750 | 7993482.5 | 8419028 | 6707557 | 8692334 | 8762740 | 7791095 | **1.00875537** | 7 |
| sp\|P35580 | P35580 | MYH10 | Myosin-10 OS=Homo sapiens (Human) GN=MYH10 PE=1 | 89361688 | 86446720 | 90559608 | 75726432 | 82256968 | 80866624 | 93224248 | 88723008 | **1.008700521** | 8 |
| sp\|Q5VIR6 | Q5VIR6 | VPS53 | Vacuolar protein sorting-associated protein 53 homolog | 435091.41 | 358141.47 | 332209.12 | 394273.53 | 388482.34 | 442044.69 | 387090.94 | 314901.59 | **1.008425281** | 8 |
| sp\|P51784 | P51784 | USP11 | Ubiquitin carboxyl-terminal hydrolase 11 OS=Homo sapi | 3171952.25 | 3487613 | 2602518 | 3293139.75 | 3204038.75 | 2919249 | 2760026 | 3776637.5 | **1.008341409** | 8 |
| sp\|P98175 | P98175 | RBM10 | RNA-binding protein 10 OS=Homo sapiens (Human) GN= | 564865.44 | 883665.75 | 698063.25 | 601568.75 | 689133.5 | 537341.38 | 796111.25 | 747480.19 | **1.007970098** | 8 |

| | | | | | | | | | | | | |
|---|---|---|---|---|---|---|---|---|---|---|---|---|
| sp\|P54578 | P54578 | USP14 | Ubiquitin carboxyl-terminal hydrolase 14 OS=Homo sapi | 31504680 | 18274550 | 32860668 | 33710096 | 15564328 | 31953024 | 35241456 | 34497496 | **1.007789515** | 6 |
| sp\|Q0316 | Q03167 | TGFBR3 | Transforming growth factor beta receptor type 3 OS=Ho | 1073537.12 | 840774.31 | 455273 | 435762.34 | 700805.44 | 665716.25 | 830587.88 | 629692.38 | **1.00764796** | 8 |
| sp\|Q1278 | Q12788 | TBL3 | Transducin beta-like protein 3 OS=Homo sapiens (Huma | 5229901 | 4172950.5 | 2963006.5 | 4593100.5 | 3861297.5 | 4320439.5 | 4629853.5 | 4272371 | **1.007370913** | 8 |
| sp\|O9485 | O94855 | SEC24D | Protein transport protein Sec24D OS=Homo sapiens (Hu | 5631138 | 4227071 | 4370563.5 | 4287157 | 4907803 | 4395005.5 | 4990927.5 | 4349625.5 | **1.00688229** | 5 |
| sp\|P25787 | P25787 | PSMA2 | Proteasome subunit alpha type-2 OS=Homo sapiens (Hu | 35886360 | 31984284 | 32017350 | 40082640 | 29819906 | 36588636 | 37986308 | 36529688 | **1.00681503** | 7 |
| sp\|Q1685 | Q16851 | UGP2 | UTP--glucose-1-phosphate uridylyltransferase OS=Homo | 73278840 | 71582792 | 72636816 | 75364448 | 76709976 | 79735960 | 72120360 | 66221688 | **1.006573342** | 7 |
| sp\|P55265 | P55265 | ADAR | Double-stranded RNA-specific adenosine deaminase OS | 9975868 | 7463183 | 9353866 | 9063214 | 8847121 | 8156735 | 9093710 | 9987578 | **1.006386997** | 8 |
| sp\|Q1320 | Q13206 | DDX10 | Probable ATP-dependent RNA helicase DDX10 OS=Homo | 2814501.75 | 2516281.5 | 1934002.75 | 2049485 | 2567215.25 | 2624904.25 | 2723085.5 | 1457099.12 | **1.00623056** | 7 |
| sp\|P09525 | P09525 | ANXA4 | Annexin A4 OS=Homo sapiens (Human) GN=ANXA4 PE=1 | 124106632 | 120498744 | 126768264 | 133406936 | 122409928 | 125790984 | 127898080 | 131624920 | **1.005830922** | 7 |
| sp\|Q9UBT | Q9UBT2 | UBA2 | SUMO-activating enzyme subunit 2 OS=Homo sapiens (H | 22845178 | 23822088 | 24112882 | 24007100 | 24303508 | 20624380 | 22818474 | 27589390 | **1.005786686** | 8 |
| sp\|Q7L5N | Q7L5N1 | COPS6 | COP9 signalosome complex subunit 6 OS=Homo sapiens | 3865346.75 | 4994232 | 3156421.75 | 4589000.5 | 3254737.25 | 4386286.5 | 4262098 | 4794648.5 | **1.005586826** | 8 |
| sp\|P46781 | P46781 | RPS9 | 40S ribosomal protein S9 OS=Homo sapiens (Human) GN | 150152816 | 129397920 | 162345696 | 155650688 | 149070352 | 149753024 | 150225008 | 151717024 | **1.005385831** | 6 |
| sp\|O7516 | O75165 | DNAJC13 | DnaJ homolog subfamily C member 13 OS=Homo sapien | 293066.38 | 265127.53 | 318310.41 | 283440 | 232770.55 | 323697.34 | 307078.19 | 302342.03 | **1.005124203** | 6 |
| sp\|Q7L57 | Q7L576 | CYFIP1 | Cytoplasmic FMR1-interacting protein 1 OS=Homo sapie | 14548535 | 13230706 | 15541588 | 13780528 | 13252278 | 14524634 | 13545737 | 16057488 | **1.004882196** | 8 |
| sp\|O9577 | O95777 | LSM8 | U6 snRNA-associated Sm-like protein LSm8 OS=Homo sa | 2676424 | 1916256.62 | 2550365 | 3083662.5 | 2383404.5 | 2393642.5 | 2611683.5 | 2884413.5 | **1.004540648** | 7 |
| sp\|O7603 | O76031 | CLPX | ATP-dependent Clp protease ATP-binding subunit clpX-lil | 12899161 | 11870541 | 11449280 | 11706373 | 12941669 | 11549652 | 11507579 | 12134101 | **1.004332696** | 7 |
| sp\|P43487 | P43487 | RANBP1 | Ran-specific GTPase-activating protein OS=Homo sapien | 28221300 | 21267092 | 31943226 | 26728542 | 29292888 | 24253550 | 28857598 | 26219762 | **1.004286588** | 8 |
| sp\|Q9UJ7 | Q9UJ70 | NAGK | N-acetyl-D-glucosamine kinase OS=Homo sapiens (Huma | 1836299.25 | 1220327.12 | 1990270.62 | 1549478.12 | 1318236.75 | 2184289.5 | 1526717.38 | 1595193.25 | **1.00425412** | 8 |
| sp\|Q969G | Q969G3 | SMARCE1 | SWI/SNF-related matrix-associated actin-dependent regu | 3033479 | 2734762.75 | 1948932.88 | 2731413 | 2347284.5 | 2592946.25 | 2799412.5 | 2748681.75 | **1.003803133** | 8 |
| sp\|Q9NUL | Q9NUL7 | DDX28 | Probable ATP-dependent RNA helicase DDX28 OS=Homo | 2160004 | 2203495.5 | 2028832.75 | 1813378.38 | 2202804.5 | 1963030.88 | 2244727 | 1825961.62 | **1.003755113** | 7 |
| sp\|Q1526 | Q15269 | PWP2 | Periodic tryptophan protein 2 homolog OS=Homo sapier | 7536001 | 6830027 | 6724134.5 | 7250778.5 | 6590060 | 7289951 | 6566177 | 7981569 | **1.003063272** | 7 |
| sp\|O6028 | O60287 | URB1 | Nucleolar pre-ribosomal-associated protein 1 OS=Homo | 953897.38 | 1049250.75 | 1020542.25 | 1160724.75 | 1086545.5 | 861532.5 | 1057052.62 | 1191132.12 | **1.002831366** | 6 |
| sp\|P10619 | P10619 | CTSA | Lysosomal protective protein OS=Homo sapiens (Human | 10775071 | 13437834 | 13599469 | 12487641 | 14582046 | 10504762 | 11882708 | 13455268 | **1.002480496** | 8 |
| sp\|P35268 | P35268 | RPL22 | 60S ribosomal protein L22 OS=Homo sapiens (Human) G | 107827272 | 95869456 | 129050760 | 115613168 | 115480864 | 102460272 | 106450560 | 125081104 | **1.002480467** | 7 |
| sp\|P33316 | P33316 | DUT | Deoxyuridine 5'-triphosphate nucleotidohydrolase, mito | 10411362 | 6318178 | 10653020 | 10732536 | 11486076 | 10757818 | 6347451.5 | 9615539 | **1.002408193** | 6 |
| sp\|Q9P2E | Q9P2E9 | RRBP1 | Ribosome-binding protein 1 OS=Homo sapiens (Human) | 334230848 | 361027968 | 378188064 | 271774272 | 339901024 | 308817088 | 359739168 | 339932992 | **1.002355836** | 7 |
| sp\|P25786 | P25786 | PSMA1 | Proteasome subunit alpha type-1 OS=Homo sapiens (Hu | 47198944 | 53405528 | 51238904 | 47935848 | 53085976 | 52695888 | 51147112 | 43275328 | **1.002127749** | 8 |
| sp\|P78406 | P78406 | RAE1 | mRNA export factor OS=Homo sapiens (Human) GN=RAE | 9453560 | 8855410 | 13124011 | 10537928 | 11501142 | 9524600 | 10458100 | 10559092 | **1.00171607** | 6 |
| sp\|Q29RF | Q29RF7 | PDS5A | Sister chromatid cohesion protein PDS5 homolog A OS=H | 15080212 | 13188133 | 13827845 | 13076388 | 14291621 | 13412881 | 13469362 | 14093020 | **1.001709291** | 6 |
| sp\|P08708 | P08708 | RPS17 | 40S ribosomal protein S17 OS=Homo sapiens (Human) G | 55038448 | 53760420 | 45420672 | 58129616 | 56319504 | 54041124 | 52939504 | 49382236 | **1.00156917** | 7 |
| sp\|Q0076 | Q00765 | REEP5 | Receptor expression-enhancing protein 5 OS=Homo sapi | 6832556 | 8591562 | 7882165 | 8700313 | 10329236 | 6804912 | 7396270.5 | 7523078 | **1.001465339** | 6 |
| sp\|Q9BRJ7 | Q9BRJ7 | NUDT16L1 | Tudor-interacting repair regulator protein OS=Homo sap | 756944.62 | 820631.88 | 728833.56 | 839310.38 | 710553.31 | 744326.56 | 778691.44 | 915979.56 | **1.001217664** | 7 |
| sp\|O9488 | O94888 | UBXN7 | UBX domain-containing protein 7 OS=Homo sapiens (Hu | 1570507.62 | 1248719.5 | 1527828.75 | 864809.62 | 1454628.25 | 1410418.62 | 931227.25 | 1421565.5 | **1.001146256** | 7 |
| sp\|P61160 | P61160 | ACTR2 | Actin-related protein 2 OS=Homo sapiens (Human) GN=/ | 9280721 | 13949041 | 6370508 | 9813242 | 13886949 | 7875364 | 8723496 | 8972824 | **1.00114481** | 6 |
| sp\|Q9UKV | Q9UKV3 | ACIN1 | Apoptotic chromatin condensation inducer in the nucleu | 23764560 | 20081450 | 16637933 | 17946012 | 23920600 | 15601640 | 24217660 | 14774609 | **1.001078083** | 8 |
| sp\|Q8N1G | Q8N1G2 | CMTR1 | Cap-specific mRNA (nucleoside-2'-O-)-methyltransferase | 2887705 | 2982167.25 | 3201476.5 | 2647209.5 | 2182852.75 | 3174400.75 | 3141438 | 3227341.5 | **1.000637856** | 6 |
| sp\|Q9H9T | Q9H9T3 | ELP3 | Elongator complex protein 3 OS=Homo sapiens (Human) | 2041544 | 2985243 | 2641280 | 3020393.5 | 3146551.5 | 2360207.75 | 1974206.5 | 3208094.25 | **1.000056089** | 7 |
| sp\|P62314 | P62314 | SNRPD1 | Small nuclear ribonucleoprotein Sm D1 OS=Homo sapier | 28704662 | 29572562 | 27613256 | 38062468 | 29889014 | 30992162 | 31014530 | 32059028 | **1.000014409** | 4 |
| sp\|Q9Y26 | Q9Y266 | NUDC | Nuclear migration protein nudC OS=Homo sapiens (Hum | 62000524 | 67485216 | 82065560 | 60088796 | 71291120 | 64099008 | 73256768 | 62830552 | **0.999401237** | 6 |
| sp\|Q96ST | Q96ST3 | SIN3A | Paired amphipathic helix protein Sin3a OS=Homo sapien | 974092.38 | 782596.5 | 0 | 269849.09 | 595820.12 | 679605.25 | 154311.3 | 595541.12 | **0.999378161** | 7 |
| sp\|P49257 | P49257 | LMAN1 | Protein ERGIC-53 OS=Homo sapiens (Human) GN=LMAN | 41362048 | 44004216 | 41235680 | 40840832 | 42716740 | 42948504 | 43003428 | 38646280 | **0.999236611** | 7 |
| sp\|Q1688 | Q16881 | TXNRD1 | Thioredoxin reductase 1, cytoplasmic OS=Homo sapiens | 23304482 | 24507990 | 23658148 | 22318934 | 22147040 | 23991878 | 26307770 | 21247146 | **0.998979417** | 7 |
| sp\|P01019 | P01019 | AGT | Angiotensinogen OS=Homo sapiens (Human) GN=AGT PI | 6026215 | 5538812.5 | 6464385 | 6134570.5 | 5895347 | 4397167 | 6787344.5 | 7052169.5 | **0.998677577** | 7 |
| sp\|Q8TD1 | Q8TD19 | NEK9 | Serine/threonine-protein kinase Nek9 OS=Homo sapiens | 1035002.5 | 1077416 | 1225366.5 | 1472439.5 | 1000332.12 | 1019903.38 | 1475232.62 | 1307073.25 | **0.99840275** | 7 |
| sp\|P00374 | P00374 | DHFR | Dihydrofolate reductase OS=Homo sapiens (Human) GN | 1119478.62 | 922309.12 | 1341644.12 | 1055976.25 | 1287878.25 | 984175 | 934606.12 | 1225242.88 | **0.998309266** | 8 |

| | | | | | | | | | | | | | |
|---|---|---|---|---|---|---|---|---|---|---|---|---|---|
| sp\|Q9Y3T | Q9Y3T9 | NOC2L | Nucleolar complex protein 2 homolog OS=Homo sapiens | 8153721.5 | 8813866 | 6340069.5 | 9410169 | 9551283 | 7288066 | 5671190 | 10140294 | **0.997952401** | 7 |
| sp\|P19388 | P19388 | POLR2E | DNA-directed RNA polymerases I, II, and III subunit RPAB | 5870056.5 | 5852452 | 7259855 | 6040332 | 7141829.5 | 4778744 | 6739133 | 6306515 | **0.997743089** | 7 |
| sp\|O0018 | O00186 | STXBP3 | Syntaxin-binding protein 3 OS=Homo sapiens (Human) G | 1547944.75 | 1459455.25 | 1471720.12 | 1606778.38 | 1722217 | 1459707.88 | 1421345.12 | 1468453 | **0.997670763** | 6 |
| sp\|Q9261 | Q92616 | GCN1 | eIF-2-alpha kinase activator GCN1 OS=Homo sapiens (Hu | 206232560 | 122458824 | 123429000 | 210794800 | 167463408 | 161627984 | 162908400 | 169347904 | **0.997635462** | 8 |
| sp\|Q1469 | Q14690 | PDCD11 | Protein RRP5 homolog OS=Homo sapiens (Human) GN=F | 29409520 | 24003944 | 25907788 | 28127722 | 26806372 | 26065080 | 28557482 | 25746686 | **0.997455965** | 8 |
| sp\|P55899 | P55899 | FCGRT | IgG receptor FcRn large subunit p51 OS=Homo sapiens (I | 1574697.25 | 1638105.25 | 766508.44 | 1849147.5 | 1346164.25 | 1050645.38 | 1194110.88 | 2222250.25 | **0.997377063** | 7 |
| sp\|Q9Y2T | Q9Y2T2 | AP3M1 | AP-3 complex subunit mu-1 OS=Homo sapiens (Human) | 4602570 | 6066981.5 | 6627458.5 | 6705616.5 | 5496045.5 | 6541489 | 5319964.5 | 6572942.5 | **0.996992621** | 6 |
| sp\|A2RRP | A2RRP1 | NBAS | Neuroblastoma-amplified sequence OS=Homo sapiens (I | 898114.81 | 1735969.5 | 636563.81 | 1201549.88 | 734193.38 | 1391088 | 909197.06 | 1421718.75 | **0.99642216** | 7 |
| sp\|P61923 | P61923 | COPZ1 | Coatomer subunit zeta-1 OS=Homo sapiens (Human) GN | 22311624 | 17086846 | 19171874 | 21360694 | 21480348 | 20388778 | 20455196 | 17290710 | **0.996046517** | 8 |
| sp\|Q0683 | Q06830 | PRDX1 | Peroxiredoxin-1 OS=Homo sapiens (Human) GN=PRDX1 | 567022208 | 619309312 | 569940736 | 560327424 | 701628480 | 518698432 | 578659584 | 507988160 | **0.995845193** | 7 |
| sp\|P05362 | P05362 | ICAM1 | Intercellular adhesion molecule 1 OS=Homo sapiens (Hu | 61211968 | 72768360 | 55447584 | 61322444 | 53547320 | 53287956 | 70956232 | 71843776 | **0.995553059** | 6 |
| sp\|Q9UBQ | Q9UBQ5 | EIF3K | Eukaryotic translation initiation factor 3 subunit K OS=H | 10477782 | 10065763 | 10480502 | 11709666 | 10142119 | 11023604 | 10720658 | 10650649 | **0.995397475** | 6 |
| sp\|Q9H0D | Q9H0D6 | XRN2 | 5'-3' exoribonuclease 2 OS=Homo sapiens (Human) GN= | 31811376 | 30437756 | 29307878 | 29049288 | 32864968 | 29404676 | 30646090 | 27090110 | **0.995021371** | 7 |
| sp\|Q8NFH | Q8NFH4 | NUP37 | Nucleoporin Nup37 OS=Homo sapiens (Human) GN=NUF | 736530 | 567810.56 | 574286.38 | 895191.38 | 828533.81 | 712598.88 | 599801.94 | 618708.94 | **0.994889806** | 7 |
| sp\|P62333 | P62333 | PSMC6 | 26S proteasome regulatory subunit 10B OS=Homo sapie | 33827520 | 28961414 | 34225340 | 32886550 | 31493106 | 33918216 | 34776144 | 29013316 | **0.99461095** | 7 |
| sp\|Q9Y6W | Q9Y6W5 | WASF2 | Wiskott-Aldrich syndrome protein family member 2 OS= | 3040984.5 | 3480502.5 | 2881292 | 2767242 | 3258949.5 | 1886423.62 | 3447554 | 3509882 | **0.994477259** | 6 |
| sp\|P78346 | P78346 | RPP30 | Ribonuclease P protein subunit p30 OS=Homo sapiens (H | 5145017.5 | 6848621 | 6282608 | 5810796.5 | 5969361 | 5659047 | 5845412 | 6478626 | **0.994412058** | 8 |
| sp\|Q7L5D | Q7L5D6 | GET4 | Golgi to ER traffic protein 4 homolog OS=Homo sapiens ( | 1653093.88 | 2467742.5 | 2565288 | 2429153.5 | 2546133 | 2388487.75 | 1703135.62 | 2426168.25 | **0.994366243** | 6 |
| sp\|Q96PY | Q96PY5 | FMNL2 | Formin-like protein 2 OS=Homo sapiens (Human) GN=FN | 3060876.5 | 2588016.75 | 2251877.5 | 2520248.25 | 1951547.62 | 3254722.75 | 2414327.25 | 2741158.5 | **0.99431314** | 8 |
| sp\|O4358 | O43583 | DENR | Density-regulated protein OS=Homo sapiens (Human) GI | 785798.5 | 634420.88 | 477852.28 | 617233.62 | 0 | 850689.75 | 796056.12 | 853934.38 | **0.994185585** | 4 |
| sp\|Q96HV | Q96HV5 | TMEM41A | Transmembrane protein 41A OS=Homo sapiens (Human | 0 | 873740.75 | 917608.44 | 1119626.75 | 1004188.94 | 807308.69 | 0 | 1082331 | **0.994109429** | 7.5 |
| sp\|Q1483 | Q14839 | CHD4 | Chromodomain-helicase-DNA-binding protein 4 OS=Hon | 16444557 | 15754959 | 20067856 | 14251258 | 18185394 | 14486140 | 17825610 | 15624682 | **0.994034694** | 8 |
| sp\|Q6FI81 | Q6FI81 | CIAPIN1 | Anamorsin OS=Homo sapiens (Human) GN=CIAPIN1 PE= | 1971907.38 | 1002938.31 | 1801148.25 | 633551.75 | 1153621.75 | 1986184.25 | 1152722.88 | 1080755.25 | **0.993296746** | 6 |
| sp\|Q9ULU | Q9ULU4 | ZMYND8 | Protein kinase C-binding protein 1 OS=Homo sapiens (Hu | 714763.31 | 843408.19 | 762086.75 | 911858.69 | 609199.94 | 810490.38 | 951407.19 | 838779.62 | **0.99311912** | 8 |
| sp\|Q1410 | Q14103 | HNRNPD | Heterogeneous nuclear ribonucleoprotein D0 OS=Homo | 152449120 | 146711728 | 143521712 | 153536144 | 158807616 | 139695392 | 162178400 | 131373848 | **0.993016911** | 8 |
| sp\|Q1590 | Q15907 | RAB11B | Ras-related protein Rab-11B OS=Homo sapiens (Human) | 26023276 | 27932760 | 28447880 | 26406116 | 28633236 | 30624440 | 21502680 | 27289180 | **0.993010792** | 6 |
| sp\|O9581 | O95819 | MAP4K4 | Mitogen-activated protein kinase kinase kinase kinase 4 | 10017893 | 7218212 | 9274119 | 6662127.5 | 9413975 | 6692268.5 | 7253890.5 | 9580300 | **0.993008711** | 7 |
| sp\|P11233 | P11233 | RALA | Ras-related protein Ral-A OS=Homo sapiens (Human) GN | 4590848 | 4766889.5 | 4678357.5 | 4582173.5 | 3632049.5 | 6624863 | 3680433.25 | 4548876 | **0.992907678** | 4 |
| sp\|Q9BY7 | Q9BY77 | POLDIP3 | Polymerase delta-interacting protein 3 OS=Homo sapien | 13007983 | 11102967 | 11087098 | 12468073 | 12194057 | 11772118 | 11592912 | 11766472 | **0.992855261** | 8 |
| sp\|Q0702 | Q07021 | C1QBP | Complement component 1 Q subcomponent-binding pro | 48411712 | 46805980 | 45282700 | 40434072 | 38998996 | 49667088 | 46399236 | 44483252 | **0.992340365** | 7 |
| sp\|Q8NDI | Q8NDI1 | EHBP1 | EH domain-binding protein 1 OS=Homo sapiens (Human | 5884508 | 3886339.25 | 5186388 | 4740863 | 4676892 | 5178594.5 | 5385493 | 4302795.5 | **0.992165576** | 7 |
| sp\|Q9UKF | Q9UKF6 | CPSF3 | Cleavage and polyadenylation specificity factor subunit 3 | 6678881.5 | 8041008.5 | 6433623.5 | 7670272 | 7161375 | 7244159 | 7042267.5 | 7144309 | **0.991962367** | 8 |
| sp\|P13010 | P13010 | XRCC5 | X-ray repair cross-complementing protein 5 OS=Homo sa | 151426944 | 153224208 | 144234064 | 152708368 | 142235968 | 156912544 | 148670992 | 148895328 | **0.991890286** | 6 |
| sp\|O4325 | O43251 | RBFOX2 | RNA binding protein fox-1 homolog 2 OS=Homo sapiens | 374077.72 | 331846.59 | 590873.75 | 983040.88 | 834091.31 | 334320.5 | 611700.19 | 480821.12 | **0.991707388** | 7 |
| sp\|P46459 | P46459 | NSF | Vesicle-fusing ATPase OS=Homo sapiens (Human) GN=N | 28499642 | 27246312 | 25421668 | 23458394 | 24236782 | 26830010 | 23157436 | 29481430 | **0.991203354** | 7 |
| sp\|P55735 | P55735 | SEC13 | Protein SEC13 homolog OS=Homo sapiens (Human) GN= | 26907014 | 27377996 | 27256816 | 27409640 | 28760656 | 23896130 | 27355500 | 27979452 | **0.991191234** | 6 |
| sp\|Q9BY4 | Q9BY42 | RTFDC1 | Protein RTF2 homolog OS=Homo sapiens (Human) GN=R | 857381.88 | 758530.31 | 887509.62 | 635121.62 | 801345.69 | 770622.31 | 771422.25 | 767082.06 | **0.991056004** | 8 |
| sp\|Q9NR5 | Q9NR50 | EIF2B3 | Translation initiation factor eIF-2B subunit gamma OS=H | 2246771.5 | 2272425.75 | 2216781.25 | 2633446 | 2092090.38 | 2414574.75 | 2273166.5 | 2504539 | **0.990922188** | 7 |
| sp\|P60660 | P60660 | MYL6 | Myosin light polypeptide 6 OS=Homo sapiens (Human) G | 35735060 | 35396984 | 47275968 | 39368304 | 38147456 | 36424296 | 41914464 | 39827112 | **0.990727455** | 6 |
| sp\|Q0482 | Q04828 | AKR1C1 | Aldo-keto reductase family 1 member C1 OS=Homo sapi | 567500864 | 533451168 | 568572544 | 545882048 | 614884672 | 526322240 | 616433280 | 436805728 | **0.990538665** | 8 |
| sp\|Q9P03 | Q9P032 | NDUFAF4 | NADH dehydrogenase [ubiquinone] 1 alpha subcomplex | 758430.31 | 838160.12 | 824802.38 | 838962.12 | 847106.56 | 846939.25 | 901121.38 | 634174.25 | **0.990487695** | 4 |
| sp\|P18085 | P18085 | ARF4 | ADP-ribosylation factor 4 OS=Homo sapiens (Human) GN | 63763828 | 66101404 | 52818532 | 66861704 | 64587152 | 63668936 | 56187852 | 62712700 | **0.990427284** | 5 |
| sp\|P62879 | P62879 | GNB2 | Guanine nucleotide-binding protein G(I)/G(S)/G(T) subun | 13114058 | 15309894 | 18844914 | 13967316 | 14901941 | 13139992 | 14650239 | 17938562 | **0.990112904** | 8 |
| sp\|Q7KZF | Q7KZF4 | SND1 | Staphylococcal nuclease domain-containing protein 1 OS | 155537664 | 150372560 | 157145312 | 144424128 | 146011328 | 146841008 | 148684512 | 159925984 | **0.990095418** | 7 |
| sp\|Q9290 | Q92905 | COPS5 | COP9 signalosome complex subunit 5 OS=Homo sapiens | 5983078.5 | 5931438 | 6594543 | 7668722 | 5865092.5 | 5057298 | 7319172 | 7676649.5 | **0.990084358** | 7 |

| | | | | | | | | | | | | | |
|---|---|---|---|---|---|---|---|---|---|---|---|---|---|
| sp\|P02787 | P02787 | TF | Serotransferrin OS=Homo sapiens (Human) GN=TF PE=1 | 116961784 | 115811072 | 112758848 | 104605896 | 116320632 | 123034480 | 111286112 | 94887864 | **0.989761993** | 8 |
| sp\|P0DMV | P0DMV8 | HSPA1A | Heat shock 70 kDa protein 1A OS=Homo sapiens (Humar | 819714880 | 537569408 | 960171392 | 506081152 | 726536576 | 656932480 | 730407424 | 680733440 | **0.989755079** | 8 |
| sp\|Q9966 | Q99661 | KIF2C | Kinesin-like protein KIF2C OS=Homo sapiens (Human) GI | 1682697.25 | 320987 | 1170724.88 | 1206499.75 | 1481634.75 | 1764022 | 347069.12 | 742712.81 | **0.98962083** | 7 |
| sp\|P23193 | P23193 | TCEA1 | Transcription elongation factor A protein 1 OS=Homo sa | 15625946 | 13168651 | 7512090 | 12105035 | 14899683 | 10046642 | 12723356 | 10232421 | **0.989473211** | 7 |
| sp\|P83731 | P83731 | RPL24 | 60S ribosomal protein L24 OS=Homo sapiens (Human) G | 92767112 | 89584792 | 88396928 | 91091472 | 95135664 | 81042840 | 93252248 | 88459552 | **0.989083582** | 7 |
| sp\|Q6PKG | Q6PKG0 | LARP1 | La-related protein 1 OS=Homo sapiens (Human) GN=LAR | 7749688 | 7282134 | 7105788 | 6250041 | 6740225 | 7027633 | 7156465 | 7153070.5 | **0.989070688** | 6 |
| sp\|Q9288 | Q92888 | ARHGEF1 | Rho guanine nucleotide exchange factor 1 OS=Homo sap | 1738164.88 | 1425980.88 | 1066991 | 1271765.75 | 1338409.62 | 1301444 | 1516700.12 | 1284907.25 | **0.988834707** | 7 |
| sp\|Q9BTV | Q9BTV4 | TMEM43 | Transmembrane protein 43 OS=Homo sapiens (Human) | 14369456 | 15070755 | 14416134 | 16016652 | 15015492 | 15896240 | 13940311 | 14335871 | **0.98855773** | 5 |
| sp\|Q9Y3C | Q9Y3C4 | TPRKB | EKC/KEOPS complex subunit TPRKB OS=Homo sapiens (H | 1073992.75 | 1649635.25 | 1561425.88 | 1640483.88 | 1573997.5 | 1322651.25 | 1517600.25 | 1442771.5 | **0.988436955** | 5 |
| sp\|Q9UHX | Q9UHX1 | PUF60 | Poly(U)-binding-splicing factor PUF60 OS=Homo sapiens | 19166910 | 18323040 | 18718474 | 19361710 | 20704646 | 18598148 | 17402720 | 17959644 | **0.988024687** | 5 |
| sp\|Q9UHB | Q9UHB9 | SRP68 | Signal recognition particle subunit SRP68 OS=Homo sapi | 28474018 | 28726646 | 33611392 | 30232934 | 30150394 | 30558980 | 29264454 | 29570218 | **0.987600115** | 7 |
| sp\|P51398 | P51398 | DAP3 | 28S ribosomal protein S29, mitochondrial OS=Homo sap | 13253777 | 12540226 | 9087993 | 11526817 | 11418484 | 11646501 | 11749294 | 11008646 | **0.987375501** | 6 |
| sp\|Q9Y3B | Q9Y3B9 | RRP15 | RRP15-like protein OS=Homo sapiens (Human) GN=RRP1 | 1858488.88 | 1926534 | 1767467.5 | 2031239.88 | 1963223.62 | 1629818.75 | 2237540.5 | 1656834.5 | **0.987300064** | 7 |
| sp\|Q8WW | Q8WWI1 | LMO7 | LIM domain only protein 7 OS=Homo sapiens (Human) G | 13446841 | 13348212 | 15580741 | 12504124 | 12891363 | 13098295 | 14884692 | 13271862 | **0.986630702** | 6 |
| sp\|Q9956 | Q99567 | NUP88 | Nuclear pore complex protein Nup88 OS=Homo sapiens | 909430.06 | 1185998.25 | 1120353.5 | 901779.12 | 832185.88 | 1137871.62 | 1252759.88 | 839415.5 | **0.986562907** | 7 |
| sp\|O9478 | O94788 | ALDH1A2 | Retinal dehydrogenase 2 OS=Homo sapiens (Human) GN | 83688144 | 88767632 | 107302832 | 94710048 | 100968184 | 95902504 | 87098864 | 85457392 | **0.986536358** | 8 |
| sp\|P13798 | P13798 | APEH | Acylamino-acid-releasing enzyme OS=Homo sapiens (Hu | 30160156 | 31399398 | 29121560 | 35037284 | 28671776 | 33616656 | 28510300 | 33157424 | **0.985982624** | 6 |
| sp\|Q96AX | Q96AX1 | VPS33A | Vacuolar protein sorting-associated protein 33A OS=Hor | 302390.72 | 252800.31 | 136802.7 | 327735.56 | 215046.8 | 266292.19 | 244917.28 | 278997.19 | **0.985804242** | 6 |
| sp\|P47755 | P47755 | CAPZA2 | F-actin-capping protein subunit alpha-2 OS=Homo sapie | 38073180 | 39380648 | 41381880 | 39243824 | 38871796 | 35956396 | 41593220 | 39389336 | **0.985647832** | 8 |
| sp\|O4315 | O43159 | RRP8 | Ribosomal RNA-processing protein 8 OS=Homo sapiens ( | 2561833.75 | 2494961.5 | 2686066.25 | 2913748 | 2948797.5 | 2612213.5 | 2349570 | 2591673.25 | **0.985515539** | 8 |
| sp\|Q96BP | Q96BP3 | PPWD1 | Peptidylprolyl isomerase domain and WD repeat-contain | 1556113.88 | 1989448 | 1454746.88 | 1361952.38 | 1395696.62 | 1263559.38 | 3192772.75 | 417612.75 | **0.985442339** | 5 |
| sp\|Q9261 | Q92614 | MYO18A | Unconventional myosin-XVIIIa OS=Homo sapiens (Huma | 3162055.25 | 3228034.5 | 2054145.88 | 2765169.25 | 2248241 | 2334695.25 | 2946528 | 3515139.25 | **0.985297937** | 8 |
| sp\|P49585 | P49585 | PCYT1A | Choline-phosphate cytidylyltransferase A OS=Homo sapi | 2116387 | 1528909.75 | 7289489.5 | 1419934.5 | 1862684.5 | 1431868.75 | 7075635.5 | 1801968.25 | **0.985223159** | 8 |
| sp\|Q0626 | Q06265 | EXOSC9 | Exosome complex component RRP45 OS=Homo sapiens | 4194132.75 | 3554047.75 | 4805892 | 3115337 | 4050091.5 | 3508793.5 | 4002276 | 3875547.5 | **0.985149345** | 7 |
| sp\|Q9HDC | Q9HDC9 | APMAP | Adipocyte plasma membrane-associated protein OS=Ho | 68948056 | 66101000 | 67431848 | 72919912 | 73355856 | 67455544 | 55507164 | 74868528 | **0.984699668** | 6 |
| sp\|Q86YS | Q86YS6 | RAB43 | Ras-related protein Rab-43 OS=Homo sapiens (Human) ( | 618848.69 | 640507.19 | 797957.69 | 882900.88 | 926106.12 | 858798.69 | 512341.94 | 597496.88 | **0.984534863** | 7 |
| sp\|Q9Y6E | Q9Y6E2 | BZW2 | Basic leucine zipper and W2 domain-containing protein 2 | 11609054 | 13116616 | 18327262 | 15286548 | 14821320 | 15486551 | 10757605 | 16368998 | **0.984487246** | 8 |
| sp\|P26196 | P26196 | DDX6 | Probable ATP-dependent RNA helicase DDX6 OS=Homo | 22390864 | 18079884 | 18103576 | 20198470 | 20680842 | 16896676 | 19954286 | 20009662 | **0.984368613** | 7 |
| sp\|Q9275 | Q92754 | TFAP2C | Transcription factor AP-2 gamma OS=Homo sapiens (Hu | 12335869 | 12463233 | 17282760 | 15572058 | 16945198 | 12822371 | 14047058 | 12934233 | **0.984301848** | 7 |
| sp\|P82650 | P82650 | MRPS22 | 28S ribosomal protein S22, mitochondrial OS=Homo sap | 5518806.5 | 6304342 | 6277298 | 5131746 | 5512435 | 5476886 | 6087808 | 5781495 | **0.983920222** | 6 |
| sp\|Q9BRX | Q9BRX5 | GINS3 | DNA replication complex GINS protein PSF3 OS=Homo sa | 625547 | 485666.5 | 678835.31 | 577080.69 | 522886.25 | 642318.44 | 607928.31 | 555674.19 | **0.983810641** | 7 |
| sp\|Q9H4L | Q9H4L4 | SENP3 | Sentrin-specific protease 3 OS=Homo sapiens (Human) G | 0 | 635890 | 496773.34 | 0 | 695168.62 | 0 | 419134.81 | 0 | **0.983790497** | 8 |
| sp\|P23229 | P23229 | ITGA6 | Integrin alpha-6 OS=Homo sapiens (Human) GN=ITGA6 F | 5303032 | 4912014 | 5243460 | 6318642.5 | 6659713.5 | 5289067 | 5738952 | 3733435 | **0.983653461** | 7 |
| sp\|Q9UL1 | Q9UL15 | BAG5 | BAG family molecular chaperone regulator 5 OS=Homo s | 376913.31 | 259111.97 | 658495.5 | 730303.75 | 563423.75 | 537317.19 | 261581.31 | 629354.56 | **0.983629337** | 7 |
| sp\|Q1350 | Q13509 | TUBB3 | Tubulin beta-3 chain OS=Homo sapiens (Human) GN=TU | 665815232 | 837245120 | 829315584 | 794057984 | 823044928 | 736391040 | 765955008 | 749418560 | **0.983487774** | 5 |
| sp\|P04406 | P04406 | GAPDH | Glyceraldehyde-3-phosphate dehydrogenase OS=Homo | 1620308608 | 1712007808 | 1783149824 | 1607295616 | 1808198528 | 1999189376 | 1789072000 | 1015082048 | **0.983456218** | 4 |
| sp\|O0048 | O00487 | PSMD14 | 26S proteasome non-ATPase regulatory subunit 14 OS=H | 13814659 | 14962385 | 14531664 | 17464670 | 13795290 | 15184666 | 14864244 | 15909055 | **0.983214311** | 8 |
| sp\|Q9NRZ | Q9NRZ9 | HELLS | Lymphoid-specific helicase OS=Homo sapiens (Human) ( | 1058260.88 | 739468.94 | 1783367.5 | 1276094 | 1055499.75 | 669759.81 | 1944438.5 | 1105072.62 | **0.983031214** | 7 |
| sp\|P07306 | P07306 | ASGR1 | Asialoglycoprotein receptor 1 OS=Homo sapiens (Humar | 39274040 | 50588984 | 43076224 | 40540024 | 52830320 | 40607788 | 45561152 | 31522490 | **0.982951727** | 7 |
| sp\|Q0786 | Q07866 | KLC1 | Kinesin light chain 1 OS=Homo sapiens (Human) GN=KLC | 6678794 | 6925952.5 | 7348505 | 7588140 | 7185687 | 7848228 | 7019641 | 5997896.5 | **0.982834089** | 8 |
| sp\|Q9Y6D | Q9Y6D0 | SELENOK | Selenoprotein K OS=Homo sapiens (Human) GN=SELENC | 991749.75 | 824054.81 | 0 | 715397.12 | 991458.88 | 604748.06 | 890703.62 | 0 | **0.98250194** | 7.5 |
| sp\|Q8N3U | Q8N3U4 | STAG2 | Cohesin subunit SA-2 OS=Homo sapiens (Human) GN=ST | 1702853.38 | 1903174.5 | 1816934.88 | 1836647.5 | 1658497 | 1642329.62 | 1980015.5 | 1849753.12 | **0.982228382** | 7 |
| sp\|P04183 | P04183 | TK1 | Thymidine kinase, cytosolic OS=Homo sapiens (Human) ( | 3018618.5 | 2543637 | 3865274.25 | 2870707 | 3436887.5 | 3170569.25 | 2749504.75 | 2717923 | **0.981838677** | 8 |
| sp\|Q9H2M | Q9H2M9 | RAB3GAP2 | Rab3 GTPase-activating protein non-catalytic subunit OS | 4202468 | 4558467 | 5221414 | 3432139.5 | 4614203.5 | 4076976.5 | 3593930.75 | 4808799.5 | **0.981591291** | 8 |
| sp\|Q9HC5 | Q9HC52 | CBX8 | Chromobox protein homolog 8 OS=Homo sapiens (Huma | 767689.44 | 669340.38 | 860504.06 | 764950.62 | 610562.25 | 636971.19 | 745041.06 | 1012695.44 | **0.9813176** | 5 |

| | | | | | | | | | | | | | |
|---|---|---|---|---|---|---|---|---|---|---|---|---|---|
| sp\|Q9BXJ8 | Q9BXJ8 | TMEM120A | Transmembrane protein 120A OS=Homo sapiens (Huma | 1473833.75 | 1618307 | 1428290.75 | 1532878.25 | 1228385.38 | 1278482.25 | 2066912.38 | 1366266.88 | **0.981289102** | 4 |
| sp\|P61026 | P61026 | RAB10 | Ras-related protein Rab-10 OS=Homo sapiens (Human) ( | 9021466 | 26432710 | 22417370 | 27724280 | 20115734 | 25006550 | 18768626 | 20076696 | **0.980977811** | 5 |
| sp\|Q9NV3 | Q9NV31 | IMP3 | U3 small nucleolar ribonucleoprotein protein IMP3 OS=H | 3315387 | 3182658.5 | 3180941.25 | 3483324.5 | 3285911.25 | 2657981.75 | 3184206.5 | 3779864.5 | **0.980676095** | 8 |
| sp\|Q1331( | Q13310 | PABPC4 | Polyadenylate-binding protein 4 OS=Homo sapiens (Hum | 77199520 | 79027720 | 85738200 | 82761416 | 81002360 | 79291112 | 82899440 | 75253896 | **0.980660522** | 7 |
| sp\|A6NDG | A6NDG6 | PGP | Glycerol-3-phosphate phosphatase OS=Homo sapiens (H | 4364955.5 | 3833966 | 3711041 | 3929181 | 4470090 | 3849833.75 | 3606836.25 | 3603450.5 | **0.980495599** | 6 |
| sp\|Q1563 | Q15637 | SF1 | Splicing factor 1 OS=Homo sapiens (Human) GN=SF1 PE= | 13295868 | 11746054 | 19648368 | 13002663 | 15224693 | 11828741 | 16740641 | 12763136 | **0.980314026** | 8 |
| sp\|Q9NY6 | Q9NY65 | TUBA8 | Tubulin alpha-8 chain OS=Homo sapiens (Human) GN=TU | 509182560 | 530895776 | 726062464 | 686488320 | 591142656 | 510652640 | 622312320 | 679564096 | **0.980038805** | 7 |
| sp\|Q8NI27 | Q8NI27 | THOC2 | THO complex subunit 2 OS=Homo sapiens (Human) GN= | 7487064 | 7537483.5 | 8480075 | 7698578.5 | 6770710 | 7232277 | 8150028.5 | 8412820 | **0.979573714** | 6 |
| sp\|Q7RTS | Q7RTS9 | DYM | Dymeclin OS=Homo sapiens (Human) GN=DYM PE=1 SV= | 269273.97 | 351981.34 | 418630.97 | 299559.34 | 276971.84 | 377933.19 | 312517.09 | 344285.91 | **0.979291739** | 8 |
| sp\|Q5R3I4 | Q5R3I4 | TTC38 | Tetratricopeptide repeat protein 38 OS=Homo sapiens (H | 17626886 | 17113858 | 15933853 | 16502287 | 18224816 | 15685414 | 18551500 | 13314738 | **0.979153305** | 8 |
| sp\|Q9HCE | Q9HCE1 | MOV10 | Putative helicase MOV-10 OS=Homo sapiens (Human) GI | 12255934 | 12859745 | 13116023 | 12267703 | 13435336 | 12617008 | 9948294 | 13434774 | **0.978930583** | 6 |
| sp\|O1460 | O14602 | EIF1AY | Eukaryotic translation initiation factor 1A, Y-chromosoma | 11261614 | 8531517 | 10460100 | 10547198 | 10286687 | 8574050 | 11081576 | 9991021 | **0.978747895** | 6 |
| sp\|Q9BUL | Q9BUL8 | PDCD10 | Programmed cell death protein 10 OS=Homo sapiens (Hu | 2473547.5 | 1910105.62 | 2034317.38 | 2804040.25 | 2264553.25 | 3074283 | 2807081.5 | 877779.06 | **0.978495586** | 6 |
| sp\|Q9UNF | Q9UNF0 | PACSIN2 | Protein kinase C and casein kinase substrate in neurons p | 8575768 | 9485361 | 8798730 | 9397754 | 9653077 | 8565730 | 8959756 | 8296632 | **0.978420587** | 6 |
| sp\|Q9Y28 | Q9Y285 | FARSA | Phenylalanine--tRNA ligase alpha subunit OS=Homo sapi | 5752777 | 7806953 | 5213804.5 | 5731041.5 | 5014424.5 | 5322292.5 | 7535932 | 6094654 | **0.978074585** | 7 |
| sp\|O1506 | O15067 | PFAS | Phosphoribosylformylglycinamidine synthase OS=Homo | 25645200 | 17710024 | 22223052 | 19652972 | 21909694 | 24723828 | 21871560 | 14856782 | **0.978066917** | 7 |
| sp\|P04075 | P04075 | ALDOA | Fructose-bisphosphate aldolase A OS=Homo sapiens (Hu | 984794432 | 936387200 | 1125132032 | 931869760 | 1077155456 | 1102910336 | 893618944 | 817209536 | **0.978058037** | 6 |
| sp\|P52294 | P52294 | KPNA1 | Importin subunit alpha-5 OS=Homo sapiens (Human) GN | 5293387 | 5941960 | 5336653 | 5063252 | 4898496 | 5308140.5 | 5394027 | 5557771 | **0.977961084** | 8 |
| sp\|O1481 | O14818 | PSMA7 | Proteasome subunit alpha type-7 OS=Homo sapiens (Hu | 88732240 | 79598344 | 82098000 | 88608472 | 83128608 | 83721936 | 82898776 | 81752776 | **0.977775409** | 7 |
| sp\|O43491 | O43491 | EPB41L2 | Band 4.1-like protein 2 OS=Homo sapiens (Human) GN=E | 38768180 | 34104352 | 34612528 | 36875836 | 35277324 | 37526688 | 36961040 | 31383828 | **0.977750097** | 8 |
| sp\|Q96EY | Q96EY7 | PTCD3 | Pentatricopeptide repeat domain-containing protein 3, n | 12118410 | 10628318 | 12364659 | 12938458 | 12039336 | 12732941 | 10121011 | 12081109 | **0.977618076** | 5 |
| sp\|O4368 | O43684 | BUB3 | Mitotic checkpoint protein BUB3 OS=Homo sapiens (Hur | 22723064 | 24976166 | 29629786 | 25764688 | 28506636 | 23656026 | 22198878 | 26343336 | **0.976828575** | 7 |
| sp\|O1503 | O15031 | PLXNB2 | Plexin-B2 OS=Homo sapiens (Human) GN=PLXNB2 PE=1 | 1500870 | 2141296.25 | 2085880.38 | 2270883.5 | 1874127.38 | 2180046.5 | 1573546.75 | 2185272.75 | **0.976754798** | 8 |
| sp\|Q96KR | Q96KR6 | FAM210B | Protein FAM210B, mitochondrial OS=Homo sapiens (Hur | 1989243 | 2708763.5 | 2290431.75 | 2291113.25 | 1831506.62 | 2286555.75 | 1933942.5 | 3011433.25 | **0.976710795** | 5 |
| sp\|P22626 | P22626 | HNRNPA2E | Heterogeneous nuclear ribonucleoproteins A2/B1 OS=H | 214800368 | 220577808 | 197965712 | 232007840 | 249550880 | 190236048 | 217888672 | 187452800 | **0.976629933** | 6 |
| sp\|O0046 | O00469 | PLOD2 | Procollagen-lysine,2-oxoglutarate 5-dioxygenase 2 OS=H | 29095144 | 24352032 | 23667220 | 24247284 | 22976272 | 31456060 | 23862484 | 20694960 | **0.976599599** | 5 |
| sp\|Q9NRV | Q9NRV9 | HEBP1 | Heme-binding protein 1 OS=Homo sapiens (Human) GN= | 749746.12 | 687818.69 | 485811.5 | 810767.69 | 573738.38 | 775580.75 | 754684.06 | 565558.44 | **0.976379309** | 8 |
| sp\|O9601 | O96019 | ACTL6A | Actin-like protein 6A OS=Homo sapiens (Human) GN=AC | 6314214 | 7379138 | 7792427 | 6159674 | 6887289 | 7604688 | 6606762 | 5890130 | **0.976249838** | 7 |
| sp\|Q9284 | Q92841 | DDX17 | Probable ATP-dependent RNA helicase DDX17 OS=Homo | 192428368 | 198407760 | 204171264 | 198399104 | 203083696 | 167801648 | 205662064 | 197978480 | **0.976203109** | 8 |
| sp\|P29401 | P29401 | TKT | Transketolase OS=Homo sapiens (Human) GN=TKT PE=1 | 337289024 | 344092992 | 406202784 | 341105152 | 368861760 | 320611872 | 363666880 | 340979328 | **0.975802929** | 7 |
| sp\|P53621 | P53621 | COPA | Coatomer subunit alpha OS=Homo sapiens (Human) GN= | 112600504 | 123651712 | 120588520 | 121317208 | 110654336 | 112570768 | 115310264 | 128027792 | **0.975751142** | 5 |
| sp\|Q6P6C | Q6P6C2 | ALKBH5 | RNA demethylase ALKBH5 OS=Homo sapiens (Human) G | 3337865.75 | 1941845.62 | 2779566 | 2287375.25 | 2318016.5 | 2927306.75 | 1759625.75 | 3089305 | **0.97560577** | 8 |
| sp\|O6068 | O60684 | KPNA6 | Importin subunit alpha-7 OS=Homo sapiens (Human) GN | 6844869 | 7471944 | 7186015 | 6495091 | 6277938 | 7137303.5 | 6857618 | 7041260 | **0.975576774** | 6 |
| sp\|Q9UDY | Q9UDY4 | DNAJB4 | DnaJ homolog subfamily B member 4 OS=Homo sapiens | 358693.78 | 475278.16 | 336261.91 | 461491.25 | 361098.69 | 513987.94 | 360941.5 | 355707.31 | **0.975492404** | 7 |
| sp\|P51570 | P51570 | GALK1 | Galactokinase OS=Homo sapiens (Human) GN=GALK1 PE | 22260314 | 21739576 | 27641666 | 21130128 | 22244280 | 19922300 | 24102326 | 24200926 | **0.975187989** | 8 |
| sp\|P07814 | P07814 | EPRS | Bifunctional glutamate/proline--tRNA ligase OS=Homo s | 70445208 | 75036568 | 67451984 | 65186824 | 68567800 | 64516204 | 70131232 | 67764904 | **0.974326086** | 6 |
| sp\|O9535 | O95352 | ATG7 | Ubiquitin-like modifier-activating enzyme ATG7 OS=Hom | 244346.22 | 267881.22 | 198234.59 | 251989.38 | 244928.94 | 262635.81 | 204866.55 | 224545.92 | **0.973531973** | 7 |
| sp\|Q9P26 | Q9P260 | RELCH | RAB11-binding protein RELCH OS=Homo sapiens (Humar | 650432.12 | 674908.38 | 1004454.19 | 740280 | 421831.31 | 729705.31 | 881425.81 | 953868.06 | **0.972885285** | 8 |
| sp\|Q7Z6Z | Q7Z6Z7 | HUWE1 | E3 ubiquitin-protein ligase HUWE1 OS=Homo sapiens (H | 15385012 | 14561401 | 12605661 | 13511364 | 12888098 | 14034134 | 13796902 | 13817591 | **0.972768117** | 7 |
| sp\|Q9297 | Q92979 | EMG1 | Ribosomal RNA small subunit methyltransferase NEP1 O | 13526139 | 13461182 | 15296075 | 14190844 | 13667806 | 13152981 | 13650446 | 14415880 | **0.971896443** | 7 |
| sp\|Q8WU | Q8WUM0 | NUP133 | Nuclear pore complex protein Nup133 OS=Homo sapien | 9303519 | 9852342 | 9976742 | 10500112 | 9299521 | 9676099 | 10274320 | 9266546 | **0.971835667** | 4 |
| sp\|Q5JTZ9 | Q5JTZ9 | AARS2 | Alanine--tRNA ligase, mitochondrial OS=Homo sapiens (I | 5821232 | 4950957.5 | 5035904 | 4163798.5 | 5505770 | 4604911 | 4042764.25 | 5241855 | **0.971129838** | 7 |
| sp\|Q0151 | Q01518 | CAP1 | Adenylyl cyclase-associated protein 1 OS=Homo sapiens | 76544864 | 86039008 | 84099688 | 84549888 | 80541216 | 78570200 | 82148944 | 80371296 | **0.97101201** | 4 |
| sp\|Q1540 | Q15404 | RSU1 | Ras suppressor protein 1 OS=Homo sapiens (Human) GN | 2252965.75 | 1859227.5 | 1107638.88 | 1986989.75 | 1758856.88 | 2186583 | 1585865.62 | 1465629.5 | **0.970876638** | 6 |
| sp\|Q9H8S | Q9H8S9 | MOB1A | MOB kinase activator 1A OS=Homo sapiens (Human) GN | 3244982.5 | 2906572.5 | 3318076.5 | 3425353 | 3261224 | 3136161.5 | 2927120 | 3194846.25 | **0.970869856** | 5 |

| | | | | | | | | | | | | | |
|---|---|---|---|---|---|---|---|---|---|---|---|---|---|
| sp\|Q9Y33 | Q9Y333 | LSM2 | U6 snRNA-associated Sm-like protein LSm2 OS=Homo sa | 4381719.5 | 3959266 | 3802511.25 | 4519644.5 | 3621773.25 | 4357011.5 | 4211476.5 | 3987465.5 | **0.970868968** | 6 |
| sp\|Q9BWF | Q9BWF3 | RBM4 | RNA-binding protein 4 OS=Homo sapiens (Human) GN=R | 4395440.5 | 4046402 | 5045451 | 4624596 | 4595349 | 3762999.5 | 4870829 | 4348001 | **0.970477349** | 6 |
| sp\|P49721 | P49721 | PSMB2 | Proteasome subunit beta type-2 OS=Homo sapiens (Hun | 35394264 | 34887420 | 34481696 | 33618160 | 37391660 | 35153516 | 31435560 | 30306580 | **0.97041351** | 7 |
| sp\|P62263 | P62263 | RPS14 | 40S ribosomal protein S14 OS=Homo sapiens (Human) G | 95454248 | 104035560 | 125917704 | 110796016 | 119308432 | 91995696 | 121763128 | 90087264 | **0.970085047** | 6 |
| sp\|P46783 | P46783 | RPS10 | 40S ribosomal protein S10 OS=Homo sapiens (Human) G | 59274788 | 52154356 | 55371208 | 56738688 | 58743984 | 46092832 | 54798908 | 57153668 | **0.969805507** | 7 |
| sp\|P11177 | P11177 | PDHB | Pyruvate dehydrogenase E1 component subunit beta, mi | 40336264 | 39153284 | 35318900 | 40415148 | 40697524 | 33670348 | 40496796 | 35659384 | **0.969724036** | 7 |
| sp\|Q1467 | Q14671 | PUM1 | Pumilio homolog 1 OS=Homo sapiens (Human) GN=PUN | 1433676.5 | 1402313.62 | 1415875 | 1311236.25 | 1295037.5 | 1165646.5 | 1307048.75 | 1625876.75 | **0.969532845** | 4 |
| sp\|P23634 | P23634 | ATP2B4 | Plasma membrane calcium-transporting ATPase 4 OS=H | 4509130 | 5532156.5 | 4483818 | 6286388 | 4911544 | 5752954 | 4750519.5 | 4757340 | **0.969289324** | 7 |
| sp\|Q7Z3U | Q7Z3U7 | MON2 | Protein MON2 homolog OS=Homo sapiens (Human) GN= | 1355380.75 | 1412215.38 | 1525895.5 | 1380131.5 | 1264331 | 1459139 | 1417363.5 | 1357288.25 | **0.969067142** | 7 |
| sp\|Q9UGP | Q9UGP8 | SEC63 | Translocation protein SEC63 homolog OS=Homo sapiens | 7972505 | 8438025 | 8334888.5 | 7968923.5 | 8795783 | 6943712 | 7259433 | 8698165 | **0.968905106** | 8 |
| sp\|P31948 | P31948 | STIP1 | Stress-induced-phosphoprotein 1 OS=Homo sapiens (Hu | 104847864 | 96839984 | 106931936 | 111110528 | 100711984 | 90202968 | 109944160 | 105724224 | **0.968677564** | 6 |
| sp\|Q9Y61 | Q9Y617 | PSAT1 | Phosphoserine aminotransferase OS=Homo sapiens (Hu | 180777072 | 214020400 | 208282624 | 182109104 | 205082864 | 169350928 | 191855776 | 193972000 | **0.968252706** | 6 |
| sp\|P63173 | P63173 | RPL38 | 60S ribosomal protein L38 OS=Homo sapiens (Human) G | 36290444 | 36808532 | 35049656 | 35519248 | 35231084 | 33215014 | 34300656 | 36346824 | **0.96816058** | 4 |
| sp\|Q1506 | Q15061 | WDR43 | WD repeat-containing protein 43 OS=Homo sapiens (Hu | 8855386 | 8534306 | 10837447 | 9952576 | 8323667.5 | 8225421 | 9672206 | 10736303 | **0.967990398** | 5 |
| sp\|P40227 | P40227 | CCT6A | T-complex protein 1 subunit zeta OS=Homo sapiens (Hu | 152197856 | 153100352 | 164807600 | 167985776 | 162493472 | 155723856 | 149185280 | 149714192 | **0.967128882** | 4 |
| sp\|Q0916 | Q09161 | NCBP1 | Nuclear cap-binding protein subunit 1 OS=Homo sapiens | 5698962 | 6107009.5 | 5420632 | 5469159 | 4789139.5 | 4753368.5 | 5405672 | 7000766.5 | **0.967094474** | 4 |
| sp\|Q9UBF | Q9UBF2 | COPG2 | Coatomer subunit gamma-2 OS=Homo sapiens (Human) | 19082678 | 17930608 | 18289088 | 18456618 | 16362676 | 15133759 | 18031428 | 21776748 | **0.966724315** | 5 |
| sp\|Q9Y3B | Q9Y3B8 | REXO2 | Oligoribonuclease, mitochondrial OS=Homo sapiens (Hu | 2793058.5 | 3107236.25 | 3773570 | 3100526.5 | 3444262.75 | 2732138.75 | 3200071.25 | 2972774.75 | **0.96671906** | 7 |
| sp\|Q9Y2W | Q9Y2W1 | THRAP3 | Thyroid hormone receptor-associated protein 3 OS=Hon | 23974008 | 22572600 | 23070710 | 20077096 | 21199804 | 18480738 | 25918340 | 21072996 | **0.966301848** | 6 |
| sp\|O7568 | O75688 | PPM1B | Protein phosphatase 1B OS=Homo sapiens (Human) GN= | 963552.81 | 1038266.31 | 841217.38 | 1216460.75 | 1241562.88 | 1005691.25 | 779642.44 | 895738 | **0.966285805** | 7 |
| sp\|Q9H2G | Q9H2G2 | SLK | STE20-like serine/threonine-protein kinase OS=Homo sa | 661041.94 | 614086.62 | 894944.38 | 710379.56 | 811500.06 | 973869.44 | 0 | 997486.25 | **0.966117563** | 5 |
| sp\|O7569 | O75695 | RP2 | Protein XRP2 OS=Homo sapiens (Human) GN=RP2 PE=1 S | 778646.38 | 828446.81 | 659580.25 | 845896.38 | 762091.69 | 654319.19 | 825311.06 | 764891.5 | **0.965958553** | 4 |
| sp\|Q9BUF | Q9BUF5 | TUBB6 | Tubulin beta-6 chain OS=Homo sapiens (Human) GN=TU | 382568096 | 545189312 | 518286880 | 475268576 | 483734208 | 452346592 | 484077312 | 435292544 | **0.965720207** | 6 |
| sp\|Q96G2 | Q96G23 | CERS2 | Ceramide synthase 2 OS=Homo sapiens (Human) GN=CE | 9979092 | 8489623 | 8737537 | 9741220 | 10004627 | 9212913 | 8158361.5 | 8304103.5 | **0.965695434** | 6 |
| sp\|Q9NQR | Q9NQR4 | NIT2 | Omega-amidase NIT2 OS=Homo sapiens (Human) GN=N | 6586783 | 6001434 | 6312672 | 5983661 | 4437761 | 6384310 | 6500446 | 6702267.5 | **0.965449827** | 6 |
| sp\|Q9UKN | Q9UKN8 | GTF3C4 | General transcription factor 3C polypeptide 4 OS=Homo | 628832.56 | 727387.94 | 744690.38 | 866777 | 642843.25 | 653652.12 | 799121.62 | 769428 | **0.965413179** | 8 |
| sp\|Q96KR | Q96KR1 | ZFR | Zinc finger RNA-binding protein OS=Homo sapiens (Hum | 2432766.5 | 3021284 | 3361697.25 | 2507854 | 2610306.75 | 2119243.5 | 3532664 | 2668935 | **0.965342079** | 8 |
| sp\|Q9289 | Q92896 | GLG1 | Golgi apparatus protein 1 OS=Homo sapiens (Human) GI | 19090416 | 21649552 | 23583392 | 19231252 | 18268590 | 20030524 | 21113112 | 21214604 | **0.96495966** | 6 |
| sp\|Q5T8P | Q5T8P6 | RBM26 | RNA-binding protein 26 OS=Homo sapiens (Human) GN= | 2621799.25 | 2816416 | 3114380.5 | 2734749.25 | 2846293 | 2641976.25 | 2445619.25 | 2957312.75 | **0.964903726** | 7 |
| sp\|O1492 | O14929 | HAT1 | Histone acetyltransferase type B catalytic subunit OS=H | 10960664 | 10852904 | 11105196 | 11300272 | 10985236 | 12433788 | 10454912 | 8771091 | **0.964404267** | 6 |
| sp\|Q0817 | Q08170 | SRSF4 | Serine/arginine-rich splicing factor 4 OS=Homo sapiens ( | 33052148 | 40018848 | 42798360 | 44628232 | 28411894 | 39535272 | 46728424 | 40099072 | **0.964342604** | 7 |
| sp\|O1540 | O15400 | STX7 | Syntaxin-7 OS=Homo sapiens (Human) GN=STX7 PE=1 SV | 5481624.5 | 5123007 | 4573777 | 5132280 | 5139369 | 5162969 | 5028052 | 4247627.5 | **0.963926826** | 7 |
| sp\|O1466 | O14662 | STX16 | Syntaxin-16 OS=Homo sapiens (Human) GN=STX16 PE=1 | 565351.25 | 1009062 | 1527578.25 | 1308600 | 823211.5 | 864466.5 | 1880923.12 | 682162 | **0.963762597** | 7 |
| sp\|Q5JS54 | Q5JS54 | PSMG4 | Proteasome assembly chaperone 4 OS=Homo sapiens (H | 1044188.62 | 772505.81 | 1242884 | 1074644.88 | 1050620.12 | 823110.88 | 967253.44 | 1142052.25 | **0.963430466** | 7 |
| sp\|Q1502 | Q15024 | EXOSC7 | Exosome complex component RRP42 OS=Homo sapiens | 2975274.75 | 2601361.5 | 2402845 | 2948312.5 | 2491858.5 | 2877893.75 | 2684880.25 | 2472746.75 | **0.963358157** | 6 |
| sp\|Q9HCU | Q9HCU5 | PREB | Prolactin regulatory element-binding protein OS=Homo | 4491608 | 5036151.5 | 3675512.5 | 4397617 | 4435248 | 4161853.25 | 4819581 | 3539192 | **0.963353286** | 6 |
| sp\|Q52LJ0 | Q52LJ0 | FAM98B | Protein FAM98B OS=Homo sapiens (Human) GN=FAM98 | 359820.47 | 301165.41 | 111358.91 | 248031.3 | 186942.34 | 312243.84 | 304195.34 | 179526.34 | **0.96327998** | 8 |
| sp\|Q1512 | Q15126 | PMVK | Phosphomevalonate kinase OS=Homo sapiens (Human) | 750693.75 | 1202328.12 | 1295238.12 | 1459746 | 1144903.75 | 696420.5 | 1391655.12 | 1301454.5 | **0.963132562** | 7 |
| sp\|Q9H6R | Q9H6R4 | NOL6 | Nucleolar protein 6 OS=Homo sapiens (Human) GN=NOL | 3912705.75 | 3024365.5 | 3834092 | 3139188.5 | 3123188.5 | 3087258.5 | 4061868.25 | 3122294.25 | **0.962923853** | 7 |
| sp\|P30101 | P30101 | PDIA3 | Protein disulfide-isomerase A3 OS=Homo sapiens (Huma | 582674880 | 591835712 | 598675904 | 575964288 | 582085760 | 558586560 | 579023616 | 542312896 | **0.962904913** | 2 |
| sp\|P31930 | P31930 | UQCRC1 | Cytochrome b-c1 complex subunit 1, mitochondrial OS=I | 63729688 | 69763840 | 84169944 | 74810984 | 79131968 | 67125952 | 70776344 | 64585260 | **0.96288588** | 7 |
| sp\|Q9Y4E | Q9Y4E8 | USP15 | Ubiquitin carboxyl-terminal hydrolase 15 OS=Homo sapi | 1404019.5 | 1288822.75 | 1407834.12 | 1423757.5 | 1126442.75 | 964749 | 1244493.12 | 1982210.75 | **0.96261368** | 4 |
| sp\|Q8IX12 | Q8IX12 | CCAR1 | Cell division cycle and apoptosis regulator protein 1 OS= | 9295637 | 9933007 | 9952369 | 7624613 | 12063542 | 7152331.5 | 4428996 | 11777734 | **0.962423612** | 8 |
| sp\|Q9NRF | Q9NRF8 | CTPS2 | CTP synthase 2 OS=Homo sapiens (Human) GN=CTPS2 PI | 6371110 | 6213003.5 | 6060600.5 | 5726838 | 5813540 | 7483768 | 4888024.5 | 5268911 | **0.962361507** | 5 |
| sp\|Q8N2K | Q8N2K0 | ABHD12 | Monoacylglycerol lipase ABHD12 OS=Homo sapiens (Hui | 4140467 | 4092362.25 | 4960858.5 | 4485873 | 4799004 | 4367244 | 3856485.75 | 3990723.5 | **0.962323527** | 5 |

| | | | | | | | | | | | | |
|---|---|---|---|---|---|---|---|---|---|---|---|---|
| sp\|Q1472 | Q14728 | MFSD10 | Major facilitator superfamily domain-containing protein | 1969894 | 2768312 | 3511053 | 2171886 | 2715532.75 | 2209311.5 | 2526313 | 2569072.25 | **0.961528652** | 8 |
| sp\|P14868 | P14868 | DARS | Aspartate--tRNA ligase, cytoplasmic OS=Homo sapiens ( | 115980152 | 107167472 | 109268600 | 110329768 | 97123808 | 110298848 | 102913832 | 115288864 | **0.961330785** | 5 |
| sp\|P48556 | P48556 | PSMD8 | 26S proteasome non-ATPase regulatory subunit 8 OS=Ho | 22023140 | 22280668 | 24155216 | 27284178 | 21153990 | 24793040 | 22342388 | 23735284 | **0.961161733** | 7 |
| sp\|Q1554 | Q15542 | TAF5 | Transcription initiation factor TFIID subunit 5 OS=Homo | 316746.72 | 285414.16 | 295243.25 | 310816.19 | 302721.81 | 306481.59 | 266187.47 | 285765.47 | **0.961046856** | 5 |
| sp\|P52594 | P52594 | AGFG1 | Arf-GAP domain and FG repeat-containing protein 1 OS= | 5369304.5 | 4151320.75 | 3724366.75 | 5367309 | 4912824 | 4371258 | 4757141 | 3836629.25 | **0.960539605** | 7 |
| sp\|P35610 | P35610 | SOAT1 | Sterol O-acyltransferase 1 OS=Homo sapiens (Human) G | 2019593.5 | 2434630 | 2183964.5 | 2166179.25 | 2408699 | 1616262.12 | 2371088.25 | 2060522.88 | **0.960497445** | 7 |
| sp\|Q9952 | Q99523 | SORT1 | Sortilin OS=Homo sapiens (Human) GN=SORT1 PE=1 SV= | 5604814 | 6504765 | 6955841 | 7093879 | 6166643 | 6645438.5 | 5873284 | 6432699 | **0.960196391** | 5 |
| sp\|Q9Y27 | Q9Y276 | BCS1L | Mitochondrial chaperone BCS1 OS=Homo sapiens (Hum | 770519.5 | 1121578.5 | 793455.88 | 1042417.38 | 1072033.5 | 1030022.12 | 365660.47 | 1111551.75 | **0.960111436** | 8 |
| sp\|P48739 | P48739 | PITPNB | Phosphatidylinositol transfer protein beta isoform OS=H | 12917382 | 13054009 | 12755664 | 12103201 | 13122340 | 12026757 | 12186194 | 11457101 | **0.959908445** | 5 |
| sp\|Q1324 | Q13243 | SRSF5 | Serine/arginine-rich splicing factor 5 OS=Homo sapiens ( | 32977502 | 29987218 | 33051996 | 30710578 | 29191494 | 28747224 | 33633484 | 30060696 | **0.959800325** | 5 |
| sp\|Q96RP | Q96RP9 | GFM1 | Elongation factor G, mitochondrial OS=Homo sapiens (H | 8959631 | 10654930 | 10346147 | 10359156 | 10507132 | 9607872 | 9301823 | 9275252 | **0.959628212** | 6 |
| sp\|Q9H8Y | Q9H8Y8 | GORASP2 | Golgi reassembly-stacking protein 2 OS=Homo sapiens ( | 20441144 | 22086332 | 22660408 | 17235632 | 17361438 | 20642840 | 20801972 | 20186606 | **0.958377655** | 6 |
| sp\|Q9287 | Q92879 | CELF1 | CUGBP Elav-like family member 1 OS=Homo sapiens (Hu | 2553496 | 4044022.5 | 2866537.5 | 3440123.5 | 2064125.38 | 2391683.25 | 3259239.5 | 4648918 | **0.958136558** | 6 |
| sp\|O7571 | O75717 | WDHD1 | WD repeat and HMG-box DNA-binding protein 1 OS=Ho | 0 | 864321.25 | 2463135 | 2234808 | 1898965 | 836581.75 | 1595463.5 | 997256.62 | **0.957931272** | 7 |
| sp\|Q68E0 | Q68E01 | INTS3 | Integrator complex subunit 3 OS=Homo sapiens (Human | 1261114.88 | 1194429.38 | 574292.62 | 1154056.25 | 935051.44 | 1066582.75 | 1072233 | 932665.88 | **0.957608846** | 4 |
| sp\|Q8NEJ | Q8NEJ9 | NGDN | Neuroguidin OS=Homo sapiens (Human) GN=NGDN PE= | 2932949 | 2236067 | 2259133.75 | 2686267.25 | 2775381 | 2248143 | 2396337.5 | 2265150.5 | **0.957545254** | 8 |
| sp\|P20618 | P20618 | PSMB1 | Proteasome subunit beta type-1 OS=Homo sapiens (Hun | 40679288 | 42796716 | 44868804 | 42286752 | 36532344 | 42460524 | 41492828 | 42898392 | **0.957525607** | 6 |
| sp\|Q1500 | Q15006 | EMC2 | ER membrane protein complex subunit 2 OS=Homo sapi | 1433341.12 | 1292817.75 | 1964469.62 | 1154754.12 | 1861742.5 | 1244687.38 | 1445782.5 | 1044171.25 | **0.95740245** | 7 |
| sp\|P61758 | P61758 | VBP1 | Prefoldin subunit 3 OS=Homo sapiens (Human) GN=VBP | 7931096 | 8725420 | 6113731 | 5139900.5 | 8495362 | 4912044 | 8557206 | 4752248.5 | **0.957245407** | 6 |
| sp\|Q96BM | Q96BM9 | ARL8A | ADP-ribosylation factor-like protein 8A OS=Homo sapien | 1218903.38 | 732865.88 | 1062241.25 | 2228053.25 | 1030750 | 2203025.5 | 1038856.5 | 743573.88 | **0.95691432** | 6 |
| sp\|P25789 | P25789 | PSMA4 | Proteasome subunit alpha type-4 OS=Homo sapiens (Hu | 29802510 | 28521086 | 33813828 | 31268668 | 35124508 | 28272238 | 24648510 | 30037094 | **0.956859974** | 6 |
| sp\|P62888 | P62888 | RPL30 | 60S ribosomal protein L30 OS=Homo sapiens (Human) G | 54254208 | 50621324 | 67600432 | 55561756 | 63563004 | 47642248 | 55238532 | 51645616 | **0.956374235** | 6 |
| sp\|P60510 | P60510 | PPP4C | Serine/threonine-protein phosphatase 4 catalytic subuni | 2496298.5 | 4684642.5 | 4722143.5 | 1499410.75 | 2575307.75 | 3918338.5 | 4263733 | 2059823.62 | **0.956329596** | 7 |
| sp\|Q9Y39 | Q9Y399 | MRPS2 | 28S ribosomal protein S2, mitochondrial OS=Homo sapie | 9394981 | 9141851 | 12418996 | 10714292 | 9757939 | 7890800.5 | 9808924 | 12387281 | **0.956199418** | 7 |
| sp\|Q96AG | Q96AG4 | LRRC59 | Leucine-rich repeat-containing protein 59 OS=Homo sap | 119425760 | 134000040 | 111284832 | 118951768 | 123498312 | 117508784 | 114620360 | 106813384 | **0.956123197** | 5 |
| sp\|Q9UP8 | Q9UP83 | COG5 | Conserved oligomeric Golgi complex subunit 5 OS=Hom | 894888.81 | 774849 | 723736.62 | 924361.12 | 457181.28 | 821741.94 | 974938.88 | 918194.5 | **0.956062033** | 7 |
| sp\|O9497 | O94973 | AP2A2 | AP-2 complex subunit alpha-2 OS=Homo sapiens (Huma | 14818094 | 16379768 | 16726762 | 16439185 | 13261401 | 14720266 | 15072108 | 18480312 | **0.956035511** | 5 |
| sp\|Q9NTK | Q9NTK5 | OLA1 | Obg-like ATPase 1 OS=Homo sapiens (Human) GN=OLA1 | 63425328 | 57440228 | 62213508 | 60276340 | 60307180 | 56647696 | 56872284 | 58796636 | **0.955901501** | 3 |
| sp\|O6027 | O60271 | SPAG9 | C-Jun-amino-terminal kinase-interacting protein 4 OS=H | 3509017.5 | 2041224 | 3472591.25 | 2947672.5 | 2916936.25 | 2323912 | 2792023.5 | 3408905 | **0.955830728** | 5 |
| sp\|P16435 | P16435 | POR | NADPH--cytochrome P450 reductase OS=Homo sapiens | 93609656 | 91492424 | 91624840 | 96787136 | 86658320 | 85033512 | 80731392 | 104495888 | **0.955570764** | 4 |
| sp\|Q96KP | Q96KP4 | CNDP2 | Cytosolic non-specific dipeptidase OS=Homo sapiens (H | 52313240 | 62493332 | 64857512 | 62974516 | 56656540 | 47500364 | 64280528 | 63416944 | **0.955554376** | 7 |
| sp\|Q9Y4Y | Q9Y4Y9 | LSM5 | U6 snRNA-associated Sm-like protein LSm5 OS=Homo sa | 2964290.5 | 2522018.5 | 2173868.5 | 2917003.5 | 2723195 | 2477129.5 | 2509234 | 2394967.5 | **0.955313708** | 5 |
| sp\|Q1543 | Q15435 | PPP1R7 | Protein phosphatase 1 regulatory subunit 7 OS=Homo sa | 20496644 | 11797860 | 8703688 | 14913977 | 11346660 | 11223154 | 9205777 | 21632656 | **0.955216869** | 7 |
| sp\|Q8TB2 | Q8TB22 | SPATA20 | Spermatogenesis-associated protein 20 OS=Homo sapie | 627787.31 | 636025.44 | 870588.81 | 665893.31 | 601494.69 | 827004.25 | 570377.31 | 675684 | **0.955099507** | 6 |
| sp\|Q9Y26 | Q9Y262 | EIF3L | Eukaryotic translation initiation factor 3 subunit L OS=H | 44593612 | 44594656 | 51521676 | 47587672 | 43763568 | 44277652 | 42716348 | 49056600 | **0.954946599** | 3 |
| sp\|Q1326 | Q13263 | TRIM28 | Transcription intermediary factor 1-beta OS=Homo sapie | 106299456 | 109501224 | 118365312 | 97452984 | 112514528 | 98008688 | 109871592 | 91625608 | **0.954592914** | 7 |
| sp\|Q96T2 | Q96T23 | RSF1 | Remodeling and spacing factor 1 OS=Homo sapiens (Hu | 420032.5 | 361730.5 | 301634.53 | 339294.75 | 322321.25 | 356501.28 | 337786.78 | 340432 | **0.953854413** | 6 |
| sp\|P12270 | P12270 | TPR | Nucleoprotein TPR OS=Homo sapiens (Human) GN=TPR | 48243980 | 48425368 | 51215840 | 42807100 | 41956324 | 44187504 | 47584672 | 48159052 | **0.953827519** | 3 |
| sp\|Q9Y67 | Q9Y679 | AUP1 | Ancient ubiquitous protein 1 OS=Homo sapiens (Human) | 12211826 | 11573808 | 14497777 | 11325828 | 14424064 | 12603249 | 11111741 | 9177710 | **0.953789354** | 6 |
| sp\|O0030 | O00303 | EIF3F | Eukaryotic translation initiation factor 3 subunit F OS=H | 31719614 | 37075720 | 35336276 | 32172218 | 31277426 | 32034020 | 33875208 | 32814226 | **0.9537581** | 5 |
| sp\|P09601 | P09601 | HMOX1 | Heme oxygenase 1 OS=Homo sapiens (Human) GN=HM | 1549128.38 | 845995.12 | 1291836.5 | 1112578.12 | 1260067.12 | 1376866 | 1031954.19 | 908529.88 | **0.953720353** | 7 |
| sp\|Q3ZCQ | Q3ZCQ8 | TIMM50 | Mitochondrial import inner membrane translocase subu | 11206068 | 12068128 | 10670039 | 13074798 | 10554101 | 10564866 | 11743706 | 11944961 | **0.952968003** | 4 |
| sp\|Q1528 | Q15287 | RNPS1 | RNA-binding protein with serine-rich domain 1 OS=Hom | 4334277 | 4909856 | 4835169 | 4588584 | 4799919 | 4579943 | 4712922 | 3695792 | **0.952897184** | 5 |
| sp\|P35609 | P35609 | ACTN2 | Alpha-actinin-2 OS=Homo sapiens (Human) GN=ACTN2 | 73577016 | 91631856 | 79950032 | 79913408 | 71149360 | 68731080 | 82723208 | 87146496 | **0.952865355** | 6 |
| sp\|P59998 | P59998 | ARPC4 | Actin-related protein 2/3 complex subunit 4 OS=Homo s | 33776884 | 33276564 | 35215248 | 34835228 | 31561156 | 30082318 | 31598108 | 37377240 | **0.95269937** | 4 |

| | | | | | | | | | | | | |
|---|---|---|---|---|---|---|---|---|---|---|---|---|
| sp\|Q9Y44( | Q9Y446 | PKP3 | Plakophilin-3 OS=Homo sapiens (Human) GN=PKP3 PE=1 | 8549606 | 8888742 | 8138330.5 | 8032439 | 7715667.5 | 7915039 | 8802993 | 7570704.5 | **0.952253626** | 3 |
| sp\|O9600{ | O96008 | TOMM40 | Mitochondrial import receptor subunit TOM40 homolog | 20106306 | 20698908 | 25197018 | 21556132 | 17733910 | 19497940 | 21636848 | 24506384 | **0.952222931** | 6 |
| sp\|Q1563: | Q15631 | TSN | Translin OS=Homo sapiens (Human) GN=TSN PE=1 SV=1 | 26107256 | 33194882 | 36827404 | 34803496 | 31427768 | 31689324 | 29690492 | 31865312 | **0.952188217** | 4 |
| sp\|P12814 | P12814 | ACTN1 | Alpha-actinin-1 OS=Homo sapiens (Human) GN=ACTN1 F | 307354112 | 346960320 | 331665024 | 339214976 | 309084736 | 304286688 | 323488096 | 324950624 | **0.95216982** | 3 |
| sp\|P48444 | P48444 | ARCN1 | Coatomer subunit delta OS=Homo sapiens (Human) GN= | 37445888 | 41723176 | 45618740 | 47357736 | 38470004 | 38616824 | 43086640 | 43697140 | **0.951930605** | 6 |
| sp\|Q86TB! | Q86TB9 | PATL1 | Protein PAT1 homolog 1 OS=Homo sapiens (Human) GN: | 1014987.38 | 1032588 | 1085811.88 | 1256509.88 | 827339.88 | 1021715.5 | 1303353.25 | 1026187.88 | **0.95186661** | 6 |
| sp\|Q9953( | Q99536 | VAT1 | Synaptic vesicle membrane protein VAT-1 homolog OS=I | 35203984 | 35207312 | 39002912 | 36980076 | 37474244 | 33319742 | 33159228 | 35380516 | **0.951770289** | 5 |
| sp\|Q9NZU | Q9NZU5 | LMCD1 | LIM and cysteine-rich domains protein 1 OS=Homo sapie | 755839.38 | 718173.81 | 738421.19 | 0 | 801149.44 | 613025 | 691342.19 | 0 | **0.95167416** | 6.5 |
| sp\|Q8TB6: | Q8TB61 | SLC35B2 | Adenosine 3'-phospho 5'-phosphosulfate transporter 1 ( | 2263237 | 1781177.88 | 1905220.62 | 1837633.62 | 1874074.62 | 1902849.12 | 1533003.5 | 2098556.75 | **0.951358413** | 7 |
| sp\|P49720 | P49720 | PSMB3 | Proteasome subunit beta type-3 OS=Homo sapiens (Hun | 34789512 | 34709572 | 40405384 | 37851784 | 35196900 | 34232716 | 32781126 | 38345688 | **0.951272302** | 5 |
| sp\|P55010 | P55010 | EIF5 | Eukaryotic translation initiation factor 5 OS=Homo sapie | 20054148 | 16242922 | 22211140 | 9634802 | 15922411 | 14291188 | 17989462 | 16614942 | **0.95120543** | 6 |
| sp\|P62829 | P62829 | RPL23 | 60S ribosomal protein L23 OS=Homo sapiens (Human) G | 68852816 | 70845728 | 73667424 | 69741368 | 71429144 | 67133920 | 67769760 | 62859308 | **0.95084831** | 3 |
| sp\|Q1469( | Q14696 | MESD | LRP chaperone MESD OS=Homo sapiens (Human) GN=M | 11458819 | 9543434 | 9911196 | 10307067 | 8446024 | 10160650 | 9684383 | 10892314 | **0.950579343** | 6 |
| sp\|O1457! | O14579 | COPE | Coatomer subunit epsilon OS=Homo sapiens (Human) GI | 9403329 | 8751087 | 8662210 | 9714138 | 8364014 | 8361985 | 8393740 | 9604570 | **0.950549761** | 3 |
| sp\|Q9BWS | Q9BWS9 | CHID1 | Chitinase domain-containing protein 1 OS=Homo sapien | 1952958 | 1979855.88 | 1493027.12 | 2000475.88 | 1413950.62 | 1831722.25 | 1824046.5 | 1988451 | **0.950426771** | 5 |
| sp\|P29083 | P29083 | GTF2E1 | General transcription factor IIE subunit 1 OS=Homo sapi | 1214791.25 | 863709.12 | 914324.75 | 830889.06 | 977465.38 | 880625.56 | 845849.56 | 928340.56 | **0.949935296** | 7 |
| sp\|Q0061( | Q00610 | CLTC | Clathrin heavy chain 1 OS=Homo sapiens (Human) GN=C | 406366336 | 418714464 | 432439488 | 393491808 | 382258752 | 388315456 | 388241664 | 409180288 | **0.949718154** | 2 |
| sp\|P49916 | P49916 | LIG3 | DNA ligase 3 OS=Homo sapiens (Human) GN=LIG3 PE=1 ! | 4026397.5 | 4388674 | 4401632 | 3643869 | 3549250.5 | 3512787.5 | 4146327.25 | 4424172 | **0.949695841** | 6 |
| sp\|P52565 | P52565 | ARHGDIA | Rho GDP-dissociation inhibitor 1 OS=Homo sapiens (Hun | 61508740 | 60051976 | 57926016 | 50385128 | 52764144 | 52591224 | 55454760 | 57483488 | **0.949631747** | 4 |
| sp\|P48643 | P48643 | CCT5 | T-complex protein 1 subunit epsilon OS=Homo sapiens ( | 134619264 | 146268224 | 159672912 | 144448736 | 144501472 | 140588512 | 138438096 | 131971656 | **0.949557369** | 4 |
| sp\|P67775 | P67775 | PPP2CA | Serine/threonine-protein phosphatase 2A catalytic subur | 25985816 | 33185824 | 31652830 | 24288392 | 30438846 | 26356594 | 29802620 | 22683610 | **0.949343697** | 6 |
| sp\|Q9NTZ( | Q9NTZ6 | RBM12 | RNA-binding protein 12 OS=Homo sapiens (Human) GN= | 13219468 | 15163902 | 16454086 | 16021418 | 15048808 | 13546868 | 14751542 | 14385256 | **0.948628691** | 4 |
| sp\|Q9260( | Q92600 | CNOT9 | CCR4-NOT transcription complex subunit 9 OS=Homo sa | 1317737.5 | 1540395.5 | 1553941.12 | 1483530.38 | 1351898.5 | 1002531.75 | 1481647.5 | 1756190.88 | **0.94854881** | 6 |
| sp\|O7591! | O75915 | ARL6IP5 | PRA1 family protein 3 OS=Homo sapiens (Human) GN=A | 2204880 | 2565283 | 2019823.5 | 2369232.5 | 2091046.38 | 2290670 | 2315960.5 | 1987567.75 | **0.948251661** | 5 |
| sp\|O1526( | O15260 | SURF4 | Surfeit locus protein 4 OS=Homo sapiens (Human) GN=S | 7162578 | 8495492 | 7161650 | 10286764 | 6954101.5 | 8388242 | 7268217 | 8782534 | **0.948246105** | 7 |
| sp\|O4381: | O43813 | LANCL1 | LanC-like protein 1 OS=Homo sapiens (Human) GN=LAN( | 7691082 | 6857711 | 7821570 | 6238101 | 6940810.5 | 6231638 | 6308051.5 | 7641619.5 | **0.948045288** | 5 |
| sp\|P55786 | P55786 | NPEPPS | Puromycin-sensitive aminopeptidase OS=Homo sapiens | 95675768 | 99412616 | 98046400 | 100105936 | 94899152 | 94578112 | 86915872 | 96338776 | **0.947846683** | 1 |
| sp\|P06744 | P06744 | GPI | Glucose-6-phosphate isomerase OS=Homo sapiens (Hun | 207053696 | 215135696 | 208494080 | 205579136 | 220311648 | 207250288 | 196990112 | 167919728 | **0.947635071** | 6 |
| sp\|P22102 | P22102 | GART | Trifunctional purine biosynthetic protein adenosine-3 O! | 74930592 | 73092312 | 67976448 | 71671992 | 65844768 | 66984236 | 71298664 | 68430016 | **0.947462059** | 2 |
| sp\|Q9H90( | Q9H900 | ZWILCH | Protein zwilch homolog OS=Homo sapiens (Human) GN= | 252554.47 | 649020.25 | 695290.69 | 569782.75 | 543669.69 | 660736.75 | 220423.77 | 627914.31 | **0.94742864** | 6 |
| sp\|P23526 | P23526 | AHCY | Adenosylhomocysteinase OS=Homo sapiens (Human) GI | 199817248 | 220062624 | 269420704 | 218811136 | 215417424 | 216379104 | 208274048 | 220187104 | **0.947303805** | 6 |
| sp\|P50148 | P50148 | GNAQ | Guanine nucleotide-binding protein G(q) subunit alpha ( | 2907249.25 | 2297138.5 | 2419141.25 | 854456 | 2571844.5 | 2304609.75 | 2180747 | 973535.19 | **0.947245889** | 7 |
| sp\|O4340: | O43402 | EMC8 | ER membrane protein complex subunit 8 OS=Homo sapi | 1641823.62 | 1496588 | 1591813.88 | 1975805.12 | 2229839.75 | 1853646.12 | 2268640 | 0 | **0.947225897** | 5 |
| sp\|Q6ZRS: | Q6ZRS2 | SRCAP | Helicase SRCAP OS=Homo sapiens (Human) GN=SRCAP P | 2399995 | 1544745.5 | 1724964.75 | 1479010.12 | 1824880 | 1583401.88 | 1450281.5 | 1910848.5 | **0.946941028** | 8 |
| sp\|Q1359( | Q13596 | SNX1 | Sorting nexin-1 OS=Homo sapiens (Human) GN=SNX1 PE | 6609713.5 | 6667062 | 6869939 | 6082636 | 6435451 | 5750579.5 | 6321373.5 | 6329739.5 | **0.946921789** | 3 |
| sp\|Q5SY1( | Q5SY16 | NOL9 | Polynucleotide 5'-hydroxyl-kinase NOL9 OS=Homo sapie | 1645278.88 | 1677808.38 | 2043154.75 | 1055672 | 868714.88 | 845284.5 | 2017527.25 | 2347702.25 | **0.946638163** | 7 |
| sp\|P63279 | P63279 | UBE2I | SUMO-conjugating enzyme UBC9 OS=Homo sapiens (Hu | 1588073.5 | 1478111.5 | 1125150 | 1941350.88 | 1551819.88 | 1882473.88 | 1288916.5 | 1081123.38 | **0.946458657** | 6 |
| sp\|Q5T28( | Q5T280 | SPOUT1 | Putative methyltransferase C9orf114 OS=Homo sapiens | 1583190.12 | 1351140.12 | 1595406.38 | 1340590.62 | 1299036.5 | 1342520.75 | 1447254 | 1467157.5 | **0.94644958** | 5 |
| sp\|P40429 | P40429 | RPL13A | 60S ribosomal protein L13a OS=Homo sapiens (Human) ( | 84665552 | 80737400 | 96923744 | 88827816 | 87696760 | 80752784 | 76371944 | 87514240 | **0.946408822** | 5 |
| sp\|P01111 | P01111 | NRAS | GTPase NRas OS=Homo sapiens (Human) GN=NRAS PE=1 | 9094299 | 8886957 | 9146840 | 10672980 | 6923466 | 10670861 | 8532716 | 9629840 | **0.945922359** | 6 |
| sp\|P33897 | P33897 | ABCD1 | ATP-binding cassette sub-family D member 1 OS=Homo | 952226 | 981188.31 | 1114857.25 | 953421.56 | 885292.62 | 1109460.25 | 795071.62 | 994970.69 | **0.945798457** | 6 |
| sp\|Q1653: | Q16531 | DDB1 | DNA damage-binding protein 1 OS=Homo sapiens (Hum | 57062116 | 63240848 | 60691828 | 63874448 | 57012832 | 57424340 | 55454208 | 61693236 | **0.94574809** | 3 |
| sp\|Q1355! | Q13555 | CAMK2G | Calcium/calmodulin-dependent protein kinase type II sul | 3042427.5 | 2959837.25 | 3030874.25 | 3019772.5 | 3044661 | 2786051.25 | 3183150.5 | 2383276.5 | **0.945592212** | 8 |
| sp\|P50579 | P50579 | METAP2 | Methionine aminopeptidase 2 OS=Homo sapiens (Huma | 8113672 | 6360312 | 6203578 | 4567231 | 7196045 | 5615580 | 5997595.5 | 5047397 | **0.945011413** | 6 |
| sp\|P21399 | P21399 | ACO1 | Cytoplasmic aconitate hydratase OS=Homo sapiens (Hur | 15830051 | 13508824 | 13434904 | 14102265 | 14477999 | 14463838 | 12577716 | 12228474 | **0.945002908** | 6 |

| | | | | | | | | | | | | | |
|---|---|---|---|---|---|---|---|---|---|---|---|---|---|
| sp\|O6083( | O60830 | TIMM17B | Mitochondrial import inner membrane translocase subu | 1106718.62 | 2074737.38 | 1631585.25 | 1012535.62 | 1749999.88 | 1011985.69 | 1671656.62 | 1069833.75 | **0.944709179** | 7 |
| sp\|P56537 | P56537 | EIF6 | Eukaryotic translation initiation factor 6 OS=Homo sapie | 26218964 | 30284352 | 31467028 | 27861126 | 28467440 | 25625136 | 28830654 | 26501098 | **0.944685654** | 5 |
| sp\|O7535: | O75352 | MPDU1 | Mannose-P-dolichol utilization defect 1 protein OS=Hom | 15009107 | 15277328 | 16897166 | 16527914 | 14203813 | 15932522 | 14940527 | 15096536 | **0.944466601** | 3 |
| sp\|Q8TCJ2 | Q8TCJ2 | STT3B | Dolichyl-diphosphooligosaccharide--protein glycosyltran | 7128489 | 6324564 | 6956961.5 | 7536427 | 6859494.5 | 6772278 | 6814833 | 5943296.5 | **0.944302766** | 3 |
| sp\|Q9H97 | Q9H974 | QTRT2 | Queuine tRNA-ribosyltransferase accessory subunit 2 OS | 580960.5 | 521608.34 | 446594.56 | 562479.88 | 488213.06 | 602523.81 | 488743.75 | 414373 | **0.944218959** | 6 |
| sp\|Q1308! | Q13085 | ACACA | Acetyl-CoA carboxylase 1 OS=Homo sapiens (Human) GN | 20776554 | 20805454 | 22968614 | 19159148 | 20300550 | 16217830 | 19297276 | 23203216 | **0.943962359** | 6 |
| sp\|Q1419 | Q14197 | MRPL58 | Peptidyl-tRNA hydrolase ICT1, mitochondrial OS=Homo s | 1607051.75 | 1302070.88 | 1607866.75 | 1327267.75 | 1827547.88 | 1904856 | 547862.12 | 1234218.5 | **0.943573217** | 8 |
| sp\|Q1669 | Q16698 | DECR1 | 2,4-dienoyl-CoA reductase, mitochondrial OS=Homo sap | 6100524 | 9833531 | 8047176 | 7835570 | 7262204.5 | 7370413.5 | 8848313 | 6532125.5 | **0.943308427** | 6 |
| sp\|P22695 | P22695 | UQCRC2 | Cytochrome b-c1 complex subunit 2, mitochondrial OS=I | 44110280 | 41582792 | 46224844 | 40405216 | 43013456 | 40235788 | 39497208 | 39724316 | **0.942826225** | 2 |
| sp\|Q8IY37 | Q8IY37 | DHX37 | Probable ATP-dependent RNA helicase DHX37 OS=Homo | 1369711.62 | 1146632 | 1074177.88 | 881855 | 965317.44 | 1203863.75 | 1187692.75 | 855892.25 | **0.941952492** | 7 |
| sp\|Q1401 | Q14012 | CAMK1 | Calcium/calmodulin-dependent protein kinase type 1 OS | 1211869.25 | 1018715.06 | 1412013.88 | 1126171.88 | 938130.75 | 1214333.12 | 922986.19 | 1414226.5 | **0.94147474** | 7 |
| sp\|Q1362( | Q13620 | CUL4B | Cullin-4B OS=Homo sapiens (Human) GN=CUL4B PE=1 SV | 6505381 | 6606373 | 4014925.5 | 6590888.5 | 5741922 | 5734340 | 5192200 | 5659504.5 | **0.941410456** | 4 |
| sp\|Q0966( | Q09666 | AHNAK | Neuroblast differentiation-associated protein AHNAK OS | 127468504 | 128942136 | 125129408 | 80372864 | 94271840 | 93748800 | 125576504 | 121244208 | **0.941392502** | 5 |
| sp\|Q96EE: | Q96EE3 | SEH1L | Nucleoporin SEH1 OS=Homo sapiens (Human) GN=SEH1I | 2160449.5 | 2158518.5 | 2547927 | 2337553.5 | 2241339.25 | 2246924.75 | 2076613.25 | 2099102.5 | **0.941281789** | 4 |
| sp\|P62195 | P62195 | PSMC5 | 26S proteasome regulatory subunit 8 OS=Homo sapiens | 33854196 | 38881496 | 34574384 | 35706272 | 32553384 | 33453580 | 33379484 | 35220536 | **0.941199981** | 2 |
| sp\|Q9Y2S | Q9Y2S7 | POLDIP2 | Polymerase delta-interacting protein 2 OS=Homo sapien | 5705543.5 | 6261256.5 | 6589727.5 | 7060601.5 | 5449664 | 6293863 | 5751363 | 6614708.5 | **0.941151465** | 6 |
| sp\|Q9NZ0 | Q9NZ08 | ERAP1 | Endoplasmic reticulum aminopeptidase 1 OS=Homo sap | 3009949.5 | 3247917.5 | 2651063.75 | 3255818 | 2617871 | 2814522 | 2869065 | 3143739.5 | **0.940849477** | 4 |
| sp\|Q0476( | Q04760 | GLO1 | Lactoylglutathione lyase OS=Homo sapiens (Human) GN | 23613168 | 23996840 | 20169202 | 23904386 | 26053776 | 20308884 | 20924208 | 18963536 | **0.940739759** | 6 |
| sp\|P30533 | P30533 | LRPAP1 | Alpha-2-macroglobulin receptor-associated protein OS=I | 15016891 | 5841729 | 12309492 | 11824168 | 13415748 | 9652710 | 12125886 | 7091679 | **0.939850637** | 7 |
| sp\|P62277 | P62277 | RPS13 | 40S ribosomal protein S13 OS=Homo sapiens (Human) G | 171487360 | 140660256 | 170987952 | 177539808 | 158214976 | 159496128 | 148753792 | 154366464 | **0.939691992** | 4 |
| sp\|Q0658 | Q06587 | RING1 | E3 ubiquitin-protein ligase RING1 OS=Homo sapiens (Hu | 3235982.75 | 2481596 | 3215713.5 | 2302278.25 | 2706559.5 | 2141763 | 2816742.25 | 2890817.75 | **0.939505698** | 6 |
| sp\|Q9Y4L1 | Q9Y4L1 | HYOU1 | Hypoxia up-regulated protein 1 OS=Homo sapiens (Hum | 124302376 | 120490392 | 118147120 | 119597400 | 113195824 | 104776560 | 120555760 | 114793168 | **0.939453433** | 3 |
| sp\|Q7Z4W | Q7Z4W1 | DCXR | L-xylulose reductase OS=Homo sapiens (Human) GN=DC | 23803276 | 26428388 | 27333266 | 25997736 | 25453454 | 23418706 | 21817550 | 26590406 | **0.939335764** | 4 |
| sp\|Q9NZT | Q9NZT2 | OGFR | Opioid growth factor receptor OS=Homo sapiens (Huma | 3871340.75 | 8155689.5 | 4123898.75 | 3528105 | 7531812 | 3425000.5 | 3789258.75 | 3736342.25 | **0.939193128** | 5 |
| sp\|Q0065: | Q00653 | NFKB2 | Nuclear factor NF-kappa-B p100 subunit OS=Homo sapie | 586705.75 | 823531.38 | 596755.19 | 662626.25 | 593020.38 | 600242.69 | 694022.94 | 619844.56 | **0.939134376** | 8 |
| sp\|Q9NP9 | Q9NP92 | MRPS30 | 39S ribosomal protein S30, mitochondrial OS=Homo sap | 5475574.5 | 769104.06 | 1115273 | 1145398.5 | 1222117.62 | 1530023.25 | 1435857.75 | 3798876.25 | **0.939041287** | 4 |
| sp\|P19338 | P19338 | NCL | Nucleolin OS=Homo sapiens (Human) GN=NCL PE=1 SV= | 1131345408 | 1133409152 | 1297164672 | 1137623680 | 1137165568 | 1015208896 | 1178114304 | 1082515456 | **0.939028392** | 5 |
| sp\|Q1657( | Q16576 | RBBP7 | Histone-binding protein RBBP7 OS=Homo sapiens (Huma | 24594906 | 28930052 | 27019140 | 25512456 | 24438986 | 23960986 | 23439596 | 27731938 | **0.938852926** | 3 |
| sp\|P07305 | P07305 | H1F0 | Histone H1.0 OS=Homo sapiens (Human) GN=H1F0 PE=1 | 111034728 | 112972576 | 92146648 | 114525312 | 107406600 | 82174648 | 121565192 | 93115864 | **0.938662104** | 6 |
| sp\|O1497 | O14979 | HNRNPDL | Heterogeneous nuclear ribonucleoprotein D-like OS=Ho | 65467564 | 59374876 | 71200208 | 70747656 | 66826408 | 57948728 | 68944728 | 56701172 | **0.938643692** | 4 |
| sp\|Q9UNX | Q9UNX3 | RPL26L1 | 60S ribosomal protein L26-like 1 OS=Homo sapiens (Hun | 36245356 | 42329008 | 61562504 | 41922080 | 54636872 | 37085344 | 42277676 | 36888460 | **0.938642972** | 7 |
| sp\|Q9H3S | Q9H3S7 | PTPN23 | Tyrosine-protein phosphatase non-receptor type 23 OS= | 1304948 | 1972212.12 | 3557797.25 | 1950284.88 | 2327560.5 | 2115338.25 | 2053465.62 | 1748620.5 | **0.938503986** | 6 |
| sp\|P00167 | P00167 | CYB5A | Cytochrome b5 OS=Homo sapiens (Human) GN=CYB5A P | 15286100 | 12641403 | 12072188 | 11487194 | 13430064 | 14385041 | 11135922 | 9368468 | **0.938481615** | 6 |
| sp\|P42785 | P42785 | PRCP | Lysosomal Pro-X carboxypeptidase OS=Homo sapiens (H | 2358963.5 | 2444455.5 | 1326669.5 | 2229384.5 | 2089478 | 2313882.25 | 2051475.25 | 1389953 | **0.938430987** | 5 |
| sp\|P48651 | P48651 | PTDSS1 | Phosphatidylserine synthase 1 OS=Homo sapiens (Huma | 1271923 | 1220807.5 | 1237813.12 | 1236246.62 | 1315529.38 | 1339273.5 | 903640.5 | 1100933.62 | **0.938106257** | 8 |
| sp\|P30046 | P30046 | DDT | D-dopachrome decarboxylase OS=Homo sapiens (Huma | 12937510 | 10482756 | 9960483 | 11095158 | 11116181 | 11911737 | 10322318 | 8372432 | **0.938095945** | 7 |
| sp\|Q1505( | Q15050 | RRS1 | Ribosome biogenesis regulatory protein homolog OS=Hc | 6186736 | 7135070.5 | 8833442 | 11143933 | 9529639 | 6350256 | 8334593 | 7018193.5 | **0.937941418** | 7 |
| sp\|P22061 | P22061 | PCMT1 | Protein-L-isoaspartate(D-aspartate) O-methyltransferase | 11572216 | 10089673 | 7373358.5 | 8703018 | 9660730 | 8286076.5 | 10990867 | 6452447 | **0.937778142** | 6 |
| sp\|Q3LXA: | Q3LXA3 | TKFC | Triokinase/FMN cyclase OS=Homo sapiens (Human) GN= | 119509256 | 126857672 | 125685008 | 113920232 | 118058928 | 106725256 | 116297872 | 114599168 | **0.937669385** | 3 |
| sp\|Q1468: | Q14683 | SMC1A | Structural maintenance of chromosomes protein 1A OS= | 27923900 | 26818122 | 35014256 | 26190804 | 27212976 | 24393576 | 29495178 | 27614346 | **0.937635291** | 7 |
| sp\|Q1542 | Q15424 | SAFB | Scaffold attachment factor B1 OS=Homo sapiens (Huma | 14552388 | 13023969 | 12665273 | 11492975 | 12044476 | 12165341 | 11518495 | 12762268 | **0.937294873** | 5 |
| sp\|Q96SY( | Q96SY0 | INTS14 | Integrator complex subunit 14 OS=Homo sapiens (Huma | 459522.97 | 540197.44 | 591551.75 | 498335.28 | 551433.5 | 510877.88 | 426950.06 | 468897.88 | **0.937094347** | 6 |
| sp\|Q1543( | Q15436 | SEC23A | Protein transport protein Sec23A OS=Homo sapiens (Hu | 55598008 | 52635128 | 48559528 | 52258068 | 46686352 | 48805336 | 48364584 | 51785384 | **0.935857311** | 2 |
| sp\|Q6P2Q | Q6P2Q9 | PRPF8 | Pre-mRNA-processing-splicing factor 8 OS=Homo sapien | 71226040 | 75123944 | 76451208 | 78466824 | 69127704 | 67159632 | 70006408 | 75568024 | **0.935584772** | 2 |
| sp\|O7564: | O75643 | SNRNP200 | U5 small nuclear ribonucleoprotein 200 kDa helicase OS= | 99781888 | 98413408 | 101164376 | 100045976 | 84796472 | 94221080 | 94109136 | 100510024 | **0.935481794** | 3 |

| | | | | | | | | | | | | | |
|---|---|---|---|---|---|---|---|---|---|---|---|---|---|
| sp\|Q96Q1 | Q96Q11 | TRNT1 | CCA tRNA nucleotidyltransferase 1, mitochondrial OS=H | 2538649.5 | 4113697 | 4131917.75 | 3797420 | 3820503 | 2961608.5 | 3370518.75 | 3486196.25 | **0.935339585** | 5 |
| sp\|Q9BWH | Q9BWH6 | RPAP1 | RNA polymerase II-associated protein 1 OS=Homo sapie | 0 | 346843.41 | 424929.84 | 463863.53 | 289871.53 | 267169.28 | 316886.09 | 281304.31 | **0.934927827** | 4 |
| sp\|P42765 | P42765 | ACAA2 | 3-ketoacyl-CoA thiolase, mitochondrial OS=Homo sapier | 52029840 | 45175968 | 45931028 | 49817220 | 40443784 | 43406368 | 48950448 | 47596760 | **0.934923907** | 4 |
| sp\|Q9GZT | Q9GZT9 | EGLN1 | Egl nine homolog 1 OS=Homo sapiens (Human) GN=EGLI | 559002.81 | 527576.56 | 658060.56 | 622375.06 | 620060.75 | 562798 | 561939.19 | 467822.88 | **0.934772627** | 6 |
| sp\|P38919 | P38919 | EIF4A3 | Eukaryotic initiation factor 4A-III OS=Homo sapiens (Hun | 87308312 | 91006032 | 104726880 | 95293872 | 88596456 | 87000056 | 85665080 | 92384920 | **0.934744135** | 3 |
| sp\|P27797 | P27797 | CALR | Calreticulin OS=Homo sapiens (Human) GN=CALR PE=1 S | 395153696 | 354263904 | 386148192 | 337083744 | 350599968 | 351407264 | 311607712 | 361695904 | **0.93390234** | 4 |
| sp\|Q3MHI | Q3MHD2 | LSM12 | Protein LSM12 homolog OS=Homo sapiens (Human) GN= | 7555489.5 | 3922040.25 | 3592517.5 | 3607405.75 | 4551478 | 5816879.5 | 3404473.25 | 3664392 | **0.933597464** | 8 |
| sp\|P41235 | P41235 | HNF4A | Hepatocyte nuclear factor 4-alpha OS=Homo sapiens (Hu | 6292228 | 5593396 | 6676560 | 5934779 | 6236326 | 5083644.5 | 5971987 | 5574710 | **0.933449077** | 4 |
| sp\|P31939 | P31939 | ATIC | Bifunctional purine biosynthesis protein PURH OS=Hom | 240639200 | 235495696 | 244732144 | 240237104 | 235966512 | 226817376 | 225471712 | 208639936 | **0.933192872** | 1 |
| sp\|P45985 | P45985 | MAP2K4 | Dual specificity mitogen-activated protein kinase kinase | 1668977 | 1905933.5 | 1619393.62 | 1747094.75 | 1614713.25 | 1659251 | 1945489.75 | 1256148.62 | **0.932895911** | 5 |
| sp\|Q1449 | Q14498 | RBM39 | RNA-binding protein 39 OS=Homo sapiens (Human) GN= | 25986650 | 25194166 | 29984984 | 23401198 | 24720588 | 22264146 | 27566248 | 22974800 | **0.932663114** | 4 |
| sp\|Q1505 | Q15056 | EIF4H | Eukaryotic translation initiation factor 4H OS=Homo sapi | 25421400 | 26954876 | 27713380 | 23483282 | 23510758 | 24340370 | 31973436 | 16758763 | **0.932515084** | 6 |
| sp\|P41240 | P41240 | CSK | Tyrosine-protein kinase CSK OS=Homo sapiens (Human) | 3337211.5 | 2861426 | 3656729 | 1388993.75 | 2818606.5 | 2563559.75 | 2751308 | 2341413.25 | **0.931568117** | 4 |
| sp\|Q9UBS | Q9UBS4 | DNAJB11 | DnaJ homolog subfamily B member 11 OS=Homo sapien | 15990568 | 19282498 | 16852484 | 21405754 | 19226324 | 15404162 | 18747304 | 15119171 | **0.931534697** | 4 |
| sp\|P25788 | P25788 | PSMA3 | Proteasome subunit alpha type-3 OS=Homo sapiens (Hu | 25831912 | 27377004 | 43649024 | 32942778 | 28125396 | 27589028 | 34125692 | 31051448 | **0.93136283** | 7 |
| sp\|O6086 | O60869 | EDF1 | Endothelial differentiation-related factor 1 OS=Homo sa | 4939452.5 | 4488831 | 5867779 | 5067675 | 4491877 | 4374164.5 | 5426128.5 | 4671796 | **0.931261562** | 5 |
| sp\|P17066 | P17066 | HSPA6 | Heat shock 70 kDa protein 6 OS=Homo sapiens (Human) | 347879744 | 319968160 | 361451936 | 265954224 | 280756192 | 279896032 | 334815520 | 309884128 | **0.930591075** | 5 |
| sp\|P15104 | P15104 | GLUL | Glutamine synthetase OS=Homo sapiens (Human) GN=G | 5185594 | 6722591.5 | 5367713 | 3117176 | 3387828 | 1789982 | 6909631 | 6888514.5 | **0.930509791** | 7 |
| sp\|Q1536 | Q15365 | PCBP1 | Poly(rC)-binding protein 1 OS=Homo sapiens (Human) G | 204787120 | 206126112 | 277539744 | 209454688 | 239074128 | 185669600 | 205638768 | 204978640 | **0.930341915** | 5 |
| sp\|P04350 | P04350 | TUBB4A | Tubulin beta-4A chain OS=Homo sapiens (Human) GN=T | 1035261952 | 1159734528 | 1088613760 | 1105084672 | 1076908160 | 1021440512 | 1021415808 | 963112384 | **0.930316859** | 1 |
| sp\|P10620 | P10620 | MGST1 | Microsomal glutathione S-transferase 1 OS=Homo sapie | 11863491 | 11316691 | 8785853 | 10889494 | 8600101 | 9712951 | 8339418 | 13215349 | **0.930284141** | 5 |
| sp\|O1546 | O15460 | P4HA2 | Prolyl 4-hydroxylase subunit alpha-2 OS=Homo sapiens ( | 14343512 | 8618454 | 8218542.5 | 11913102 | 11468446 | 10813314 | 5942614 | 11863345 | **0.930247397** | 6 |
| sp\|P20290 | P20290 | BTF3 | Transcription factor BTF3 OS=Homo sapiens (Human) GN | 16846800 | 12087971 | 9054874 | 12144164 | 13683064 | 11094931 | 11369211 | 10487778 | **0.93021027** | 6 |
| sp\|Q1057 | Q10570 | CPSF1 | Cleavage and polyadenylation specificity factor subunit 1 | 6106570.5 | 6424641 | 4536625 | 6611561.5 | 4573448.5 | 3945084.25 | 5571458.5 | 7936692 | **0.930204528** | 6 |
| sp\|P52907 | P52907 | CAPZA1 | F-actin-capping protein subunit alpha-1 OS=Homo sapie | 59451084 | 61200012 | 60818140 | 59063096 | 56276336 | 55528640 | 58313428 | 53594288 | **0.930073268** | 0 |
| sp\|Q1502 | Q15029 | EFTUD2 | 116 kDa U5 small nuclear ribonucleoprotein component | 59794072 | 56876588 | 59370408 | 57221420 | 53661404 | 56586352 | 52313304 | 54379012 | **0.930025543** | 0 |
| sp\|Q8N1B | Q8N1B4 | VPS52 | Vacuolar protein sorting-associated protein 52 homolog | 311905.72 | 252282.14 | 212727.28 | 254110 | 232370.19 | 250486.97 | 257741.3 | 218114.42 | **0.929863728** | 6 |
| sp\|P04632 | P04632 | CAPNS1 | Calpain small subunit 1 OS=Homo sapiens (Human) GN= | 5713045 | 5524690 | 5698087 | 6132433 | 5379656.5 | 5190716 | 5456768 | 5423182.5 | **0.929863269** | 0 |
| sp\|P12268 | P12268 | IMPDH2 | Inosine-5'-monophosphate dehydrogenase 2 OS=Homo | 43538168 | 40863816 | 46745200 | 44435136 | 41827152 | 44621260 | 37086332 | 39636016 | **0.929312017** | 4 |
| sp\|O9539 | O95394 | PGM3 | Phosphoacetylglucosamine mutase OS=Homo sapiens (H | 7041236 | 7101381.5 | 7376475 | 8374115.5 | 6542289.5 | 5357089.5 | 7154939 | 8723728 | **0.929242723** | 6 |
| sp\|Q0902 | Q09028 | RBBP4 | Histone-binding protein RBBP4 OS=Homo sapiens (Hum | 37632152 | 41847140 | 46129812 | 37393180 | 37088024 | 37537736 | 36860656 | 39944420 | **0.929010516** | 3 |
| sp\|P11717 | P11717 | IGF2R | Cation-independent mannose-6-phosphate receptor OS | 45369380 | 51221280 | 48447208 | 44082616 | 41021044 | 40179496 | 43455168 | 50986432 | **0.928731443** | 3 |
| sp\|O6031 | O60313 | OPA1 | Dynamin-like 120 kDa protein, mitochondrial OS=Homo | 4566691 | 4898320 | 2745897.5 | 4979807 | 5446454 | 4721935 | 2850911.5 | 2942058 | **0.928487153** | 8 |
| sp\|Q8NC5 | Q8NC51 | SERBP1 | Plasminogen activator inhibitor 1 RNA-binding protein O | 50212916 | 48987176 | 58936340 | 53238948 | 54776464 | 44443280 | 56676300 | 40325844 | **0.928310043** | 6 |
| sp\|Q9BRJ | Q9BRJ2 | MRPL45 | 39S ribosomal protein L45, mitochondrial OS=Homo sapi | 2821496.75 | 2842133.25 | 2551120.25 | 3049433 | 2326484.25 | 2506696.75 | 2597380.5 | 3013429 | **0.927185777** | 4 |
| sp\|Q9BYD | Q9BYD3 | MRPL4 | 39S ribosomal protein L4, mitochondrial OS=Homo sapie | 4138163 | 3383059 | 3430948 | 4073353 | 3986519.75 | 3412684 | 3629678 | 2901057.5 | **0.92708515** | 5 |
| sp\|O7553 | O75533 | SF3B1 | Splicing factor 3B subunit 1 OS=Homo sapiens (Human) ( | 98963632 | 93583776 | 98091952 | 93919672 | 83140520 | 87474976 | 91673744 | 94022256 | **0.926545644** | 2 |
| sp\|P00558 | P00558 | PGK1 | Phosphoglycerate kinase 1 OS=Homo sapiens (Human) G | 355171328 | 327496928 | 332163232 | 356206944 | 369565056 | 281023296 | 321805280 | 297487776 | **0.926218681** | 4 |
| sp\|Q9BSC | Q9BSC4 | NOL10 | Nucleolar protein 10 OS=Homo sapiens (Human) GN=NC | 4868114 | 5332112.5 | 6745386 | 5939731 | 5213377 | 5366858 | 5231451 | 5383690.5 | **0.926155052** | 6 |
| sp\|O7508 | O75083 | WDR1 | WD repeat-containing protein 1 OS=Homo sapiens (Hum | 119909072 | 120643224 | 120425040 | 121354288 | 121256808 | 120783352 | 112673608 | 91985256 | **0.926124272** | 6 |
| sp\|Q9UHR | Q9UHR5 | SAP30BP | SAP30-binding protein OS=Homo sapiens (Human) GN=S | 1098979.38 | 1335238.75 | 1444362.25 | 1016285.88 | 1525967 | 0 | 1595770.5 | 1411086.62 | **0.926036357** | 5 |
| sp\|P60900 | P60900 | PSMA6 | Proteasome subunit alpha type-6 OS=Homo sapiens (Hu | 65771084 | 65252628 | 67505752 | 67888000 | 63119520 | 66995216 | 61631128 | 54919240 | **0.925859365** | 2 |
| sp\|P62979 | P62979 | RPS27A | Ubiquitin-40S ribosomal protein S27a OS=Homo sapiens | 258783536 | 302635712 | 308039392 | 292891168 | 254627184 | 289737952 | 223288656 | 308233984 | **0.925614448** | 5 |
| sp\|P40818 | P40818 | USP8 | Ubiquitin carboxyl-terminal hydrolase 8 OS=Homo sapie | 810667.25 | 711028.75 | 678269.81 | 542606.06 | 758829.56 | 577084 | 700344.75 | 501544.69 | **0.925336917** | 6 |
| sp\|P32322 | P32322 | PYCR1 | Pyrroline-5-carboxylate reductase 1, mitochondrial OS=H | 55117308 | 55245892 | 47982400 | 54257532 | 49782248 | 44587248 | 55299424 | 47021544 | **0.925153182** | 5 |

| | | | | | | | | | | | | |
|---|---|---|---|---|---|---|---|---|---|---|---|---|
| sp\|P52888 | P52888 | THOP1 | Thimet oligopeptidase OS=Homo sapiens (Human) GN=T | 18134114 | 17599638 | 23079712 | 23496224 | 19212754 | 18331852 | 19327190 | 19254942 | **0.924881868** | 8 |
| sp\|Q96I25 | Q96I25 | RBM17 | Splicing factor 45 OS=Homo sapiens (Human) GN=RBM1 | 3119959.75 | 4286129.5 | 4442418 | 3777895.5 | 3940858 | 3232801 | 4047170.75 | 3229292 | **0.924724774** | 6 |
| sp\|P07237 | P07237 | P4HB | Protein disulfide-isomerase OS=Homo sapiens (Human) | 1262118528 | 1241633664 | 1384620800 | 1018453824 | 1184401536 | 1108674432 | 1205005440 | 1037993088 | **0.924441531** | 4 |
| sp\|P30050 | P30050 | RPL12 | 60S ribosomal protein L12 OS=Homo sapiens (Human) G | 138408976 | 158525392 | 154825392 | 142466704 | 132847560 | 131611056 | 144195648 | 140657536 | **0.924414905** | 3 |
| sp\|Q96FZ | Q96FZ2 | HMCES | Embryonic stem cell-specific 5-hydroxymethylcytosine-bi | 378222.34 | 299608.97 | 328575.72 | 299761.25 | 245958.03 | 427676.69 | 302161.56 | 231328.23 | **0.924172274** | 6 |
| sp\|P31943 | P31943 | HNRNPH1 | Heterogeneous nuclear ribonucleoprotein H OS=Homo s | 192045968 | 205238592 | 210081168 | 187340368 | 194114944 | 171419920 | 201393376 | 167176848 | **0.923744126** | 4 |
| sp\|Q9BQ5 | Q9BQ52 | ELAC2 | Zinc phosphodiesterase ELAC protein 2 OS=Homo sapier | 2569585.5 | 3367592.5 | 2164690.25 | 2183126.25 | 2221646 | 3432868.75 | 2020060.5 | 1823067.62 | **0.923446568** | 6 |
| sp\|P53675 | P53675 | CLTCL1 | Clathrin heavy chain 2 OS=Homo sapiens (Human) GN=C | 75676616 | 76239696 | 67958856 | 74950816 | 60054472 | 69961536 | 67442432 | 74795072 | **0.923437983** | 2 |
| sp\|O6070 | O60701 | UGDH | UDP-glucose 6-dehydrogenase OS=Homo sapiens (Huma | 595521728 | 617987456 | 659366592 | 630459968 | 569298176 | 544353152 | 629508352 | 568143424 | **0.923289299** | 2 |
| sp\|Q9Y64 | Q9Y646 | CPQ | Carboxypeptidase Q OS=Homo sapiens (Human) GN=CP | 988240.5 | 1142035.5 | 1058356.62 | 888670 | 744921.06 | 1067599.5 | 907940.12 | 1043263.88 | **0.923091787** | 6 |
| sp\|P62269 | P62269 | RPS18 | 40S ribosomal protein S18 OS=Homo sapiens (Human) G | 223093456 | 230521008 | 272849440 | 249362272 | 239165360 | 210591184 | 235716416 | 215020608 | **0.922801202** | 4 |
| sp\|P09417 | P09417 | QDPR | Dihydropteridine reductase OS=Homo sapiens (Human) | 21540038 | 22538148 | 26554672 | 21779328 | 23264224 | 21206240 | 21916464 | 18800936 | **0.921825007** | 5 |
| sp\|P62847 | P62847 | RPS24 | 40S ribosomal protein S24 OS=Homo sapiens (Human) G | 59873876 | 56846728 | 56372920 | 68002288 | 56935744 | 58479064 | 51862048 | 54770104 | **0.920990531** | 4 |
| sp\|Q53GS | Q53GS9 | USP39 | U4/U6.U5 tri-snRNP-associated protein 2 OS=Homo sapi | 6474050 | 10737409 | 10940478 | 11214783 | 8649214 | 9497107 | 8955828 | 9152635 | **0.920950082** | 4 |
| sp\|Q0821 | Q08211 | DHX9 | ATP-dependent RNA helicase A OS=Homo sapiens (Huma | 177245392 | 185377344 | 183874960 | 181329232 | 167426704 | 161465520 | 161869392 | 179502896 | **0.920911945** | 1 |
| sp\|O0056 | O00566 | MPHOSPH | U3 small nucleolar ribonucleoprotein protein MPP10 OS | 3629310 | 5672197.5 | 5215116.5 | 4431921.5 | 3384641.5 | 3669589.5 | 5263308.5 | 5131129 | **0.920844742** | 6 |
| sp\|P28066 | P28066 | PSMA5 | Proteasome subunit alpha type-5 OS=Homo sapiens (Hu | 59356648 | 66123760 | 60569300 | 67786824 | 60871380 | 59377160 | 54345376 | 59140496 | **0.920806829** | 3 |
| sp\|Q1500 | Q15005 | SPCS2 | Signal peptidase complex subunit 2 OS=Homo sapiens (H | 19090636 | 16860192 | 20240294 | 17883052 | 18306856 | 15980513 | 18473462 | 15437383 | **0.920674647** | 4 |
| sp\|Q9BVJ | Q9BVJ6 | UTP14A | U3 small nucleolar RNA-associated protein 14 homolog A | 8201701.5 | 6954094.5 | 5328125.5 | 5201306 | 5869015 | 5200343 | 6488138.5 | 6087472 | **0.920566832** | 6 |
| sp\|Q96PK | Q96PK6 | RBM14 | RNA-binding protein 14 OS=Homo sapiens (Human) GN= | 32021204 | 30850640 | 34170960 | 29736700 | 32352434 | 27337960 | 30158452 | 26785888 | **0.919980993** | 4 |
| sp\|Q9H3P | Q9H3P7 | ACBD3 | Golgi resident protein GCP60 OS=Homo sapiens (Human | 9330609 | 10171091 | 10636082 | 7932022.5 | 10510772 | 7955627 | 8414363 | 8137956.5 | **0.919855485** | 6 |
| sp\|Q1573 | Q15738 | NSDHL | Sterol-4-alpha-carboxylate 3-dehydrogenase, decarboxyl | 4944609.5 | 4818278.5 | 3897039.75 | 4967027 | 3965683.75 | 4972611 | 3349128 | 4846018 | **0.919819744** | 7 |
| sp\|P04179 | P04179 | SOD2 | Superoxide dismutase [Mn], mitochondrial OS=Homo sa | 94667752 | 105178280 | 87314144 | 96556888 | 89323280 | 77908376 | 97660632 | 88011600 | **0.919698187** | 5 |
| sp\|P51812 | P51812 | RPS6KA3 | Ribosomal protein S6 kinase alpha-3 OS=Homo sapiens ( | 83554416 | 87498928 | 80068928 | 85268280 | 77392576 | 69815496 | 78027352 | 84088168 | **0.919537098** | 2 |
| sp\|P17987 | P17987 | TCP1 | T-complex protein 1 subunit alpha OS=Homo sapiens (H | 217259776 | 211849728 | 210319024 | 206374896 | 198493584 | 200699328 | 184001184 | 194534336 | **0.919514405** | 0 |
| sp\|Q8WU | Q8WUM4 | PDCD6IP | Programmed cell death 6-interacting protein OS=Homo | 40210052 | 42665344 | 46492652 | 40855472 | 40225528 | 39447504 | 38497936 | 38331060 | **0.919391327** | 1 |
| sp\|Q1322 | Q13228 | SELENBP1 | Methanethiol oxidase OS=Homo sapiens (Human) GN=S | 8594916 | 8500453 | 8027718 | 6230479 | 8370424 | 4376987.5 | 9036089 | 7039141.5 | **0.919277954** | 7 |
| sp\|P46063 | P46063 | RECQL | ATP-dependent DNA helicase Q1 OS=Homo sapiens (Hur | 5186105 | 5277518.5 | 5539843.5 | 5258091 | 5586087 | 5348906 | 5175602 | 3429455.5 | **0.919031921** | 7 |
| sp\|O9523 | O95232 | LUC7L3 | Luc7-like protein 3 OS=Homo sapiens (Human) GN=LUC7 | 6159057 | 6114411 | 7390953 | 6311067 | 6506630 | 5868257 | 5670858 | 5824167 | **0.918939887** | 3 |
| sp\|O7582 | O75822 | EIF3J | Eukaryotic translation initiation factor 3 subunit J OS=Ho | 6499785 | 7472806 | 7851507 | 7163242 | 6863933 | 5244289 | 8362816 | 6158641.5 | **0.918665856** | 5 |
| sp\|Q1317 | Q13177 | PAK2 | Serine/threonine-protein kinase PAK 2 OS=Homo sapien | 31341028 | 26623300 | 24206372 | 24472520 | 25467542 | 24647376 | 25172338 | 22656476 | **0.918424369** | 6 |
| sp\|Q8NB3 | Q8NB37 | GATD1 | Glutamine amidotransferase-like class 1 domain-containi | 1154298.75 | 1068897.38 | 1631820.5 | 1270264.12 | 1485444.75 | 1049031.5 | 1048291.81 | 1123621.25 | **0.918269562** | 4 |
| sp\|Q96QD | Q96QD8 | SLC38A2 | Sodium-coupled neutral amino acid transporter 2 OS=Ho | 28695220 | 26636404 | 17769520 | 26444508 | 20875904 | 24297966 | 21486868 | 24739420 | **0.918173282** | 4 |
| sp\|Q9HCM | Q9HCM4 | EPB41L5 | Band 4.1-like protein 5 OS=Homo sapiens (Human) GN=E | 1104247.5 | 1242806.75 | 1522624.25 | 1440416.25 | 913948.12 | 1238775.38 | 1244703.38 | 1477806.5 | **0.918106665** | 6 |
| sp\|Q0146 | Q01469 | FABP5 | Fatty acid-binding protein 5 OS=Homo sapiens (Human) | 13628867 | 35366476 | 19727610 | 19383966 | 21038590 | 22384518 | 20484352 | 16979642 | **0.918056186** | 6 |
| sp\|Q1361 | Q13619 | CUL4A | Cullin-4A OS=Homo sapiens (Human) GN=CUL4A PE=1 SV | 7476622 | 7051199 | 4639348 | 7609697.5 | 6667553.5 | 5785875.5 | 6094647 | 6021624.5 | **0.917571909** | 4 |
| sp\|Q9H2U | Q9H2U2 | PPA2 | Inorganic pyrophosphatase 2, mitochondrial OS=Homo s | 2951347.25 | 2617273.5 | 2087989.25 | 2341188.5 | 1441909.38 | 2432234 | 2519703.25 | 2772652.5 | **0.916851758** | 7 |
| sp\|P57081 | P57081 | WDR4 | tRNA (guanine-N(7)-)-methyltransferase non-catalytic sul | 486838.19 | 687436.5 | 662837.94 | 0 | 367381.91 | 603898.69 | 0 | 712599.44 | **0.91659053** | 7.5 |
| sp\|P50991 | P50991 | CCT4 | T-complex protein 1 subunit delta OS=Homo sapiens (Hu | 159376992 | 185463440 | 191504672 | 173524800 | 177036352 | 159572512 | 161404352 | 152563152 | **0.916472672** | 4 |
| sp\|Q1581 | Q15813 | TBCE | Tubulin-specific chaperone E OS=Homo sapiens (Human | 4407495 | 3896225.75 | 3939400.5 | 3977814 | 3960373.25 | 3616349.75 | 3702617 | 3584041 | **0.916308509** | 2 |
| sp\|Q9NQ2 | Q9NQ29 | LUC7L | Putative RNA-binding protein Luc7-like 1 OS=Homo sapie | 3557948 | 4126567.5 | 6316077.5 | 4260008 | 4349441 | 3080326.5 | 5160063.5 | 4134664.75 | **0.915878713** | 8 |
| sp\|Q1071 | Q10713 | PMPCA | Mitochondrial-processing peptidase subunit alpha OS=H | 6566204 | 11260054 | 6683034 | 11838749 | 11010878 | 12296327 | 4817933.5 | 5164279 | **0.915851765** | 6 |
| sp\|A6NCE | A6NCE7 | MAP1LC3B | Microtubule-associated proteins 1A/1B light chain 3 beta | 1484838.75 | 1090164.88 | 1311560.75 | 1123479.75 | 1600892.75 | 831300.19 | 982862.81 | 1171376.5 | **0.915447475** | 6 |
| sp\|P06730 | P06730 | EIF4E | Eukaryotic translation initiation factor 4E OS=Homo sapi | 10000098 | 13282420 | 9804374 | 10469448 | 11682036 | 9967072 | 10941802 | 7270553 | **0.915170168** | 7 |
| sp\|Q5VW3 | Q5VW36 | FOCAD | Focadhesin OS=Homo sapiens (Human) GN=FOCAD PE=1 | 0 | 283500.97 | 326593.97 | 0 | 0 | 253945.27 | 0 | 304246.78 | **0.914926536** | 7 |

| | | | | | | | | | | | | | |
|---|---|---|---|---|---|---|---|---|---|---|---|---|---|
| sp\|Q9UPN | Q9UPN6 | SCAF8 | Protein SCAF8 OS=Homo sapiens (Human) GN=SCAF8 PE | 4658678.5 | 3802161.25 | 4422882.5 | 4337695 | 3593389.5 | 4047663.5 | 4133732.5 | 3978201.5 | **0.914732323** | 3 |
| sp\|O7560( | O75600 | GCAT | 2-amino-3-ketobutyrate coenzyme A ligase, mitochondri | 905987.38 | 465066.31 | 804582.69 | 827965.62 | 422623.06 | 785881.19 | 790240.19 | 748662.81 | **0.914704162** | 3 |
| sp\|Q1299( | Q12996 | CSTF3 | Cleavage stimulation factor subunit 3 OS=Homo sapiens | 1518307.38 | 1766888.25 | 1612724.88 | 2065394 | 1559448.62 | 1507062.88 | 1541661.75 | 1759869.38 | **0.914513142** | 4 |
| sp\|Q1400{ | Q14008 | CKAP5 | Cytoskeleton-associated protein 5 OS=Homo sapiens (Hu | 18263336 | 15562023 | 22058322 | 17599926 | 18926168 | 14500986 | 16835164 | 16880588 | **0.913712714** | 5 |
| sp\|P47897 | P47897 | QARS | Glutamine--tRNA ligase OS=Homo sapiens (Human) GN= | 53055176 | 56496344 | 58147984 | 52232660 | 50156564 | 49082964 | 51054312 | 50648404 | **0.913655558** | 0 |
| sp\|Q9H50 | Q9H501 | ESF1 | ESF1 homolog OS=Homo sapiens (Human) GN=ESF1 PE= | 2534325.75 | 3165465.5 | 3360825 | 2362895 | 2964826.5 | 2722508 | 2174679.5 | 2571836.75 | **0.913366348** | 6 |
| sp\|P55060 | P55060 | CSE1L | Exportin-2 OS=Homo sapiens (Human) GN=CSE1L PE=1 S | 116914032 | 110504312 | 124597456 | 107529376 | 106727664 | 109643040 | 98101376 | 105043720 | **0.912893491** | 1 |
| sp\|O1480 | O14802 | POLR3A | DNA-directed RNA polymerase III subunit RPC1 OS=Hom | 2872059.75 | 2844095.5 | 2365740.5 | 2351725 | 1859523.25 | 2364922.5 | 2760767.25 | 2538646.5 | **0.912804838** | 5 |
| sp\|Q9UNN | Q9UNM6 | PSMD13 | 26S proteasome non-ATPase regulatory subunit 13 OS=H | 56128408 | 71630304 | 74311272 | 51520800 | 62064456 | 53292884 | 62661584 | 53324960 | **0.912272443** | 6 |
| sp\|O1456 | O14562 | UBFD1 | Ubiquitin domain-containing protein UBFD1 OS=Homo s | 4281834.5 | 4281667 | 2986904.75 | 3853834.25 | 4290095 | 2637737 | 4365885.5 | 2758406.5 | **0.912224397** | 8 |
| sp\|Q9NUD | Q9NUD5 | ZCCHC3 | Zinc finger CCHC domain-containing protein 3 OS=Homo | 11474515 | 12963523 | 13128268 | 11932880 | 10508051 | 11529392 | 11553632 | 11561487 | **0.91218797** | 3 |
| sp\|Q1508 | Q15084 | PDIA6 | Protein disulfide-isomerase A6 OS=Homo sapiens (Huma | 319307040 | 320905952 | 320891360 | 299833344 | 294845888 | 266840240 | 300917952 | 287204704 | **0.911868039** | 1 |
| sp\|Q1478 | Q14789 | GOLGB1 | Golgin subfamily B member 1 OS=Homo sapiens (Humar | 986712.12 | 1053819.62 | 1447586 | 883608.25 | 885959.69 | 877796.31 | 958028.38 | 1263964.12 | **0.911710503** | 5 |
| sp\|O1517 | O15173 | PGRMC2 | Membrane-associated progesterone receptor componen | 16097532 | 13564292 | 9852270 | 10081950 | 12656242 | 9866738 | 11712016 | 10973527 | **0.91153486** | 7 |
| sp\|O7582 | O75821 | EIF3G | Eukaryotic translation initiation factor 3 subunit G OS=H | 30995814 | 30260236 | 39049348 | 34852412 | 34512964 | 28875848 | 30785324 | 29025106 | **0.911521443** | 3 |
| sp\|Q9H4A | Q9H4A4 | RNPEP | Aminopeptidase B OS=Homo sapiens (Human) GN=RNPE | 14684774 | 14661328 | 20507360 | 14404136 | 15274714 | 15251111 | 14115964 | 13924078 | **0.911423222** | 6 |
| sp\|Q7Z4H | Q7Z4H8 | KDELC2 | KDEL motif-containing protein 2 OS=Homo sapiens (Hum | 2534670.75 | 2727648 | 2921718 | 2827151 | 2846689 | 3011327.75 | 2176640.25 | 1999779.25 | **0.911294629** | 7 |
| sp\|Q9UPY | Q9UPY3 | DICER1 | Endoribonuclease Dicer OS=Homo sapiens (Human) GN= | 1434797 | 2244558.5 | 1306204 | 1310186.75 | 758989 | 1459424 | 1604243.62 | 1913897.88 | **0.911179433** | 7 |
| sp\|P63244 | P63244 | RACK1 | Receptor of activated protein C kinase 1 OS=Homo sapie | 199445216 | 201958832 | 220877504 | 193183904 | 193055088 | 177998112 | 183290128 | 188645952 | **0.911122935** | 0 |
| sp\|Q9984 | Q99848 | EBNA1BP2 | Probable rRNA-processing protein EBP2 OS=Homo sapie | 3822704.25 | 9796874 | 25881244 | 3139434 | 10485911 | 3881345.5 | 20602616 | 3872436.25 | **0.910930472** | 6 |
| sp\|P11142 | P11142 | HSPA8 | Heat shock cognate 71 kDa protein OS=Homo sapiens (H | 1191415424 | 987959360 | 1473473280 | 995618880 | 1207006080 | 1015191104 | 1040447040 | 971395456 | **0.910846464** | 7 |
| sp\|Q9NQZ | Q9NQZ2 | UTP3 | Something about silencing protein 10 OS=Homo sapiens | 5751462.5 | 5853271 | 5159261.5 | 5900079 | 4574653 | 5730110 | 5340670.5 | 4996967 | **0.910798319** | 2 |
| sp\|P01034 | P01034 | CST3 | Cystatin-C OS=Homo sapiens (Human) GN=CST3 PE=1 SV | 1924314.5 | 1720177.12 | 2273293.75 | 2049706.25 | 2463029 | 1359883 | 1880824.38 | 1548460.38 | **0.910223331** | 5 |
| sp\|P49247 | P49247 | RPIA | Ribose-5-phosphate isomerase OS=Homo sapiens (Huma | 4049446.25 | 3309299.5 | 2632099.25 | 3620874 | 3526261 | 2807748 | 2875260.25 | 3177949.75 | **0.910040752** | 5 |
| sp\|Q68EM | Q68EM7 | ARHGAP17 | Rho GTPase-activating protein 17 OS=Homo sapiens (Hu | 3203610.25 | 2484256.5 | 4053112 | 3135276.5 | 2818537 | 2694354.25 | 2973434.75 | 3231038 | **0.909997804** | 6 |
| sp\|P31350 | P31350 | RRM2 | Ribonucleoside-diphosphate reductase subunit M2 OS=I | 5571188 | 2336093.75 | 8975295 | 5261514 | 6630821.5 | 5712319.5 | 3710115.5 | 4096696.5 | **0.909947183** | 8 |
| sp\|P62753 | P62753 | RPS6 | 40S ribosomal protein S6 OS=Homo sapiens (Human) GN | 65549060 | 60012504 | 73780528 | 65663396 | 60466300 | 62592392 | 58303780 | 59711168 | **0.90969301** | 2 |
| sp\|Q9Y27 | Q9Y277 | VDAC3 | Voltage-dependent anion-selective channel protein 3 OS | 15625865 | 17265084 | 15774572 | 16861182 | 14130912 | 14634101 | 12546850 | 18241122 | **0.908835364** | 4 |
| sp\|P35250 | P35250 | RFC2 | Replication factor C subunit 2 OS=Homo sapiens (Humar | 5724868 | 5360119.5 | 6679143 | 6424180.5 | 5707843.5 | 5711106 | 4961659.5 | 5600751 | **0.908759607** | 3 |
| sp\|Q9Y3A | Q9Y3A4 | RRP7A | Ribosomal RNA-processing protein 7 homolog A OS=Hor | 1887143.75 | 1755528 | 1837494 | 2041485.5 | 1813134.25 | 1566608.62 | 1922746 | 1532774 | **0.908744987** | 4 |
| sp\|P35606 | P35606 | COPB2 | Coatomer subunit beta' OS=Homo sapiens (Human) GN= | 53488476 | 61553712 | 58326372 | 56119784 | 54764112 | 48177600 | 51096824 | 54501712 | **0.908718257** | 2 |
| sp\|Q8NAV | Q8NAV1 | PRPF38A | Pre-mRNA-splicing factor 38A OS=Homo sapiens (Humar | 2124191.5 | 1277352.62 | 2566836 | 2524027 | 2289581 | 1316044.25 | 2861084.5 | 1249537.38 | **0.908605419** | 7 |
| sp\|Q9HAS | Q9HAS0 | C17orf75 | Protein Njmu-R1 OS=Homo sapiens (Human) GN=C17orf | 166473.25 | 177414.25 | 106099.6 | 184605.98 | 149880.67 | 162593.95 | 138346.77 | 125726.64 | **0.908531858** | 4 |
| sp\|O9477 | O94776 | MTA2 | Metastasis-associated protein MTA2 OS=Homo sapiens ( | 19065560 | 14808998 | 18569748 | 17601688 | 16644300 | 17035944 | 14885376 | 15072776 | **0.908522991** | 4 |
| sp\|Q5QJE | Q5QJE6 | DNTTIP2 | Deoxynucleotidyltransferase terminal-interacting proteir | 8724837 | 9447546 | 10095184 | 9434986 | 8829867 | 7796497.5 | 9653956 | 7968373.5 | **0.908391907** | 4 |
| sp\|Q86TU | Q86TU7 | SETD3 | Histone-lysine N-methyltransferase setd3 OS=Homo sap | 978531.06 | 1469132.38 | 1997135.5 | 1742975.75 | 1098306.75 | 1612110.5 | 1403742.25 | 1506750.75 | **0.908389612** | 6 |
| sp\|Q1358 | Q13586 | STIM1 | Stromal interaction molecule 1 OS=Homo sapiens (Huma | 1437559.5 | 2067911.75 | 2144330.75 | 1928599.88 | 1382857.12 | 2054728.38 | 1602434.88 | 1841327.5 | **0.908020977** | 4 |
| sp\|P28482 | P28482 | MAPK1 | Mitogen-activated protein kinase 1 OS=Homo sapiens (H | 23053636 | 22307348 | 21031314 | 18524712 | 21114082 | 20668744 | 18561176 | 16760247 | **0.907995336** | 4 |
| sp\|Q9UJX | Q9UJX4 | ANAPC5 | Anaphase-promoting complex subunit 5 OS=Homo sapie | 258340.03 | 454618.41 | 353614 | 444771.59 | 301883.56 | 273108.28 | 377555.88 | 419391.03 | **0.907760723** | 6 |
| sp\|Q9Y3U | Q9Y3U8 | RPL36 | 60S ribosomal protein L36 OS=Homo sapiens (Human) G | 38006692 | 33807024 | 34879620 | 37250324 | 36476196 | 30471296 | 32654356 | 31006162 | **0.907355072** | 2 |
| sp\|Q8WTT | Q8WTT2 | NOC3L | Nucleolar complex protein 3 homolog OS=Homo sapiens | 8339087.5 | 7550157 | 9018938 | 8223000 | 7449453 | 7955670 | 7225410 | 7430459.5 | **0.907332315** | 1 |
| sp\|Q0275 | Q02750 | MAP2K1 | Dual specificity mitogen-activated protein kinase kinase | 13134872 | 9389255 | 12773900 | 12669653 | 12881386 | 10567315 | 10649443 | 9414024 | **0.90711429** | 6 |
| sp\|Q9BVG | Q9BVG4 | PBDC1 | Protein PBDC1 OS=Homo sapiens (Human) GN=PBDC1 PI | 6307015 | 5961302 | 6800666 | 6841079 | 5137092 | 6118145 | 5570745 | 6676446 | **0.907077258** | 3 |
| sp\|Q1415 | Q14152 | EIF3A | Eukaryotic translation initiation factor 3 subunit A OS=H | 106093232 | 104685920 | 119838424 | 95393808 | 102356616 | 93427528 | 92126704 | 98108336 | **0.906124105** | 2 |
| sp\|Q96PU | Q96PU8 | QKI | Protein quaking OS=Homo sapiens (Human) GN=QKI PE= | 4210542 | 3917999 | 4359660.5 | 4198764 | 4229892 | 3694515 | 4938486 | 2255998.75 | **0.906030024** | 7 |

| | | | | | | | | | | | | | |
|---|---|---|---|---|---|---|---|---|---|---|---|---|---|
| sp\|P23396 | P23396 | RPS3 | 40S ribosomal protein S3 OS=Homo sapiens (Human) GN | 293728736 | 301986048 | 350998176 | 328697664 | 310382912 | 273965024 | 268701056 | 301675136 | **0.9053744** | 3 |
| sp\|P00390 | P00390 | GSR | Glutathione reductase, mitochondrial OS=Homo sapiens | 64550992 | 60850040 | 62101656 | 58135088 | 53729872 | 55401088 | 54469632 | 58573852 | **0.904479953** | 1 |
| sp\|Q9BXR( | Q9BXR0 | QTRT1 | Queuine tRNA-ribosyltransferase catalytic subunit 1 OS= | 7773795.5 | 6621562.5 | 7847218 | 6904695 | 6146803.5 | 7046197.5 | 7227805 | 5930357.5 | **0.904069664** | 4 |
| sp\|P62913 | P62913 | RPL11 | 60S ribosomal protein L11 OS=Homo sapiens (Human) G | 122743464 | 123261512 | 153617552 | 139926480 | 118947848 | 118257336 | 127128128 | 123374296 | **0.903917162** | 4 |
| sp\|Q9BXY( | Q9BXY0 | MAK16 | Protein MAK16 homolog OS=Homo sapiens (Human) GN | 3729516.75 | 4838272 | 2497534.5 | 3020368.5 | 1483904.12 | 2724153 | 3868655 | 4655185.5 | **0.903888701** | 7 |
| sp\|Q8IXH7 | Q8IXH7 | NELFCD | Negative elongation factor C/D OS=Homo sapiens (Huma | 3539071.75 | 4640794.5 | 3765887.75 | 3678479 | 3820667 | 3215434 | 3656314.5 | 3422503.75 | **0.903399178** | 4 |
| sp\|Q8WYA | Q8WYA6 | CTNNBL1 | Beta-catenin-like protein 1 OS=Homo sapiens (Human) G | 9037426 | 8845261 | 9368045 | 9258366 | 8238770.5 | 8300064 | 8787335 | 7651249.5 | **0.903265783** | 0 |
| sp\|Q9980( | Q99808 | SLC29A1 | Equilibrative nucleoside transporter 1 OS=Homo sapiens | 1672604.5 | 2241815 | 2105480.25 | 2488554 | 2035355 | 2443947.75 | 1295082.75 | 1910335.5 | **0.903186551** | 5 |
| sp\|Q6P1M | Q6P1M0 | SLC27A4 | Long-chain fatty acid transport protein 4 OS=Homo sapie | 1417709.25 | 1607196.12 | 1913106.25 | 1184680.25 | 1534865.25 | 1182324.75 | 1464041.12 | 1348192.88 | **0.903103425** | 5 |
| sp\|P17844 | P17844 | DDX5 | Probable ATP-dependent RNA helicase DDX5 OS=Homo s | 225821744 | 212839968 | 263529008 | 235213344 | 229225776 | 185065728 | 217449168 | 214522336 | **0.902772924** | 4 |
| sp\|Q96ST2 | Q96ST2 | IWS1 | Protein IWS1 homolog OS=Homo sapiens (Human) GN=I' | 4999714 | 4527380.5 | 4676622 | 4567203 | 4170561.25 | 4317223 | 4366394 | 4090668.25 | **0.902717978** | 0 |
| sp\|P19525 | P19525 | EIF2AK2 | Interferon-induced, double-stranded RNA-activated prot | 7567086.5 | 7700518.5 | 9523308 | 8988644 | 8130761 | 7732116 | 7113432 | 7507213 | **0.902425156** | 4 |
| sp\|P46778 | P46778 | RPL21 | 60S ribosomal protein L21 OS=Homo sapiens (Human) G | 43167088 | 39506844 | 39360888 | 42983752 | 38064656 | 38595920 | 39097668 | 33133832 | **0.90227466** | 0 |
| sp\|P11216 | P11216 | PYGB | Glycogen phosphorylase, brain form OS=Homo sapiens ( | 52800808 | 58593232 | 62064276 | 61115160 | 49957780 | 51161092 | 52072032 | 58387344 | **0.901970042** | 1 |
| sp\|P52735 | P52735 | VAV2 | Guanine nucleotide exchange factor VAV2 OS=Homo sap | 1473866.5 | 1737302.75 | 1625405.5 | 1839980.75 | 1425684.75 | 1504180.75 | 1643225.38 | 1448842 | **0.901952044** | 3 |
| sp\|P19387 | P19387 | POLR2C | DNA-directed RNA polymerase II subunit RPB3 OS=Homo | 3164350.25 | 3121052.25 | 2947397.75 | 3683548 | 2597896 | 2989032 | 3008919 | 3050075.5 | **0.901642033** | 3 |
| sp\|P57772 | P57772 | EEFSEC | Selenocysteine-specific elongation factor OS=Homo sapi | 1759403.5 | 1297117.25 | 1647221.25 | 1942441.75 | 862473.81 | 825816.38 | 2309084.5 | 1994845.75 | **0.901603185** | 8 |
| sp\|P61247 | P61247 | RPS3A | 40S ribosomal protein S3a OS=Homo sapiens (Human) G | 231037392 | 216215808 | 260271344 | 241244368 | 226366528 | 221509568 | 199718160 | 207746592 | **0.90152706** | 2 |
| sp\|Q9Y678 | Q9Y678 | COPG1 | Coatomer subunit gamma-1 OS=Homo sapiens (Human) | 85588640 | 73522296 | 68969088 | 77853912 | 66731688 | 66614060 | 68559992 | 73861120 | **0.901393496** | 2 |
| sp\|O0042! | O00425 | IGF2BP3 | Insulin-like growth factor 2 mRNA-binding protein 3 OS= | 76921608 | 75226648 | 74082448 | 69741568 | 73564872 | 62777552 | 68816240 | 61572604 | **0.90120357** | 1 |
| sp\|Q8NFH | Q8NFH5 | NUP35 | Nucleoporin NUP35 OS=Homo sapiens (Human) GN=NUI | 4645880 | 4040318.25 | 5047714.5 | 1993839.62 | 3610717.25 | 2099699.75 | 4571659.5 | 3885625.25 | **0.900809055** | 5 |
| sp\|Q9249! | Q92499 | DDX1 | ATP-dependent RNA helicase DDX1 OS=Homo sapiens (H | 37004740 | 36821688 | 38455040 | 37444684 | 33091232 | 31218792 | 35330332 | 35223084 | **0.900734028** | 0 |
| sp\|Q7L0Y3 | Q7L0Y3 | TRMT10C | tRNA methyltransferase 10 homolog C OS=Homo sapien | 1508840.38 | 1650863 | 1816164.75 | 1557266.25 | 1403850.25 | 1524471 | 1337899 | 1617962.5 | **0.900667644** | 3 |
| sp\|P08758 | P08758 | ANXA5 | Annexin A5 OS=Homo sapiens (Human) GN=ANXA5 PE=1 | 338761600 | 335976224 | 348873120 | 322040544 | 325976096 | 277558400 | 306459424 | 301548800 | **0.900339152** | 1 |
| sp\|P22314 | P22314 | UBA1 | Ubiquitin-like modifier-activating enzyme 1 OS=Homo sa | 146651584 | 148670592 | 143458720 | 137288480 | 126064032 | 129019440 | 136398720 | 127006424 | **0.900045442** | 0 |
| sp\|P78371 | P78371 | CCT2 | T-complex protein 1 subunit beta OS=Homo sapiens (Hu | 142781824 | 122198544 | 124358448 | 135951296 | 128434560 | 124265600 | 115448816 | 104473456 | **0.899736015** | 3 |
| sp\|P11586 | P11586 | MTHFD1 | C-1-tetrahydrofolate synthase, cytoplasmic OS=Homo sa | 146428448 | 142615824 | 168701120 | 148580816 | 153084240 | 129251784 | 134399168 | 128797440 | **0.89973454** | 3 |
| sp\|P36578 | P36578 | RPL4 | 60S ribosomal protein L4 OS=Homo sapiens (Human) GN | 335186752 | 323102432 | 389252384 | 292161088 | 328680320 | 311837152 | 304651136 | 259874720 | **0.89948566** | 4 |
| sp\|O0023 | O00232 | PSMD12 | 26S proteasome non-ATPase regulatory subunit 12 OS=H | 29144292 | 29569784 | 30866734 | 30702428 | 25507390 | 27618052 | 26497970 | 28563782 | **0.899436994** | 0 |
| sp\|P61158 | P61158 | ACTR3 | Actin-related protein 3 OS=Homo sapiens (Human) GN=/ | 36010832 | 39670256 | 26283442 | 33104256 | 30437508 | 32893104 | 30731610 | 27403934 | **0.899291092** | 4 |
| sp\|Q6Y7W | Q6Y7W6 | GIGYF2 | GRB10-interacting GYF protein 2 OS=Homo sapiens (Hun | 4835312.5 | 4352319.5 | 5396114 | 3286878.5 | 3951490 | 4543229 | 3649793.25 | 3922072.5 | **0.899049988** | 5 |
| sp\|P62906 | P62906 | RPL10A | 60S ribosomal protein L10a OS=Homo sapiens (Human) ( | 131907592 | 125544960 | 151635152 | 136139760 | 143770480 | 122648544 | 109994392 | 113649952 | **0.898823703** | 3 |
| sp\|Q8N0U | Q8N0U8 | VKORC1L1 | Vitamin K epoxide reductase complex subunit 1-like prot | 1864795.5 | 2229609 | 2516660.25 | 2325701.25 | 2175699.5 | 2089855.88 | 1800782.88 | 1963443.38 | **0.898510898** | 3 |
| sp\|P62873 | P62873 | GNB1 | Guanine nucleotide-binding protein G(I)/G(S)/G(T) subun | 40884512 | 36543048 | 38026348 | 31278196 | 34312932 | 31021064 | 32508228 | 33931036 | **0.898053367** | 3 |
| sp\|Q0288( | Q02880 | TOP2B | DNA topoisomerase 2-beta OS=Homo sapiens (Human) ( | 11519202 | 14835622 | 18671780 | 14743457 | 10271595 | 12252046 | 12497821 | 18650272 | **0.897970206** | 5 |
| sp\|Q9NP7 | Q9NP79 | VTA1 | Vacuolar protein sorting-associated protein VTA1 homolo | 10173262 | 10698912 | 11180798 | 11592222 | 10217292 | 11029226 | 8453493 | 9491438 | **0.89795566** | 3 |
| sp\|O9490( | O94906 | PRPF6 | Pre-mRNA-processing factor 6 OS=Homo sapiens (Huma | 12468990 | 13446292 | 15986824 | 12664652 | 10703787 | 10833506 | 14556882 | 12897614 | **0.897832138** | 5 |
| sp\|Q8WW | Q8WWI5 | SLC44A1 | Choline transporter-like protein 1 OS=Homo sapiens (Hu | 1309706.25 | 1497926.25 | 1221680 | 1404850.5 | 1356788.5 | 1291138.75 | 1093685.75 | 1131411.38 | **0.896738721** | 3 |
| sp\|Q9NV8 | Q9NV88 | INTS9 | Integrator complex subunit 9 OS=Homo sapiens (Human | 234847.23 | 213136.47 | 168243.92 | 253265.98 | 205615.38 | 251813.17 | 172497.06 | 149595.95 | **0.896523632** | 5 |
| sp\|Q9971 | Q99714 | HSD17B10 | 3-hydroxyacyl-CoA dehydrogenase type-2 OS=Homo sap | 19377650 | 36180016 | 16477466 | 20055238 | 19856660 | 19758370 | 20771944 | 22162980 | **0.896401589** | 6 |
| sp\|Q1324 | Q13247 | SRSF6 | Serine/arginine-rich splicing factor 6 OS=Homo sapiens ( | 58326328 | 59009128 | 59590316 | 58619132 | 43023308 | 52476300 | 61037480 | 54414516 | **0.89558976** | 4 |
| sp\|O4339( | O43396 | TXNL1 | Thioredoxin-like protein 1 OS=Homo sapiens (Human) G | 26049912 | 28086608 | 30829278 | 24718050 | 26148318 | 25970920 | 24818656 | 21259668 | **0.895278236** | 4 |
| sp\|P05388 | P05388 | RPLP0 | 60S acidic ribosomal protein P0 OS=Homo sapiens (Hum | 150248832 | 155692176 | 172528832 | 163652192 | 152432784 | 142948656 | 132189808 | 147240256 | **0.895174866** | 1 |
| sp\|P67812 | P67812 | SEC11A | Signal peptidase complex catalytic subunit SEC11A OS=H | 11824822 | 16049476 | 15612419 | 14875542 | 13609050 | 13459732 | 12672218 | 12485956 | **0.894875505** | 4 |
| sp\|Q9BRZ | Q9BRZ2 | TRIM56 | E3 ubiquitin-protein ligase TRIM56 OS=Homo sapiens (H | 1112527 | 1003695.12 | 966641.62 | 352159.66 | 556408.94 | 714687.5 | 865160.12 | 937265.38 | **0.894760117** | 4 |

| | | | | | | | | | | | | |
|---|---|---|---|---|---|---|---|---|---|---|---|---|
| sp\|Q9UHA | Q9UHA3 | RSL24D1 | Probable ribosome biogenesis protein RLP24 OS=Homo s | 2327262.25 | 2039059.38 | 2135483.75 | 2295333 | 2174462 | 1508726.12 | 1866576.12 | 2318411.75 | **0.894401753** | 5 |
| sp\|P35998 | P35998 | PSMC2 | 26S proteasome regulatory subunit 7 OS=Homo sapiens | 35727372 | 28813220 | 45077108 | 35873856 | 39810420 | 35183300 | 24053240 | 31050836 | **0.894194822** | 5 |
| sp\|O4377€ | O43776 | NARS | Asparagine--tRNA ligase, cytoplasmic OS=Homo sapiens | 41854488 | 46838516 | 38939772 | 47339684 | 37453772 | 34890980 | 41088932 | 42996108 | **0.89402522** | 3 |
| sp\|P05023 | P05023 | ATP1A1 | Sodium/potassium-transporting ATPase subunit alpha-1 | 180542080 | 192485536 | 176587792 | 175266816 | 184181936 | 184590256 | 122490816 | 156763184 | **0.893974456** | 6 |
| sp\|Q6UB3 | Q6UB35 | MTHFD1L | Monofunctional C1-tetrahydrofolate synthase, mitochon | 21949184 | 20822144 | 23079300 | 23046810 | 18385784 | 19789864 | 19546240 | 21750104 | **0.89397393** | 1 |
| sp\|Q0551 | Q05519 | SRSF11 | Serine/arginine-rich splicing factor 11 OS=Homo sapiens | 10422966 | 9968082 | 6733976 | 9652336 | 8060620.5 | 7813972.5 | 9029123 | 7952068 | **0.893369834** | 4 |
| sp\|P17980 | P17980 | PSMC3 | 26S proteasome regulatory subunit 6A OS=Homo sapien | 33462456 | 35365656 | 39949608 | 36258584 | 33709696 | 34188056 | 34323396 | 27335168 | **0.893268185** | 3 |
| sp\|P50453 | P50453 | SERPINB9 | Serpin B9 OS=Homo sapiens (Human) GN=SERPINB9 PE= | 11560710 | 12357258 | 8918888 | 12192212 | 7301231.5 | 12103367 | 12390927 | 8424688 | **0.893205551** | 6 |
| sp\|P61225 | P61225 | RAP2B | Ras-related protein Rap-2b OS=Homo sapiens (Human) ( | 3122610.25 | 3166787.25 | 3448590.5 | 3988247 | 3313599.25 | 3060141.5 | 2726173.25 | 3159808.25 | **0.893159869** | 3 |
| sp\|O9545€ | O95456 | PSMG1 | Proteasome assembly chaperone 1 OS=Homo sapiens (H | 4460629 | 4636674 | 5684583.5 | 4860080 | 4597130 | 3968327.5 | 4427659.5 | 4545715 | **0.89292648** | 2 |
| sp\|Q1534 | Q15349 | RPS6KA2 | Ribosomal protein S6 kinase alpha-2 OS=Homo sapiens ( | 15623772 | 15642304 | 11263666 | 16203180 | 13249410 | 13429726 | 11444132 | 14309770 | **0.892736752** | 4 |
| sp\|P22392 | P22392 | NME2 | Nucleoside diphosphate kinase B OS=Homo sapiens (Hu | 206073792 | 212069088 | 234343152 | 195859072 | 208313744 | 189546784 | 192853936 | 166608816 | **0.892706608** | 2 |
| sp\|O9598 | O95983 | MBD3 | Methyl-CpG-binding domain protein 3 OS=Homo sapien | 1007537.38 | 1086515.5 | 985852.5 | 1094052.75 | 992873.12 | 1081811.5 | 914084 | 735656.81 | **0.892300621** | 3 |
| sp\|Q0158 | Q01581 | HMGCS1 | Hydroxymethylglutaryl-CoA synthase, cytoplasmic OS=H | 11607522 | 12346373 | 19462446 | 12758158 | 14266583 | 8080641.5 | 14665139 | 13111553 | **0.892289516** | 7 |
| sp\|Q9NVX | Q9NVX2 | NLE1 | Notchless protein homolog 1 OS=Homo sapiens (Human | 1184372.5 | 1091316.62 | 1389467 | 1355458.62 | 1194544 | 1091316.38 | 1105561.12 | 1088383.88 | **0.892282243** | 3 |
| sp\|Q6P1M | Q6P1M3 | LLGL2 | Lethal(2) giant larvae protein homolog 2 OS=Homo sapie | 2002176 | 2308620.75 | 2042525.5 | 2006821.25 | 1902291.5 | 1896597.38 | 1794670.62 | 1865713.25 | **0.892242191** | 0 |
| sp\|Q8IY81 | Q8IY81 | FTSJ3 | pre-rRNA processing protein FTSJ3 OS=Homo sapiens (H | 23169408 | 24974328 | 30644472 | 24917408 | 26503000 | 23098676 | 20746390 | 22164608 | **0.89207005** | 3 |
| sp\|Q9H58 | Q9H583 | HEATR1 | HEAT repeat-containing protein 1 OS=Homo sapiens (Hu | 22451048 | 23955046 | 23883880 | 24518380 | 19985974 | 21333634 | 20229660 | 23002058 | **0.891813036** | 1 |
| sp\|Q1355 | Q13557 | CAMK2D | Calcium/calmodulin-dependent protein kinase type II sul | 5257823 | 5018281 | 6067068.5 | 5408386 | 5433696.5 | 4821788.5 | 4866981 | 4272541 | **0.891660568** | 3 |
| sp\|Q1304 | Q13045 | FLII | Protein flightless-1 homolog OS=Homo sapiens (Human) | 6864999 | 6419807 | 7006550 | 6637478 | 5769424 | 7348747 | 4744724 | 6147374.5 | **0.891619351** | 4 |
| sp\|P07910 | P07910 | HNRNPC | Heterogeneous nuclear ribonucleoproteins C1/C2 OS=H | 325177760 | 351747040 | 294536416 | 328197600 | 269859744 | 314326016 | 285932768 | 288144480 | **0.891205441** | 1 |
| sp\|Q6NU | Q6NUM9 | RETSAT | All-trans-retinol 13,14-reductase OS=Homo sapiens (Hur | 4138741 | 4378153 | 4429769.5 | 4595456.5 | 3399337.5 | 3141542 | 4932204 | 4153058.5 | **0.890778424** | 5 |
| sp\|P55036 | P55036 | PSMD4 | 26S proteasome non-ATPase regulatory subunit 4 OS=H | 21682984 | 20397172 | 23896696 | 22989436 | 20749902 | 15032490 | 19530918 | 23932936 | **0.89074466** | 5 |
| sp\|P18124 | P18124 | RPL7 | 60S ribosomal protein L7 OS=Homo sapiens (Human) GN | 228747056 | 233941472 | 285017760 | 243330784 | 227300544 | 237544752 | 210689424 | 207123824 | **0.890641298** | 2 |
| sp\|P46779 | P46779 | RPL28 | 60S ribosomal protein L28 OS=Homo sapiens (Human) G | 125504488 | 123151648 | 176002224 | 135469792 | 141882144 | 115374968 | 131348208 | 109910592 | **0.890003315** | 5 |
| sp\|Q1497 | Q14974 | KPNB1 | Importin subunit beta-1 OS=Homo sapiens (Human) GN= | 115175536 | 100686232 | 117584024 | 98985336 | 102621808 | 92081416 | 94379712 | 95731184 | **0.889885337** | 2 |
| sp\|Q8IWA | Q8IWA0 | WDR75 | WD repeat-containing protein 75 OS=Homo sapiens (Hu | 3629494 | 4181013 | 3558236.25 | 4193391 | 3266940 | 3941775 | 2594775 | 4042011.5 | **0.889691689** | 4 |
| sp\|Q9NW | Q9NWT1 | PAK1IP1 | p21-activated protein kinase-interacting protein 1 OS=H | 1490934.75 | 1978497.25 | 1933121.25 | 1372090.88 | 1578093.5 | 1747092.25 | 1115788.25 | 1582306.25 | **0.889091757** | 6 |
| sp\|Q53FA | Q53FA7 | TP53I3 | Quinone oxidoreductase PIG3 OS=Homo sapiens (Huma | 3762192.75 | 2522637 | 3821331.75 | 4044971.5 | 3028152 | 2911254 | 3766528.5 | 2867236 | **0.888492144** | 5 |
| sp\|P30154 | P30154 | PPP2R1B | Serine/threonine-protein phosphatase 2A 65 kDa regulat | 18299184 | 15615770 | 17986242 | 17553414 | 15638948 | 16397960 | 13406586 | 16259827 | **0.888397775** | 3 |
| sp\|P68371 | P68371 | TUBB4B | Tubulin beta-4B chain OS=Homo sapiens (Human) GN=T | 1288070144 | 1449623552 | 1419460224 | 1403876480 | 1293615232 | 1225620480 | 1286210432 | 1134823808 | **0.888373125** | 1 |
| sp\|P07437 | P07437 | TUBB | Tubulin beta chain OS=Homo sapiens (Human) GN=TUBE | 1593546368 | 1736410368 | 1770685056 | 1718359936 | 1572482688 | 1500038400 | 1554064384 | 1430241920 | **0.888227872** | 0 |
| sp\|P56182 | P56182 | RRP1 | Ribosomal RNA processing protein 1 homolog A OS=Hon | 8468564 | 8414824 | 3477407 | 7643252 | 5946214.5 | 3181029 | 8401510 | 7336497 | **0.887916325** | 4 |
| sp\|Q9BUQ | Q9BUQ8 | DDX23 | Probable ATP-dependent RNA helicase DDX23 OS=Homo | 4698489.5 | 8249601 | 10287458 | 10893890 | 5888589 | 7096042 | 10034176 | 7284543.5 | **0.887894786** | 5 |
| sp\|P54725 | P54725 | RAD23A | UV excision repair protein RAD23 homolog A OS=Homo s | 6745325 | 5817389 | 9467699 | 7373006.5 | 8095845 | 6523453 | 6254192.5 | 5231517 | **0.887822163** | 5 |
| sp\|Q6RFH | Q6RFH5 | WDR74 | WD repeat-containing protein 74 OS=Homo sapiens (Hu | 1867617 | 994346.38 | 1261531.62 | 1388389.12 | 1223948.88 | 1196308 | 1491832.12 | 979463.38 | **0.887455591** | 5 |
| sp\|Q1500 | Q15008 | PSMD6 | 26S proteasome non-ATPase regulatory subunit 6 OS=H | 27839232 | 23764666 | 22976278 | 25288922 | 22412342 | 22568052 | 22845260 | 20726594 | **0.886683166** | 0 |
| sp\|P12694 | P12694 | BCKDHA | 2-oxoisovalerate dehydrogenase subunit alpha, mitocho | 919066.69 | 908294 | 574467.44 | 634055.44 | 831880.56 | 1036125.5 | 0 | 823507.88 | **0.886566918** | 8 |
| sp\|Q9UNS | Q9UNS2 | COPS3 | COP9 signalosome complex subunit 3 OS=Homo sapiens | 4112941.5 | 4508061.5 | 4990348.5 | 5662979.5 | 3672569 | 4701180.5 | 4105897.5 | 4608138 | **0.886556581** | 4 |
| sp\|Q8WXF | Q8WXF1 | PSPC1 | Paraspeckle component 1 OS=Homo sapiens (Human) GI | 40768804 | 36590652 | 37773472 | 42138928 | 33273530 | 34467224 | 37318180 | 34340536 | **0.886359922** | 1 |
| sp\|P62249 | P62249 | RPS16 | 40S ribosomal protein S16 OS=Homo sapiens (Human) G | 174707216 | 159386464 | 200842304 | 176416960 | 161778256 | 161630848 | 155511408 | 151514400 | **0.886247702** | 2 |
| sp\|Q6GM | Q6GMV2 | SMYD5 | SET and MYND domain-containing protein 5 OS=Homo s | 190219.2 | 173227.72 | 132050.91 | 274741.88 | 109646.58 | 200709.75 | 120196.17 | 251831.06 | **0.885936613** | 6 |
| sp\|Q1316 | Q13162 | PRDX4 | Peroxiredoxin-4 OS=Homo sapiens (Human) GN=PRDX4 | 95303248 | 71625928 | 75145672 | 53487132 | 68638472 | 75967120 | 64651932 | 52551620 | **0.885801158** | 5 |
| sp\|Q6UW | Q6UWP7 | LCLAT1 | Lysocardiolipin acyltransferase 1 OS=Homo sapiens (Hur | 1415740.75 | 1464399.62 | 1448405.12 | 1457775 | 1270129.62 | 1205193.75 | 1283692.75 | 1366472.38 | **0.885794091** | 0 |
| sp\|Q7Z7K | Q7Z7K6 | CENPV | Centromere protein V OS=Homo sapiens (Human) GN=C | 4821284.5 | 5302146.5 | 4686782 | 5248611 | 4593934 | 5410247.5 | 3498292.5 | 4265038 | **0.885770372** | 4 |

| | | | | | | | | | | | | | |
|---|---|---|---|---|---|---|---|---|---|---|---|---|---|
| sp\|P37268 | P37268 | FDFT1 | Squalene synthase OS=Homo sapiens (Human) GN=FDFT | 8746155 | 7463092.5 | 10023320 | 9252384 | 7479882.5 | 6337531.5 | 8176278.5 | 9434624 | **0.885680131** | 5 |
| sp\|P51149 | P51149 | RAB7A | Ras-related protein Rab-7a OS=Homo sapiens (Human) ( | 57537432 | 59177688 | 54534120 | 59815256 | 56260512 | 54765384 | 44646128 | 48975496 | **0.885672717** | 2 |
| sp\|P46926 | P46926 | GNPDA1 | Glucosamine-6-phosphate isomerase 1 OS=Homo sapier | 7443483 | 8895079 | 7401934 | 9260493 | 7609329.5 | 7272239 | 5435326 | 8907783 | **0.885569748** | 5 |
| sp\|Q9BUN | Q9BUN8 | DERL1 | Derlin-1 OS=Homo sapiens (Human) GN=DERL1 PE=1 SV= | 2501511.5 | 2951541.75 | 2852386.5 | 2630654.5 | 2513437.75 | 2674516.75 | 2179043.75 | 2316742.5 | **0.88548439** | 3 |
| sp\|Q0766( | Q07666 | KHDRBS1 | KH domain-containing, RNA-binding, signal transduction- | 19652720 | 14573754 | 15697457 | 20729520 | 16090758 | 16560344 | 15169165 | 14724340 | **0.885230744** | 6 |
| sp\|Q9NQX | Q9NQX3 | GPHN | Gephyrin OS=Homo sapiens (Human) GN=GPHN PE=1 SV | 4119933.25 | 2667898.5 | 3729043.5 | 3039042 | 3097662.5 | 2993562.5 | 2196230 | 3709605 | **0.885005402** | 5 |
| sp\|Q9292; | Q92922 | SMARCC1 | SWI/SNF complex subunit SMARCC1 OS=Homo sapiens ( | 3655972.5 | 1794817.12 | 2869910 | 5517241 | 1204256.25 | 1338060.88 | 5630204 | 4072206.75 | **0.884866341** | 7 |
| sp\|P55196 | P55196 | AFDN | Afadin OS=Homo sapiens (Human) GN=AFDN PE=1 SV=3 | 13030212 | 15923538 | 16019042 | 14566000 | 12918880 | 11578132 | 15711848 | 12461436 | **0.884638304** | 2 |
| sp\|P55212 | P55212 | CASP6 | Caspase-6 OS=Homo sapiens (Human) GN=CASP6 PE=1 S | 4441215.5 | 4261469 | 5134332 | 5185880.5 | 3943044.5 | 3971071.25 | 4342003.5 | 4571639 | **0.884605444** | 3 |
| sp\|Q3ZCM | Q3ZCM7 | TUBB8 | Tubulin beta-8 chain OS=Homo sapiens (Human) GN=TU | 406764864 | 625085120 | 649458944 | 569735552 | 504843136 | 456380928 | 561478720 | 467906784 | **0.884304857** | 4 |
| sp\|Q9Y4R{ | Q9Y4R8 | TELO2 | Telomere length regulation protein TEL2 homolog OS=H | 745494.44 | 595042.12 | 570799.12 | 721334.75 | 556919.38 | 590294.62 | 610684.31 | 569702.06 | **0.884121439** | 3 |
| sp\|P29144 | P29144 | TPP2 | Tripeptidyl-peptidase 2 OS=Homo sapiens (Human) GN= | 5630168.5 | 6014807 | 5343148.5 | 3709157.75 | 5899421.5 | 3650731.25 | 5584020.5 | 3161079.25 | **0.8839447** | 5 |
| sp\|P50990 | P50990 | CCT8 | T-complex protein 1 subunit theta OS=Homo sapiens (Hu | 213112160 | 204073328 | 237789152 | 207634864 | 198469280 | 188700528 | 184625696 | 190492928 | **0.883700479** | 0 |
| sp\|Q7Z5K. | Q7Z5K2 | WAPL | Wings apart-like protein homolog OS=Homo sapiens (Hu | 855523.94 | 836947.88 | 869158.56 | 846157.25 | 981793.88 | 619405.44 | 1111555.12 | 298526.84 | **0.883646989** | 8 |
| sp\|Q5JRX3 | Q5JRX3 | PITRM1 | Presequence protease, mitochondrial OS=Homo sapiens | 5234069 | 5360399 | 4255894 | 3847730.5 | 4258113 | 3975685 | 4022493.5 | 4258928.5 | **0.883256942** | 6 |
| sp\|P83111 | P83111 | LACTB | Serine beta-lactamase-like protein LACTB, mitochondrial | 738134.19 | 926797.31 | 786663.94 | 974679.06 | 606036.25 | 837305.88 | 804134.69 | 778749.62 | **0.883241095** | 5 |
| sp\|Q1388! | Q13885 | TUBB2A | Tubulin beta-2A chain OS=Homo sapiens (Human) GN=T | 1097320832 | 1240313088 | 1218750848 | 1189396480 | 1083554048 | 1040621760 | 1115024768 | 952226688 | **0.883190152** | 1 |
| sp\|P11413 | P11413 | G6PD | Glucose-6-phosphate 1-dehydrogenase OS=Homo sapie | 81030856 | 78614064 | 84883040 | 86675472 | 78285520 | 78849056 | 67129280 | 68094536 | **0.882715467** | 1 |
| sp\|P50851 | P50851 | LRBA | Lipopolysaccharide-responsive and beige-like anchor pro | 6193516.5 | 5784762 | 6079072.5 | 7199313 | 7549279 | 3477141 | 7056367 | 4206873.5 | **0.882525915** | 7 |
| sp\|Q1562! | Q15629 | TRAM1 | Translocating chain-associated membrane protein 1 OS= | 5275944 | 4244842.5 | 4140005.5 | 4461447 | 4551616 | 4357225.5 | 4019809.5 | 3058674 | **0.882193696** | 5 |
| sp\|P43490 | P43490 | NAMPT | Nicotinamide phosphoribosyltransferase OS=Homo sapi | 13689852 | 14900290 | 13285506 | 13797760 | 11335652 | 12990442 | 11944573 | 12822144 | **0.881799997** | 0 |
| sp\|P78347 | P78347 | GTF2I | General transcription factor II-I OS=Homo sapiens (Huma | 32226444 | 31462330 | 33506662 | 35089132 | 30178904 | 29787410 | 27649480 | 28877592 | **0.880627179** | 0 |
| sp\|Q9H3U | Q9H3U1 | UNC45A | Protein unc-45 homolog A OS=Homo sapiens (Human) G | 14259086 | 11265309 | 12922317 | 10339377 | 11023273 | 13483130 | 10087668 | 8364601.5 | **0.880551677** | 4 |
| sp\|P48723 | P48723 | HSPA13 | Heat shock 70 kDa protein 13 OS=Homo sapiens (Human | 1525070.38 | 1785376.25 | 1636608 | 1429847.75 | 1487461.12 | 1150702.62 | 1610932.88 | 1365284.12 | **0.880424445** | 3 |
| sp\|Q9UDX | Q9UDX5 | MTFP1 | Mitochondrial fission process protein 1 OS=Homo sapier | 598434.5 | 517104.56 | 375010.38 | 705963.19 | 397732.12 | 726179.12 | 426046.78 | 383720.19 | **0.880340128** | 7 |
| sp\|P52732 | P52732 | KIF11 | Kinesin-like protein KIF11 OS=Homo sapiens (Human) GI | 411782.88 | 394035.12 | 838366.38 | 641038.56 | 715361 | 570614.69 | 0 | 724859.81 | **0.879929684** | 8 |
| sp\|Q9Y281 | Q9Y281 | CFL2 | Cofilin-2 OS=Homo sapiens (Human) GN=CFL2 PE=1 SV=: | 95313664 | 105360544 | 91511912 | 113181792 | 94399944 | 93046152 | 90737416 | 78496120 | **0.879891135** | 2 |
| sp\|Q9Y4F1 | Q9Y4F1 | FARP1 | FERM, ARHGEF and pleckstrin domain-containing proteir | 15030720 | 16267460 | 16499938 | 16009886 | 14327083 | 12770459 | 14672968 | 14370736 | **0.879846453** | 0 |
| sp\|P62917 | P62917 | RPL8 | 60S ribosomal protein L8 OS=Homo sapiens (Human) GN | 224307296 | 199924544 | 243645664 | 195700336 | 226379456 | 167344960 | 186544672 | 179368016 | **0.879639413** | 3 |
| sp\|Q96P1: | Q96P11 | NSUN5 | Probable 28S rRNA (cytosine-C(5))-methyltransferase OS | 1148783 | 1271794.25 | 1613080 | 1306012.75 | 1027775 | 1127849.5 | 1315101.38 | 1226001.38 | **0.879591297** | 4 |
| sp\|P15848 | P15848 | ARSB | Arylsulfatase B OS=Homo sapiens (Human) GN=ARSB PE: | 4376384 | 5428423 | 4428280.5 | 5330271 | 4753641.5 | 4507221.5 | 2500869.75 | 5440262.5 | **0.879296633** | 8 |
| sp\|O7519( | O75190 | DNAJB6 | DnaJ homolog subfamily B member 6 OS=Homo sapiens | 1425900.12 | 1138786.12 | 1706143.75 | 1745277.75 | 1755726.38 | 1312888.12 | 1655385.75 | 565331.38 | **0.879194964** | 7 |
| sp\|O7561( | O75616 | ERAL1 | GTPase Era, mitochondrial OS=Homo sapiens (Human) G | 5518898.5 | 4098618 | 6210709.5 | 6793635 | 6249470.5 | 4579741 | 4144716.75 | 4914392.5 | **0.879163777** | 6 |
| sp\|Q1282 | Q12824 | SMARCB1 | SWI/SNF-related matrix-associated actin-dependent regu | 2513167.25 | 3117820.75 | 2876703.75 | 2604746.75 | 2565055.25 | 2316542.75 | 2199423 | 2686249.25 | **0.878949319** | 3 |
| sp\|P15880 | P15880 | RPS2 | 40S ribosomal protein S2 OS=Homo sapiens (Human) GN | 205067600 | 209944624 | 237281824 | 217564816 | 197701008 | 189804400 | 188028128 | 188896016 | **0.878797221** | 0 |
| sp\|P61254 | P61254 | RPL26 | 60S ribosomal protein L26 OS=Homo sapiens (Human) G | 33591104 | 62729500 | 64390040 | 43797996 | 63901808 | 44663144 | 38990836 | 32074244 | **0.878349355** | 6 |
| sp\|P35658 | P35658 | NUP214 | Nuclear pore complex protein Nup214 OS=Homo sapien | 4606264 | 4377155.5 | 4506170 | 3674249 | 3688474.75 | 3870890.75 | 4199536 | 3315041.5 | **0.878238455** | 3 |
| sp\|O1492! | O14925 | TIMM23 | Mitochondrial import inner membrane translocase subu | 1772115.75 | 1643045 | 2109361.75 | 1811987.88 | 1523048.25 | 1660159.12 | 1672029.62 | 1586101.12 | **0.877983916** | 2 |
| sp\|P62241 | P62241 | RPS8 | 40S ribosomal protein S8 OS=Homo sapiens (Human) GN | 185906752 | 188012048 | 219675328 | 218192560 | 188256192 | 169965248 | 170666304 | 183773264 | **0.877891962** | 2 |
| sp\|Q1415 | Q14157 | UBAP2L | Ubiquitin-associated protein 2-like OS=Homo sapiens (H | 9198056 | 7587633.5 | 9656160 | 8602646 | 8146878 | 7228089 | 8044799 | 7337870 | **0.877673813** | 2 |
| sp\|Q0875: | Q08752 | PPID | Peptidyl-prolyl cis-trans isomerase D OS=Homo sapiens ( | 10817828 | 9637292 | 11946551 | 11662112 | 10091155 | 10404283 | 10754028 | 7421619.5 | **0.877616103** | 3 |
| sp\|Q0370 | Q03701 | CEBPZ | CCAAT/enhancer-binding protein zeta OS=Homo sapiens | 6142437.5 | 5340105.5 | 6242797.5 | 4832512.5 | 4539717 | 6106883 | 4475378 | 4666214 | **0.877219654** | 2 |
| sp\|Q1344: | Q13443 | ADAM9 | Disintegrin and metalloproteinase domain-containing pr | 667563.19 | 555601.25 | 488050.75 | 482163.84 | 389480.09 | 539163.12 | 475932.5 | 518845.06 | **0.876921291** | 4 |
| sp\|Q1372 | Q13724 | MOGS | Mannosyl-oligosaccharide glucosidase OS=Homo sapien | 36138200 | 35738532 | 29814176 | 33124228 | 28229330 | 31283848 | 28490326 | 30161502 | **0.876496583** | 2 |
| sp\|P84098 | P84098 | RPL19 | 60S ribosomal protein L19 OS=Homo sapiens (Human) G | 55706248 | 58125440 | 93318480 | 55767280 | 67147192 | 50027504 | 48474532 | 64769828 | **0.876393171** | 6 |

| | | | | | | | | | | | | |
|---|---|---|---|---|---|---|---|---|---|---|---|---|
| sp\|Q6AI08 | Q6AI08 | HEATR6 | HEAT repeat-containing protein 6 OS=Homo sapiens (Hu | 0 | 382763.88 | 508020.59 | 238810.7 | 299245.75 | 215582.27 | 195688.22 | 279233.16 | **0.876198329** | 6 |
| sp\|P26599 | P26599 | PTBP1 | Polypyrimidine tract-binding protein 1 OS=Homo sapien | 187571008 | 173706704 | 173094656 | 204954944 | 158131120 | 175234576 | 166367888 | 147685616 | **0.875686843** | 2 |
| sp\|Q9Y2Z | Q9Y2Z0 | SUGT1 | Protein SGT1 homolog OS=Homo sapiens (Human) GN=S | 11358162 | 11856210 | 14319730 | 9606948 | 11956059 | 9453793 | 8691251 | 11175106 | **0.875589513** | 4 |
| sp\|Q9Y6N | Q9Y6N1 | COX11 | Cytochrome c oxidase assembly protein COX11, mitocho | 2514679.5 | 1809128.88 | 1880076.88 | 2252575 | 2117583.75 | 2097029.62 | 1499754 | 1683409 | **0.874807679** | 4 |
| sp\|Q1504 | Q15042 | RAB3GAP1 | Rab3 GTPase-activating protein catalytic subunit OS=Hor | 345585.19 | 329904 | 272551.59 | 381016.34 | 229622.02 | 294310.09 | 298865.88 | 339729.91 | **0.874701232** | 4 |
| sp\|Q9BQP | Q9BQP7 | MGME1 | Mitochondrial genome maintenance exonuclease 1 OS= | 7184719 | 8233440.5 | 6402271 | 8069551 | 4380516 | 8038653 | 8139389 | 5582844.5 | **0.874587443** | 5 |
| sp\|Q0520 | Q05209 | PTPN12 | Tyrosine-protein phosphatase non-receptor type 12 OS= | 154656.08 | 144095.58 | 83887.21 | 143766.44 | 108699.59 | 148322.52 | 113337.57 | 89861.91 | **0.874272317** | 6 |
| sp\|Q1320 | Q13200 | PSMD2 | 26S proteasome non-ATPase regulatory subunit 2 OS=Hc | 62791488 | 62844836 | 70352104 | 61907860 | 58113104 | 49075348 | 58971308 | 59278916 | **0.87414471** | 0 |
| sp\|Q9BRJ | Q9BRJ6 | C7orf50 | Uncharacterized protein C7orf50 OS=Homo sapiens (Hur | 7926343 | 7371969 | 6554282 | 6194963 | 6930355.5 | 5785896 | 6591714 | 5205925 | **0.874011612** | 4 |
| sp\|O4314 | O43143 | DHX15 | Pre-mRNA-splicing factor ATP-dependent RNA helicase D | 45245964 | 54146088 | 49992764 | 50970468 | 41702040 | 44178332 | 42050984 | 47165136 | **0.873929993** | 1 |
| sp\|Q1416 | Q14166 | TTLL12 | Tubulin--tyrosine ligase-like protein 12 OS=Homo sapien | 23144500 | 21703320 | 19771650 | 21340462 | 19008170 | 16062522 | 21221842 | 18829130 | **0.873914884** | 1 |
| sp\|P14324 | P14324 | FDPS | Farnesyl pyrophosphate synthase OS=Homo sapiens (Hu | 34375696 | 36097372 | 40697836 | 36527880 | 30906842 | 29894134 | 33989396 | 34108548 | **0.872714836** | 0 |
| sp\|Q9NY6 | Q9NY61 | AATF | Protein AATF OS=Homo sapiens (Human) GN=AATF PE=1 | 5843683.5 | 5587668 | 4495292.5 | 5697685.5 | 4623776 | 4474826.5 | 4600887.5 | 5167287.5 | **0.872479191** | 3 |
| sp\|P22234 | P22234 | PAICS | Multifunctional protein ADE2 OS=Homo sapiens (Humar | 128265880 | 129227744 | 142482720 | 119471288 | 125556824 | 124651904 | 90830656 | 112042512 | **0.87223787** | 2 |
| sp\|Q14651 | Q14651 | PLS1 | Plastin-1 OS=Homo sapiens (Human) GN=PLS1 PE=1 SV= | 12050602 | 10268841 | 9901947 | 9753508 | 9367973 | 9211040 | 8627760 | 9397108 | **0.872042167** | 0 |
| sp\|P01009 | P01009 | SERPINA1 | Alpha-1-antitrypsin OS=Homo sapiens (Human) GN=SERI | 17513736 | 18831216 | 30123794 | 18823466 | 22467092 | 10224383 | 19025696 | 22605136 | **0.871384447** | 7 |
| sp\|Q9279 | Q92797 | SYMPK | Symplekin OS=Homo sapiens (Human) GN=SYMPK PE=1 | 10475085 | 10335622 | 7787343 | 8928955 | 6776649 | 7711907 | 8487200 | 9724414 | **0.871377026** | 3 |
| sp\|P63172 | P63172 | DYNLT1 | Dynein light chain Tctex-type 1 OS=Homo sapiens (Huma | 2471295 | 2101117.75 | 1672840.12 | 2075301.25 | 1645340.5 | 1971542 | 1840109 | 1787511.38 | **0.870675531** | 3 |
| sp\|Q9Y2G | Q9Y2G5 | POFUT2 | GDP-fucose protein O-fucosyltransferase 2 OS=Homo sa | 606915.56 | 527164.62 | 510299.28 | 535565.38 | 415390.03 | 416928.5 | 537457.69 | 527704.62 | **0.870426079** | 5 |
| sp\|Q1334 | Q13347 | EIF3I | Eukaryotic translation initiation factor 3 subunit I OS=Ho | 24142602 | 29614740 | 24149240 | 23488084 | 23041206 | 22569260 | 20390342 | 22245852 | **0.870328425** | 0 |
| sp\|O4324 | O43242 | PSMD3 | 26S proteasome non-ATPase regulatory subunit 3 OS=Hc | 40279244 | 39460900 | 42437268 | 39570156 | 32319440 | 39224976 | 31862386 | 37267096 | **0.869712601** | 0 |
| sp\|P43686 | P43686 | PSMC4 | 26S proteasome regulatory subunit 6B OS=Homo sapien | 22545788 | 22633000 | 22819180 | 24058194 | 18936882 | 22770880 | 20281218 | 17955384 | **0.868430339** | 2 |
| sp\|Q9BVL | Q9BVL2 | NUP58 | Nucleoporin p58/p45 OS=Homo sapiens (Human) GN=NI | 439729.16 | 521223.59 | 0 | 420718.12 | 0 | 0 | 690622.94 | 509155.44 | **0.868353242** | 8 |
| sp\|P48147 | P48147 | PREP | Prolyl endopeptidase OS=Homo sapiens (Human) GN=PF | 11542617 | 12629561 | 12112469 | 11124916 | 9985484 | 11385985 | 7935913.5 | 11850150 | **0.868127228** | 3 |
| sp\|P19971 | P19971 | TYMP | Thymidine phosphorylase OS=Homo sapiens (Human) GI | 311639.47 | 834768 | 232239.31 | 413244.31 | 394269.62 | 408446.06 | 417635.75 | 334810.84 | **0.867888835** | 7 |
| sp\|Q0231 | Q02318 | CYP27A1 | Sterol 26-hydroxylase, mitochondrial OS=Homo sapiens | 19485856 | 16286425 | 16498492 | 22351360 | 15643088 | 17768320 | 12304731 | 19037154 | **0.867749157** | 4 |
| sp\|Q66K7 | Q66K74 | MAP1S | Microtubule-associated protein 1S OS=Homo sapiens (H | 1699462.5 | 1549673.5 | 736025.12 | 2037668.88 | 0 | 1722365.75 | 1900704.12 | 1602555 | **0.867636123** | 8 |
| sp\|P49368 | P49368 | CCT3 | T-complex protein 1 subunit gamma OS=Homo sapiens ( | 173425616 | 183729168 | 216940528 | 175327968 | 159825936 | 155381984 | 159262288 | 175712112 | **0.867576892** | 2 |
| sp\|P62244 | P62244 | RPS15A | 40S ribosomal protein S15a OS=Homo sapiens (Human) | 110112832 | 105334704 | 111060576 | 110870512 | 106952624 | 99391608 | 84588832 | 88487840 | **0.867488449** | 1 |
| sp\|Q9UIG | Q9UIG0 | BAZ1B | Tyrosine-protein kinase BAZ1B OS=Homo sapiens (Huma | 4137273 | 2661602.25 | 3770078.75 | 3748424.25 | 2797334.25 | 2905397.75 | 3473033.25 | 3243863.75 | **0.867451344** | 4 |
| sp\|P29372 | P29372 | MPG | DNA-3-methyladenine glycosylase OS=Homo sapiens (Hu | 815047.88 | 921965.69 | 997906.38 | 1161919.12 | 852939.25 | 878451.06 | 770574.19 | 878170.31 | **0.86740426** | 3 |
| sp\|O0074 | O00743 | PPP6C | Serine/threonine-protein phosphatase 6 catalytic subuni | 3312594 | 2861622 | 2490283.5 | 2218404.75 | 2505602.75 | 2789885.25 | 2042151.12 | 2095774.5 | **0.866810311** | 4 |
| sp\|P68032 | P68032 | ACTC1 | Actin, alpha cardiac muscle 1 OS=Homo sapiens (Human | 2775437056 | 3380167168 | 3855241216 | 2985291520 | 2389935360 | 3043584256 | 2923911424 | 2904670208 | **0.866572989** | 4 |
| sp\|P30519 | P30519 | HMOX2 | Heme oxygenase 2 OS=Homo sapiens (Human) GN=HM( | 2158598 | 2204190.75 | 1735461.62 | 2168271.5 | 1715633.25 | 2125143.25 | 1740918.75 | 1578834.75 | **0.866208317** | 2 |
| sp\|Q1309 | Q13098 | GPS1 | COP9 signalosome complex subunit 1 OS=Homo sapiens | 5169309 | 5050875.5 | 4826273 | 3495989.25 | 3504863.5 | 4732485 | 3112659.5 | 4710394.5 | **0.866142571** | 3 |
| sp\|P11021 | P11021 | HSPA5 | Endoplasmic reticulum chaperone BiP OS=Homo sapiens | 1337861760 | 1095786368 | 1506477824 | 986931264 | 1095208064 | 1015897344 | 1095759744 | 1060590912 | **0.866126752** | 4 |
| sp\|Q9BV2 | Q9BV20 | MRI1 | Methylthioribose-1-phosphate isomerase OS=Homo sap | 868404.38 | 1019374.56 | 1310116.75 | 1170075 | 715482.56 | 1115561.25 | 1026429.25 | 924720.19 | **0.865892543** | 5 |
| sp\|Q9Y6W | Q9Y6W3 | CAPN7 | Calpain-7 OS=Homo sapiens (Human) GN=CAPN7 PE=1 S | 1853966.12 | 2107001 | 1934873 | 2256586 | 2486170.5 | 2163609.5 | 1165741.75 | 1242985.5 | **0.86581677** | 7 |
| sp\|Q9UBV | Q9UBV2 | SEL1L | Protein sel-1 homolog 1 OS=Homo sapiens (Human) GN= | 3175768.25 | 2504451.25 | 2580812.25 | 2837259.75 | 2748036.25 | 2357958.25 | 2318115.75 | 2181926.5 | **0.865541939** | 2 |
| sp\|Q86X5 | Q86X55 | CARM1 | Histone-arginine methyltransferase CARM1 OS=Homo sa | 13004795 | 12032572 | 12019221 | 9549663 | 11858404 | 9994655 | 9636942 | 8848202 | **0.865510573** | 3 |
| sp\|Q8N76 | Q8N766 | EMC1 | ER membrane protein complex subunit 1 OS=Homo sapi | 12339024 | 11642097 | 12384750 | 12518458 | 10857203 | 10335780 | 11139733 | 9960232 | **0.865163721** | 0 |
| sp\|P12004 | P12004 | PCNA | Proliferating cell nuclear antigen OS=Homo sapiens (Hur | 133265504 | 158308736 | 198368112 | 151196192 | 147092112 | 126885984 | 137474560 | 143218960 | **0.865135346** | 3 |
| sp\|P08865 | P08865 | RPSA | 40S ribosomal protein SA OS=Homo sapiens (Human) GN | 244981344 | 239673568 | 308394784 | 250870080 | 236041040 | 219748848 | 230158320 | 217174048 | **0.865126111** | 0 |
| sp\|Q9NW | Q9NW13 | RBM28 | RNA-binding protein 28 OS=Homo sapiens (Human) GN= | 4830250.5 | 4425956 | 6214788 | 4230375 | 4447447 | 3463228 | 3582031 | 5549112 | **0.86500677** | 5 |
| sp\|P52895 | P52895 | AKR1C2 | Aldo-keto reductase family 1 member C2 OS=Homo sapi | 468265376 | 433634592 | 463287424 | 443049504 | 427015712 | 380699360 | 468560896 | 287172512 | **0.864625915** | 4 |

| | | | | | | | | | | | | | |
|---|---|---|---|---|---|---|---|---|---|---|---|---|---|
| sp\|Q9983 | Q99832 | CCT7 | T-complex protein 1 subunit eta OS=Homo sapiens (Hum | 181621120 | 191424016 | 225798336 | 188652528 | 177480416 | 165438496 | 170460352 | 167509360 | 0.864624867 | 0 |
| sp\|B5ME1 | B5ME19 | EIF3CL | Eukaryotic translation initiation factor 3 subunit C-like pr | 50223740 | 59983476 | 63787708 | 58318068 | 54494728 | 52591064 | 42995492 | 50739104 | 0.864438903 | 3 |
| sp\|Q9982 | Q99828 | CIB1 | Calcium and integrin-binding protein 1 OS=Homo sapien | 289663.56 | 225163.09 | 309279.78 | 272852.72 | 259661.52 | 242040.17 | 205548.52 | 240882.34 | 0.864328038 | 3 |
| sp\|P54727 | P54727 | RAD23B | UV excision repair protein RAD23 homolog B OS=Homo | 22876402 | 21898378 | 27463822 | 24276308 | 22298752 | 19858424 | 21961190 | 19284028 | 0.864139997 | 2 |
| sp\|Q1670 | Q16706 | MAN2A1 | Alpha-mannosidase 2 OS=Homo sapiens (Human) GN=M | 4479574.5 | 4860985.5 | 4257376 | 4417665.5 | 3671764.75 | 3746810 | 3806570.25 | 4342309 | 0.864109589 | 1 |
| sp\|O9543 | O95433 | AHSA1 | Activator of 90 kDa heat shock protein ATPase homolog | 28572988 | 29003464 | 28166676 | 28271634 | 25640718 | 26991998 | 22549804 | 23292992 | 0.863708438 | 0 |
| sp\|P6282 | P62820 | RAB1A | Ras-related protein Rab-1A OS=Homo sapiens (Human) | 64060488 | 77395520 | 72528984 | 72275664 | 73061680 | 64421568 | 52749064 | 56929672 | 0.863415837 | 4 |
| sp\|Q9NRZ | Q9NRZ7 | AGPAT3 | 1-acyl-sn-glycerol-3-phosphate acyltransferase gamma | 691469.31 | 662936.06 | 1086710 | 876807.5 | 724907.25 | 679175.12 | 585004.38 | 875633.06 | 0.863407596 | 5 |
| sp\|P68363 | P68363 | TUBA1B | Tubulin alpha-1B chain OS=Homo sapiens (Human) GN= | 1397653248 | 1626600320 | 1771606912 | 1707002496 | 1372903040 | 1296894976 | 1352892544 | 1587266432 | 0.862690328 | 1 |
| sp\|O1465 | O14656 | TOR1A | Torsin-1A OS=Homo sapiens (Human) GN=TOR1A PE=1 S | 112036.06 | 132399.36 | 66262.59 | 190698.09 | 101034.67 | 133735.5 | 83802.81 | 113916.2 | 0.862569892 | 7 |
| sp\|P10515 | P10515 | DLAT | Dihydrolipoyllysine-residue acetyltransferase componen | 18349064 | 18738282 | 19099644 | 20207682 | 13492180 | 15977386 | 18589586 | 17834302 | 0.862539916 | 1 |
| sp\|P43034 | P43034 | PAFAH1B1 | Platelet-activating factor acetylhydrolase IB subunit alph | 4622964 | 6228000 | 5097531.5 | 5968573 | 4698533.5 | 5158005 | 3934219.75 | 5113473.5 | 0.862534684 | 5 |
| sp\|Q9266 | Q92665 | MRPS31 | 28S ribosomal protein S31, mitochondrial OS=Homo sap | 4691076 | 4418028 | 3502826.75 | 5389793.5 | 2920312.5 | 4067657 | 4029063.25 | 4493933.5 | 0.86163781 | 4 |
| sp\|Q1558 | Q15582 | TGFBI | Transforming growth factor-beta-induced protein ig-h3 | 97324.63 | 69631.78 | 77000.71 | 112687.49 | 54917.07 | 89855.19 | 93369.43 | 69149.59 | 0.86161762 | 4 |
| sp\|P82933 | P82933 | MRPS9 | 28S ribosomal protein S9, mitochondrial OS=Homo sapie | 5435452 | 5701078.5 | 5493202.5 | 6517293 | 4766284 | 4206426 | 5064040 | 5902768 | 0.861428937 | 3 |
| sp\|P21266 | P21266 | GSTM3 | Glutathione S-transferase Mu 3 OS=Homo sapiens (Hum | 11794921 | 13873197 | 11849441 | 13095062 | 9862942 | 11789147 | 10367899 | 11542311 | 0.860700318 | 0 |
| sp\|P6228 | P62280 | RPS11 | 40S ribosomal protein S11 OS=Homo sapiens (Human) G | 211099312 | 242271696 | 288163648 | 232337920 | 221405200 | 196140416 | 207011296 | 213544304 | 0.860586114 | 2 |
| sp\|P07384 | P07384 | CAPN1 | Calpain-1 catalytic subunit OS=Homo sapiens (Human) G | 18833530 | 18254726 | 17490802 | 17596874 | 13989126 | 14049669 | 16203235 | 17867376 | 0.860527939 | 2 |
| sp\|Q1330 | Q13308 | PTK7 | Inactive tyrosine-protein kinase 7 OS=Homo sapiens (Hu | 8084359 | 5637132 | 7615752.5 | 6801165 | 8016620 | 7279290.5 | 3146830.25 | 5765688 | 0.860333972 | 6 |
| sp\|Q7L1Q | Q7L1Q6 | BZW1 | Basic leucine zipper and W2 domain-containing protein 1 | 25438120 | 26165680 | 33321746 | 26267312 | 25640062 | 22761602 | 19609336 | 27621242 | 0.860057415 | 4 |
| sp\|P26373 | P26373 | RPL13 | 60S ribosomal protein L13 OS=Homo sapiens (Human) G | 203224848 | 185020192 | 266068144 | 183892208 | 186631568 | 170792320 | 183522384 | 179809504 | 0.85987967 | 2 |
| sp\|Q9UPN | Q9UPN3 | MACF1 | Microtubule-actin cross-linking factor 1, isoforms 1/2/3/ | 2759336.25 | 3277816 | 2804827 | 5575593.5 | 4140617.5 | 3034972.25 | 3555713 | 1665869.5 | 0.85986542 | 8 |
| sp\|O0042 | O00429 | DNM1L | Dynamin-1-like protein OS=Homo sapiens (Human) GN= | 14522029 | 15951432 | 15801374 | 16120155 | 12077616 | 13114965 | 13240654 | 15189863 | 0.85941352 | 1 |
| sp\|Q1424 | Q14247 | CTTN | Src substrate cortactin OS=Homo sapiens (Human) GN= | 28301340 | 26674970 | 28777264 | 26183286 | 24638920 | 21460000 | 26310312 | 22048750 | 0.859202109 | 1 |
| sp\|Q9UKS | Q9UKS6 | PACSIN3 | Protein kinase C and casein kinase substrate in neurons | 7100297.5 | 6322806.5 | 5945589 | 6370331 | 5504343 | 5279002.5 | 5285409 | 6043593 | 0.859098134 | 1 |
| sp\|P19784 | P19784 | CSNK2A2 | Casein kinase II subunit alpha' OS=Homo sapiens (Huma | 657422.38 | 1500606.25 | 1062923.38 | 1342486 | 933289.88 | 1580918.25 | 552976.44 | 852583.69 | 0.858950697 | 6 |
| sp\|Q1580 | Q15800 | MSMO1 | Methylsterol monooxygenase 1 OS=Homo sapiens (Hum | 1024201 | 1314271.12 | 1393539.88 | 1514485.5 | 1139017.12 | 1380365.75 | 799633.06 | 1186751.88 | 0.85881444 | 4 |
| sp\|P42704 | P42704 | LRPPRC | Leucine-rich PPR motif-containing protein, mitochondria | 290146880 | 310710144 | 330055104 | 322609440 | 275429856 | 251320592 | 264295840 | 285301280 | 0.858658994 | 0 |
| sp\|Q8WU | Q8WU76 | SCFD2 | Sec1 family domain-containing protein 2 OS=Homo sapie | 533222.62 | 383537.5 | 578203.75 | 485822.97 | 410974.03 | 568813.69 | 369081.62 | 351439.31 | 0.858400619 | 4 |
| sp\|Q9NW | Q9NWH9 | SLTM | SAFB-like transcription modulator OS=Homo sapiens (Hu | 991616.5 | 832546.81 | 892071.88 | 911636.38 | 865774.12 | 752163.75 | 801243.81 | 694722.44 | 0.858328102 | 1 |
| sp\|P28288 | P28288 | ABCD3 | ATP-binding cassette sub-family D member 3 OS=Homo | 36453496 | 36998600 | 34362328 | 34544956 | 31588466 | 33463950 | 28717436 | 28395254 | 0.858145814 | 0 |
| sp\|Q9Y4C | Q9Y4C2 | TCAF1 | TRPM8 channel-associated factor 1 OS=Homo sapiens (H | 1907727 | 2111136.25 | 2528515.75 | 2721923.25 | 2025937.25 | 1695598.75 | 2130662.5 | 2098539.5 | 0.857749352 | 4 |
| sp\|Q7L2H | Q7L2H7 | EIF3M | Eukaryotic translation initiation factor 3 subunit M OS=H | 8807625 | 9512522 | 10006371 | 10580965 | 7469925 | 9402049 | 8732973 | 7741664 | 0.857074486 | 1 |
| sp\|P31689 | P31689 | DNAJA1 | DnaJ homolog subfamily A member 1 OS=Homo sapiens | 22174568 | 17330162 | 22444366 | 20775778 | 20273070 | 18010292 | 17572872 | 15019896 | 0.856769271 | 3 |
| sp\|Q1653 | Q16537 | PPP2R5E | Serine/threonine-protein phosphatase 2A 56 kDa regulat | 3238755.5 | 3044806.5 | 2621828.75 | 2460939 | 2660312.5 | 2835168.25 | 1702581.88 | 2540048.25 | 0.856750692 | 5 |
| sp\|O7569 | O75694 | NUP155 | Nuclear pore complex protein Nup155 OS=Homo sapien | 12108680 | 11194421 | 11103927 | 12189230 | 8952943 | 10712219 | 9177852 | 11063956 | 0.856441519 | 0 |
| sp\|P27361 | P27361 | MAPK3 | Mitogen-activated protein kinase 3 OS=Homo sapiens (H | 638275.5 | 1116222.25 | 1001321.94 | 1028593.75 | 694623.56 | 571317.69 | 784867.25 | 1188654.5 | 0.856001346 | 6 |
| sp\|P62424 | P62424 | RPL7A | 60S ribosomal protein L7a OS=Homo sapiens (Human) G | 274047584 | 263890368 | 290579520 | 267685952 | 262466320 | 231662400 | 224711776 | 219469024 | 0.855962953 | 0 |
| sp\|P60709 | P60709 | ACTB | Actin, cytoplasmic 1 OS=Homo sapiens (Human) GN=AC | 3475984896 | 4308818432 | 4980549632 | 4235479808 | 3177673472 | 3760276480 | 3671382272 | 3941283072 | 0.855876621 | 3 |
| sp\|O4381 | O43818 | RRP9 | U3 small nucleolar RNA-interacting protein 2 OS=Homo | 6979982.5 | 6643371.5 | 8363436.5 | 7242584 | 5409544.5 | 5876773 | 6226909.5 | 7499317.5 | 0.855733143 | 3 |
| sp\|Q96SU | Q96SU4 | OSBPL9 | Oxysterol-binding protein-related protein 9 OS=Homo sa | 1984659.62 | 1576579.75 | 1792688.25 | 1707730.38 | 1387624.12 | 1313190 | 1586778.25 | 1753705.12 | 0.855506949 | 3 |
| sp\|Q7Z4G | Q7Z4G4 | TRMT11 | tRNA (guanine(10)-N2)-methyltransferase homolog OS= | 1333625.5 | 1273972.25 | 1317801.5 | 1560726.12 | 1157688.5 | 1130944.5 | 1081250.38 | 1318040.88 | 0.854505492 | 2 |
| sp\|Q9UBB | Q9UBB6 | NCDN | Neurochondrin OS=Homo sapiens (Human) GN=NCDN P | 271565 | 284278.25 | 355749.03 | 265472.22 | 196693.33 | 210584.94 | 285798.84 | 312624.41 | 0.854414962 | 6 |
| sp\|P27635 | P27635 | RPL10 | 60S ribosomal protein L10 OS=Homo sapiens (Human) G | 65914660 | 78244440 | 77228296 | 68558120 | 66151520 | 64804600 | 65559488 | 51121740 | 0.854082351 | 1 |
| sp\|Q1455 | Q14554 | PDIA5 | Protein disulfide-isomerase A5 OS=Homo sapiens (Huma | 1687492.62 | 1526869.75 | 867108.31 | 941098.5 | 1041202.75 | 1106161.25 | 1202456.12 | 939582.81 | 0.854025654 | 7 |

| | | | | | | | | | | | | | |
|---|---|---|---|---|---|---|---|---|---|---|---|---|---|
| sp\|Q1420 | Q14204 | DYNC1H1 | Cytoplasmic dynein 1 heavy chain 1 OS=Homo sapiens (H | 240655728 | 239126976 | 237862032 | 231478880 | 194746176 | 197494096 | 201091440 | 217157600 | **0.853934407** | 0 |
| sp\|Q9HB7 | Q9HB71 | CACYBP | Calcyclin-binding protein OS=Homo sapiens (Human) GN | 44255660 | 41604472 | 54381456 | 47311132 | 44610316 | 38554928 | 37049276 | 39882692 | **0.853611784** | 2 |
| sp\|Q9BQE | Q9BQE3 | TUBA1C | Tubulin alpha-1C chain OS=Homo sapiens (Human) GN= | 1349248256 | 1583740288 | 1726240640 | 1667018880 | 1322416640 | 1236218624 | 1304061440 | 1535048576 | **0.853230102** | 1 |
| sp\|Q6DKJ | Q6DKJ4 | NXN | Nucleoredoxin OS=Homo sapiens (Human) GN=NXN PE= | 1970292.88 | 1816861.62 | 1717790.62 | 1774811.62 | 1424313.38 | 1583321.12 | 1644803.25 | 1556851.88 | **0.852952901** | 0 |
| sp\|Q9H0C | Q9H0C8 | ILKAP | Integrin-linked kinase-associated serine/threonine phosp | 4731414.5 | 5552552.5 | 6789635 | 5696915 | 5112547.5 | 4226504.5 | 4947336 | 5134243.5 | **0.852884961** | 3 |
| sp\|Q9H77 | Q9H773 | DCTPP1 | dCTP pyrophosphatase 1 OS=Homo sapiens (Human) GN | 5776590.5 | 5016784 | 3913788 | 3985312 | 4611519.5 | 4914809 | 3254830.5 | 3159183.25 | **0.852767901** | 4 |
| sp\|P63261 | P63261 | ACTG1 | Actin, cytoplasmic 2 OS=Homo sapiens (Human) GN=ACT | 3472197888 | 4304758272 | 4977362432 | 4231772928 | 3161525248 | 3746012672 | 3653937664 | 3922569472 | **0.852700284** | 3 |
| sp\|O6050 | O60506 | SYNCRIP | Heterogeneous nuclear ribonucleoprotein Q OS=Homo s | 103559552 | 110916296 | 114218880 | 106651464 | 90468072 | 91312384 | 93746424 | 95433856 | **0.852105159** | 0 |
| sp\|P54707 | P54707 | ATP12A | Potassium-transporting ATPase alpha chain 2 OS=Homo | 15395022 | 15940920 | 14674173 | 14718093 | 15025159 | 14664478 | 9239833 | 12810465 | **0.851991796** | 2 |
| sp\|P36551 | P36551 | CPOX | Oxygen-dependent coproporphyrinogen-III oxidase, mitc | 22014780 | 21988576 | 21885908 | 21581748 | 19938668 | 19341184 | 17560812 | 17632902 | **0.851408533** | 0 |
| sp\|O7522 | O75223 | GGCT | Gamma-glutamylcyclotransferase OS=Homo sapiens (Hu | 2556251.25 | 3891090.75 | 2939693.75 | 2606122.75 | 2314500.75 | 2820853.25 | 2129244 | 2945766.75 | **0.851349105** | 5 |
| sp\|O4326 | O43264 | ZW10 | Centromere/kinetochore protein zw10 homolog OS=Hor | 4362004 | 4003377 | 4352239.5 | 3496438.5 | 3615069.5 | 4059966.25 | 3078884.5 | 3046424 | **0.851134454** | 3 |
| sp\|P30419 | P30419 | NMT1 | Glycylpeptide N-tetradecanoyltransferase 1 OS=Homo sa | 15670342 | 14400212 | 13980639 | 14626876 | 12076200 | 14097209 | 11617791 | 12115133 | **0.850510827** | 1 |
| sp\|Q0108 | Q01082 | SPTBN1 | Spectrin beta chain, non-erythrocytic 1 OS=Homo sapier | 428147936 | 468422752 | 463083520 | 400470080 | 375425120 | 368017536 | 388515392 | 364716064 | **0.85032297** | 0 |
| sp\|P27695 | P27695 | APEX1 | DNA-(apurinic or apyrimidinic site) lyase OS=Homo sapie | 58563084 | 60696072 | 62110648 | 58030552 | 53377228 | 44459816 | 57796784 | 47920584 | **0.850267792** | 0 |
| sp\|Q9NTN | Q9NTN3 | SLC35D1 | UDP-glucuronic acid/UDP-N-acetylgalactosamine transpo | 157996.05 | 92079.25 | 80691.86 | 232329.28 | 130925.2 | 115906.48 | 152331.44 | 79541.86 | **0.850129651** | 6 |
| sp\|Q9Y5K | Q9Y5K5 | UCHL5 | Ubiquitin carboxyl-terminal hydrolase isozyme L5 OS=Hc | 11423443 | 9342676 | 9379038 | 9572643 | 9885520 | 8171643 | 7367822.5 | 8329234 | **0.849851188** | 3 |
| sp\|Q96T3 | Q96T37 | RBM15 | RNA-binding protein 15 OS=Homo sapiens (Human) GN= | 4342438.5 | 3987288.5 | 4843603.5 | 9818713 | 10251334 | 3238384.5 | 3352951.5 | 2695387 | **0.849774706** | 4 |
| sp\|P52756 | P52756 | RBM5 | RNA-binding protein 5 OS=Homo sapiens (Human) GN=R | 672447.94 | 598221.25 | 647609 | 782769.5 | 535339.38 | 117641.71 | 772279.69 | 869959.88 | **0.849751994** | 7 |
| sp\|P46776 | P46776 | RPL27A | 60S ribosomal protein L27a OS=Homo sapiens (Human) | 113314568 | 102285024 | 148161456 | 123756856 | 111062280 | 98254752 | 100105144 | 104691264 | **0.84943227** | 2 |
| sp\|P28370 | P28370 | SMARCA1 | Probable global transcription activator SNF2L1 OS=Homo | 11227887 | 8640569 | 11261910 | 11630383 | 8684621 | 9170024 | 10284918 | 8173411.5 | **0.849212779** | 3 |
| sp\|Q8TEQ | Q8TEQ6 | GEMIN5 | Gem-associated protein 5 OS=Homo sapiens (Human) GI | 9198179 | 7482072 | 9158092 | 8363798 | 6290560 | 8495509 | 6050622.5 | 8205285 | **0.849127442** | 3 |
| sp\|P53618 | P53618 | COPB1 | Coatomer subunit beta OS=Homo sapiens (Human) GN= | 83661840 | 92792656 | 86113096 | 75296984 | 64956052 | 74199424 | 65499616 | 82234128 | **0.849124887** | 1 |
| sp\|Q9H0A | Q9H0A0 | NAT10 | RNA cytidine acetyltransferase OS=Homo sapiens (Huma | 30962330 | 33376230 | 33538814 | 30228022 | 27038848 | 26475902 | 27922028 | 27259794 | **0.848493314** | 0 |
| sp\|O4361 | O43617 | TRAPPC3 | Trafficking protein particle complex subunit 3 OS=Homo | 1868464.62 | 1635208.62 | 1890137.75 | 1988160.25 | 1637627.88 | 1204317.25 | 1579364.38 | 1841237.5 | **0.848357005** | 2 |
| sp\|Q96HW | Q96HW7 | INTS4 | Integrator complex subunit 4 OS=Homo sapiens (Human | 834688.38 | 820242.94 | 778934.62 | 795923.38 | 900715.56 | 614978.88 | 559069.94 | 663501.88 | **0.847815752** | 4 |
| sp\|Q1501 | Q15019 | SEPT2 | Septin-2 OS=Homo sapiens (Human) GN=SEPT2 PE=1 SV= | 28946102 | 22935496 | 25977552 | 25662770 | 23763156 | 22874266 | 22099686 | 19003476 | **0.847555609** | 1 |
| sp\|P61962 | P61962 | DCAF7 | DDB1- and CUL4-associated factor 7 OS=Homo sapiens ( | 2201288.75 | 2497748 | 2502344 | 2018549.75 | 1927353.12 | 1962949.75 | 1563320.12 | 2350835.75 | **0.84647696** | 2 |
| sp\|Q9H0S | Q9H0S4 | DDX47 | Probable ATP-dependent RNA helicase DDX47 OS=Homo | 11731758 | 10459206 | 11845148 | 11122182 | 10107843 | 9881802 | 9115729 | 9116048 | **0.846387643** | 0 |
| sp\|P36507 | P36507 | MAP2K2 | Dual specificity mitogen-activated protein kinase kinase | 21002092 | 18925198 | 22616734 | 21160746 | 21118746 | 17318206 | 17171974 | 15214536 | **0.846110228** | 2 |
| sp\|P42566 | P42566 | EPS15 | Epidermal growth factor receptor substrate 15 OS=Hom | 388260.94 | 768475 | 650970.31 | 626069.5 | 403552.38 | 377382.03 | 681022.25 | 597234.56 | **0.846089135** | 5 |
| sp\|O1542 | O15427 | SLC16A3 | Monocarboxylate transporter 4 OS=Homo sapiens (Hum | 36273944 | 39351340 | 41629340 | 40083192 | 36820020 | 41100380 | 23708232 | 31395944 | **0.845471098** | 4 |
| sp\|P50570 | P50570 | DNM2 | Dynamin-2 OS=Homo sapiens (Human) GN=DNM2 PE=1 | 21151596 | 19104964 | 21943778 | 20462800 | 15528935 | 20511478 | 16892458 | 16946474 | **0.845350741** | 2 |
| sp\|P39023 | P39023 | RPL3 | 60S ribosomal protein L3 OS=Homo sapiens (Human) GN | 128696912 | 130784448 | 134435072 | 138651808 | 116858424 | 122242256 | 106490872 | 104589456 | **0.845302018** | 0 |
| sp\|P18077 | P18077 | RPL35A | 60S ribosomal protein L35a OS=Homo sapiens (Human) | 34911112 | 38579648 | 39372040 | 43367656 | 33935104 | 33413726 | 31413528 | 33257044 | **0.84502987** | 0 |
| sp\|P11940 | P11940 | PABPC1 | Polyadenylate-binding protein 1 OS=Homo sapiens (Hun | 171461648 | 174996000 | 221869936 | 176351472 | 164193968 | 160606688 | 154862704 | 149604832 | **0.845019323** | 0 |
| sp\|P07947 | P07947 | YES1 | Tyrosine-protein kinase Yes OS=Homo sapiens (Human) | 9683576 | 9409619 | 10980876 | 8931671 | 9499566 | 7629257.5 | 7585264.5 | 8216047.5 | **0.844238151** | 2 |
| sp\|Q9BQL | Q9BQL6 | FERMT1 | Fermitin family homolog 1 OS=Homo sapiens (Human) G | 1301247 | 1357143.75 | 1350371.88 | 1150061.88 | 1272586.75 | 954209.75 | 1045300.06 | 1082880.88 | **0.844180187** | 1 |
| sp\|P18621 | P18621 | RPL17 | 60S ribosomal protein L17 OS=Homo sapiens (Human) G | 80035816 | 51055320 | 59695012 | 49956920 | 56420916 | 55897412 | 45358700 | 45532828 | **0.844094319** | 4 |
| sp\|P35269 | P35269 | GTF2F1 | General transcription factor IIF subunit 1 OS=Homo sapi | 1006629.38 | 988267.62 | 1008164.69 | 938053.44 | 851439.62 | 826042.31 | 862237.19 | 785361.19 | **0.843690225** | 0 |
| sp\|Q71U3 | Q71U36 | TUBA1A | Tubulin alpha-1A chain OS=Homo sapiens (Human) GN= | 1252111360 | 1483764224 | 1599840768 | 1561764736 | 1235835776 | 1133651072 | 1210538880 | 1394192896 | **0.843447999** | 1 |
| sp\|P11172 | P11172 | UMPS | Uridine 5'-monophosphate synthase OS=Homo sapiens ( | 12831994 | 17220872 | 16494385 | 14470810 | 12194356 | 11582234 | 12741810 | 14944908 | **0.843411068** | 2 |
| sp\|Q9NV5 | Q9NV56 | MRGBP | MRG/MORF4L-binding protein OS=Homo sapiens (Huma | 1073956.75 | 0 | 0 | 531668.12 | 0 | 419883.19 | 0 | 934278.94 | **0.84338637** | 7 |
| sp\|Q1312 | Q13126 | MTAP | S-methyl-5'-thioadenosine phosphorylase OS=Homo sap | 12663474 | 12569873 | 11854998 | 13603259 | 10480485 | 11032192 | 10950688 | 10286503 | **0.84333232** | 0 |
| sp\|Q96LI5 | Q96LI5 | CNOT6L | CCR4-NOT transcription complex subunit 6-like OS=Hom | 1036388.88 | 1053628.25 | 908095.62 | 890371.06 | 926888.88 | 860425 | 668218.19 | 821489.38 | **0.842750442** | 2 |

| | | | | | | | | | | | | | |
|---|---|---|---|---|---|---|---|---|---|---|---|---|---|
| sp\|O14874 | O14874 | BCKDK | [3-methyl-2-oxobutanoate dehydrogenase [lipoamide]] k | 1236843.38 | 1136701.62 | 1426498.25 | 1290492 | 1203733.5 | 1092939.5 | 975031.5 | 1015508.5 | **0.84219297** | 1 |
| sp\|P46087 | P46087 | NOP2 | Probable 28S rRNA (cytosine(4447)-C(5))-methyltransfera | 39992968 | 44684488 | 45419468 | 37732588 | 36857416 | 32387912 | 37389472 | 34703664 | **0.842155008** | 0 |
| sp\|P55081 | P55081 | MFAP1 | Microfibrillar-associated protein 1 OS=Homo sapiens (Hu | 1415924.25 | 1224570 | 1909699.88 | 1462051.88 | 1643135.75 | 0 | 1504471.12 | 1914977.5 | **0.842045446** | 6 |
| sp\|Q9NXJ5 | Q9NXJ5 | PGPEP1 | Pyroglutamyl-peptidase 1 OS=Homo sapiens (Human) GN | 200573.59 | 210381.45 | 93983.27 | 239298.05 | 128514.14 | 216625.53 | 138662.48 | 142710.86 | **0.841819943** | 6 |
| sp\|P62701 | P62701 | RPS4X | 40S ribosomal protein S4, X isoform OS=Homo sapiens (H | 138679376 | 140540480 | 143324432 | 135508624 | 128074904 | 119024600 | 114782256 | 107835672 | **0.841707698** | 0 |
| sp\|Q8IZL8 | Q8IZL8 | PELP1 | Proline-, glutamic acid- and leucine-rich protein 1 OS=Ho | 14079022 | 14376972 | 13853670 | 12648531 | 11072985 | 11109105 | 10664140 | 13407339 | **0.841613685** | 1 |
| sp\|Q9Y4P3 | Q9Y4P3 | TBL2 | Transducin beta-like protein 2 OS=Homo sapiens (Huma | 8455015 | 8420003 | 7020111.5 | 7742599 | 5951404 | 6941927 | 6471211 | 7260496 | **0.841559722** | 1 |
| sp\|P15927 | P15927 | RPA2 | Replication protein A 32 kDa subunit OS=Homo sapiens ( | 2497419.5 | 1234194.25 | 1124644 | 1098538.25 | 978105.38 | 1801331.88 | 938820.5 | 1292531 | **0.841471103** | 6 |
| sp\|P46976 | P46976 | GYG1 | Glycogenin-1 OS=Homo sapiens (Human) GN=GYG1 PE=1 | 3235655 | 2999270 | 3813665 | 2873080.75 | 3109930.5 | 2527677 | 2488470 | 2741672.5 | **0.841048361** | 2 |
| sp\|Q06787 | Q06787 | FMR1 | Synaptic functional regulator FMR1 OS=Homo sapiens (H | 1092670.25 | 970751.25 | 866333.12 | 1160565.38 | 889345.06 | 1088224.38 | 785676.5 | 676775.88 | **0.841015329** | 3 |
| sp\|P27816 | P27816 | MAP4 | Microtubule-associated protein 4 OS=Homo sapiens (Hu | 42506108 | 46690080 | 50005120 | 39289112 | 39234140 | 26322762 | 46754784 | 37771244 | **0.840845856** | 3 |
| sp\|P49662 | P49662 | CASP4 | Caspase-4 OS=Homo sapiens (Human) GN=CASP4 PE=1 S | 214574.48 | 240999.02 | 222734.83 | 206946.69 | 138971.47 | 225503.44 | 169625.05 | 210229.16 | **0.840807568** | 4 |
| sp\|Q8N4Q | Q8N4Q0 | ZADH2 | Prostaglandin reductase 3 OS=Homo sapiens (Human) G | 686615.69 | 800630.44 | 994844 | 725317.75 | 1014929.19 | 938982.06 | 0 | 742052.75 | **0.840542925** | 7 |
| sp\|P99999 | P99999 | CYCS | Cytochrome c OS=Homo sapiens (Human) GN=CYCS PE=1 | 14012147 | 18022102 | 19459130 | 27260760 | 17422896 | 15372840 | 19240480 | 14153024 | **0.840454113** | 5 |
| sp\|A6NHR | A6NHR9 | SMCHD1 | Structural maintenance of chromosomes flexible hinge d | 9082833 | 8899691 | 9135584 | 9213219 | 5696533 | 7984082 | 8648396 | 8189848.5 | **0.840014996** | 0 |
| sp\|Q96NL6 | Q96NL6 | SCLT1 | Sodium channel and clathrin linker 1 OS=Homo sapiens ( | 9801677 | 11607399 | 9412562 | 8252802 | 10978216 | 7284341.5 | 7464427.5 | 7095605 | **0.840001546** | 3 |
| sp\|O00170 | O00170 | AIP | AH receptor-interacting protein OS=Homo sapiens (Hum | 4626880 | 4876096.5 | 3688871 | 5634282.5 | 4585746.5 | 3879714.25 | 3764887.5 | 3582055 | **0.839917883** | 3 |
| sp\|P50914 | P50914 | RPL14 | 60S ribosomal protein L14 OS=Homo sapiens (Human) G | 80575536 | 84893584 | 89052080 | 80419352 | 73979192 | 68669680 | 71006256 | 67596896 | **0.839707292** | 0 |
| sp\|Q93052 | Q93052 | LPP | Lipoma-preferred partner OS=Homo sapiens (Human) GN | 1664890 | 1454763.5 | 1739421.5 | 1137090.88 | 1491507.12 | 1291254.75 | 1808839.38 | 442298.56 | **0.839519772** | 7 |
| sp\|P35251 | P35251 | RFC1 | Replication factor C subunit 1 OS=Homo sapiens (Human | 4311137.5 | 3789961.75 | 4569709.5 | 4426961 | 3844171 | 3587138.25 | 3506080 | 3410464.75 | **0.839165237** | 1 |
| sp\|Q969V3 | Q969V3 | NCLN | Nicalin OS=Homo sapiens (Human) GN=NCLN PE=1 SV=2 | 7724414 | 7056467.5 | 7264822.5 | 6493529.5 | 5938582.5 | 5401884 | 6693593 | 5914052 | **0.839129457** | 1 |
| sp\|P60842 | P60842 | EIF4A1 | Eukaryotic initiation factor 4A-I OS=Homo sapiens (Huma | 206985072 | 247007536 | 254138192 | 244593600 | 200282416 | 181993648 | 196927312 | 220181008 | **0.839051025** | 1 |
| sp\|O95219 | O95219 | SNX4 | Sorting nexin-4 OS=Homo sapiens (Human) GN=SNX4 PE | 1645869.25 | 2795958.5 | 1674538.75 | 2007721.25 | 1841677.75 | 1521849.75 | 1634910.88 | 1817118.12 | **0.838931916** | 4 |
| sp\|Q16658 | Q16658 | FSCN1 | Fascin OS=Homo sapiens (Human) GN=FSCN1 PE=1 SV=3 | 30741070 | 32260600 | 34327392 | 33743004 | 29060174 | 30444614 | 23995938 | 26452538 | **0.838876409** | 0 |
| sp\|O14773 | O14773 | TPP1 | Tripeptidyl-peptidase 1 OS=Homo sapiens (Human) GN=T | 23942180 | 23234432 | 17957916 | 20879750 | 18138764 | 18182796 | 16386685 | 19439728 | **0.83879066** | 3 |
| sp\|P07992 | P07992 | ERCC1 | DNA excision repair protein ERCC-1 OS=Homo sapiens (H | 134795.36 | 145210.14 | 124694.98 | 148568.06 | 107254.42 | 111687.62 | 114762.91 | 130330.26 | **0.83871606** | 1 |
| sp\|P05386 | P05386 | RPLP1 | 60S acidic ribosomal protein P1 OS=Homo sapiens (Hum | 21269024 | 17689356 | 17777534 | 18465812 | 16177649 | 13474716 | 17800564 | 15614184 | **0.838639169** | 2 |
| sp\|Q99873 | Q99873 | PRMT1 | Protein arginine N-methyltransferase 1 OS=Homo sapien | 33662312 | 40635108 | 44271420 | 38094372 | 29692478 | 30208874 | 29655834 | 41815180 | **0.838565508** | 3 |
| sp\|Q9NUC | Q9NUQ9 | FAM49B | Protein FAM49B OS=Homo sapiens (Human) GN=FAM49 | 2960909.25 | 1848374.25 | 2906750.5 | 2755678.75 | 2202440.25 | 2371879.5 | 2073938.25 | 2131129.5 | **0.838390788** | 4 |
| sp\|Q86U42 | Q86U42 | PABPN1 | Polyadenylate-binding protein 2 OS=Homo sapiens (Hum | 12754799 | 10842412 | 14396548 | 12785362 | 11008996 | 9350195 | 11447886 | 10758581 | **0.838251178** | 2 |
| sp\|Q9UHD | Q9UHD8 | SEPT9 | Septin-9 OS=Homo sapiens (Human) GN=SEPT9 PE=1 SV= | 55807492 | 61782280 | 73753496 | 62423028 | 52817900 | 46498492 | 61102192 | 52214172 | **0.837907789** | 1 |
| sp\|P78310 | P78310 | CXADR | Coxsackievirus and adenovirus receptor OS=Homo sapie | 4847731.5 | 5330680 | 4730253 | 5760323 | 4394840 | 4275071 | 4307776 | 4331965.5 | **0.837469784** | 0 |
| sp\|Q02543 | Q02543 | RPL18A | 60S ribosomal protein L18a OS=Homo sapiens (Human) ( | 77716312 | 82018656 | 95893696 | 85814960 | 71954736 | 68935920 | 69423080 | 75460448 | **0.836958619** | 0 |
| sp\|Q9NQ8 | Q9NQ88 | TIGAR | Fructose-2,6-bisphosphatase TIGAR OS=Homo sapiens (H | 4447220 | 4945592 | 5914832 | 5238071 | 4098221 | 4685076 | 4047766.5 | 4351104 | **0.836289586** | 1 |
| sp\|O95292 | O95292 | VAPB | Vesicle-associated membrane protein-associated protein | 16413952 | 14619954 | 18351248 | 19303506 | 17389688 | 13167426 | 13425259 | 13448973 | **0.836111026** | 2 |
| sp\|Q5T6V5 | Q5T6V5 | C9orf64 | Queuosine salvage protein OS=Homo sapiens (Human) G | 6261425.5 | 6341251.5 | 5617648 | 6735773 | 5752204.5 | 5040653.5 | 5499542.5 | 4563581 | **0.835706828** | 1 |
| sp\|P12955 | P12955 | PEPD | Xaa-Pro dipeptidase OS=Homo sapiens (Human) GN=PEP | 30482752 | 33118672 | 32325958 | 27029880 | 26258896 | 22521550 | 28363490 | 25586284 | **0.835495345** | 1 |
| sp\|Q7Z406 | Q7Z406 | MYH14 | Myosin-14 OS=Homo sapiens (Human) GN=MYH14 PE=1 | 34050356 | 29480972 | 32276930 | 27991276 | 23527606 | 26313258 | 27697626 | 25892730 | **0.835473419** | 0 |
| sp\|Q9Y3F4 | Q9Y3F4 | STRAP | Serine-threonine kinase receptor-associated protein OS= | 22671800 | 23136136 | 23867264 | 22921056 | 20563678 | 19775386 | 19156722 | 17865712 | **0.835471123** | 0 |
| sp\|Q9NRG | Q9NRG9 | AAAS | Aladin OS=Homo sapiens (Human) GN=AAAS PE=1 SV=1 | 22112014 | 25646500 | 29128080 | 39176084 | 25220320 | 20034118 | 22322358 | 29372784 | **0.835320894** | 5 |
| sp\|Q9NXF1 | Q9NXF1 | TEX10 | Testis-expressed protein 10 OS=Homo sapiens (Human) | 4373321.5 | 4244315.5 | 3386547.25 | 4780278 | 3365560 | 3885099 | 3380947.5 | 3388122.5 | **0.835280201** | 2 |
| sp\|Q13769 | Q13769 | THOC5 | THO complex subunit 5 homolog OS=Homo sapiens (Hur | 667385.5 | 531871.56 | 108145.7 | 625630.38 | 313400.47 | 605313.25 | 356667.19 | 338864.03 | **0.835083945** | 5 |
| sp\|Q7KZ85 | Q7KZ85 | SUPT6H | Transcription elongation factor SPT6 OS=Homo sapiens ( | 9432315 | 12324832 | 10349345 | 10340950 | 9090549 | 8216643.5 | 8791314 | 9348359 | **0.835076599** | 0 |
| sp\|Q7L2E3 | Q7L2E3 | DHX30 | ATP-dependent RNA helicase DHX30 OS=Homo sapiens ( | 13586786 | 11182710 | 15337368 | 12516144 | 11861474 | 9149616 | 9438067 | 13458565 | **0.834382596** | 3 |
| sp\|O15523 | O15523 | DDX3Y | ATP-dependent RNA helicase DDX3Y OS=Homo sapiens ( | 116265448 | 124751136 | 138589664 | 123941160 | 114586656 | 102702160 | 102753496 | 99836904 | **0.833842473** | 0 |

| | | | | | | | | | | | | | |
|---|---|---|---|---|---|---|---|---|---|---|---|---|---|
| sp\|O1512 | O15127 | SCAMP2 | Secretory carrier-associated membrane protein 2 OS=Hc | 1646368.38 | 1816672.25 | 1850116 | 1918084.12 | 1615803.62 | 1681662.88 | 1438124.5 | 1289190.62 | **0.833160149** | 1 |
| sp\|Q0702 | Q07020 | RPL18 | 60S ribosomal protein L18 OS=Homo sapiens (Human) G | 217081744 | 209485920 | 252304624 | 204302176 | 187195696 | 188322432 | 171231136 | 189008992 | **0.833083707** | 0 |
| sp\|Q9NXG | Q9NXG2 | THUMPD1 | THUMP domain-containing protein 1 OS=Homo sapiens | 5849348 | 6410818.5 | 6850174 | 6748559 | 4825407.5 | 4615246 | 6940611 | 5151477.5 | **0.832701407** | 4 |
| sp\|Q1499 | Q14999 | CUL7 | Cullin-7 OS=Homo sapiens (Human) GN=CUL7 PE=1 SV=2 | 258719.92 | 193321.38 | 612095.81 | 471380.19 | 152728.7 | 193575.52 | 194895.34 | 736839.38 | **0.832318164** | 6 |
| sp\|Q9UJX2 | Q9UJX2 | CDC23 | Cell division cycle protein 23 homolog OS=Homo sapiens | 552373.94 | 713413.94 | 826673.56 | 694441.5 | 527813.94 | 553060.19 | 593005.25 | 645295.19 | **0.832169121** | 3 |
| sp\|Q1618 | Q16186 | ADRM1 | Proteasomal ubiquitin receptor ADRM1 OS=Homo sapie | 23703928 | 26433742 | 27764254 | 23411318 | 23014358 | 20978430 | 18466092 | 21837122 | **0.832033408** | 0 |
| sp\|Q1424 | Q14240 | EIF4A2 | Eukaryotic initiation factor 4A-II OS=Homo sapiens (Hum | 151757696 | 188857328 | 206584960 | 183286304 | 154872112 | 135849312 | 152096208 | 164530848 | **0.831430364** | 3 |
| sp\|P49643 | P49643 | PRIM2 | DNA primase large subunit OS=Homo sapiens (Human) G | 1262308.5 | 1433231.25 | 1576102 | 1624176.88 | 1298023 | 1331998.75 | 1157806.5 | 1113894.38 | **0.831389657** | 2 |
| sp\|Q9264 | Q92643 | PIGK | GPI-anchor transamidase OS=Homo sapiens (Human) GN | 1194858.75 | 1203693.5 | 1273911.62 | 1078773.25 | 1030144.38 | 1151920.38 | 812381.06 | 953687.69 | **0.830969579** | 1 |
| sp\|Q9946 | Q99460 | PSMD1 | 26S proteasome non-ATPase regulatory subunit 1 OS=Hc | 58676312 | 68994584 | 69330112 | 65000576 | 48821220 | 59337816 | 53953732 | 55536048 | **0.830715649** | 1 |
| sp\|P35749 | P35749 | MYH11 | Myosin-11 OS=Homo sapiens (Human) GN=MYH11 PE=1 | 40050520 | 35292136 | 38999736 | 30752976 | 28203228 | 30218670 | 31157660 | 30940360 | **0.830625537** | 2 |
| sp\|Q1401 | Q14011 | CIRBP | Cold-inducible RNA-binding protein OS=Homo sapiens (H | 6016021 | 6510456 | 9571612 | 8680160 | 5584768 | 6245707 | 7170251.5 | 6549903.5 | **0.830152164** | 5 |
| sp\|O7590 | O75909 | CCNK | Cyclin-K OS=Homo sapiens (Human) GN=CCNK PE=1 SV= | 1252527.25 | 1218465.62 | 1066226.38 | 716437.69 | 940679.69 | 1169452.38 | 775980.88 | 643320.56 | **0.829740988** | 4 |
| sp\|O9595 | O95954 | FTCD | Formimidoyltransferase-cyclodeaminase OS=Homo sapi | 7216027 | 7260831 | 8612286 | 8583309 | 5695198 | 7073178 | 5172844 | 8337823 | **0.829712905** | 2 |
| sp\|O0057 | O00571 | DDX3X | ATP-dependent RNA helicase DDX3X OS=Homo sapiens ( | 149644080 | 155698848 | 173807232 | 157649232 | 143786192 | 127809216 | 129722008 | 127035272 | **0.829700365** | 0 |
| sp\|Q9NXE | Q9NXE4 | SMPD4 | Sphingomyelin phosphodiesterase 4 OS=Homo sapiens ( | 7665642 | 1737569 | 12108688 | 2450052.25 | 2793219.5 | 7144687 | 2559888.75 | 7371160.5 | **0.829187721** | 8 |
| sp\|Q9NSD | Q9NSD9 | FARSB | Phenylalanine--tRNA ligase beta subunit OS=Homo sapie | 45698840 | 45784060 | 45783732 | 40578296 | 39747528 | 37928932 | 36581396 | 33157132 | **0.828896217** | 0 |
| sp\|Q9UQ1 | Q9UQ16 | DNM3 | Dynamin-3 OS=Homo sapiens (Human) GN=DNM3 PE=1 | 9103655 | 8828788 | 10616991 | 8842174 | 6675008 | 9483884 | 7990173 | 6842228 | **0.828830175** | 3 |
| sp\|P11171 | P11171 | EPB41 | Protein 4.1 OS=Homo sapiens (Human) GN=EPB41 PE=1 | 3669404.25 | 3090552.5 | 3199055.5 | 2989325 | 2684397 | 2358554.5 | 2896403.5 | 2791917.5 | **0.828776104** | 0 |
| sp\|Q9Y2R | Q9Y2R4 | DDX52 | Probable ATP-dependent RNA helicase DDX52 OS=Homc | 1355998.38 | 1263127.25 | 1154046.38 | 1463404.88 | 1000679 | 1388646.5 | 910859.12 | 1039388.12 | **0.8287041** | 3 |
| sp\|P17858 | P17858 | PFKL | ATP-dependent 6-phosphofructokinase, liver type OS=H | 28496380 | 29504786 | 29690608 | 29854676 | 25410798 | 27807028 | 22801442 | 21377076 | **0.828577503** | 0 |
| sp\|O7547 | O75475 | PSIP1 | PC4 and SFRS1-interacting protein OS=Homo sapiens (Hu | 3960883 | 3088521.5 | 3458910 | 3169721.25 | 3235555.5 | 3108427 | 2709533.5 | 2274201 | **0.828168401** | 3 |
| sp\|P62081 | P62081 | RPS7 | 40S ribosomal protein S7 OS=Homo sapiens (Human) GN | 136920096 | 117456600 | 130681464 | 134550160 | 102153272 | 115007688 | 110211320 | 102815088 | **0.827907005** | 0 |
| sp\|Q8N4V | Q8N4V1 | MMGT1 | Membrane magnesium transporter 1 OS=Homo sapiens | 2669283.5 | 1982077.25 | 2497893.75 | 1463576.75 | 2365820 | 1539325 | 1689564.5 | 1535692.12 | **0.827881264** | 5 |
| sp\|P42695 | P42695 | NCAPD3 | Condensin-2 complex subunit D3 OS=Homo sapiens (Hu | 494825.53 | 483332.5 | 730072.19 | 512452.19 | 410009.66 | 468040.91 | 440903.09 | 518706.31 | **0.827520388** | 3 |
| sp\|P08559 | P08559 | PDHA1 | Pyruvate dehydrogenase E1 component subunit alpha, s | 14747962 | 15029851 | 14872784 | 15456052 | 12752655 | 11972674 | 10587079 | 14412348 | **0.827275465** | 0 |
| sp\|Q9Y22 | Q9Y221 | NIP7 | 60S ribosome subunit biogenesis protein NIP7 homolog | 1173901.62 | 975668.81 | 1220053.25 | 1117319.88 | 956099.94 | 897872.31 | 889044.75 | 967430.88 | **0.826943292** | 0 |
| sp\|P28161 | P28161 | GSTM2 | Glutathione S-transferase Mu 2 OS=Homo sapiens (Hum | 4832088.5 | 5991911 | 5436814 | 5616864 | 4679542 | 4581659 | 4068376.5 | 4761991 | **0.826941914** | 0 |
| sp\|Q9H0A | Q9H0A8 | COMMD4 | COMM domain-containing protein 4 OS=Homo sapiens ( | 239949.17 | 246052.03 | 339309.94 | 247097.08 | 174841.75 | 233100.8 | 227194.84 | 251418.39 | **0.826696181** | 3 |
| sp\|Q9NVP | Q9NVP1 | DDX18 | ATP-dependent RNA helicase DDX18 OS=Homo sapiens ( | 46018400 | 49059200 | 55179948 | 45781912 | 39782792 | 41434628 | 36378176 | 44439760 | **0.826544595** | 0 |
| sp\|Q3YEC | Q3YEC7 | RABL6 | Rab-like protein 6 OS=Homo sapiens (Human) GN=RABL | 2067654.75 | 1746801 | 2035481.38 | 1966197.88 | 1323720.75 | 2120423.5 | 1463830.62 | 1551766.5 | **0.826462358** | 4 |
| sp\|Q9HD2 | Q9HD26 | GOPC | Golgi-associated PDZ and coiled-coil motif-containing prc | 1488948.12 | 1915574.38 | 1832755.75 | 2070017.5 | 1595306 | 1227393.75 | 1666334 | 1548055.75 | **0.82617287** | 3 |
| sp\|Q9UBQ | Q9UBQ7 | GRHPR | Glyoxylate reductase/hydroxypyruvate reductase OS=Hc | 21431340 | 15520260 | 23014918 | 18870696 | 17111508 | 16374894 | 21148638 | 10477488 | **0.825911073** | 4 |
| sp\|O1535 | O15355 | PPM1G | Protein phosphatase 1G OS=Homo sapiens (Human) GN= | 54025392 | 83384808 | 55610276 | 51671096 | 49295676 | 49717868 | 55540764 | 47534112 | **0.825890399** | 2 |
| sp\|Q8WX | Q8WX92 | NELFB | Negative elongation factor B OS=Homo sapiens (Human) | 4716700 | 4710984.5 | 4321206 | 3818034 | 3386213.5 | 3903067.5 | 3866646.5 | 3351085 | **0.825814018** | 2 |
| sp\|P55884 | P55884 | EIF3B | Eukaryotic translation initiation factor 3 subunit B OS=Hc | 83169928 | 87629896 | 100418304 | 85238608 | 73946488 | 76540032 | 68964344 | 74669552 | **0.82512234** | 0 |
| sp\|Q6NUC | Q6NUQ4 | TMEM214 | Transmembrane protein 214 OS=Homo sapiens (Human) | 17253834 | 12241553 | 18920656 | 17120396 | 11162565 | 15000512 | 11456498 | 16452269 | **0.825065335** | 2 |
| sp\|P29992 | P29992 | GNA11 | Guanine nucleotide-binding protein subunit alpha-11 OS | 6092282 | 5253886 | 5489826 | 4091010 | 4747988 | 5512482 | 4302742 | 2694585 | **0.824666398** | 5 |
| sp\|Q9278 | Q92785 | DPF2 | Zinc finger protein ubi-d4 OS=Homo sapiens (Human) GN | 2932354.75 | 3282933.5 | 2701408.5 | 1380848.62 | 1113628.25 | 2478555 | 2835326.5 | 2064516.75 | **0.824665121** | 4 |
| sp\|P52943 | P52943 | CRIP2 | Cysteine-rich protein 2 OS=Homo sapiens (Human) GN=( | 1386193 | 1269494.62 | 1223904.38 | 3038511 | 1337483.88 | 1053762 | 2536991.75 | 774046.94 | **0.824255518** | 5 |
| sp\|Q0420 | Q04206 | RELA | Transcription factor p65 OS=Homo sapiens (Human) GN= | 1720511.88 | 2010628.5 | 1475277.25 | 1481918.12 | 1552627.88 | 1211612.62 | 1580243.62 | 1164156 | **0.823618958** | 4 |
| sp\|P46108 | P46108 | CRK | Adapter molecule crk OS=Homo sapiens (Human) GN=CF | 7950068 | 7564741.5 | 6628328 | 6802444 | 5520227.5 | 5862449 | 6510337 | 5932582 | **0.823116837** | 0 |
| sp\|Q6PL1 | Q6PL18 | ATAD2 | ATPase family AAA domain-containing protein 2 OS=Hor | 3309695.25 | 4957085.5 | 5953618.5 | 6117288 | 4608268 | 4448165.5 | 4467551 | 3212845.25 | **0.822946559** | 3 |
| sp\|P20042 | P20042 | EIF2S2 | Eukaryotic translation initiation factor 2 subunit 2 OS=Hc | 37920956 | 31230628 | 32332284 | 30691388 | 25386882 | 30270698 | 27182516 | 25932494 | **0.822942155** | 0 |
| sp\|P17812 | P17812 | CTPS1 | CTP synthase 1 OS=Homo sapiens (Human) GN=CTPS1 PI | 16002856 | 14434942 | 20712134 | 12396230 | 14230226 | 14663075 | 12322202 | 11020803 | **0.822021415** | 3 |

| | | | | | | | | | | | | |
|---|---|---|---|---|---|---|---|---|---|---|---|---|
| sp\|Q96G0: | Q96G03 | PGM2 | Phosphoglucomutase-2 OS=Homo sapiens (Human) GN= | 4608740.5 | 4113018 | 3894562.5 | 3836637 | 3264173.5 | 4035740.75 | 3206761 | 3017916 | **0.822015789** | 2 |
| sp\|Q9UJY4 | Q9UJY4 | GGA2 | ADP-ribosylation factor-binding protein GGA2 OS=Homo | 1200020.88 | 964068.44 | 1856390.5 | 1131890.12 | 1062154.5 | 892891.56 | 1208635.38 | 1070356.5 | **0.821765127** | 5 |
| sp\|Q8N6T: | Q8N6T3 | ARFGAP1 | ADP-ribosylation factor GTPase-activating protein 1 OS=I | 9885669 | 8635814 | 8678555 | 9615895 | 8648876 | 4337646 | 9669273 | 7587663.5 | **0.821477443** | 4 |
| sp\|Q9NQV | Q9NQW6 | ANLN | Anillin OS=Homo sapiens (Human) GN=ANLN PE=1 SV=2 | 627580.38 | 0 | 1003214.69 | 855894.88 | 560120 | 645672.81 | 395155.81 | 441383.97 | **0.821305684** | 5 |
| sp\|O1498( | O14980 | XPO1 | Exportin-1 OS=Homo sapiens (Human) GN=XPO1 PE=1 S\ | 59483300 | 60040336 | 66522920 | 65046908 | 48418032 | 52220924 | 46982072 | 58535576 | **0.821035326** | 0 |
| sp\|P46379 | P46379 | BAG6 | Large proline-rich protein BAG6 OS=Homo sapiens (Hum | 10848491 | 10758056 | 13967995 | 10773980 | 7941329.5 | 8879049 | 10323086 | 10904534 | **0.820910719** | 3 |
| sp\|A3KN8: | A3KN83 | SBNO1 | Protein strawberry notch homolog 1 OS=Homo sapiens ( | 443606.91 | 350009.94 | 368345.56 | 405698.59 | 0 | 430857.09 | 468162.81 | 387397.03 | **0.820596373** | 7 |
| sp\|P61313 | P61313 | RPL15 | 60S ribosomal protein L15 OS=Homo sapiens (Human) G | 106018336 | 114289568 | 140838704 | 92036000 | 100201024 | 91394792 | 90696840 | 89515936 | **0.820438793** | 1 |
| sp\|O6050⁄ | O60504 | SORBS3 | Vinexin OS=Homo sapiens (Human) GN=SORBS3 PE=1 S\ | 2443285.5 | 1921169 | 2274841.5 | 0 | 1919602.25 | 517000.47 | 1256760.25 | 1753100.88 | **0.820337555** | 4 |
| sp\|O1516( | O15160 | POLR1C | DNA-directed RNA polymerases I and III subunit RPAC1 ( | 4717192.5 | 4197451 | 3851157 | 4187413.5 | 4143272.5 | 3216684.5 | 3704693.5 | 2837790.75 | **0.820047529** | 1 |
| sp\|P30876 | P30876 | POLR2B | DNA-directed RNA polymerase II subunit RPB2 OS=Homo | 17915256 | 18863028 | 18769488 | 16127494 | 14788898 | 14556953 | 13878788 | 15546990 | **0.819970853** | 0 |
| sp\|Q9P00( | Q9P000 | COMMD9 | COMM domain-containing protein 9 OS=Homo sapiens ( | 276771.47 | 221132.17 | 210167.77 | 284755.09 | 282208 | 241439.11 | 290287.62 | 0 | **0.819815678** | 7 |
| sp\|Q8IWA | Q8IWA4 | MFN1 | Mitofusin-1 OS=Homo sapiens (Human) GN=MFN1 PE=1 | 1412418.25 | 1458015.25 | 1639949.5 | 1780406.5 | 1446699.12 | 1322785.62 | 816179.88 | 1571536.75 | **0.819801929** | 3 |
| sp\|Q1352: | Q13523 | PRPF4B | Serine/threonine-protein kinase PRP4 homolog OS=Hom | 3913482.5 | 3161754.5 | 2123815.5 | 5823456.5 | 4377436 | 3347418.5 | 2788960 | 1799547.5 | **0.819660817** | 6 |
| sp\|P13489 | P13489 | RNH1 | Ribonuclease inhibitor OS=Homo sapiens (Human) GN=F | 55242352 | 58950144 | 56956868 | 50650080 | 47989640 | 43413820 | 47968460 | 42420256 | **0.819624129** | 0 |
| sp\|P63000 | P63000 | RAC1 | Ras-related C3 botulinum toxin substrate 1 OS=Homo sa | 17923844 | 17832778 | 17013824 | 19593150 | 17549248 | 15792795 | 11898573 | 14066651 | **0.819573242** | 1 |
| sp\|Q9Y5U! | Q9Y5U9 | IER3IP1 | Immediate early response 3-interacting protein 1 OS=Ho | 1151997.75 | 1314775.12 | 1288066.5 | 1317764.12 | 1362237.75 | 1350013.88 | 0 | 1439661.12 | **0.818497396** | 4 |
| sp\|Q1416( | Q14160 | SCRIB | Protein scribble homolog OS=Homo sapiens (Human) GN | 7019872 | 7758448.5 | 6765690 | 7171259 | 6382144 | 6199758 | 5603343 | 5293310 | **0.817633106** | 0 |
| sp\|Q9Y3E7 | Q9Y3E7 | CHMP3 | Charged multivesicular body protein 3 OS=Homo sapien: | 1784543.75 | 1710893.12 | 1674490.25 | 2244600.5 | 1574192.88 | 1525738.62 | 1612815.38 | 1341187.62 | **0.816496318** | 0 |
| sp\|Q1276! | Q12769 | NUP160 | Nuclear pore complex protein Nup160 OS=Homo sapien | 6294896 | 5847696.5 | 7812060.5 | 6178321 | 4207536.5 | 5350292 | 6015662.5 | 5759499 | **0.816324617** | 1 |
| sp\|P00491 | P00491 | PNP | Purine nucleoside phosphorylase OS=Homo sapiens (Hu | 21199856 | 22514740 | 19042864 | 20961054 | 18265148 | 17406158 | 14692381 | 17975632 | **0.816298758** | 0 |
| sp\|Q8TCG: | Q8TCG1 | CIP2A | Protein CIP2A OS=Homo sapiens (Human) GN=CIP2A PE= | 135833.09 | 108786.73 | 69474.55 | 200383.88 | 61250.88 | 184795.45 | 79746.09 | 94116.62 | **0.816184241** | 5 |
| sp\|P49327 | P49327 | FASN | Fatty acid synthase OS=Homo sapiens (Human) GN=FASI | 1085860096 | 991758400 | 1101906048 | 980856448 | 891188544 | 847301376 | 830306624 | 823857280 | **0.8154671** | 0 |
| sp\|Q6S8J3 | Q6S8J3 | POTEE | POTE ankyrin domain family member E OS=Homo sapier | 1478981376 | 1960382336 | 1910881792 | 1488003584 | 1061140992 | 1556840192 | 1667516288 | 1290459648 | **0.815407138** | 4 |
| sp\|P31949 | P31949 | S100A11 | Protein S100-A11 OS=Homo sapiens (Human) GN=S100A | 1211400.38 | 2807145.5 | 737837.62 | 1115843.25 | 1245044.12 | 1378613.12 | 1434234.75 | 729878.69 | **0.815324558** | 7 |
| sp\|O7502 | O75027 | ABCB7 | ATP-binding cassette sub-family B member 7, mitochond | 227500.33 | 194282.36 | 117064.87 | 204222.89 | 164968.28 | 186342.88 | 129792.74 | 124566.4 | **0.815091355** | 4 |
| sp\|Q9NV0 | Q9NV06 | DCAF13 | DDB1- and CUL4-associated factor 13 OS=Homo sapiens | 1308980.12 | 1879677.38 | 2108132.25 | 1496265 | 1798048.38 | 764141.62 | 1177581.88 | 1797037.38 | **0.815069135** | 4 |
| sp\|Q9266 | Q92667 | AKAP1 | A-kinase anchor protein 1, mitochondrial OS=Homo sapi | 974860.25 | 1033416.38 | 1480838.88 | 1047770 | 1064244.5 | 831826.31 | 1164767.5 | 634040.12 | **0.814408568** | 6 |
| sp\|Q9982! | Q99829 | CPNE1 | Copine-1 OS=Homo sapiens (Human) GN=CPNE1 PE=1 S\ | 35527636 | 29047576 | 29514540 | 23878390 | 26081120 | 28458372 | 22605322 | 18905520 | **0.814205703** | 2 |
| sp\|P42766 | P42766 | RPL35 | 60S ribosomal protein L35 OS=Homo sapiens (Human) G | 55399744 | 47427456 | 67187840 | 61293120 | 43263836 | 52349764 | 45396120 | 47296268 | **0.814091418** | 1 |
| sp\|O1514⁄ | O15144 | ARPC2 | Actin-related protein 2/3 complex subunit 2 OS=Homo s | 5249063.5 | 3986183 | 3224831.5 | 4944359.5 | 3967926 | 3636423.25 | 3312084.75 | 3252288 | **0.814086752** | 4 |
| sp\|Q9NX5 | Q9NX58 | LYAR | Cell growth-regulating nucleolar protein OS=Homo sapie | 8882915 | 8736120 | 7967394.5 | 8897901 | 7101971 | 6456457.5 | 6575815.5 | 7935858 | **0.813995852** | 0 |
| sp\|Q96IJ6 | Q96IJ6 | GMPPA | Mannose-1-phosphate guanyltransferase alpha OS=Hom | 8175113.5 | 5693302 | 7711556.5 | 7299249 | 6932589 | 5786640 | 5740892 | 5040597 | **0.813758723** | 3 |
| sp\|Q0034: | Q00341 | HDLBP | Vigilin OS=Homo sapiens (Human) GN=HDLBP PE=1 SV=2 | 115282048 | 122037792 | 119310472 | 116086688 | 101286712 | 87995032 | 101788032 | 93566504 | **0.813671351** | 0 |
| sp\|O1514: | O15143 | ARPC1B | Actin-related protein 2/3 complex subunit 1B OS=Homo | 27980596 | 28309152 | 27073864 | 27061584 | 19492190 | 24888814 | 22458416 | 22960952 | **0.813223569** | 0 |
| sp\|Q53H1: | Q53H12 | AGK | Acylglycerol kinase, mitochondrial OS=Homo sapiens (Hu | 5301732.5 | 4668344 | 8741955 | 5451099.5 | 4618424.5 | 5386313 | 4335956 | 5289054 | **0.812384268** | 3 |
| sp\|Q8N1F | Q8N1F7 | NUP93 | Nuclear pore complex protein Nup93 OS=Homo sapiens | 21345616 | 20587962 | 25495810 | 22632438 | 16378916 | 17096060 | 20071244 | 19614162 | **0.812335095** | 0 |
| sp\|Q5TA4! | Q5TA45 | INTS11 | Integrator complex subunit 11 OS=Homo sapiens (Huma | 6200637 | 4501682 | 5051217 | 4717192.5 | 4655601 | 4238305 | 4264858.5 | 3467676 | **0.812205609** | 1 |
| sp\|P24534 | P24534 | EEF1B2 | Elongation factor 1-beta OS=Homo sapiens (Human) GN | 77589008 | 71184608 | 83149264 | 88119624 | 64060924 | 59783064 | 80058904 | 55808836 | **0.811491364** | 2 |
| sp\|P36954 | P36954 | POLR2I | DNA-directed RNA polymerase II subunit RPB9 OS=Homo | 576205.75 | 422973.12 | 383573.12 | 508833.66 | 293962.53 | 513738.19 | 317178.59 | 409547.34 | **0.811185394** | 4 |
| sp\|Q9951! | Q99519 | NEU1 | Sialidase-1 OS=Homo sapiens (Human) GN=NEU1 PE=1 S | 19191762 | 20315410 | 18870272 | 20647120 | 15079847 | 13525011 | 17915530 | 17560786 | **0.810901962** | 0 |
| sp\|O4343 | O43432 | EIF4G3 | Eukaryotic translation initiation factor 4 gamma 3 OS=Ho | 11047253 | 11566786 | 12993974 | 10153368 | 9735808 | 8618600 | 8825332 | 9928001 | **0.810896441** | 0 |
| sp\|Q9H2J4 | Q9H2J4 | PDCL3 | Phosducin-like protein 3 OS=Homo sapiens (Human) GN | 1754134.5 | 1626086.62 | 2385243.75 | 1883688.75 | 1621960.25 | 1138947.5 | 1612622.38 | 1827147 | **0.810635717** | 2 |
| sp\|Q96GQ | Q96GQ7 | DDX27 | Probable ATP-dependent RNA helicase DDX27 OS=Homo | 26565416 | 26460074 | 27938854 | 24197162 | 18853706 | 17611040 | 24614412 | 24156128 | **0.810517928** | 1 |
| sp\|Q0287 | Q02878 | RPL6 | 60S ribosomal protein L6 OS=Homo sapiens (Human) GN | 212298960 | 216697168 | 242989024 | 209223168 | 185637232 | 152923200 | 199758528 | 175435168 | **0.809972071** | 0 |

| | | | | | | | | | | | | | |
|---|---|---|---|---|---|---|---|---|---|---|---|---|---|
| sp\|Q9Y2A | Q9Y2A7 | NCKAP1 | Nck-associated protein 1 OS=Homo sapiens (Human) GN | 2872685 | 3182829.5 | 4197542 | 2841532 | 3014408 | 3366607.5 | 1210903.62 | 3007829.5 | **0.809475504** | 7 |
| sp\|Q9996 | Q99961 | SH3GL1 | Endophilin-A2 OS=Homo sapiens (Human) GN=SH3GL1 P | 12545096 | 12882110 | 14702070 | 12247638 | 10946533 | 9726973 | 10627360 | 11093156 | **0.809402822** | 0 |
| sp\|O0042 | O00422 | SAP18 | Histone deacetylase complex subunit SAP18 OS=Homo s | 2640192.75 | 1629152.38 | 1429569.25 | 1832985.12 | 1252349.75 | 1700380 | 1849960.38 | 1288172 | **0.808675438** | 5 |
| sp\|Q9Y2Y | Q9Y2Y0 | ARL2BP | ADP-ribosylation factor-like protein 2-binding protein OS | 0 | 1030570.31 | 1397749.75 | 1654933 | 0 | 1474356.38 | 0 | 1826811 | **0.808465048** | 8 |
| sp\|O0020 | O00203 | AP3B1 | AP-3 complex subunit beta-1 OS=Homo sapiens (Human | 14712538 | 15592856 | 17408268 | 15233910 | 11972177 | 11917345 | 10855873 | 16096932 | **0.807693218** | 3 |
| sp\|P05198 | P05198 | EIF2S1 | Eukaryotic translation initiation factor 2 subunit 1 OS=Ho | 40436200 | 41777316 | 47050616 | 42765648 | 34085004 | 37101948 | 31708894 | 35961520 | **0.807170514** | 0 |
| sp\|P35613 | P35613 | BSG | Basigin OS=Homo sapiens (Human) GN=BSG PE=1 SV=2 | 80065464 | 80947424 | 73774584 | 83144440 | 64883032 | 83874496 | 52311956 | 55545868 | **0.807139335** | 4 |
| sp\|Q6P1J9 | Q6P1J9 | CDC73 | Parafibromin OS=Homo sapiens (Human) GN=CDC73 PE= | 4325520.5 | 5362488 | 5077415.5 | 5570359.5 | 3624781.5 | 4430504.5 | 4761749.5 | 3591679.75 | **0.806888766** | 2 |
| sp\|Q1363 | Q13630 | TSTA3 | GDP-L-fucose synthase OS=Homo sapiens (Human) GN= | 1798427.5 | 1986875.25 | 2014635.75 | 2363139 | 1806048.25 | 1840798.88 | 611185.62 | 2323810.25 | **0.806294317** | 5 |
| sp\|P49761 | P49761 | CLK3 | Dual specificity protein kinase CLK3 OS=Homo sapiens (H | 520547.69 | 420230.28 | 307560.88 | 386203.34 | 315150.12 | 451526.06 | 306542.66 | 244658.22 | **0.806266775** | 4 |
| sp\|Q0648 | Q06481 | APLP2 | Amyloid-like protein 2 OS=Homo sapiens (Human) GN=A | 0 | 536694.25 | 375258.53 | 824741.56 | 341253.88 | 449438.62 | 399804.31 | 209228.66 | **0.805971113** | 6 |
| sp\|Q9NX4 | Q9NX47 | MARCH5 | E3 ubiquitin-protein ligase MARCH5 OS=Homo sapiens ( | 907802.69 | 1021688.44 | 1081513.62 | 1299432.75 | 189907.25 | 286041.66 | 1215411.5 | 1781984.12 | **0.805798606** | 7 |
| sp\|Q96GV | Q96GW9 | MARS2 | Methionine--tRNA ligase, mitochondrial OS=Homo sapie | 3080574.5 | 3389281 | 2991186.25 | 2951402.5 | 2520189.5 | 2502083.5 | 2651319 | 2325859.5 | **0.805598905** | 0 |
| sp\|Q8N6H | Q8N6H7 | ARFGAP2 | ADP-ribosylation factor GTPase-activating protein 2 OS= | 3044127 | 2900967.5 | 3522923.5 | 3184250.25 | 3389534.75 | 3160328.5 | 1733242.5 | 1909246.25 | **0.805575079** | 5 |
| sp\|P07355 | P07355 | ANXA2 | Annexin A2 OS=Homo sapiens (Human) GN=ANXA2 PE=1 | 919369.5 | 9129413 | 1691203.75 | 389025.22 | 3411624.75 | 2221013.5 | 1600997.25 | 2529655.5 | **0.804953563** | 5 |
| sp\|O6048 | O60488 | ACSL4 | Long-chain-fatty-acid--CoA ligase 4 OS=Homo sapiens (H | 69853768 | 70793800 | 68476064 | 70136440 | 57659136 | 56935928 | 51668576 | 58365952 | **0.804374182** | 0 |
| sp\|Q9H7B | Q9H7B2 | RPF2 | Ribosome production factor 2 homolog OS=Homo sapie | 8307239 | 7699021 | 10077733 | 8898751 | 7037050.5 | 6985183.5 | 6926702 | 7187328.5 | **0.804289809** | 0 |
| sp\|Q8IXI1\| | Q8IXI1 | RHOT2 | Mitochondrial Rho GTPase 2 OS=Homo sapiens (Human) | 3700495 | 4527903.5 | 4907607 | 5087601 | 2279262.25 | 4445436.5 | 3709133.25 | 4201680.5 | **0.80310736** | 3 |
| sp\|Q8IXT5 | Q8IXT5 | RBM12B | RNA-binding protein 12B OS=Homo sapiens (Human) GN | 926301.38 | 887869.94 | 884818 | 880137.25 | 734310.62 | 418991.88 | 965465.81 | 754187.81 | **0.802697548** | 4 |
| sp\|Q1539 | Q15392 | DHCR24 | Delta(24)-sterol reductase OS=Homo sapiens (Human) G | 23749944 | 31154112 | 32803744 | 27567174 | 19346774 | 19559736 | 24407328 | 29202148 | **0.802567832** | 3 |
| sp\|Q8WV | Q8WVM0 | TFB1M | Dimethyladenosine transferase 1, mitochondrial OS=Ho | 13433737 | 11331779 | 12512054 | 14352581 | 10980517 | 9875474 | 10470143 | 10106827 | **0.802495445** | 0 |
| sp\|P62899 | P62899 | RPL31 | 60S ribosomal protein L31 OS=Homo sapiens (Human) G | 13017226 | 6898154 | 5276919 | 7845439 | 8118295 | 7586151.5 | 6682353 | 4117256.5 | **0.802235795** | 6 |
| sp\|Q6YP2 | Q6YP21 | KYAT3 | Kynurenine--oxoglutarate transaminase 3 OS=Homo sap | 3651274 | 2811748 | 3634412.5 | 3600446.5 | 2913663 | 2297720 | 2917698 | 2858787 | **0.802158232** | 3 |
| sp\|E9PAV | E9PAV3 | NACA | Nascent polypeptide-associated complex subunit alpha, | 29641504 | 26449020 | 31172722 | 26568932 | 26313524 | 23439532 | 18759212 | 22776036 | **0.801955173** | 0 |
| sp\|P37198 | P37198 | NUP62 | Nuclear pore glycoprotein p62 OS=Homo sapiens (Huma | 661261.44 | 618259.19 | 739712.44 | 846674.38 | 492912.53 | 635630.19 | 755435 | 411224.62 | **0.800864082** | 4 |
| sp\|P41227 | P41227 | NAA10 | N-alpha-acetyltransferase 10 OS=Homo sapiens (Human | 3194666.25 | 3378145.5 | 3459229.5 | 4084871 | 3227927.75 | 2874149.5 | 2387375.5 | 2802024.75 | **0.799854621** | 1 |
| sp\|Q96BV | Q96BW9 | TAMM41 | Phosphatidate cytidylyltransferase, mitochondrial OS=H | 1896807 | 2109524.75 | 1726200.38 | 2133589.5 | 1533429.5 | 1689574 | 1432064.5 | 1628556.62 | **0.798821187** | 0 |
| sp\|P14373 | P14373 | TRIM27 | Zinc finger protein RFP OS=Homo sapiens (Human) GN= | 868206.31 | 1069463.25 | 215897.73 | 1568764.12 | 694771.38 | 628515.94 | 825875.94 | 823575.12 | **0.798622705** | 4 |
| sp\|P29084 | P29084 | GTF2E2 | Transcription initiation factor IIE subunit beta OS=Homo | 514792.53 | 275145.91 | 247866.19 | 510188.31 | 363619.53 | 288331.19 | 334138.66 | 247387.58 | **0.796823376** | 6 |
| sp\|P62910 | P62910 | RPL32 | 60S ribosomal protein L32 OS=Homo sapiens (Human) G | 50740140 | 53358072 | 53210152 | 63307440 | 47543688 | 43705464 | 36923160 | 47569528 | **0.796596784** | 0 |
| sp\|Q9NY3 | Q9NY33 | DPP3 | Dipeptidyl peptidase 3 OS=Homo sapiens (Human) GN=D | 35779044 | 36298820 | 29108486 | 37089196 | 26380682 | 27322948 | 27107934 | 29320748 | **0.796469912** | 1 |
| sp\|Q1457 | Q14571 | ITPR2 | Inositol 1,4,5-trisphosphate receptor type 2 OS=Homo s | 5090339.5 | 5244939.5 | 2682405.75 | 5640520 | 3662785.5 | 3545500 | 4359708.5 | 3285585.75 | **0.796088367** | 4 |
| sp\|Q8N7H | Q8N7H5 | PAF1 | RNA polymerase II-associated factor 1 homolog OS=Hom | 2477944.75 | 2191090.25 | 2272737.5 | 2417208 | 2294206.25 | 1764877.75 | 1816948.25 | 1572704.88 | **0.795891938** | 2 |
| sp\|O1467 | O14672 | ADAM10 | Disintegrin and metalloproteinase domain-containing pr | 2089121 | 2259437.5 | 2650773.5 | 2432098.5 | 1916005.5 | 1566751 | 1915271 | 2108343.25 | **0.79588889** | 1 |
| sp\|Q1297 | Q12972 | PPP1R8 | Nuclear inhibitor of protein phosphatase 1 OS=Homo sa | 4603452.5 | 5147779.5 | 4682315 | 4324411.5 | 3798082 | 3482623.5 | 4440145 | 3207431.75 | **0.795837258** | 1 |
| sp\|Q8TAT | Q8TAT6 | NPLOC4 | Nuclear protein localization protein 4 homolog OS=Hom | 10577380 | 11492571 | 11491641 | 12535177 | 8462041 | 8634489 | 8448848 | 11135127 | **0.79572833** | 1 |
| sp\|Q1534 | Q15345 | LRRC41 | Leucine-rich repeat-containing protein 41 OS=Homo sap | 400886.34 | 403740.25 | 350657.94 | 366400.75 | 342385.47 | 314096.16 | 286243.41 | 268059.97 | **0.795686878** | 0 |
| sp\|Q9Y32 | Q9Y320 | TMX2 | Thioredoxin-related transmembrane protein 2 OS=Hom | 8150787 | 7825736.5 | 9913347 | 7109021 | 6859075.5 | 7052625 | 7825939 | 4514035 | **0.795532011** | 2 |
| sp\|O9545 | O95453 | PARN | Poly(A)-specific ribonuclease PARN OS=Homo sapiens (H | 115739.36 | 105172.91 | 0 | 131339.41 | 0 | 128000.53 | 72332.3 | 79811.37 | **0.795295568** | 5.5 |
| sp\|Q5T16 | Q5T160 | RARS2 | Probable arginine--tRNA ligase, mitochondrial OS=Homo | 917660.94 | 1102787.5 | 583099.12 | 991665.12 | 698161.5 | 758969.5 | 609024.62 | 789313.12 | **0.794241953** | 4 |
| sp\|Q9BZJ0 | Q9BZJ0 | CRNKL1 | Crooked neck-like protein 1 OS=Homo sapiens (Human) | 3872869.5 | 4416584 | 3907703.25 | 4295695.5 | 3460358.5 | 3504731.75 | 2868675.75 | 3264634.5 | **0.794186494** | 0 |
| sp\|P46777 | P46777 | RPL5 | 60S ribosomal protein L5 OS=Homo sapiens (Human) GN | 100635688 | 114163592 | 124499096 | 99301104 | 100104160 | 83880040 | 84644800 | 79508384 | **0.793747827** | 1 |
| sp\|Q96EP | Q96EP5 | DAZAP1 | DAZ-associated protein 1 OS=Homo sapiens (Human) GN | 8426536 | 7854692 | 7934981 | 8835307 | 3649686 | 7270833 | 8008816 | 7304651.5 | **0.793730203** | 2 |
| sp\|Q9Y4W | Q9Y4W2 | LAS1L | Ribosomal biogenesis protein LAS1L OS=Homo sapiens ( | 3250265 | 1425348.5 | 3238617.75 | 3242728 | 1394847 | 1872858.75 | 1901362 | 3685611.5 | **0.793646284** | 6 |
| sp\|P07900 | P07900 | HSP90AA1 | Heat shock protein HSP 90-alpha OS=Homo sapiens (Hu | 1275204352 | 1198563840 | 1610048000 | 1196786304 | 1163906560 | 1032594304 | 1002108288 | 987373120 | **0.792709217** | 0 |

| | | | | | | | | | | | | | |
|---|---|---|---|---|---|---|---|---|---|---|---|---|---|
| sp\|Q9Y37( | Q9Y376 | CAB39 | Calcium-binding protein 39 OS=Homo sapiens (Human) ( | 4111076.75 | 5107735 | 5224096 | 4810515.5 | 4008952 | 3420101 | 4165574.5 | 3663748 | **0.792501952** | 1 |
| sp\|P42696 | P42696 | RBM34 | RNA-binding protein 34 OS=Homo sapiens (Human) GN= | 2825433 | 2021484.62 | 2506218.75 | 2100126 | 1848443.88 | 1078929.75 | 2316809.75 | 2244713 | **0.79220232** | 4 |
| sp\|Q9Y3Z3 | Q9Y3Z3 | SAMHD1 | Deoxynucleoside triphosphate triphosphohydrolase SAM | 7674487 | 9225862 | 9975399 | 8889388 | 7905059.5 | 6452362.5 | 7259407.5 | 6713940.5 | **0.792133714** | 1 |
| sp\|P08238 | P08238 | HSP90AB1 | Heat shock protein HSP 90-beta OS=Homo sapiens (Hum | 1746678016 | 1626274688 | 2348065792 | 1763845120 | 1612052608 | 1491480448 | 1374711552 | 1450172160 | **0.79205408** | 0 |
| sp\|P49005 | P49005 | POLD2 | DNA polymerase delta subunit 2 OS=Homo sapiens (Hun | 834987.56 | 1072925.88 | 822415.38 | 994077.75 | 667469.56 | 632108.88 | 850815.69 | 796929.81 | **0.791353974** | 2 |
| sp\|Q9UKX | Q9UKX7 | NUP50 | Nuclear pore complex protein Nup50 OS=Homo sapiens | 6728471 | 7191485.5 | 6128513 | 6930268.5 | 4930165 | 5824639 | 7097447 | 3488266.25 | **0.791012435** | 3 |
| sp\|Q5JTV8 | Q5JTV8 | TOR1AIP1 | Torsin-1A-interacting protein 1 OS=Homo sapiens (Huma | 6535383.5 | 6110358 | 6057154 | 6464022 | 4451045 | 4219225 | 5611253 | 5622973 | **0.790899243** | 0 |
| sp\|Q7Z2K6 | Q7Z2K6 | ERMP1 | Endoplasmic reticulum metallopeptidase 1 OS=Homo sa | 7754561 | 7246455 | 7632805 | 7670933.5 | 6854409 | 7190868.5 | 5253836.5 | 4646536 | **0.790161491** | 0 |
| sp\|Q9BSD | Q9BSD7 | NTPCR | Cancer-related nucleoside-triphosphatase OS=Homo sap | 1509258.38 | 1952404.25 | 1435850 | 2321072.25 | 1254089.5 | 1370001.12 | 1640859.5 | 1434906.25 | **0.789608554** | 2 |
| sp\|Q9Y5Y2 | Q9Y5Y2 | NUBP2 | Cytosolic Fe-S cluster assembly factor NUBP2 OS=Homo | 6728815 | 6615088.5 | 5708794 | 6970473 | 4041923 | 7416292 | 5294232 | 3791202.25 | **0.789436831** | 4 |
| sp\|O60264 | O60264 | SMARCA5 | SWI/SNF-related matrix-associated actin-dependent regu | 10935396 | 11250508 | 11629110 | 8585286 | 7286800 | 9973727 | 7060805.5 | 9131984 | **0.788987731** | 2 |
| sp\|Q9Y2Z2 | Q9Y2Z2 | MTO1 | Protein MTO1 homolog, mitochondrial OS=Homo sapien | 472123.5 | 498972.69 | 482254.81 | 461106.22 | 330815.12 | 400273.25 | 348682.59 | 430444.06 | **0.788847619** | 0 |
| sp\|P80404 | P80404 | ABAT | 4-aminobutyrate aminotransferase, mitochondrial OS=H | 52730048 | 60739608 | 51224540 | 57820920 | 40946484 | 38447776 | 44522316 | 51416964 | **0.787962378** | 1 |
| sp\|Q9NP5 | Q9NP50 | SINHCAF | SIN3-HDAC complex-associated factor OS=Homo sapiens | 344169.38 | 253384.72 | 197154.67 | 317632.91 | 209314.91 | 224050.44 | 230482.28 | 212564.59 | **0.787898391** | 4 |
| sp\|O60841 | O60841 | EIF5B | Eukaryotic translation initiation factor 5B OS=Homo sapi | 39825664 | 35612792 | 38773680 | 31590590 | 27914764 | 27605688 | 29282568 | 30018072 | **0.787509913** | 0 |
| sp\|P51956 | P51956 | NEK3 | Serine/threonine-protein kinase Nek3 OS=Homo sapiens | 259142.86 | 1206470.88 | 1379925.5 | 913231.12 | 1101740.5 | 723443.56 | 605141.44 | 529584 | **0.7874675** | 5 |
| sp\|Q9Y3E5 | Q9Y3E5 | PTRH2 | Peptidyl-tRNA hydrolase 2, mitochondrial OS=Homo sap | 8886294 | 6755965 | 6652161 | 7446253.5 | 6352275 | 5839030 | 7539320 | 3683038.75 | **0.78726071** | 3 |
| sp\|P55854 | P55854 | SUMO3 | Small ubiquitin-related modifier 3 OS=Homo sapiens (Hu | 24098652 | 31685238 | 25572134 | 22033568 | 18903406 | 19928334 | 23254942 | 19304702 | **0.787229956** | 1 |
| sp\|P33992 | P33992 | MCM5 | DNA replication licensing factor MCM5 OS=Homo sapier | 26617662 | 24078108 | 23754590 | 19772454 | 17909322 | 18303586 | 17134812 | 20759756 | **0.786513084** | 1 |
| sp\|P51532 | P51532 | SMARCA4 | Transcription activator BRG1 OS=Homo sapiens (Human) | 15033486 | 12153002 | 19155808 | 13278401 | 10659401 | 13472977 | 12603758 | 10102969 | **0.785618206** | 3 |
| sp\|Q1333( | Q13330 | MTA1 | Metastasis-associated protein MTA1 OS=Homo sapiens ( | 5101490.5 | 6523360 | 6148890.5 | 6276306.5 | 4546056 | 3711091.5 | 4309520.5 | 6325729.5 | **0.785545122** | 3 |
| sp\|P07711 | P07711 | CTSL | Cathepsin L1 OS=Homo sapiens (Human) GN=CTSL PE=1 | 960438.75 | 578602.56 | 267879.53 | 895528.94 | 432290.84 | 577136.5 | 569762.81 | 543108 | **0.785323807** | 4 |
| sp\|Q15691 | Q15691 | MAPRE1 | Microtubule-associated protein RP/EB family member 1 | 64879444 | 65969884 | 66641040 | 67815760 | 54258880 | 49313748 | 53089828 | 51662972 | **0.785226597** | 0 |
| sp\|O75794 | O75794 | CDC123 | Cell division cycle protein 123 homolog OS=Homo sapier | 514259.06 | 585008.75 | 697831.31 | 593337.19 | 524786.94 | 399779.62 | 482340.31 | 469430.91 | **0.784935274** | 1 |
| sp\|P52298 | P52298 | NCBP2 | Nuclear cap-binding protein subunit 2 OS=Homo sapiens | 1170927.75 | 1542778.38 | 1743088.75 | 1493086.5 | 1738163.62 | 0 | 1633927.75 | 1295178.5 | **0.784430743** | 7 |
| sp\|Q9H4N | Q9H4M9 | EHD1 | EH domain-containing protein 1 OS=Homo sapiens (Hum | 8130933 | 7020326 | 9213319 | 7757482.5 | 7373206.5 | 6937480 | 5674997 | 5182951 | **0.783531134** | 1 |
| sp\|Q13867 | Q13867 | BLMH | Bleomycin hydrolase OS=Homo sapiens (Human) GN=BL | 8054806.5 | 10739966 | 7291117 | 7897765 | 5612480.5 | 8263795 | 7139235.5 | 5611008 | **0.783509584** | 3 |
| sp\|Q58FF6 | Q58FF6 | HSP90AB4 | Putative heat shock protein HSP 90-beta 4 OS=Homo sap | 143177056 | 158000960 | 248847616 | 174804848 | 142976592 | 130385536 | 143717056 | 150742096 | **0.783384937** | 2 |
| sp\|Q9H3N | Q9H3N1 | TMX1 | Thioredoxin-related transmembrane protein 1 OS=Homo | 21537034 | 23719036 | 22239056 | 22048800 | 19646498 | 17424474 | 16590779 | 16483339 | **0.783359555** | 0 |
| sp\|Q969N | Q969N2 | PIGT | GPI transamidase component PIG-T OS=Homo sapiens (H | 2466718.75 | 3051548.25 | 3412513.5 | 2919938.25 | 2353168.5 | 2923745.75 | 1767634.38 | 2237624.75 | **0.783258263** | 2 |
| sp\|Q16181 | Q16181 | SEPT7 | Septin-7 OS=Homo sapiens (Human) GN=SEPT7 PE=1 SV= | 13832454 | 18860844 | 16694698 | 19817444 | 11754531 | 15173549 | 15304215 | 11968121 | **0.783181438** | 2 |
| sp\|Q9H8Y | Q9H8Y5 | ANKZF1 | Ankyrin repeat and zinc finger domain-containing protein | 0 | 322482.53 | 168085.11 | 378758.38 | 154239.09 | 317488.88 | 208966.12 | 0 | **0.783013593** | 5.5 |
| sp\|Q9H0U | Q9H0U4 | RAB1B | Ras-related protein Rab-1B OS=Homo sapiens (Human) ( | 32193880 | 33514996 | 37981488 | 38282332 | 29349028 | 30280354 | 24588364 | 26936120 | **0.782924246** | 0 |
| sp\|P25205 | P25205 | MCM3 | DNA replication licensing factor MCM3 OS=Homo sapier | 46601672 | 46868292 | 58712564 | 47699072 | 40626660 | 39287256 | 38620996 | 37949888 | **0.782887469** | 0 |
| sp\|Q9H8H | Q9H8H0 | NOL11 | Nucleolar protein 11 OS=Homo sapiens (Human) GN=NC | 4944884 | 4572594 | 4247498 | 5044816.5 | 2917865.25 | 3901923.25 | 3081427 | 4820804.5 | **0.78267849** | 2 |
| sp\|Q8NHH | Q8NHH9 | ATL2 | Atlastin-2 OS=Homo sapiens (Human) GN=ATL2 PE=1 SV= | 4015220.5 | 3623551 | 4078366 | 4278922.5 | 3128336 | 3421003.5 | 2727254 | 3241395 | **0.782566988** | 0 |
| sp\|P53582 | P53582 | METAP1 | Methionine aminopeptidase 1 OS=Homo sapiens (Huma | 3589412 | 3932908.5 | 3418134 | 3726602 | 2471466 | 2941950 | 2966537.75 | 3081589.25 | **0.78144807** | 0 |
| sp\|Q9BY4 | Q9BY44 | EIF2A | Eukaryotic translation initiation factor 2A OS=Homo sapi | 14251810 | 15775457 | 14535354 | 14815956 | 12210053 | 11597506 | 12707363 | 9849664 | **0.780830197** | 0 |
| sp\|P41091 | P41091 | EIF2S3 | Eukaryotic translation initiation factor 2 subunit 3 OS=Hc | 105682056 | 123043840 | 130922000 | 126019656 | 112183392 | 89315024 | 97062944 | 80450552 | **0.780393729** | 1 |
| sp\|P61011 | P61011 | SRP54 | Signal recognition particle 54 kDa protein OS=Homo sapi | 15115372 | 12914662 | 11635938 | 16332779 | 12215726 | 10849254 | 10889957 | 9680651 | **0.779224308** | 1 |
| sp\|Q9Y31( | Q9Y316 | MEMO1 | Protein MEMO1 OS=Homo sapiens (Human) GN=MEMO | 443654.56 | 333918.06 | 367913.78 | 341962.75 | 285919.06 | 338776.03 | 319752.75 | 214386.25 | **0.77907476** | 1 |
| sp\|P61513 | P61513 | RPL37A | 60S ribosomal protein L37a OS=Homo sapiens (Human) ( | 15390637 | 16643804 | 19321868 | 13436378 | 13466334 | 14029942 | 12097856 | 10883500 | **0.779063724** | 2 |
| sp\|P61353 | P61353 | RPL27 | 60S ribosomal protein L27 OS=Homo sapiens (Human) G | 30312782 | 31888448 | 26730824 | 37052456 | 28383584 | 22782736 | 23988532 | 22970536 | **0.778868672** | 1 |
| sp\|Q00613 | Q00613 | HSF1 | Heat shock factor protein 1 OS=Homo sapiens (Human) ( | 223183.36 | 186111.39 | 134668.78 | 188612.25 | 103458.94 | 156631.84 | 170331.39 | 140105.78 | **0.778797178** | 3 |
| sp\|Q05682 | Q05682 | CALD1 | Caldesmon OS=Homo sapiens (Human) GN=CALD1 PE=1 | 11096465 | 11251069 | 11060078 | 13072205 | 9279454 | 9148868 | 8904537 | 8854140 | **0.778552958** | 0 |

| | | | | | | | | | | | | | |
|---|---|---|---|---|---|---|---|---|---|---|---|---|---|
| sp\|Q9297: | Q92973 | TNPO1 | Transportin-1 OS=Homo sapiens (Human) GN=TNPO1 PE | 39134320 | 41365224 | 42026288 | 37357932 | 31146530 | 29211862 | 33714380 | 30277112 | **0.777751792** | 0 |
| sp\|Q9UHI( | Q9UHI6 | DDX20 | Probable ATP-dependent RNA helicase DDX20 OS=Homc | 1437391.62 | 1229296.5 | 1446787.75 | 1243154.75 | 1241612.62 | 873257.56 | 1128697.75 | 919329.31 | **0.777148461** | 1 |
| sp\|P33993 | P33993 | MCM7 | DNA replication licensing factor MCM7 OS=Homo sapier | 40994952 | 33177840 | 36956728 | 36963284 | 28531210 | 30195908 | 27481500 | 28816850 | **0.776712068** | 0 |
| sp\|P46060 | P46060 | RANGAP1 | Ran GTPase-activating protein 1 OS=Homo sapiens (Hum | 19703654 | 18101388 | 20045532 | 19523172 | 15787931 | 14773833 | 14114688 | 15363347 | **0.775971206** | 0 |
| sp\|Q96CU | Q96CU9 | FOXRED1 | FAD-dependent oxidoreductase domain-containing prote | 1321120 | 1254879.25 | 1474816.62 | 1046768.44 | 938108.12 | 966978.12 | 957096.94 | 1090528.38 | **0.775408766** | 1 |
| sp\|Q1413 | Q14137 | BOP1 | Ribosome biogenesis protein BOP1 OS=Homo sapiens (H | 8471403 | 6890054 | 6945184 | 7626479 | 5092404.5 | 6046936 | 5214247 | 6852958.5 | **0.775279891** | 0 |
| sp\|P28332 | P28332 | ADH6 | Alcohol dehydrogenase 6 OS=Homo sapiens (Human) GN | 3264939.5 | 3260190.5 | 3081604.5 | 2867277.75 | 1631818.25 | 2771728.75 | 2719308.5 | 2547460 | **0.775236973** | 0 |
| sp\|Q6UWI | Q6UWE0 | LRSAM1 | E3 ubiquitin-protein ligase LRSAM1 OS=Homo sapiens (H | 181555.8 | 186483.62 | 137712.8 | 193233.78 | 105515.05 | 149036.52 | 146410.34 | 140455.94 | **0.7745761** | 3 |
| sp\|Q0519: | Q05193 | DNM1 | Dynamin-1 OS=Homo sapiens (Human) GN=DNM1 PE=1 | 4286109 | 4452687 | 5434818 | 4528565 | 2589232.75 | 4563336 | 3830836.25 | 3498541 | **0.774345385** | 3 |
| sp\|Q1356 | Q13564 | NAE1 | NEDD8-activating enzyme E1 regulatory subunit OS=Hon | 3377913.75 | 4273534.5 | 3345630.25 | 2977709 | 2503128.25 | 3161344 | 2486179.75 | 2666132.75 | **0.774021412** | 1 |
| sp\|Q1304: | Q13043 | STK4 | Serine/threonine-protein kinase 4 OS=Homo sapiens (Hu | 256183.86 | 237837.78 | 421965.38 | 240696.45 | 204059.98 | 233156.12 | 219752.7 | 236472.67 | **0.772416563** | 0 |
| sp\|P48507 | P48507 | GCLM | Glutamate--cysteine ligase regulatory subunit OS=Homo | 2524352 | 2511744 | 3176304.25 | 2750784.5 | 2886238.25 | 1684232.62 | 1949982.5 | 1942330.25 | **0.771927484** | 3 |
| sp\|O7553 | O75534 | CSDE1 | Cold shock domain-containing protein E1 OS=Homo sapi | 12999393 | 14274156 | 19493680 | 15304931 | 14911782 | 10254108 | 12616544 | 10121518 | **0.771746174** | 2 |
| sp\|O6088 | O60884 | DNAJA2 | DnaJ homolog subfamily A member 2 OS=Homo sapiens | 23330714 | 22227784 | 21534626 | 21702910 | 17592316 | 17601104 | 16298975 | 17030512 | **0.771688823** | 0 |
| sp\|P08754 | P08754 | GNAI3 | Guanine nucleotide-binding protein G(k) subunit alpha C | 5453175 | 6985395 | 8032301 | 7277344 | 4781873 | 6324870 | 3303289.5 | 7000202 | **0.771589614** | 3 |
| sp\|Q1444 | Q14444 | CAPRIN1 | Caprin-1 OS=Homo sapiens (Human) GN=CAPRIN1 PE=1 | 25451188 | 25234792 | 27137376 | 21656672 | 18708038 | 21070054 | 17190598 | 19760372 | **0.77130117** | 0 |
| sp\|Q9BXB | Q9BXB5 | OSBPL10 | Oxysterol-binding protein-related protein 10 OS=Homo s | 1600668.38 | 1628411.75 | 1960496 | 1907123.5 | 1286756.75 | 1242519.88 | 1497690.25 | 1446309.75 | **0.771242537** | 0 |
| sp\|P49207 | P49207 | RPL34 | 60S ribosomal protein L34 OS=Homo sapiens (Human) G | 41167984 | 41551192 | 43289820 | 41160800 | 34734008 | 29537050 | 29952002 | 34687604 | **0.771136097** | 0 |
| sp\|P19838 | P19838 | NFKB1 | Nuclear factor NF-kappa-B p105 subunit OS=Homo sapie | 4162690.25 | 2556247.75 | 2584382.25 | 3532954.75 | 2948327 | 3208969.75 | 1988743.25 | 1752385 | **0.771129085** | 4 |
| sp\|P00492 | P00492 | HPRT1 | Hypoxanthine-guanine phosphoribosyltransferase OS=H | 8384802.5 | 7983611 | 7424119 | 10893610 | 7747084.5 | 7301609 | 6002775 | 5695953 | **0.771127014** | 1 |
| sp\|Q8NE7 | Q8NE71 | ABCF1 | ATP-binding cassette sub-family F member 1 OS=Homo s | 28721624 | 27562658 | 24104300 | 27518678 | 20270596 | 21316560 | 19413242 | 22167278 | **0.770732905** | 0 |
| sp\|Q86TG | Q86TG7 | PEG10 | Retrotransposon-derived protein PEG10 OS=Homo sapie | 26636548 | 30454232 | 44249880 | 27168516 | 28924276 | 23465164 | 26401870 | 20161888 | **0.770008812** | 2 |
| sp\|Q6PD7 | Q6PD74 | AAGAB | Alpha- and gamma-adaptin-binding protein p34 OS=Hon | 448401.09 | 435124.78 | 622187.06 | 390221.03 | 338543.41 | 397515.34 | 400792.5 | 322963.09 | **0.769971091** | 2 |
| sp\|O7538( | O75380 | NDUFS6 | NADH dehydrogenase [ubiquinone] iron-sulfur protein 6, | 1836165.5 | 1171321.75 | 0 | 1147739.88 | 1049585.88 | 639260.69 | 710170.75 | 799065.31 | **0.769652905** | 4 |
| sp\|Q9Y5B | Q9Y5B9 | SUPT16H | FACT complex subunit SPT16 OS=Homo sapiens (Human | 71592832 | 78210192 | 85825416 | 78701536 | 61064744 | 56931776 | 62562768 | 61202616 | **0.769134103** | 0 |
| sp\|Q9BQG | Q9BQG0 | MYBBP1A | Myb-binding protein 1A OS=Homo sapiens (Human) GN= | 46699360 | 46246892 | 49326560 | 45084400 | 33181160 | 35452440 | 37870748 | 37579272 | **0.769031619** | 0 |
| sp\|Q1456( | Q14566 | MCM6 | DNA replication licensing factor MCM6 OS=Homo sapier | 38352044 | 36589468 | 41264516 | 28014754 | 29101198 | 27129664 | 26594346 | 28020400 | **0.768582769** | 2 |
| sp\|Q9278( | Q92786 | PROX1 | Prospero homeobox protein 1 OS=Homo sapiens (Huma | 17164176 | 13302678 | 11917652 | 6962239.5 | 10734669 | 10897469 | 9175688 | 7117938.5 | **0.768556551** | 4 |
| sp\|P61221 | P61221 | ABCE1 | ATP-binding cassette sub-family E member 1 OS=Homo s | 32964392 | 37944908 | 39243576 | 38208216 | 32188934 | 26849572 | 25237580 | 29697904 | **0.768220215** | 0 |
| sp\|P52789 | P52789 | HK2 | Hexokinase-2 OS=Homo sapiens (Human) GN=HK2 PE=1 | 34303296 | 33501280 | 29883276 | 33417690 | 26756536 | 30120908 | 20517530 | 23222332 | **0.767452729** | 1 |
| sp\|Q9UNC | Q9UNQ2 | DIMT1 | Probable dimethyladenosine transferase OS=Homo sapi | 3795955.25 | 3995903.75 | 4652825 | 3714168 | 3700587.5 | 2931275.75 | 2834755.75 | 2932359 | **0.767318** | 0 |
| sp\|Q9H0V | Q9H0V9 | LMAN2L | VIP36-like protein OS=Homo sapiens (Human) GN=LMAN | 1154860.75 | 926528.94 | 1068103.75 | 1298669.5 | 801047.38 | 735056.88 | 1067411.25 | 808985.25 | **0.767170809** | 1 |
| sp\|Q9NVD | Q9NVD7 | PARVA | Alpha-parvin OS=Homo sapiens (Human) GN=PARVA PE= | 3678190 | 3742839.5 | 3889739 | 4110405.5 | 3042994 | 2616993 | 2867566.25 | 3298871 | **0.766895196** | 0 |
| sp\|O6023 | O60231 | DHX16 | Pre-mRNA-splicing factor ATP-dependent RNA helicase D | 1832216 | 1856927.88 | 1727902.62 | 1634811.38 | 1280455.25 | 1388051.25 | 1289611.25 | 1448893.62 | **0.766749906** | 0 |
| sp\|Q9Y6E( | Q9Y6E0 | STK24 | Serine/threonine-protein kinase 24 OS=Homo sapiens (H | 4989419 | 5302218.5 | 6068367 | 4980781.5 | 4343814 | 4968028 | 3597668.5 | 3452348.5 | **0.766694301** | 0 |
| sp\|O9557: | O95573 | ACSL3 | Long-chain-fatty-acid--CoA ligase 3 OS=Homo sapiens (H | 60535212 | 58763704 | 49182280 | 53868824 | 43974684 | 45913212 | 39975600 | 40609232 | **0.766686362** | 0 |
| sp\|Q9NR1 | Q9NR19 | ACSS2 | Acetyl-coenzyme A synthetase, cytoplasmic OS=Homo sa | 1015164.5 | 1184309.38 | 782389.56 | 1261148.25 | 835056.44 | 1028392.75 | 751579.25 | 636111.12 | **0.766233939** | 3 |
| sp\|O1478 | O14787 | TNPO2 | Transportin-2 OS=Homo sapiens (Human) GN=TNPO2 PE | 19680050 | 21764642 | 21412326 | 18693146 | 15830882 | 14755750 | 17205588 | 14690367 | **0.766185915** | 0 |
| sp\|P51665 | P51665 | PSMD7 | 26S proteasome non-ATPase regulatory subunit 7 OS=Hc | 18421650 | 20195046 | 23499654 | 17743888 | 19388870 | 14338648 | 14606288 | 12838943 | **0.765997579** | 2 |
| sp\|Q1346 | Q13464 | ROCK1 | Rho-associated protein kinase 1 OS=Homo sapiens (Hum | 3201505.75 | 3894689.25 | 4031483 | 3628713.75 | 2872486 | 2665798.5 | 3027722 | 2723873.75 | **0.765084069** | 0 |
| sp\|Q58FF8 | Q58FF8 | HSP90AB2 | Putative heat shock protein HSP 90-beta 2 OS=Homo sap | 814942464 | 777977920 | 1118193792 | 766427328 | 714933760 | 663404224 | 616447232 | 665676736 | **0.765041035** | 0 |
| sp\|O9534 | O95347 | SMC2 | Structural maintenance of chromosomes protein 2 OS=H | 7958178.5 | 7726560.5 | 7944088.5 | 7750003 | 6028378.5 | 6549645.5 | 6558934 | 4859888.5 | **0.764746363** | 0 |
| sp\|Q1311 | Q13112 | CHAF1B | Chromatin assembly factor 1 subunit B OS=Homo sapien | 2337416.25 | 1885604.12 | 2634474 | 2055442.25 | 1902031.88 | 1797603.75 | 1450073.25 | 1662017.38 | **0.764251621** | 1 |
| sp\|Q9Y28 | Q9Y287 | ITM2B | Integral membrane protein 2B OS=Homo sapiens (Huma | 423268.41 | 443866.97 | 235610.39 | 522650.12 | 243857.69 | 420956.56 | 322296.34 | 254070.02 | **0.763617416** | 4 |
| sp\|Q9P0L( | Q9P0L0 | VAPA | Vesicle-associated membrane protein-associated protein | 10416534 | 11504690 | 12301326 | 10996620 | 9675938 | 7895470 | 8511558 | 8436472 | **0.763380619** | 0 |

| | | | | | | | | | | | | | |
|---|---|---|---|---|---|---|---|---|---|---|---|---|---|
| sp\|Q8NFH | Q8NFH3 | NUP43 | Nucleoporin Nup43 OS=Homo sapiens (Human) GN=NUF | 4119081.75 | 4145258.75 | 4756777 | 4554049.5 | 3607235 | 3393943.75 | 3231047.25 | 3184237.75 | **0.763376174** | 0 |
| sp\|Q9NRX | Q9NRX1 | PNO1 | RNA-binding protein PNO1 OS=Homo sapiens (Human) G | 6240657 | 4733138 | 7522140 | 6794167 | 5154740 | 4653782 | 4928849.5 | 4559527 | **0.763021774** | 2 |
| sp\|Q9953‹ | Q99538 | LGMN | Legumain OS=Homo sapiens (Human) GN=LGMN PE=1 S | 1474387.5 | 1561609.12 | 1509843.88 | 1580666.75 | 1108656.62 | 1073337.12 | 1420390.5 | 1068495.38 | **0.762404977** | 0 |
| sp\|P23378 | P23378 | GLDC | Glycine dehydrogenase (decarboxylating), mitochondrial | 33880720 | 35770660 | 31388660 | 32643076 | 25492600 | 27303304 | 25917980 | 23055686 | **0.76127467** | 0 |
| sp\|Q9Y312 | Q9Y312 | AAR2 | Protein AAR2 homolog OS=Homo sapiens (Human) GN=/ | 1243644.38 | 0 | 1212702.12 | 3208649 | 962755.44 | 1187828 | 1065237.5 | 1094100.25 | **0.760798696** | 4 |
| sp\|Q32P2ξ | Q32P28 | P3H1 | Prolyl 3-hydroxylase 1 OS=Homo sapiens (Human) GN=P | 8230310.5 | 7323720 | 9140122 | 8455710 | 6663479 | 6401264 | 6356046.5 | 5791807.5 | **0.76056415** | 0 |
| sp\|P49792 | P49792 | RANBP2 | E3 SUMO-protein ligase RanBP2 OS=Homo sapiens (Hum | 32891236 | 27391668 | 31820700 | 25569710 | 23933500 | 18224046 | 24952042 | 22307768 | **0.759877945** | 0 |
| sp\|P52948 | P52948 | NUP98 | Nuclear pore complex protein Nup98-Nup96 OS=Homo s | 13971899 | 12282072 | 15139047 | 14102694 | 11564145 | 9819747 | 10167218 | 10598503 | **0.759511167** | 0 |
| sp\|P67870 | P67870 | CSNK2B | Casein kinase II subunit beta OS=Homo sapiens (Human) | 5333634.5 | 5290838 | 3204235.5 | 4270164.5 | 2691206.75 | 3827043 | 3750453.5 | 3476630.5 | **0.759458013** | 3 |
| sp\|G2XKQ | G2XKQ0 | SUMO1P1 | Small ubiquitin-related modifier 5 OS=Homo sapiens (Hu | 12242972 | 12038478 | 12100406 | 11086120 | 10033766 | 9125568 | 8761732 | 8126511 | **0.759408343** | 0 |
| sp\|Q9NR7 | Q9NR77 | PXMP2 | Peroxisomal membrane protein 2 OS=Homo sapiens (Hu | 514250.78 | 435178.28 | 590475.75 | 695429.62 | 447950.03 | 583135.19 | 0 | 662080.62 | **0.757455268** | 6 |
| sp\|Q9H07 | Q9H078 | CLPB | Caseinolytic peptidase B protein homolog OS=Homo sap | 1391619 | 2352579.5 | 2515058.5 | 1485664.25 | 1504798.25 | 342432.47 | 1948617.5 | 2064328.88 | **0.756647732** | 6 |
| sp\|Q9H2U | Q9H2U1 | DHX36 | ATP-dependent DNA/RNA helicase DHX36 OS=Homo sap | 1016117.19 | 1206463.88 | 1149727.12 | 1283120.25 | 740406.44 | 912070.12 | 900253.12 | 961427.38 | **0.754851483** | 0 |
| sp\|Q9UJX! | Q9UJX5 | ANAPC4 | Anaphase-promoting complex subunit 4 OS=Homo sapie | 311315.19 | 296580.72 | 370848.16 | 327796.75 | 239354.61 | 276609.22 | 247348.52 | 222739.3 | **0.754704051** | 0 |
| sp\|P62191 | P62191 | PSMC1 | 26S proteasome regulatory subunit 4 OS=Homo sapiens | 38415808 | 35315044 | 38832740 | 38393908 | 31771140 | 28774762 | 27716734 | 25651656 | **0.754611675** | 0 |
| sp\|Q9NPL | Q9NPL8 | TIMMDC1 | Complex I assembly factor TIMMDC1, mitochondrial OS= | 654698.5 | 448771.97 | 331205.62 | 430150.84 | 376856.16 | 442795.03 | 300241.34 | 286493 | **0.754164104** | 3 |
| sp\|P42677 | P42677 | RPS27 | 40S ribosomal protein S27 OS=Homo sapiens (Human) G | 13059649 | 12186841 | 8336236.5 | 14592511 | 12786658 | 9562319 | 8682707 | 5297426 | **0.754103392** | 4 |
| sp\|Q9HCS | Q9HCS7 | XAB2 | Pre-mRNA-splicing factor SYF1 OS=Homo sapiens (Huma | 2046659.25 | 2024320 | 2135505 | 2080116.25 | 1734568.38 | 2010961.75 | 1100312.62 | 1399042 | **0.753612383** | 0 |
| sp\|Q9NRN | Q9NRN7 | AASDHPPT | L-aminoadipate-semialdehyde dehydrogenase-phospho | 3542075 | 3324213.5 | 3459554.75 | 3716457.75 | 3123998 | 2204684.5 | 1996120.25 | 3247255.75 | **0.752872232** | 0 |
| sp\|P14635 | P14635 | CCNB1 | G2/mitotic-specific cyclin-B1 OS=Homo sapiens (Human) | 1374973.62 | 2175033.25 | 3905304.5 | 1673481.88 | 1594771.5 | 1752134 | 1356368.75 | 2164013.75 | **0.752266791** | 5 |
| sp\|Q9NZN | Q9NZM5 | NOP53 | Ribosome biogenesis protein NOP53 OS=Homo sapiens ( | 1357880.75 | 1311973.25 | 1056586.75 | 1445349.12 | 973939.06 | 965020.19 | 1043408 | 907664.12 | **0.752163461** | 0 |
| sp\|Q9BTC | Q9BTC0 | DIDO1 | Death-inducer obliterator 1 OS=Homo sapiens (Human) | 307105.12 | 215798.08 | 215575.73 | 162425.78 | 173438.16 | 144795.83 | 199916.22 | 159127.58 | **0.751775168** | 2 |
| sp\|Q9HAV | Q9HAV4 | XPO5 | Exportin-5 OS=Homo sapiens (Human) GN=XPO5 PE=1 S | 20240640 | 22856308 | 23998472 | 18290390 | 15036152 | 14510118 | 15724145 | 18872416 | **0.751211835** | 1 |
| sp\|Q1425ξ | Q14258 | TRIM25 | E3 ubiquitin/ISG15 ligase TRIM25 OS=Homo sapiens (Hu | 25526478 | 26178746 | 30541560 | 19255804 | 24453262 | 17197302 | 16985198 | 17505816 | **0.750144203** | 1 |
| sp\|O76021 | O76021 | RSL1D1 | Ribosomal L1 domain-containing protein 1 OS=Homo sa | 110099720 | 110734160 | 117304464 | 112502400 | 77941912 | 79383240 | 87261280 | 93412936 | **0.750041741** | 0 |
| sp\|P68104 | P68104 | EEF1A1 | Elongation factor 1-alpha 1 OS=Homo sapiens (Human) ( | 2389877760 | 2009700224 | 2346902528 | 2381638656 | 1842476544 | 1916792320 | 1490350976 | 1592463488 | **0.749561131** | 0 |
| sp\|Q9983( | Q99836 | MYD88 | Myeloid differentiation primary response protein MyD8 | 661466.88 | 1989969.25 | 1019722.19 | 1615783.12 | 1371002.75 | 733174.56 | 902334.12 | 954554.25 | **0.749216863** | 5 |
| sp\|Q0715 | Q07157 | TJP1 | Tight junction protein ZO-1 OS=Homo sapiens (Human) ( | 11028540 | 12184586 | 12317298 | 9048499 | 5286194.5 | 6758254.5 | 11707735 | 9646729 | **0.749208611** | 3 |
| sp\|Q6NYC | Q6NYC1 | JMJD6 | Bifunctional arginine demethylase and lysyl-hydroxylase | 4003608.25 | 3869001 | 3286289.25 | 3314056.75 | 2709937.5 | 2957972.25 | 2400302.25 | 2759052.25 | **0.748103208** | 0 |
| sp\|Q86XP | Q86XP3 | DDX42 | ATP-dependent RNA helicase DDX42 OS=Homo sapiens ( | 17380206 | 16669130 | 16310378 | 15275774 | 14230206 | 11806701 | 11834509 | 11210281 | **0.747792063** | 0 |
| sp\|P50613 | P50613 | CDK7 | Cyclin-dependent kinase 7 OS=Homo sapiens (Human) G | 2843366 | 2726135.5 | 3409396 | 2825995.75 | 2744036.5 | 1963669 | 2056627.25 | 2062792.5 | **0.747751383** | 1 |
| sp\|Q9297 | Q92974 | ARHGEF2 | Rho guanine nucleotide exchange factor 2 OS=Homo sa | 637824.12 | 586508.56 | 460183.53 | 436638.22 | 355399.28 | 387998.81 | 460422.72 | 381254.09 | **0.747269919** | 2 |
| sp\|Q1466 | Q14669 | TRIP12 | E3 ubiquitin-protein ligase TRIP12 OS=Homo sapiens (Hu | 10182334 | 10008964 | 11646597 | 8798170 | 6797282 | 7379336 | 7205232.5 | 8981672 | **0.747206269** | 1 |
| sp\|Q9GZN | Q9GZM8 | NDEL1 | Nuclear distribution protein nudE-like 1 OS=Homo sapie | 671478.69 | 724873.12 | 845497 | 967211.31 | 540949.25 | 685151.25 | 652802 | 518811.03 | **0.747170025** | 1 |
| sp\|Q9Y57 | Q9Y570 | PPME1 | Protein phosphatase methylesterase 1 OS=Homo sapien | 4348379.5 | 3819504.5 | 5540139.5 | 2953554.5 | 2446822 | 3403341.75 | 2277890.5 | 4308368 | **0.74641323** | 3 |
| sp\|Q5JPH | Q5JPH6 | EARS2 | Probable glutamate--tRNA ligase, mitochondrial OS=Hor | 11050217 | 8980367 | 3751853.5 | 6604604.5 | 6962646.5 | 6493416.5 | 5936589 | 3263649 | **0.745590867** | 4 |
| sp\|P49790 | P49790 | NUP153 | Nuclear pore complex protein Nup153 OS=Homo sapien | 10239994 | 9359290 | 9216814 | 8398623 | 6977963.5 | 6479873.5 | 8000741 | 6281937 | **0.745417788** | 0 |
| sp\|P51114 | P51114 | FXR1 | Fragile X mental retardation syndrome-related protein 1 | 5537924.5 | 4994291 | 6992136.5 | 5321543 | 4205873.5 | 3994166.5 | 4067726 | 4755002 | **0.745112765** | 0 |
| sp\|O6051ξ | O60518 | RANBP6 | Ran-binding protein 6 OS=Homo sapiens (Human) GN=R | 7239816.5 | 7845921.5 | 9351359 | 7614971.5 | 5606496 | 5089246.5 | 6469368.5 | 6681703.5 | **0.744002357** | 0 |
| sp\|Q9UI1( | Q9UI10 | EIF2B4 | Translation initiation factor eIF-2B subunit delta OS=Hor | 4298234 | 5929226.5 | 4089176.5 | 4914885 | 4127521 | 2954456 | 3323577 | 3881412 | **0.742893152** | 1 |
| sp\|P05534 | P05534 | HLA-A | HLA class I histocompatibility antigen, A-24 alpha chain ( | 32262212 | 27099754 | 32127826 | 29528410 | 25306858 | 22914530 | 22390072 | 19289608 | **0.742872283** | 0 |
| sp\|P16152 | P16152 | CBR1 | Carbonyl reductase [NADPH] 1 OS=Homo sapiens (Huma | 25908888 | 18207474 | 18363820 | 18546902 | 16598574 | 14452973 | 15509376 | 13622946 | **0.742762371** | 0 |
| sp\|Q96EN | Q96EN8 | MOCOS | Molybdenum cofactor sulfurase OS=Homo sapiens (Hum | 520534.72 | 407804.53 | 492270.78 | 502905.72 | 330608.31 | 397862.41 | 355630.28 | 341927.56 | **0.741365679** | 0 |
| sp\|O4329( | O43290 | SART1 | U4/U6.U5 tri-snRNP-associated protein 1 OS=Homo sapi | 2921951.5 | 2337072.5 | 2513149 | 2432429.5 | 2401587 | 615215.06 | 2810722.5 | 1736935.38 | **0.741279236** | 4 |
| sp\|P49841 | P49841 | GSK3B | Glycogen synthase kinase-3 beta OS=Homo sapiens (Hur | 426751.34 | 363088.94 | 136455.91 | 247494.84 | 241826.2 | 256739.31 | 198918.5 | 172525.3 | **0.74119608** | 5 |

| | | | | | | | | | | | | | |
|---|---|---|---|---|---|---|---|---|---|---|---|---|---|
| sp\|O1461 | O14617 | AP3D1 | AP-3 complex subunit delta-1 OS=Homo sapiens (Human | 12208131 | 11411183 | 9692545 | 7364918.5 | 4410682 | 5145563 | 12655178 | 7934029 | **0.74109735** | 5 |
| sp\|P13797 | P13797 | PLS3 | Plastin-3 OS=Homo sapiens (Human) GN=PLS3 PE=1 SV= | 33907068 | 33598540 | 29790168 | 33559796 | 24826726 | 25679074 | 23625428 | 22843134 | **0.741079348** | 0 |
| sp\|Q0894 | Q08945 | SSRP1 | FACT complex subunit SSRP1 OS=Homo sapiens (Human | 31253050 | 32244644 | 37108508 | 34109692 | 26476602 | 26493876 | 23704160 | 23012060 | **0.739977259** | 0 |
| sp\|P21333 | P21333 | FLNA | Filamin-A OS=Homo sapiens (Human) GN=FLNA PE=1 SV | 501139200 | 479343328 | 515031776 | 407681728 | 356736448 | 341492608 | 377123712 | 332581056 | **0.739773413** | 0 |
| sp\|Q5C9Z | Q5C9Z4 | NOM1 | Nucleolar MIF4G domain-containing protein 1 OS=Homo | 468581.66 | 395886.69 | 480091.31 | 358498.72 | 324142.78 | 289403.06 | 314792 | 331499.25 | **0.739749796** | 0 |
| sp\|P19367 | P19367 | HK1 | Hexokinase-1 OS=Homo sapiens (Human) GN=HK1 PE=1 | 5798330 | 3332920.5 | 5068339 | 6001227 | 4200952 | 4192356.25 | 2129668.5 | 4412715 | **0.739360795** | 3 |
| sp\|P49915 | P49915 | GMPS | GMP synthase [glutamine-hydrolyzing] OS=Homo sapier | 31399652 | 32640914 | 34031040 | 35356208 | 20959500 | 25372406 | 26950656 | 25361240 | **0.73930464** | 0 |
| sp\|P57737 | P57737 | CORO7 | Coronin-7 OS=Homo sapiens (Human) GN=CORO7 PE=1 | 564749.88 | 753357.56 | 625037.06 | 1033085.62 | 451215.22 | 513772.81 | 346064.19 | 888898.56 | **0.739173616** | 3 |
| sp\|P10319 | P10319 | HLA-B | HLA class I histocompatibility antigen, B-58 alpha chain | 18874968 | 18210420 | 17554618 | 17087276 | 15645072 | 12579072 | 13682516 | 11104592 | **0.739066789** | 0 |
| sp\|Q9980 | Q99805 | TM9SF2 | Transmembrane 9 superfamily member 2 OS=Homo sap | 10912910 | 12525535 | 12277341 | 12007759 | 8575915 | 7537819.5 | 8688237 | 10436590 | **0.738389436** | 0 |
| sp\|P78344 | P78344 | EIF4G2 | Eukaryotic translation initiation factor 4 gamma 2 OS=Hc | 39443752 | 39026616 | 44015100 | 40000592 | 32242244 | 29183228 | 28217104 | 30318722 | **0.738286706** | 0 |
| sp\|Q9BYG | Q9BYG3 | NIFK | MKI67 FHA domain-interacting nucleolar phosphoprotei | 12051695 | 11245275 | 13381156 | 12302015 | 8370722 | 9136319 | 8105798 | 10531513 | **0.737938913** | 0 |
| sp\|P42345 | P42345 | MTOR | Serine/threonine-protein kinase mTOR OS=Homo sapier | 6473963 | 6471649 | 1299576 | 5829790.5 | 3258556.25 | 3778952.75 | 3859161 | 3913991.25 | **0.737767228** | 4 |
| sp\|Q969X | Q969X6 | UTP4 | U3 small nucleolar RNA-associated protein 4 homolog O | 6388302.5 | 6646386.5 | 8478021 | 7675049 | 4947048.5 | 5192368.5 | 5531787.5 | 5861755 | **0.737739389** | 0 |
| sp\|O4371 | O43719 | HTATSF1 | HIV Tat-specific factor 1 OS=Homo sapiens (Human) GN= | 1963516.5 | 0 | 9474046 | 2161120.75 | 1853699.62 | 1989558.75 | 4688561 | 1494631.38 | **0.737310412** | 7 |
| sp\|Q9BS2 | Q9BS26 | ERP44 | Endoplasmic reticulum resident protein 44 OS=Homo sa | 5202546.5 | 6549636 | 5548327.5 | 5734913 | 5137219.5 | 3965008.5 | 3726343.25 | 4152465.25 | **0.737170596** | 0 |
| sp\|O0054 | O00541 | PES1 | Pescadillo homolog OS=Homo sapiens (Human) GN=PES | 24717098 | 25894824 | 32878626 | 27255250 | 20779018 | 18276034 | 22252020 | 20330914 | **0.73716554** | 0 |
| sp\|Q5JRA | Q5JRA6 | MIA3 | Transport and Golgi organization protein 1 homolog OS= | 1064885.38 | 923821 | 1070394.25 | 0 | 608809.44 | 846132 | 799619.38 | 0 | **0.737001195** | 3.5 |
| sp\|Q9Y6A | Q9Y6A4 | CFAP20 | Cilia- and flagella-associated protein 20 OS=Homo sapie | 9512878 | 8889578 | 10632407 | 10471621 | 8903820 | 3564285.5 | 8386494 | 8255717.5 | **0.736849095** | 1 |
| sp\|Q1432 | Q14320 | FAM50A | Protein FAM50A OS=Homo sapiens (Human) GN=FAM50 | 2464564.5 | 2406287.25 | 2015965.88 | 2276022.25 | 1901576.62 | 692407.69 | 1972136.5 | 2183412.5 | **0.73662024** | 1 |
| sp\|P28070 | P28070 | PSMB4 | Proteasome subunit beta type-4 OS=Homo sapiens (Hun | 34277988 | 35658992 | 22255444 | 24398900 | 19471244 | 24212914 | 19692792 | 22429054 | **0.73595531** | 2 |
| sp\|O1516 | O15164 | TRIM24 | Transcription intermediary factor 1-alpha OS=Homo sap | 1676369 | 2340560.75 | 1959039.5 | 1155391 | 1583931.25 | 639016.19 | 1372522.62 | 1651533.5 | **0.73576476** | 3 |
| sp\|P68400 | P68400 | CSNK2A1 | Casein kinase II subunit alpha OS=Homo sapiens (Humar | 35654200 | 36847412 | 30785508 | 31052556 | 23162220 | 26394030 | 24426068 | 24780352 | **0.735171268** | 0 |
| sp\|P02768 | P02768 | ALB | Serum albumin OS=Homo sapiens (Human) GN=ALB PE= | 101516592 | 108991080 | 140356064 | 95916016 | 112175000 | 50013884 | 85068848 | 81035736 | **0.734799342** | 3 |
| sp\|Q9NU2 | Q9NU22 | MDN1 | Midasin OS=Homo sapiens (Human) GN=MDN1 PE=1 SV | 5361800 | 4747883 | 6307622.5 | 5627035 | 3917064 | 4496390 | 3592631.5 | 4176172.75 | **0.734077676** | 0 |
| sp\|O9575 | O95757 | HSPA4L | Heat shock 70 kDa protein 4L OS=Homo sapiens (Human | 18974366 | 19919288 | 23764592 | 18759998 | 14904218 | 14197387 | 16130656 | 14510635 | **0.733777751** | 0 |
| sp\|P49736 | P49736 | MCM2 | DNA replication licensing factor MCM2 OS=Homo sapier | 21551656 | 21693140 | 22743762 | 21558796 | 17388280 | 17844762 | 13703865 | 15289428 | **0.733618231** | 0 |
| sp\|Q9NY9 | Q9NY93 | DDX56 | Probable ATP-dependent RNA helicase DDX56 OS=Homo | 6132089.5 | 6129116 | 6330537 | 6015389 | 4258348.5 | 4436848 | 4520256 | 4834295.5 | **0.733516948** | 0 |
| sp\|P0DJD | P0DJD0 | RGPD1 | RANBP2-like and GRIP domain-containing protein 1 OS= | 7805805 | 7172979.5 | 7683153 | 6584397 | 5964882 | 3259365.75 | 6361946.5 | 5864881.5 | **0.733462026** | 0 |
| sp\|O4317 | O43172 | PRPF4 | U4/U6 small nuclear ribonucleoprotein Prp4 OS=Homo s | 4926930 | 4985241.5 | 4413859.5 | 3028559 | 3164997 | 3211027 | 3403622 | 2947416.5 | **0.73335426** | 3 |
| sp\|Q9UJU | Q9UJU6 | DBNL | Drebrin-like protein OS=Homo sapiens (Human) GN=DBN | 8118702.5 | 7600706.5 | 8026273 | 7652866.5 | 6360265.5 | 7038643 | 6935586.5 | 2683719.5 | **0.733098044** | 0 |
| sp\|Q96P7 | Q96P70 | IPO9 | Importin-9 OS=Homo sapiens (Human) GN=IPO9 PE=1 S | 9510006 | 10220554 | 10761080 | 9131842 | 7538646.5 | 7729855 | 6613304 | 7146270 | **0.732597794** | 0 |
| sp\|O1529 | O15294 | OGT | UDP-N-acetylglucosamine--peptide N-acetylglucosaminy | 1251318 | 1154459 | 2028512 | 1270752.25 | 1428878.88 | 1513079.38 | 515113.5 | 721635.12 | **0.732458662** | 6 |
| sp\|Q6DKI1 | Q6DKI1 | RPL7L1 | 60S ribosomal protein L7-like 1 OS=Homo sapiens (Huma | 13170817 | 12953163 | 13711300 | 12417032 | 9603700 | 9222787 | 9908176 | 9476461 | **0.73128102** | 0 |
| sp\|Q8WW | Q8WWM7 | ATXN2L | Ataxin-2-like protein OS=Homo sapiens (Human) GN=AT | 21250236 | 17287960 | 29060650 | 20942984 | 16357461 | 15848010 | 15168652 | 17317308 | **0.730631285** | 1 |
| sp\|Q1284 | Q12849 | GRSF1 | G-rich sequence factor 1 OS=Homo sapiens (Human) GN | 4604466 | 4250210 | 3270053.5 | 4291662.5 | 2808412.75 | 2957480.75 | 2957564.75 | 3267373 | **0.730418185** | 0 |
| sp\|Q1468 | Q14684 | RRP1B | Ribosomal RNA processing protein 1 homolog B OS=Hon | 6974577 | 8248952 | 8036366.5 | 6840018 | 5254662 | 5080734.5 | 5323985 | 6321427 | **0.730261517** | 0 |
| sp\|P31153 | P31153 | MAT2A | S-adenosylmethionine synthase isoform type-2 OS=Hom | 14409679 | 13418867 | 15962286 | 11394647 | 8991863 | 9586628 | 10277577 | 11429845 | **0.730009302** | 1 |
| sp\|Q8N68 | Q8N684 | CPSF7 | Cleavage and polyadenylation specificity factor subunit 7 | 8482876 | 10261396 | 14598376 | 13606343 | 9692332 | 6438925.5 | 7553937.5 | 10511535 | **0.72838051** | 3 |
| sp\|Q9BRR | Q9BRR6 | ADPGK | ADP-dependent glucokinase OS=Homo sapiens (Human) | 1721128.88 | 1823418 | 1491410.5 | 1771907.5 | 1137269.75 | 1717526.62 | 1038393.88 | 1064638.25 | **0.728250132** | 1 |
| sp\|P04114 | P04114 | APOB | Apolipoprotein B-100 OS=Homo sapiens (Human) GN=A | 99800480 | 88058520 | 96376104 | 95673960 | 74020296 | 63405252 | 65621464 | 73550904 | **0.728063482** | 0 |
| sp\|Q9HCE | Q9HCE5 | METTL14 | N6-adenosine-methyltransferase non-catalytic subunit ( | 684196.12 | 735486.69 | 0 | 0 | 477324.53 | 0 | 0 | 555205 | **0.72729593** | 6 |
| sp\|P00352 | P00352 | ALDH1A1 | Retinal dehydrogenase 1 OS=Homo sapiens (Human) GN | 519545344 | 551261248 | 566780800 | 552090752 | 425634912 | 389986144 | 390682272 | 384961856 | **0.726711909** | 0 |
| sp\|P08962 | P08962 | CD63 | CD63 antigen OS=Homo sapiens (Human) GN=CD63 PE= | 559617.38 | 597837.06 | 702796.62 | 785900.5 | 0 | 441151.25 | 613803.94 | 866436.12 | **0.726107808** | 6 |
| sp\|Q9BQB | Q9BQB6 | VKORC1 | Vitamin K epoxide reductase complex subunit 1 OS=Hon | 3785297.5 | 4145018 | 3371576 | 2977143.5 | 2439028.25 | 2886053 | 2277651 | 2760616.75 | **0.725773766** | 0 |

| | | | | | | | | | | | | | |
|---|---|---|---|---|---|---|---|---|---|---|---|---|---|
| sp\|Q2TAA | Q2TAA2 | IAH1 | Isoamyl acetate-hydrolyzing esterase 1 homolog OS=Hor | 832441.12 | 828523.25 | 775619 | 740723.81 | 500325.34 | 667406.31 | 590637.88 | 547547.19 | **0.725745604** | 0 |
| sp\|Q1382 | Q13823 | GNL2 | Nucleolar GTP-binding protein 2 OS=Homo sapiens (Hum | 2762553 | 2982917.75 | 3020607 | 3106660.75 | 2653437.5 | 1590287.62 | 2036017.5 | 2326775.75 | **0.724897493** | 0 |
| sp\|Q0279 | Q02790 | FKBP4 | Peptidyl-prolyl cis-trans isomerase FKBP4 OS=Homo sapi | 38033808 | 34125484 | 37920208 | 31383200 | 27495226 | 25064290 | 26256380 | 23705186 | **0.724721655** | 0 |
| sp\|Q8TD3 | Q8TD30 | GPT2 | Alanine aminotransferase 2 OS=Homo sapiens (Human) | 21598154 | 18637576 | 13835724 | 21812898 | 12788005 | 14843254 | 14731316 | 12622611 | **0.72459189** | 2 |
| sp\|Q9H30 | Q9H307 | PNN | Pinin OS=Homo sapiens (Human) GN=PNN PE=1 SV=5 | 24068234 | 23258260 | 27230810 | 21739236 | 17152812 | 14955422 | 23207970 | 14453578 | **0.724530518** | 1 |
| sp\|Q9UK5 | Q9UK58 | CCNL1 | Cyclin-L1 OS=Homo sapiens (Human) GN=CCNL1 PE=1 SV | 129033.96 | 156285.83 | 114436.8 | 200587.59 | 79412.8 | 107855.48 | 115123.4 | 132397.59 | **0.724233339** | 3 |
| sp\|P33908 | P33908 | MAN1A1 | Mannosyl-oligosaccharide 1,2-alpha-mannosidase IA OS | 16795932 | 15717419 | 16148584 | 16244370 | 4861737 | 13543724 | 13949475 | 14558814 | **0.722791877** | 0 |
| sp\|P33991 | P33991 | MCM4 | DNA replication licensing factor MCM4 OS=Homo sapier | 16968404 | 15746743 | 15558346 | 15334241 | 11946651 | 11899713 | 11341028 | 10768636 | **0.722491199** | 0 |
| sp\|P46734 | P46734 | MAP2K3 | Dual specificity mitogen-activated protein kinase kinase | 4883458.5 | 5162144 | 4831616.5 | 4908671 | 3496035.25 | 3133276.75 | 4348636.5 | 3316325 | **0.72244784** | 0 |
| sp\|P30520 | P30520 | ADSS | Adenylosuccinate synthetase isozyme 2 OS=Homo sapie | 14860739 | 13958521 | 15461312 | 14532304 | 11476428 | 10493289 | 10150188 | 10362930 | **0.722339016** | 0 |
| sp\|P46940 | P46940 | IQGAP1 | Ras GTPase-activating-like protein IQGAP1 OS=Homo sap | 61369028 | 64574084 | 59734044 | 54427048 | 44918964 | 43318540 | 43657208 | 41535912 | **0.722313983** | 0 |
| sp\|O1511 | O15118 | NPC1 | NPC intracellular cholesterol transporter 1 OS=Homo sap | 1385563.12 | 1399287.5 | 834085.12 | 1234263.38 | 1277585.5 | 1787972.88 | 438937.41 | 0 | **0.722100145** | 6 |
| sp\|Q53H8 | Q53H82 | LACTB2 | Endoribonuclease LACTB2 OS=Homo sapiens (Human) G | 540591.5 | 652279.12 | 821686.38 | 760582.06 | 392314.28 | 542186.12 | 496306.03 | 572852.5 | **0.722003073** | 2 |
| sp\|Q96B2 | Q96B26 | EXOSC8 | Exosome complex component RRP43 OS=Homo sapiens | 4474467.5 | 4322572 | 4793609 | 3660479 | 2416101.5 | 2813206.25 | 1885050.12 | 5335976 | **0.721711312** | 4 |
| sp\|P22413 | P22413 | ENPP1 | Ectonucleotide pyrophosphatase/phosphodiesterase fan | 1874759.62 | 2065025.75 | 1993585.38 | 2684448 | 1450717 | 1940491.12 | 1180829.88 | 1643651 | **0.721260122** | 1 |
| sp\|O6025 | O60256 | PRPSAP2 | Phosphoribosyl pyrophosphate synthase-associated prot | 3583594.5 | 2480614 | 2695676.5 | 3219496.25 | 2342851 | 2673894 | 1573179.88 | 2046990.5 | **0.72098176** | 1 |
| sp\|Q96Q1 | Q96Q15 | SMG1 | Serine/threonine-protein kinase SMG1 OS=Homo sapien | 156664.09 | 203044.17 | 171522.27 | 208863.72 | 99338.12 | 143416.11 | 128159.48 | 162666.72 | **0.72096281** | 1 |
| sp\|O7511 | O75116 | ROCK2 | Rho-associated protein kinase 2 OS=Homo sapiens (Hum | 30152360 | 30356356 | 36257176 | 28732698 | 23601880 | 20904590 | 24119552 | 21842986 | **0.720876689** | 0 |
| sp\|Q2TAA | Q2TAA5 | ALG11 | GDP-Man:Man(3)GlcNAc(2)-PP-Dol alpha-1,2-mannosyltr | 623064.06 | 494114.94 | 621532.62 | 612772.38 | 476003.47 | 527293.44 | 691332.88 | 0 | **0.720663968** | 5 |
| sp\|P60891 | P60891 | PRPS1 | Ribose-phosphate pyrophosphokinase 1 OS=Homo sapie | 12188372 | 10125830 | 8927116 | 9738466 | 8521463 | 8478815 | 5510985 | 6979094 | **0.719631831** | 0 |
| sp\|O1532 | O15320 | CTAGE5 | Endoplasmic reticulum export factor CTAGE5 OS=Homo | 449801.5 | 451128.53 | 563486.5 | 502571.31 | 324935.94 | 316540.59 | 321177.5 | 451073.84 | **0.718727305** | 1 |
| sp\|Q1500 | Q15007 | WTAP | Pre-mRNA-splicing regulator WTAP OS=Homo sapiens (H | 547997.25 | 1566256.38 | 693729.81 | 632829.75 | 552485.94 | 586944.12 | 720598.19 | 611748.5 | **0.718369936** | 6 |
| sp\|Q96P4 | Q96P48 | ARAP1 | Arf-GAP with Rho-GAP domain, ANK repeat and PH doma | 5244575.5 | 5844574 | 6699203 | 6630778 | 4392479.5 | 3968471.75 | 4362461 | 4812123 | **0.718106456** | 0 |
| sp\|Q8TED | Q8TED0 | UTP15 | U3 small nucleolar RNA-associated protein 15 homolog | 5875404 | 5941703 | 5973573 | 5794791 | 3313759.75 | 3867569.5 | 4871745.5 | 4869215.5 | **0.717487908** | 0 |
| sp\|P57088 | P57088 | TMEM33 | Transmembrane protein 33 OS=Homo sapiens (Human) | 20117580 | 20214144 | 25873234 | 19119162 | 15961314 | 15354748 | 14851895 | 15036694 | **0.717319452** | 0 |
| sp\|Q9P28 | Q9P289 | STK26 | Serine/threonine-protein kinase 26 OS=Homo sapiens (H | 5886163 | 6293988 | 7098770.5 | 5884047 | 5436236.5 | 5045286 | 3839521.5 | 3719476.5 | **0.716947227** | 0 |
| sp\|Q96SB | Q96SB4 | SRPK1 | SRSF protein kinase 1 OS=Homo sapiens (Human) GN=SF | 6030820.5 | 4825223 | 6049478 | 4967027.5 | 4654721 | 4711212 | 3666640 | 2637035.5 | **0.71640523** | 0 |
| sp\|P31146 | P31146 | CORO1A | Coronin-1A OS=Homo sapiens (Human) GN=CORO1A PE | 860382.12 | 726548.25 | 597239.75 | 703880.12 | 532102.88 | 553056.25 | 463010.41 | 520235.16 | **0.716194155** | 0 |
| sp\|Q96FZ | Q96FZ7 | CHMP6 | Charged multivesicular body protein 6 OS=Homo sapien | 1213091.12 | 959359.56 | 1288233.5 | 888686.44 | 962598.81 | 722136.56 | 664242.62 | 765776.56 | **0.716139143** | 2 |
| sp\|Q1594 | Q15942 | ZYX | Zyxin OS=Homo sapiens (Human) GN=ZYX PE=1 SV=1 | 14864110 | 10026600 | 12491095 | 11202955 | 8090354 | 6871080 | 10573860 | 9250665 | **0.715984992** | 1 |
| sp\|Q8NBS | Q8NBS9 | TXNDC5 | Thioredoxin domain-containing protein 5 OS=Homo sapi | 138351920 | 123639752 | 132944400 | 128304024 | 101648112 | 90247536 | 96698032 | 86013720 | **0.71593787** | 0 |
| sp\|P32969 | P32969 | RPL9 | 60S ribosomal protein L9 OS=Homo sapiens (Human) GN | 75170696 | 91976344 | 94452288 | 98761064 | 58773892 | 66112164 | 57397680 | 75618624 | **0.715678986** | 1 |
| sp\|Q9943 | Q99439 | CNN2 | Calponin-2 OS=Homo sapiens (Human) GN=CNN2 PE=1 S | 4378569 | 3470662.75 | 5117093 | 5149983.5 | 3014565.5 | 3558006 | 3530764 | 2860795.25 | **0.715605551** | 2 |
| sp\|P20340 | P20340 | RAB6A | Ras-related protein Rab-6A OS=Homo sapiens (Human) | 17701116 | 15398130 | 22188116 | 19074842 | 12884286 | 12700830 | 15431011 | 12195649 | **0.715575563** | 1 |
| sp\|P61769 | P61769 | B2M | Beta-2-microglobulin OS=Homo sapiens (Human) GN=B2 | 1729236 | 2294405 | 1239450.12 | 2449846.25 | 1317255 | 1695844.25 | 1476626.88 | 1027144.25 | **0.715274884** | 3 |
| sp\|O9548 | O95487 | SEC24B | Protein transport protein Sec24B OS=Homo sapiens (Hu | 2275056.75 | 3087600.25 | 2515435 | 2345978.25 | 1970754.25 | 1869839.62 | 1685475 | 1782775.62 | **0.714866419** | 0 |
| sp\|P61086 | P61086 | UBE2K | Ubiquitin-conjugating enzyme E2 K OS=Homo sapiens (H | 8786116 | 11672624 | 11762441 | 10807411 | 6366212.5 | 8141871 | 6834075 | 9391954 | **0.714271861** | 1 |
| sp\|Q9UDY | Q9UDY2 | TJP2 | Tight junction protein ZO-2 OS=Homo sapiens (Human) | 40246864 | 45112148 | 48627776 | 38767388 | 32248238 | 24353614 | 31989672 | 34751728 | **0.713981305** | 0 |
| sp\|P53985 | P53985 | SLC16A1 | Monocarboxylate transporter 1 OS=Homo sapiens (Hum | 2173767.5 | 3240974.5 | 2000014.75 | 3082134 | 2348986.5 | 2626045.75 | 1790098.5 | 728732.81 | **0.713912695** | 4 |
| sp\|P35579 | P35579 | MYH9 | Myosin-9 OS=Homo sapiens (Human) GN=MYH9 PE=1 SV | 205155280 | 199750912 | 220502896 | 177569056 | 140630496 | 145142992 | 150420896 | 136819584 | **0.713610915** | 0 |
| sp\|Q1389 | Q13895 | BYSL | Bystin OS=Homo sapiens (Human) GN=BYSL PE=1 SV=3 | 10331434 | 14050437 | 13634354 | 11194468 | 10673269 | 7043645 | 8799964 | 8596206 | **0.713525493** | 1 |
| sp\|Q8WYF | Q8WYP5 | AHCTF1 | Protein ELYS OS=Homo sapiens (Human) GN=AHCTF1 PE | 2529577 | 4056371 | 3592874.75 | 4544260 | 2377776 | 2029133.25 | 3331142.5 | 2763350.25 | **0.713261087** | 2 |
| sp\|Q9973 | Q99733 | NAP1L4 | Nucleosome assembly protein 1-like 4 OS=Homo sapiens | 64172424 | 60663292 | 57186992 | 55039020 | 47662488 | 41892420 | 41683788 | 37809072 | **0.713095992** | 0 |
| sp\|Q0463 | Q04637 | EIF4G1 | Eukaryotic translation initiation factor 4 gamma 1 OS=Hc | 53447000 | 60295836 | 67790704 | 51699920 | 38979272 | 44681304 | 46336916 | 36224056 | **0.712683112** | 0 |
| sp\|Q5VTL | Q5VTL8 | PRPF38B | Pre-mRNA-splicing factor 38B OS=Homo sapiens (Humar | 371384.34 | 256953.19 | 246418.36 | 385273.28 | 282238.66 | 354150.78 | 260530.19 | 0 | **0.711824497** | 6 |

| | | | | | | | | | | | | |
|---|---|---|---|---|---|---|---|---|---|---|---|---|
| sp\|Q9UBM | Q9UBM7 | DHCR7 | 7-dehydrocholesterol reductase OS=Homo sapiens (Hum | 16355172 | 15663450 | 16971176 | 15540774 | 10469011 | 11979108 | 11382983 | 12070020 | **0.711308153** | 0 |
| sp\|P02671 | P02671 | FGA | Fibrinogen alpha chain OS=Homo sapiens (Human) GN=F | 7134156 | 9076730 | 10771640 | 8324650 | 8003439 | 4101321.75 | 7703937 | 5282066 | **0.710642045** | 2 |
| sp\|O1482 | O14828 | SCAMP3 | Secretory carrier-associated membrane protein 3 OS=Ho | 2858181.5 | 2571871.75 | 3255087.5 | 2393068.5 | 2499560.75 | 2137784 | 1236124.62 | 1997705 | **0.710509631** | 1 |
| sp\|Q9BVP | Q9BVP2 | GNL3 | Guanine nucleotide-binding protein-like 3 OS=Homo sap | 19646774 | 18889246 | 22506454 | 19848592 | 12967335 | 12609298 | 15856757 | 16002843 | **0.710044209** | 0 |
| sp\|Q9UHC | Q9UHQ9 | CYB5R1 | NADH-cytochrome b5 reductase 1 OS=Homo sapiens (Hu | 2092120.38 | 0 | 1212510.12 | 949341.81 | 1388277.12 | 0 | 1629157.62 | 0 | **0.709321669** | 7 |
| sp\|P08237 | P08237 | PFKM | ATP-dependent 6-phosphofructokinase, muscle type OS | 1687654.62 | 1821586 | 2001450.5 | 2287858.5 | 1100445.75 | 1767905.25 | 1401794.12 | 1261325.62 | **0.709294806** | 1 |
| sp\|Q9979 | Q99797 | MIPEP | Mitochondrial intermediate peptidase OS=Homo sapien | 863756.38 | 962321.69 | 96014.95 | 1091210 | 924371.19 | 958193.56 | 138134.28 | 113199.48 | **0.708159284** | 6 |
| sp\|Q96SZ | Q96SZ6 | CDK5RAP1 | CDK5 regulatory subunit-associated protein 1 OS=Homo | 1590651.38 | 1524696 | 1662849.88 | 1552698.25 | 258373.28 | 2344169.25 | 1730015.38 | 148463.11 | **0.707802082** | 8 |
| sp\|P41250 | P41250 | GARS | Glycine--tRNA ligase OS=Homo sapiens (Human) GN=GA | 77853208 | 78626176 | 65711752 | 75889072 | 53932188 | 43457328 | 54968464 | 58394500 | **0.707032786** | 0 |
| sp\|Q9Y5J1 | Q9Y5J1 | UTP18 | U3 small nucleolar RNA-associated protein 18 homolog ( | 8993434 | 2813378.5 | 9755438 | 10205179 | 6743609 | 6431064.5 | 5142987 | 4103879 | **0.705802763** | 4 |
| sp\|P52306 | P52306 | RAP1GDS1 | Rap1 GTPase-GDP dissociation stimulator 1 OS=Homo sa | 5653818 | 4248838 | 5754039.5 | 4467534 | 3997804.5 | 3931671 | 2696608 | 3577189.25 | **0.705779705** | 0 |
| sp\|Q9UN8 | Q9UN86 | G3BP2 | Ras GTPase-activating protein-binding protein 2 OS=Hon | 5662050 | 6248755.5 | 5690887.5 | 5973106 | 6700572 | 3446458.25 | 3934645.5 | 2548649 | **0.705428061** | 4 |
| sp\|Q7Z47 | Q7Z478 | DHX29 | ATP-dependent RNA helicase DHX29 OS=Homo sapiens ( | 1001985.5 | 2014455.5 | 1277962 | 1385650.12 | 934972.75 | 265649.88 | 1713430.5 | 1092294.75 | **0.70533634** | 4 |
| sp\|Q6L8Q | Q6L8Q7 | PDE12 | 2',5'-phosphodiesterase 12 OS=Homo sapiens (Human) ( | 7975544 | 7486400 | 7219715.5 | 8751607 | 6035564 | 7195903.5 | 3553612.5 | 5377880 | **0.70507976** | 0 |
| sp\|Q9BW | Q9BWQ6 | YIPF2 | Protein YIPF2 OS=Homo sapiens (Human) GN=YIPF2 PE= | 260361.25 | 0 | 0 | 0 | 0 | 183524.2 | 0 | 0 | **0.704882927** | 7.5 |
| sp\|Q9Y6G | Q9Y6G9 | DYNC1LI1 | Cytoplasmic dynein 1 light intermediate chain 1 OS=Hon | 9477859 | 8752802 | 11905398 | 9639579 | 7744662 | 6906228 | 6583204 | 6779408.5 | **0.704287949** | 0 |
| sp\|Q5T6F | Q5T6F2 | UBAP2 | Ubiquitin-associated protein 2 OS=Homo sapiens (Huma | 3730432.5 | 1256399.75 | 3635752 | 4341949 | 2635266 | 2546337.75 | 2338793.75 | 1609921.75 | **0.704253603** | 4 |
| sp\|Q1276 | Q12765 | SCRN1 | Secernin-1 OS=Homo sapiens (Human) GN=SCRN1 PE=1 | 9181346 | 9822604 | 10911294 | 7171023 | 7010609.5 | 6069776 | 6929062.5 | 6106188 | **0.704186161** | 0 |
| sp\|O9575 | O95758 | PTBP3 | Polypyrimidine tract-binding protein 3 OS=Homo sapien | 51323932 | 46345200 | 59908580 | 63316072 | 45167932 | 41754904 | 35621840 | 32815416 | **0.70332487** | 0 |
| sp\|O7582 | O75828 | CBR3 | Carbonyl reductase [NADPH] 3 OS=Homo sapiens (Huma | 5381395 | 4185942 | 3278605.5 | 4405365 | 3142794.5 | 3096046.5 | 3205620.25 | 2687989.25 | **0.703277157** | 0 |
| sp\|P53801 | P53801 | PTTG1IP | Pituitary tumor-transforming gene 1 protein-interacting | 2350220.5 | 2535119.5 | 2134204 | 2118527.75 | 2831589 | 3087946.5 | 0 | 501679.38 | **0.702688166** | 8 |
| sp\|Q8IYD1 | Q8IYD1 | GSPT2 | Eukaryotic peptide chain release factor GTP-binding subu | 24827030 | 28007120 | 27838378 | 24910928 | 19860874 | 16443347 | 18999926 | 18848120 | **0.702309527** | 0 |
| sp\|Q9H4L | Q9H4L7 | SMARCAD | SWI/SNF-related matrix-associated actin-dependent regu | 1579359.5 | 614743.06 | 1846263.5 | 1663288.38 | 1718938.25 | 907365.5 | 591375.88 | 787822.38 | **0.702269405** | 5 |
| sp\|P54709 | P54709 | ATP1B3 | Sodium/potassium-transporting ATPase subunit beta-3 ( | 13687248 | 16882300 | 16036162 | 18502040 | 12563389 | 12982181 | 9303912 | 10861518 | **0.702082317** | 0 |
| sp\|Q0053 | Q00534 | CDK6 | Cyclin-dependent kinase 6 OS=Homo sapiens (Human) G | 6393364.5 | 4427774 | 5387173.5 | 5968211.5 | 3037934.5 | 4439908.5 | 4564546.5 | 3515948 | **0.701567922** | 2 |
| sp\|Q6UN1 | Q6UN15 | FIP1L1 | Pre-mRNA 3'-end-processing factor FIP1 OS=Homo sapie | 3229980.25 | 3348164.5 | 3880434.5 | 3109398 | 2731608.5 | 1940071.62 | 2699996.5 | 2130800.75 | **0.70036065** | 0 |
| sp\|P57105 | P57105 | SYNJ2BP | Synaptojanin-2-binding protein OS=Homo sapiens (Hum | 889939.31 | 1576722.5 | 1669290.12 | 1806136.75 | 772070.88 | 898768.75 | 1435264 | 1055345.75 | **0.700334445** | 3 |
| sp\|P26641 | P26641 | EEF1G | Elongation factor 1-gamma OS=Homo sapiens (Human) ( | 163511200 | 156312528 | 202428272 | 169955200 | 122316776 | 120869696 | 110915184 | 130648592 | **0.700296455** | 0 |
| sp\|P54687 | P54687 | BCAT1 | Branched-chain-amino-acid aminotransferase, cytosolic | 2360789.75 | 1971313.75 | 1256824.62 | 1599727.88 | 1167324.12 | 1148898.75 | 1357125.62 | 1360674.38 | **0.700273162** | 2 |
| sp\|Q6ZRP | Q6ZRP7 | QSOX2 | Sulfhydryl oxidase 2 OS=Homo sapiens (Human) GN=QS( | 5723631.5 | 5448659.5 | 4996983 | 5807908 | 3825324.75 | 4593517 | 3442995 | 3518072.25 | **0.699812606** | 0 |
| sp\|Q27J81 | Q27J81 | INF2 | Inverted formin-2 OS=Homo sapiens (Human) GN=INF2 I | 3266214.5 | 2502958.5 | 3204788.75 | 3469517 | 1822250.5 | 2375121.75 | 2186756.75 | 2317908.5 | **0.699325138** | 0 |
| sp\|Q9HBU | Q9HBU6 | ETNK1 | Ethanolamine kinase 1 OS=Homo sapiens (Human) GN=E | 494402.25 | 0 | 593937.56 | 761963.31 | 679491.94 | 612522.56 | 0 | 0 | **0.698271805** | 7 |
| sp\|Q0620 | Q06203 | PPAT | Amidophosphoribosyltransferase OS=Homo sapiens (Hu | 10175898 | 9345854 | 8854320 | 9438886 | 7663561 | 6460886 | 6063435 | 6205588.5 | **0.697963766** | 0 |
| sp\|Q8IZ73 | Q8IZ73 | RPUSD2 | RNA pseudouridylate synthase domain-containing protei | 200886.95 | 174792.69 | 87893.39 | 157767.66 | 158324.16 | 112359.14 | 99968.6 | 63002.39 | **0.69793319** | 4 |
| sp\|Q0781 | Q07817 | BCL2L1 | Bcl-2-like protein 1 OS=Homo sapiens (Human) GN=BCL2 | 3417835.5 | 3781545.5 | 4737124 | 4290983 | 2928235.5 | 2616020.5 | 2950181.5 | 2817757.5 | **0.697100808** | 0 |
| sp\|Q9NUC | Q9NUQ6 | SPATS2L | SPATS2-like protein OS=Homo sapiens (Human) GN=SPA | 22380356 | 25066622 | 26015326 | 22231746 | 14959429 | 19259306 | 17799440 | 14689638 | **0.697094678** | 0 |
| sp\|O9600 | O96005 | CLPTM1 | Cleft lip and palate transmembrane protein 1 OS=Homo | 2614127.25 | 1601545 | 2953485.5 | 1966675.25 | 1353905.25 | 2332168.5 | 1365565.75 | 1308904.5 | **0.696219381** | 2 |
| sp\|P16188 | P16188 | HLA-A | HLA class I histocompatibility antigen, A-30 alpha chain ( | 32521048 | 29980186 | 32187150 | 29936564 | 24135672 | 22032834 | 21493768 | 19061126 | **0.695875115** | 0 |
| sp\|Q9H32 | Q9H324 | ADAMTS1( | A disintegrin and metalloproteinase with thrombospondi | 4276926 | 3600084.75 | 3590160 | 0 | 1631890.88 | 2284149.75 | 2255031.5 | 1805309.38 | **0.695584088** | 4 |
| sp\|Q1541 | Q15417 | CNN3 | Calponin-3 OS=Homo sapiens (Human) GN=CNN3 PE=1 S | 17412926 | 17760714 | 16685979 | 17957782 | 12668847 | 11058787 | 12860520 | 11923127 | **0.694830806** | 0 |
| sp\|Q9UHJ | Q9UHJ6 | SHPK | Sedoheptulokinase OS=Homo sapiens (Human) GN=SHP | 4123986.5 | 4351940.5 | 4035600.5 | 4296967.5 | 2707231.5 | 2330639.5 | 2620006 | 3998288.5 | **0.693468719** | 0 |
| sp\|P51948 | P51948 | MNAT1 | CDK-activating kinase assembly factor MAT1 OS=Homo s | 760634.44 | 688008.31 | 417672.16 | 725313.94 | 441291.06 | 543997.44 | 467409.28 | 344005.19 | **0.693271712** | 3 |
| sp\|P11766 | P11766 | ADH5 | Alcohol dehydrogenase class-3 OS=Homo sapiens (Huma | 65439656 | 69821032 | 62245768 | 66801504 | 42434556 | 43769324 | 49011820 | 47285416 | **0.690486643** | 0 |
| sp\|Q1575 | Q15758 | SLC1A5 | Neutral amino acid transporter B(0) OS=Homo sapiens (H | 46529300 | 51774936 | 45641916 | 46633300 | 34823644 | 33682896 | 28997690 | 33804044 | **0.688994919** | 0 |
| sp\|Q9C0B | Q9C0B1 | FTO | Alpha-ketoglutarate-dependent dioxygenase FTO OS=Ho | 4527116 | 5353995 | 4747911 | 5235686.5 | 4002754.25 | 2940654 | 2928128.25 | 3787334 | **0.687594812** | 0 |

| | | | | | | | | | | | | | |
|---|---|---|---|---|---|---|---|---|---|---|---|---|---|
| sp\|Q1321 | Q13217 | DNAJC3 | DnaJ homolog subfamily C member 3 OS=Homo sapiens | 4145983.75 | 4076668.5 | 3038305.5 | 3758208.5 | 2718845.25 | 3222989.75 | 1155030.25 | 3227371.25 | 0.687404103 | 2 |
| sp\|Q8NEZ | Q8NEZ5 | FBXO22 | F-box only protein 22 OS=Homo sapiens (Human) GN=FE | 2548801 | 3147987.5 | 3300317 | 3076986.5 | 2286991.75 | 1965329.12 | 1920631.5 | 2125418 | 0.687287323 | 0 |
| sp\|P07942 | P07942 | LAMB1 | Laminin subunit beta-1 OS=Homo sapiens (Human) GN= | 10946356 | 9059160 | 9409702 | 8161909 | 4931848 | 8037785.5 | 9131278 | 3708673.25 | 0.68684295 | 2 |
| sp\|Q96EK | Q96EK9 | KTI12 | Protein KTI12 homolog OS=Homo sapiens (Human) GN=I | 329939.53 | 274760.09 | 299606.56 | 488451.56 | 222794.5 | 274327.78 | 234068.86 | 224533.25 | 0.686210073 | 0 |
| sp\|P26640 | P26640 | VARS | Valine--tRNA ligase OS=Homo sapiens (Human) GN=VAR | 94195520 | 99902864 | 99929232 | 90508984 | 68751368 | 66539900 | 63777940 | 64682048 | 0.685893764 | 0 |
| sp\|Q9BR7 | Q9BR76 | CORO1B | Coronin-1B OS=Homo sapiens (Human) GN=CORO1B PE= | 15159609 | 12774097 | 12668817 | 12473922 | 10521993 | 8271679 | 10203083 | 7392688.5 | 0.685604386 | 0 |
| sp\|Q1435 | Q14353 | GAMT | Guanidinoacetate N-methyltransferase OS=Homo sapier | 5257445 | 4610790 | 3718102 | 3216198.5 | 4147929.75 | 2976780.5 | 1675236.25 | 2713469 | 0.685218936 | 2 |
| sp\|P69849 | P69849 | NOMO3 | Nodal modulator 3 OS=Homo sapiens (Human) GN=NON | 33811128 | 34207436 | 38746584 | 33470598 | 24387884 | 24779284 | 22646204 | 24215318 | 0.684766137 | 0 |
| sp\|P53396 | P53396 | ACLY | ATP-citrate synthase OS=Homo sapiens (Human) GN=AC | 145445072 | 154704640 | 162041392 | 137539936 | 110639056 | 95450416 | 105419272 | 97826488 | 0.682531343 | 0 |
| sp\|P15907 | P15907 | ST6GAL1 | Beta-galactoside alpha-2,6-sialyltransferase 1 OS=Homo | 1455907.5 | 1328072.25 | 1338392.25 | 0 | 836552 | 0 | 1072698.75 | 903229.25 | 0.682247987 | 3.5 |
| sp\|Q86WJ | Q86WJ1 | CHD1L | Chromodomain-helicase-DNA-binding protein 1-like OS= | 1576742.88 | 1241468.75 | 1647633 | 1378617.12 | 1125657.25 | 1063892.25 | 996585.56 | 799545.69 | 0.681958565 | 0 |
| sp\|P00338 | P00338 | LDHA | L-lactate dehydrogenase A chain OS=Homo sapiens (Hur | 1474436864 | 1362272512 | 1508999552 | 1448700928 | 1057448384 | 1135094784 | 897151424 | 860935552 | 0.68180026 | 0 |
| sp\|O7513 | O75131 | CPNE3 | Copine-3 OS=Homo sapiens (Human) GN=CPNE3 PE=1 S\ | 22232636 | 18922826 | 17830668 | 17517912 | 14422130 | 11352288 | 13215940 | 13130825 | 0.681286657 | 0 |
| sp\|Q5U4P | Q5U4P2 | ASPHD1 | Aspartate beta-hydroxylase domain-containing protein 1 | 175431.3 | 205516.19 | 174671.55 | 260253.3 | 145607.94 | 136752.98 | 128369.57 | 144776.52 | 0.680874915 | 0 |
| sp\|O1537 | O15372 | EIF3H | Eukaryotic translation initiation factor 3 subunit H OS=H | 16752896 | 16199580 | 22373416 | 18796880 | 16878346 | 13309424 | 12415995 | 7838648 | 0.680525183 | 2 |
| sp\|Q96GG | Q96GG9 | DCUN1D1 | DCN1-like protein 1 OS=Homo sapiens (Human) GN=DCL | 1736462.25 | 1646005.5 | 2246985.5 | 1620919 | 1174827.38 | 1773602.75 | 1118145.25 | 866354.88 | 0.68036924 | 3 |
| sp\|Q9269 | Q92692 | NECTIN2 | Nectin-2 OS=Homo sapiens (Human) GN=NECTIN2 PE=1 | 2657892.5 | 2257272 | 2160240 | 2549662 | 2262257 | 838039.56 | 1074455.88 | 2371458 | 0.680121061 | 4 |
| sp\|P51148 | P51148 | RAB5C | Ras-related protein Rab-5C OS=Homo sapiens (Human) ( | 35192480 | 36177536 | 37318488 | 36632964 | 26383040 | 24770588 | 22438076 | 25242780 | 0.680109315 | 0 |
| sp\|P02763 | P02763 | ORM1 | Alpha-1-acid glycoprotein 1 OS=Homo sapiens (Human) | 1055486.25 | 1588800.75 | 1500949.5 | 1391752.5 | 1042750.12 | 793735.5 | 1156903 | 772066 | 0.68005456 | 1 |
| sp\|Q9NVA | Q9NVA2 | SEPT11 | Septin-11 OS=Homo sapiens (Human) GN=SEPT11 PE=1 S | 13986905 | 14542694 | 11824750 | 10283133 | 9142985 | 9308220 | 9531144 | 6453191 | 0.680040528 | 0 |
| sp\|Q1498 | Q14980 | NUMA1 | Nuclear mitotic apparatus protein 1 OS=Homo sapiens (I | 21238602 | 18899430 | 22360176 | 15579408 | 15188098 | 14400033 | 12328801 | 11131394 | 0.679430658 | 0 |
| sp\|Q1504 | Q15041 | ARL6IP1 | ADP-ribosylation factor-like protein 6-interacting protein | 13641814 | 18252556 | 17565070 | 17293546 | 13050873 | 11573560 | 9353822 | 11373964 | 0.6794036 | 0 |
| sp\|Q9ULK | Q9ULK4 | MED23 | Mediator of RNA polymerase II transcription subunit 23 | 218595.16 | 490844.56 | 472759.81 | 549471.88 | 326181.62 | 519879.44 | 123307.98 | 205496.28 | 0.678457422 | 4 |
| sp\|P16383 | P16383 | GCFC2 | GC-rich sequence DNA-binding factor 2 OS=Homo sapier | 150862.67 | 178144.28 | 70052.05 | 196555.34 | 147053 | 97827.39 | 87108.45 | 72076.1 | 0.678400288 | 4 |
| sp\|Q68CZ | Q68CZ2 | TNS3 | Tensin-3 OS=Homo sapiens (Human) GN=TNS3 PE=1 SV= | 6361378 | 5328333 | 5248884.5 | 5281693 | 3973697 | 3840311.25 | 3638746 | 3616082.25 | 0.678156654 | 0 |
| sp\|Q9NW | Q9NW68 | BSDC1 | BSD domain-containing protein 1 OS=Homo sapiens (Hu | 283992.91 | 284186.78 | 0 | 269088.66 | 256550.45 | 310944.16 | 0 | 0 | 0.67779298 | 6 |
| sp\|Q9H1A | Q9H1A4 | ANAPC1 | Anaphase-promoting complex subunit 1 OS=Homo sapie | 1621613.62 | 1559168.5 | 720204.06 | 1483079.62 | 695349.75 | 1497745.25 | 673584.38 | 782560.19 | 0.677785099 | 3 |
| sp\|Q96LD | Q96LD4 | TRIM47 | E3 ubiquitin-protein ligase TRIM47 OS=Homo sapiens (H | 5674105 | 4884013 | 5171499 | 5326138 | 3431284 | 3697843.5 | 3618467 | 3496316 | 0.676485384 | 0 |
| sp\|Q1330 | Q13309 | SKP2 | S-phase kinase-associated protein 2 OS=Homo sapiens (I | 191588.86 | 182082.16 | 1302076 | 433834.78 | 171490.58 | 250096.56 | 168294.23 | 836675.81 | 0.676227478 | 5 |
| sp\|P18283 | P18283 | GPX2 | Glutathione peroxidase 2 OS=Homo sapiens (Human) GN | 4004912 | 4136259 | 4320697 | 3976924 | 2767693 | 2458850.25 | 2966655.75 | 2919153.5 | 0.675983521 | 0 |
| sp\|Q2NL8 | Q2NL82 | TSR1 | Pre-rRNA-processing protein TSR1 homolog OS=Homo sa | 12339642 | 10006615 | 12261009 | 9582760 | 7293233 | 8361626 | 7205096.5 | 6999234 | 0.67569975 | 0 |
| sp\|Q96T8 | Q96T88 | UHRF1 | E3 ubiquitin-protein ligase UHRF1 OS=Homo sapiens (Hu | 2593244 | 3507374 | 7152386 | 5285486 | 3346570.5 | 1693564.25 | 4476515.5 | 3008696.5 | 0.675640074 | 4 |
| sp\|Q0053 | Q00535 | CDK5 | Cyclin-dependent-like kinase 5 OS=Homo sapiens (Huma | 2443427 | 2328293.25 | 2537777 | 2008669.75 | 1536139 | 1439168.5 | 1768907.5 | 1549193.25 | 0.675391228 | 0 |
| sp\|O1523 | O15230 | LAMA5 | Laminin subunit alpha-5 OS=Homo sapiens (Human) GN= | 5449758.5 | 5176525.5 | 4266156.5 | 4882441.5 | 2703537.25 | 3538898.25 | 3705737 | 3395554.75 | 0.674781637 | 0 |
| sp\|P30408 | P30408 | TM4SF1 | Transmembrane 4 L6 family member 1 OS=Homo sapien | 3299265.5 | 5471342 | 3797124.25 | 6312858 | 4327778 | 4727281 | 2956247.5 | 711597.62 | 0.673861584 | 4 |
| sp\|O9516 | O95163 | ELP1 | Elongator complex protein 1 OS=Homo sapiens (Human) | 19353242 | 17903518 | 22266994 | 20916434 | 11138308 | 13654404 | 13944695 | 15279208 | 0.671512789 | 0 |
| sp\|Q9257 | Q92576 | PHF3 | PHD finger protein 3 OS=Homo sapiens (Human) GN=PH | 431289.09 | 504688.56 | 406167.28 | 402247.19 | 0 | 336013.25 | 445113.84 | 390081.78 | 0.671413759 | 3 |
| sp\|Q1484 | Q14847 | LASP1 | LIM and SH3 domain protein 1 OS=Homo sapiens (Huma | 62598096 | 61772456 | 66267616 | 62360732 | 53148216 | 36873168 | 45780264 | 33944496 | 0.670936293 | 0 |
| sp\|A6NKT | A6NKT7 | RGPD3 | RanBP2-like and GRIP domain-containing protein 3 OS=H | 10757464 | 9625138 | 10078660 | 8722685 | 7447685 | 3409721.5 | 8144887.5 | 7275686.5 | 0.670631279 | 0 |
| sp\|Q0325 | Q03252 | LMNB2 | Lamin-B2 OS=Homo sapiens (Human) GN=LMNB2 PE=1 S | 170529360 | 179167760 | 223909968 | 179709840 | 134192264 | 118117632 | 121517216 | 130552976 | 0.669545671 | 0 |
| sp\|Q8N54 | Q8N543 | OGFOD1 | Prolyl 3-hydroxylase OGFOD1 OS=Homo sapiens (Humar | 1383279.5 | 1245235 | 1341157 | 634398.19 | 756897.94 | 1048297.06 | 228882.77 | 1048134.06 | 0.669453774 | 3 |
| sp\|Q9Y6M | Q9Y6M5 | SLC30A1 | Zinc transporter 1 OS=Homo sapiens (Human) GN=SLC3C | 1134524.75 | 1277975.75 | 1243997.38 | 1159604.62 | 882089.5 | 831984.5 | 742721.19 | 765689.81 | 0.669106399 | 0 |
| sp\|Q9Y37 | Q9Y371 | SH3GLB1 | Endophilin-B1 OS=Homo sapiens (Human) GN=SH3GLB1 | 3143940.5 | 2793483 | 3343507 | 2784908 | 2256094.5 | 2499633 | 2138932.75 | 1174430.25 | 0.668755056 | 0 |
| sp\|Q8TER | Q8TER5 | ARHGEF40 | Rho guanine nucleotide exchange factor 40 OS=Homo sa | 295745.22 | 197163.41 | 129514.01 | 0 | 108056.67 | 125505.89 | 87891.17 | 94714.85 | 0.668626996 | 4 |
| sp\|Q9961 | Q99615 | DNAJC7 | DnaJ homolog subfamily C member 7 OS=Homo sapiens | 11489683 | 11104128 | 14044066 | 12664199 | 9832206 | 6815315 | 9553868 | 6711817 | 0.667582558 | 0 |

| | | | | | | | | | | | | | |
|---|---|---|---|---|---|---|---|---|---|---|---|---|---|
| sp\|Q9BU2 | Q9BU23 | LMF2 | Lipase maturation factor 2 OS=Homo sapiens (Human) G | 821556.62 | 537115.5 | 660111.69 | 1163289.5 | 0 | 775263.69 | 0 | 1348710.5 | **0.667481225** | 6 |
| sp\|O6030( | O60306 | AQR | RNA helicase aquarius OS=Homo sapiens (Human) GN=A | 1560907.75 | 2197838 | 1192520.75 | 3345460.5 | 977588.69 | 1285569.75 | 1640950 | 1632887.75 | **0.667371144** | 5 |
| sp\|Q9259( | Q92598 | HSPH1 | Heat shock protein 105 kDa OS=Homo sapiens (Human) | 65044104 | 65314248 | 85178544 | 66143516 | 48186552 | 47870884 | 48464508 | 43252644 | **0.666622811** | 0 |
| sp\|Q9NP5 | Q9NP58 | ABCB6 | ATP-binding cassette sub-family B member 6, mitochond | 519138.53 | 404222.62 | 524965.12 | 550745.31 | 400013.44 | 333816.91 | 266006.62 | 331525.09 | **0.665990189** | 0 |
| sp\|P61289 | P61289 | PSME3 | Proteasome activator complex subunit 3 OS=Homo sapie | 5206587 | 13661101 | 17734608 | 15076302 | 13772925 | 11295912 | 4439115.5 | 4877585 | **0.665372878** | 3 |
| sp\|P49770 | P49770 | EIF2B2 | Translation initiation factor eIF-2B subunit beta OS=Hom | 1155338.62 | 2590108 | 3030635.25 | 2158246.5 | 2092132.5 | 1136192 | 2314726 | 388954.12 | **0.663956413** | 3 |
| sp\|P15170 | P15170 | GSPT1 | Eukaryotic peptide chain release factor GTP-binding subu | 34527668 | 42271176 | 43591520 | 38936880 | 30410832 | 23299548 | 26425590 | 25613666 | **0.66372601** | 0 |
| sp\|Q9UI2( | Q9UI26 | IPO11 | Importin-11 OS=Homo sapiens (Human) GN=IPO11 PE=1 | 1282147.25 | 1986573 | 1537502.12 | 1956703.88 | 1147341.62 | 1194066 | 684431.56 | 1462484.12 | **0.663665865** | 1 |
| sp\|Q8NBJ! | Q8NBJ5 | COLGALT1 | Procollagen galactosyltransferase 1 OS=Homo sapiens (H | 23282682 | 21539422 | 28307482 | 28341766 | 19675224 | 16835576 | 17098798 | 13717945 | **0.663512821** | 0 |
| sp\|P32780 | P32780 | GTF2H1 | General transcription factor IIH subunit 1 OS=Homo sapi | 490321.47 | 492529.78 | 364816.03 | 448411.56 | 219039.66 | 346182.09 | 355510.16 | 270469.31 | **0.663223236** | 0 |
| sp\|P43007 | P43007 | SLC1A4 | Neutral amino acid transporter A OS=Homo sapiens (Hu | 8014230 | 8725560 | 7970980 | 7569459.5 | 4855168 | 5857242.5 | 5180947.5 | 5504738 | **0.662885498** | 0 |
| sp\|O0044: | O00443 | PIK3C2A | Phosphatidylinositol 4-phosphate 3-kinase C2 domain-co | 0 | 349887.75 | 669756.5 | 748090.25 | 229632.03 | 245121.38 | 187126.3 | 509654.94 | **0.662732243** | 5 |
| sp\|Q9275( | Q92759 | GTF2H4 | General transcription factor IIH subunit 4 OS=Homo sapi | 350227.12 | 295902.78 | 219394 | 340130.09 | 167359.3 | 238588.78 | 209552.36 | 182743.11 | **0.662083447** | 1 |
| sp\|Q1514( | Q15149 | PLEC | Plectin OS=Homo sapiens (Human) GN=PLEC PE=1 SV=3 | 67242528 | 73133840 | 66023684 | 59914356 | 38166116 | 41319692 | 51583924 | 45184360 | **0.661827099** | 0 |
| sp\|P53384 | P53384 | NUBP1 | Cytosolic Fe-S cluster assembly factor NUBP1 OS=Homo | 2660115 | 2068014.62 | 2777929.25 | 2437427.25 | 1757250.62 | 1659738.88 | 1652268 | 1494707.25 | **0.660127109** | 0 |
| sp\|Q9C0C! | Q9C0C9 | UBE2O | (E3-independent) E2 ubiquitin-conjugating enzyme OS=H | 3641948.5 | 2549027 | 4342388 | 2899022.75 | 2188788.25 | 2700901.75 | 2405565.25 | 1571302.62 | **0.660088067** | 1 |
| sp\|Q9NSI2 | Q9NSI2 | FAM207A | Protein FAM207A OS=Homo sapiens (Human) GN=FAM2 | 619346.62 | 506823.66 | 290155.56 | 848817.88 | 326005.44 | 0 | 831160.88 | 334116.06 | **0.658361042** | 5 |
| sp\|P37837 | P37837 | TALDO1 | Transaldolase OS=Homo sapiens (Human) GN=TALDO1 P | 262722688 | 272854944 | 295747392 | 285821824 | 201313120 | 174230512 | 187402272 | 172298912 | **0.658145182** | 0 |
| sp\|Q96TA: | Q96TA1 | FAM129B | Niban-like protein 1 OS=Homo sapiens (Human) GN=FAM | 4895307 | 3932353.75 | 4650277 | 3571763.5 | 1151712.75 | 3223807.5 | 2485846.5 | 4348763.5 | **0.657497166** | 2 |
| sp\|P18858 | P18858 | LIG1 | DNA ligase 1 OS=Homo sapiens (Human) GN=LIG1 PE=1 S | 1209899.5 | 2785123 | 884413.88 | 815077.38 | 833121.62 | 928849.88 | 862827.94 | 1117029.25 | **0.657093625** | 6 |
| sp\|Q1539: | Q15397 | PUM3 | Pumilio homolog 3 OS=Homo sapiens (Human) GN=PUM | 6851922 | 6370550 | 7021629 | 6633583 | 4277733.5 | 3794521 | 3816209 | 5726856.5 | **0.655388314** | 0 |
| sp\|O0041( | O00410 | IPO5 | Importin-5 OS=Homo sapiens (Human) GN=IPO5 PE=1 SV | 78907000 | 74833800 | 90383208 | 79080272 | 54890716 | 53839468 | 49494508 | 53525412 | **0.655158725** | 0 |
| sp\|P55268 | P55268 | LAMB2 | Laminin subunit beta-2 OS=Homo sapiens (Human) GN= | 326881.56 | 318466.38 | 304656.28 | 314965.75 | 137796.89 | 140894.45 | 286706.06 | 261535.81 | **0.653717661** | 0 |
| sp\|O1527( | O15270 | SPTLC2 | Serine palmitoyltransferase 2 OS=Homo sapiens (Humar | 0 | 0 | 0 | 707843.62 | 0 | 0 | 0 | 461704.5 | **0.65226907** | 7.5 |
| sp\|Q1336: | Q13362 | PPP2R5C | Serine/threonine-protein phosphatase 2A 56 kDa regulat | 1642538.75 | 1714783.62 | 1312611 | 1342512.75 | 1220055.38 | 1308464.75 | 135533.11 | 1254426 | **0.651727959** | 0 |
| sp\|P53365 | P53365 | ARFIP2 | Arfaptin-2 OS=Homo sapiens (Human) GN=ARFIP2 PE=1 | 540816.88 | 627546.19 | 903850.38 | 683016.06 | 0 | 574441.81 | 681767.38 | 539187.25 | **0.651632263** | 3 |
| sp\|Q9UPN | Q9UPN9 | TRIM33 | E3 ubiquitin-protein ligase TRIM33 OS=Homo sapiens (H | 3237657.5 | 2126315.75 | 4006593.5 | 3933396.5 | 2402354.75 | 2684875 | 2089495 | 1487244.62 | **0.651232209** | 2 |
| sp\|Q7Z2T! | Q7Z2T5 | TRMT1L | TRMT1-like protein OS=Homo sapiens (Human) GN=TRM | 568211.5 | 612137.25 | 403834.56 | 450959.22 | 116003.55 | 429034.59 | 453097.06 | 323374.81 | **0.649345189** | 3 |
| sp\|Q9BXJ( | Q9BXJ9 | NAA15 | N-alpha-acetyltransferase 15, NatA auxiliary subunit OS= | 12589870 | 12205853 | 14036600 | 11996560 | 7594748.5 | 8802851 | 7952818 | 8646844 | **0.64918329** | 0 |
| sp\|Q9Y29( | Q9Y296 | TRAPPC4 | Trafficking protein particle complex subunit 4 OS=Homo | 1087318.25 | 618110.81 | 1304460.5 | 1359163.25 | 462627.16 | 738643.62 | 646248.56 | 988732.38 | **0.64916856** | 3 |
| sp\|Q7Z41: | Q7Z417 | NUFIP2 | Nuclear fragile X mental retardation-interacting protein 2 | 4287052 | 3572976.75 | 4486313.5 | 3664578.5 | 2215673.25 | 3195288.25 | 2077167.88 | 2903896.5 | **0.649058605** | 0 |
| sp\|Q7Z2W | Q7Z2W4 | ZC3HAV1 | Zinc finger CCCH-type antiviral protein 1 OS=Homo sapie | 8963853 | 8360221.5 | 10661535 | 8356376 | 7503242 | 4857958.5 | 4279877 | 6943539 | **0.648963346** | 0 |
| sp\|P02786 | P02786 | TFRC | Transferrin receptor protein 1 OS=Homo sapiens (Humai | 95329944 | 92870008 | 108416440 | 100400848 | 66955380 | 70895024 | 56764364 | 62689508 | **0.648093458** | 0 |
| sp\|Q1289( | Q12899 | TRIM26 | Tripartite motif-containing protein 26 OS=Homo sapiens | 1604179.25 | 715592 | 1387636.75 | 777245.81 | 699663.75 | 556507.19 | 1232108.38 | 413591.06 | **0.647066753** | 2 |
| sp\|Q5SRE! | Q5SRE5 | NUP188 | Nucleoporin NUP188 homolog OS=Homo sapiens (Huma | 3098563.5 | 3509904.5 | 3225097.25 | 4130109 | 2365890.75 | 1996525.75 | 2319806 | 2329560 | **0.645373298** | 0 |
| sp\|P29692 | P29692 | EEF1D | Elongation factor 1-delta OS=Homo sapiens (Human) GN | 136499760 | 127102360 | 135129872 | 126047048 | 93015528 | 85011920 | 83747320 | 76822320 | **0.645218391** | 0 |
| sp\|O9537: | O95373 | IPO7 | Importin-7 OS=Homo sapiens (Human) GN=IPO7 PE=1 SV | 41216464 | 40072860 | 39652764 | 35737000 | 22934834 | 27223646 | 23635938 | 27242640 | **0.644866263** | 0 |
| sp\|Q96N6 | Q96N67 | DOCK7 | Dedicator of cytokinesis protein 7 OS=Homo sapiens (Hu | 1623431.12 | 1794241 | 2106428.5 | 1772557.75 | 1161128.62 | 1538716.75 | 600343.19 | 1395699 | **0.643566866** | 0 |
| sp\|Q9Y3A! | Q9Y3A5 | SBDS | Ribosome maturation protein SBDS OS=Homo sapiens (H | 7598524 | 7459606.5 | 8262285 | 7305416 | 3816259.5 | 7302322.5 | 3492883.5 | 5097631.5 | **0.643544878** | 0 |
| sp\|P49589 | P49589 | CARS | Cysteine--tRNA ligase, cytoplasmic OS=Homo sapiens (H | 14002405 | 14990464 | 10023090 | 14446228 | 8585324 | 7644909.5 | 10152620 | 8012775 | **0.643363664** | 1 |
| sp\|P49427 | P49427 | CDC34 | Ubiquitin-conjugating enzyme E2 R1 OS=Homo sapiens ( | 1475881.75 | 1407706.5 | 1846693 | 1577480 | 957924.5 | 968165.94 | 926688.19 | 1205243 | **0.643337861** | 0 |
| sp\|P04156 | P04156 | PRNP | Major prion protein OS=Homo sapiens (Human) GN=PRN | 1858048.38 | 2168157.5 | 1717953.38 | 1911095 | 1559805.62 | 835530.69 | 1605210 | 921257.56 | **0.642931469** | 0 |
| sp\|Q9H93 | Q9H936 | SLC25A22 | Mitochondrial glutamate carrier 1 OS=Homo sapiens (Hu | 2610549 | 3695327 | 4321609.5 | 4641276 | 1113894.38 | 3065789.25 | 2158700.5 | 3434341.75 | **0.640047058** | 2 |
| sp\|Q9H48 | Q9H488 | POFUT1 | GDP-fucose protein O-fucosyltransferase 1 OS=Homo sa | 4422867 | 4002309.5 | 3455086.25 | 3739277 | 2068431.62 | 3254105.75 | 2638025.75 | 2031844 | **0.639737616** | 0 |
| sp\|Q9Y6N | Q9Y6N7 | ROBO1 | Roundabout homolog 1 OS=Homo sapiens (Human) GN= | 1430876 | 1307659.38 | 1174356.5 | 1238495.12 | 686670.5 | 858348.44 | 925039.81 | 820549.06 | **0.638780936** | 0 |

| | | | | | | | | | | | | | |
|---|---|---|---|---|---|---|---|---|---|---|---|---|---|
| sp\|Q86X1( | Q86X10 | RALGAPB | Ral GTPase-activating protein subunit beta OS=Homo sa | 828568 | 913625.62 | 714018.5 | 769985.88 | 661424.75 | 423812.97 | 394150.56 | 580891.19 | 0.638609121 | 0 |
| sp\|Q1653! | Q16539 | MAPK14 | Mitogen-activated protein kinase 14 OS=Homo sapiens ( | 956565.5 | 825620.25 | 1234279.88 | 981838.75 | 352506.16 | 852878.75 | 661067.5 | 685197.75 | 0.638183069 | 1 |
| sp\|Q6ZMZ | Q6ZMZ3 | SYNE3 | Nesprin-3 OS=Homo sapiens (Human) GN=SYNE3 PE=1 S | 262853.59 | 362501.03 | 317099.66 | 348456.53 | 167463.56 | 367026.25 | 289228.72 | 0 | 0.638090969 | 5 |
| sp\|P52788 | P52788 | SMS | Spermine synthase OS=Homo sapiens (Human) GN=SMS | 1646571.75 | 1976673.5 | 1817964.25 | 2102918.75 | 648130.25 | 1176707.75 | 1296307.75 | 1692344 | 0.638044528 | 1 |
| sp\|Q1640: | Q16401 | PSMD5 | 26S proteasome non-ATPase regulatory subunit 5 OS=Hc | 10961903 | 10612798 | 12619810 | 11019413 | 7470794 | 8102712.5 | 7805074 | 5366044 | 0.635747176 | 0 |
| sp\|P09543 | P09543 | CNP | 2',3'-cyclic-nucleotide 3'-phosphodiesterase OS=Homo s | 32677164 | 28280808 | 26496060 | 36983604 | 18737480 | 21602598 | 19724332 | 19015640 | 0.635499456 | 0 |
| sp\|P48059 | P48059 | LIMS1 | LIM and senescent cell antigen-like-containing domain pı | 587532.19 | 0 | 425202 | 489326.03 | 0 | 400008.97 | 554204.19 | 0 | 0.635269577 | 5 |
| sp\|Q1584: | Q15843 | NEDD8 | NEDD8 OS=Homo sapiens (Human) GN=NEDD8 PE=1 SV= | 4919065 | 5483951 | 4955714 | 6237191 | 3554035.75 | 3939972.5 | 3422746.75 | 2796365 | 0.634986579 | 0 |
| sp\|Q1278! | Q12789 | GTF3C1 | General transcription factor 3C polypeptide 1 OS=Homo | 1793507 | 2199914 | 1640546.25 | 2036389.25 | 1205848.25 | 1490192.75 | 753945.94 | 1418695 | 0.634739981 | 0 |
| sp\|Q96PZ( | Q96PZ0 | PUS7 | Pseudouridylate synthase 7 homolog OS=Homo sapiens | 6631863.5 | 6907905 | 5567167 | 6918115 | 3521699.5 | 4530482 | 3087201.25 | 5373661 | 0.634505733 | 0 |
| sp\|Q9BWL | Q9BWU0 | SLC4A1AP | Kanadaptin OS=Homo sapiens (Human) GN=SLC4A1AP P | 1010782.69 | 1033446.38 | 863916.12 | 1602095.25 | 980326.81 | 509780.59 | 642738.12 | 719913.88 | 0.632507166 | 1 |
| sp\|Q96AY: | Q96AY3 | FKBP10 | Peptidyl-prolyl cis-trans isomerase FKBP10 OS=Homo sa | 14669966 | 12036406 | 15680082 | 14026260 | 8507104 | 9592574 | 8694885 | 8869561 | 0.63220011 | 0 |
| sp\|P07339 | P07339 | CTSD | Cathepsin D OS=Homo sapiens (Human) GN=CTSD PE=1 : | 134543456 | 155481760 | 135283424 | 159520528 | 97507688 | 81467800 | 93533528 | 96938904 | 0.631719381 | 0 |
| sp\|Q6EMK | Q6EMK4 | VASN | Vasorin OS=Homo sapiens (Human) GN=VASN PE=1 SV=: | 2216472 | 1984262.75 | 1977878.5 | 2262901.5 | 964693.38 | 922359.06 | 1853793.75 | 1587570 | 0.631215646 | 0 |
| sp\|Q8IUF8 | Q8IUF8 | RIOX2 | Ribosomal oxygenase 2 OS=Homo sapiens (Human) GN= | 2574606.25 | 2877299.25 | 2537204 | 2816768 | 1826234.75 | 1469276.88 | 1709511 | 1812293.25 | 0.630889614 | 0 |
| sp\|Q1326! | Q13268 | DHRS2 | Dehydrogenase/reductase SDR family member 2, mitoch | 402697792 | 398837760 | 338931360 | 495089472 | 252274160 | 232064960 | 233036560 | 314344928 | 0.630807117 | 0 |
| sp\|Q96RS( | Q96RS6 | NUDCD1 | NudC domain-containing protein 1 OS=Homo sapiens (H | 1714723.12 | 1550331.5 | 1259935.62 | 1425713 | 1346088.25 | 1559267.25 | 623361.06 | 221662.73 | 0.630241358 | 4 |
| sp\|Q6P16: | Q6P161 | MRPL54 | 39S ribosomal protein L54, mitochondrial OS=Homo sapi | 1006233.88 | 1147213.88 | 1262891 | 1159650.25 | 0 | 1280207 | 1017130.94 | 578941.12 | 0.628558996 | 5 |
| sp\|Q1502: | Q15021 | NCAPD2 | Condensin complex subunit 1 OS=Homo sapiens (Humar | 7844895.5 | 9020445 | 10311968 | 7713380.5 | 5913319 | 6073304.5 | 4402596 | 5540715 | 0.628532572 | 0 |
| sp\|Q969G( | Q969G6 | RFK | Riboflavin kinase OS=Homo sapiens (Human) GN=RFK PE | 392837.22 | 305659.22 | 104917.44 | 332880.81 | 130303.88 | 228480.38 | 175096.61 | 179648.31 | 0.627943778 | 4 |
| sp\|P84095 | P84095 | RHOG | Rho-related GTP-binding protein RhoG OS=Homo sapien | 4479357.5 | 4938547 | 3016161.5 | 4145640.75 | 3799179.75 | 2632259.75 | 1562474.25 | 2407694 | 0.627369827 | 1 |
| sp\|O7584- | O75844 | ZMPSTE24 | CAAX prenyl protease 1 homolog OS=Homo sapiens (Hu | 5422180.5 | 6835148.5 | 6264574.5 | 7026512 | 4655122.5 | 3228446.5 | 4737574 | 3401139.5 | 0.627134098 | 0 |
| sp\|Q9UG5 | Q9UG56 | PISD | Phosphatidylserine decarboxylase proenzyme, mitochon | 1267930.88 | 1291581.62 | 1161887.75 | 1093607.38 | 1233208 | 0 | 922926.94 | 862114.75 | 0.626842141 | 2 |
| sp\|Q969Z( | Q969Z0 | TBRG4 | FAST kinase domain-containing protein 4 OS=Homo sapi | 4585920 | 4810206 | 3241291.5 | 3599688.25 | 3340901.5 | 2355814.5 | 2495570.5 | 1983498.88 | 0.626699458 | 1 |
| sp\|Q9Y3D! | Q9Y3D8 | AK6 | Adenylate kinase isoenzyme 6 OS=Homo sapiens (Huma | 1241032.12 | 1427036 | 1289590.25 | 1309869.62 | 0 | 1006655.75 | 1168290.25 | 1124524.38 | 0.626379278 | 0 |
| sp\|Q8TF0! | Q8TF05 | PPP4R1 | Serine/threonine-protein phosphatase 4 regulatory subu | 2668186.25 | 2635419.75 | 2782600 | 2618584.75 | 1573108.25 | 1461942.75 | 1712030.62 | 1957465.88 | 0.626312803 | 0 |
| sp\|Q969E: | Q969E2 | SCAMP4 | Secretory carrier-associated membrane protein 4 OS=Hc | 1876257.38 | 2140082.25 | 1776312.62 | 1648298.75 | 1094023.88 | 1184241.75 | 1208555.75 | 1171881.38 | 0.626089697 | 0 |
| sp\|Q9274: | Q92747 | ARPC1A | Actin-related protein 2/3 complex subunit 1A OS=Homo | 5335311 | 4476872 | 1304436.75 | 3903037 | 1751309.25 | 3351555 | 2865522.5 | 1435062.88 | 0.6260762 | 4 |
| sp\|Q1388! | Q13888 | GTF2H2 | General transcription factor IIH subunit 2 OS=Homo sapi | 637582.12 | 784529 | 808483.81 | 558714.5 | 498837.88 | 638878.44 | 0 | 607119.56 | 0.625544037 | 3 |
| sp\|Q1473! | Q14738 | PPP2R5D | Serine/threonine-protein phosphatase 2A 56 kDa regulat | 1701437.75 | 1832380.25 | 1349831.38 | 1603426.75 | 1246293.62 | 1331785.38 | 145953.59 | 1333676.25 | 0.625506586 | 0 |
| sp\|Q9NW\ | Q9NWY4 | HPF1 | Histone PARylation factor 1 OS=Homo sapiens (Human) ( | 343258.38 | 0 | 0 | 290682.47 | 0 | 395972.06 | 0 | 0 | 0.624619884 | 7 |
| sp\|Q1504: | Q15043 | SLC39A14 | Zinc transporter ZIP14 OS=Homo sapiens (Human) GN=S | 14938179 | 18026184 | 14402900 | 13763356 | 11134174 | 8171707.5 | 8081830.5 | 10786417 | 0.624468223 | 0 |
| sp\|Q86UT( | Q86UT6 | NLRX1 | NLR family member X1 OS=Homo sapiens (Human) GN=I | 302404.59 | 337210.94 | 312675.09 | 316306.34 | 221962.56 | 150746.77 | 199857.77 | 219555.27 | 0.62440822 | 0 |
| sp\|O0050! | O00505 | KPNA3 | Importin subunit alpha-4 OS=Homo sapiens (Human) GN | 18644188 | 17814838 | 21085734 | 18546900 | 11919399 | 11867638 | 11790097 | 11922052 | 0.624236427 | 0 |
| sp\|Q9BTE: | Q9BTE3 | MCMBP | Mini-chromosome maintenance complex-binding proteir | 1857610.62 | 1676752 | 2249511 | 1977485.62 | 1749536.88 | 988912.38 | 1681980.5 | 423980.12 | 0.624170294 | 2 |
| sp\|Q9NX4( | Q9NX40 | OCIAD1 | OCIA domain-containing protein 1 OS=Homo sapiens (Hu | 453185.22 | 307805.81 | 214249.41 | 404432.59 | 424835.97 | 0 | 436203.62 | 0 | 0.6240896 | 6 |
| sp\|O0076- | O00764 | PDXK | Pyridoxal kinase OS=Homo sapiens (Human) GN=PDXK P | 10143159 | 7002034.5 | 8292583.5 | 10783196 | 5534769.5 | 5292875 | 6666377.5 | 5062961 | 0.622760272 | 0 |
| sp\|Q9UHD | Q9UHD1 | CHORDC1 | Cysteine and histidine-rich domain-containing protein 1 | 14195454 | 10741618 | 8260885 | 6044166 | 7308340.5 | 8041384 | 5384360 | 3672705.25 | 0.621953857 | 2 |
| sp\|O6056! | O60568 | PLOD3 | Multifunctional procollagen lysine hydroxylase and glyco | 20014716 | 21297818 | 17984132 | 19882530 | 12541995 | 12450998 | 12030877 | 12024093 | 0.619455179 | 0 |
| sp\|Q1431! | Q14315 | FLNC | Filamin-C OS=Homo sapiens (Human) GN=FLNC PE=1 SV: | 72562120 | 78856944 | 82376464 | 71746160 | 49865880 | 42533072 | 53083548 | 43404744 | 0.618204492 | 0 |
| sp\|Q3KQV | Q3KQV9 | UAP1L1 | UDP-N-acetylhexosamine pyrophosphorylase-like proteir | 4211084.5 | 4544782 | 4977239.5 | 5240225 | 858070.69 | 3090967.5 | 2962409.5 | 4768412 | 0.615593524 | 2 |
| sp\|Q9H9Y( | Q9H9Y6 | POLR1B | DNA-directed RNA polymerase I subunit RPA2 OS=Homo | 283779.81 | 902325.06 | 639388.81 | 896953.25 | 305115.22 | 454505.81 | 383388.62 | 532742 | 0.615531429 | 4 |
| sp\|P01892 | P01892 | HLA-A | HLA class I histocompatibility antigen, A-2 alpha chain O: | 41130984 | 38565328 | 38414880 | 34616300 | 25064516 | 24961074 | 22926720 | 20878100 | 0.614364898 | 0 |
| sp\|P08243 | P08243 | ASNS | Asparagine synthetase [glutamine-hydrolyzing] OS=Hom | 18840436 | 23968970 | 16341537 | 20220462 | 13028664 | 11197610 | 12838422 | 11660106 | 0.613883577 | 0 |
| sp\|O7571! | O75718 | CRTAP | Cartilage-associated protein OS=Homo sapiens (Human) | 6014807.5 | 6457736 | 7183754 | 5353707 | 3250412.5 | 4039999.5 | 4854658 | 3190535 | 0.613178818 | 0 |

| | | | | | | | | | | | | | |
|---|---|---|---|---|---|---|---|---|---|---|---|---|---|
| sp\|P51648 | P51648 | ALDH3A2 | Fatty aldehyde dehydrogenase OS=Homo sapiens (Huma | 14513682 | 15365325 | 9455888 | 16098046 | 8875405 | 8200221 | 7739082.5 | 9150938 | **0.612733979** | 0 |
| sp\|Q9300 | Q93009 | USP7 | Ubiquitin carboxyl-terminal hydrolase 7 OS=Homo sapie | 32152840 | 30371080 | 31556038 | 28493568 | 17534690 | 19040276 | 18628772 | 19753942 | **0.611532379** | 0 |
| sp\|O4350 | O43505 | B4GAT1 | Beta-1,4-glucuronyltransferase 1 OS=Homo sapiens (Hu | 1040601.06 | 1434362.88 | 1232294.12 | 1965370.5 | 849558.44 | 623872.62 | 986836.31 | 1008442.19 | **0.611481877** | 0 |
| sp\|Q969X | Q969X1 | TMBIM1 | Protein lifeguard 3 OS=Homo sapiens (Human) GN=TMB | 955736.38 | 1192501.62 | 1192153.25 | 1181459.5 | 0 | 947483.62 | 774946.19 | 1041191 | **0.611170285** | 1 |
| sp\|P18074 | P18074 | ERCC2 | General transcription and DNA repair factor IIH helicase s | 1955667 | 1834074.5 | 1294140.88 | 1391575.88 | 872911.75 | 923634.44 | 1166849.12 | 981291.81 | **0.609174974** | 0 |
| sp\|Q9BV5 | Q9BV57 | ADI1 | 1,2-dihydroxy-3-keto-5-methylthiopentene dioxygenase | 5708006.5 | 5722267.5 | 5399134.5 | 6665773.5 | 3513640.5 | 3574401.25 | 3162856 | 4054269 | **0.608855328** | 0 |
| sp\|Q96K7 | Q96K76 | USP47 | Ubiquitin carboxyl-terminal hydrolase 47 OS=Homo sapi | 1333696.75 | 1194731.25 | 1240575.62 | 1313435.88 | 556527.5 | 968469.88 | 664803.75 | 899691.38 | **0.607875905** | 0 |
| sp\|Q7LGA | Q7LGA3 | HS2ST1 | Heparan sulfate 2-O-sulfotransferase 1 OS=Homo sapier | 3911172 | 4243016 | 4621879.5 | 3785906.5 | 2230869 | 2756062 | 2340447.5 | 2680664 | **0.604278361** | 0 |
| sp\|Q96C8 | Q96C86 | DCPS | m7GpppX diphosphatase OS=Homo sapiens (Human) GN | 436339.31 | 1250593 | 354956.25 | 598436.94 | 407081.84 | 378120.88 | 393613.38 | 416550.38 | **0.604230986** | 4 |
| sp\|P31151 | P31151 | S100A7 | Protein S100-A7 OS=Homo sapiens (Human) GN=S100A7 | 1178719.75 | 4391272.5 | 0 | 1752162.38 | 890214.62 | 0 | 1318234.25 | 2209516 | **0.603369513** | 6.5 |
| sp\|Q1654 | Q16543 | CDC37 | Hsp90 co-chaperone Cdc37 OS=Homo sapiens (Human) | 15912719 | 14469581 | 23291272 | 16224993 | 13034290 | 11176370 | 7727290 | 10211071 | **0.603002665** | 0 |
| sp\|Q96QE | Q96QE5 | TEFM | Transcription elongation factor, mitochondrial OS=Homo | 522722.84 | 625345.5 | 419039.41 | 402593.59 | 466681.28 | 354070.84 | 0 | 365593.09 | **0.602297001** | 2 |
| sp\|Q1341 | Q13418 | ILK | Integrin-linked protein kinase OS=Homo sapiens (Humar | 2568415 | 2236094.5 | 1985526 | 1227658.5 | 1205278.5 | 1075293.12 | 1564964 | 970758.31 | **0.600708125** | 1 |
| sp\|Q8N3C | Q8N3C0 | ASCC3 | Activating signal cointegrator 1 complex subunit 3 OS=H | 1632129.12 | 1545217.5 | 2329451 | 1599890.38 | 1595627.5 | 1481631.75 | 266262.34 | 918100.62 | **0.599663614** | 1 |
| sp\|O1525 | O15254 | ACOX3 | Peroxisomal acyl-coenzyme A oxidase 3 OS=Homo sapie | 283763.47 | 315835.69 | 395826.59 | 0 | 298057.06 | 0 | 0 | 298642.38 | **0.599441435** | 5 |
| sp\|Q8TB3 | Q8TB37 | NUBPL | Iron-sulfur protein NUBPL OS=Homo sapiens (Human) G | 2269957.25 | 2976265.5 | 2698843.25 | 2486409 | 1744952.12 | 1969248.38 | 1978404.62 | 545787 | **0.598035476** | 0 |
| sp\|Q9NZQ | Q9NZQ3 | NCKIPSD | NCK-interacting protein with SH3 domain OS=Homo sapi | 0 | 437579.47 | 439578.56 | 536625 | 418814.12 | 0 | 0 | 426234.22 | **0.597721377** | 3 |
| sp\|Q5VYK | Q5VYK3 | ECPAS | Proteasome adapter and scaffold protein ECM29 OS=Ho | 9273790 | 10672137 | 11772617 | 11894861 | 3805571.75 | 5818599.5 | 7453289 | 8978172 | **0.597422564** | 0 |
| sp\|Q1622 | Q16222 | UAP1 | UDP-N-acetylhexosamine pyrophosphorylase OS=Homo | 5225346.5 | 5209315 | 5275594.5 | 6351064 | 1144781 | 3669241.75 | 3308487.5 | 5050065.5 | **0.597089193** | 0 |
| sp\|P13645 | P13645 | KRT10 | Keratin, type I cytoskeletal 10 OS=Homo sapiens (Humar | 56087152 | 489580192 | 76171240 | 78493352 | 98673872 | 99555024 | 115023624 | 102601032 | **0.593794929** | 4 |
| sp\|P28340 | P28340 | POLD1 | DNA polymerase delta catalytic subunit OS=Homo sapie | 4049123 | 4438887 | 4600509 | 5501467 | 2801278 | 3585919.5 | 1537956.5 | 3062206.5 | **0.591036513** | 0 |
| sp\|P52434 | P52434 | POLR2H | DNA-directed RNA polymerases I, II, and III subunit RPAB | 288386.91 | 230289.41 | 76370.72 | 3704635 | 931692.25 | 314581.38 | 130142.3 | 1162579.75 | **0.590507776** | 6 |
| sp\|Q0040 | Q00403 | GTF2B | Transcription initiation factor IIB OS=Homo sapiens (Hun | 773018.56 | 678518.94 | 588133.62 | 796295.62 | 738099.31 | 419970.34 | 0 | 516413.91 | **0.590445415** | 2 |
| sp\|Q9BV4 | Q9BV44 | THUMPD3 | THUMP domain-containing protein 3 OS=Homo sapiens | 461204.22 | 481029.66 | 0 | 637291.69 | 562165.5 | 0 | 370238.62 | 0 | **0.590306442** | 5 |
| sp\|Q9UKV | Q9UKV5 | AMFR | E3 ubiquitin-protein ligase AMFR OS=Homo sapiens (Hu | 127296.86 | 102969.74 | 154188.7 | 118651.73 | 141391.05 | 83417.05 | 0 | 72004.07 | **0.589958304** | 3 |
| sp\|Q9BV3 | Q9BV38 | WDR18 | WD repeat-containing protein 18 OS=Homo sapiens (Hu | 1903418.5 | 1668933 | 3182774.5 | 3506222.5 | 1003495.38 | 1471133.62 | 2609996.5 | 964127.75 | **0.589469625** | 2 |
| sp\|Q9Y28 | Q9Y289 | SLC5A6 | Sodium-dependent multivitamin transporter OS=Homo s | 1773319.75 | 1849679.25 | 2159359.25 | 1673089.25 | 1618421.75 | 1534699.25 | 1230081.25 | 0 | **0.587919404** | 0 |
| sp\|P11388 | P11388 | TOP2A | DNA topoisomerase 2-alpha OS=Homo sapiens (Human) | 18780986 | 20584902 | 39931556 | 26109680 | 15665502 | 14859672 | 14274302 | 17118840 | **0.5874206** | 0 |
| sp\|Q86Y5 | Q86Y56 | DNAAF5 | Dynein assembly factor 5, axonemal OS=Homo sapiens ( | 4704538 | 5438402.5 | 6594639.5 | 7528023 | 1855549.38 | 5487527 | 3513210.75 | 3385675 | **0.58691977** | 2 |
| sp\|P42574 | P42574 | CASP3 | Caspase-3 OS=Homo sapiens (Human) GN=CASP3 PE=1 S | 5412106.5 | 3867123.25 | 5182606 | 3323360.25 | 3020338.25 | 0 | 4056919.5 | 3355079 | **0.586574179** | 3 |
| sp\|Q9GZR | Q9GZR7 | DDX24 | ATP-dependent RNA helicase DDX24 OS=Homo sapiens ( | 1469531.12 | 1184899.75 | 1267454 | 1362700.38 | 587012.44 | 849157.19 | 891008.31 | 767052.44 | **0.585520005** | 0 |
| sp\|O1473 | O14735 | CDIPT | CDP-diacylglycerol--inositol 3-phosphatidyltransferase C | 1357516.25 | 1437535.38 | 1371724.75 | 1356907.75 | 1527069.5 | 418305.59 | 923353.94 | 365406.94 | **0.585503424** | 4 |
| sp\|O7515 | O75153 | CLUH | Clustered mitochondria protein homolog OS=Homo sapi | 11468904 | 10837798 | 11773674 | 9436496 | 6439644.5 | 6853426.5 | 5158761 | 7013510 | **0.585183191** | 0 |
| sp\|O4384 | O43847 | NRDC | Nardilysin OS=Homo sapiens (Human) GN=NRDC PE=1 S | 3708414.5 | 3251083.25 | 4024075.75 | 4104276.5 | 3482003.25 | 2750115.5 | 1343585 | 1248062.62 | **0.584825961** | 1 |
| sp\|P40938 | P40938 | RFC3 | Replication factor C subunit 3 OS=Homo sapiens (Humar | 1429451.25 | 1102961.38 | 2323153.5 | 1490736.62 | 634340.5 | 1424047.75 | 836605.12 | 814110.62 | **0.584451158** | 1 |
| sp\|Q1455 | Q14558 | PRPSAP1 | Phosphoribosyl pyrophosphate synthase-associated prot | 10538943 | 10201313 | 10475510 | 11266403 | 6801514 | 5896314.5 | 5845908 | 6253808 | **0.583716535** | 0 |
| sp\|Q9ULT | Q9ULT8 | HECTD1 | E3 ubiquitin-protein ligase HECTD1 OS=Homo sapiens (H | 3246104.5 | 3349699 | 4942891 | 3939214.5 | 2512323.75 | 1981371.88 | 2702903 | 1811448 | **0.581993771** | 0 |
| sp\|Q9299 | Q92990 | GLMN | Glomulin OS=Homo sapiens (Human) GN=GLMN PE=1 S | 189895.98 | 468370 | 405736.53 | 1200665.38 | 253464.03 | 428201.38 | 272164.56 | 362544.56 | **0.581266037** | 5 |
| sp\|Q9GZU | Q9GZU2 | PEG3 | Paternally-expressed gene 3 protein OS=Homo sapiens ( | 36431120 | 33147644 | 31067776 | 27603368 | 20753456 | 19835590 | 18098436 | 15586921 | **0.579138061** | 0 |
| sp\|Q7Z4V | Q7Z4V5 | HDGFL2 | Hepatoma-derived growth factor-related protein 2 OS=H | 5416236 | 9093704 | 7006656.5 | 5279888 | 2993741 | 3673823.5 | 5372408 | 3440514.5 | **0.577705893** | 1 |
| sp\|O7544 | O75448 | MED24 | Mediator of RNA polymerase II transcription subunit 24 | 512321.56 | 318002.66 | 430605.53 | 611217.38 | 384113.09 | 359278.16 | 0 | 336250.44 | **0.576686347** | 3 |
| sp\|Q8WU | Q8WUA2 | PPIL4 | Peptidyl-prolyl cis-trans isomerase-like 4 OS=Homo sapie | 1060011.25 | 925611 | 578021.19 | 995226 | 604376.44 | 751833.88 | 0 | 688602.31 | **0.574568038** | 3 |
| sp\|Q9GZL | Q9GZL7 | WDR12 | Ribosome biogenesis protein WDR12 OS=Homo sapiens | 8374901.5 | 7901121 | 4681626 | 9306189 | 4051368.5 | 5766342 | 3038672.5 | 4517426 | **0.574078188** | 1 |
| sp\|A0AVT | A0AVT1 | UBA6 | Ubiquitin-like modifier-activating enzyme 6 OS=Homo sa | 9019639 | 11630988 | 9121114 | 10290996 | 4915378 | 6425801 | 4007917.5 | 7640115 | **0.573830278** | 0 |
| sp\|P26358 | P26358 | DNMT1 | DNA (cytosine-5)-methyltransferase 1 OS=Homo sapiens | 7026095 | 3347052.5 | 5891045 | 4085603.5 | 2613141 | 4068536.5 | 2382139 | 2601279 | **0.573229112** | 1 |

| | | | | | | | | | | | | |
|---|---|---|---|---|---|---|---|---|---|---|---|---|
| sp\|P13693 | P13693 | TPT1 | Translationally-controlled tumor protein OS=Homo sapie | 16797654 | 16846142 | 19135656 | 18359604 | 9330478 | 12250817 | 7958285 | 11134857 | **0.571759583** | 0 |
| sp\|Q9283 | Q92835 | INPP5D | Phosphatidylinositol 3,4,5-trisphosphate 5-phosphatase | 473222.66 | 508102.31 | 364571.28 | 276936.66 | 291157.19 | 0 | 403384.66 | 228809.06 | **0.568974726** | 3 |
| sp\|Q1499 | Q14997 | PSME4 | Proteasome activator complex subunit 4 OS=Homo sapie | 8489039 | 6722042 | 7401766 | 8257730 | 3189709.5 | 5227106.5 | 5107012.5 | 4027646.5 | **0.568550274** | 0 |
| sp\|Q6P3X | Q6P3X3 | TTC27 | Tetratricopeptide repeat protein 27 OS=Homo sapiens (I | 1378575.75 | 1416896.5 | 1648757.75 | 1523560.75 | 1173797.38 | 1222478.88 | 84388.72 | 911025.94 | **0.568332749** | 0 |
| sp\|P86397 | P86397 | HTD2 | Hydroxyacyl-thioester dehydratase type 2, mitochondrial | 153338.16 | 211030.92 | 112812.46 | 177437.36 | 100096.09 | 103315.74 | 82258.34 | 85507.7 | **0.567013678** | 0 |
| sp\|Q9NVU | Q9NVU7 | SDAD1 | Protein SDA1 homolog OS=Homo sapiens (Human) GN=S | 3109146.5 | 3024995.5 | 3982932.5 | 3423809 | 2445255.25 | 1201706.75 | 2830751 | 1195910.38 | **0.566700347** | 0 |
| sp\|Q9UET | Q9UET6 | FTSJ1 | Putative tRNA (cytidine(32)/guanosine(34)-2'-O)-methylt | 553687.56 | 534792.5 | 455241.12 | 610955.38 | 0 | 352989.81 | 499431.25 | 368428.78 | **0.566604688** | 1 |
| sp\|Q9UH6 | Q9UH65 | SWAP70 | Switch-associated protein 70 OS=Homo sapiens (Human | 3188650.5 | 3377247 | 4137871.25 | 2538154.5 | 1648005 | 1498626.75 | 1759679.38 | 2588059.75 | **0.565957885** | 1 |
| sp\|Q9Y52 | Q9Y520 | PRRC2C | Protein PRRC2C OS=Homo sapiens (Human) GN=PRRC2C | 2054987.75 | 986314.88 | 1165393.75 | 0 | 500847.44 | 679453.81 | 545015.19 | 647659.44 | **0.564094878** | 4 |
| sp\|P11908 | P11908 | PRPS2 | Ribose-phosphate pyrophosphokinase 2 OS=Homo sapie | 12730218 | 10010100 | 11209776 | 10054986 | 6202084.5 | 7789412.5 | 5076535.5 | 5730720.5 | **0.563542959** | 0 |
| sp\|P41567 | P41567 | EIF1 | Eukaryotic translation initiation factor 1 OS=Homo sapie | 8763047 | 7217261 | 3804605.25 | 7511610 | 3656879.75 | 3991477.5 | 3749223.5 | 3975543 | **0.56318981** | 2 |
| sp\|Q9H5Q | Q9H5Q4 | TFB2M | Dimethyladenosine transferase 2, mitochondrial OS=Hoi | 1774140.25 | 1633165 | 1663790 | 1902246.75 | 1007293.5 | 1303677.38 | 361264.91 | 1240814.25 | **0.561144146** | 0 |
| sp\|P01130 | P01130 | LDLR | Low-density lipoprotein receptor OS=Homo sapiens (Hu | 2435564.5 | 1684276.12 | 1752523.88 | 1468393 | 967128.25 | 677757.69 | 1253719.25 | 1216719.5 | **0.560613083** | 0 |
| sp\|P36639 | P36639 | NUDT1 | 7,8-dihydro-8-oxoguanine triphosphatase OS=Homo sap | 522471.06 | 0 | 422741 | 365184.16 | 344264.38 | 389462.06 | 0 | 0 | **0.559927165** | 4 |
| sp\|O4365 | O43657 | TSPAN6 | Tetraspanin-6 OS=Homo sapiens (Human) GN=TSPAN6 P | 1836501.75 | 3209617.25 | 2878096.75 | 2700721.25 | 1588508.62 | 1136598.62 | 2132961.75 | 1085212.75 | **0.55937101** | 1 |
| sp\|Q8TBE | Q8TBE9 | NANP | N-acylneuraminate-9-phosphatase OS=Homo sapiens (H | 367402.5 | 393298.06 | 457321.53 | 377126.91 | 306623.16 | 292845.62 | 291645.66 | 0 | **0.558640252** | 0 |
| sp\|O0062 | O00629 | KPNA4 | Importin subunit alpha-3 OS=Homo sapiens (Human) GN | 9349741 | 9700531 | 8385595.5 | 11919882 | 6111125.5 | 3703085 | 5199102.5 | 6964006 | **0.558427149** | 0 |
| sp\|P33947 | P33947 | KDELR2 | ER lumen protein-retaining receptor 2 OS=Homo sapien | 304273.59 | 249953.55 | 427124 | 0 | 133683.55 | 0 | 101982.5 | 311301.84 | **0.557362057** | 5.5 |
| sp\|Q6PI48 | Q6PI48 | DARS2 | Aspartate--tRNA ligase, mitochondrial OS=Homo sapiens | 3921323 | 3225205 | 4165439 | 3146685.5 | 2097947.5 | 1172686.5 | 1833277.5 | 2954758 | **0.55735965** | 0 |
| sp\|Q9Y30 | Q9Y305 | ACOT9 | Acyl-coenzyme A thioesterase 9, mitochondrial OS=Hom | 5604297 | 7224541 | 4848099 | 4813294 | 3858716 | 3737319 | 2380345.75 | 2537995.75 | **0.556436103** | 0 |
| sp\|Q9957 | Q99570 | PIK3R4 | Phosphoinositide 3-kinase regulatory subunit 4 OS=Hom | 634615 | 766208.19 | 875932.38 | 691566.94 | 143909.77 | 594605.44 | 418876.62 | 491734.38 | **0.555575145** | 0 |
| sp\|Q9BXW | Q9BXW9 | FANCD2 | Fanconi anemia group D2 protein OS=Homo sapiens (Hu | 344574.09 | 381439.09 | 383173.66 | 444372.03 | 194585.8 | 239634.27 | 205282 | 221681.92 | **0.554329808** | 0 |
| sp\|P27105 | P27105 | STOM | Erythrocyte band 7 integral membrane protein OS=Hom | 1006218.69 | 3761164 | 2178583.75 | 3567740 | 1677413.75 | 465644.75 | 1590392.38 | 2082350.25 | **0.553163736** | 3 |
| sp\|O1522 | O15228 | GNPAT | Dihydroxyacetone phosphate acyltransferase OS=Homo | 241859.67 | 322021.97 | 146441.95 | 291386.16 | 150582.64 | 146521.53 | 131532.95 | 125337.21 | **0.553028789** | 2 |
| sp\|P08195 | P08195 | SLC3A2 | 4F2 cell-surface antigen heavy chain OS=Homo sapiens ( | 150806912 | 153629024 | 149530416 | 149879648 | 84683768 | 82743280 | 81601304 | 84750400 | **0.552754762** | 0 |
| sp\|Q9Y6D | Q9Y6D9 | MAD1L1 | Mitotic spindle assembly checkpoint protein MAD1 OS=I | 1326596.88 | 1418944.75 | 2008223.5 | 1244294.5 | 958409.31 | 673968.62 | 1061546.88 | 613186.19 | **0.551363475** | 0 |
| sp\|Q1453 | Q14534 | SQLE | Squalene monooxygenase OS=Homo sapiens (Human) G | 605802.75 | 542148.38 | 565574.5 | 520331.75 | 378545.28 | 0 | 391731.47 | 461305.22 | **0.551325246** | 0 |
| sp\|Q9ULX | Q9ULX3 | NOB1 | RNA-binding protein NOB1 OS=Homo sapiens (Human) ( | 4806573.5 | 5215462 | 6715218 | 5506101.5 | 1177523.25 | 3720428 | 3530333.5 | 3822605 | **0.550766274** | 0 |
| sp\|Q53EP | Q53EP0 | FNDC3B | Fibronectin type III domain-containing protein 3B OS=Hc | 1105862.25 | 594623.94 | 1040033 | 1019162.12 | 709878.75 | 402453.38 | 344704.81 | 608476.38 | **0.549385214** | 2 |
| sp\|P98194 | P98194 | ATP2C1 | Calcium-transporting ATPase type 2C member 1 OS=Hon | 606376.5 | 593226.88 | 595178.62 | 798647.75 | 345725.75 | 556902.94 | 180231.33 | 341821.94 | **0.5493428** | 0 |
| sp\|P23368 | P23368 | ME2 | NAD-dependent malic enzyme, mitochondrial OS=Homo | 1240614.75 | 959254.5 | 195511.47 | 1074376.75 | 465192.25 | 870208.81 | 365164.56 | 203407.56 | **0.548733794** | 4 |
| sp\|P62070 | P62070 | RRAS2 | Ras-related protein R-Ras2 OS=Homo sapiens (Human) ( | 2209509 | 1881018.25 | 2348770.75 | 2364724.5 | 1615238 | 1180065.5 | 886872.44 | 1139846.88 | **0.547706781** | 0 |
| sp\|Q1047 | Q10471 | GALNT2 | Polypeptide N-acetylgalactosaminyltransferase 2 OS=Ho | 11121428 | 11843253 | 9744134 | 9890820 | 6637459 | 5301084 | 3617276 | 7771228 | **0.547587955** | 0 |
| sp\|Q68D8 | Q68D85 | NCR3LG1 | Natural cytotoxicity triggering receptor 3 ligand 1 OS=Hc | 1005984.25 | 1186125.5 | 1145864.25 | 1228706.5 | 794024.81 | 486616.88 | 669333.38 | 550061.94 | **0.547451702** | 0 |
| sp\|Q8NCA | Q8NCA5 | FAM98A | Protein FAM98A OS=Homo sapiens (Human) GN=FAM98 | 1897128.5 | 6048053 | 3061769 | 6528352 | 2670685.5 | 2139549.75 | 2188145 | 2526340.25 | **0.543174006** | 4 |
| sp\|O1530 | O15305 | PMM2 | Phosphomannomutase 2 OS=Homo sapiens (Human) GN | 5327686 | 4795229 | 4760420 | 6725572 | 2949995.25 | 3042974.5 | 3448508.5 | 2244548 | **0.54079673** | 0 |
| sp\|Q8TC0 | Q8TC07 | TBC1D15 | TBC1 domain family member 15 OS=Homo sapiens (Hum | 521263.91 | 369690.12 | 526437.75 | 997571.88 | 273326.75 | 338443.31 | 280703.41 | 412934.44 | **0.540549712** | 1 |
| sp\|P23921 | P23921 | RRM1 | Ribonucleoside-diphosphate reductase large subunit OS | 9267612 | 7967805 | 13124496 | 10337463 | 4611356 | 6469422.5 | 5803426.5 | 5066472.5 | **0.539363459** | 0 |
| sp\|P27707 | P27707 | DCK | Deoxycytidine kinase OS=Homo sapiens (Human) GN=D( | 508680.38 | 0 | 606341.12 | 535162 | 0 | 469743.72 | 420246.03 | 0 | **0.539327747** | 3 |
| sp\|P04899 | P04899 | GNAI2 | Guanine nucleotide-binding protein G(i) subunit alpha-2 | 3330541.5 | 3700115 | 3630368.5 | 2460331.5 | 1542403.88 | 2210474.5 | 872613.44 | 2425153.5 | **0.53734119** | 0 |
| sp\|P13639 | P13639 | EEF2 | Elongation factor 2 OS=Homo sapiens (Human) GN=EEF2 | 849825408 | 886948736 | 1125279360 | 862916160 | 532977632 | 489952032 | 503207648 | 469109952 | **0.535641212** | 0 |
| sp\|Q9NVV | Q9NVV4 | MTPAP | Poly(A) RNA polymerase, mitochondrial OS=Homo sapie | 4707357 | 4048169.25 | 1428421.75 | 3710212 | 2052100.25 | 1811658.75 | 2094744.12 | 1479804.75 | **0.535354989** | 4 |
| sp\|Q8IUR | Q8IUR0 | TRAPPC5 | Trafficking protein particle complex subunit 5 OS=Homo | 299252.09 | 349440.75 | 381916.38 | 284114.62 | 0 | 355611 | 343661.38 | 0 | **0.531877767** | 5 |
| sp\|P60602 | P60602 | ROMO1 | Reactive oxygen species modulator 1 OS=Homo sapiens | 1270591.75 | 1385268.75 | 1466718.75 | 1308630.12 | 769414.5 | 800873.56 | 693546.25 | 616514.19 | **0.530332805** | 0 |
| sp\|O4376 | O43765 | SGTA | Small glutamine-rich tetratricopeptide repeat-containing | 15661828 | 15548346 | 13086128 | 14457576 | 6429843 | 9290970 | 8366993 | 7016797 | **0.529405106** | 0 |

| | | | | | | | | | | | | | |
|---|---|---|---|---|---|---|---|---|---|---|---|---|---|
| sp\|Q1418 | Q14186 | TFDP1 | Transcription factor Dp-1 OS=Homo sapiens (Human) GN | 613207.12 | 319140.56 | 384771.22 | 379985.88 | 190736.44 | 242304.03 | 255395.5 | 208782.69 | **0.528676055** | **0** |
| sp\|Q5UIP0 | Q5UIP0 | RIF1 | Telomere-associated protein RIF1 OS=Homo sapiens (Hu | 5464809.5 | 3942584.5 | 7622612 | 5408602.5 | 2703900.5 | 4377344 | 2422641.25 | 2347038 | **0.528148782** | **1** |
| sp\|Q9Y2Z4 | Q9Y2Z4 | YARS2 | Tyrosine--tRNA ligase, mitochondrial OS=Homo sapiens ( | 6568024 | 18053724 | 16780722 | 17400396 | 7400516.5 | 7441198 | 7963024 | 8204348.5 | **0.527339722** | **4** |
| sp\|Q7Z4H | Q7Z4H7 | HAUS6 | HAUS augmin-like complex subunit 6 OS=Homo sapiens | 1126893 | 987954.94 | 1164088.38 | 727033.19 | 615396.88 | 769522.31 | 529099.31 | 196074.09 | **0.526737057** | **1** |
| sp\|P10646 | P10646 | TFPI | Tissue factor pathway inhibitor OS=Homo sapiens (Huma | 6561906 | 6643786 | 2979076.5 | 7148337 | 2013022.38 | 3826370.5 | 2241110 | 4065265.75 | **0.520538024** | **2** |
| sp\|Q9NPI6 | Q9NPI6 | DCP1A | mRNA-decapping enzyme 1A OS=Homo sapiens (Human | 591717.06 | 634329.62 | 747540.06 | 602329.19 | 634463.88 | 0 | 0 | 704617.81 | **0.519846814** | **6** |
| sp\|P52292 | P52292 | KPNA2 | Importin subunit alpha-1 OS=Homo sapiens (Human) GN | 44686916 | 36794916 | 75948832 | 40622764 | 30312354 | 22656036 | 26386954 | 23314232 | **0.51839333** | **0** |
| sp\|Q6YHU | Q6YHU6 | THADA | Thyroid adenoma-associated protein OS=Homo sapiens | 444270.5 | 496277 | 511313.84 | 469675.38 | 250390.06 | 92637.18 | 347072.12 | 305695.56 | **0.51822841** | **0** |
| sp\|Q9NRG | Q9NRG7 | SDR39U1 | Epimerase family protein SDR39U1 OS=Homo sapiens (H | 525289.06 | 558580.19 | 535597.88 | 761044.44 | 304803.38 | 314369.22 | 331292.59 | 280230.94 | **0.51698809** | **0** |
| sp\|Q9985 | Q99856 | ARID3A | AT-rich interactive domain-containing protein 3A OS=Ho | 14711373 | 14665136 | 16159298 | 16111656 | 9172704 | 6221197.5 | 5828045 | 10588823 | **0.516011008** | **0** |
| sp\|Q9257 | Q92572 | AP3S1 | AP-3 complex subunit sigma-1 OS=Homo sapiens (Huma | 480109.75 | 2778067.5 | 4919238 | 2552513.75 | 2819043.75 | 334338.88 | 2359345.5 | 0 | **0.513771166** | **4** |
| sp\|Q8IX18 | Q8IX18 | DHX40 | Probable ATP-dependent RNA helicase DHX40 OS=Homo | 1222257.88 | 1190726.38 | 1274818.5 | 1244780.12 | 163003.08 | 772295.38 | 844756.94 | 748581.38 | **0.512639492** | **0** |
| sp\|P42224 | P42224 | STAT1 | Signal transducer and activator of transcription 1-alpha/b | 12807733 | 11196102 | 13555261 | 11269576 | 7223336.5 | 6731875.5 | 5993548.5 | 4994392.5 | **0.510830051** | **0** |
| sp\|O6057 | O60573 | EIF4E2 | Eukaryotic translation initiation factor 4E type 2 OS=Hon | 783180.69 | 1169670.25 | 1601307 | 1627137 | 656799.69 | 537499 | 660534.06 | 788348.44 | **0.51013911** | **1** |
| sp\|P09382 | P09382 | LGALS1 | Galectin-1 OS=Homo sapiens (Human) GN=LGALS1 PE=1 | 10486776 | 22855588 | 22159192 | 8420959 | 5084225 | 13302905 | 10836407 | 3320586.5 | **0.509118321** | **4** |
| sp\|O7536 | O75369 | FLNB | Filamin-B OS=Homo sapiens (Human) GN=FLNB PE=1 SV | 543341696 | 548454784 | 581111616 | 455204800 | 282733664 | 245495696 | 293837120 | 250635472 | **0.504062521** | **0** |
| sp\|O1539 | O15397 | IPO8 | Importin-8 OS=Homo sapiens (Human) GN=IPO8 PE=1 S | 4721424 | 4715181.5 | 730845.38 | 3259897 | 1138123.12 | 2347086 | 537359.44 | 2743568.5 | **0.503907184** | **3** |
| sp\|P58107 | P58107 | EPPK1 | Epiplakin OS=Homo sapiens (Human) GN=EPPK1 PE=1 S | 69151328 | 52320180 | 53116256 | 54155804 | 26961042 | 28928644 | 30041768 | 29330990 | **0.503893705** | **0** |
| sp\|O6034 | O60346 | PHLPP1 | PH domain leucine-rich repeat-containing protein phospl | 255077.19 | 253766.55 | 229879.14 | 217391.78 | 118048.3 | 114644.62 | 127890.27 | 120728.95 | **0.503404205** | **0** |
| sp\|P61960 | P61960 | UFM1 | Ubiquitin-fold modifier 1 OS=Homo sapiens (Human) GN | 809515.06 | 762365.81 | 0 | 755613.75 | 689166.88 | 482459.75 | 0 | 0 | **0.503385323** | **3** |
| sp\|Q96LJ7 | Q96LJ7 | DHRS1 | Dehydrogenase/reductase SDR family member 1 OS=Ho | 2553551.5 | 2224121.25 | 2816590 | 2726767.25 | 1048705.38 | 1416268.75 | 1287663.5 | 1425219.75 | **0.501680295** | **0** |
| sp\|Q9HC0 | Q9HC07 | TMEM165 | Transmembrane protein 165 OS=Homo sapiens (Human) | 4189672 | 4746691 | 4901360 | 4393790 | 2877159.5 | 2054701.5 | 2782083.75 | 1415991.88 | **0.500777781** | **0** |
| sp\|Q5H9R | Q5H9R7 | PPP6R3 | Serine/threonine-protein phosphatase 6 regulatory subu | 3367180 | 5142832.5 | 5486514 | 4293808 | 2645800.5 | 2348866.25 | 1314534.5 | 2841418.25 | **0.500298095** | **0** |
| sp\|P51452 | P51452 | DUSP3 | Dual specificity protein phosphatase 3 OS=Homo sapiens | 6774215.5 | 6200326 | 6351996.5 | 6326669 | 3387347 | 2761326.5 | 2919554.5 | 3762842.5 | **0.50017413** | **0** |
| sp\|Q7Z3C | Q7Z3C6 | ATG9A | Autophagy-related protein 9A OS=Homo sapiens (Humai | 1313296.25 | 954250.12 | 896003.19 | 2193186.25 | 634785.12 | 667045.62 | 527712.06 | 847512.62 | **0.499754984** | **0** |
| sp\|Q0079 | Q00796 | SORD | Sorbitol dehydrogenase OS=Homo sapiens (Human) GN= | 39362304 | 38041568 | 33703268 | 36166468 | 18797136 | 20056092 | 17657158 | 16947168 | **0.498782878** | **0** |
| sp\|Q9NPQ | Q9NPQ8 | RIC8A | Synembryn-A OS=Homo sapiens (Human) GN=RIC8A PE= | 1441744.25 | 1576886.25 | 2145430 | 1443347.5 | 565385.75 | 835709.75 | 759455.25 | 1128614.75 | **0.497799667** | **0** |
| sp\|P02753 | P02753 | RBP4 | Retinol-binding protein 4 OS=Homo sapiens (Human) GN | 12004500 | 13202914 | 11329716 | 13244775 | 6076951.5 | 4328975 | 7495109 | 6873433.5 | **0.497660124** | **0** |
| sp\|P11166 | P11166 | SLC2A1 | Solute carrier family 2, facilitated glucose transporter me | 82950768 | 93611344 | 75058304 | 84206328 | 48909872 | 48580628 | 29493046 | 38495560 | **0.492751423** | **0** |
| sp\|P37059 | P37059 | HSD17B2 | Estradiol 17-beta-dehydrogenase 2 OS=Homo sapiens (H | 548857.75 | 802267.62 | 704279.25 | 659698.12 | 0 | 398978.66 | 383775.78 | 542595.81 | **0.488139999** | **0** |
| sp\|Q1682 | Q16822 | PCK2 | Phosphoenolpyruvate carboxykinase [GTP], mitochondri | 120748896 | 129852416 | 104527872 | 119293736 | 56902824 | 52352752 | 60217876 | 60850196 | **0.485481705** | **0** |
| sp\|Q0165 | Q01650 | SLC7A5 | Large neutral amino acids transporter small subunit 1 OS | 11035589 | 14008644 | 15839688 | 11503494 | 7225027 | 6338944 | 5499581.5 | 6338920.5 | **0.484896478** | **0** |
| sp\|Q9H6S | Q9H6S3 | EPS8L2 | Epidermal growth factor receptor kinase substrate 8-like | 27074966 | 28272836 | 31107334 | 26568768 | 12752338 | 13787138 | 13673544 | 14413086 | **0.483314627** | **0** |
| sp\|P33527 | P33527 | ABCC1 | Multidrug resistance-associated protein 1 OS=Homo sap | 1139468.88 | 901081.38 | 1612572 | 1679515.5 | 1270645.25 | 720807.69 | 60945.5 | 522043.56 | **0.482770838** | **2** |
| sp\|P46821 | P46821 | MAP1B | Microtubule-associated protein 1B OS=Homo sapiens (H | 1505540.5 | 1517658.5 | 1233462.62 | 489745.09 | 164461.7 | 489366.56 | 1177350.75 | 459215.31 | **0.482553321** | **1** |
| sp\|Q96IV0 | Q96IV0 | NGLY1 | Peptide-N(4)-(N-acetyl-beta-glucosaminyl)asparagine am | 650478.31 | 0 | 751115.25 | 678936.25 | 521469.16 | 0 | 482279 | 0 | **0.482448343** | **3** |
| sp\|Q96S5 | Q96S59 | RANBP9 | Ran-binding protein 9 OS=Homo sapiens (Human) GN=R | 689171.75 | 431094.56 | 849143.88 | 572211.25 | 0 | 0 | 585320.88 | 639179.06 | **0.481779041** | **4** |
| sp\|Q63HN | Q63HN8 | RNF213 | E3 ubiquitin-protein ligase RNF213 OS=Homo sapiens (H | 6148996.5 | 6145423.5 | 5566351.5 | 5797114 | 2809473.5 | 2413116.25 | 2201210.75 | 3849545.5 | **0.476515367** | **0** |
| sp\|P07307 | P07307 | ASGR2 | Asialoglycoprotein receptor 2 OS=Homo sapiens (Humar | 1099253 | 528889.94 | 0 | 1568414.75 | 253147.22 | 1244192.75 | 0 | 0 | **0.468422633** | **5** |
| sp\|Q9BQA | Q9BQA1 | WDR77 | Methylosome protein 50 OS=Homo sapiens (Human) GN | 14358896 | 16055612 | 15385650 | 13788929 | 6599625 | 6117519 | 6335772 | 8740132 | **0.466411711** | **0** |
| sp\|Q0632 | Q06323 | PSME1 | Proteasome activator complex subunit 1 OS=Homo sapie | 90094392 | 88367288 | 81252576 | 91233768 | 44652032 | 34055556 | 41615092 | 43295164 | **0.466216741** | **0** |
| sp\|P27708 | P27708 | CAD | CAD protein OS=Homo sapiens (Human) GN=CAD PE=1 S | 51467000 | 44194260 | 45432232 | 39399248 | 19271902 | 24983402 | 20561250 | 19330902 | **0.466209644** | **0** |
| sp\|Q86XA | Q86XA9 | HEATR5A | HEAT repeat-containing protein 5A OS=Homo sapiens (H | 902550.25 | 911342 | 627297.12 | 782700.5 | 385911.03 | 440198.19 | 340140.28 | 332870.06 | **0.465003341** | **0** |
| sp\|Q5TFE | Q5TFE4 | NT5DC1 | 5'-nucleotidase domain-containing protein 1 OS=Homo s | 7688923.5 | 5545434.5 | 8157330 | 6024684 | 4710427.5 | 2066443 | 3093001 | 2878400.5 | **0.464987563** | **0** |
| sp\|Q8TBK | Q8TBK6 | ZCCHC10 | Zinc finger CCHC domain-containing protein 10 OS=Hom | 663707.62 | 529745.69 | 673524.88 | 0 | 0 | 0 | 868105.94 | 0 | **0.464979154** | **5.5** |

| | | | | | | | | | | | | | |
|---|---|---|---|---|---|---|---|---|---|---|---|---|---|
| sp\|P43155 | P43155 | CRAT | Carnitine O-acetyltransferase OS=Homo sapiens (Humar | 4412984.5 | 10912211 | 17586490 | 14495674 | 5583215 | 5224281 | 4387310 | 6842780.5 | **0.46485581** | 3 |
| sp\|Q9NRK | Q9NRK6 | ABCB10 | ATP-binding cassette sub-family B member 10, mitochon | 2445374.5 | 1798665 | 1515028.88 | 2554905.5 | 801777.12 | 1174419.25 | 855030.38 | 1027853.56 | **0.464167962** | 0 |
| sp\|Q8WZA | Q8WZA9 | IRGQ | Immunity-related GTPase family Q protein OS=Homo sa | 1303933.62 | 1097600.88 | 1034798.62 | 1063823.62 | 749165.81 | 0 | 693030 | 641617.75 | **0.463053551** | 0 |
| sp\|Q6IA69 | Q6IA69 | NADSYN1 | Glutamine-dependent NAD(+) synthetase OS=Homo sap | 683886.12 | 554288.12 | 441649.56 | 637311.62 | 407881.5 | 445018.38 | 0 | 218217.52 | **0.462259301** | 1 |
| sp\|P49321 | P49321 | NASP | Nuclear autoantigenic sperm protein OS=Homo sapiens | 63751996 | 68786920 | 121382616 | 89406848 | 44457960 | 39468760 | 37800608 | 36693444 | **0.461426381** | 0 |
| sp\|Q9UBT | Q9UBT7 | CTNNAL1 | Alpha-catulin OS=Homo sapiens (Human) GN=CTNNAL1 | 1614327.25 | 1769323 | 1759766.25 | 1560101.75 | 708851.88 | 645553.88 | 563778.75 | 1168846.5 | **0.460509078** | 0 |
| sp\|Q9Y6M | Q9Y6M7 | SLC4A7 | Sodium bicarbonate cotransporter 3 OS=Homo sapiens ( | 338211.78 | 454468.66 | 410012.5 | 407467.03 | 194683.25 | 279069.41 | 0 | 266463.44 | **0.459715875** | 0 |
| sp\|Q9BPX | Q9BPX3 | NCAPG | Condensin complex subunit 3 OS=Homo sapiens (Humar | 4590599.5 | 4782368 | 6669734 | 4851070.5 | 3518070.5 | 2403318 | 1184560.25 | 2416470.75 | **0.455753968** | 0 |
| sp\|Q1289 | Q12894 | IFRD2 | Interferon-related developmental regulator 2 OS=Homo | 148396.11 | 153278.61 | 143293.53 | 241790.16 | 83399.29 | 0 | 110473.37 | 118570.03 | **0.454952841** | 0 |
| sp\|P11169 | P11169 | SLC2A3 | Solute carrier family 2, facilitated glucose transporter me | 65940264 | 59021760 | 61228184 | 75741312 | 35966604 | 46671852 | 17519410 | 19003452 | **0.454933099** | 0 |
| sp\|P11532 | P11532 | DMD | Dystrophin OS=Homo sapiens (Human) GN=DMD PE=1 S | 2149162.5 | 1548482.75 | 1685492.25 | 1917954.25 | 683521.38 | 670882.5 | 1238291.25 | 728733.25 | **0.454922153** | 0 |
| sp\|Q9GZZ | Q9GZZ9 | UBA5 | Ubiquitin-like modifier-activating enzyme 5 OS=Homo sa | 1246448.38 | 1168758.12 | 1127462.25 | 992769.06 | 554177.88 | 652427.56 | 0 | 844633.06 | **0.452269127** | 0 |
| sp\|P40763 | P40763 | STAT3 | Signal transducer and activator of transcription 3 OS=Ho | 10882320 | 10875215 | 10514083 | 11247133 | 4357549 | 5088127.5 | 5094986.5 | 5101906 | **0.451358749** | 0 |
| sp\|Q7Z5L | Q7Z5L9 | IRF2BP2 | Interferon regulatory factor 2-binding protein 2 OS=Hom | 2743832 | 3803171 | 4016500.5 | 2311156.25 | 1404967.25 | 1662499.62 | 1705525.38 | 988933.06 | **0.447540007** | 0 |
| sp\|P02751 | P02751 | FN1 | Fibronectin OS=Homo sapiens (Human) GN=FN1 PE=1 S\ | 30846546 | 29423080 | 32775110 | 24040270 | 13590262 | 12696342 | 11859486 | 14237429 | **0.447397329** | 0 |
| sp\|Q9Y58 | Q9Y587 | AP4S1 | AP-4 complex subunit sigma-1 OS=Homo sapiens (Huma | 352995.38 | 501721.25 | 489408.16 | 379870.12 | 0 | 255561.92 | 297962.12 | 216929.95 | **0.446900386** | 0 |
| sp\|Q8NHC | Q8NHQ9 | DDX55 | ATP-dependent RNA helicase DDX55 OS=Homo sapiens ( | 1182073.12 | 1756553.5 | 2529669 | 1822398 | 517660.53 | 1154047.38 | 463313.44 | 1119848.25 | **0.446441693** | 0 |
| sp\|Q8TD1 | Q8TD16 | BICD2 | Protein bicaudal D homolog 2 OS=Homo sapiens (Humar | 403495.66 | 275586.31 | 318080.91 | 395713.44 | 113757.53 | 236496.44 | 127895.32 | 137925.64 | **0.442304116** | 0 |
| sp\|P23258 | P23258 | TUBG1 | Tubulin gamma-1 chain OS=Homo sapiens (Human) GN= | 3048370 | 839939.38 | 512957.41 | 992793.69 | 552631.56 | 854417.81 | 560156.25 | 402971.03 | **0.439404908** | 4 |
| sp\|O75191 | O75191 | XYLB | Xylulose kinase OS=Homo sapiens (Human) GN=XYLB PE | 1338449.62 | 1183201.25 | 1729559.5 | 1653875.25 | 774000.69 | 583044.38 | 0 | 1227867.75 | **0.437743495** | 1 |
| sp\|O7512 | O75127 | PTCD1 | Pentatricopeptide repeat-containing protein 1, mitochon | 1262392.75 | 1327184.62 | 1341871.62 | 1437480.12 | 449002.22 | 370094.94 | 1020094.5 | 501087.31 | **0.435893066** | 0 |
| sp\|Q9BRP | Q9BRP1 | PDCD2L | Programmed cell death protein 2-like OS=Homo sapiens | 1371953.38 | 1104931.25 | 1680119.5 | 1233371.75 | 728797.88 | 545142.44 | 595295.38 | 461257.19 | **0.432343299** | 0 |
| sp\|Q0241 | Q02413 | DSG1 | Desmoglein-1 OS=Homo sapiens (Human) GN=DSG1 PE= | 337599.22 | 1604252.38 | 400756.19 | 213159.83 | 211702.83 | 0 | 0 | 890357.25 | **0.431205119** | 3 |
| sp\|Q9Y25 | Q9Y256 | RCE1 | CAAX prenyl protease 2 OS=Homo sapiens (Human) GN= | 322536.66 | 318155.12 | 549076.88 | 396699.84 | 0 | 0 | 351570.28 | 331868.28 | **0.430792392** | 4 |
| sp\|Q9H22 | Q9H223 | EHD4 | EH domain-containing protein 4 OS=Homo sapiens (Hum | 6324350 | 6411867 | 6657297 | 6062660.5 | 2313800 | 3977509.5 | 1337988.75 | 3333007.5 | **0.430634452** | 0 |
| sp\|O95251 | O95251 | KAT7 | Histone acetyltransferase KAT7 OS=Homo sapiens (Hum | 253488.28 | 300594.5 | 117973.78 | 538165.38 | 75236.95 | 104688.81 | 208783.91 | 130585.04 | **0.429090477** | 2 |
| sp\|P25098 | P25098 | GRK2 | Beta-adrenergic receptor kinase 1 OS=Homo sapiens (Hu | 655394.5 | 749427.69 | 799629.12 | 608266.06 | 0 | 385997.88 | 341279.84 | 479205.88 | **0.428938795** | 0 |
| sp\|Q9UNH | Q9UNH7 | SNX6 | Sorting nexin-6 OS=Homo sapiens (Human) GN=SNX6 PE | 4286440.5 | 4529243 | 3994646 | 4609498 | 1646892.62 | 1956661 | 1780319.25 | 2051257.75 | **0.426819991** | 0 |
| sp\|O1474 | O14744 | PRMT5 | Protein arginine N-methyltransferase 5 OS=Homo sapier | 42380724 | 48927844 | 45697724 | 43061252 | 22133544 | 17752580 | 20540970 | 16261883 | **0.425890059** | 0 |
| sp\|Q9Y5L | Q9Y5L3 | ENTPD2 | Ectonucleoside triphosphate diphosphohydrolase 2 OS= | 496433 | 375092.09 | 285443.88 | 396828.69 | 193004.55 | 286075.81 | 179195.14 | 0 | **0.42365587** | 1 |
| sp\|Q969V | Q969V6 | MRTFA | Myocardin-related transcription factor A OS=Homo sapi | 224977.77 | 195349.95 | 189638.69 | 197372.16 | 95746.39 | 86125.62 | 98517.36 | 59008.99 | **0.420391608** | 0 |
| sp\|Q6P99 | Q6P996 | PDXDC1 | Pyridoxal-dependent decarboxylase domain-containing p | 21470830 | 21810040 | 22757064 | 18969016 | 8260570 | 7673204 | 9960399 | 9797706 | **0.419870128** | 0 |
| sp\|Q9UBU | Q9UBU8 | MORF4L1 | Mortality factor 4-like protein 1 OS=Homo sapiens (Hum | 441661 | 422681 | 361222.03 | 206642.81 | 0 | 311294.12 | 0 | 289005.12 | **0.419142838** | 2 |
| sp\|Q9Y3A | Q9Y3A2 | UTP11 | Probable U3 small nucleolar RNA-associated protein 11 ( | 644609.94 | 0 | 0 | 809486.81 | 606251.88 | 0 | 0 | 0 | **0.416926783** | 5 |
| sp\|O14841 | O14841 | OPLAH | 5-oxoprolinase OS=Homo sapiens (Human) GN=OPLAH P | 4577309 | 4172249.75 | 3787859 | 4364001 | 1207921.5 | 1132488.5 | 3164918.75 | 1525540 | **0.415992814** | 0 |
| sp\|Q14C8 | Q14C86 | GAPVD1 | GTPase-activating protein and VPS9 domain-containing p | 1943452.88 | 1645685 | 1756774.12 | 2117908.5 | 608821.75 | 1124592 | 535036.19 | 830917.25 | **0.415252107** | 0 |
| sp\|P28676 | P28676 | GCA | Grancalcin OS=Homo sapiens (Human) GN=GCA PE=1 SV | 746975.31 | 596069.25 | 662158.94 | 788852.38 | 0 | 363478.88 | 346327.5 | 443791.41 | **0.412875705** | 0 |
| sp\|Q9Y24 | Q9Y241 | HIGD1A | HIG1 domain family member 1A, mitochondrial OS=Hom | 6206260 | 3557546.5 | 4966685 | 9553948 | 2477501.5 | 2408320.25 | 2492425.5 | 2619538 | **0.411695121** | 0 |
| sp\|Q9H1C | Q9H1C4 | UNC93B1 | Protein unc-93 homolog B1 OS=Homo sapiens (Human) ( | 358114.78 | 473596.28 | 377253.97 | 494217 | 198689.48 | 285142.31 | 205421.77 | 0 | **0.404685787** | 0 |
| sp\|Q9BT2 | Q9BT22 | ALG1 | Chitobiosyldiphosphodolichol beta-mannosyltransferase | 3351571.25 | 2373904 | 1105960.75 | 3300398.5 | 1459041.25 | 697996.94 | 1426472.25 | 504991.31 | **0.403530254** | 2 |
| sp\|Q5BKZ | Q5BKZ1 | ZNF326 | DBIRD complex subunit ZNF326 OS=Homo sapiens (Hum | 5973456 | 6497264.5 | 6701995.5 | 4783974 | 2093807 | 2754698 | 2476980.75 | 2334614.5 | **0.403231843** | 0 |
| sp\|Q1413 | Q14139 | UBE4A | Ubiquitin conjugation factor E4 A OS=Homo sapiens (Hu | 724910.06 | 745170 | 798037.62 | 640556.19 | 132285.2 | 161554.97 | 133580.23 | 738103.44 | **0.400706264** | 2 |
| sp\|Q8WT | Q8WTW3 | COG1 | Conserved oligomeric Golgi complex subunit 1 OS=Hom | 816812 | 1827762.75 | 1839727 | 7448182 | 833748.19 | 963037.88 | 1353244.12 | 1615184.25 | **0.399348077** | 4 |
| sp\|Q9Y5A | Q9Y5A9 | YTHDF2 | YTH domain-containing family protein 2 OS=Homo sapie | 2455090.75 | 2078257.62 | 2434823 | 2761635.5 | 2130347.25 | 0 | 1713060.62 | 0 | **0.395013788** | 1 |
| sp\|Q9NVI | Q9NVI1 | FANCI | Fanconi anemia group I protein OS=Homo sapiens (Hum | 353429.09 | 243772.11 | 352423.84 | 348492.06 | 206469.12 | 0 | 0 | 298768.25 | **0.389207853** | 1 |

| | | | | | | | | | | | | | |
|---|---|---|---|---|---|---|---|---|---|---|---|---|---|
| sp\|Q9BQ6 | Q9BQ67 | GRWD1 | Glutamate-rich WD repeat-containing protein 1 OS=Hom | 5526077 | 5807613 | 6038835 | 5487794.5 | 1757704.75 | 2496665.5 | 2100981.75 | 2501467.5 | **0.387432009** | 0 |
| sp\|P35790 | P35790 | CHKA | Choline kinase alpha OS=Homo sapiens (Human) GN=CH | 3604254.5 | 3456229.75 | 2772762 | 3681560 | 848807.88 | 1260396.75 | 2235049.75 | 792544.88 | **0.380086785** | 0 |
| sp\|Q86UK | Q86UK7 | ZNF598 | E3 ubiquitin-protein ligase ZNF598 OS=Homo sapiens (H | 448398.25 | 408936.56 | 488338.59 | 0 | 0 | 262921.31 | 0 | 243697.64 | **0.376479872** | 3 |
| sp\|Q9H9A | Q9H9A6 | LRRC40 | Leucine-rich repeat-containing protein 40 OS=Homo sap | 1160211.5 | 1149188.25 | 1879265.75 | 1148961.75 | 417066.09 | 934216.62 | 372638.53 | 263361.31 | **0.372315723** | 0 |
| sp\|Q9BRF | Q9BRF8 | CPPED1 | Serine/threonine-protein phosphatase CPPED1 OS=Hom | 56399672 | 55895568 | 53089008 | 125088640 | 28382936 | 27451570 | 27319732 | 24364076 | **0.370149224** | 0 |
| sp\|Q0818 | Q08188 | TGM3 | Protein-glutamine gamma-glutamyltransferase E OS=Ho | 0 | 1784518.12 | 0 | 0 | 0 | 658441.69 | 0 | 0 | **0.368974505** | 7.5 |
| sp\|Q86YD | Q86YD1 | PTOV1 | Prostate tumor-overexpressed gene 1 protein OS=Homo | 1297251.75 | 0 | 1162065.25 | 2686698.5 | 409812.03 | 550234.62 | 470035.53 | 464120.16 | **0.368091068** | 4 |
| sp\|Q96AC | Q96AC1 | FERMT2 | Fermitin family homolog 2 OS=Homo sapiens (Human) G | 2560481.5 | 3485984.25 | 3659097 | 2970235.75 | 1300153.62 | 983553.5 | 1129784.25 | 1060522.88 | **0.352957192** | 0 |
| sp\|Q8IY26 | Q8IY26 | PLPP6 | Phospholipid phosphatase 6 OS=Homo sapiens (Human) | 834416.44 | 771704.31 | 772832.44 | 954428.94 | 0 | 536941.12 | 639393.88 | 0 | **0.352895334** | 0 |
| sp\|P61201 | P61201 | COPS2 | COP9 signalosome complex subunit 2 OS=Homo sapiens | 1449235.75 | 3119200 | 3831899 | 3406743 | 1166816.75 | 676647.62 | 1205554.25 | 1106848.12 | **0.351980975** | 0 |
| sp\|P18827 | P18827 | SDC1 | Syndecan-1 OS=Homo sapiens (Human) GN=SDC1 PE=1 S | 5724860.5 | 4849878 | 3969196.25 | 5160199.5 | 890166.12 | 2297961.25 | 2150932.25 | 1592595.25 | **0.351786827** | 0 |
| sp\|P33121 | P33121 | ACSL1 | Long-chain-fatty-acid--CoA ligase 1 OS=Homo sapiens (H | 12575073 | 12297796 | 14314784 | 13515850 | 4632165.5 | 5192998 | 3986259.75 | 4652179 | **0.350329697** | 0 |
| sp\|O75312 | O75312 | ZPR1 | Zinc finger protein ZPR1 OS=Homo sapiens (Human) GN= | 1307986 | 1677572.38 | 13566551 | 1449214.62 | 1676838.75 | 1329235.75 | 1908681.75 | 1315313.5 | **0.346089529** | 7 |
| sp\|Q9Y5B | Q9Y5B6 | PAXBP1 | PAX3- and PAX7-binding protein 1 OS=Homo sapiens (Hu | 644164.44 | 492341.88 | 0 | 0 | 0 | 391472 | 0 | 0 | **0.34445211** | 5 |
| sp\|O00411 | O00411 | POLRMT | DNA-directed RNA polymerase, mitochondrial OS=Homo | 2871825 | 3539364.75 | 3314463 | 4224334 | 1063224.62 | 1247380.62 | 742020.38 | 1750810.75 | **0.344332683** | 0 |
| sp\|Q9963 | Q99638 | RAD9A | Cell cycle checkpoint control protein RAD9A OS=Homo s | 604135.38 | 565102.44 | 457267.38 | 651646.56 | 296691.09 | 168348.53 | 162694.5 | 156595.77 | **0.344283425** | 0 |
| sp\|Q1354 | Q13546 | RIPK1 | Receptor-interacting serine/threonine-protein kinase 1 | 494931.47 | 499786.72 | 629296.5 | 397443.16 | 0 | 0 | 402075.62 | 287128.88 | **0.340944284** | 1 |
| sp\|P36969 | P36969 | GPX4 | Phospholipid hydroperoxide glutathione peroxidase OS= | 2242180.75 | 2592721 | 3846336 | 3143701.5 | 819866.56 | 1205258.12 | 704909.56 | 1255805.62 | **0.337070642** | 0 |
| sp\|Q96MG | Q96MG7 | NSMCE3 | Non-structural maintenance of chromosomes element 3 | 300593.38 | 343725.31 | 403094.06 | 452078.31 | 0 | 0 | 278289.5 | 221748.59 | **0.333471871** | 0 |
| sp\|Q0143 | Q01433 | AMPD2 | AMP deaminase 2 OS=Homo sapiens (Human) GN=AMPI | 1621789 | 1062394.5 | 1527862.5 | 1704183 | 254658.94 | 0 | 724753.19 | 991617.94 | **0.333156487** | 0 |
| sp\|Q68CP | Q68CP9 | ARID2 | AT-rich interactive domain-containing protein 2 OS=Hom | 5493061.5 | 6898644 | 5771704 | 6859730 | 2047145.62 | 2686074.75 | 1872450.5 | 1651551 | **0.329983449** | 0 |
| sp\|Q6GYQ | Q6GYQ0 | RALGAPA1 | Ral GTPase-activating protein subunit alpha-1 OS=Homo | 639639.44 | 499532.19 | 555385.31 | 498931.56 | 0 | 356866 | 365139.53 | 0 | **0.329158566** | 0 |
| sp\|Q8TEX | Q8TEX9 | IPO4 | Importin-4 OS=Homo sapiens (Human) GN=IPO4 PE=1 SV | 63644864 | 53600444 | 58740752 | 62097472 | 19271332 | 20848556 | 17067954 | 20817354 | **0.327637932** | 0 |
| sp\|Q8NCF | Q8NCF5 | NFATC2IP | NFATC2-interacting protein OS=Homo sapiens (Human) | 424559.53 | 0 | 0 | 380987.78 | 263837.06 | 0 | 0 | 0 | **0.32752522** | 5 |
| sp\|Q9UHN | Q9UHN6 | CEMIP2 | Cell surface hyaluronidase OS=Homo sapiens (Human) G | 1357431.12 | 2306269 | 1835930.75 | 1667185.62 | 700760.25 | 915933.25 | 701698.25 | 0 | **0.323489761** | 0 |
| sp\|O9560 | O95602 | POLR1A | DNA-directed RNA polymerase I subunit RPA1 OS=Homo | 1580754.5 | 1597350 | 3364008.5 | 2551375.5 | 705714.31 | 656538.12 | 579820.44 | 991950.62 | **0.322651037** | 0 |
| sp\|P11047 | P11047 | LAMC1 | Laminin subunit gamma-1 OS=Homo sapiens (Human) G | 10113994 | 9437985 | 9928170 | 9266569 | 3758921.75 | 3746137.75 | 2885545.75 | 2043468.62 | **0.320906505** | 0 |
| sp\|Q9Y5X | Q9Y5X2 | SNX8 | Sorting nexin-8 OS=Homo sapiens (Human) GN=SNX8 PE | 669435.38 | 1046382.94 | 1398129.5 | 934885.5 | 239515.81 | 407071.38 | 531222.62 | 119517.7 | **0.320420083** | 0 |
| sp\|Q96EP | Q96EP0 | RNF31 | E3 ubiquitin-protein ligase RNF31 OS=Homo sapiens (Hu | 1015856.12 | 1009533.5 | 1145656.25 | 539828.75 | 255469.38 | 331589.44 | 318718.97 | 280034.19 | **0.319550538** | 0 |
| sp\|Q9BRT | Q9BRT2 | UQCC2 | Ubiquinol-cytochrome-c reductase complex assembly fac | 0 | 620941.62 | 728917.44 | 1011571.88 | 0 | 745383 | 0 | 0 | **0.315648867** | 4.5 |
| sp\|P42357 | P42357 | HAL | Histidine ammonia-lyase OS=Homo sapiens (Human) GN | 0 | 216356.84 | 153409.91 | 0 | 0 | 0 | 115695.51 | 0 | **0.312887814** | 5 |
| sp\|Q9BTY | Q9BTY7 | HGH1 | Protein HGH1 homolog OS=Homo sapiens (Human) GN= | 122638.51 | 0 | 0 | 124686.38 | 0 | 0 | 0 | 77128.84 | **0.311852317** | 5 |
| sp\|Q9UL4 | Q9UL46 | PSME2 | Proteasome activator complex subunit 2 OS=Homo sapie | 73854160 | 81908656 | 67116240 | 85194376 | 19127420 | 22776788 | 22152390 | 31534392 | **0.31028638** | 0 |
| sp\|Q9995 | Q99956 | DUSP9 | Dual specificity protein phosphatase 9 OS=Homo sapien | 10367735 | 8405520 | 12808894 | 17200410 | 3926879 | 2741348.75 | 4033612.25 | 4043956.25 | **0.302275989** | 0 |
| sp\|P19174 | P19174 | PLCG1 | 1-phosphatidylinositol 4,5-bisphosphate phosphodiester | 2141372 | 2350197 | 2862382.5 | 1433357.88 | 519973.81 | 562961.25 | 1022378.88 | 523955.84 | **0.299212155** | 0 |
| sp\|Q8WV | Q8WVM8 | SCFD1 | Sec1 family domain-containing protein 1 OS=Homo sapie | 37523244 | 36719424 | 41046020 | 38477300 | 10797205 | 11315010 | 10926541 | 12487064 | **0.296072107** | 0 |
| sp\|Q96E2 | Q96E29 | MTERF3 | Transcription termination factor 3, mitochondrial OS=Hc | 473145.94 | 502285.47 | 471832.88 | 413381.06 | 289991.94 | 0 | 0 | 258570.47 | **0.294823734** | 0 |
| sp\|Q9BVC | Q9BVC6 | TMEM109 | Transmembrane protein 109 OS=Homo sapiens (Human) | 3014757.25 | 3603563 | 4338445.5 | 3299675.25 | 2194444.75 | 0 | 2008005.75 | 0 | **0.294775568** | 0 |
| sp\|Q9UN3 | Q9UN36 | NDRG2 | Protein NDRG2 OS=Homo sapiens (Human) GN=NDRG2 I | 2316438 | 1523394.75 | 2276980.5 | 1926590.62 | 0 | 1287291 | 1010985.56 | 0 | **0.285734323** | 0 |
| sp\|Q8N2G | Q8N2G8 | GHDC | GH3 domain-containing protein OS=Homo sapiens (Hum | 708517.06 | 849211.12 | 792588.56 | 787279 | 524994.19 | 352170.59 | 0 | 0 | **0.279565901** | 0 |
| sp\|Q8IVL6 | Q8IVL6 | P3H3 | Prolyl 3-hydroxylase 3 OS=Homo sapiens (Human) GN=P | 2102218 | 1329325.88 | 1955483.25 | 2373191 | 234210.62 | 901213.38 | 274393.56 | 747688.06 | **0.27802126** | 0 |
| sp\|Q9BZE | Q9BZE9 | ASPSCR1 | Tether containing UBX domain for GLUT4 OS=Homo sapi | 23004536 | 20800084 | 21335154 | 21146610 | 7316352.5 | 4995120.5 | 5584675.5 | 5982014 | **0.276731523** | 0 |
| sp\|Q5ZPR | Q5ZPR3 | CD276 | CD276 antigen OS=Homo sapiens (Human) GN=CD276 PI | 758477.19 | 0 | 526913.31 | 592446.25 | 0 | 0 | 0 | 516415.84 | **0.275005716** | 2.5 |
| sp\|Q9BTZ | Q9BTZ2 | DHRS4 | Dehydrogenase/reductase SDR family member 4 OS=Ho | 3648862 | 5283953.5 | 4700827 | 3719917 | 934333.12 | 1495713.25 | 949526 | 1361877.75 | **0.273226373** | 0 |
| sp\|P12277 | P12277 | CKB | Creatine kinase B-type OS=Homo sapiens (Human) GN=C | 60755000 | 54599724 | 60378348 | 52534916 | 18067792 | 23001320 | 10370327 | 10806339 | **0.272687285** | 0 |

| | | | | | | | | | | | | | |
|---|---|---|---|---|---|---|---|---|---|---|---|---|---|
| sp\|O0056( | O00560 | SDCBP | Syntenin-1 OS=Homo sapiens (Human) GN=SDCBP PE=1 | 1812650.62 | 1801842.88 | 1792427.12 | 1813684.25 | 429976.72 | 604499.31 | 610743.19 | 304428.25 | **0.270011655** | **0** |
| sp\|P02679 | P02679 | FGG | Fibrinogen gamma chain OS=Homo sapiens (Human) GN | 22486724 | 32423168 | 35476368 | 27485028 | 4844163.5 | 5199120 | 11883405 | 9372688 | **0.265538597** | **0** |
| sp\|Q86U3: | Q86U38 | NOP9 | Nucleolar protein 9 OS=Homo sapiens (Human) GN=NOF | 15312048 | 13942227 | 14534857 | 15181250 | 3849829.75 | 3278744.25 | 4427945 | 3907451.5 | **0.262232836** | **0** |
| sp\|Q96ER: | Q96ER3 | SAAL1 | Protein SAAL1 OS=Homo sapiens (Human) GN=SAAL1 PE | 932465.12 | 688032.69 | 1335736.5 | 786835.12 | 0 | 543189.88 | 407342.84 | 0 | **0.25394472** | **0** |
| sp\|O7578? | O75787 | ATP6AP2 | Renin receptor OS=Homo sapiens (Human) GN=ATP6AP2 | 0 | 533608.31 | 703626.12 | 631516.69 | 0 | 0 | 474093.75 | 0 | **0.2536955** | **2.5** |
| sp\|P15291 | P15291 | B4GALT1 | Beta-1,4-galactosyltransferase 1 OS=Homo sapiens (Hun | 2771415.25 | 1950107.25 | 2016189 | 2549153.75 | 1203468 | 423576.75 | 321389.47 | 352827.91 | **0.247797515** | **0** |
| sp\|Q8NBT | Q8NBT2 | SPC24 | Kinetochore protein Spc24 OS=Homo sapiens (Human) G | 356446.06 | 208700.23 | 172668.09 | 0 | 0 | 0 | 182222.78 | 0 | **0.246976455** | **3.5** |
| sp\|P04264 | P04264 | KRT1 | Keratin, type II cytoskeletal 1 OS=Homo sapiens (Human | 6946205 | 428274752 | 16810610 | 14165467 | 19092360 | 28787024 | 35131776 | 32094942 | **0.246904407** | **4** |
| sp\|Q1505( | Q15054 | POLD3 | DNA polymerase delta subunit 3 OS=Homo sapiens (Hun | 1273264.25 | 980689.81 | 1112931.25 | 1388295.5 | 0 | 609253 | 562082.69 | 0 | **0.246328318** | **0** |
| sp\|P07996 | P07996 | THBS1 | Thrombospondin-1 OS=Homo sapiens (Human) GN=THB | 874252.19 | 861989.69 | 1088211.62 | 796893.75 | 361016 | 510843.41 | 0 | 0 | **0.240755539** | **0** |
| sp\|O1489: | O14893 | GEMIN2 | Gem-associated protein 2 OS=Homo sapiens (Human) GI | 373874.5 | 367851.88 | 284973.69 | 406780.09 | 36638.55 | 154128.22 | 153394.64 | 0 | **0.240088018** | **0** |
| sp\|Q9UGV | Q9UGV2 | NDRG3 | Protein NDRG3 OS=Homo sapiens (Human) GN=NDRG3 I | 427472.22 | 450253.81 | 0 | 383809.81 | 0 | 0 | 0 | 302654.19 | **0.239909308** | **2.5** |
| sp\|Q9Y6K! | Q9Y6K5 | OAS3 | 2'-5'-oligoadenylate synthase 3 OS=Homo sapiens (Hum | 1815187.38 | 1243525.12 | 2135001.25 | 2273465.5 | 647615.38 | 0 | 599897.69 | 543843.38 | **0.239897341** | **0** |
| sp\|P53611 | P53611 | RABGGTB | Geranylgeranyl transferase type-2 subunit beta OS=Hom | 1756332.62 | 985459 | 1005788.06 | 896073.94 | 1107288.38 | 0 | 0 | 0 | **0.238451976** | **3** |
| sp\|Q8NBP | Q8NBP7 | PCSK9 | Proprotein convertase subtilisin/kexin type 9 OS=Homo | 188226.3 | 242359.38 | 189447.23 | 239010.62 | 49620.05 | 0 | 76169.76 | 73226.7 | **0.231672206** | **0** |
| sp\|Q1590( | Q15904 | ATP6AP1 | V-type proton ATPase subunit S1 OS=Homo sapiens (Hur | 0 | 584546.31 | 589813.62 | 587100.44 | 391144.88 | 0 | 0 | 0 | **0.222057156** | **2.5** |
| sp\|O9581( | O95810 | CAVIN2 | Caveolae-associated protein 2 OS=Homo sapiens (Huma | 3331654.25 | 2484506.75 | 2849240.25 | 0 | 0 | 1919478.12 | 0 | 0 | **0.221510587** | **2.5** |
| sp\|Q0295: | Q02952 | AKAP12 | A-kinase anchor protein 12 OS=Homo sapiens (Human) ( | 837260.38 | 1101216.25 | 1178182.12 | 1025109.56 | 179522.47 | 287272.12 | 252941.7 | 191936.12 | **0.220116709** | **0** |
| sp\|Q1367: | Q13671 | RIN1 | Ras and Rab interactor 1 OS=Homo sapiens (Human) GN | 186425.77 | 229558.75 | 268966.12 | 191011.94 | 0 | 0 | 188497.39 | 0 | **0.215188861** | **1** |
| sp\|Q5VSL! | Q5VSL9 | STRIP1 | Striatin-interacting protein 1 OS=Homo sapiens (Human) | 1320849.75 | 1268047.88 | 1212785.62 | 1538254.38 | 138979.55 | 987704.88 | 0 | 0 | **0.210992058** | **0** |
| sp\|P31431 | P31431 | SDC4 | Syndecan-4 OS=Homo sapiens (Human) GN=SDC4 PE=1 S | 4440049.5 | 3510577.5 | 5569966.5 | 4592496 | 1038944.81 | 904977.44 | 808313.56 | 932796.12 | **0.203445797** | **0** |
| sp\|P48730 | P48730 | CSNK1D | Casein kinase I isoform delta OS=Homo sapiens (Human) | 335250.97 | 267167.25 | 468346.84 | 342534.16 | 0 | 0 | 284302.94 | 0 | **0.201162596** | **1** |
| sp\|Q1303: | Q13033 | STRN3 | Striatin-3 OS=Homo sapiens (Human) GN=STRN3 PE=1 SV | 3703242 | 3925962.25 | 4359087 | 2838769.75 | 883016.25 | 897822.81 | 0 | 1166816.75 | **0.198802434** | **0** |
| sp\|P16591 | P16591 | FER | Tyrosine-protein kinase Fer OS=Homo sapiens (Human) ( | 459986.12 | 219861.91 | 225687.34 | 0 | 177624.39 | 0 | 0 | 0 | **0.196154006** | **2.5** |
| sp\|Q9269( | Q92696 | RABGGTA | Geranylgeranyl transferase type-2 subunit alpha OS=Hor | 4514926.5 | 3909135 | 5364122.5 | 3853056.5 | 532119.62 | 693700.88 | 1167492 | 1059599 | **0.195729518** | **0** |
| sp\|O1509: | O15091 | KIAA0391 | Mitochondrial ribonuclease P catalytic subunit OS=Homo | 4429534.5 | 4471404.5 | 4954169 | 4889825 | 546887.5 | 745375.62 | 538936.19 | 1783292.5 | **0.192825006** | **0** |
| sp\|Q9UG6 | Q9UG63 | ABCF2 | ATP-binding cassette sub-family F member 2 OS=Homo s | 701474.88 | 0 | 602353.12 | 702343.38 | 0 | 0 | 0 | 373105.03 | **0.185978643** | **2.5** |
| sp\|Q96CM | Q96CM3 | RPUSD4 | Mitochondrial RNA pseudouridine synthase RPUSD4 OS= | 81537.97 | 87364.84 | 0 | 97264.95 | 0 | 46388.61 | 0 | 0 | **0.174283354** | **2.5** |
| sp\|Q8IVL1 | Q8IVL1 | NAV2 | Neuron navigator 2 OS=Homo sapiens (Human) GN=NAV | 367571.5 | 266000.38 | 360766.22 | 357585.03 | 65766.34 | 0 | 82966.41 | 80207.97 | **0.16934448** | **0** |
| sp\|O1496! | O14965 | AURKA | Aurora kinase A OS=Homo sapiens (Human) GN=AURKA | 1696403.25 | 633749 | 2452773 | 1726261.12 | 552520.38 | 545282.19 | 0 | 0 | **0.168654346** | **0** |
| sp\|P53350 | P53350 | PLK1 | Serine/threonine-protein kinase PLK1 OS=Homo sapiens | 3271548.75 | 2961362.75 | 5618156.5 | 2017188.75 | 1225012.25 | 523489.34 | 0 | 457238.47 | **0.159049555** | **0** |
| sp\|P37802 | P37802 | TAGLN2 | Transgelin-2 OS=Homo sapiens (Human) GN=TAGLN2 PE | 272813.56 | 264881.28 | 210051.31 | 259444.7 | 160086.27 | 0 | 0 | 0 | **0.158943332** | **0** |
| sp\|P24390 | P24390 | KDELR1 | ER lumen protein-retaining receptor 1 OS=Homo sapiens | 762013.19 | 759004.12 | 742454.81 | 654501.38 | 0 | 0 | 0 | 462709.22 | **0.158572112** | **0** |
| sp\|Q0351! | Q03519 | TAP2 | Antigen peptide transporter 2 OS=Homo sapiens (Humai | 1130323.88 | 864153.62 | 767473.06 | 800080.75 | 345500.47 | 78458.64 | 64306.08 | 65684.34 | **0.155515065** | **0** |
| sp\|Q9P2T: | Q9P2T1 | GMPR2 | GMP reductase 2 OS=Homo sapiens (Human) GN=GMPR | 1520199.38 | 1417331.88 | 1346760.12 | 1580169 | 0 | 346287.03 | 543888.56 | 0 | **0.15179156** | **0** |
| sp\|Q1313( | Q13136 | PPFIA1 | Liprin-alpha-1 OS=Homo sapiens (Human) GN=PPFIA1 PE | 844283.94 | 644967.19 | 770971.88 | 611427.44 | 426574.97 | 0 | 0 | 0 | **0.148546969** | **0** |
| sp\|Q9NRL: | Q9NRL2 | BAZ1A | Bromodomain adjacent to zinc finger domain protein 1A | 229853.58 | 575769.62 | 457321.5 | 523888.44 | 37120.83 | 80272.68 | 72879.73 | 67714.2 | **0.144382502** | **0** |
| sp\|P08833 | P08833 | IGFBP1 | Insulin-like growth factor-binding protein 1 OS=Homo sa | 979960.38 | 920840.12 | 0 | 1119141 | 429432.28 | 0 | 0 | 0 | **0.142198874** | **2.5** |
| sp\|O1551: | O15511 | ARPC5 | Actin-related protein 2/3 complex subunit 5 OS=Homo s | 7778252 | 8446174 | 204266880 | 9890020 | 7658335.5 | 7566049 | 6198799 | 7307756 | **0.124710366** | **0** |
| sp\|Q8TB3( | Q8TB36 | GDAP1 | Ganglioside-induced differentiation-associated protein 1 | 891773.12 | 752467.62 | 925093.69 | 934619.31 | 0 | 0 | 0 | 420951.34 | **0.120136101** | **0** |
| sp\|P05109 | P05109 | S100A8 | Protein S100-A8 OS=Homo sapiens (Human) GN=S100A8 | 9107840 | 179673392 | 6674114 | 6203620 | 8810619 | 6987605 | 6826463.5 | 397023.53 | **0.114161604** | **6** |
| sp\|Q9BQE | Q9BQE5 | APOL2 | Apolipoprotein L2 OS=Homo sapiens (Human) GN=APOL | 5189391 | 1126557 | 6365086 | 4992765 | 695461.88 | 648572.19 | 317313.81 | 84213.09 | **0.098765465** | **0** |
| sp\|Q08AN | Q08AM6 | VAC14 | Protein VAC14 homolog OS=Homo sapiens (Human) GN= | 1122765.38 | 1289716.62 | 57731700 | 1241387.75 | 1032410.38 | 1358097.62 | 1332338.75 | 1603496.38 | **0.086768652** | **7** |
| sp\|P06702 | P06702 | S100A9 | Protein S100-A9 OS=Homo sapiens (Human) GN=S100A9 | 0 | 17452006 | 0 | 368128.94 | 398062.16 | 366674.56 | 279282.94 | 384963.69 | **0.080189255** | **6** |
| sp\|P29508 | P29508 | SERPINB3 | Serpin B3 OS=Homo sapiens (Human) GN=SERPINB3 PE= | 1149473.25 | 22834246 | 840196 | 625632.31 | 859790.75 | 0 | 960538.88 | 0 | **0.071526994** | **4** |

| | | | | | | | | | | | | | |
|---|---|---|---|---|---|---|---|---|---|---|---|---|---|
| sp\|Q0351 | Q03518 | TAP1 | Antigen peptide transporter 1 OS=Homo sapiens (Huma | 2923946.5 | 2528307 | 2519480 | 1410694 | 135654.94 | 159990.25 | 146757.78 | 199201.64 | **0.068383647** | 0 |
| sp\|Q5VTQ | Q5VTQ0 | TTC39B | Tetratricopeptide repeat protein 39B OS=Homo sapiens | 227778.31 | 303120.31 | 104360.66 | 234099.94 | 0 | 57757.77 | 0 | 0 | **0.066437174** | 0 |
| sp\|P48594 | P48594 | SERPINB4 | Serpin B4 OS=Homo sapiens (Human) GN=SERPINB4 PE= | 0 | 11807634 | 0 | 0 | 0 | 0 | 305695.78 | 371713.91 | **0.057370485** | 7 |
| sp\|Q8N8S | Q8N8S7 | ENAH | Protein enabled homolog OS=Homo sapiens (Human) GN | 4142273.5 | 2484229.5 | 2111110.75 | 2026275 | 174556.45 | 245456.64 | 112527.33 | 72062.19 | **0.056169534** | 0 |
| sp\|P23141 | P23141 | CES1 | Liver carboxylesterase 1 OS=Homo sapiens (Human) GN= | 104360584 | 74363048 | 68794792 | 102723080 | 1216856.88 | 1812302.5 | 184397.44 | 178761.88 | **0.009685656** | 0 |
| sp\|O43731 | O43731 | KDELR3 | ER lumen protein-retaining receptor 3 OS=Homo sapiens | 304273.59 | 0 | 141598.03 | 0 | 0 | 0 | 0 | 0 | **0** | 4 |
| sp\|O6067 | O60678 | PRMT3 | Protein arginine N-methyltransferase 3 OS=Homo sapier | 656485.69 | 633645.31 | 906438.94 | 718076 | 0 | 0 | 0 | 0 | **0** | 0 |
| sp\|P00533 | P00533 | EGFR | Epidermal growth factor receptor OS=Homo sapiens (Hu | 537630.25 | 868266.12 | 610048.56 | 739461.56 | 0 | 0 | 0 | 0 | **0** | 0 |
| sp\|P01834 | P01834 | IGKC | Immunoglobulin kappa constant OS=Homo sapiens (Hur | 0 | 402878.62 | 0 | 0 | 0 | 0 | 0 | 0 | **0** | 6 |
| sp\|P01857 | P01857 | IGHG1 | Immunoglobulin heavy constant gamma 1 OS=Homo sap | 0 | 2765675.25 | 0 | 0 | 0 | 0 | 0 | 0 | **0** | 6 |
| sp\|P02792 | P02792 | FTL | Ferritin light chain OS=Homo sapiens (Human) GN=FTL P | 3172909.25 | 3838858.5 | 3096547.25 | 3651949.5 | 0 | 0 | 0 | 0 | **0** | 0 |
| sp\|P02795 | P02795 | MT2A | Metallothionein-2 OS=Homo sapiens (Human) GN=MT2/ | 2619599.75 | 0 | 3921167.5 | 0 | 0 | 0 | 0 | 0 | **0** | 4 |
| sp\|P05156 | P05156 | CFI | Complement factor I OS=Homo sapiens (Human) GN=CF | 1084334 | 0 | 958651.5 | 811417.69 | 0 | 0 | 0 | 0 | **0** | 2 |
| sp\|P07108 | P07108 | DBI | Acyl-CoA-binding protein OS=Homo sapiens (Human) GN | 2388485.25 | 1874691.5 | 1623671.62 | 1780177.5 | 0 | 0 | 0 | 0 | **0** | 0 |
| sp\|P09455 | P09455 | RBP1 | Retinol-binding protein 1 OS=Homo sapiens (Human) GN | 997347.69 | 1512074.88 | 1383124 | 1459047.75 | 0 | 0 | 0 | 0 | **0** | 0 |
| sp\|P09564 | P09564 | CD7 | T-cell antigen CD7 OS=Homo sapiens (Human) GN=CD7 F | 1595628.88 | 1805110.12 | 2153795.25 | 1258746.25 | 0 | 0 | 0 | 0 | **0** | 0 |
| sp\|P1790 | P17900 | GM2A | Ganglioside GM2 activator OS=Homo sapiens (Human) G | 0 | 1314624.38 | 0 | 0 | 0 | 0 | 0 | 0 | **0** | 6 |
| sp\|P22531 | P22531 | SPRR2E | Small proline-rich protein 2E OS=Homo sapiens (Human) | 0 | 18492110 | 0 | 0 | 0 | 0 | 0 | 0 | **0** | 6 |
| sp\|P22532 | P22532 | SPRR2D | Small proline-rich protein 2D OS=Homo sapiens (Human | 0 | 23172992 | 0 | 0 | 0 | 0 | 0 | 0 | **0** | 6 |
| sp\|P3026 | P30260 | CDC27 | Cell division cycle protein 27 homolog OS=Homo sapiens | 570096.56 | 667211.56 | 0 | 0 | 0 | 0 | 0 | 0 | **0** | 4 |
| sp\|P30825 | P30825 | SLC7A1 | High affinity cationic amino acid transporter 1 OS=Homo | 2536007.25 | 3373480.5 | 2338910.75 | 2615352.75 | 0 | 0 | 0 | 0 | **0** | 0 |
| sp\|P35325 | P35325 | SPRR2B | Small proline-rich protein 2B OS=Homo sapiens (Human) | 0 | 18761962 | 0 | 0 | 0 | 0 | 0 | 0 | **0** | 6 |
| sp\|P47929 | P47929 | LGALS7 | Galectin-7 OS=Homo sapiens (Human) GN=LGALS7 PE=1 | 0 | 2098488.25 | 0 | 0 | 0 | 0 | 0 | 0 | **0** | 6 |
| sp\|Q1374 | Q13740 | ALCAM | CD166 antigen OS=Homo sapiens (Human) GN=ALCAM F | 753567 | 675433.81 | 278793.56 | 504454.44 | 0 | 0 | 0 | 0 | **0** | 0 |
| sp\|Q1512 | Q15125 | EBP | 3-beta-hydroxysteroid-Delta(8),Delta(7)-isomerase OS=H | 1948023.38 | 0 | 0 | 0 | 0 | 0 | 0 | 0 | **0** | 6 |
| sp\|Q5RL7 | Q5RL73 | RBM48 | RNA-binding protein 48 OS=Homo sapiens (Human) GN= | 0 | 3467758.75 | 943134.88 | 741968.94 | 0 | 0 | 0 | 0 | **0** | 2 |
| sp\|Q5T74 | Q5T749 | KPRP | Keratinocyte proline-rich protein OS=Homo sapiens (Hur | 0 | 3551472 | 0 | 0 | 0 | 0 | 0 | 0 | **0** | 6 |
| sp\|Q6NZI2 | Q6NZI2 | CAVIN1 | Caveolae-associated protein 1 OS=Homo sapiens (Huma | 635062.75 | 514969.16 | 0 | 808924.5 | 0 | 0 | 0 | 0 | **0** | 2 |
| sp\|Q6PJG | Q6PJG6 | BRAT1 | BRCA1-associated ATM activator 1 OS=Homo sapiens (H | 0 | 0 | 0 | 238084.25 | 0 | 0 | 0 | 0 | **0** | 6 |
| sp\|Q8IYS1 | Q8IYS1 | PM20D2 | Peptidase M20 domain-containing protein 2 OS=Homo s | 0 | 0 | 0 | 513230.22 | 0 | 0 | 0 | 0 | **0** | 6 |
| sp\|Q8N7R | Q8N7R7 | CCNYL1 | Cyclin-Y-like protein 1 OS=Homo sapiens (Human) GN=C | 384717.03 | 330790.31 | 487626.44 | 272434.16 | 0 | 0 | 0 | 0 | **0** | 0 |
| sp\|Q8N8R | Q8N8R5 | C2orf69 | UPF0565 protein C2orf69 OS=Homo sapiens (Human) GN | 477324.5 | 0 | 633908.12 | 448838.88 | 0 | 0 | 0 | 0 | **0** | 2 |
| sp\|Q8WV | Q8WVV4 | POF1B | Protein POF1B OS=Homo sapiens (Human) GN=POF1B PI | 0 | 4655884 | 0 | 0 | 0 | 0 | 0 | 0 | **0** | 6 |
| sp\|Q96DB | Q96DB5 | RMDN1 | Regulator of microtubule dynamics protein 1 OS=Homo | 0 | 637595.19 | 0 | 668866.31 | 0 | 0 | 0 | 0 | **0** | 4 |
| sp\|Q96RM | Q96RM1 | SPRR2F | Small proline-rich protein 2F OS=Homo sapiens (Human) | 0 | 5058778 | 0 | 0 | 0 | 0 | 0 | 0 | **0** | 6 |
| sp\|Q9BW | Q9BW85 | YJU2 | YJU2 splicing factor homolog OS=Homo sapiens (Human | 0 | 0 | 323867.88 | 0 | 0 | 0 | 0 | 0 | **0** | 6 |
| sp\|Q9BYK | Q9BYK8 | HELZ2 | Helicase with zinc finger domain 2 OS=Homo sapiens (Hu | 0 | 289860.12 | 253323.33 | 0 | 0 | 0 | 0 | 0 | **0** | 4 |
| sp\|Q9H0E | Q9H0E2 | TOLLIP | Toll-interacting protein OS=Homo sapiens (Human) GN= | 0 | 0 | 1945686.75 | 0 | 0 | 0 | 0 | 0 | **0** | 6 |
| sp\|Q9NNV | Q9NNW5 | WDR6 | WD repeat-containing protein 6 OS=Homo sapiens (Hum | 662794.56 | 0 | 0 | 726481.12 | 0 | 0 | 0 | 0 | **0** | 4 |
| sp\|Q9NZB | Q9NZB2 | FAM120A | Constitutive coactivator of PPAR-gamma-like protein 1 C | 0 | 866773.5 | 901267.94 | 717989.31 | 0 | 0 | 0 | 0 | **0** | 2 |
| sp\|Q9P0I2 | Q9P0I2 | EMC3 | ER membrane protein complex subunit 3 OS=Homo sapi | 0 | 0 | 0 | 713168 | 0 | 0 | 0 | 0 | **0** | 6 |
| sp\|Q9P2W | Q9P2W9 | STX18 | Syntaxin-18 OS=Homo sapiens (Human) GN=STX18 PE=1 | 0 | 0 | 441428.91 | 0 | 0 | 0 | 0 | 0 | **0** | 6 |
| sp\|Q9UER | Q9UER7 | DAXX | Death domain-associated protein 6 OS=Homo sapiens (H | 0 | 0 | 0 | 445563.84 | 0 | 0 | 0 | 0 | **0** | 6 |
| sp\|Q9UKY | Q9UKY3 | CES1P1 | Putative inactive carboxylesterase 4 OS=Homo sapiens (I | 16969488 | 18823458 | 18171810 | 17437006 | 0 | 0 | 0 | 0 | **0** | 0 |
| sp\|Q9Y54 | Q9Y547 | HSPB11 | Intraflagellar transport protein 25 homolog OS=Homo sa | 0 | 981516.69 | 923559.44 | 955230.19 | 0 | 0 | 0 | 0 | **0** | 2 |
