## Supplementary material for "E cadherin appears to be an essential on/off switch for initiating bile canaliculi formation": SI

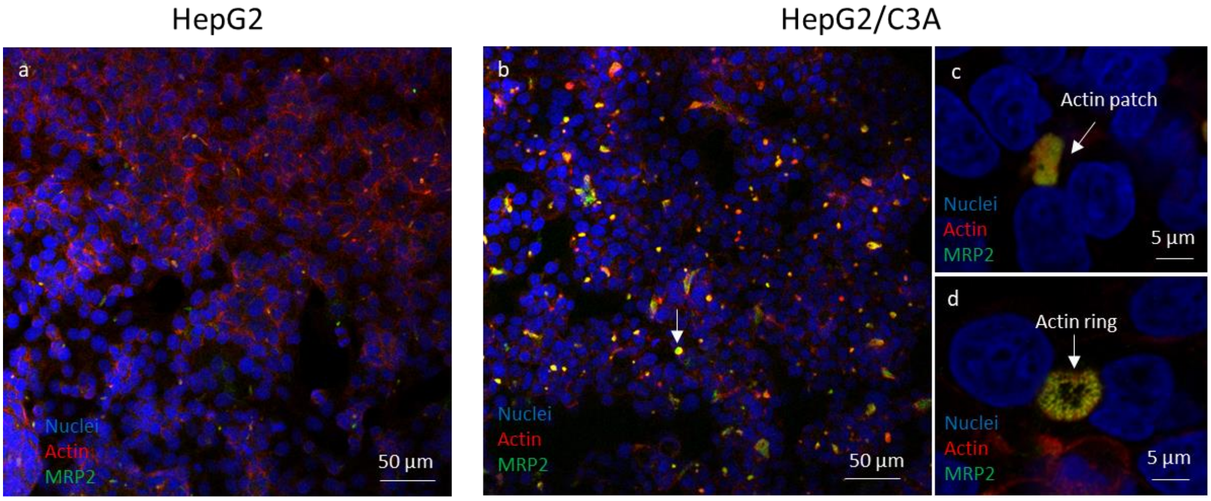


**Figure S1.** Comparison between HepG2 and HepG2/C3A in their capacity to form BC. Fluorescence confocal microscope images acquired at 20x magnification of HepG2 (a) and HepG2/C3A (b) cells labelled forMRP2 and actin. In c and d, HepG2/C3A cells at higher magnification (63x) in airyscan mode. White arrows pinpoint BCs.

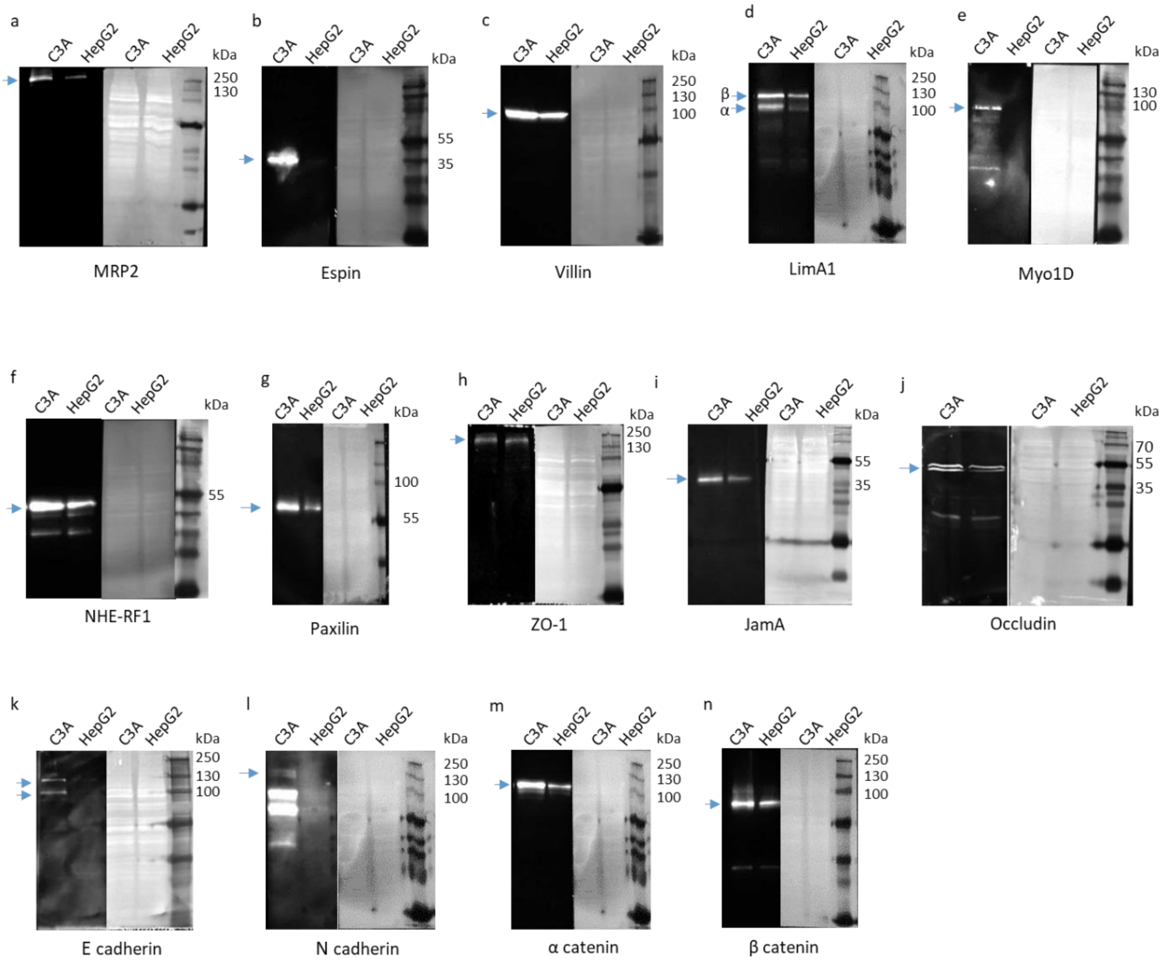


**Figure S2.** Validation by Western blot of the shotgun results. Representative immunoblots of 50 µg of protein extracts from HepG2 and HepG2/C3A cells on the left and the corresponding ink stained of the membrane on the right: (a) MRP2, (b) Espin, (c) Villin, (d) LimA1, (e) Myo1D, (f) NHE-RF1, (g) Paxilin, (h) ZO-1, (i) JamA, (j) Occludin, (k) E cadherin, (l) N cadherin, (m) α catenin, (n) β catenin.

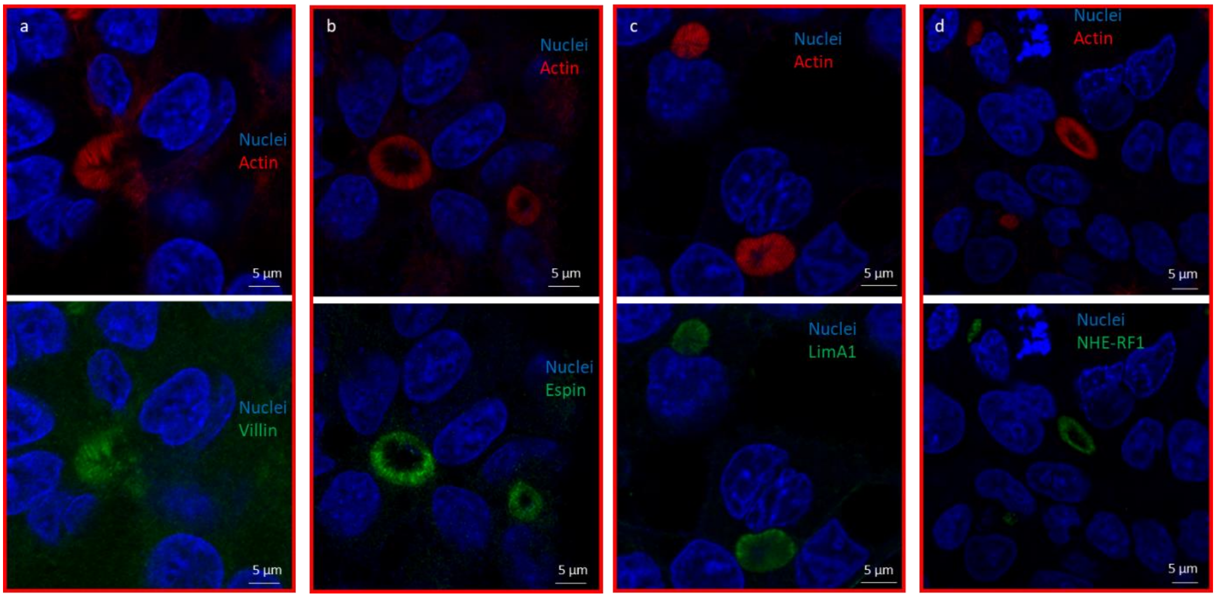


**Figure S3.** Fluorescence confocal microscope images acquired at 63x magnification of HepG2/C3A cells immunolabelled with antibody directed against the indicated proteins (a-d).

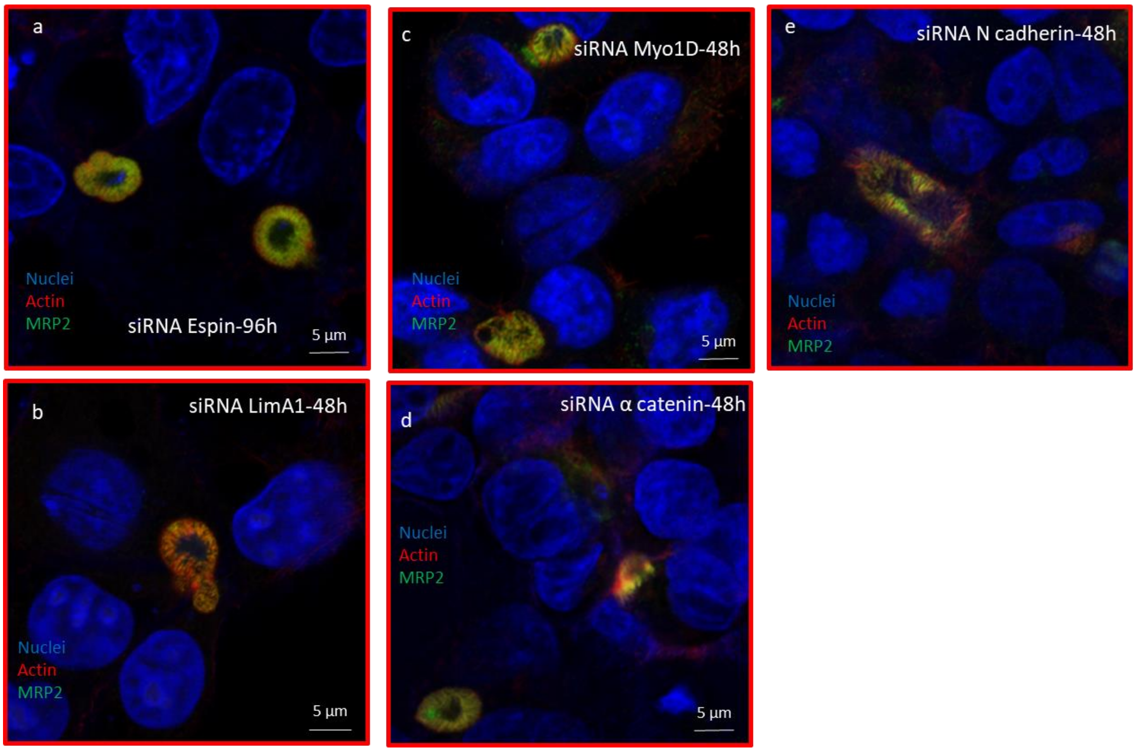


**Figure S4**. Immunofluorescence labelling of BC (MRP2, actin) on the HepG2/C3A cell line after siRNA treatment against: (a) Espin, (b) LimA1, (c) Myo1D, (d) α catenin, (e) N cadherin. Confocal images acquired at 63x magnification in airyscan mode of HepG2/C3A cells labelled for MRP2 and Actin.

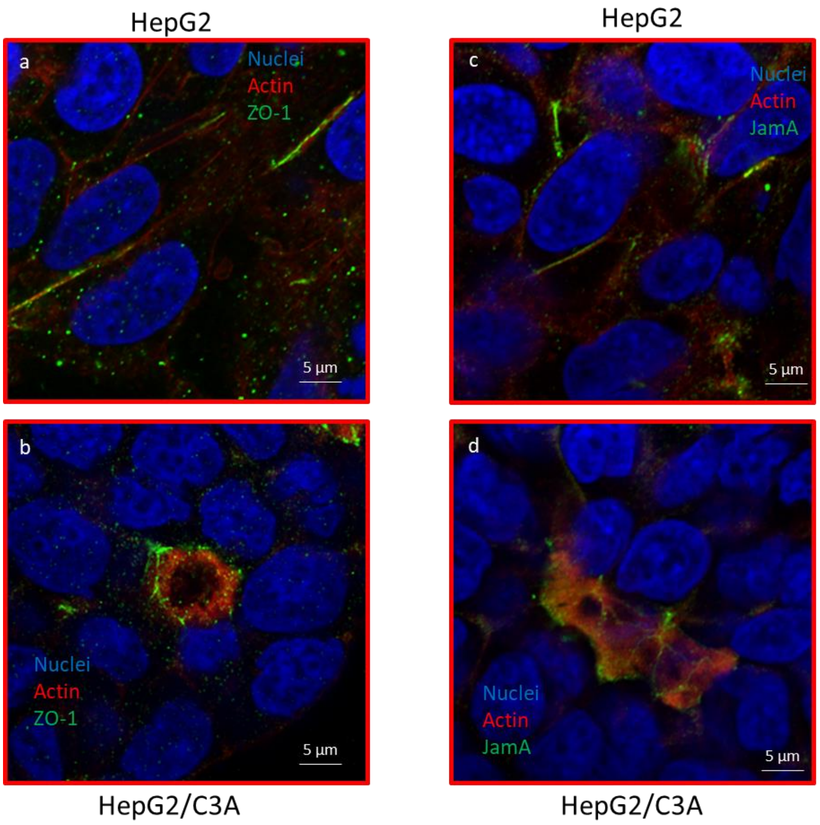


**Figure S5**. Comparison of localization of proteins of the tight junction between HepG2 and HepG2/C3A cell lines**:** confocal images acquired at 63x magnification in airyscan mode for HepG2 (a, c) and HepG2/C3A (b, d) cells, labeled with actin and ZO-1 (a, b) and actin and JamA (e, d).
